# A Cellular Platform for Production of C_4_ Monomers

**DOI:** 10.1101/2023.01.09.523327

**Authors:** Matthew A. Davis, Vivian Yaci Yu, Beverly Fu, Miao Wen, Edward J. Koleski, Joshua Silverman, Charles A. Berdan, Daniel K. Nomura, Michelle C. Y. Chang

**Affiliations:** Department of Molecular & Cell Biology, University of California, Berkeley, Berkeley, California 94720, United States; Department of Chemistry, University of California, Berkeley, Berkeley, California 94720, United States; Calysta, 1900 Alameda de las Pulgas Suite 200, San Mateo, California 94404, United States; Department of Nutritional Sciences and Toxicology, University of California, Berkeley, Berkeley, CA 94720, United States; Department of Molecular & Cell Biology, Department of Chemistry, Department of Nutritional Sciences & Toxicology, University of California, Berkeley, Berkeley, California 94720, United States; Department of Chemistry, Department of Molecular & Cell Biology, and Department of Chemical & Biomolecular Engineering, University of California, Berkeley, Berkeley, California 94720, United States

**Author notes:** Matthew A. Davis and Vivian Yaci Yu contributed equally to this paper.

**Keywords:** metabolic engineering, biofuels, bioplastics, adaptive laboratory evolution

## Abstract

Living organisms carry out a wide range of remarkable functions, including the synthesis of thousands of simple and complex chemical structures for cellular growth and maintenance. The manipulation of this reaction network has allowed for the genetic engineering of cells for targeted chemical synthesis, but it remains challenging to alter the program underlying their fundamental chemical behavior. By taking advantage of the unique ability of living systems to use evolution to find solutions to complex problems, we have achieved ~95% theoretical yield of three C_4_ commodity chemicals, *n*-butanol, 1,3-butanediol, and 4-hydroxy-2-butanone. Genomic sequencing of the evolved strains identified *pcnB* and *rpoBC* as two gene loci that are able to alter carbon flow by remodeling the transcriptional landscape of the cell, highlighting the potential of synthetic pathways as a tool to identify metabolic control points.

## INTRODUCTION

Living systems rely on a dynamic and complex network of chemical reactions to carry out the tasks needed to coordinate cellular growth and maintenance, allowing transformation of simple carbon sources into the thousands of molecules needed for life. As such, cells possess an enormous synthetic potential that can be engineered for targeted chemical synthesis, enabling the reduction of traditionally multi-stage synthetic routes into a single fermentation step that can be carried out in water and under ambient temperature and pressure. Their ability to utilize building blocks including sugars from renewable plant biomass, CO_2_, and CH_4_ for biosynthesis opens a route to shifting industrial chemical production away from its traditional reliance on petrochemical feedstocks towards a universal fermentation platform.

A major challenge in the development of cell-based chemical synthesis is that the reaction network used to produce target compounds is also used to carry out basic cell functions. These reactions are thus subject to many levels of regulation in order to maintain the necessary coordination between parts of the metabolic network.^1–3^ In particular, key hubs of the metabolic map, such as the central carbon pathways of glycolysis and the tricarboxylic acid (TCA) cycle, form many connections with the rest of the network and are difficult to manipulate as their behavior is affected by multiple inputs and outputs.^4^ As a result, the construction of high-yielding pathways can be difficult to achieve as evolution drives the cell to direct carbon flux to cell growth and biomass in competition with engineered biosynthesis.

Since these central carbon pathways are closely tied to cell state, they are correspondingly subject to homeostatic mechanisms to ensure robustness to change. Therefore, many simultaneous alterations are needed to rationally engineer carbon flow to insufficiently active nodes.^5–8^ However, an advantage that living systems provide is that evolution can be used to solve this multi-dimensional problem if product titers can be tied to cell growth.^9,10^ In this work, we demonstrate the design and evolution of synthetic pathways to selectively produce three industrially-relevant C_4_ compounds: 1,3-butanediol (butylene glycol, BDO), 4-hydroxy-2-butanone (HB), and *n*-butanol (Figure 1A). These three compounds are used for various purposes, ranging from pharmaceutical precursors (BDO and HB) to a drop-in gasoline replacement (*n*-butanol).^11–13^ In particular, BDO is valuable as a humectant or solvent for a variety of different high-value products, as well as a co-monomer for production of various polymers. These three compounds can also be further dehydrated to produce the C_4_ monomers 1,3-butadiene (from BDO),^14^ methyl vinyl ketone (from HB),^15^ and 1-butene (from *n*-butanol).^16^ Using a genetic selection, the yields of these pathways were improved from 11–20% to near quantitative yields. Genome sequencing of the evolved strains showed that two gene loci, *pcnB* and *rpoBC*, were mutated in the most successful daughter cells. Subsequent characterization demonstrated that mutations at these two loci are sufficient to capture the majority of the evolved phenotype and likely operate by large-scale shifts in the transcriptome. Taken together, these results highlight the possibility of synthetic pathways to be used not only for scalable chemical production but also as a platform for discovery and study of cellular function.

**Figure 1.**
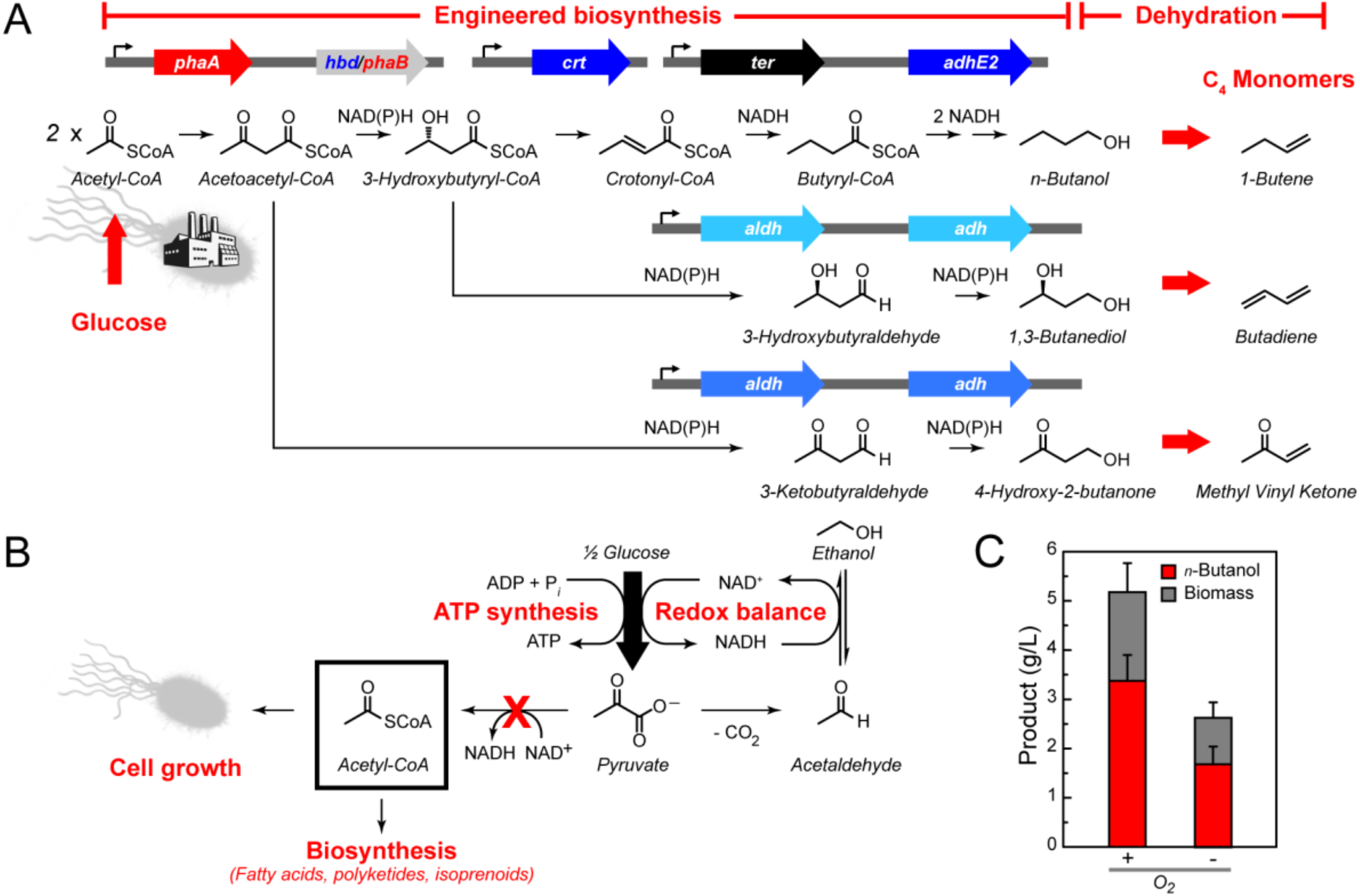
Synthetic pathways for production of C_4_ monomers. (A) Design of a platform for production of C_4_ monomers based on *n*-butanol formation. Identification of selective aldehyde and alcohol dehydrogenases enables the formation of three different C_4_ products from glucose, *n*-butanol, 1,3-butanediol, and 4-hydroxy-2-butanone via engineered microbes. Chemical dehydration of these compounds produces the industrially-relevant C_4_ monomers 1-butene, butadiene, and methyl vinyl ketone, respectively. (*phaA*, acetoacetyl-CoA synthase; *phaB*, R-specific NAD(P)H-dependent acetoacetyl-CoA dehydrogenase; *hbd*, S-specific NADH-dependent acetoacetyl-CoA dehydrogenase; *crt*, crotonase; *ter*, trans-enoyl-CoA reductase; *adhE2*, bifunctional aldehyde/alcohol dehydrogenase; *aldh*, aldehyde dehydrogenase; *adh*, alcohol dehydrogenase. Genes derived from the poly(hydroxyl)alkanote pathway of *Ralstonia eutrophus* are labeled in red. Genes derived from the acetone-butanol-ethanol pathway of *Clostridium acetobutylicum* are labeled in royal blue. Gene from *Treponema denticola* is labeled in black. Light blue *aldh* and *adh* genes denote their general function.) (B) Anaerobic fermentation pathways can operate at near quantitative yields in the absence of O_2_. Under these conditions, substrate-level phosphorylation pathways such as glycolysis serve as the only route to ATP synthesis but require the use of NAD^+^. In Baker’s yeast (*Saccharomyces cerevisiae*), decarboxylation of pyruvate and subsequent reduction to ethanol allow for the stoichiometric regeneration of NAD^+^ and is required for cell survival. Because of the low ATP yield under anaerobic growth, cell growth as well as flux to anabolic pathways utilizing the key building block, acetyl-CoA, are greatly reduced. As a result, acetyl-CoA is not readily available for the downstream biosynthesis of a broad range of target compounds during anaerobic growth. (C) Production of *n*-butanol and biomass in *E. coli* DH1 containing a synthetic butanol pathway (pBT33-Bu1 pCWori-ter.adhE2 pBBR1-aceEF.lpd) under aerobic and anaerobic conditions. Data are mean ± s.d. of biological replicates (n = 3).

## RESULTS AND DISCUSSION

### Implementation of a genetic selection for C_4_ production

Anaerobic cell culture is often preferred for industrial fermentations both because limitations in culture oxygenation on large-scale can be eliminated and because theoretical product yields can be increased.^17^ Under anaerobic conditions, carbon assimilation pathways like glycolysis serve as the primary route for cellular ATP synthesis since the lack of oxygen as a terminal electron acceptor makes aerobic respiration unavailable (Figure 1B).^9,18^ The relatively low ATP yield from substrate-level phosphorylation then results in a minimal loss of carbon to competing cell growth or biomass accumulation.^19^ In addition to 2 mol of ATP, 2 mol of NADH is also generated from 1 mol of glucose through glycolysis. Fermentative pathways provide a mechanism to oxidize NADH to NAD^+^, which is needed for glycolysis to remain operational. High flux through fermentation pathways is thus driven by cell survival. Lactate and ethanol production provide the paradigms for this process, resulting in rapid and near-quantitative yield from sugar via pyruvate (Figure 1B).

Like ethanol and lactate, the C_4_ alcohol, *n*-butanol, can serve to balance glucose fermentation because its biosynthesis recycles the 4 NADH produced per molecule of glucose. However, a major challenge for production of longer-chain target compounds is that they typically require building blocks from pathways downstream of glycolysis and whose intracellular concentrations are regulated at many levels. One of the most important of these building blocks is acetyl coenzyme A (CoA), which is a two-carbon intermediate that serves as a central point of many metabolic decision points.^20,21^ Its synthesis and usage are tightly controlled with flux dropping under anaerobic conditions as both biosynthesis and cell growth are greatly reduced during fermentative growth (Figure 1B). Indeed, *n*-butanol titers are greatly lowered when our first-generation *Escherichia coli* production strain was cultured anaerobically (Figure 1C).^11^ In order to reduce carbon flow to competing native pathways, the major fermentation pathways were knocked out of *E. coli* DH1 to generate DH1 *ΔldhA ΔadhE ΔfrdBC ΔpoxB ΔackA-pta* (DH1Δ5),^10,22^ a selection strain that would require the production of *n*-butanol under anaerobic conditions (Figure 2A, Supporting Information Table S1, Figure S1).^13^ In order to provide a means to increase flux to acetyl-CoA, the pyruvate dehydrogenase complex (PDHc, *aceEF-lpd*) was overexpressed for the oxidative decarboxylation of pyruvate to produce acetyl-CoA and NADH to stoichiometrically balance *n*-butanol production from glucose.

**Figure 2.**
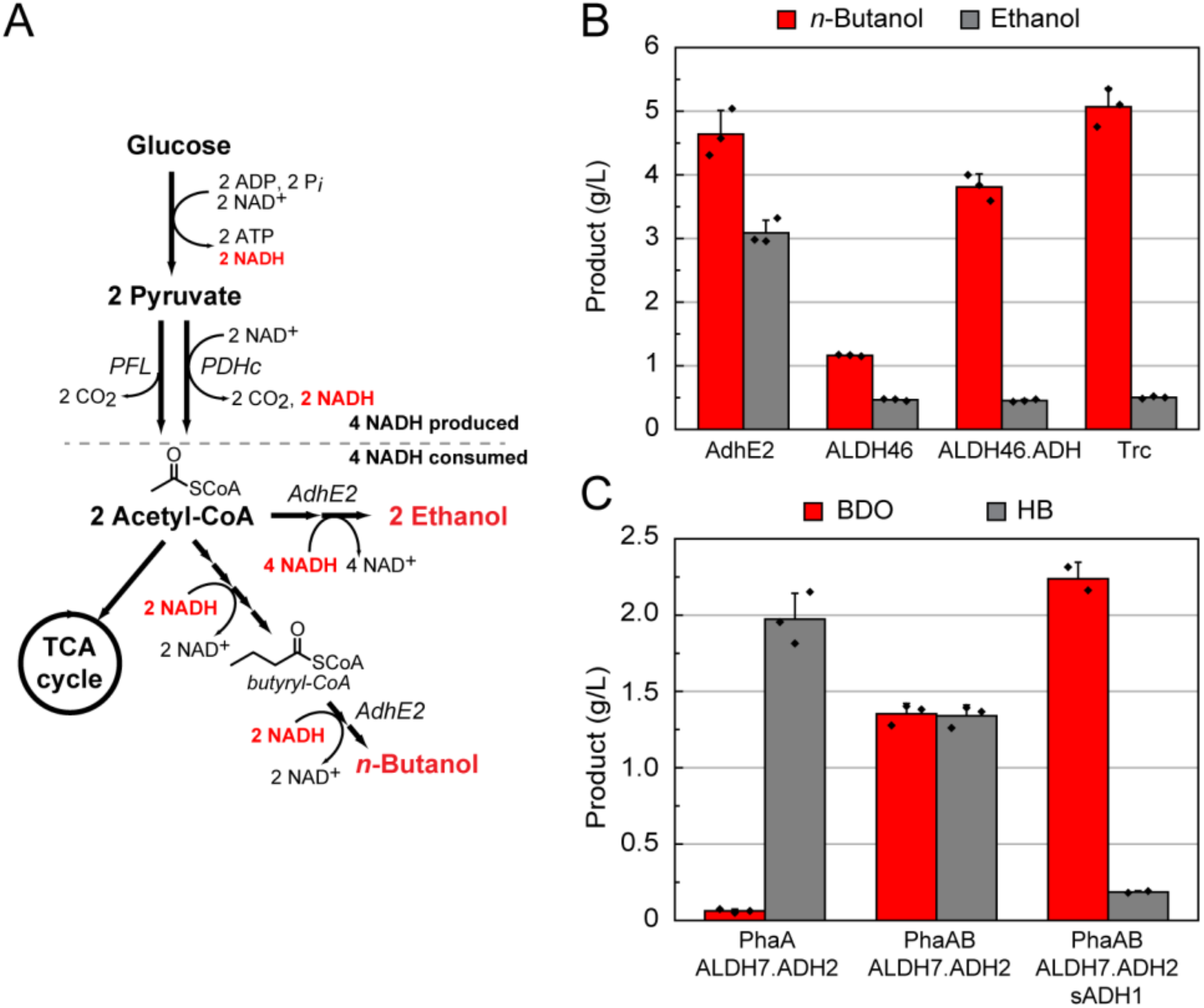
Production of C4 monomer precursors in engineered *E. coli*. (A) Design of a host for the anaerobic production of target compounds from acetyl-CoA. Deletion of the major fermentation pathways of *E. coli* in DH1Δ5 allows the synthetic *n*-butanol pathway to be the major mechanism for balanced NAD^+^ regeneration production via the acetyl-CoA intermediate. However, the promiscuity of AdhE2 towards acetyl-CoA and butyryl-CoA leads to ethanol fermentation as a pathway short-circuit that also maintains stoichiometric redox balance. (PDHc, pyruvate dehydrogenase complex; PFL, pyruvate formate lyase)(B) Screening of AdhE, ALDH, and ADH candidates in *E. coli* DH1Δ5 pBBR1-aceEF.lpd pBT33-Bu2 yields a C4-selective fermentation pathway under anaerobic conditions. When AdhE2 is included, high levels of ethanol are produced along with the target n-butanol product. Replacement with ALDH46 reduces ethanol production to background levels but concomitantly drops n-butanol titers. Addition of the ADH domain from AdhE2 and tuning the promoter for expression allows for high n-butanol yields with very little ethanol being formed. All strains were grown anaerobically in TB with 2.5% (w/v) glucose media for 3 d post induction. (AdhE2, pCWori-ter-adhE2; ALDH46, pCWori-ter.aldh46; ALDH46.ADH, pCWori-ter-aldh46.ADHAdhE2; Trc, pCWori.trc-ter-aldh46.ADHAdhE2). (C) Screening of ALDH, ADH, and sADH candidates in E. coli pT533-phaA/phaAB pCWori.trc-ter-ALDH.ADH led to identification of the ALDH7.ADH2 pair for production of HB and BDO under anaerobic conditions. In the absence of PhaB, HB is selectively produced. Addition of PhaB leads to a 1:1 ratio of both products being formed. The inclusion of sADH then allows for HB to be converted to BDO. All strains were grown anaerobically in TB with 2.5% (w/v) glucose media for 5 d post induction. Data are mean ± s.d. of biological replicates (n = 3).

The fermentation knockout strain DH1Δ5 was found competent to grow under anaerobic conditions when the synthetic *n*-butanol pathway consisting of *phaA*, *hbd*, *crt*, *ter*, and *adhE2* was expressed (Figure 1). However, increased ethanol production, rather than *n*-butanol, was observed (Figure S2). The most likely source of ethanol was the promiscuity of AdhE2, which is a bifunctional aldehyde-alcohol dehydrogenase that produces both *n*-butanol and ethanol from the respective two-step reduction of butyryl-CoA and acetyl-CoA in its native host (Figure 2A).^23,24^ In this case, direct reduction of two equivalents of acetyl-CoA to ethanol would also regenerate the necessary four NAD^+^ per glucose and creates a short circuit in the pathway to circumvent *n*-butanol production (Figure 2A). Biochemical analysis of different AdhE2 constructs was carried out in order to assess the selectivity of each domain for the C_4_ and C_2_ substrates (Figure S3). Although there is a ten-fold preference for butyryl-CoA over acetyl-CoA, the higher *k*_cat_/*K*_M_ for the C_4_ substrate arises directly from a 10-fold lower *K*_M_ (10 ± 1 μM) with no change in *k_cat_* within error. Given that the *K_M_* for acetyl-CoA (100 ± 10 μM) is well within the expected physiological range (0.5–1.0 mM), it is likely that AdhE2 is capable of producing both *n*-butanol and ethanol at competitive rates under intracellular conditions. Although the mechanism of substrate channeling between the aldehyde dehydrogenase (ALDH) and alcohol dehydrogenase (ADH) domains is not yet fully understand, the high *K*_*M*s_ (4.0–4.5 mM) measured for the aldehyde intermediate imply that substrate selection is controlled by the ALDH domain. We thus set out to identify enzymes that could efficiently carry out the reduction of butyryl-CoA while excluding acetyl-CoA.

### Identifying C_4_-selective dehydrogenases

Enzymes that tailor acyl-CoA substrates are typically permissive to a broad range of chain lengths, making the exclusion of a smaller substrate, like acetyl-CoA, challenging.^25,26^ We thus initiated a search for acylating ALDH candidates with characterized substrate selectivity and found three C_2_-specific bifunctional AdhE2 homologs, four C_4_-specific monofunctional ALDHs, and one atypical C_2_-specific ALDH (SI Appendix, Figure S4). These sequences were arranged in a biased phylogenetic tree with branching guided by characterized substrate preference (Figure S4). Next, the entire ALDH gene family from the Pfam database was assembled into a second phylogenetic tree based on the first tree (Figure S4) to generate a full family tree of < 1,200 sequences.^27^ This tree was dominated by monofunctional ALDH sequences (67%) as the majority of ALDH domains found in the Pfam database are derived from standalone enzymes. Approximately 40 mutations were selected from the natural sequence diversity in the C_4_ branch and used to design 95 AdhE2 variants (Table S1). Each variant contained 3–5 mutations, and every mutation was present in multiple variants, ensuring that each mutation can be evaluated in multiple contexts. These AdhE2 variants were synthesized, cloned into expression vectors, and cotransformed into DH1Δ5 with the appropriate butanol production plasmids for in vivo screening (Figure S5). Around two-thirds of the variants (66 variants, 69%) remained active; however, only mild improvements in the *n*-butanol:ethanol ratio were observed.

Given the modest gains using this approach, we turned our attention to screening wild-type ALDH sequences falling within the C_4_-selective branch since it seemed likely that the sequence information derived mostly from monofunctional ALDHs did not accurately predict the selectivity of their bifunctional counterparts. The C_4_ branch of the tree was widely sampled to incorporate the full diversity of the branch in a small number of sequences comprising 15 bifunctional AdhE2 homologs and 18 monofunctional ALDHs (Table S1). We found that all bifunctional enzymes except one yielded lower *n*-butanol:ethanol ratios compared to AdhE2 (Figure S5), possibly because a large majority of sequenced AdhE2 homologs are thought to be involved in ethanol generation and likely display a natural preference for acetyl-CoA. In contrast, 15 out of 16 monofunctional ALDHs produced more *n*-butanol than ethanol (Figure S6). We next sought to improve overall *n*-butanol production to the levels observed using AdhE2 (Figure 2B). We reasoned that the bottleneck was the reduction of butyraldehyde to *n*-butanol due to the absence of the ADH domain. In our biochemical studies of AdhE2, we characterized a truncation mutant consisting of solely the ADH domain of AdhE2 (ADH_AdhE2_) that surprisingly showed an order of magnitude decrease in KM for butyraldehyde to 300 ± 50 μM. We thus supplemented our pathway containing ALDH46 with ADH_AdhE2_ domain, which more than doubled *n*-butanol titers. Increasing the promoter strength for expression of ALDH46-ADH_AdhE2_ then improved *n*-butanol titers in the two-protein system beyond that observed in the original bifunctional AdhE2-dependent pathway, with no ethanol production above background (Figure 2B).

### Developing a platform for the production of C_4_ commodity chemicals

With a family of C_4_-selective monofunctional ALDHs in hand, we set out to explore the possibility of producing other important C_4_ commodity chemicals from our *n*-butanol pathway. In particular, reduction of 3-hydroxybutyryl-CoA, an intermediate in the butanol production pathway, yields BDO (Figure 1A).^28^ Upon chemical dehydration, BDOs can be used to produce butadiene for synthetic rubber production, which is currently produced from fossil fuel sources at the level of >10 million metric tonnes per year.^29,30^ We therefore set out to screen our 16-member ALDH library for potential candidate enzymes to construct a BDO pathway that can reduce either stereoisomer, (*R*)-3- or (*S*)-3-hydroxybutyryl-CoA (Figure S7). In this screen, we found that all candidates were competent to produce 1,3-butanediol at levels from 150–700 mg L^-1^. Interestingly, we found very little sensitivity to the stereochemistry of the substrate, although preferences between butyryl-CoA and 3-hydroxybutyryl-CoA reduction were observed.

We hypothesized that the differences in *n*-butanol compared to BDO production might arise from limitations in ADH activity for reduction of 3-hydroxybutyraldehyde. As such, we generated an ADH sequence similarity network with the goal of identifying a subfamily with the desired substrate selectivity within the larger superfamily (Figure S8).^31,32^ A list of candidates within the subfamilies defined by the known C_4_-selective ADHs (*bdhA*, *bdhB*, *dhaT*, and *yqhD*) was then generated and screened by co-expression with ALDH46, which showed no stereochemical preference for reduction of 3-hydroxybutyryl-CoA. While several hits were found, it was interesting to note that they appeared to all be highly specific for the (*R*)-isomer.

During this analysis, we identified HB as a side-product that appears to arise from the reduction of an earlier pathway intermediate, acetoacetyl-CoA (Figure 1A). HB is also an interesting product as its dehydration produces methyl vinyl ketone, a reagent used in the production of fine chemicals,^33^ as well as a potential monomer unit for polymers.^34^ We therefore set out to characterize the selectivity of ALDH-ADH pairs by examining partitioning between BDO and HB (Figure S9). This screen indicated HB production is highly specific to the ALDH7-ADH2 pair, providing an even distribution of products at high titer (3.4 ± 0.1 g L^-1^). On the other end, the ALDH3-ADH22 pair was found to capture a large fraction of the C_4_ product pool as BDO (81%), producing 2.9 ± 0.1 g L^-1^ of total products under screening conditions.

A selective pathway for production of HB over BDO was engineered by simply removing the PhaB ketoreductase, forming a truncated pathway by eliminating production of 3-hydroxybutyryl-CoA required for BDO formation (Figure 1A). With this change, the PhaA-ALDH7-ADH2 pathway generated 2.0 ± 0.2 g L^-1^ HB (Figure 2C). To selectively produce BDO over HB, we pursued an approach to redirect HB to BDO production by adding a secondary alcohol dehydrogenase (sADH). Specifically, we set out to find an sADH that would catalyze the reduction of HB, resulting from promiscuous ALDH activity on acetoacetyl-CoA, directly to BDO (Figure S10).

A number of sADHs have been reported to reduce 4-hydroxy-2-butanone or similar substrates.^35^ Several of these sADHs were co-expressed with the ALDH7-ADH2 pair, which consistently produced an even mixture of butanediol and hydroxybutanone. Several of the sADHs enabled a shift in the product profile, producing high levels of BDO (>2 g L^-1^) and minimal amounts of HB (<250 mg L^-1^) (Figure S10). With these sADHs in hand, we could now control the product profile between HB, BDO, or a mixture of the two (Figure 2C).

### Adaptive evolution of C_4_ pathways

With highly specific pathways established for production of *n*-butanol, HB, and BDO in place, we next set out to develop a genetic selection for increasing titers under anaerobic conditions with the longterm goal of gaining new insight into the manipulation of central carbon homeostasis. In contrast to our results with the promiscuous *n*-butanol pathway containing the ethanol shortcircuit (Figure 3A), growth of the fermentation-deficient strain, DH1Δ5, depends solely on *n*-butanol production. Using a set of control plasmids with low, medium, and high *n*-butanol productivity, we observe that rescue of DH1Δ5 growth under anaerobic conditions is directly correlated to product titer and thus the capacity of the synthetic pathway to recycle NADH. Indeed, strains complemented with a very low-flux pathway do not grow significantly, if at all, while strains complemented with high flux pathway variants grow similarly to wild-type.

**Figure 3.**
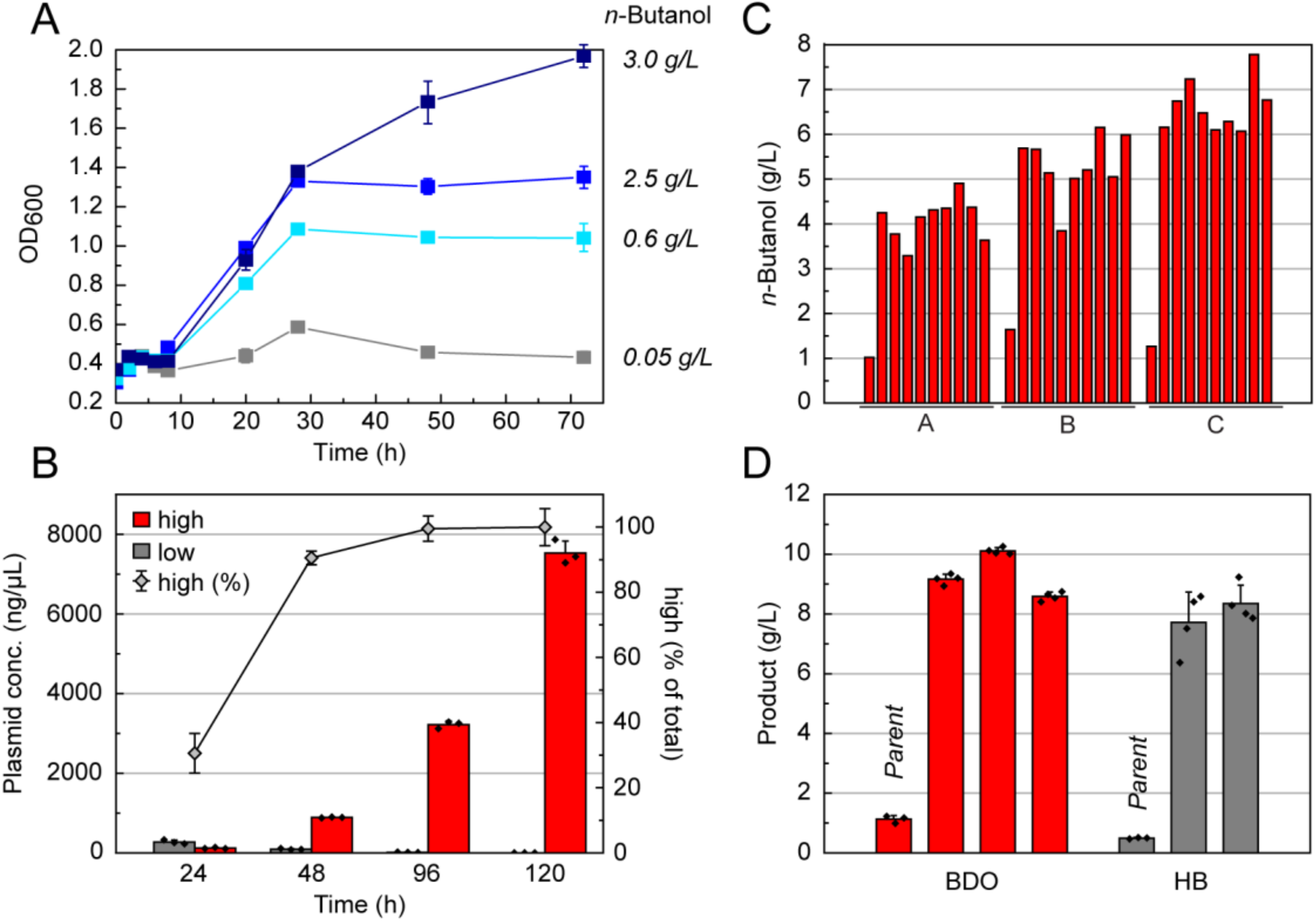
Development of a genetic selection for evolving C_4_ monomer synthesis in *E. coli*. (A) The *n*-butanol pathway complements the deletion of the native fermentation pathways of *E. coli* under anaerobic conditions. *n*-Butanol pathway variants displaying a range of yields were transformed into DH1Δ*5* and cultured anaerobically. Growth was monitored by OD_600_ and *n*-butanol production was quantified at the end of the experiment. All strains were grown in TB with 2.5% (*w*/*v*) glucose media. All strains contained pCDF3-ter.aldh46 and one of the following variants of the pBT-Bu2 plasmid in order of increasing butanol production: pBT-0.03HBD, pBT-0.3crt, pTT-Bu2, pBT-Bu2. (B) Enrichment of a medium-producing *n*-butanol strain, DH1Δ*5* pCDF3-ter.aldh46 pBT-0.3crt, was observed when mixed (0.1%) with a low-producing strain DH1Δ5 pCDF3-ter.aldh46 pBT-0.03HBD (99.9%). The mixed culture was grown anaerobically in TB with 2.5% (*w*/*v*) glucose media. Changes in the genetic population were monitored using qPCR. (C) A representative adaptive evolution for *n*-butanol production. *E. coli* BW25113Δ*5* pBBR1-aceEF.lpd pT5T33-Bu2 containing either pCWOri.trc-ter-ALDH46.ADH2 [A], pCWOri.trc-ter-ALDH46.ADH8 [B], or pCWOri.trc-ter-ALDH21.ADH2 [C] was subjected to multiple rounds of dilution in M9 containing 10% (v/v) LB and 2.5% (w/v) glucose under anaerobic conditions. Individual clones were then isolated and characterized for their *n*-butanol titers compared to the parent strain. (D) Characterization of BDO and HB strains after adaptive evolution. Data are mean ± s.d. of biological replicates (n = 3).

Given the dependence of growth rate on pathway titer, we then tested our ability to enrich cell cultures for high producing variants. To do so, cultures of the low-production strain were seeded with either 0.1% or 1% of the medium-production inoculated strain. Throughout the course of extended anaerobic growth, we observed a significant lag phase dependent on the seeding level (Figure S11). In this simulated selection, we tracked *n*-butanol production as well as the abundance of the two sub-populations using qPCR. In agreement with the growth curves, the abundance of the low-production strain was largely static while the abundance of the medium-production strain was enriched >40-fold as *n*-butanol production initiated (Figure 3B).

In order to select for variants with improved *n*-butanol productivity under anaerobic conditions, we turned to adaptive evolution after efforts using synthetic mutagenesis methods such as chemical mutagens or UV irradiation appeared to find only local minima in the evolutionary trajectory (Figure S12). In this approach, the natural mutation frequency is utilized, which requires longer evolution times but selects for more advantageous mutations and minimizes the occurrence of neutral mutations.^36,37^ Since every evolutionary trajectory has the potential to yield different results, we evolved two different host strains, DH1Δ5 and BW25113Δ5, using media ranging in richness from M9, 10% (*v*/*v*) LB in M9, and LB, by diluting the culture every 24 h over the course of 4–70 days (Figure S13). Using this approach, we were able to evolve strains six-fold from 11% to 66% carbon conversion as well as from 43% to >95% yield under these various conditions (Figure 3C). Although the redox balance is not stoichiometric as it is with *n*-butanol, we were also able to evolve BDO and HB production in DH1Δ5 from 20% to ~95% theoretical carbon conversion in TB (Figure 3D, Figure S14). Isolation of pathway plasmids from the evolved strains and transformation into a clean background showed no improvement in product titers, indicating that the relevant mutations were generated on the chromosome (Figure S14). Culture of the evolved strains in rich media in shake-flasks with an oleyl alcohol overlay further yielded titers of up to 47 ± 6 g L^-1^ and >95% yield (Figure S15). Taken together, the evolved strains demonstrate robust production of a range of C_4_ products from acetyl-CoA under anaerobic conditions.

### Identifying players in transcriptional re-programming

We took a genome scale approach to explore key factors responsible for the evolution of this large shift in central carbon flow. A total of 31 isolated strains from three independent selections for *n*-butanol (21 strains), BDO (8 strains), and HB (2 strains) production carried out under different growth conditions were sequenced to identify the changes between the genomes of the parental and evolved strains. Interestingly, we found mutations only in a handful of genes, which consistently appeared regardless of selection conditions (Figure 4A, Table S2). In addition, a few mutations mapped to the non-coding portions of the genome (0–1 mutation per strain with a total number of six distinct mutations from all 31 strains that were sequenced) along with rearrangements that appeared to be mostly associated with mobile elements. Of the mutations in coding regions, the most striking is the finding that poly(A) polymerase (*pcnB*) and/or the RNA polymerase *β* and *β′* subunits (*rpoBC*) were mutated in nearly all of the most successful evolved strains. These two gene loci are involved in regulating the transcriptional landscape of the cell by forming part of the transcription complex (*rpoBC*)^38,39^ as well as by controlling the lifetime of mRNAs by polyadenylation (*pcnB*).^40^ Mutations in *rne* (ribonuclease E) also occurred frequently (12%) in the evolved *n*-butanol strains.

**Figure 4.**
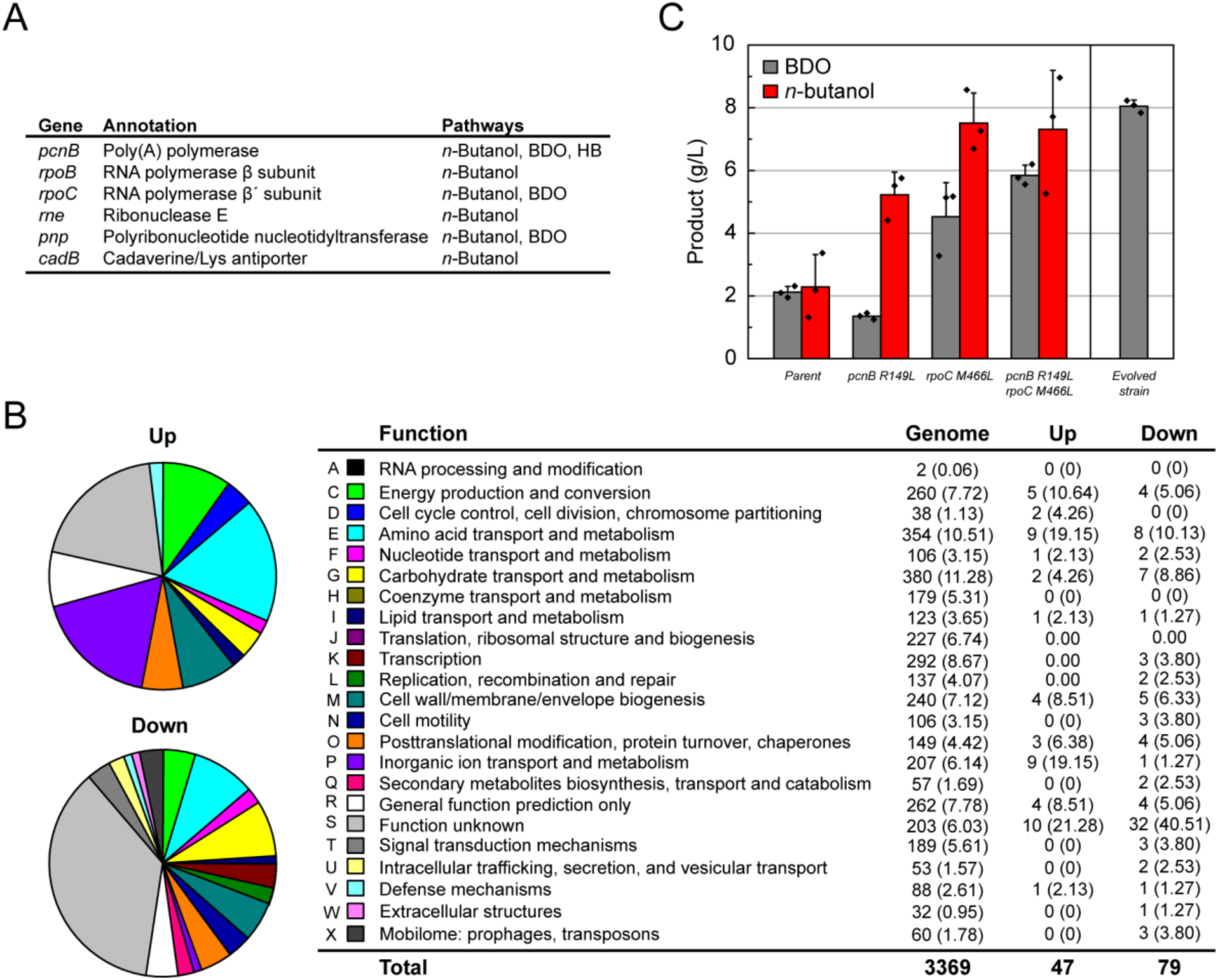
Characterization of evolved *E. coli* strains for C_4_ monomer synthesis. (A) List of genes found mutated in more than one evolved strain carrying the *n*-butanol, BDO, or HB pathways. (B) Clusters of orthologous groups (COG) categories for genes differentially expressed between the parent (DH1Δ*5* pT533-phaA.phaB pCWori.trc-ALDH7.ADH2) and evolved BDO strain (DH1Δ*5*.2406). COG categories were identified by the IMG-ER annotation pipeline. COG categories represented by genes that are upregulated and downregulated 24 h after induction with IPTG. Comparison of COG category representation in the differentially expressed genes compared to the entire genome. The number of the open reading frames represented by each COG is given, and the percentage of total genes with COG categories is in parentheses. Since some genes fall into multiple COG categories, the percentage was calculated by dividing the total number of unique genes. (C) Generating the *pcnB* and *rpoC* mutations found in DH1Δ*5*.2406 in a clean genetic background (DH1Δ*5* parent) captures the majority of the improvement observed in the evolved strain, indicating that these two gene loci play an important role in enabling the increases in BDO production. Introduction of the *n*-butanol pathway into DH1Δ*5.pcnB*(R149L).*rpoC*(M466L) shows that some aspects of this phenotype can be transferred to other pathways.

The discovery that genes involved in RNA metabolism appear to drive metabolic network evolution led us to the hypothesis that the phenotypic changes were largely being controlled by alterations in the global transcriptional program. This model is consistent with pathway enzymatic activity measure in cell lysates, which showed no significant increase between parent and evolved strains at the end of a production growth (Figure S16). This result suggests that yield increases were not derived from overexpression of heterologous pathway genes. To further characterize this phenomenon, we performed an RNA-Seq experiment on the evolved BDO strain with the largest improvement in production titer (DH1Δ5.2406) containing point mutations in *pcnB* and *rpoC*. We found 126 differentially-expressed genes (β value > 2) between the parental and evolved strain (Figure S17), indicating that alterations in acetyl-CoA and central carbon homeostasis may require changes at many metabolic nodes. This data is supported by metabolomics experiment that suggest that acetyl-CoA levels are higher in the evolved strains (Figure S18). These genes fall into a broad range of categories but dominated by energy production and conversion, amino acid transport and metabolism, cell envelope biogenesis, and carbohydrate transport and metabolism (Figure 4B).

In order to validate the impact of the *pcnB* and *rpoC* mutations, the two mutations observed in this BDO strain (*pcnB* R149L / *rpoC* M466L) were introduced into a clean genetic background. These experiments show that these mutations in *rpoC* and *pcnB* are synergistic, as both are required to achieve a substantive increase in BDO titer compared to the parent (Figure 4C). Indeed, the double mutant demonstrates a 2.75-fold increase in BDO titers (parent, 2.1 ± 0.1 g L^-1^; double mutant, 5.8 ± 0.2 g L^-1^), which recapitulates 73% of the improvement observed in the fully evolved strain (8.1 ± 0.1 g L^-1^). We were also interested in the generality of these mutations and thus tested their ability to increase the yields of other synthetic pathways. When the *n*-butanol pathway is introduced into the double mutant, we observe a 3.2-fold increase in product titer from 2.3 ± 0.6 to 7.3 ± 1.1 g L^-1^. (Figure 4C). Altogether, these data show mutations in only two genes, *pcnB* and *rpoC*, are capable of driving a large shift in central carbon metabolism that can be generalized to related pathways utilizing the acetyl-CoA building block.

## CONCLUSIONS

In this work we have demonstrated the anaerobic production of three industrially relevant C_4_ chemicals BDO, HB, and *n*-butanol, at >95% of theoretical yields. Furthermore, BDO, HB and *n*-butanol serve as bioproduct precursors to 1,3-butadiene, methyl vinyl ketone, and 1-butene respectively via dehydration. Overall, six C_4_ chemicals can be accessed from glucose using a single platform. Notably, the production of advanced products from acetyl-CoA can be difficult when switching to anaerobic metabolism. Indeed, acetyl-CoA represents a metabolic checkpoint where carbon is differentiated from pyruvate towards different cell fates, including lowered cell growth and TCA flux, which is one of the major consumers of acetyl-CoA. As such, rational engineering of central carbon pathways for the purpose of rerouting flux to a synthetic product can be quite challenging as it opposes the cell’s evolutionary impetus to direct carbon towards biomass. However, using a selection design in which fitness is driven by product titer, strains could be identified with up to 5-fold improvements in yield and near quantitative production from the acetyl-CoA building block.

In our system, we found that use of natural adaptive evolution allowed us to rapidly reach high production strains compared to the use of mutagens that increase the mutation rate but appeared to only find local minima in the evolutionary trajectory. Genome-level characterization of these strains revealed that mutations in two gene loci, *pcnB* and *rpoBC*, were sufficient to enable the needed shifts in carbon flow. Interestingly, these mutations have been previously observed in studies of *E. coli* evolution for both growth in M9 (*rpoBC*)^41^ and production of *n*-butanol (*pcnB*).^13^ Physiological studies suggest that this effect may rely on remodeling the transcriptome by influencing RNA metabolism along with *rne*. Interestingly, a wide range of mutations were identified within these three genes, some of which have been found to be important for activity in biochemical studies.^40^ This finding suggests that these four genes could be used for diversity generation at the phenotypic level by inducing pleiotropic changes in the transcriptional landscape. Furthermore, it was found that mutations found in the evolved BDO strain could be translated to significant increases in *n*-butanol yields, indicating that these strains could be relevant to the production of other acetyl-CoA-derived products such as fatty acids, polyketides, and isoprenoids.

In conclusion, living systems offer a unique advantage for chemical synthesis to increase product yields through evolution. In particular, central carbon metabolism plays an essential role in cell fitness and thus represents a key regulator and reporter of cellular state.^42^ These pathways are subject to tight homeostasis with multiple mechanisms to ensure robustness and reduce sensitivity to change. ^43^ In this regard, engineered pathways provide an interesting platform where product titer can be treated as a synthetic phenotype or marker for quantitative assessment of genetic traits that lead to large shifts in the regulatory and metabolic network. ^44^ By using evolution to solve difficult design challenges, we can also take advantage of synthetic pathways to identify new strategies to alter behaviors that are hard-wired into the systems-level behavior of the host.

## ASSOCIATED CONTENT

### Supporting Information

The Supporting Information is available free of charge.

## AUTHOR INFORMATION

**Corresponding Author**, Michelle C. Y. Chang – *Department of Chemistry, Department of Molecular & Cell Biology, and Department of Chemical & Biomolecular Engineering, University of California, Berkeley, Berkeley, California 94720, United States*

## Author Contributions

The manuscript was written through contributions of all authors. / All authors have given approval to the final version of the manuscript.

## Notes

The authors declare no competing financial interest.

## ACKNOWLEDGMENT

This work was funded by the generous support of the National Science Foundation through a CAREER Award (029504-003) to M.C.Y.C. and the Center for Sustainable Polymers, a National Science Foundation-supported center for Chemical Innovation (CHE-1901635).

## ABBREVIATIONS

BDO: 1,3-butanediol
HB: 4-hydroxy-2-butanone
TCA: tricarboxylic acid
PDHc: pyruvate dehydrogenase complex
ALDH: aldehyde dehydrogenase
ADH: alcohol dehydrogenase
sADH: secondary alcohol dehydrogenase
qPCR: real-time polymerase chain reaction
LB: Luria broth
TB: terrific broth
RNA-seq: RNA sequencing

## Notes

### Competing Interest Statement

The authors have declared no competing interest.

