## Supporting Information for "A Cellular Platform for Production of C_4_ Monomers"

**Materials and Metho**ds

| *Commercial materials* | *S2* |
| --- | --- |
| *Bacterial strains* | *S3* |
| *Gene and plasmid construction* | *S3* |
| *Expression and purification of AdhE2 constructs* | *S9* |
| *AdhE2 biochemical assays* | *S9* |
| *ALDH domain phylogenetic analysis* | *S10* |
| *Production of C_4_ compounds in shake flasks* | *S11* |
| *Quantification of n-butanol titers* | *S11* |
| *Quantification of 1,3-butanediol and 4-hydroxy-2-butanone titers* | *S12* |
| *Anaerobic growth competition and enrichment validation* | *S12* |
| *Adaptive evolution* | *S12* |
| *Genome sequencing* | *S13* |
| *Cell lysate enzyme assays* | *S13* |
| *RNA sequencing and analysis* | *S14* |
| *Generation of chromosomal point mutations* | *S15* |

**Supplementary Results**

| **Table S1.** | *Strains, plasmids, oligonucleotides, and sequences* | *S17* |
| --- | --- | --- |
| **Figure S1** | *Fermentation pathways of E. coli and gene knockouts* | *S139* |
| **Figure S2** | *Dependence of the n-butanol:ethanol ratio on oxygen* | *S140* |
| **Figure S3.** | *Biochemical characterization of AdhE2 constructs* | *S141* |
| **Figure S4.** | *Phylogenetic analysis of ALDH domains* | *S143* |
| **Figure S5.** | *Design and screening of AdhE2 mutants* | *S144* |
| **Figure S6.** | *Screening of monofunctional ALDHs* *for C_4_ selectivity* | *S145* |
| **Figure S7.** | *Screening ALDHs for production of BDO* | *S146* |
| **Figure S8.** | *Identification and screening of ADHs for BDO production* | *S150* |
| **Figure S9.** | *Screening of ALDH-ADH pairs for BDO vs. HB production* | *S153* |
| **Figure S10.** | *Screening of sADHs to increase BDO selectivity* | *S155* |
| **Figure S11.** | *Development and validation of a genetic selection for n-butanol production* | *S157* |
| **Figure S12.** | *Selection of n-butanol strains using EMS mutagenesis under anaerobic conditions* | *S158* |
| **Figure S13.** | *Characterization of adaptive evolution of n-butanol strains under anaerobic conditions* | *S159* |
| **Figure S14** | *Characterization of adaptive evolution of BDO and HB strains under anaerobic conditions* | *S162* |
| **Figure S15.** | *Production titers of C_4_ monomers of parent and evolved strains with an organic overlay* | *S164* |
| **Table S2.** | *Genome sequencing of evolved n-butanol, BDO, and HB strains* | *S166* |
| **Figure S16.** | *Cell lysate enzyme activities of n-butanol pathway enzymes for parent and evolved strains* | *S170* |
| **Figure S17.** | *RNA-Seq data for the WT and the BDO evolved strain* | *S171* |
| **Figure S18.** | *Acetyl-CoA pool analysis between parent, mutant, and the evolved strain* | *S172* |

**Literature Cited**	*S173*

**Materials and Methods**

**Commercial materials.** Terrific Broth (TB), LB Broth Miller (LB), LB Agar Miller, glycerol, and dimethylsulfoxide (DMSO) were purchased from EMD Biosciences (Darmstadt, Germany). Carbenicillin (Cb), kanamycin (Km), chloramphenicol (Cm), isopropyl-ß-d-thiogalactopyranoside (IPTG), phenylmethanesulfonyl fluoride (PMSF), tris (hydroxymethyl) aminomethane hydrochloride (Tris-HCl), sodium chloride, dithiothreitol (DTT), 4-(2-hydroxyethyl)-1-piperazineethanesulfonic acid (HEPES), and magnesium chloride hexahydrate were purchased from Fisher Scientific (Pittsburgh, PA). Tris-(2-carboxyethyl)phosphine hydrochloride (TCEP) was purchased from Biosynth, Inc. (Itasca, IL). Imidazole was purchased from Acros Organics (Morris Plains, NJ). Sodium hydroxide was purchased from Avantor Performance Materials (Center Valley, PA). Coenzyme A sodium salt hydrate (CoA), acetyl-CoA, butyryl-CoA, acetoacetyl-CoA, β-nicotinamide adenine dinucleotide reduced dipotassium salt (NADH), β-nicotinamide adenine dinucleotide hydrate (NAD^+^), formic acid, trichloroacetic acid (TCA), β-mercaptoethanol (BME), lysozyme from chicken egg white, and bovine serum albumin (BSA) were purchased from Sigma-Aldrich (St. Louis, MO). Sodium phosphate dibasic hepthydrate, and N,N,N',N'-tetramethyl-ethane-1,2-diamine (TEMED) were purchased from Sigma-Aldrich (St. Louis, MO). Acrylamide/Bis-acrylamide (30%, 37.5:1), electrophoresis grade sodium dodecyl sulfate (SDS), Bio-Rad protein assay dye reagent concentrate, and ammonium persulfate were purchased from Bio-Rad Laboratories (Hercules, CA). Restriction enzymes, T4 DNA ligase, Phusion DNA polymerase, Q5 DNA polymerase, T5 exonuclease, and Taq DNA ligase were purchased from New England Biolabs (Ipswich, MA). Deoxynucleotides (dNTPs) and Platinum Taq High-Fidelity polymerase (Pt Taq HF) were purchased from Invitrogen (Carlsbad, CA). PageRuler™ Plus prestained protein ladder was purchased from Fermentas (Glen Burnie, Maryland). **Oligonucleotides were purchased from Integrated DNA Technologies (Coralville, IA), resuspended at a stock concentration of 100** µM in 10 mM Tris-HCl, pH 8.5, and stored at either 4 °C for immediate use or –20 °C for longer term use. Amicon Ultra 10,000 centrifugal concentrators were purchased from EMD Millipore (Billerica, MA). cOmplete EDTA-free protease inhibitor was purchased from Roche Applied Science (Penzberg, Germany). TEV protease was purchased from the QB3 MacroLab at UC Berkeley. Amicon Ultra spin concentrators and MultiScreen_HTS_ 0.22μm filter plates were purchased from Merck Millipore (Cork, Ireland). d-(+)-glucose was purchased from MP Biochemicals (Santa Ana, CA). 2,4-Pentanediol, 1,3-butanediol, and 4-hydroxy-2-butanone were purchased from Sigma-Aldrich (St. Louis, MO). DNA purification kits, Ni-NTA agarose, genomic DNA isolation, and RNeasy RNA isolation kit were purchased from Qiagen (Valencia, CA). Genome library prep Kapa Biosystem Hyper Plus Kit was purchased from Kapa Biosystem (Wilmington, MA). Illumina TruSeq RNA Sample Prep Kit was purchased from Illumina (Hayward, CA).

**Bacterial strains.** *E. coli* DH10B was used for DNA construction and BL21(DE3) Star-T1^R^ was used for heterologous production of proteins for purification. *E. coli* DH1 (ATCC 39936), DH1Δ5, BW25113Δ5-T1R, DH1Δ5_2406_pcnB(R149L), DH1Δ5_2406_rpoC(M466L), and DH1Δ5_2406_pcnB(R149L)_rpoC(M466L) were used for production and evolution experiments. Genotypes are listed in *Table S1A*.

DH1Δ5 and BW25113Δ5-T1R were constructed with the lamda-red recombinase system [*1*]. The pKD3 template plasmid was used for gene disruption with subsequence recycling of the Cm^R^ marker using the pCP20 plasmid. The following primers were used.

| **Name** | **Sequence** | | **Note** |
| --- | --- | --- | --- |
| adhE F1 | GCTTTGCAAAAATTTGATTTGGATCACGTAATCAGTACCC | Δ*adhE* verification | |
| adhE R1 | CATGAGCAGAAAGCGTCAGGCAGTGTTGTATCCAC | Δ*adhE* verification | |
| adhE KF1 | TAGTAGCTTAAATGTGATTCAACATCACTGGAGAAAGTCTTgtgtaggctggagctgcttc | Δ*adhE* | |
| adhE KR1 | ATCTGAATCAGCTCCCCTGGAATGCAGGGGAGCGGCAAGAcatatgaatatcctccttag | Δ*adhE* | |
| fabB KO F2 | attgtgcattcgaaacttactctatgtgcgacttacagaggtattgaATGgtgtaggctggagctgcttc | Δ*fabB* | |
| fabB KO R2 | gtaacgtcggatgcgacgctggcgcgtctactccgacctactgcgaaTTAcatatgaatatcctccttag | Δ*fabB* | |
| frdBC KF1 | ATGGCTGAGATGAAAAACCTGAAAATTGAGGTGGTGCGCTATAACgtgtaggctggagctgcttc | Δ*frdBC* [*2*] | |
| frdBC KR1 | TTACCAGTACAGGGCAACAAACAGGATTACGATGGTGGCAACCACcatatgaatatcctccttag | Δ*frdBC* [*2*] | |
| poxB KF1 | CAGATGAACTAAACTTGTTACCGTTATCACATTCAGGAGATGGAGAACCgtgtaggctggagctgcttc | Δ*poxB* from promoter | |
| poxB KR1 | CATGGCATGTCCTTATTATGACGGGAAATGCCACCCTTTattccggggatccgtcgacc | Δ*poxB* from promoter | |
| poxB F1 | AGTGGTTTCCGGTGAATATACGGTGAGCAGCAC | Δ*poxB* verification | |
| poxB R1 | GTTCGCAGTGACTGAGCAGAGCGACCAGGT | Δ*poxB* verification | |
| poxB KF2 | AAACAAACGGTTGCAGCTTATATCGCCAAAACACTCGAATgtgtaggctggagctgcttc | Δ*poxB* | |
| poxB KR2 | CCTTAGCCAGTTTGTTTTCGCCAGTTCGATCACTTCATCAcatatgaatatcctccttag | Δ*poxB* | |
| ackA KF1 | GGCTCCCTGACGTTTTTTTAGCCACGTATCAATTATAGGTACTTCCgtgtaggctggagctgcttc | Δ*ackA-pta* from promoter | |
| pta KR1 | CCGCAGCGCAAAGCTGCGGATGATGACGAGAattccggggatccgtcgacc | Δ*ackA-pta* from promoter | |
| ackA F1 | ATACCCACTATCAGGTATCCTTTAGCAGCCTGAAGGCC | Δ*ackA-pta* verification | |
| pta R1 | TCACCAACGTATCGGGCATTGCCCATCTTC | Δ*ackA-pta* verification | |
| ackA KF2 | ATGTCGAGTAAGTTAGTACTGGTTCTGAACTGCGGTAGTTCTTCAgtgtaggctggagctgcttc | Δ*ackA* [*3*] | |
| ackA KR2 | TCAGGCAGTCAGGCGGCTCGCGTCTTGCGCGATAACCAGTTCTTCcatatgaatatcctccttag | Δ*ackA* [*3*] | |
| pta KR2 | TGTGCAGACTGAATCGCAGTCAGCGCGATGGTGTAGACGAcatatgaatatcctccttag | Δ*ackA-pta* from promoter | |
| ackA F2 | ttcaaaacattttgtcttccatacccactatcaggtatcctttagcagcctgaaggcct | Δ*ackA-pta* verification | |
| pta R2 | ttgttggtttggattcagtgattgcggacatagcgcaaatattcccttgcacaaaacaa | Δ*ackA-pta* verification | |

**Gene and plasmid construction.** Plasmid construction was carried out using standard molecular biology techniques using the Gibson protocol [*4*] and Golden Gate Assembly [*5*]. PCR amplifications were carried out with Q5 DNA polymerase or Phusion DNA polymerase, following manufacturer instructions. Primer sequences are listed in *Table S1B*. Constructs were verified by sequencing (Quintara Biosciences; Berkeley, CA). Synthetic genes were assembled using gBlock sequences (*Table S1D*). gBlocks **were resuspended at** 10 ng/μL in 10 mM Tris-HCl, pH 8.5 and used directly for assembly of vectors.

*Constructs for AdhE2 biochemical characterization*

*pCWori-strep.adhe2.* The synthetic gene for AdhE2 was amplified from pCWori-ter.adhE2 using the adhE2 F6 and adhE2 R1 and inserted into the NdeI-HindIII of pCWOri.

*pCWori-strep.adhe2(C244A)*. The synthetic gene for AdhE2 was amplified by SOE-PCR with primers adhE2 F6/adhE2 R9 (5'-end), adhE2 F9/adhE2 R1 (3'-end) and inserted into the NdeI-HindIII site of pCWori.

*pCWori-strep.adhe2(H721A/H735A)*. The pCWori-strep.adhE2(H735A) cloning intermediate was made by SOE-PCR with adhE2 F6/adhE2 R9 (5'-end), adhE2 F9/adhE2 R1 (3'-end) and inserted into the NdeI-HindIII sites of pCWori. pCWori-strep.adhE2(H735A, H721A) was made by SOE-PCR amplified from pCWOri-strep.adhE2(H735A) with adhE2 F6/adhE2 R11 (5'-end), adhE2 F11/adhE2 R1 (3'-end) and inserted into the NdeI-HindIII sites of pCWori.

*pCWori-strep.aldh* The sequence coding residues M1-N448 of the synthetic *adhE2* gene were amplified using primers adhE2 F6 and adhE2 AADH R1 from pCWori-ter.adhE2 and inserted into the NdeI-HindII site of pCWori.

*Constructs for the front end of the pathway*

*pBT33-Bu1.* Contains the phaA.phaB operon driven by the *P*_BAD_ promoter and the *crt* gene driven by the *P*_Trc_ promoter in the pBAD33 backbone [*6*].

*pT5T33-phaA.phaB-crt.* Contains the phaA.phaB operon driven by the *P*_T5_ promoter and the *crt* gene driven by the *P*_Trc_ promoter in the pBAD33 backbone [*6*].

*pT533-phaA.phaB.* The oligos trc.crt delete GO1 and trc.crt delete GO2 were inserted into the XbaI restriction site of pT5T33-phaA.phaB-crt to remove the *crt* gene and generate pT533-phaA.phaB.

*pT533-phaA.* The *phaA* gene was amplified from pT533-phaA.phaB using primers phaA GF1 and phaA GR1 and inserted into pT533-phaA.phaB cut with NdeI and XbaI using Gibson assembly.

*pBT33-Bu2.* Contains the *phaA.hbd* operon driven by the *P*_BAD_ promoter and the *crt* gene driven by the *P*_Trc_ promoter in the pBAD33 backbone [*6*]

*pT5T33-phaA.hbd-crt*. Contains the *phaA.hbd* operon driven by the *P*_T5_ promoter and the *crt* gene driven by the *P*_Trc_ promoter in the pBAD33 backbone [*6*]

*pTT33-phaA.hbd-crt*. Contains the *phaA.hbd* operon driven by the *P*_Trc_ promoter and the *crt* gene driven by the *P*_Trc_ promoter in the pBAD33 backbone [*6*].

*pT533-phaA.hbd.* The oligos trc.crt delete GO1 and trc.crt delete GO2 were inserted into the XbaI restriction site of pT5T33-phaA.HBD-crt to remove the *crt* gene and generate pT533-phaA.HBD.

*pBT-0.03HBD*. This plasmid was made with the parent plasmid pBT33-Bu2 with the 30K RBS (CGACTACGGGTAATTCGGAACGACAGTCGCCCCAAA) in front of the *hbd* gene.

*pBT-0.3crt*. This plasmid was made with the parent plasmid pBT33-Bu2 with the 300K RBS (GAGTCGAGAGGCTGCACACAAGACAGACCGAGTC) in front of the *crt* gene.

*PDHc overexpression*

*pBBR1-aceEF.lpd.* Contains the *aceEF-lpd* operon driven by the *P*_Lac_ promoter [*6*].

*AdhE2 screening*

*pCDF3-ter.adhE2.* The pCDF3 cloning intermediate was constructed by QuickChange mutagenesis of pCDFDuet to modify the origin using CloDF13-cop3 QCF1/QCR1 primers. The Sp resistance marker and CloDF13cop3 origin were amplified using aadA CloDF F1/R1 primers and inserted into the Bsu36I and HindIII cut sites of pCWori-ter.adhE2 to replace the origin and resistance markers of pCWori.

*pCDF3-ter.adhE2.1-96*. AdhE2 variant containing plasmids (*Table S1.D1*) were constructed by Calysta Inc. (Menlo Park) using pCDF3-ter.adhE2 as the parent plasmid.

*pCDF3-ter.adhE29-45*. AdhE2 homologs containing plasmids (*Table S1.D2*) were constructed by Calysta Inc. (Menlo Park) using pCDF3-ter.adhE2 as the parent plasmid.

*ALDH screening*

*pCDF3-ter.aldh1-15,46.* Monofunctional ALDH containing plasmids were constructed by Calysta Inc. (Menlo Park) using pCDF3-ter.adhE2 as the parent plasmid. The *aldhX* gene (*Table S1.D3*) was inserted upstream of the ADH domain of AdhE2, separated by a stop codon, leaving some residual expression of the ADH domain.

*pCDF3-aldh1-15,46.* (*Table S1.B1*) The oligos ter delete GO5 and ter delete GO6 were inserted into the BamHI-EcoRI restriction sites of pCDF3-ter.aldh1-15,46 to remove the *ter* gene and generate pCDF3-aldh1-15,46.

*pCWori-ter.adhE2*. Contians the *ter.adhE2* operon driven by a double *P*_Tac_. It also contains the *lacIq* gene [*6*].

*pCWori-ter-aldh46.adh.* The *aldh46* gene was amplified from pCDF3-ter.aldh46 using the aldh46 GF3 and aldh46.ADH GR1 primers. The ADH domain of adhE2 was amplified from pCWori-ter.adhE2 using primers aldh46.ADH GF1 and ADH GR1. The two amplicons were inserted into the EcoRI and HindIII cut pCWori-ter.adhE2 using Gibson assembly.

*pCWori.trc-ter-*adhE2. The *rrnb* terminator and *P*_trc_ promoter from pBT33-Bu2 was amplified with primers pCWoriTrc F1/R1 and inserted between the *ter* and *adhE2* genes at the EcoRI site of pCWori-ter.adhE2 using Gibson assembly.

*pCWori.trc-ter-aldh46.adh.* The *aldh46* gene was amplified from pCDF3-ter.aldh46 using primers aldh GF1 and aldh46.ADH GR1 primers. The sequence encoding the ADH domain of AdhE2 was amplified from pCWori-ter.adhE2 using primers aldh46.ADH GF1 and ADH GR1. The two amplicons were inserted into EcoRI and HindII digested pCWori.trc-ter-adhE2 using Gibson assembly.

*ADH screening:*

*pCWori.trc-ter-aldh46.adhX.* The cloning intermediate pCWOri.trc-ter-aldh46 was constructed by Gibson assembly with the aldh46.x G1 gBlock and HindIII digested pCWori.trc-ter-aldh46.adh. gBlocks containing *adh/dhaT* genes (*Table S1.E1)* were inserted into SpeI/XbaI digested pCWori.trc-ter-aldh46 using Gibson assembly.

*ALDH.ADH screening*

*ALDH.ADH (X.Y) combinations 3.12, 6.8, 7.2, 7.8, 7.12, 9.8, 9.12.* These plasmids were made by Gibson assembly using the pCWori.trc-ter-aldh46.adhY plasmid digested with EcoRI and XbaI to remove the *aldh46* gene and generate the backbone. Inserts were obtained by PCR amplification using primers aldhX GF2 and aldhX.adhY GR1 from pCDF3-aldhX. For the plasmid containing *aldh7.adh2*, the extra oligos aldh7.adh2 GO1 and aldh7.adh2 GO2 were also included to reinsert the 5’ end of *adh*2 removed by EcoRI digestion.

*ALDH.ADH (X.Y) combinations 3.2, 3.8, 6.2, 6.12, 9.2, 14.2, 14.12, 14.8.* These plasmids were constructed by Golden Gate assembly. *aldhX* was amplified with primers aldhX GGF1 and aldhX.(adhY) GGR1 from pCDF3-aldhX. *adhY* was amplified with (aldhX).adhY GGF1 and adhY GGR1 from pCWori.trc-ter-aldh46.adhY and inserted into pCWori.trc-ter-RFP digested with BsaI.

*ALDH.ADH combinations 3.15, 3.17, 3.18, 3.19, 3.21, 3.22, 7.15, 7.18, 7.19, 7.20, 7.21, 7.22, 46.19, 46.20, 46.21*. The appropriate *aldh* gene was amplified with primers aldh3 GGF1 or aldh round 1 library GGF1 and aldh3/7/46.(adh) GGR1 from pCDF3-aldh3/7/46 (see *Table S1.B3*)*.* The adh gene was amplified with adh15-22 GGF1 and adh15-22 GGR1 from the appropriate gBlock (*Table S1.E1)*. The *aldh* and *adh* genes were inserted into pCWori.trc-ter-RFP digested with BsaI via Golden Gate assembly.

*pCWori.trc-ter-RFP.* Prior to inserting the BsaI-RFP-BsaI Golden Gate cassette, two BsaI cut sites on the backbone of pCWori.trc-ter-aldh46.adh2 were removed. The first BsaI cut site in the *rrnB* terminator was removed by amplifying the regions of the *rrnB* terminator flanking the cut site with primers rrnB-1 GF1/GR1 and rnnB-2 GF1/GR1 and inserting into pCWori.trc-ter-aldh46.adh2 digested with XhoI and SfiI using Gibson assembly. The second BsaI site was removed by inserting oligos BsaI delete GO1 and BsaI delete GO2 into the BsaI cut site using Gibson assembly. The gene for RFP was amplified from pET28a(gg)-RFP with RFP BsaI GF2 and RFP BsaI GR2 and inserted into the BsaI removed pCWori.trc-ter-aldh46.adh2 cut with EcoRI and SpeI using Gibson assembly.

*sADH screening*

*pCWori.trc-sadhX-aldh7.adh2.* The *sadhX* gene was amplified from the corresponding gBlock (*Table S1.E2*) using primers sadhX GF1 and sadhX GR1 and inserted into pCWori.trc-ter-aldh7.adh2 cut with NdeI and XhoI.

*Constructs for chromosomal mutations*. The pCRISPR-Gibson1 plasmids were constructed to clone constructs with specific guide sequence to target the *E. coli* chromosome for introduction of point mutants. The parent plasmid, pCRISPR-Gibson1 (#2786), was generated from pCRISPR (Addgene 42875) to introduce cut sites between sgRNA promoter and the sgRNA to facilitate the use of Gibson assembly to introduce guide sequences for the target DNA. All guide sequences were generated using the Benchling CRISPR tool (see *Table S1.C* for guide sequences).

pCRISPR-PcnB2409 (#2784) was constructed by insertion of the annealed oligonucleotides, P1155 and P1156, and inserted into the XbaI-HindIII site of pCRISPR-Gibson1 using the Gibson protocol.

pCRISPR-RpoC2406 (#2794) was constructed by insertion of the annealed oligonucleotides, P1232 and P1233, and inserted into the XbaI-HindIII site of pCRISPR-Gibson1 using the Gibson protocol.

**Expression and purification of AdhE2 constructs.** TB (1 L) containing carbenicillin (50 µg/mL) in a 2.8 L Fernbach baffled shake flask was inoculated to OD_600_ = 0.05 with an overnight TB culture of freshly transformed *E. coli* containing the appropriate overexpression plasmid. The cultures were grown at 37 °C at 200 rpm to OD_600_ = 0.6 to 0.8 at which point cultures were cooled on ice for 20 min, followed by induction of protein expression with 1 mM IPTG and overnight growth at 16 °C. Cell pellets were harvested by centrifugation at 9,800 × g for 7 min and resuspended at 20 mL/L of culture with Buffer W (100 mM Tris-HCl, 150 mM sodium chloride, 1 mM EDTA, pH 8.0) supplemented with 2 mg/mL lysozyme and 2 µL/50 mL final volume Benzonase and frozen at –80 °C.

Frozen cell suspensions were thawed and frozen twice before finally thawing and adding 0.5 mM PMSF as a 50 mM stock solution in ethanol dropwise. The cell suspension was lysed with a Misonix 3000 probe sonicator at full power with a 15 second on, 60 second off cycle for a total sonication time of 2.5 min on ice. The lysate was centrifuged at 15,300 × g for 20 min at 4 °C to separate the soluble and insoluble fractions. DNA was precipitated in the soluble fraction by addition of 0.5% polyethylenimine as a 15% *v*/*v* stock solution added dropwise. The precipitated DNA was removed by centrifugation at 15,300 × g for 20 min at 4 °C. The lysate was loaded onto a Strep-tactin Superflow High Capacity column (IBA, 1 mL resin/L expression culture) by gravity flow. The column was washed with 20 column volumes Buffer W. The protein was then eluted with 2.5 mM desthiobiotin in Buffer W. Fractions containing ALDH protein by A280 were pooled and concentrated in an Amicon Ultra 10,000 MWCO concentrator. Concentrated protein was supplemented with glycerol to 10% *v*/*v* and stored at –80 °C.

**AdhE2 biochemical assays.** AdhE2 activity was measured using three different assays as described below.

*NADH oxidation assay.* Reactions were measured by monitoring the oxidation of NADH at 340 nm at 25 °C. The assay mixture (400 μL in a 1 cm cuvette) contained 100 μM NADH in 100 mM Tris, 1 mM DTT, pH 7.5. The reaction was initiated by the addition of substrate. Addition of either butyryl-CoA or acetyl-CoA monitored the additive reduction of the acyl-CoA substrate in the ALDH active and the aldehyde intermediate in the ADH active site. Addition of either butyraldehyde or acetaldehyde monitors solely the activity of the ADH domain.

*CPM free CoA release assay.* The activity of the ALDH domain was monitored by release of free CoA using the 7-diethylamino-3-(4'-maleimidylphenyl)-4-methylcoumarin (CPM) fluorescence-based thiol quantification assay [*7*] with butyryl-CoA or acetyl-CoA as a substrate. Assay conditions were the same as above except a final assay volume of 200 μL was used and the reaction was initiated by substrate addition. The reaction of CPM with released free CoA was monitored by fluorescence with excitation at 405 nm and emission at 530 nm.

*k*_cat_ and *K*_M_ were determined by fitting to initial rate data with Origin (OriginLab, Northampton, MA) using the equation:

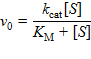

where *v*_o_ is the initial rate and [*S*] is the substrate concentration.

**ALDH domain phylogenetic analysis.** A literature search for ALDH domains of characterized substrate specificity was performed since our biochemical characterization suggested that this domain controls the overall selectivity of acyl-CoA reduction. The set of identified homologs consisted of three C_2_-specific bifunctional AdhE2 homologs, four C_4_-specific monofunctional ALDHs, and one atypical C_2_-specific ALDH (*Figure S5A*). The sequences were all of bacterial origin except for an AdhE2 homolog identified from the protozoan parasite *Giardia intestinalis*. These sequences were then placed in a phylogenetic tree such that the branching pattern was biased by their characterized substrate preference. Next, the entire ALDH gene family was assembled into a second phylogenetic tree with the branching pattern again biased by the characterized substrate specificity from the first tree. The full family tree comprised of more than 1,200 sequences, of which approximately 33% were derived from bifunctional ALDH domains.

**Production of C_4_ compounds in shake flasks.** Overnight cultures of freshly transformed *E. coli* strains were grown for 12–16 h in TB at 37 °C and used to inoculate TB (50 ml) with glucose replacing the standard glycerol supplement (1.5% (*w*/*v*) glucose for aerobic cultures and 2.5% (w/v) glucose for anaerobic cultures) and appropriate antibiotics to an optical density at 600 nm OD_600_ of 0.05 in a 250 mL-baffled flask (Kimble Glass; Chicago, IL) or a 250 mL-baffled anaerobic flask with GL45 threaded top (Chemglass). The cultures were grown at 37 °C in a rotary shaker (200 rpm) and induced with IPTG (1.0 mM) at OD_600_ = 0.35–0.45. The growth temperature was then reduced to 30 °C, and the culture flasks were sealed with Parafilm M (Pechiney Plastic Packaging) to prevent product evaporation for aerobic cultures. Anaerobic cultures were sealed and the headspace was sparged with argon for 3 min immediately follow induction. Aerobic cultures were unsealed for 10 to 30 min every 24 h then resealed with Parafilm M, and additional glucose (1% (*w/v*)) was added 1 day post-induction. Samples were quantified after 3 d of cell culture, unless otherwise noted. For cultures grown with an oleyl alcohol layer, cultures (40 mL) were grown at 37 °C for 3 h before induction with IPTG (1 mM). Oleyl alcohol (10 mL) was then added. Cultures were sealed and the headspace was sparged with argon for 3 min. At this time, the growth temperature was reduced to 30 °C. Cultures were grown for 5 d before harvesting. Both the aqueous and organic layers were reserved for quantification.

**Quantification of *n-*butanol titers.** Samples (2 mL) were removed from cell culture and cleared of biomass by centrifugation at 20,817 × *g* for 2 min using an Eppendorf 5417R centrifuge. The supernatant or cleared medium sample was then mixed in a 9:1 ratio with an aqueous solution containing the hexanol internal standard (10 g L^−1^). These samples were then analyzed on a Trace GC Ultra (Thermo Scientific) using an HP-5MS column (0.25 mm × 30 m, 0.25 μM film thickness, J & W Scientific). The oven program was as follows: 75 °C for 3 min, ramp to 300 °C at 45 °C min^−1^, 300 °C for 1 min. Alcohols were quantified by flame ionization detection (FID) (flow: 350 mL min^−1^ air, 35 mL min^−1^ H_2_ and 30 mL min^−1^ helium). Samples containing *n*-butanol levels below 500 mg L^−1^ were requantified after extraction of the cleared medium sample or standard (500 μL) with toluene (500 μL) containing the isobutanol internal standard (100 mg L^−1^) using a Digital Vortex Mixer (Fisher) for 5 min set at 2,000. The organic layer was then quantified using the same GC parameters with a DSQII single-quadrupole mass spectrometer (Thermo Scientific) using single-ion monitoring (*m/z* 41 and 56) concurrent with full scan mode (*m/z* 35–80). Samples were quantified relative to a standard curve of 2, 4, 8, 16, 31, 63, 125, 250, 500 mg L^−1^ *n*-butanol for MS detection or 125, 250, 500, 1,000, 2,000, 4,000, 8,000 mg L^−1^ *n*-butanol/ethanol for FID detection. Standard curves were prepared freshly during each run and normalized for injection volume using the internal isobutanol standard (100 or 1,000 mg L^−1^ for MS and FID, respectively). Standard curve was normalized for injection volume using the internal standard.

**Quantification of 1,3-butanediol (BDO) and 4-hydroxy-2-butanone (HB) titers.** Samples (2 mL) were removed from cell culture and cleared of biomass by centrifugation at 20,817 × *g* for 2 min using an Eppendorf 5417R centrifuge. The cleared medium samples, or standards prepared in TB medium, diluted 1:1000 into water, and filtered through a 0.22 µm filter (EMD Millipore MSGVN2210). Supernatants were diluted 1- to 1,000-fold with water containing 2,4-pentanediol (10 μM) added as internal standard and analyzed on an Agilent 1290 HPLC using a Rezex ROA-Organic Acid H^+^ (8%) column (150 × 4.6 mm, Phenomenex) with isocratic elution (0.5% *v/v* formic acid, 0.6 mL min^−1^, 55 °C). Samples were detected with an Agilent 6460C triple quadrupole MS with Jet Stream ESI source, operating in positive MRM mode (*m/z* 91→73 transition; fragmentor, 50 V; collision energy, 0 V; cell accelerator voltage, 7 V; delta EMV, +400). Samples were quantified relative to a standard curve of 0.3125, 0.625, 1.25, 2.5, 5, 10 g L^−1^ 1,3-butanediol and 4-hydroxy-2-butanone.

**Anaerobic growth competition and enrichment validation.** DH1Δ*5* transformed with *n*-butanol production plasmids capable of a range of titers were mixed at various ratios and cultured anaerobically as described above. Flasks were sampled with a syringe to collect culture media supernatants for quantification of metabolites and to measure growth. Pelleted cells were used as template for qPCR of *n*-butanol plasmids to determine the relative abundance of different subpopulations and compared to a standard curve of purified plasmids. The qPCR reactions were performed using Bio-Rad Sybr-Green mastermix according to the manufacturer protocol, and OD-normalized boiled cell pellet was used as template.

**Adaptive evolution.** Host strains were transformed with appropriate synthetic pathways and plated on LB agar plate with appropriate antibiotics over night at 37 °C. Colonies were picked and grown in 5 ml TB media with 2.5% (*w/v*) glucose replaced with the standard glycerol carbon source overnight at 37 °C at 200 rpm. Overnight cultures were then inoculated to fresh 30 mL TB media with 2.5% glucose with initial OD_600_ of 0.05 and grown at 37 °C at 200 rpm. Once cultures reached OD_600_ ~0.3–0.4, cultures were induced with 1 mM IPTG and sparged with argon for 3 min. Growth temperature was then lowered to 30 °C. Cultures were then serially transferred to fresh media every 24–72 h to an initial OD_600_ of 0.05 to approximate continuous growth with limited time spent in stationary phase. The growth time of 24­72 h was chosen such that the cultures would be in late-log or early-stationary phase. Growth media was TB with 2.5% glucose, 1 mM IPTG, and appropriate antibiotics. Culture OD_600_ was monitored daily and cultures were transferred when the majority of cultures were in late log-phase growth, usually OD_600_ 1.5–2.0. Culture supernatant samples (2 mL) were collected for metabolite quantification. All cultures were transferred simultaneously, the headspace was sparged with argon for 3 min, and growth was continued at 30 °C in a rotary shaker (200 rpm). Selections were continued (from three weeks or three months) until no improved strains were isolated from the culture. Final cultures were stored as 15% glycerol stocks at –80 °C in addition to being streaked on LB agar plates. Individual colonies were picked and cultured for metabolite production in TB to confirm butanediol, hydroxybutanone, and butanol production relative to wild type strains.

**Genome sequencing.** Cells were grown on 10 mL LB media with 2.5% (*w/v*) glucose with appropriate antibiotics overnight at 37 °C. Cell were then spun down at 8000 × *g* in a Beckman centrifuge. Cell pellets were then processed using the Qiagen Genomic DNA Isolation Kit according to manufacturer specifications. Genomic libraries were then prepared for sequencing using the Kapa Biosystem Hyper Plus Kit with no modifications to the standard protocol. For each library, 1 μg of genomic DNA was used with 3 μl of adapter (40 μM) per ligation. A double-sided selection to obtain 600 bp fragments was then performed using 0.55 vol of right and 0.6 vol of left Ampure XP beads (Beckman Coulter). No PCR amplification was carried out after the size selection. Libraries were sequenced at the UC Davis DNA Core Facility with PE300 sequencing using an Illumina MiSeq. Sequencing results were mapped against the the *E. coli* genome (DH1- Accession ID: NC_017625, BW25113 - Accession ID: NZ_CP009273) and compared against reads obtained from our DH1Δ*5* or BW25113Δ*5* parent strain using Breseq v. 0.25d [*8*].

**Cell lysate enzyme assays.** Biomass was harvested at the end of production and stored at –80 °C. Frozen cell pellets (from 2 mL culture) were thawed and resuspended in 500 µL of 100 mM Tris-HCl pH 7.5 containing DTT (5 mM) and PMSF (0.5 mM).

*PhaA.* Thiolysis activity was measured by monitoring the enolate form of acetoacetyl CoA as previously described [9]. Assays were performed at 30 °C in a 96 well plate in a total volume of 100 µL containing 100 mM Tris-HCl, pH 7.5, 10 mM MgCl_2_, 1 mM DTT, 10 µM CoA, and 20 µM acetoacetyl CoA.

*Hbd, Ter, Aldh, and Adh* activities were assayed as described [*6*]. Briefly, all assays were perfomed at 30 °C in a 96 well plate in a total volume of 100 µL. The mixture for the Hbd assays contained 100 mM Tris-HCl, pH 7.5, 100 µM acetoacetyl CoA, 100 µM NADH. The Hbd activity was monitored by the oxidation of NADH at 340 nm. The mixture for the Ter assays contained 100 mM Tris-HCl, pH 7.5, 100 µM NADH, and 50 µM crotonyl CoA. The Ter activity was monitored by the oxidation of NADH at 340 nm. The mixture to assay the aldehyde domain of AdhE2 assays contained 100 mM Tris-HCl, pH 7.5, 0.5 mM DTT, 400 µM NAD^+^, 400 µM CoA, and 10 mM butyraldehyde. The acitivity of the aldehyde domain was monitored by the reduction of NAD^+^ at 340 nm. The mixture to assay the alcohol domain of AdhE2 contained 100 mM Tris-HCl, pH 7.5, 0.5 mM DTT, 400 µM NADH, and 10 mM butyraldehyde. The activity of the alcohol domain was monitored by the oxidation of NADH at 340 nm.

**RNA sequencing and analysis.** Cells with synthetic pathways were harvested after 24 h post induction with IPTG for RNA isolation. RNA was isolated using the RNeasy RNA isolation kit (Qiagen). In house rRNA removal method was used to remove rRNA before sequencing. 5 μg of total RNA was treated with 4.5 μL of TURBO DNase (ThermoFischer) in a 50 μL reaction including 5 μL of 10× buffer to remove genomic DNA. The reaction was incubated at 37 °C for 30 min. The reaction was diluted with 100 μL of Bufffer RLT and 200 μL of 70% ethanol and transferred to an RNeasy column (Qiagen) for RNA cleanup following the manufacture instructions. 1 μg of DNase treated RNA was combined with 1 μL of 0.5 μM DNA probes (see *Table* *S1E*) with Hybridization buffer (200 mM NaCl, 100 mM Tris-HCl pH 7.5) up to 20 μL. Hybridization of oligos occurred by holding at 95 °C for 2 min, followed by a gradient to 45 °C at –0.1 °C/s. 5U of RNase H (Epicentre) in 2.5 μL of 10× Digestion buffer (0.5 M Tris-HCl pH 7.5, 1 M NaCl, 200 mM MgCl_2_ ) were added, and the resulting mixture was incubated at 45 °C for 30 min. Following cleanup with the Qiagen RNeasy Kit, the sample was treated with 3 U of TURBO DNase. Finally, the Qiagen RNeasy Kit was used to clean up samples one last time before RNA-Seq library prep. RNA-Seq libraries were prepared using the Illumina TruSeq RNA Sample Prep Kit. Samples were sequenced with Illumina HiSeq4000 at UC Davis DNA core. Reads were mapped using the Kallisto [*10*] and Sleuth [*11*]. Functional enrichment analysis of differentially expressed genes is based on clusters of orthologous groups (COG) categories provided by the IMG-ER annotations [*12*].

**Generation of chromosomal point mutations.** Point mutations were made using the CRISPR Cas9 system [*13,14*]. Briefly, cells were transformed with the pKD46-Cas9-RecA-Cure, which allows for the expression of the Cas9 protein for double stranded DNA breaks and the RecA protein to assist homologous recombination. A single transformant was picked and inoculated in liquid culture to make electro-competent cells. Then cells that carried the pKD46-Cas9-RecA-Cure plasmid were transformed with both the pCRISPR plasmid containing the specific guide RNA and the double stranded DNA repair fragment that carried the desired sequence. The repair fragment also carries a silent mutation to remove the PAM site and a phosphatioate modification at both the 5′- and 3′-ends. Transformations were recovered and plated on plates with appropriate selection markers. Colonies were validated by Sanger sequencing.

DH1Δ*5* _2406_*pcnB*(R149L) - CGC 🡪 CTC mutation at position 446 that corresponds to the *pcnB*(R149L) mutation was made in the strain DH1Δ5 using the CRISPR Cas9 system. DH1Δ*5* was transformed with pKD46-Cas9-RecA-Cure and plated on appropriate antibiotic resistant LB agar platse and incubated at 30 °C overnight. A single colony was picked and inoculated in 10 mL LB liquid media with appropriate antibiotics overnight at 30 °C. The overnight culture was then diluted in fresh LB media with 0.2% *w/v* of arabinose to an OD_600_ ~ 0.01 to induce RecA expression. Once cultures reached an OD_600_ of 0.4,cells were harvested to make electro-competent cells. DH1_ pKD46-Cas9-RecA-Cure electro-competent cells were then transformed with pCRISPR_gibson_1guide_2409pcnB (#2784) plasmid and repair fragments (P1227_2406_pcnB RF_R and P1226_2406_pcnB RF_F). Cells were recovered at 30 °C for 1.5 h and plated on appropriate antibiotic selection LB agar plate. Plate was incubated at 30 °C overnight. Colonies were picked and validated by Sanger sequencing (Quintara Biosciences). Once sequences were confirmed, a single colony was inoculated in 10 mL LB media with 0.05 mM IPTG to induce the guide to target and cure the pCRISPR_gibson_1guide_2409pcnB (#2784) plasmid. Once the pCRISPR_gibson_1guide_2409pcnB (#2784) plasmid was cured, cells were grown at 37 °C to cure the pKD46-Cas9-RecA-Cure plasmid, which contains a temperature sensitive origin of replication.

DH1Δ*5* _2406_*rpoC*(M466L) – ATG 🡪CTG mutation at position 1396 that corresponds to the *rpoC*(M466L) mutation was made in the strain DH1Δ5 using the CRISPR Cas9 system as described above. The pCRISPR_gibson_1guide_2406_rpoC (#2794) plasmid and repair fragments (P1231_2406_rpoC_RF_R and P1230_2406_rpoC_RF_F) were used.

DH1Δ*5* _2406_*pcnB*(R149L)_*rpoC*(M466L) – the double mutant was made using the CRISPR Cas9 system as described above in a sequencial manner. Once the pcnB(R149L) mutation was confirmed and the pCRISPR_gibson_1guide_2409pcnB (#2784) plasmid was cured, cells were grown up to make electro-compotent cells. Cells were then transformed with the pCRISPR_gibson_1guide_2406_rpoC (#2794) construct and repair fragments (P1231_2406_rpoC_RF_R and P1230_2406_rpoC_RF_F). Once the desired mutations were confirmed with sequencing, cells were growing in IPTG-containing media to cure the pCRISPR_gibson_1guide_2406_rpoC (#2794) plasmid. Finally, cells were grown at 37 °C to cure the pKD46-Cas9-RecA-Cure plasmid, which contains a temperature sensitive origin of replication.

**Metabolomics.** Five replicates of cultures were grown as described previously for production in shake flasks and harvested 24 h after induction. Cultures were centrifuged at 20,817 × g for 1 min at 4 °C with an Eppendorf 5417R Centrifuge (Hamburg, Germany). The supernatants were decanted immediately and cell pellets were flash frozen with liquid nitrogen and stored at –80 °C until extraction. Pellets were extracted with 90% *v/v* methanol with 0.1% *v/v* formic acid containing D_3_^15^N-serine (0.01 mg/mL; Cambridge Isotope Laboratories, Inc., DNLM6863) to a final concentration of 1 mg biomass/μL of extraction buffer. The mass of the biomass was calculated using the standard value for *E. coli* of 23.8 mg/OD_600_. Samples were vortexed for 15 s, incubated at –80 °C for 30 min, and then thawed at –20 °C for 30 min. The vortex-freeze-thaw cycle was repeated for a total of five times. At the end of this procedure, the lysed cells were centrifuged at 20,817 × g at 4 °C for 5 min and the supernatant collected for LC-MS/MS analysis. Metabolomic analyses were performed as previously reported [*15*]. Briefly, supernatant was collected and frozen at –80 °C until analysis. 20 μL of supernatant was analyzed by single-reaction monitoring (SRM)-based targeted LC-MS/MS. Metabolite separation was performed using normal-phase chromatography using a Luna-5 mm NH_2_ column (50 mm × 4.60 mm, Phenomenex). Mobile phases were run as follows: Buffer A, acetonitrile; Buffer B, 95:5 water/ acetonitrile with 0.1% formic acid or 0.2% ammonium hydroxide with 50 mM ammonium acetate for positive and negative ionization modes, respectively. Flow rate began at 0.2 mL/min for 2 min, followed by a gradient starting at 0 %B and increasing linearly to 100 %B over the course of 13 min with a flow rate of 0.7 mL/min, followed by an isocratic gradient of 100 %B for 10 min before equilibrating for 5 min at 0 %B with a flow rate of 0.7 mL/min. MS analysis was performed using an electrospray ionization (ESI) source on an Agilent 6430 QQQ LC-MS/MS.Drying gas temperature 350 °C, drying gas flow rate was 10 L/min, the nebulizer pressure was 35 psi, capillary voltage was 3.0 kV, and fragmentor voltage was 100 V. Representative metabolites were quantified by SRM of the transitions from precursor to product ions at associated collision energies. Data was analyzed using Agilent Qualitative Analysis software by calculating area under the curve.

**Supplementary Results**

**Table S1.** Strains, oligonucleotides, and sequences. (A) *E. coli* strains. (B) Plasmids and gene sequences (C). Oligos for sequencing and cloning (D) aldehyde dehydrogenases sequences. (E) gBlock sequences. (F) Sequence for DNA probes for rRNA depleteion. (G) Open reading frame for *pncB* and *rpoC*

**A. *E. coli* strains**

A1. Parent and constructed strains

| **Organism** | **Name** | **Description** | **Source** |
| --- | --- | --- | --- |
| *E. coli* | DH10B | F- endA1 recA1 galE15 galK16 nupG rpsL ΔlacX74 Φ80lacZΔM15 araD139 Δ(ara,leu)7697 mcrA Δ(mrr-hsdRMS-mcrBC) λ- | Invitrogen |
| *E. coli* | BL21 (DE3) Star T1^R^ | RNaseE mutation to increase mRNA stability, Δ*fhuA* | Invitrogen |
| *E. coli* | DH1Δ*5* | DH1 Δ*ackA-pta* Δ*adhE* Δ*ldhA* Δ*poxB* Δ*frdBC* | This study |
| *E. coli* | BW25113Δ*5*-T1^R^ | BW25113 Δ*ackA-pta* Δ*adhE* Δ*ldhA* Δ*poxB* Δ*frdBC* ∆*fhuA*, P1 transduced *fhuA*:Km^R^ from 1637 parent to 1435 then recycled Km marker | This study |
| *E. coli* | DH1Δ5_2406_pcnB(R149L) | DH1 Δ*ackA-pta* Δ*adhE* Δ*ldhA* Δ*poxB* Δ*frdBC* *pcnB*(R19L) | This study |
| *E. coli* | DH1Δ5_2406_rpoC(M466L) | DH1 Δ*ackA-pta* Δ*adhE* Δ*ldhA* Δ*poxB* Δ*frdBC* *rpoC*(M466L) | This study |
| *E. coli* | DH1Δ5_2406_pcnB(R149L)_rpoC(M466L) | DH1 Δ*ackA-pta* Δ*adhE* Δ*ldhA* Δ*poxB* Δ*frdBC* *pcnB*(R19L) *rpoC*(M466L) | This study |

*A2. Strains isolated from evolutions. All strains contained the pBBR1-AceEF.Lpd plasmid (#339) for overexpression of the pyruvate dehydrogenase complex. Parent strains for n-butanol production contained the pT5T33-Bu2 plasmid (#499) and one of the following three plasmids: pCWori.trc-ter-aldh46.adh2 (#1866), pCWori.trc-ter-aldh46.adh8 (#1867), or pCWori.trc-aldh21.adh2 (#2456). The parent strain for HB production contained the pT533-phaA (#2080) and pCWori.trc-aldh7.adh2 (#2076) plasmids. The parent strain for BDO production contained the pT533-phaA.phaB (#1319) plasmid and one of the following three plasmids: pCWori.trc-aldh7.adh2 (#2076), pCWori.trc-aldh3.adh22 (#2468), or pCWori.trc-sadh1-aldh7.adh2 (#2430). An identifier number was used during isolation of individual clones from an evolution experiment consisting of plasmid combination (A, B, C, D), flask number-dilution number-clone.*

| **Product** | **Parent** | **Plasmids** | **Media** | **Identifier** | **No.** |
| --- | --- | --- | --- | --- | --- |
| *n*-Butanol | DH1Δ*5* | 339-499-1866 | LB/2.5% Glc | A1-D45-2 | 2616 |
|  | DH1Δ*5* | 339-499-1866 | LB/2.5% Glc | A3-D26-2 | 2619 |
|  | DH1Δ*5* | 339-499-1866 | LB/2.5% Glc | A3-D26-3 | 2620 |
|  | DH1Δ*5* | 339-499-1866 | LB/2.5% Glc | A3-D35-1 | 2621 |
|  | DH1Δ*5* | 339-499-1866 | LB/2.5% Glc | A3-D35-2 | 2622 |
|  | DH1Δ*5* | 339-499-2456 | LB/2.5% Glc | C1-D41-1 | 2625 |
|  | DH1Δ*5* | 339-499-2456 | LB/2.5% Glc | C1-D41-3 | 2626 |
|  | DH1Δ*5* | 339-499-2456 | LB/2.5% Glc | C3-D35-1 | 2628 |
|  | DH1Δ*5* | 339-499-2456 | LB/2.5% Glc | C3-D41-1 | 2629 |
|  | DH1Δ*5* | 339-499-2456 | LB/2.5% Glc | C3-D41-6 | 2630 |
|  | DH1Δ*5* | 339-499-2456 | M9/10% LB/2.5% Glc | D15-12-1 | 2685 |
|  | DH1Δ*5* | 339-499-2456 | M9/10% LB/2.5% Glc | D15-12-2 | 2686 |
|  | DH1Δ*5* | 339-499-2456 | M9/10% LB/2.5% Glc | D15-12-3 | 2687 |
|  | BW25113Δ*5* | 339-499-2456 | M9/10% LB/2.5% Glc | C1-D4-3 | 2726 |
|  | BW25113Δ*5* | 339-499-2456 | M9/10% LB/2.5% Glc | D4-C3-3 | 2727 |
|  | BW25113Δ*5* | 339-499-1866 | M9/10% LB/2.5% Glc | D17-A3-1 | 2728 |
|  | BW25113Δ*5* | 339-499-1867 | M9/10% LB/2.5% Glc | D17-B3-1 | 2729 |
|  | BW25113Δ*5* | 339-499-2456 | M9/10% LB/2.5% Glc | D17-C1-3 | 2730 |
|  | BW25113Δ*5* | 339-499-2456 | M9/10% LB/2.5% Glc | D17-C3-2 | 2731 |
|  | BW25113Δ*5* | 339-499-2456 | M9/2.5% Glc | C1-D9-1 | 2748 |
|  | DH1Δ5 | 339-499-2456 | M9/2.5% Glc | C1-D11-2 | 2750 |
| HB | DH1Δ*5* | 339-2080-2076 | TB/2.5% Glc | A3-D17-4 | 2403 |
|  | DH1Δ*5* | 339-2080-2076 | TB/2.5% Glc | A3-D26-2 | 2404 |
| BDO | DH1Δ*5* | 339-1319-2076 | TB/2.5% Glc | B1-D17-2 | 2405 |
|  | DH1Δ*5* | 339-1319-2076 | TB/2.5% Glc | B1-D26-3 | 2406 |
|  | DH1Δ*5* | 339-1319-2076 | TB/2.5% Glc | B3-D26-4 | 2407 |
|  | DH1Δ*5* | 339-1319-2430 | TB/2.5% Glc | C1-D17-4 | 2408 |
|  | DH1Δ*5* | 339-1319-2430 | TB/2.5% Glc | C3-D17-3 | 2409 |
|  | DH1Δ*5* | 339-1319-2430 | TB/2.5% Glc | C2-D26-1 | 2410 |
|  | DH1Δ*5* | 339-1319-2430 | TB/2.5% Glc | C3-D26-2 | 2411 |
|  | DH1Δ*5* | 339-1319-2468 | TB/2.5% Glc | D3-D17-2 | 2412 |

**B. Plasmids**

*B1. Monofunctional ALDHs (BDO).* All constructs were constructed using pCDF3-ter.adhE2 as the parent using Gibson assembly. pCDF3-ter.adhE2 contains the *ter* gene driven by two *P*_tac_ promoters, the Sp resistant marker, and the CloDF13cop3 origin. This backbone plasmid was digested with EcoRI and HindIII for *aldhX* insertion. The *aldhX* gene was inserted upstream of the ADH domain of AdhE2, separated by a stop codon, leaving some residual expression of the ADH domain. The pCDF3-ter.aldhX plasmids were used to screen for *n*-butanol production. A second set of plasmids, pCDF3-aldhX, was generated without the *ter* gene to screen for BDO and HB production and are listed below. The oligos ter delete GO5 and ter delete GO6 were inserted into the BamHI-EcoRI restriction sites of pCDF3-ter.aldhX to remove the *ter* gene and generate pCDF3-aldhX.

| **Name** | **Marker** | **Origin** | **Promoter** | **Description** | **Number** |
| --- | --- | --- | --- | --- | --- |
| pCDF3-aldh1 | Sp | CloDF13cop3 | double *P*_Tac_ | Codon-optimized coenzyme A acylating aldehyde dehydrogenase *Clostridium beijerinckii*, from Calysta Biosystems, contains unannotated C-terminal adhE2 fragment after aldh1, GI:4884855 | 1647 |
| pCDF3-aldh2 | Sp | CloDF13cop3 | double *P*_Tac_ | Codon-optimized ethanolamine utilization protein EutE *Escherichia coli* CFT073, from Calysta Biosystems, contains unannotated C-terminal adhE2 fragment after aldh2, GI:26250354 | 1648 |
| pCDF3-aldh3 | Sp | CloDF13cop3 | double *P*_Tac_ | Codon-optimized butyraldehyde dehydrogenase from *Clostridium saccharoperbutylacetonicum* N1-4(HMT), from Calysta Biosystems, contains unannotated C-terminal adhE2 fragment after aldh3, GI:31075383 | 1649 |
| pCDF3-aldh4 | Sp | CloDF13cop3 | double *P*_Tac_ | Codon-optimized ethanolamine utilization protein eutE *Vibrio shilonii* AK1, from Calysta Biosystems, contains unannotated C-terminal adhE2 fragment after aldh4, GI:149190407 | 1650 |
| pCDF3-aldh5 | Sp | CloDF13cop3 | double *P*_Tac_ | Codon-optimized hypothetical protein RUMGNA_01022 from *Ruminococcus gnavus* ATCC 29149, from Calysta Biosystems, contains unannotated C-terminal adhE2 fragment after aldh5, GI:154503198 | 1651 |
| pCDF3-aldh6 | Sp | CloDF13cop3 | double *P*_Tac_ | Codon-optimized hypothetical protein CLOBOL_07248 from *Clostridium bolteae* ATCC BAA-613, from Calysta Biosystems, contains unannotated C-terminal adhE2 fragment after aldh6, GI:160942363 | 1652 |
| pCDF3-aldh7 | Sp | CloDF13cop3 | double *P*_Tac_ | Codon-optimized ethanolamine utilization protein EutE from *Clostridium botulinum* B str. Eklund 17B, from Calysta Biosystems, contains unannotated C-terminal adhE2 fragment after aldh7, GI:187934965 | 1653 |
| pCDF3-aldh8 | Sp | CloDF13cop3 | double *P*_Tac_ | Codon-optimized coenzyme A acylating aldehyde dehydrogenase from *Clostridium saccharobutylicum*, from Calysta Biosystems, contains unannotated C-terminal adhE2 fragment after aldh8, GI:189310620 | 1654 |
| pCDF3-aldh9 | Sp | CloDF13cop3 | double *P*_Tac_ | Codon-optimized ethanolamine utilization protein EutE from *Clostridium botulinum* E1 str. 'BoNT E Beluga', from Calysta Biosystems, contains unannotated C-terminal adhE2 fragment after aldh9, GI:251780016 | 1655 |
| pCDF3-aldh10 | Sp | CloDF13cop3 | double *P*_Tac_ | Codon-optimized aldehyde dehydrogenase from *Clostridium carboxidivorans* P7, from Calysta Biosystems, contains unannotated C-terminal adhE2 fragment after aldh10, GI:255526882 | 1656 |
| pCDF3-aldh11 | Sp | CloDF13cop3 | double *P*_Tac_ | Codon-optimized aldehyde dehydrogenase from *Clostridium saccharolyticum* WM1, from Calysta Biosystems, contains unannotated C-terminal adhE2 fragment after aldh11, GI:302386203 | 1657 |
| pCDF3-aldh12 | Sp | CloDF13cop3 | double *P*_Tac_ | Codon-optimized aldehyde dehydrogenase from *Geobacillus* sp. Y4.1MC1, from Calysta Biosystems, contains unannotated C-terminal adhE2 fragment after aldh12, GI:312110932 | 1658 |
| pCDF3-aldh13 | Sp | CloDF13cop3 | double *P*_Tac_ | Codon-optimized acetaldehyde dehydrogenase (acetylating) from *Clostridium* sp. DL-VIII, from Calysta Biosystems, contains unannotated C-terminal adhE2 fragment after aldh13, GI:359413662 | 1659 |
| pCDF3-aldh14 | Sp | CloDF13cop3 | double *P*_Tac_ | Codon-optimized hypothetical protein HMPREF9942_01197 from *Fusobacterium nucleatum* subsp. *animalis* F0419, from Calysta Biosystems, contains unannotated C-terminal adhE2 fragment after aldh14, GI:371960349 | 1660 |
| pCDF3-aldh15 | Sp | CloDF13cop3 | double *P*_Tac_ | Codon-optimized hypothetical protein HMPREF0402_00608 from *Fusobacterium* sp. 12_1B, from Calysta Biosystems, contains unannotated C-terminal adhE2 fragment after aldh15, GI:373496187 | 1661 |
| pCDF3-aldh46 | Sp | CloDF13cop3 | double *P*_Tac_ | Codon-optimized aldehyde dehydrogenase from *Clostridium beijerinckii* NCIMB 8052, from Calysta Biosystems, contains unannotated C-terminal adhE2 fragment after aldh46, GI:150018649 | 1662 |

*B2.* *Primary ADH screening (n-butanol)*. pCWori.trc-ter-aldh46 was constructed as a parent plasmid for the construction of the pCWori.trc-ter-aldh46.ADH series of constructs. pCWori.trc-ter-aldh46 was constructed by Gibson assembly with the aldh46.x G1 gBlock and HindIII digested pCWori-ter-trc.aldh46.adh. pCWori.trc-ter-aldh46 contains the *ter* gene driven by two *P*_tac_ promoters, the *aldh46* gene driven by the *P*_trc_ promter, the Cb resistance marker, and the ColE1 origin. pCWori.trc-ter-aldh46 was then digested with SpeI and XbaI for the insertion of corresponding ADHs. All ADHs were synthesized on gBlocks along with the apporiate overhang sequences for Gibson assembly.

| **Name** | **Marker** | **Origin** | **Promoter** | **Description** | **Number** |
| --- | --- | --- | --- | --- | --- |
| pCWori.trc-ter-aldh46.adh2 | Cb | ColE1 | double *P*_Tac_, *P*_Trc_ | Codon-optimized Ter from *Treponema denticola* under double tac promoter, codon-optimized ALDH from *Clostridium beijerinckii* NCIMB 8052 in operon with ADH from *Campylobacter fetus* subsp. *fetus* (strain 82-40) under Trc promoter | 1866 |
| pCWori.trc-ter-aldh46.adh3 | Cb | ColE1 | double *P*_Tac_, *P*_Trc_ | Codon-optimized Ter from *Treponema denticola* under double tac promoter, codon-optimized ALDH from *Clostridium beijerinckii* NCIMB 8052 in operon with codon-optimized ADH from *Olsenella* sp. oral taxon 809 str. F0356 under Trc promoter | 1911 |
| pCWori.trc-ter-aldh46.adh4 | Cb | ColE1 | double *P*_Tac_, *P*_Trc_ | Codon-optimized Ter from *Treponema denticola* under double tac promoter, codon-optimized ALDH from *Clostridium beijerinckii* NCIMB 8052 in operon with codon-optimized ADH from *Anaerofustis stercorihominis* DSM 17244 under Trc promoter | 1912 |
| pCWori.trc-ter-aldh46.adh5 | Cb | ColE1 | double *P*_Tac_, *P*_Trc_ | Codon-optimized Ter from *Treponema denticola* under double tac promoter, codon-optimized ALDH from *Clostridium beijerinckii* NCIMB 8052 in operon with codon-optimized ADH from *Bacillus subtilis* (strain 168) under Trc promoter | 1917 |
| pCWori.trc-ter-aldh46.adh6 | Cb | ColE1 | double *P*_Tac_, *P*_Trc_ | Codon-optimized Ter from *Treponema denticola* under double tac promoter, codon-optimized ALDH from *Clostridium beijerinckii* NCIMB 8052 in operon with codon-optimized ADH from *Faecalibacterium prausnitzii* M21/2 under Trc promoter | 1918 |
| pCWori.trc-ter-aldh46.adh7 | Cb | ColE1 | double *P*_Tac_, *P*_Trc_ | Codon-optimized Ter from *Treponema denticola* under double tac promoter, codon-optimized ALDH from *Clostridium beijerinckii* NCIMB 8052 in operon with codon-optimized ADH from *Erysipelotrichaceae bacterium* 3_1_53 under Trc promoter | 1919 |
| pCWori.trc-ter-aldh46.adh8 | Cb | ColE1 | double *P*_Tac_, *P*_Trc_ | Codon-optimized Ter from *Treponema denticola* under double tac promoter, codon-optimized ALDH from *Clostridium beijerinckii* NCIMB 8052 in operon with codon-optimized ADH from *Olsenella uli* (strain ATCC 49627) under Trc promoter | 1913 |
| pCWori.trc-ter-aldh46.adh9 | Cb | ColE1 | double *P*_Tac_, *P*_Trc_ | Codon-optimized Ter from *Treponema denticola* under double tac promoter, codon-optimized ALDH from *Clostridium beijerinckii* NCIMB 8052 in operon with codon-optimized ADH from *Streptococcus gallolyticus* (strain ATCC 43143 / F-1867) under Trc promoter | 1914 |
| pCWori.trc-ter-aldh46.adh10 | Cb | ColE1 | double *P*_Tac_, *P*_Trc_ | Codon-optimized Ter from *Treponema denticola* under double tac promoter, codon-optimized ALDH from *Clostridium beijerinckii* NCIMB 8052 in operon with codon-optimized ADH from *Desulfurispirillum indicum* (strain ATCC BAA-1389 / S5) under Trc promoter | 1920 |
| pCWori.trc-ter-aldh46.adh12 | Cb | ColE1 | double *P*_Tac_, *P*_Trc_ | Codon-optimized Ter from *Treponema denticola* under double tac promoter, codon-optimized ALDH from *Clostridium beijerinckii* NCIMB 8052 in operon with codon-optimized ADH from *Clostridium spiroforme* DSM 1552 under Trc promoter | 1921 |
| pCWori.trc-ter-aldh46.adh13 | Cb | ColE1 | double *P*_Tac_, *P*_Trc_ | Codon-optimized Ter from *Treponema denticola* under double tac promoter, codon-optimized ALDH from *Clostridium beijerinckii* NCIMB 8052 in operon with codon-optimized ADH from *Tetragenococcus halophilus* (strain DSM 20338) under Trc promoter | 1915 |
| pCWori.trc-ter-aldh46.adh14 | Cb | ColE1 | double *P*_Tac_, *P*_Trc_ | Codon-optimized Ter from *Treponema denticola* under double tac promoter, codon-optimized ALDH from *Clostridium beijerinckii* NCIMB 8052 in operon with codon-optimized ADH from *Succinatimonas hippei* YIT 12066 under Trc promoter | 1906 |
| pCWori.trc-ter-aldh46.dhaT2 | Cb | ColE1 | double *P*_Tac_, *P*_Trc_ | Codon-optimized Ter from *Treponema denticola* under double tac promoter, codon-optimized ALDH from *Clostridium beijerinckii* NCIMB 8052 in operon with codon-optimized dhaT from *Halanaerobium hydrogeniformans* (*Halanaerobium* sp. (strain *sapolanicus*)) under Trc promoter | 1907 |
| pCWori.trc-ter-aldh46.dhaT3 | Cb | ColE1 | double *P*_Tac_, *P*_Trc_ | Codon-optimized Ter from *Treponema denticola* under double tac promoter, codon-optimized ALDH from *Clostridium beijerinckii* NCIMB 8052 in operon with codon-optimized dhaT from *Citrobacter freundii* under Trc promoter | 1908 |
| pCWori.trc-ter-aldh46.dhaT4 | Cb | ColE1 | double *P*_Tac_, *P*_Trc_ | Codon-optimized Ter from *Treponema denticola* under double tac promoter, codon-optimized ALDH from *Clostridium beijerinckii* NCIMB 8052 in operon with codon-optimized dhaT from *Clostridium novyi* (strain NT)i under Trc promoter | 1909 |
| pCWori.trc-ter-aldh46.dhaT5 | Cb | ColE1 | double *P*_Tac_, *P*_Trc_ | Codon-optimized Ter from *Treponema denticola* under double tac promoter, codon-optimized ALDH from *Clostridium beijerinckii* NCIMB 8052 in operon with codon-optimized dhaT from *Pelobacter carbinolicus* (strain DSM 2380 / Gra Bd 1) under Trc promoter | 1910 |
| pCWori.trc-ter-aldh46.dhaT6 | Cb | ColE1 | double *P*_Tac_, *P*_Trc_ | Codon-optimized Ter from *Treponema denticola* under double tac promoter, codon-optimized ALDH from *Clostridium beijerinckii* NCIMB 8052 in operon with codon-optimized dhaT from *Pelotomaculum thermopropionicum* (strain DSM 13744 / JCM 10971 / SI) under Trc promoter | 1922 |
| pCWori.trc-ter-aldh46.dhaT7 | Cb | ColE1 | double *P*_Tac_, *P*_Trc_ | Codon-optimized Ter from *Treponema denticola* under double tac promoter, codon-optimized ALDH from *Clostridium beijerinckii* NCIMB 8052 in operon with codon-optimized dhaT from *Clostridium perfringens* D str. JGS1721 under Trc promoter | 1923 |
| pCWori.trc-ter-aldh46.dhaT8 | Cb | ColE1 | double *P*_Tac_, *P*_Trc_ | Codon-optimized Ter from *Treponema denticola* under double tac promoter, codon-optimized ALDH from *Clostridium beijerinckii* NCIMB 8052 in operon with codon-optimized dhaT from *Ilyobacter polytropus* (strain DSM 2926 / CuHBu1) under Trc promoter | 1924 |

*B3.* *ALDH.ADH pairs*. The pCWori.trc-ter-aldhX.adhY series of plasmids was constructed using Gibson Assembly or Golden gate assembly. For constructs made by Gibson assembly, the pCWori.trc-ter-aldh46.adhY series of plasmids were used as the parent plasmids. The parent plasmid was digested with EcoRI and XbaI to remove *aldh46* to insert the corresponding *aldhX* gene. The *aldhX* gene was amplified with specific primers to generate apporiate overhang for the Gibsion reaction. The pCDF3-aldh series of plasmid were used as PCR template for the ALDHs amplication. For constructs made by Golden Gate assembly, *aldhX* was amplified with the primers indicated in the primer table below using pCDF3-aldhX as a template. *adhY* was amplified with the primers indicated in the primer table below using pCWori.trc-ter-aldh46.adhY as a template for *adh2/8/12*. *adh15-22* were amplified using the gBlocks listed in *Table S1.E1* as a template. The two PCR products were then inserted into the Golden Gate entry vector, pCWori.trc-ter-RFP, digested with BsaI.

*Primer Table S1.B3*

| **Combination** | ***aldh* primers** | ***adh* primers** | **Assembly Method** |
| --- | --- | --- | --- |
| aldh3.adh2 | aldh3 GGF1  aldh3.(adh2) GGR1 | (aldh3).adh2 GGF1  adh2 GGR1 | Golden Gate |
| aldh3.adh8 | aldh3 GGF1  aldh3.(adh8) GGR1 | (aldh3).adh8 GGF1  adh8 GGR1 | Golden Gate |
| aldh3.adh12 | aldh3 GF1  aldh3.adh12 GR1 | from backbone | Gibson |
| aldh6.adh2 | aldh6 GGF1  aldh6.(adh2) GGR1 | (aldh6).adh2 GGF1  adh2 GGR1 | Golden Gate |
| aldh6.adh8 | aldh6 GF1  aldh6.adh8 GR1 | from backbone | Gibson |
| aldh6.adh12 | aldh6 GGF1  aldh6.(adh12) GGR1 | (aldh14).adh12 GGF1  adh12 GGR1 | Golden Gate |
| aldh7.adh2 | aldh7 GF1  aldh7.adh2 GR1  aldh7.adh2 GO1  aldh7.adh2 GO2 | from backbone | Gibson |
| aldh7.adh8 | aldh7 GF1  aldh7.adh8 GR1  aldh7.adh8 GO1  aldh7.adh8 GO2 | from backbone | Gibson |
| aldh7.adh12 | aldh7 GF1  aldh7.adh12 GR1  aldh7.adh12 GO1  aldh7.adh12 GO2 | from backbone | GIbson |
| aldh9.adh2 | aldh9 GGF1  aldh9.(adh2) GGR1 | (aldh9).adh2 GGF1  adh2 GGR1 | Golden Gate |
| aldh9.adh8 | aldh9 GF1  aldh9.adh8 GR1 | from backbone | GIbson |
| aldh9.adh12 | aldh9 GF1  aldh9.adh12 GR1 | from backbone | Gibson |
| aldh14.adh2 | aldh14 GGF1  aldh14.(adh2) GGR1 | (aldh14).adh2 GGF1  adh2 GGR1 | Golden Gate |
| aldh14.adh8 | aldh14 GGF1  aldh14.(adh8) GGR1 | (aldh14).adh8 GGF1  adh8 GGR1 | Golden Gate |
| aldh14.adh12 | aldh14 GGF1  aldh14.(adh12) GGR1 | (aldh14).adh12 GGF1  adh12 GGR1 | Golden Gate |
| aldh3.adh15 | aldh3 GGF1  aldh3.(adh) GGR1 | adh15 GGF1  adh15 GGR1 | Golden Gate |
| aldh3.adh17 | aldh3 GGF1  aldh3.(adh) GGR1 | adh17 GGF1  adh17 GGR1 | Golden Gate |
| aldh3.adh18 | aldh3 GGF1  aldh3.(adh) GGR1 | adh18 GGF1  adh18 GGR1 | Golden Gate |
| aldh3.adh19 | aldh3 GGF1  aldh3.(adh) GGR1 | adh19 GGF1  adh19 GGR1 | Golden Gate |
| aldh3.adh21 | aldh3 GGF1  aldh3.(adh) GGR1 | adh21 GGF1  adh21 GGR1 | Golden Gate |
| aldh3.adh22 | aldh3 GGF1  aldh3.(adh) GGR1 | adh22 GGF1  adh22 GGR1 | Golden Gate |
| aldh7.adh15 | aldh round 1 library GGF1  aldh7.(adh) GGR1 | adh15 GGF1  adh15 GGR1 | Golden Gate |
| aldh7.adh18 | aldh round 1 library GGF1  aldh7.(adh) GGR1 | adh18 GGF1  adh18 GGR1 | Golden Gate |
| aldh7.adh19 | aldh round 1 library GGF1  aldh7.(adh) GGR1 | adh19 GGF1  adh19 GGR1 | Golden Gate |
| aldh7.adh20 | aldh round 1 library GGF1  aldh7.(adh) GGR1 | adh20 GGF1  adh20 GGR1 | Golden Gate |
| aldh7.adh21 | aldh round 1 library GGF1  aldh7.(adh) GGR1 | adh21 GGF1  adh21 GGR1 | Golden Gate |
| aldh7.adh22 | aldh round 1 library GGF1  aldh7.(adh) GGR1 | adh22 GGF1  adh22 GGR1 | Golden Gate |
| aldh46.adh19 | aldh round 1 library GGF1 aldh46.(adh) GGR1 | adh19 GGF1  adh19 GGR1 | Golden Gate |
| aldh46.adh20 | aldh round 1 library GGF1 aldh46.(adh) GGR1 | adh20 GGF1  adh20 GGR1 | Golden Gate |
| aldh46.adh21 | aldh round 1 library GGF1 aldh46.(adh) GGR1 | adh21 GGF1  adh21 GGR1 | Golden Gate |

*Plasmid Table S1.B3*

| **Name** | **Marker** | **Origin** | **Promoter** | **Description** | **No.** |
| --- | --- | --- | --- | --- | --- |
| pCWori.trc-ter-aldh3.adh2 | Cb | ColE1 | double tac, trc | Codon-optimized Ter from *Treponema denticola* under double tac promoter, codon-optimized butyraldehyde dehydrogenase from *Clostridium saccharoperbutylacetonicum* N1-4(HMT) in operon with ADH *from Campylobacter fetus* subsp. fetus (strain 82-40) under Trc promoter | 2096 |
| pCWori.trc-ter-aldh6.adh2 | Cb | ColE1 | double tac, trc | Codon-optimized Ter from *Treponema denticola* under double tac promoter, codon-optimized hypothetical protein CLOBOL_07248 from *Clostridium bolteae* ATCC BAA-613 in operon with ADH from *Campylobacter fetus* subsp. *fetus* (strain 82-40) under Trc promoter | 2099 |
| pCWori.trc-ter-aldh7.adh2 | Cb | ColE1 | double tac, trc | Codon-optimized Ter from *Treponema denticola* under double tac promoter, codon-optimized ethanolamine utilization protein EutE from *Clostridium botulinum* B str. Eklund 17B in operon with ADH from *Campylobacter fetus* subsp. *fetus* (strain 82-40) under Trc promoter | 2076 |
| pCWori.trc-ter-aldh9.adh2 | Cb | ColE1 | double tac, trc | Codon-optimized Ter from *Treponema denticola* under double tac promoter, codon-optimized ethanolamine utilization protein EutE from Clostridium botulinum E1 str. 'BoNT E Beluga' in operon with ADH from *Campylobacter fetus* subsp. *fetus* (strain 82-40) under Trc promoter | 2103 |
| pCWOri.trc-ter-aldh14.adh2 | Cb | ColE1 | double tac, trc | Codon-optimized Ter from *Treponema denticola* under double tac promoter, codon-optimized hypothetical protein HMPREF9942_01197 from *Fusobacterium nucleatum* subsp. *animalis* F0419 in operon with ADH from *Campylobacter fetus subsp. fetus* (strain 82-40) under Trc promoter | 2106 |
| pCWori.trc-ter-aldh3.adh8 | Cb | ColE1 | double tac, trc | Codon-optimized Ter from *Treponema denticola* under double tac promoter, codon-optimized butyraldehyde dehydrogenase from *Clostridium saccharoperbutylacetonicum* N1-4(HMT) in operon with ADH from *Olsenella uli* (strain ATCC 49627) under Trc promoter | 2097 |
| pCWori.trc-ter-aldh6.adh8 | Cb | ColE1 | double tac, trc | Codon-optimized Ter from *Treponema denticola* under double tac promoter, codon-optimized hypothetical protein CLOBOL_07248 from *Clostridium bolteae* ATCC BAA-613 in operon with ADH from *Olsenella uli* (strain ATCC 49627 under Trc promoter | 2100 |
| pCWori.trc-ter-aldh7.adh8 | Cb | ColE1 | double tac, trc | Codon-optimized Ter from *Treponema denticola* under double tac promoter, codon-optimized ethanolamine utilization protein EutE from *Clostridium botulinum* B str. Eklund 17B in operon with ADH from *Olsenella uli* (strain ATCC 49627 under Trc promoter | 2101 |
| pCWori.trc-ter-aldh9.adh8 | Cb | ColE1 | double tac, trc | Codon-optimized Ter from *Treponema denticola* under double tac promoter, codon-optimized ethanolamine utilization protein EutE from *Clostridium botulinum* E1 str. 'BoNT E Beluga' in operon with ADH from *Olsenella uli* (strain ATCC 49627 under Trc promoter | 2104 |
| pCWori.trc-ter-aldh14.adh8 | Cb | ColE1 | double tac, trc | Codon-optimized Ter from *Treponema denticola* under double tac promoter, codon-optimized hypothetical protein HMPREF9942_01197 from *Fusobacterium nucleatum* subsp. animalis F0419 in operon with ADH from *Olsenella uli* (strain ATCC 49627 under Trc promoter | 2107 |
| pCWori.trc-ter-aldh3.adh12 | Cb | ColE1 | double tac, trc | Codon-optimized Ter from *Treponema denticola* under double tac promoter, codon-optimized butyraldehyde dehydrogenase from *Clostridium saccharoperbutylacetonicum* N1-4(HMT) in operon with ADH from *Clostridium spiroforme* DSM 1552 under Trc promoter | 2098 |
| pCWori.trc-ter-aldh7.adh12 | Cb | ColE1 | double tac, trc | Codon-optimized Ter from *Treponema denticola* under double tac promoter, codon-optimized ethanolamine utilization protein EutE from *Clostridium botulinum* B str. Eklund 17B in operon with ADH from *Clostridium spiroforme* DSM 1552 under Trc promoter | 2102 |
| pCWori.trc-ter-aldh9.adh12 | Cb | ColE1 | double tac, trc | Codon-optimized Ter from *Treponema denticola* under double tac promoter, codon-optimized ethanolamine utilization protein EutE from *Clostridium botulinum* E1 str. 'BoNT E Beluga' in operon with ADH from *Clostridium spiroforme* DSM 1552 under Trc promoter | 2105 |
| pCWori.trc-ter-aldh3.adh15 | Cb | ColE1 | double tac, trc | Codon-optimized Ter from *Treponema denticola* under double tac promoter, codon-optimized butyraldehyde dehydrogenase from *Clostridium saccharoperbutylacetonicum* N1-4(HMT) in operon with ADH from *Desulfuromonas acetoxidans* DSM 684 under Trc promoter | 2463 |
| pCWori.trc-ter-aldh3.adh17 | Cb | ColE1 | double tac, trc | Codon-optimized Ter from *Treponema denticola* under double tac promoter, codon-optimized butyraldehyde dehydrogenase from *Clostridium saccharoperbutylacetonicum* N1-4(HMT) in operon with ADH from *Dickeya dadantii* (strain Ech586) under Trc promoter | 2464 |
| pCWori.trc-ter-aldh3.adh18 | Cb | ColE1 | double tac, trc | Codon-optimized Ter from *Treponema denticola* under double tac promoter, codon-optimized butyraldehyde dehydrogenase from *Clostridium saccharoperbutylacetonicum* N1-4(HMT) in operon with ADH from *Haemophilus parainfluenzae* ATCC 33392 under Trc promoter | 2465 |
| pCWori.trc-ter-aldh3.adh19 | Cb | ColE1 | double tac, trc | Codon-optimized Ter from *Treponema denticola* under double tac promoter, codon-optimized butyraldehyde dehydrogenase from *Clostridium saccharoperbutylacetonicum* N1-4(HMT) in operon with ADH from *Enterococcus saccharolyticus* 30_1 under Trc promoter | 2466 |
| pCWori.trc-ter-aldh3.adh21 | Cb | ColE1 | double tac, trc | Codon-optimized Ter from *Treponema denticola* under double tac promoter, codon-optimized butyraldehyde dehydrogenase from *Clostridium saccharoperbutylacetonicum* N1-4(HMT) in operon with ADH from *Erysipelotrichaceae* bacterium 3_1_53 under Trc promoter | 2467 |
| pCWori.trc-ter-aldh3.adh22 | Cb | ColE1 | double tac, trc | Codon-optimized Ter from *Treponema denticola* under double tac promoter, codon-optimized butyraldehyde dehydrogenase from *Clostridium saccharoperbutylacetonicum* N1-4(HMT) in operon with ADH from *Clostridium* sp. SS2/1 under Trc promoter |  |
| pCWori.trc-ter-aldh7.adh15 | Cb | ColE1 | double tac, trc | Codon-optimized Ter from *Treponema denticola* under double tac promoter, codon-optimized ethanolamine utilization protein EutE from *Clostridium botulinum* B str. Eklund 17B in operon with ADH from *Desulfuromonas acetoxidans* DSM 684 under Trc promoter | 2479 |
| pCWori.trc-ter-aldh7.adh18 | Cb | ColE1 | double tac, trc | Codon-optimized Ter from *Treponema denticola* under double tac promoter, codon-optimized ethanolamine utilization protein EutE from *Clostridium botulinum* B str. Eklund 17B in operon with ADH from *Haemophilus parainfluenzae* ATCC 33392 under Trc promoter | 2482 |
| pCWori.trc-ter-aldh7.adh19 | Cb | ColE1 | double tac, trc | Codon-optimized Ter from *Treponema denticola* under double tac promoter, codon-optimized ethanolamine utilization protein EutE from *Clostridium botulinum* B str. Eklund 17B in operon with ADH from *Enterococcus saccharolyticus* 30_1 under Trc promoter | 2483 |
| pCWori.trc-ter-aldh7.adh20 | Cb | ColE1 | double tac, trc | Codon-optimized Ter from *Treponema denticola* under double tac promoter, codon-optimized ethanolamine utilization protein EutE from *Clostridium botulinum* B str. Eklund 17B in operon with ADH from *Clostridium botulinum* (strain Alaska E43 / Type E3) under Trc promoter | 2484 |
| pCWori.trc-ter-aldh7.adh21 | Cb | ColE1 | double tac, trc | Codon-optimized Ter from *Treponema denticola* under double tac promoter, codon-optimized ethanolamine utilization protein EutE from *Clostridium botulinum* B str. Eklund 17B in operon with ADH from *Erysipelotrichaceae* bacterium 3_1_53 under Trc promoter | 2485 |
| pCWori.trc-ter-aldh7.adh22 | Cb | ColE1 | double tac, trc | Codon-optimized Ter from *Treponema denticola* under double tac promoter, codon-optimized ethanolamine utilization protein EutE from *Clostridium botulinum* B str. Eklund 17B in operon with ADH from ADH from *Clostridium* sp. SS2/1 under Trc promoter | 2486 |
| pCWori.trc-ter-aldh46.adh19 | Cb | ColE1 | double tac, trc | Codon-optimized Ter from *Treponema denticola* under double tac promoter, codon-optimized codon-optimized ALDH from *Clostridium beijerinckii* NCIMB 8052 in operon with ADH from *Enterococcus saccharolyticus* 30_1 under Trc promoter | 2491 |
| pCWori.trc-ter-aldh46.adh20 | Cb | ColE1 | double tac, trc | Codon-optimized Ter from *Treponema denticola* under double tac promoter, codon-optimized codon-optimized ALDH from *Clostridium beijerinckii* NCIMB 8052 in operon with ADH from *Clostridium botulinum* (strain Alaska E43 / Type E3) under Trc promoter | 2492 |
| pCWori.trc-ter-aldh46.adh21 | Cb | ColE1 | double tac, trc | Codon-optimized Ter from *Treponema denticola* under double tac promoter, codon-optimized codon-optimized ALDH from *Clostridium beijerinckii* NCIMB 8052 in operon with ADH from *Erysipelotrichaceae* bacterium 3_1_53 under Trc promoter | 2493 |

*B4. Secondary ADHs (BDO).* The pCWori.trc-sADH-aldh7.adh2 series of plasmids (SADH1-16) were constructed by replaceing the *ter* gene in the NdeI and XhoI site of the pCWori.trc-ter-aldh7.adh2 parent plasmid with the corresponding gBlock sequences (*Table S1.E2*).

| **Name** | **Marker** | **Origin** | **Promoter** | **Description** | **No.** |
| --- | --- | --- | --- | --- | --- |
| pCWori.trc-sadh1-aldh7.adh2 | Cb | ColE1 | *P*_Trc_,  double *P*_Tac_ | SADH1 is from *Pichia kudriavzevii*. Accession # KGK36767.1 | 2430 |
| pCWori.trc-sadh2-aldh7.adh2 | Cb | ColE1 | *P*_Trc_,  double *P*_Tac_ | SADH2 is from *Pyrococcus furiosus* DSM 3638. Accession # WP_011011186.1 | 2431 |
| pCWori.trc-sadh3-aldh7.adh2 | Cb | ColE1 | *P*_Trc_,  double *P*_Tac_ | SADH3 is from *Cupriavidus necator*. Accession # WP_011614641.1 | 2432 |
| pCWori.trc-sadh4-aldh7.adh2 | Cb | ColE1 | *P*_Trc_,  double *P*_Tac_ | SADH4 is from *Thermoanaerobacter brockii*. Accession # P14941.1 | 2433 |
| pCWori.trc-sadh5-aldh7.adh2 | Cb | ColE1 | *P*_Trc_,  double *P*_Tac_ | SADH5 is from *Clostridium beijerinckii*. Accession # AAA23199.2 | 2434 |
| pCWori.trc-sadh6-aldh7.adh2 | Cb | ColE1 | *P*_Trc_,  double *P*_Tac_ | SADH6 is from *Kluyveromyces lactis* NRRL Y-1140. Accession # XP_455102.1 | 2435 |
| pCWori.trc-sadh7-aldh7.adh2 | Cb | ColE1 | *P*_Trc_,  double *P*_Tac_ | SADH7 is from *Phytomonas* sp. ADU-2003 . Accession # AAP39869.1 | 2436 |
| pCWori.trc-sadh8-aldh7.adh2 | Cb | ColE1 | *P*_Trc_,  double *P*_Tac_ | SADH8 is from *Ralstonia eutropha* H16 . Accession # Q0KDL6.1 | 2437 |
| pCWori.trc-sadh9-aldh7.adh2 | Cb | ColE1 | *P*_Trc_,  double *P*_Tac_ | SADH9 is from *Trichomonas vaginalis* G3. Accession # XP_001580601.1 | 2438 |
| pCWori.trc-sadh10-aldh7.adh2 | Cb | ColE1 | *P*_Trc_,  double *P*_Tac_ | SADH10 is from *Pseudomonas fluorescens*. Accession # AJP52792.1 | 2439 |
| pCWori.trc-sadh11-aldh7.adh2 | Cb | ColE1 | *P*_Trc_,  double *P*_Tac_ | SADH11 is *from Lactococcus lactis*. Accession # WP_011835462.1 | 2440 |
| pCWori.trc-sadh12-aldh7.adh2 | Cb | ColE1 | *P*_Trc_,  double *P*_Tac_ | SADH12 is from *Saccharomyces cerevisiae*. Accession # AAC04974.1 | 2441 |
| pCWori.trc-sadh13-aldh7.adh2 | Cb | ColE1 | *P*_Trc_,  double *P*_Tac_ | SADH13 is *from Escherichia coli*. Accession # WP_000374004.1 | 2442 |
| pCWori.trc-sadh14-aldh7.adh2 | Cb | ColE1 | *P*_Trc_,  double *P*_Tac_ | SADH14 is from *Zygoascus ofunaensis*. Accession # BAD32689.1 | 2443 |
| pCWori.trc-sadh15-2-aldh7.adh2 | Cb | ColE1 | *P*_Trc_,  double *P*_Tac_ | SADH15 is from *Candida parapsilosis*. Accession # BAA24528.1 | 2444 |
| pCWori.trc-sadh16-aldh7.adh2 | Cb | ColE1 | *P*_Trc_,  double *P*_Tac_ | SADH16 is from *Cyberlindnera jadinii*. Accession # BAN45671.1 | 2445 |

**C. Oligonucleotide sequences.** For primers used to introduce guide sequences for genome mutation, the guide sequence is in red.

| **Name** | **Sequence** |
| --- | --- |
| adhE2 F6 | gatatacatatggcaagctggagccacccgcagttcgaaaagggtgcaggtatgaaagtcacgaaccagaaggaactgaagcagaaactgaacgaactgc |
| adhE2 R1 | atcaagcttggtaccttaaaaagatttgatataaatgtctttcagctcagagatcagcgggtaacgcggg |
| adhE2 R9 | cgcgatacccgccggcacgtggtgcatagcacccag |
| adhE2 F9 | cacgtgccggcgggtatcgcgtgtgctgtcctgatcgaagaagtaattaagtacaacgc |
| adhE2 R11 | gtgagccatagacgcgcacacgcccaggaacgcgttgg |
| adhE2 F11 | cctgggcgtgtgcgcgtctatggctcacaaactgggtgctatgcaccacgtgcc |
| adhE2 AADH R1 | atcaagcttggtaccttagttttcacggcgttctgcaacagatttgatgttcagcagatgc |
| trc.crt delete GO1 | caagcttgcatgcctgcaggtcgactctagattagcccatgtgcaggccaccgttcaggg |
| trc.crt delete GO2 | gaacggtggcctgcacatgggctaatctagagtcgacctgcaggcatgcaagcttggctg |
| phaA GF1 | taacaatttcacacagagctcaaggagatatacatatgactgatgttgtgattgtaagcg |
| phaA GR1 | aaaacagccaagcttgcatgcctgcaggtcgactctagattatttgcgttcgacagccag |
| Clo DF13-cop3 QCF1 | gatttggttgctgtgctctgcgaaaaccagttaccacggttaagcagttcc |
| Clo DF13-cop3 QCR1 | ggaactgcttaaccgtggtaactggttttcgcagagcacagcaaccaaatc |
| aadA CloDF F1 | gatacaaagcttttatttgccgactaccttggtgatctcg |
| aadA CloDF R1 | gatacacctgagggatcaaaggatcttcttgagatcctttttttctgcg |
| ter delete GO5 | agcggataacaatttcacacaggaaacaggatccgaattcaaaaaaggaggtaaaaaatg |
| ter delete GO6 | cattttttacctccttttttgaattcggatcctgtttcctgtgtgaaattgttatccgct |
| aldh46 GF3 | tgagggtattaactacgaggc |
| aldh46.ADH GR1 | actttgaaccacagcattaggacctcctctggtaagctctagattagccggccagaacac |
| aldh46.ADH GF1 | gttctggccggctaatctagagcttaccagaggaggtcctaatgctgtggttcaaagtcc |
| ADH GR1 | catgtttgacagcttatcatcgataagcttttaaaaagatttgatataaatgtctttcag |
| aldh GF1 | ttgtgagcggataacaatttcacacaggaaacaggaattcaaaaaaggaggtaaaaaatg |
| aldh3 GF2 | cggataacaatttcacacaggaaacaggaattcaaaaaaggaggtaaaaaatgattaagg |
| aldh3.adh12 GR1 | cgggttggagtaggtaaagtcgcccagcatttctttcccccttaggaactagggttctagattaacccgccagaacacaac |
| aldh6 GF2 | gcggataacaatttcacacaggaaacaggaattcaaaaaaggaggtaaaaaatgaaagag |
| aldh6.adh8 GR1 | ggtcggtacgtggaacatgaagtcgtacatttggtccccctttgcgctaggattctctagattaacgaatgctaaaggcgtcc |
| aldh7 GF2 | gagcggataacaatttcacacaggaaacaggaattcaaaaaaggaggtaaaaaatggaac |
| aldh7.adh2 GR1 | ttcgatacgggttggattgcagtaggaaaagttgaccatcttatacctctttgtgtgtgaaagctctagattaaccggccagaacgcaac |
| aldh7.adh2 GO1 | ttttcctactgcaatccaacccgtatcgaattcggcaaaggtaaagaaaactccatcggt |
| aldh7.adh2 GO2 | accgatggagttttctttacctttgccgaattcgatacgggttggattgcagtaggaaaa |
| aldh7.adh8 GR1 | ggtcggtacgtggaacatgaagtcgtacattaaaagtcctcccccttgtgggttttctagattaaccggccagaacgcaac |
| aldh7.adh12 GR1 | cgggttggagtaggtaaagtcgcccagcataagtatatctccttaagttccttggtctagattaaccggccagaacgcaac |
| aldh9 GF2 | gcggataacaatttcacacaggaaacaggaattcaaaaaaggaggtaaaaaatgaatgac |
| aldh9.adh8 GR1 | tcggtacgtggaacatgaagtcgtacatttagtgcccctccttaagattagcgtctagattacttaatagaaaagttgtcaaccataacg |
| aldh9.adh12 GR1 | ggttggagtaggtaaagtcgcccagcatagtttagtacctccttcgttaggtatctagattacttaatagaaaagttgtcaaccataacg |
| aldh round 1 library GGF1 | gctcagcggtctcgaattcaaaaaaggaggtaaaaaatg |
| aldh3 GGF1 | gatacaggtctcgattcaaaaaaggaggtaaaaaatgattaaggac |
| aldh6 GGF1 | gatacaggtctctattcaaaaaaggaggtaaaaaatgaaagagg |
| aldh9 GGF1 | gatacaggtctcaattcaaaaaaggaggtaaaaaatgaatgactttaac |
| aldh14 GGF1 | gatacaggtctccattcaaaaaaggaggtaaaaaatggaatttg |
| adh2 GGR1 | gatacaggtctccactagtttagtaagccaggttcagaatttcttc |
| adh8 GGR1 | gatacaggtctcaactagtttacagagcagcacggtagatttc |
| adh12 GGR1 | gatacaggtctcgactagtttatttcatggattgtttcaggatg |
| (aldh14).adh12 GGF1 | gatacaggtctcaagacaccaggaggcaagcatgctgggcgactttacct |
| aldh14.(adh12) GGR1 | gatacaggtctccgtcttagagtatctagattatttaatgctaaaaccgccca |
| (aldh14).adh8 GGF1 | gatacaggtctcacgagaaggaggataatcctatgtacgacttcatgttccacg |
| aldh14.(adh8) GGR1 | gatacaggtctctctcgcgctagtctagattatttaatgctaaaaccgccca |
| (aldh14).adh2 GGF1 | gatacaggtctcaccgaatctaggaggaatcacatggtcaacttttcctactgcaa |
| aldh14.(adh2) GGR1 | gatacaggtctcctcgggttcttctagattatttaatgctaaaaccgccca |
| (aldh9).adh2 GGF1 | gatacaggtctccaagggaggcttccgccaatggtcaacttttcctactgcaa |
| aldh9.(adh2) GGR1 | gatacaggtctcaccttttgtatgatctagattacttaatagaaaagttgtcaaccataacg |
| (aldh6).adh12 GGF1 | gatacaggtctctaaggaggacacgacaccatgctgggcgactttacct |
| aldh6.(adh12) GGR1 | gatacaggtctcccctttatgtcggtctagattaacgaatgctaaaggcgtcc |
| (aldh6).adh2 GGF1 | gatacaggtctcaaaagggggtaaccaatatggtcaacttttcctactgcaa |
| aldh6.(adh2) GGR1 | gatacaggtctctctttagattaggatctagattaacgaatgctaaaggcgtcc |
| (aldh3).adh8 GGF1 | gatacaggtctcacaacaaggggggcaataaatgtacgacttcatgttccacg |
| aldh3.(adh8) GGR1 | gatacaggtctcagttgctatgagtctagattaacccgccagaacacaa |
| (aldh3).adh2 GGF1 | gatacaggtctcctccagaggaggaacagatggtcaacttttcctactgcaa |
| aldh3.(adh2) GGR1 | gatacaggtctcgtggatttgttttgtctagattaacccgccagaacacaa |
| adh15 GGF1 | gatacaggtctcgtctagaagagatc |
| adh15 GGR1 | tgtatcggtctctactagtttagctacg |
| adh16 GGF1 | gatacaggtctcgtctagaaagagtaataag |
| adh16 GGR1 | tgtatcggtctctactagtttaatcgttag |
| adh17 GGF1 | gatacaggtctcgtctagatagccag |
| adh17 GGR1 | tgtatcggtctctactagtttacagtgc |
| adh18 GGF1 | gatacaggtctcgtctagacacgac |
| adh18 GGR1 | tgtatcggtctctactagtttattttgc |
| adh20 GGF1 | gatacaggtctcgtctagacataaagc |
| adh20 GGR1 | tgtatcggtctctactagtttacagagc |
| adh22 GGF1 | gatacaggtctcgtctagattaaaatcc |
| adh22 GGR1 | tgtatcggtctctactagtttacaggg |
| aldh3.(adh) GGR1 | gatacaggtctcttagattaacccgccagaacacaa |
| aldh6.(adh) GGR1 | gatacaggtctcttagattaacgaatgctaaaggcgtcc |
| aldh7.(adh) GGR1 | gatacaggtctcttagattaaccggccagaacgc |
| aldh46.(adh) GGR1 | gatacaggtctcttagattagccggccagaacacaac |
| rrnB-1 GF1 | ggtattaactacgaggcagaagttg |
| rrnB-1 GR1 | gttccctactctcgcatgggctgaccccacactaccatcg |
| rrnB-2 GF1 | cgatggtagtgtggggtcagcccatgcgagagtagggaac |
| rrnB-2 GR1 | cagcttccgatggctgcc |
| BsaI delete GO1 | gctgataaatctggagccggtgagcgtggctcccgcggtatcattgcagcactggggcca |
| BsaI delte GO2 | tggccccagtgctgcaatgataccgcgggagccacgctcaccggctccagatttatcagc |
| RFP BsaI GR2 | gatctgctcatgtttgacagcttatcatcgatactaggagacctataaacgcagaaaggc |
| RFP BsaI GF2 | gataacaatttcacacaggaaacaggaattcagagacctccctatcagtgatagagattg |
| sadh1 GF1 | caggaaacaggatccatcgatgcttaggaggtcatatgttcatgaaaggtctgacgtatc |
| sadh1 GR1 | aaaatcttctctcatccgccaaaacagccctcgagttacggcgtcagaataattttcatc |
| sadh2 GF1 | ggaaacaggatccatcgatgcttaggaggtcatatgtctaagaactacaaaaagcaccag |
| sadh2 GR1 | atcttctctcatccgccaaaacagccctcgagttagtattccggtttctggtaaatagag |
| sadh3 GF1 | cacaggaaacaggatccatcgatgcttaggaggtcatatgaaaatctctctgacttccgc |
| sadh3 GR1 | gctgaaaatcttctctcatccgccaaaacagccctcgagttaagcggtcggtactgcgac |
| sadh4 GF1 | cacacaggaaacaggatccatcgatgcttaggaggtcatatgaagggtttcgctatgctg |
| sadh4 GR1 | gctgaaaatcttctctcatccgccaaaacagccctcgagttacgccaggatgactaccgg |
| sadh5 GF1 | cacacaggaaacaggatccatcgatgcttaggaggtcatatgaaaggcttcgctatgctg |
| sadh5 GR1 | cttctctcatccgccaaaacagccctcgagttacagaattacaacagctttgatcagatc |
| sadh6 GF1 | acaatttcacacaggaaacaggatccatcgatgcttaggaggtcatatgcgcgcactggc |
| sadh6 GR1 | gctgaaaatcttctctcatccgccaaaacagccctcgagttagttggtcgcgtccagttc |
| sadh7 GF1 | acacaggaaacaggatccatcgatgcttaggaggtcatatgaaagcagccgtgttcaaag |
| sadh7 GR1 | gaaaatcttctctcatccgccaaaacagccctcgagttacggcttgatggctactttcag |
| sadh8 GF1 | ttcacacaggaaacaggatccatcgatgcttaggaggtcatatgaccgcgatgatgaagg |
| sadh8 GR1 | aggctgaaaatcttctctcatccgccaaaacagccctcgagttagtgcggtttgatggcg |
| sadh9 GF1 | tttcacacaggaaacaggatccatcgatgcttaggaggtcatatgaccttcgaactgccg |
| sadh9 GR1 | aatcttctctcatccgccaaaacagccctcgagttacagtttgtcgttgtattcaatcag |
| sadh10 GF1 | ttcacacaggaaacaggatccatcgatgcttaggaggtcatatgatggcgaccatgaaag |
| sadh10 GR1 | ctgaaaatcttctctcatccgccaaaacagccctcgagttacggggtgattgcaactttc |
| sadh11 GF1 | ttcacacaggaaacaggatccatcgatgcttaggaggtcatatgtctaaagtggctgccg |
| sadh11 GR1 | gctgaaaatcttctctcatccgccaaaacagccctcgagttaatgaaattgcatgccacc |
| sadh12 GF1 | cacacaggaaacaggatccatcgatgcttaggaggtcatatgcgtgcgctggcctatttc |
| sadh12 GR1 | ctgaaaatcttctctcatccgccaaaacagccctcgagttatttcatttcaccgtggttg |
| sadh13 GF1 | tttcacacaggaaacaggatccatcgatgcttaggaggtcatatggaccgcatcatccag |
| sadh13 GR1 | aggctgaaaatcttctctcatccgccaaaacagccctcgagttattcccattcctgcagg |
| sadh14 GF1 | acacaggaaacaggatccatcgatgcttaggaggtcatatgatgaaagcgctgtgttacc |
| sadh14 GR1 | ctgaaaatcttctctcatccgccaaaacagccctcgagttattcgtcacaggtgatcacc |
| sadh15 GF1 | tttcacacaggaaacaggatccatcgatgcttaggaggtcatatgagcattccgtccagc |
| sadh15 GR1 | tctctcatccgccaaaacagccctcgagttacgggttaaagacaacacgaccctcatacg |
| sadh16 GF1 | ttcacacaggaaacaggatccatcgatgcttaggaggtcatatgctgcgcactaatctgc |
| sadh16 GR1 | aatcttctctcatccgccaaaacagccctcgagttatttggtagtatccacgacgtaacg |
| .03 HBD QF1 | cgactacgggtaattcggaac |
| .03 HBD QR1-1 | atcaacaaattcatccttaatgtcg |
| .03 HBD QR1-2 | gtttttgttaatgaagtccaggc |
| .03 HBD QR1 | cgactgtcgttccgaattacc |
| .03 HBD QF1-1 | ggttgccgtatcctggtgacg |
| .03 HBD QF1-2 | gccacccaatcggtgcttctg |
| pTT QF1 | aatgagctgttgacaattaatcatc |
| pTT QR1 | ccaccgaacttaccaacagc |
| pTT QF2 | cgtataatgtgtggaattgtgagc |
| pTT QR2 | actactgcgcccagttccgg |
| pTT QF3 | agctggcacgacaggtttcc |
| pTT QR3 | cacagcttccgatggctgcc |
| .3 crt QF1 | ggctgcacacaagacagacc |
| .3 crt QR1-1 | gatttcgccaataacgtagtcc |
| .3 crt QR1-2 | gcacttcgctgtcgttctcg |
| .3 crt QR1 | gtcttgtgtgcagcctctcg |
| .3 crt QF1-1 | tcgctcaaggcgcactcccg |
| .3 crt QF1-2 | gtaaatcactgcataattcgtgtcg |
| CmR QF1 | ggggcgaagaagttgtcc |
| CmR QR1 | atattcgcaagatgtggcg |
| CmR QF2 | ttgccatacggaattccg |
| CmR QR2 | gaggcatttcagtcagttgc |

**D. Gene sequences of dehydrogenases**

*D1. AdhE2 variants in the pCDF3-TdTer.AdhE2.1-96 plasmids*

**AdhE2.1 (76930)**

```
       1  ATGAAAGTGA CGAACCAAAA AGAACTGAAA CAAAAACTGA ATGAACTGCG TGAGGCCCAG
      61  AAAAAGTTCG CTACGTATAC CCAGGAGCAA GTTGACAAGA TCTTCAAACA GTGTGCCATT
```

121 GCGGCTGCCA AAGAGCGTAT TAACTTGGCC AAACTGGCGG TGGAAGAAAC CGGCATCGGT

181 CTGGTCGAGG ACAAGATTAT CAAGAATCAC TTTGCCGCGG AGTATATCTA CAACAAATAC

```
     241  AAAAACGAAA AGACTTGCGG CATCATTGAT CATGATGACA GCTTGGGTAT CACGAAAGTC
     301  GCAGAACCAA TCGGTATTGT CGCGGCAATT GTCCCGACCA CTAATCCGAC CAGCACCGCG
     361  ATCTTCAAGA GCTTGATCAG CCTGAAAACC CGCAACGCGA TTTTCTTCAG CCCGCACCCG
     421  CGTGCGAAGA AGAGCACCAT CGCAGCGGCG AAGCTGATTT TGGATGCCGC GGTTAAGGCC
     481  GGTGCACCGA AAAACATTAT TGGCTGGATT GACGAGCCGA GCATTGAGCT GTCTCAGGAC
     541  CTGATGAGCG AAGCGGACAT TATTCTGGCC ACCGGTGGTC CGTCTATGGT TAAGGCAGCG
     601  TACTCCAGCG GTAAGCCTGC TATCGGCGTT GGTGCGGGCA ATACGCCGGC TATCATCGAT
     661  GAAAGCGCGG ATATTGATAT GGCAGTTAGC TCCATCATTC TGTCCAAAAC CTACGACAAT
     721  GGTGTTATCT GTGCAAGCGA GCAAAGCATC CTGGTGATGA ATAGCATTTA CGAGAAGGTC
     781  AAAGAAGAGT TTGTTAAGCG TGGTTCCTAT ATTCTGAACC AGAACGAGAT TGCAAAAATC
     841  AAAGAAACGA TGTTTAAGAA TGGCGCGATT AATGCAGATA TTGTTGGTAA GTCCGCATAC
     901  ATTATCGCGA AGATGGCGGG TATCGAAGTG CCGCAAACCA CCAAAATCTT GATCGGCGAA
     961  GTGCAGAGCG TGGAGAAGAG CGAACTGTTC AGCCATGAGA AACTGAGCCC GGTTCTGGCT
    1021  ATGTACAAAG TGAAGGACTT TGACGAAGCC CTGAAGAAAG CTCAGCGTCT GATTGAGCTG
    1081  GGCGGCAGCG GTCACACGAG CAGCCTGTAT ATCGATAGCC AAAACAATAA GGACAAGGTG
    1141  AAAGAGTTCG GTCTGGCGAT GAAAACCTCT CGCACGTTTA TCAACATGCC GAGCAGCCAA
    1201  GGTGCGAGCG GCGATCTGTA TAACTTCGCA ATCGCGCCTT CTTTTACGCT GGGTTGCGGT
    1261  ACCTGGGGCG GTAATTCGGT GAGCCAGAAT GTCGAGCCGA AGCACCTGCT GAACATCAAA
    1321  TCGGTCGCGG AGCGCCGTGA GAAGAGACCG TGGTTCAAAG TCCCACAGAA AATTTACTTC
    1381  AAATACGGCT GCCTGCGTTT CGCGCTGAAA GAACTGAAAG ACATGAACAA AAAGCGTGCG
    1441  TTCATTGTTA CCGACAAAGA CCTGTTCAAA CTGGGTTACG TGAACAAAAT CACCAAAGTT
    1501  CTGGATGAAA TTGACATCAA GTACTCCATC TTCACTGATA TCAAATCCGA CCCAACGATT
    1561  GATAGCGTGA AAAAGGGCGC TAAAGAAATG CTGAACTTTG AACCGGACAC CATCATCAGC
    1621  ATCGGTGGTG GCTCTCCTAT GGATGCTGCG AAGGTCATGC ACCTGCTGTA CGAATACCCG
    1681  GAAGCGGAAA TCGAAAACCT GGCTATCAAC TTCATGGACA TCCGCAAACG TATCTGCAAC
    1741  TTCCCGAAGC TGGGCACTAA AGCTATTTCC GTTGCCATCC CGACTACCGC GGGCACTGGT
    1801  TCCGAAGCCA CGCCGTTCGC CGTGATCACC AACGATGAAA CCGGTATGAA ATACCCGCTG
    1861  ACCTCTTACG AACTGACCCC GAACATGGCA ATTATCGACA CCGAGCTGAT GCTGAACATG
    1921  CCGCGCAAGC TGACCGCTGC TACCGGCATC GACGCTCTGG TACATGCTAT TGAGGCGTAC
    1981  GTTTCCGTGA TGGCTACCGA TTACACCGAC GAACTGGCCC TGCGTGCGAT CAAAATGATT
    2041  TTCAAGTACC TGCCTCGCGC TTACAAAAAC GGCACGAATG ACATCGAGGC GCGTGAGAAA
    2101  ATGGCCCATG CAAGCAACAT CGCGGGCATG GCCTTCGCCA ACGCGTTCCT GGGCGTGTGC
    2161  CACTCTATGG CTCACAAACT GGGTGCTATG CACCACGTGC CGCACGGTAT CGCGTGTGCT
    2221  GTCCTGATCG AAGAAGTAAT TAAGTACAAC GCTACTGATT GCCCGACTAA ACAGACCGCC
    2281  TTCCCACAGT ACAAATCTCC TAACGCTAAA CGTAAGTACG CTGAGATCGC CGAATACCTG
    2341  AACCTGAAGG GTACGAGCGA CACTGAGAAA GTTACTGCGC TGATCGAAGC TATCTCTAAA
    2401  CTGAAAATTG ACCTGTCCAT CCCGCAGAAC ATCAGCGCCG CAGGCATCAA CAAAAAGGAC
    2461  TTTTACAACA CGCTGGACAA AATGAGCGAA CTGGCTTTTG ACGACCAGTG CACCACTGCA
    2521  AACCCGCGTT ACCCGCTGAT CTCTGAGCTG AAAGACATTT ATATCAAATC TTTTTAA
```

**AdhE2.2 (76931)**

```
       1  ATGAAAGTGA CGAACCAAAA AGAACTGAAA CAAAAAGACA ATGAACTGCG TGAGGCCCAG
      61  AAAAAGTTCG CTAGCTATAC CCAGGAGCAA GTTGACAAGA TCTTCAAACA GTGTGCCATT
```

121 GCGGCTGCCA AAGAGCGTAT TAACTTGGCC AAACTGGCGG TGGAAGAAAC CGGCATCGGT

181 CTGGTCGAGG ACAAGATTAT CAAGAATCAC TTTGCCGCGG AGTATATCTA CAACAAATAC

```
     241  AAAAACGGTA AGACTTGCGG CATCATTGAT CATGATGACA GCTTGGGTAT CACGAAAGTC
     301  GCAGAACCAA TCGGTATTGT CGCGGCAATT GTCCCGACCA CTAATCCGAC CAGCACCGCG
     361  ATCTTCAAGA GCTTGATCAG CCTGAAAACC CGCAACGCGA TTTTCTTCAG CCCGCACCCG
     421  CGTGCGAAGA AGAGCACCAT CGCAGCGGCG AAGCTGATTT TGGATGCCGC GGTTAAGGCC
     481  GGTGCACCGA AAAACATTAT TGGCTGGATT GACGAGCCGA GCATTGAGCT GTCTCAGGAC
     541  CTGATGAGCG AAGCGGACAT TATTCTGGCC ACCGGTGGTC CGTCTATGGT TAAGGCAGCG
     601  TACTCCAGCG GTAAGCCTGC TATCGGCGTT GGTGCGGGCA ATACGCCGGC TATCATCGAT
     661  GAAAGCGCGG ATATTGATAT GGCAGTTAGC TCCATCATTC TGTCCAAAAC CTACGACAAT
     721  GGTGTTATCT GTGCAAGCGA GCAAAGCATC CTGGTGATGA ATAGCATTTA CGAGAAGGTC
     781  AAAGAAGAGT TTGTTAAGCG TGGTTCCTAT ATTCTGAACC AGAACGAGAT TGCAAAAATC
     841  AAAGAAACGA TGTTTAAGAA TGGCGCGATT AATGCAGATA TTGTTGGTAA GTCCGCATAC
     901  ATTATCGCGA AGATGGCGGG TATCGAAGTG CCGCAAACCA CCAAAATCTT GATCGGCGAA
     961  GTGCAGAGCG TGGAGAAGAG CGAACTGTTC AGCCATGAGA AACTGAGCCC GGTTCTGGCT
    1021  ATGTACAAAG TGAAGGACTT TGACGAAGCC CTGAAGAAAG CTCAGCGTCT GATTGAGCTG
    1081  GGCGGCAGCG GTCACACGAG CAGCCTGTAT ATCGATAGCC AAAACAATAA GGACAAGGTG
    1141  AAAGAGTTCG GTCTGGCGAT GAAAACCTCT CGCACGTTTA TCAACATGCC GAGCAGCCAA
    1201  GGTGCGAGCG GCGATCTGTA TAACTTCGCA ATCGCGCCTT CTTTTACGCT GGGTTGCGGT
    1261  ACCTGGGGCG GTAATTCGGT GAGCCAGAAT GTCGAGCCGA AGCACCTGCT GAACATCAAA
    1321  TCGGTCGCGG AGCGCCGTGA GAAGAGACCG TGGTTCAAAG TCCCACAGAA AATTTACTTC
    1381  AAATACGGCT GCCTGCGTTT CGCGCTGAAA GAACTGAAAG ACATGAACAA AAAGCGTGCG
    1441  TTCATTGTTA CCGACAAAGA CCTGTTCAAA CTGGGTTACG TGAACAAAAT CACCAAAGTT
    1501  CTGGATGAAA TTGACATCAA GTACTCCATC TTCACTGATA TCAAATCCGA CCCAACGATT
    1561  GATAGCGTGA AAAAGGGCGC TAAAGAAATG CTGAACTTTG AACCGGACAC CATCATCAGC
    1621  ATCGGTGGTG GCTCTCCTAT GGATGCTGCG AAGGTCATGC ACCTGCTGTA CGAATACCCG
    1681  GAAGCGGAAA TCGAAAACCT GGCTATCAAC TTCATGGACA TCCGCAAACG TATCTGCAAC
    1741  TTCCCGAAGC TGGGCACTAA AGCTATTTCC GTTGCCATCC CGACTACCGC GGGCACTGGT
    1801  TCCGAAGCCA CGCCGTTCGC CGTGATCACC AACGATGAAA CCGGTATGAA ATACCCGCTG
    1861  ACCTCTTACG AACTGACCCC GAACATGGCA ATTATCGACA CCGAGCTGAT GCTGAACATG
    1921  CCGCGCAAGC TGACCGCTGC TACCGGCATC GACGCTCTGG TACATGCTAT TGAGGCGTAC
    1981  GTTTCCGTGA TGGCTACCGA TTACACCGAC GAACTGGCCC TGCGTGCGAT CAAAATGATT
    2041  TTCAAGTACC TGCCTCGCGC TTACAAAAAC GGCACGAATG ACATCGAGGC GCGTGAGAAA
    2101  ATGGCCCATG CAAGCAACAT CGCGGGCATG GCCTTCGCCA ACGCGTTCCT GGGCGTGTGC
    2161  CACTCTATGG CTCACAAACT GGGTGCTATG CACCACGTGC CGCACGGTAT CGCGTGTGCT
    2221  GTCCTGATCG AAGAAGTAAT TAAGTACAAC GCTACTGATT GCCCGACTAA ACAGACCGCC
    2281  TTCCCACAGT ACAAATCTCC TAACGCTAAA CGTAAGTACG CTGAGATCGC CGAATACCTG
    2341  AACCTGAAGG GTACGAGCGA CACTGAGAAA GTTACTGCGC TGATCGAAGC TATCTCTAAA
    2401  CTGAAAATTG ACCTGTCCAT CCCGCAGAAC ATCAGCGCCG CAGGCATCAA CAAAAAGGAC
    2461  TTTTACAACA CGCTGGACAA AATGAGCGAA CTGGCTTTTG ACGACCAGTG CACCACTGCA
    2521  AACCCGCGTT ACCCGCTGAT CTCTGAGCTG AAAGACATTT ATATCAAATC TTTTTAA
```

**AdhE2.3 (76932)**

```
       1  ATGAAAGTGA CGAACCAAAA AGAACTGAAA CAAAAACTGA ATGAACTGCG TGAGGCCCAG
      61  AAAAAGTTCG CTACGTATAC CCAGGAGCAA GTTGACAAGA TCTTCAAACA GTGTGCCATT
```

121 GCGGCTGCCA AAGAGCGTAT TAACTTGGCC AAACTGGCGG TGGAAGAAAC CGGCATCGGT

181 CTGGTCGAGG ACAAGATTAT CAAGAATCAC TTTGCCGCGG AGTATATCTA CAACAAATAC

```
     241  AAAAACGAAA AGACTTGCGG CATCATTGAT CATGATGACA GCTTGGGTAT CACGAAAGTC
     301  GCAGAACCAA TCGGTATTGT CGCGGCAATT GTCCCGACCA CTAATCCGAC CAGCACCGCG
     361  ATCTTCAAGA GCTTGATCAG CCTGAAAACC CGCAACGCGA TTTTCTTCAG CCCGCACCCG
     421  CGTGCGAAGA AGAGCACCAT CGCAGCGGCG AAGCTGATTT TGGATGCCGC GGTTAAGGCC
     481  GGTGCACCGA AAAACATTAT TGGCTGGATT GACGAGCCGA GCATTGAGCT GTCTCAGGAC
     541  CTGATGAGCG AAGCGGACAT TATTCTGGCC ACCGGTGGTC CGTCTATGGT TAAGGCAGCG
     601  TACTCCAGCG GTAAGCCTGC TATCGGCGTT GGTGCGGGCA ATACGCCGGC TATCATCGAT
     661  GAAAGCGCGG ATATTGATAT GGCAGTTAGC TCCATCATTC TGGGTAAAAC CTACGACAAT
     721  GGTGTTATCT GTGCAAGCGA GCAAAGCATC CTGGTGATGA ATAGCATTTA CGAGAAGGTC
     781  AAAGAAGAGT TTGTTAAGCG TGGTTCCTAT ATTCTGAACC AGAACGAGAT TGCAAAAATC
     841  AAAGAAACGA TGTTTAAGAA TGGCGCGATT AATGCAGATA TTGTTGGTAA GTCCGCATAC
     901  ATTATCGCGA AGATGGCGGG TATCGAAGTG CCGCAATACA CCAAAATCTT GATCGGCGAA
     961  GTGCAGAGCG TGGAGAAGAG CGAACTGTTC AGCCATGAGA AACTGAGCCC GGTTCTGGCT
    1021  ATGTACAAAG TGAAGGACTT TGACGAAGCC CTGAAGAAAG CTCAGCGTCT GATTGAGCTG
    1081  GGCGGCAGCG GTCACACGAG CAGCCTGTAT ATCGATAGCC AAAACAATAA GGACAAGGTG
    1141  AAAGAGTTCG GTCTGGCGAT GAAAACCTCT CGCACGTTTA TCAACATGCC GAGCAGCCAA
    1201  GGTGCGAGCG GCGATCTGTA TAACTTCGCA ATCGCGCCTT CTTTTACGCT GGGTTGCGGT
    1261  ACCTGGGAGG GTAATTCGGT GAGCCAGAAT GTCGAGCCGA AGCACCTGCT GAACATCAAA
    1321  TCGGTCGCGG AGCGCCGTGA GAAGAGACCG TGGTTCAAAG TCCCACAGAA AATTTACTTC
    1381  AAATACGGCT GCCTGCGTTT CGCGCTGAAA GAACTGAAAG ACATGAACAA AAAGCGTGCG
    1441  TTCATTGTTA CCGACAAAGA CCTGTTCAAA CTGGGTTACG TGAACAAAAT CACCAAAGTT
    1501  CTGGATGAAA TTGACATCAA GTACTCCATC TTCACTGATA TCAAATCCGA CCCAACGATT
    1561  GATAGCGTGA AAAAGGGCGC TAAAGAAATG CTGAACTTTG AACCGGACAC CATCATCAGC
    1621  ATCGGTGGTG GCTCTCCTAT GGATGCTGCG AAGGTCATGC ACCTGCTGTA CGAATACCCG
    1681  GAAGCGGAAA TCGAAAACCT GGCTATCAAC TTCATGGACA TCCGCAAACG TATCTGCAAC
    1741  TTCCCGAAGC TGGGCACTAA AGCTATTTCC GTTGCCATCC CGACTACCGC GGGCACTGGT
    1801  TCCGAAGCCA CGCCGTTCGC CGTGATCACC AACGATGAAA CCGGTATGAA ATACCCGCTG
    1861  ACCTCTTACG AACTGACCCC GAACATGGCA ATTATCGACA CCGAGCTGAT GCTGAACATG
    1921  CCGCGCAAGC TGACCGCTGC TACCGGCATC GACGCTCTGG TACATGCTAT TGAGGCGTAC
    1981  GTTTCCGTGA TGGCTACCGA TTACACCGAC GAACTGGCCC TGCGTGCGAT CAAAATGATT
    2041  TTCAAGTACC TGCCTCGCGC TTACAAAAAC GGCACGAATG ACATCGAGGC GCGTGAGAAA
    2101  ATGGCCCATG CAAGCAACAT CGCGGGCATG GCCTTCGCCA ACGCGTTCCT GGGCGTGTGC
    2161  CACTCTATGG CTCACAAACT GGGTGCTATG CACCACGTGC CGCACGGTAT CGCGTGTGCT
    2221  GTCCTGATCG AAGAAGTAAT TAAGTACAAC GCTACTGATT GCCCGACTAA ACAGACCGCC
    2281  TTCCCACAGT ACAAATCTCC TAACGCTAAA CGTAAGTACG CTGAGATCGC CGAATACCTG
    2341  AACCTGAAGG GTACGAGCGA CACTGAGAAA GTTACTGCGC TGATCGAAGC TATCTCTAAA
    2401  CTGAAAATTG ACCTGTCCAT CCCGCAGAAC ATCAGCGCCG CAGGCATCAA CAAAAAGGAC
    2461  TTTTACAACA CGCTGGACAA AATGAGCGAA CTGGCTTTTG ACGACCAGTG CACCACTGCA
2521  AACCCGCGTT ACCCGCTGAT CTCTGAGCTG AAAGACATTT ATATCAAATC TTTTTAA
```

**AdhE2.4 (76933)**

```
       1  ATGAAAGTGA CGAACCAAAA AGAACTGAAA CAAAAACTGA ATGAACTGCG TGAGGCCCAG
      61  AAAAAGTTCG CTACGTATAC CCAGGAGCAA GTTGACAAGA TCTTCAAACA GTGTGCCATT
```

121 GCGGCTGCCA AAGAGCGTAT TAACTTGGCC AAACTGGCGG TGGAAGAAAC CGGCATCGGT

181 CTGGTCGAGG ACAAGATTAT CAAGAATCAC TTTGCCGCGG AGTATATCTA CAACAAATAC

```
     241  AAAAACGAAA AGACTTGCGG CATCATTGAT CATGATGACA GCTTGGGTAT CACGAAAGTC
     301  GCAGAACCAA TCGGTATTGT CGCGGCAATT GTCCCGACCA CTAATCCGAC CAGCACCGCG
     361  ATCTTCAAGA GCTTGATCAG CCTGGCGACC CGCAACGCGA TTTTCTTCAG CCCGCACCCG
     421  CGTGCGAAGA AGAGCACCAT CGCAGCGGCG AAGCTGATTT TGGATGCCGC GGTTAAGGCC
     481  GGTGCACCGA AAAACATTAT TGGCTGGATT GACGAGCCGA GCATTGAGCT GTCTCAGGAC
     541  CTGATGAGCG AAGCGGACAT TATTCTGGCC ACCGGTGGTC CGTCTATGGT TAAGGCAGCG
     601  TACTCCAGCG GTAAGCCTGC TATCGGCGCG GGTGCGGGCA ATACGCCGGC TATCATCGAT
     661  GAAAGCGCGG ATATTGATAT GGCAGTTAGC TCCATCATTC TGTCCAAAAC CTACGACAAT
     721  GGTGTTATCT GTGCAAGCGA GCAAAGCATC CTGGTGATGA ATAGCATTTA CGAGAAGGTC
     781  AAAGAAGAGT TTGTTAAGCG TGGTTCCTAT ATTCTGAACC AGAACGCGAT TGCAAAAATC
     841  AAAGAAACGA TGTTTAAGAA TGGCGCGATT AATGCAGATA TTGTTGGTAA GTCCGCATAC
     901  ATTATCGCGA AGATGGCGGG TATCGAAGTG CCGCAAACCA CCAAAATCTT GATCGGCGAA
     961  GTGCAGAGCG TGGAGAAGAG CGAACTGTTC AGCCATGAGA AACTGAGCCC GGTTCTGGCT
    1021  ATGTACAAAG TGAAGGACTT TGACGAAGCC CTGAAGAAAG CTCAGCGTCT GATTGAGCTG
    1081  GGCGGCAGCG GTCACACGAG CAGCCTGTAT ATCGATAGCC AAAACAATAA GGACAAGGTG
    1141  AAAGAGTTCG GTCTGGCGAT GAAAACCTCT CGCACGTTTA TCAACATGCC GAGCAGCCAA
    1201  GGTGCGAGCG GCGATCTGTA TAACTTCGCA ATCGCGCCTT CTTTTACGCT GGGTTGCGGT
    1261  ACCTGGGGCG GTAATTCGGT GAGCCAGAAT GTCGAGCCGA AGCACCTGCT GAACATCAAA
    1321  TCGGTCGCGG AGCGCCGTGA GAAGAGACCG TGGTTCAAAG TCCCACAGAA AATTTACTTC
    1381  AAATACGGCT GCCTGCGTTT CGCGCTGAAA GAACTGAAAG ACATGAACAA AAAGCGTGCG
    1441  TTCATTGTTA CCGACAAAGA CCTGTTCAAA CTGGGTTACG TGAACAAAAT CACCAAAGTT
    1501  CTGGATGAAA TTGACATCAA GTACTCCATC TTCACTGATA TCAAATCCGA CCCAACGATT
    1561  GATAGCGTGA AAAAGGGCGC TAAAGAAATG CTGAACTTTG AACCGGACAC CATCATCAGC
    1621  ATCGGTGGTG GCTCTCCTAT GGATGCTGCG AAGGTCATGC ACCTGCTGTA CGAATACCCG
    1681  GAAGCGGAAA TCGAAAACCT GGCTATCAAC TTCATGGACA TCCGCAAACG TATCTGCAAC
    1741  TTCCCGAAGC TGGGCACTAA AGCTATTTCC GTTGCCATCC CGACTACCGC GGGCACTGGT
    1801  TCCGAAGCCA CGCCGTTCGC CGTGATCACC AACGATGAAA CCGGTATGAA ATACCCGCTG
    1861  ACCTCTTACG AACTGACCCC GAACATGGCA ATTATCGACA CCGAGCTGAT GCTGAACATG
    1921  CCGCGCAAGC TGACCGCTGC TACCGGCATC GACGCTCTGG TACATGCTAT TGAGGCGTAC
    1981  GTTTCCGTGA TGGCTACCGA TTACACCGAC GAACTGGCCC TGCGTGCGAT CAAAATGATT
    2041  TTCAAGTACC TGCCTCGCGC TTACAAAAAC GGCACGAATG ACATCGAGGC GCGTGAGAAA
    2101  ATGGCCCATG CAAGCAACAT CGCGGGCATG GCCTTCGCCA ACGCGTTCCT GGGCGTGTGC
    2161  CACTCTATGG CTCACAAACT GGGTGCTATG CACCACGTGC CGCACGGTAT CGCGTGTGCT
    2221  GTCCTGATCG AAGAAGTAAT TAAGTACAAC GCTACTGATT GCCCGACTAA ACAGACCGCC
    2281  TTCCCACAGT ACAAATCTCC TAACGCTAAA CGTAAGTACG CTGAGATCGC CGAATACCTG
    2341  AACCTGAAGG GTACGAGCGA CACTGAGAAA GTTACTGCGC TGATCGAAGC TATCTCTAAA
    2401  CTGAAAATTG ACCTGTCCAT CCCGCAGAAC ATCAGCGCCG CAGGCATCAA CAAAAAGGAC
    2461  TTTTACAACA CGCTGGACAA AATGAGCGAA CTGGCTTTTG ACGACCAGTG CACCACTGCA
2521  AACCCGCGTT ACCCGCTGAT CTCTGAGCTG AAAGACATTT ATATCAAATC TTTTTAA
```

**AdhE2.5 (76934)**

```
       1  ATGAAAGTGA CGAACCAAAA AGAACTGAAA CAAAAACTGA ATGAACTGCG TGAGGCCCAG
      61  AAAAAGTTCG CTTGCTATAC CCAGGACCAA GTTGACAAGA TCTTCAAACA GTGTGCCATT
```

121 GCGGCTGCCA AAGAGCGTAT TAACTTGGCC GAGCTGGCGG TGGAAGAAAC CGGCATCGGT

181 CTGGTCGAGG ACAAGATTAT CAAGAATCAC TTTGCCGCGG AGTATATCTA CAACAAATAC

```
     241  AAAAACGAAA AGACTTGCGG CATCATTGAT CATGATGACA GCTTGGGTAT CACGAAAGTC
     301  GCAGAACCAA TCGGTATTGT CGCGGCAATT GTCCCGACCA CTAATCCGAC CAGCACCGCG
     361  ATCTTCAAGA GCTTGATCAG CCTGAAAACC CGCAACGCGA TTTTCTTCAG CCCGCACCCG
     421  CGTGCGAAGA AGAGCACCAT CGCAGCGGCG AAGCTGATTT TGGATGCCGC GGTTAAGGCC
     481  GGTGCACCGA AAAACATTAT TGGCTGGATT GACGAGCCGA GCATTGAGCT GTCTCAGGAC
     541  CTGATGAGCG AAGCGGACAT TATTCTGGCC ACCGGTGGTC CGTCTATGGT TAAGGCAGCG
     601  TACTCCAGCG GTAAGCCTGC TATCGGCGTT GGTGCGGGCA ATACGCCGGC TATCATCGAT
     661  GAAAGCGCGG ATATTGATAT GGCAGTTAGC TCCATCATTC TGTCCAAAAC CTACGACAAT
     721  GGTGTTATCT GTGCAAGCGA GCAAAGCATC CTGGTGATGA ATAGCATTTA CGAGAAGGTC
     781  AAAGAAGAGT TTGTTAAGCG TGGTTCCTAT ATTCTGAACC AGAACGAGAT TGCAAAAATC
     841  AAAGAAACGA TGTTTAAGAA TGGCGCGATT AATGCAGATA TTGTTGGTAA GTCCGCATAC
     901  ATTATCGCGA AGATGGCGGG TATCGAAGTG CCGCAAACCA CCAAAATCTT GATCGGCGAA
     961  GTGCAGAGCG TGGAGAAGAG CGAACTGTTC AGCCATGAGA AACTGAGCCC GGTTCTGGCT
    1021  ATGTACAAAG TGAAGGACTT TGACGAAGCC CTGAAGAAAG CTCAGCGTCT GATTGAGCTG
    1081  GGCGGCAGCG GTCACACGAG CAGCCTGTAT ATCGATAGCC AAAACAATAA GGACAAGGTG
    1141  AAAGAGTTCG GTCTGGCGAT GAAAACCTCT CGCACGTTTA TCAACATGCC GAGCAGCCAA
    1201  GGTGCGAGCG GCGATCTGTA TAACTTCGCA ATCGCGCCTT CTTTTACGCT GGGTTGCGGT
    1261  ACCTGGGGCG GTAATTCGGT GAGCCAGAAT GTCGAGCCGA AGCACCTGCT GAACATCAAA
    1321  TCGGTCGCGG AGCGCCGTGA GAAGAGACCG TGGTTCAAAG TCCCACAGAA AATTTACTTC
    1381  AAATACGGCT GCCTGCGTTT CGCGCTGAAA GAACTGAAAG ACATGAACAA AAAGCGTGCG
    1441  TTCATTGTTA CCGACAAAGA CCTGTTCAAA CTGGGTTACG TGAACAAAAT CACCAAAGTT
    1501  CTGGATGAAA TTGACATCAA GTACTCCATC TTCACTGATA TCAAATCCGA CCCAACGATT
    1561  GATAGCGTGA AAAAGGGCGC TAAAGAAATG CTGAACTTTG AACCGGACAC CATCATCAGC
    1621  ATCGGTGGTG GCTCTCCTAT GGATGCTGCG AAGGTCATGC ACCTGCTGTA CGAATACCCG
    1681  GAAGCGGAAA TCGAAAACCT GGCTATCAAC TTCATGGACA TCCGCAAACG TATCTGCAAC
    1741  TTCCCGAAGC TGGGCACTAA AGCTATTTCC GTTGCCATCC CGACTACCGC GGGCACTGGT
    1801  TCCGAAGCCA CGCCGTTCGC CGTGATCACC AACGATGAAA CCGGTATGAA ATACCCGCTG
    1861  ACCTCTTACG AACTGACCCC GAACATGGCA ATTATCGACA CCGAGCTGAT GCTGAACATG
    1921  CCGCGCAAGC TGACCGCTGC TACCGGCATC GACGCTCTGG TACATGCTAT TGAGGCGTAC
    1981  GTTTCCGTGA TGGCTACCGA TTACACCGAC GAACTGGCCC TGCGTGCGAT CAAAATGATT
    2041  TTCAAGTACC TGCCTCGCGC TTACAAAAAC GGCACGAATG ACATCGAGGC GCGTGAGAAA
    2101  ATGGCCCATG CAAGCAACAT CGCGGGCATG GCCTTCGCCA ACGCGTTCCT GGGCGTGTGC
    2161  CACTCTATGG CTCACAAACT GGGTGCTATG CACCACGTGC CGCACGGTAT CGCGTGTGCT
    2221  GTCCTGATCG AAGAAGTAAT TAAGTACAAC GCTACTGATT GCCCGACTAA ACAGACCGCC
    2281  TTCCCACAGT ACAAATCTCC TAACGCTAAA CGTAAGTACG CTGAGATCGC CGAATACCTG
    2341  AACCTGAAGG GTACGAGCGA CACTGAGAAA GTTACTGCGC TGATCGAAGC TATCTCTAAA
    2401  CTGAAAATTG ACCTGTCCAT CCCGCAGAAC ATCAGCGCCG CAGGCATCAA CAAAAAGGAC
    2461  TTTTACAACA CGCTGGACAA AATGAGCGAA CTGGCTTTTG ACGACCAGTG CACCACTGCA
    2521  AACCCGCGTT ACCCGCTGAT CTCTGAGCTG AAAGACATTT ATATCAAATC TTTTTAA
```

**AdhE2.7 (76936)**

```
       1  ATGAAAGTGA CGAACCAAAA AGAACTGAAA CAAAAACTGA ATGAACTGCG TGAGGCCCAG
      61  AAAAAGTTCG CTACGTATAC CCAGGAGCAA GTTGACAAGT TCTTCAAACA GTGTGCCATT
```

121 GCGGCTGCCA AAGAGCGTAT TAACTTGGCC AAACTGGCGG TGGAAGAAAC CGGCATCGGT

181 CTGGTCGAGG ACAAGATTAT CAAGAATCAC TTTGCCGCGG AGTATATCTA CAACAAATAC

```
     241  AAAAACGAAA AGACTTGCGG CATCATTGAT CATGATGACA GCTTGGGTAT CACGAAAGTC
     301  GCAGAACCAA TCGGTATTGT CGCGGCAATT GTCCCGACCA CTAATCCGAC CAGCACCGCG
     361  ATCTTCAAGA GCTTGATCAG CCTGAAAACC CGCAACGCGA TTTTCTTCAG CCCGCACCCG
     421  CGTGCGAAGA AGAGCACCAT CGCAGCGGCG AAGCTGATTT TGGATGCCGC GGTTAAGGCC
     481  GGTGCACCGA AAAACATTAT TGGCTGGATT GACGAGCCGA GCATTGAGCT GTCTCAGGAC
     541  CTGATGAGCG AAGCGGACAT TATTCTGGCC ACCGGTGGTC CGTCTATGGT TAAGGCAGCG
     601  TACTCCAGCG GTAAGCCTGC TATCGGCGTT GGTGCGGGCA ATACGCCGGC TATCATCGAT
     661  GAAAGCGCGG ATATTGATAT GGCAGTTAGC TCCATCATTC TGTCCAAAAC CTACGACAAT
     721  GGTGTTATCT GTGCAAGCGA GCAAAGCATC CTGGTGATGA ATAGCATTTA CGAGAAGGTC
     781  AAAGAAGAGT TTGTTAAGCG TGGTTCCTAT ATTCTGAACC AGAACGAGAT TGCAAAAATC
     841  AAAGAAACGA TGTTTAAGAA TGGCGCGATT AATGCAGATA TTGTTGGTAA GTCCGCATAC
     901  ATTATCGCGA AGATGGCGGG TATCGAAGTG CCGCAAACCA CCAAAATCTT GATCGGCGAA
     961  GTGCAGAGCG TGGAGAAGAG CGAACTGTTC AGCCATGAGA AACTGAGCCC GGTTCTGGCT
    1021  ATGTACAAAG TGAAGGACTT TGACGAAGCC CTGAAGAAAG CTCAGCGTCT GATTGAGCTG
    1081  GGCGGCAGCG GTCACACGAG CAGCCTGTAT ATCGATAGCC AAAACAATAA GGACAAGGTG
    1141  AAAGAGTTCG GTCTGGAGAT GAAAACCTCT CGCTTCTTTA TCAACATGCC GAGCAGCCAA
    1201  GGTGCGAGCG GCGATCTGTA TAACTTCGCA ATCGCGCCTT CTTTTACGCT GGGTTGCGGT
    1261  ACCTGGGGCG GTAATTCGGT GAGCCAGAAT GTCGAGCCGA AGCACCTGCT GAACATCAAA
    1321  TCGGTCGCGG AGCGCCGTGA GAAGAGACCG TGGTTCAAAG TCCCACAGAA AATTTACTTC
    1381  AAATACGGCT GCCTGCGTTT CGCGCTGAAA GAACTGAAAG ACATGAACAA AAAGCGTGCG
    1441  TTCATTGTTA CCGACAAAGA CCTGTTCAAA CTGGGTTACG TGAACAAAAT CACCAAAGTT
    1501  CTGGATGAAA TTGACATCAA GTACTCCATC TTCACTGATA TCAAATCCGA CCCAACGATT
    1561  GATAGCGTGA AAAAGGGCGC TAAAGAAATG CTGAACTTTG AACCGGACAC CATCATCAGC
    1621  ATCGGTGGTG GCTCTCCTAT GGATGCTGCG AAGGTCATGC ACCTGCTGTA CGAATACCCG
    1681  GAAGCGGAAA TCGAAAACCT GGCTATCAAC TTCATGGACA TCCGCAAACG TATCTGCAAC
    1741  TTCCCGAAGC TGGGCACTAA AGCTATTTCC GTTGCCATCC CGACTACCGC GGGCACTGGT
    1801  TCCGAAGCCA CGCCGTTCGC CGTGATCACC AACGATGAAA CCGGTATGAA ATACCCGCTG
    1861  ACCTCTTACG AACTGACCCC GAACATGGCA ATTATCGACA CCGAGCTGAT GCTGAACATG
    1921  CCGCGCAAGC TGACCGCTGC TACCGGCATC GACGCTCTGG TACATGCTAT TGAGGCGTAC
    1981  GTTTCCGTGA TGGCTACCGA TTACACCGAC GAACTGGCCC TGCGTGCGAT CAAAATGATT
    2041  TTCAAGTACC TGCCTCGCGC TTACAAAAAC GGCACGAATG ACATCGAGGC GCGTGAGAAA
    2101  ATGGCCCATG CAAGCAACAT CGCGGGCATG GCCTTCGCCA ACGCGTTCCT GGGCGTGTGC
    2161  CACTCTATGG CTCACAAACT GGGTGCTATG CACCACGTGC CGCACGGTAT CGCGTGTGCT
    2221  GTCCTGATCG AAGAAGTAAT TAAGTACAAC GCTACTGATT GCCCGACTAA ACAGACCGCC
    2281  TTCCCACAGT ACAAATCTCC TAACGCTAAA CGTAAGTACG CTGAGATCGC CGAATACCTG
    2341  AACCTGAAGG GTACGAGCGA CACTGAGAAA GTTACTGCGC TGATCGAAGC TATCTCTAAA
    2401  CTGAAAATTG ACCTGTCCAT CCCGCAGAAC ATCAGCGCCG CAGGCATCAA CAAAAAGGAC
    2461  TTTTACAACA CGCTGGACAA AATGAGCGAA CTGGCTTTTG ACGACCAGTG CACCACTGCA
    2521  AACCCGCGTT ACCCGCTGAT CTCTGAGCTG AAAGACATTT ATATCAAATC TTTTTAA
```

**AdhE2.8 (76937)**

```
       1  ATGAAAGTGA CGAACCAAAA AGAACTGAAA CAAAAACTGA ATGAACTGCG TGAGGCCCAG
      61  AAAAAGTTCG CTACGTATAC CCAGGAGCAA GTTGACAAGA TCTTCAAACA GTGTGCCATT
```

121 GCGGCTGCCA AAGAGCGTAT TAACTTGGCC AAACTGGCGG TGGAAGAAAC CGGCATCGGT

181 CTGGTCGAGG ACAAGATTAT CAAGAATCAC TTTGCCGCGG AGTATATCTA CAACAAATAC

```
     241  AAAAACGAAA AGACTTGCGG CATCATTGAT CATGATGACA GCTTGGGTAT CACGAAAGTC
     301  GCAGAACCAA TCGGTATTGT CGCGGCAATT GTCCCGACCA CTAATCCGAC CAGCACCGCG
     361  ATCTTCAAGA GCTTGATCAG CCTGAAAACC CGCAACGCGA TTTTCTTCAG CCCGCACCCG
     421  CGTGCGAAGA AGAGCACCAT CGCAGCGGCG AAGCTGATTT TGGATGCCGC GGTTAAGGCC
     481  GGTGCACCGA AAAACATTAT TGGCTGGATT GACGAGCCGA GCATTGAGCT GTCTCAGGAC
     541  CTGATGAGCG AAGCGGACAT TATTCTGGCC ACCGGTGGTC CGTCTATGGT TAAGGCAGCG
     601  TACTCCAGCG GTAAGCCTGC TATCGGCGTT GGTGCGGGCA ATACGCCGGC TATCATCGAT
     661  GAAAGCGCGG ATATTGATAT GGCAGTTAGC TCCATCATTC TGTCCAAAAC CTACGACAAT
     721  GGTGTTATCT GTGCAAGCGA GCAAAGCATC CTGGTGATGA ATAGCATTTA CGAGAAGGTC
     781  AAAGCGGAGT TTGTTAAGCG TGGTTCCTAT ATTCTGAACC AGGAGGAGAT TGCAAAAATC
     841  AAAGAAACGA TGTTTAAGAA TGGCGCGATT AATGCAGATA TTGTTGGTAA GTCCGCATAC
     901  ATTATCGCGA AGATGGCGGG TATCGAAGTG CCGCAAACCA CCAAAATCTT GATCGGCGAA
     961  GTGCAGAGCG TGGAGAAGAG CGAACTGTTC AGCCATGAGA AACTGAGCCC GGTTCTGGCT
    1021  ATGTACAAAG TGAAGGACTT TGACGAAGCC CTGAAGCTGG CTCAGCGTCT GATTGAGCTG
    1081  GGCGGCAGCG GTCACACGAG CAGCCTGTAT ATCGATAGCC AAAACAATAA GGACAAGGTG
    1141  AAAGAGTTCG GTCTGGCGAT GAAAACCTCT CGCACGTTTA TCAACATGCC GAGCAGCCAA
    1201  GGTGCGAGCG GCGATCTGTA TAACTTCGCA ATCGCGCCTT CTTTTACGCT GGGTTGCGGT
    1261  ACCTGGGGCG GTAATTCGGT GAGCCAGAAT GTCGAGCCGA AGCACCTGCT GAACATCAAA
    1321  TCGGTCGCGG AGCGCCGTGA GAAGAGACCG TGGTTCAAAG TCCCACAGAA AATTTACTTC
    1381  AAATACGGCT GCCTGCGTTT CGCGCTGAAA GAACTGAAAG ACATGAACAA AAAGCGTGCG
    1441  TTCATTGTTA CCGACAAAGA CCTGTTCAAA CTGGGTTACG TGAACAAAAT CACCAAAGTT
    1501  CTGGATGAAA TTGACATCAA GTACTCCATC TTCACTGATA TCAAATCCGA CCCAACGATT
    1561  GATAGCGTGA AAAAGGGCGC TAAAGAAATG CTGAACTTTG AACCGGACAC CATCATCAGC
    1621  ATCGGTGGTG GCTCTCCTAT GGATGCTGCG AAGGTCATGC ACCTGCTGTA CGAATACCCG
    1681  GAAGCGGAAA TCGAAAACCT GGCTATCAAC TTCATGGACA TCCGCAAACG TATCTGCAAC
    1741  TTCCCGAAGC TGGGCACTAA AGCTATTTCC GTTGCCATCC CGACTACCGC GGGCACTGGT
    1801  TCCGAAGCCA CGCCGTTCGC CGTGATCACC AACGATGAAA CCGGTATGAA ATACCCGCTG
    1861  ACCTCTTACG AACTGACCCC GAACATGGCA ATTATCGACA CCGAGCTGAT GCTGAACATG
    1921  CCGCGCAAGC TGACCGCTGC TACCGGCATC GACGCTCTGG TACATGCTAT TGAGGCGTAC
    1981  GTTTCCGTGA TGGCTACCGA TTACACCGAC GAACTGGCCC TGCGTGCGAT CAAAATGATT
    2041  TTCAAGTACC TGCCTCGCGC TTACAAAAAC GGCACGAATG ACATCGAGGC GCGTGAGAAA
    2101  ATGGCCCATG CAAGCAACAT CGCGGGCATG GCCTTCGCCA ACGCGTTCCT GGGCGTGTGC
    2161  CACTCTATGG CTCACAAACT GGGTGCTATG CACCACGTGC CGCACGGTAT CGCGTGTGCT
    2221  GTCCTGATCG AAGAAGTAAT TAAGTACAAC GCTACTGATT GCCCGACTAA ACAGACCGCC
    2281  TTCCCACAGT ACAAATCTCC TAACGCTAAA CGTAAGTACG CTGAGATCGC CGAATACCTG
    2341  AACCTGAAGG GTACGAGCGA CACTGAGAAA GTTACTGCGC TGATCGAAGC TATCTCTAAA
    2401  CTGAAAATTG ACCTGTCCAT CCCGCAGAAC ATCAGCGCCG CAGGCATCAA CAAAAAGGAC
    2461  TTTTACAACA CGCTGGACAA AATGAGCGAA CTGGCTTTTG ACGACCAGTG CACCACTGCA
    2521  AACCCGCGTT ACCCGCTGAT CTCTGAGCTG AAAGACATTT ATATCAAATC TTTTTAA
```

**AdhE2.9 (76938)**

```
       1  ATGAAAGTGA CGAACCAAAA AGAACTGAAA CAAAAACTGA ATGAACTGCG TGAGGCCCAG
      61  AAAAAGTTCG CTACGTATAC CCAGGAGCAA GTTGACAAGA TCTTCAAACA GTGTGCCATT
```

121 GCGGCTGCCA AAGAGCGTAT TAACTTGGCC AAACTGGCGG TGGAAGAAAC CGGCATCGGT

181 CTGGTCGAGG ACAAGATTAT CAAGAATCAC TTTGCCGCGG AGTATATCTA CAACAAATAC

```
     241  AAAAACGAAA AGACTTGCGG CATCATTGAT CATGATGACA GCTTGGGTAT CACGAAAGTC
     301  GCAGAACCAA TCGGTATTGT CGCGGCAATT GTCCCGACCA CTAATCCGAC CAGCACCGCG
     361  ATCTTCAAGA GCTTGATCAG CCTGAAAACC CGCAACGCGA TTTTCTTCAG CCCGCACCCG
     421  CGTGCGAAGA AGAGCACCAT CGCAGCGGCG AAGCTGATTT TGGATGCCGC GGTTAAGGCC
     481  GGTGCACCGA AAAACATTAT TGGCTGGATT GACGAGCCGA GCATTGAGCT GTCTCAGGAC
     541  CTGATGAGCG AAGCGGACAT TATTCTGGCC ACCGGTGGTC CGTCTATGGT TAAGGCAGCG
     601  TACTCCAGCG GTAAGCCTGC TATCGGCGTT GGTGCGGGCA ATACGCCGGC TATCATCGAT
     661  GAAAGCGCGG ATATTGATAT GGCAGTTAGC TCCATCATTC TGTCCAAAAC CTACGACAAT
     721  GGTGTTATCT GTGCAAGCGA GCAAAGCATC CTGGTGATGA ATAGCATTTA CGAGAAGGTC
     781  AAAGAAGAGT TTGTTAAGCG TGGTTCCTAT ATTCTGAACC AGAACGAGAT TGCAAAAATC
     841  AAAGAAACGA TGTTTAAGAA TGGCGCGATT AATGCAGATA TTGTTGGTAA GTCCGCATAC
     901  ATTATCGCGA AGATGGCGGG TATCGAAGTG CCGCAATACA CCAAAATCTT GATCGGCGAA
     961  GTGCAGCACG TGGAGAAGAG CGAACTGTTC AGCCATGAGA AACTGAGCCC GGTTCTGGCT
    1021  ATGTACAAAG TGAAGGACTT TGACGAAGCC CTGAAGAAAG CTCAGCGTCT GATTGAGCTG
    1081  GGCGGCAGCG GTCACACGAG CAGCCTGTAT ATCGATAGCC AAAACAATAA GGACAAGGTG
    1141  AAAGAGTTCG GTCTGGCGAT GAAAACCTCT CGCACGTTTA TCAACATGCC GAGCAGCCAA
    1201  GGTGCGAGCG GCGATCTGTA TAACTTCGCA ATCGCGCCTT CTTTTACGCT GGGTTGCCCG
    1261  ACCTGGGGCG GTAATTCGGT GAGCCAGAAT GTCGAGCCGA AGCACCTGCT GAACATCAAA
    1321  TCGGTCGCGG AGCGCCGTGA GAAGAGACCG TGGTTCAAAG TCCCACAGAA AATTTACTTC
    1381  AAATACGGCT GCCTGCGTTT CGCGCTGAAA GAACTGAAAG ACATGAACAA AAAGCGTGCG
    1441  TTCATTGTTA CCGACAAAGA CCTGTTCAAA CTGGGTTACG TGAACAAAAT CACCAAAGTT
    1501  CTGGATGAAA TTGACATCAA GTACTCCATC TTCACTGATA TCAAATCCGA CCCAACGATT
    1561  GATAGCGTGA AAAAGGGCGC TAAAGAAATG CTGAACTTTG AACCGGACAC CATCATCAGC
    1621  ATCGGTGGTG GCTCTCCTAT GGATGCTGCG AAGGTCATGC ACCTGCTGTA CGAATACCCG
    1681  GAAGCGGAAA TCGAAAACCT GGCTATCAAC TTCATGGACA TCCGCAAACG TATCTGCAAC
    1741  TTCCCGAAGC TGGGCACTAA AGCTATTTCC GTTGCCATCC CGACTACCGC GGGCACTGGT
    1801  TCCGAAGCCA CGCCGTTCGC CGTGATCACC AACGATGAAA CCGGTATGAA ATACCCGCTG
    1861  ACCTCTTACG AACTGACCCC GAACATGGCA ATTATCGACA CCGAGCTGAT GCTGAACATG
    1921  CCGCGCAAGC TGACCGCTGC TACCGGCATC GACGCTCTGG TACATGCTAT TGAGGCGTAC
    1981  GTTTCCGTGA TGGCTACCGA TTACACCGAC GAACTGGCCC TGCGTGCGAT CAAAATGATT
    2041  TTCAAGTACC TGCCTCGCGC TTACAAAAAC GGCACGAATG ACATCGAGGC GCGTGAGAAA
    2101  ATGGCCCATG CAAGCAACAT CGCGGGCATG GCCTTCGCCA ACGCGTTCCT GGGCGTGTGC
    2161  CACTCTATGG CTCACAAACT GGGTGCTATG CACCACGTGC CGCACGGTAT CGCGTGTGCT
    2221  GTCCTGATCG AAGAAGTAAT TAAGTACAAC GCTACTGATT GCCCGACTAA ACAGACCGCC
    2281  TTCCCACAGT ACAAATCTCC TAACGCTAAA CGTAAGTACG CTGAGATCGC CGAATACCTG
    2341  AACCTGAAGG GTACGAGCGA CACTGAGAAA GTTACTGCGC TGATCGAAGC TATCTCTAAA
    2401  CTGAAAATTG ACCTGTCCAT CCCGCAGAAC ATCAGCGCCG CAGGCATCAA CAAAAAGGAC
    2461  TTTTACAACA CGCTGGACAA AATGAGCGAA CTGGCTTTTG ACGACCAGTG CACCACTGCA
    2521  AACCCGCGTT ACCCGCTGAT CTCTGAGCTG AAAGACATTT ATATCAAATC TTTTTAA
```

**AdhE2.10 (76939)**

```
       1  ATGAAAGTGA CGAACCAAAA AGAACTGAAA CAAAAACTGA ATGAACTGCG TGAGGCCCAG
      61  AAAAAGTTCC TGACGTATAC CCAGGAGCAA GTTGACAAGA TCTTCAAACA GTGTGCCATT
```

121 GCGGCTGCCA AAGAGCGTAT TAACTTGGCC AAACTGGCGG TGGAAGAAAC CGGCATCGGT

181 CTGGTCGAGG ACAAGATTAT CAAGAATCAC TTTGCCGCGG AGTATATCTA CAACAAATAC

```
     241  AAAAACGAAA AGACTTGCGG CATCATTGAT CATGATGACA GCTTGGGTAT CACGAAAGTC
     301  GCAGAACCAA TCGGTATTGT CGCGGCAATT GTCCCGACCA CTAATCCGAC CAGCACCGCG
     361  ATCTTCAAGA GCTTGATCAG CCTGAAAACC CGCAACGCGA TTTTCTTCAG CCCGCACCCG
     421  CGTGCGAAGA AGAGCACCAT CGCAGCGGCG AAGCTGATTT TGGATGCCGC GGTTAAGGCC
     481  GGTGCACCGA AAAACATTAT TGGCTGGATT GACGAGCCGA GCATTGAGCT GTCTCAGGAC
     541  CTGATGAGCG AAGCGGACAT TATTCTGGCC ACCGGTGGTC CGTCTATGGT TAAGGCAGCG
     601  TACTCCAGCG GTAAGCCTGC TATCGGCGTT GGTGCGGGCA ATACGCCGGC TATCATCGAT
     661  GAAAGCGCGG ATATTGATAT GGCAGTTAGC TCCATCATTC TGTCCAAAAC CTACGACAAT
     721  GGTGTTATCT GTGCAAGCGA GCAAAGCATC CTGGTGATGA ATAGCATTTA CGAGAAGGTC
     781  AAAGAAGAGT TTGTTAAGCG TGGTTCCTAT ATTCTGAACC AGAACGAGAT TGCAAAAATC
     841  AAAGAAACGA TGTTTAAGAA TGGCGCGATT AATGCAGATA TTGTTGGTAA GTCCGCATAC
     901  ATTATCGCGA AGATGGCGGG TATCGAAGTG CCGCAAACCA CCAAAATCTT GATCGGCGAA
     961  GTGCAGAGCG TGGAGAAGAG CGAACTGTTC AGCCATGAGC TGCTGAGCCC GGTTCTGGCT
    1021  ATGTACAAAG TGAAGGACTT TGACGAAGCC CTGAAGAAAG CTCAGCGTCT GATTAACCTG
    1081  GGCGGCAGCG GTCACACGAG CAGCCTGTAT ATCGATAGCC AAAACAATAA GGACAAGGTG
    1141  AAAGAGTTCG GTCTGGCGAT GAAAACCTCT CGCACGTTTA TCAACATGCC GAGCAGCCAA
    1201  GGTGCGAGCG GCGATCTGTA TAACTTCGCA ATCGCGCCTT CTTTTACGCT GGGTTGCGGT
    1261  ACCTGGGGCG GTAATTCGGT GAGCCAGAAT GTCGAGCCGA AGCACCTGCT GAACATCAAA
    1321  TCGGTCGCGG AGCGCCGTGA GAAGAGACCG TGGTTCAAAG TCCCACAGAA AATTTACTTC
    1381  AAATACGGCT GCCTGCGTTT CGCGCTGAAA GAACTGAAAG ACATGAACAA AAAGCGTGCG
    1441  TTCATTGTTA CCGACAAAGA CCTGTTCAAA CTGGGTTACG TGAACAAAAT CACCAAAGTT
    1501  CTGGATGAAA TTGACATCAA GTACTCCATC TTCACTGATA TCAAATCCGA CCCAACGATT
    1561  GATAGCGTGA AAAAGGGCGC TAAAGAAATG CTGAACTTTG AACCGGACAC CATCATCAGC
    1621  ATCGGTGGTG GCTCTCCTAT GGATGCTGCG AAGGTCATGC ACCTGCTGTA CGAATACCCG
    1681  GAAGCGGAAA TCGAAAACCT GGCTATCAAC TTCATGGACA TCCGCAAACG TATCTGCAAC
    1741  TTCCCGAAGC TGGGCACTAA AGCTATTTCC GTTGCCATCC CGACTACCGC GGGCACTGGT
    1801  TCCGAAGCCA CGCCGTTCGC CGTGATCACC AACGATGAAA CCGGTATGAA ATACCCGCTG
    1861  ACCTCTTACG AACTGACCCC GAACATGGCA ATTATCGACA CCGAGCTGAT GCTGAACATG
    1921  CCGCGCAAGC TGACCGCTGC TACCGGCATC GACGCTCTGG TACATGCTAT TGAGGCGTAC
    1981  GTTTCCGTGA TGGCTACCGA TTACACCGAC GAACTGGCCC TGCGTGCGAT CAAAATGATT
    2041  TTCAAGTACC TGCCTCGCGC TTACAAAAAC GGCACGAATG ACATCGAGGC GCGTGAGAAA
    2101  ATGGCCCATG CAAGCAACAT CGCGGGCATG GCCTTCGCCA ACGCGTTCCT GGGCGTGTGC
    2161  CACTCTATGG CTCACAAACT GGGTGCTATG CACCACGTGC CGCACGGTAT CGCGTGTGCT
    2221  GTCCTGATCG AAGAAGTAAT TAAGTACAAC GCTACTGATT GCCCGACTAA ACAGACCGCC
    2281  TTCCCACAGT ACAAATCTCC TAACGCTAAA CGTAAGTACG CTGAGATCGC CGAATACCTG
    2341  AACCTGAAGG GTACGAGCGA CACTGAGAAA GTTACTGCGC TGATCGAAGC TATCTCTAAA
    2401  CTGAAAATTG ACCTGTCCAT CCCGCAGAAC ATCAGCGCCG CAGGCATCAA CAAAAAGGAC
    2461  TTTTACAACA CGCTGGACAA AATGAGCGAA CTGGCTTTTG ACGACCAGTG CACCACTGCA
    2521  AACCCGCGTT ACCCGCTGAT CTCTGAGCTG AAAGACATTT ATATCAAATC TTTTTAA
```

**AdhE2.11 (76940)**

```
       1  ATGAAAGTGA CGAACCAAAA AGAACTGAAA CAAAAACTGA ATGAACTGCG TGAGGCCCAG
      61  AAAAAGTTCG CTACGTATAC CCAGGACCAA GTTGACAAGA TCTTCAAACA GTGTGCCATT
```

121 GCGGCTGCCA AAGAGCGTAT TAACTTGGCC AAACTGGCGG TGGAAGAAAC CGGCATCGGT

181 CTGGTCGAGG ACAAGATTAT CAAGAATCAC TTTGCCGCGG AGTATATCTA CAACAAATAC

```
     241  AAAAACGAAA AGACTTGCGG CATCATTGAT CATGATGACA GCTTGGGTAT CACGAAAGTC
     301  GCAGAACCAA TCGGTATTGT CGCGGCAATT GTCCCGACCA CTAATCCGAC CAGCACCGCG
     361  ATCTTCAAGA GCTTGATCAG CCTGAAAACC CGCAACGCGA TTTTCTTCAG CCCGCACCCG
     421  CGTGCGAAGA AGAGCACCAT CGCAGCGGCG ACCCTGATTT TGGATGCCGC GGTTAAGGCC
     481  GGTGCACCGA AAAACATTAT TGGCTGGATT GACGAGCCGA GCATTGAGCT GTCTCAGGAC
     541  CTGATGAGCG AAGCGGACAT TATTCTGGCC ACCGGTGGTC CGTCTATGGT TAAGGCAGCG
     601  TACTCCAGCG GTAAGCCTGC TATCGGCGTT GGTGCGGGCA ATACGCCGGC TATCATCGAT
     661  GAAAGCGCGG ATATTGATAT GGCAGTTAGC TCCATCATTC TGTCCAAAAC CTACGACAAT
     721  GGTGTTATCT GTGCAAGCGA GCAAAGCATC CTGGTGATGA ATAGCATTTA CGAGAAGGTC
     781  AAAGAAGAGT TTGTTAAGCG TGGTTCCTAT ATTCTGAACC AGCAGGAGAT TGCAAAAATC
     841  AAAGAAACGA TGTTTAAGAA TGGCGCGATT AATGCAGATA TTGTTGGTAA GTCCGCATAC
     901  ATTATCGCGA AGATGGCGGG TATCGAAGTG CCGCAAACCA CCAAAATCTT GATCGGCGAA
     961  GTGCAGAGCG TGGAGAAGAG CGAACTGTTC AGCCATGAGA AACTGAGCCC GGTTCTGGCT
    1021  ATGTACAAAG TGAAGGACTT TGACGAAGCC CTGAAGAAAG CTCAGCGTCT GATTGAGCTG
    1081  GGCGGCAGCG GTCACACGAG CAGCCTGTAT ATCGATAGCC AAAACAATAA GGACAAGGTG
    1141  AAAGAGTTCG GTCTGGCGAT GAAAACCTCT CGCACGTTTA TCAACATGCC GAGCAGCCAA
    1201  GGTGCGAGCG GCGATCTGTA TAACTTCGCA ATCGCGCCTT CTTTTACGCT GGGTTGCGGT
    1261  ACCTGGGGCG GTAATTCGGT GAGCCAGAAT GTCGAGCCGA AGCACCTGCT GAACATCAAA
    1321  TCGGTCGCGG AGCGCCGTGA GAAGAGACCG TGGTTCAAAG TCCCACAGAA AATTTACTTC
    1381  AAATACGGCT GCCTGCGTTT CGCGCTGAAA GAACTGAAAG ACATGAACAA AAAGCGTGCG
    1441  TTCATTGTTA CCGACAAAGA CCTGTTCAAA CTGGGTTACG TGAACAAAAT CACCAAAGTT
    1501  CTGGATGAAA TTGACATCAA GTACTCCATC TTCACTGATA TCAAATCCGA CCCAACGATT
    1561  GATAGCGTGA AAAAGGGCGC TAAAGAAATG CTGAACTTTG AACCGGACAC CATCATCAGC
    1621  ATCGGTGGTG GCTCTCCTAT GGATGCTGCG AAGGTCATGC ACCTGCTGTA CGAATACCCG
    1681  GAAGCGGAAA TCGAAAACCT GGCTATCAAC TTCATGGACA TCCGCAAACG TATCTGCAAC
    1741  TTCCCGAAGC TGGGCACTAA AGCTATTTCC GTTGCCATCC CGACTACCGC GGGCACTGGT
    1801  TCCGAAGCCA CGCCGTTCGC CGTGATCACC AACGATGAAA CCGGTATGAA ATACCCGCTG
    1861  ACCTCTTACG AACTGACCCC GAACATGGCA ATTATCGACA CCGAGCTGAT GCTGAACATG
    1921  CCGCGCAAGC TGACCGCTGC TACCGGCATC GACGCTCTGG TACATGCTAT TGAGGCGTAC
    1981  GTTTCCGTGA TGGCTACCGA TTACACCGAC GAACTGGCCC TGCGTGCGAT CAAAATGATT
    2041  TTCAAGTACC TGCCTCGCGC TTACAAAAAC GGCACGAATG ACATCGAGGC GCGTGAGAAA
    2101  ATGGCCCATG CAAGCAACAT CGCGGGCATG GCCTTCGCCA ACGCGTTCCT GGGCGTGTGC
    2161  CACTCTATGG CTCACAAACT GGGTGCTATG CACCACGTGC CGCACGGTAT CGCGTGTGCT
    2221  GTCCTGATCG AAGAAGTAAT TAAGTACAAC GCTACTGATT GCCCGACTAA ACAGACCGCC
    2281  TTCCCACAGT ACAAATCTCC TAACGCTAAA CGTAAGTACG CTGAGATCGC CGAATACCTG
    2341  AACCTGAAGG GTACGAGCGA CACTGAGAAA GTTACTGCGC TGATCGAAGC TATCTCTAAA
    2401  CTGAAAATTG ACCTGTCCAT CCCGCAGAAC ATCAGCGCCG CAGGCATCAA CAAAAAGGAC
    2461  TTTTACAACA CGCTGGACAA AATGAGCGAA CTGGCTTTTG ACGACCAGTG CACCACTGCA
    2521  AACCCGCGTT ACCCGCTGAT CTCTGAGCTG AAAGACATTT ATATCAAATC TTTTTAA
```

**AdhE2.12 (76941)**

```
       1  ATGAAAGTGA CGAACCAAAA AGAACTGAAA CAAAAACTGA ATGAACTGCG TGAGGCCCAG
      61  AAAAAGTTCG CTAGCTATAC CCAGGAGCAA GTTGACAAGA TCTTCAAACA GTGTGCCATT
```

121 GCGGCTGCCA AAGAGCGTAT TAACTTGGCC AAACTGGCGG TGGAAGAAAC CGGCATCGGT

181 CTGGTCGAGG ACAAGATTAT CAAGAATCAC TTTGCCGCGG AGTATATCTA CAACAAATAC

```
     241  AAAAACGAAA AGACTTGCGG CATCATTGAT CATGATGACA GCTTGGGTAT CACGAAAGTC
     301  GCAGAACCAA TCGGTATTGT CGCGAGCATT GTCCCGACCA CTAATCCGAC CAGCACCGCG
     361  ATCTTCAAGA GCTTGATCAG CCTGAAAACC CGCAACGCGA TTTTCTTCAG CCCGCACCCG
     421  CGTGCGAAGA AGAGCACCAT CGCAGCGGCG AAGCTGATTT TGGATGCCGC GGTTAAGGCC
     481  GGTGCACCGA AAAACATTAT TGGCTGGATT GACGAGCCGA GCATTGAGCT GTCTCAGGAC
     541  CTGATGAGCG AAGCGGACAT TATTCTGGCC ACCGGTGGTC CGTCTATGGT TAAGGCAGCG
     601  TACTCCAGCG GTAAGCCTGC TATCGGCGTT GGTGCGGGCA ATACGCCGGC TATCATCGAT
     661  GAAAGCGCGG ATATTGATAT GGCAGTTAGC TCCATCATTC TGGGTAAAAC CTACGACAAT
     721  GGTGTTATCT GTGCAAGCGA GCAAAGCATC CTGGTGATGA ATAGCATTTA CGAGAAGGTC
     781  AAAGAAGAGT TTGTTAAGCG TGGTTCCTAT ATTCTGAACC AGAACGAGAT TGCAAAAATC
     841  AAAGAAACGA TGTTTAAGAA TGGCGCGATT AATGCAGATA TTGTTGGTAA GTCCGCATAC
     901  ATTATCGCGA AGATGGCGGG TATCGAAGTG CCGCAAACCA CCAAAATCTT GATCGGCGAA
     961  GTGCAGAGCG TGGAGAAGAG CGAACTGTTC AGCCATGAGA AACTGAGCCC GGTTCTGGCT
    1021  ATGTACAAAG TGAAGGACTT TGACGAAGCC CTGAAGAAAG CTCAGCGTCT GATTGAGCTG
    1081  GGCGGCAGCG GTCACACGAG CAGCCTGTAT ATCGATAGCC AAAACAATAA GGACAAGGTG
    1141  AAAGAGTTCG GTCTGGCGAT GAAAACCTCT CGCACGTTTA TCAACATGCC GAGCAGCCAA
    1201  GGTGCGAGCG GCGATCTGTA TAACTTCGCA ATCGCGCCTT CTTTTACGCT GGGTTGCGGT
    1261  ACCTGGGGCG GTAATTCGGT GAGCCAGAAT GTCGAGCCGA AGCACCTGCT GAACATCAAA
    1321  TCGGTCGCGG AGCGCCGTGA GAAGAGACCG TGGTTCAAAG TCCCACAGAA AATTTACTTC
    1381  AAATACGGCT GCCTGCGTTT CGCGCTGAAA GAACTGAAAG ACATGAACAA AAAGCGTGCG
    1441  TTCATTGTTA CCGACAAAGA CCTGTTCAAA CTGGGTTACG TGAACAAAAT CACCAAAGTT
    1501  CTGGATGAAA TTGACATCAA GTACTCCATC TTCACTGATA TCAAATCCGA CCCAACGATT
    1561  GATAGCGTGA AAAAGGGCGC TAAAGAAATG CTGAACTTTG AACCGGACAC CATCATCAGC
    1621  ATCGGTGGTG GCTCTCCTAT GGATGCTGCG AAGGTCATGC ACCTGCTGTA CGAATACCCG
    1681  GAAGCGGAAA TCGAAAACCT GGCTATCAAC TTCATGGACA TCCGCAAACG TATCTGCAAC
    1741  TTCCCGAAGC TGGGCACTAA AGCTATTTCC GTTGCCATCC CGACTACCGC GGGCACTGGT
    1801  TCCGAAGCCA CGCCGTTCGC CGTGATCACC AACGATGAAA CCGGTATGAA ATACCCGCTG
    1861  ACCTCTTACG AACTGACCCC GAACATGGCA ATTATCGACA CCGAGCTGAT GCTGAACATG
    1921  CCGCGCAAGC TGACCGCTGC TACCGGCATC GACGCTCTGG TACATGCTAT TGAGGCGTAC
    1981  GTTTCCGTGA TGGCTACCGA TTACACCGAC GAACTGGCCC TGCGTGCGAT CAAAATGATT
    2041  TTCAAGTACC TGCCTCGCGC TTACAAAAAC GGCACGAATG ACATCGAGGC GCGTGAGAAA
    2101  ATGGCCCATG CAAGCAACAT CGCGGGCATG GCCTTCGCCA ACGCGTTCCT GGGCGTGTGC
    2161  CACTCTATGG CTCACAAACT GGGTGCTATG CACCACGTGC CGCACGGTAT CGCGTGTGCT
    2221  GTCCTGATCG AAGAAGTAAT TAAGTACAAC GCTACTGATT GCCCGACTAA ACAGACCGCC
    2281  TTCCCACAGT ACAAATCTCC TAACGCTAAA CGTAAGTACG CTGAGATCGC CGAATACCTG
    2341  AACCTGAAGG GTACGAGCGA CACTGAGAAA GTTACTGCGC TGATCGAAGC TATCTCTAAA
    2401  CTGAAAATTG ACCTGTCCAT CCCGCAGAAC ATCAGCGCCG CAGGCATCAA CAAAAAGGAC
    2461  TTTTACAACA CGCTGGACAA AATGAGCGAA CTGGCTTTTG ACGACCAGTG CACCACTGCA
    2521  AACCCGCGTT ACCCGCTGAT CTCTGAGCTG AAAGACATTT ATATCAAATC TTTTTAA
```

**AdhE2.13 (76942)**

```
       1  ATGAAAGTGA CGAACCAAAA AGAACTGAAA CAAAAACTGA ATGAACTGCG TGAGGCCCAG
      61  AAAAAGTTCG CTAGCTATAC CCAGGAGCAA GTTGACAAGA TCTTCAAACA GTGTGCCATT
```

121 GCGGCTGCCA AAGAGCGTAT TAACTTGGCC AAACTGGCGG TGGAAGAAAC CGGCATCGGT

181 CTGGTCGAGG ACAAGATTAT CAAGAATCAC TTTGCCGCGG AGTATATCTA CAACAAATAC

```
     241  AAAAACGAAA AGACTTGCGG CATCATTGAT CATGATGACA GCTTGGGTAT CACGAAAGTC
     301  GCAGAACCAA TCGGTATTGT CGCGGCAATT GTCCCGACCA CTAATCCGAC CAGCACCGCG
     361  ATCTTCAAGA GCTTGATCAG CCTGAAAACC CGCAACGCGA TTTTCTTCAG CCCGCACCCG
     421  CGTGCGAAGA AGAGCACCAT CGCAGCGGCG AAGCTGATTT TGGATGCCGC GGTTAAGGCC
     481  GGTGCACCGA AAAACATTAT TGGCTGGATT GACGAGCCGA GCATTGAGCT GTCTCAGGAC
     541  CTGATGAGCG AAGCGGACAT TATTCTGGCC ACCGGTGGTC CGTCTATGGT TAAGGCAGCG
     601  TACTCCAGCG GTAAGCCTGC TATCGGCGTT GGTGCGGGCA ATACGCCGGC TATCATCGAT
     661  GAAAGCGCGG ATATTGATAT GGCAGTTAGC TCCATCATTC TGTCCAAAAC CTACGACAAT
     721  GGTGTTATCT GTGCAAGCGA GCAAAGCATC CTGGTGATGA ATAGCATTTA CGAGAAGGTC
     781  AAAGAAGAGT TTGTTAAGCG TGGTTCCTAT ATTCTGAACC AGAACGAGAT TGCAAAAATC
     841  AAAGAAACGA TGTTTAAGAA TGGCGCGATT AATCGTGATA TTGTTGGTAA GTCCGCATAC
     901  ATTATCGCGA AGATGGCGGG TATCGAAGTG CCGCAAACCA CCAAAATCTT GATCGGCGAA
     961  GTGCAGAGCG TGGAGAAGAG CGAACTGTTC AGCAAAGAGA AACTGAGCCC GGTTCTGGCT
    1021  ATGTACAAAG TGAAGGACTT TGACGAAGCC CTGAAGAAAG CTCAGCGTCT GATTGAGCTG
    1081  GGCGGCAGCG GTCACACGAG CAGCCTGTAT ATCGATAGCC AAAACAATAA GGACAAGGTG
    1141  AAAGAGTTCG GTCTGGCGAT GAAAACCTCT CGCACGTTTA TCAACATGCC GAGCAGCCAA
    1201  GGTGCGAGCG GCGATCTGTA TAACTTCGCA ATCGCGCCTT CTTTTACGCT GGGTTGCGGT
    1261  ACCTGGGGCG GTAATTCGGT GAGCCAGAAT GTCGAGCCGA AGCACCTGCT GAACATCAAA
    1321  TCGGTCGCGG AGCGCCGTGA GAAGAGACCG TGGTTCAAAG TCCCACAGAA AATTTACTTC
    1381  AAATACGGCT GCCTGCGTTT CGCGCTGAAA GAACTGAAAG ACATGAACAA AAAGCGTGCG
    1441  TTCATTGTTA CCGACAAAGA CCTGTTCAAA CTGGGTTACG TGAACAAAAT CACCAAAGTT
    1501  CTGGATGAAA TTGACATCAA GTACTCCATC TTCACTGATA TCAAATCCGA CCCAACGATT
    1561  GATAGCGTGA AAAAGGGCGC TAAAGAAATG CTGAACTTTG AACCGGACAC CATCATCAGC
    1621  ATCGGTGGTG GCTCTCCTAT GGATGCTGCG AAGGTCATGC ACCTGCTGTA CGAATACCCG
    1681  GAAGCGGAAA TCGAAAACCT GGCTATCAAC TTCATGGACA TCCGCAAACG TATCTGCAAC
    1741  TTCCCGAAGC TGGGCACTAA AGCTATTTCC GTTGCCATCC CGACTACCGC GGGCACTGGT
    1801  TCCGAAGCCA CGCCGTTCGC CGTGATCACC AACGATGAAA CCGGTATGAA ATACCCGCTG
    1861  ACCTCTTACG AACTGACCCC GAACATGGCA ATTATCGACA CCGAGCTGAT GCTGAACATG
    1921  CCGCGCAAGC TGACCGCTGC TACCGGCATC GACGCTCTGG TACATGCTAT TGAGGCGTAC
    1981  GTTTCCGTGA TGGCTACCGA TTACACCGAC GAACTGGCCC TGCGTGCGAT CAAAATGATT
    2041  TTCAAGTACC TGCCTCGCGC TTACAAAAAC GGCACGAATG ACATCGAGGC GCGTGAGAAA
    2101  ATGGCCCATG CAAGCAACAT CGCGGGCATG GCCTTCGCCA ACGCGTTCCT GGGCGTGTGC
    2161  CACTCTATGG CTCACAAACT GGGTGCTATG CACCACGTGC CGCACGGTAT CGCGTGTGCT
    2221  GTCCTGATCG AAGAAGTAAT TAAGTACAAC GCTACTGATT GCCCGACTAA ACAGACCGCC
    2281  TTCCCACAGT ACAAATCTCC TAACGCTAAA CGTAAGTACG CTGAGATCGC CGAATACCTG
    2341  AACCTGAAGG GTACGAGCGA CACTGAGAAA GTTACTGCGC TGATCGAAGC TATCTCTAAA
    2401  CTGAAAATTG ACCTGTCCAT CCCGCAGAAC ATCAGCGCCG CAGGCATCAA CAAAAAGGAC
    2461  TTTTACAACA CGCTGGACAA AATGAGCGAA CTGGCTTTTG ACGACCAGTG CACCACTGCA
    2521  AACCCGCGTT ACCCGCTGAT CTCTGAGCTG AAAGACATTT ATATCAAATC TTTTTAA
```

**AdhE2.14 (76943)**

```
       1  ATGAAAGTGA CGAACCAAAA AGAACTGAAA CAAAAAGACA ATGAACTGCG TGAGGCCCAG
      61  AAAAAGTTCG CTACGTATAC CCAGGAGCAA GTTGACAAGA TCTTCAAACA GTGTGCCATT
```

121 GCGGCTGCCA AAGAGCGTAT TAACTTGGCC AAACTGGCGG TGGAAGAAAC CGGCATCGGT

181 CTGGTCGAGG ACAAGATTAT CAAGAATCAC TTTGCCGCGG AGTATATCTA CAACAAATAC

```
     241  AAAAACGAAA AGACTTGCGG CATCATTGAT CATGATGACA GCTTGGGTAT CACGAAAGTC
     301  GCAGAACCAA TCGGTATTGT CGCGGCAATT GTCCCGACCA CTAATCCGAC CAGCACCGCG
     361  ATCTTCAAGA GCTTGATCAG CCTGAAAACC CGCAACGCGA TTTTCTTCAG CCCGCACCCG
     421  CGTGCGAAGA AGAGCACCAT CGCAGCGGCG AAGCTGATTT TGGATGCCGC GGTTAAGGCC
     481  GGTGCACCGA AAAACATTAT TGGCTGGATT GACGAGCCGA GCATTGAGCT GTCTCAGGAC
     541  CTGATGAGCG AAGCGGACAT TATTCTGGCC ACCGGTGGTC CGTCTATGGT TAAGGCAGCG
     601  TACTCCAGCG GTAAGCCTGC TATCGGCGTT GGTGCGGGCA ATACGCCGGC TATCATCGAT
     661  GAAAGCGCGG ATATTGATAT GGCAGTTAGC TCCATCATTC TGTCCAAAAC CTACGACAAT
     721  GGTGTTATCT GTGCAAGCGA GCAAAGCATC CTGGTGATGA ATAGCATTTA CGAGAAGGTC
     781  AAAGAAGAGT TTGTTAAGCG TGGTTCCTAT ATTCTGAACC AGAACGAGAT TGCAAAAATC
     841  AAAGAAACGA TGTTTAAGAA TGGCGCGATT AATGCAGATA TTGTTGGTAA GTCCGCATAC
     901  ATTATCGCGA AGATGGCGGG TATCGAAGTG CCGCAAACCA CCAAAATCTT GATCGGCGAA
     961  GTGCAGAGCG TGGAGAAGAG CGAACTGTTC AGCCATGAGA AACTGAGCCC GGTTCTGGCT
    1021  ATGTACAAAG TGAAGGACTT TGACGAAGCC CTGAAGAAAG CTCAGCGTCT GATTGAGCTG
    1081  GGCGGCAGCG GTCACACGAG CAGCCTGTAT AGCGATAGCC AAAACAATAA GGACAAGGTG
    1141  AAAGAGTTCG GTCTGGAGAT GAAAACCTCT CGCACGTTTA TCAACATGCC GAGCAGCCAA
    1201  GGTGCGAGCG GCGATCTGTA TAACTTCGCA ATCGCGCCTT CTTTTACGCT GGGTTGCGGT
    1261  ACCTGGGGCG GTAATTCGGT GAGCCAGAAT GTCGAGCCGA AGCACCTGCT GAACATCAAA
    1321  TCGGTCGCGG AGCGCCGTGA GAAGAGACCG TGGTTCAAAG TCCCACAGAA AATTTACTTC
    1381  AAATACGGCT GCCTGCGTTT CGCGCTGAAA GAACTGAAAG ACATGAACAA AAAGCGTGCG
    1441  TTCATTGTTA CCGACAAAGA CCTGTTCAAA CTGGGTTACG TGAACAAAAT CACCAAAGTT
    1501  CTGGATGAAA TTGACATCAA GTACTCCATC TTCACTGATA TCAAATCCGA CCCAACGATT
    1561  GATAGCGTGA AAAAGGGCGC TAAAGAAATG CTGAACTTTG AACCGGACAC CATCATCAGC
    1621  ATCGGTGGTG GCTCTCCTAT GGATGCTGCG AAGGTCATGC ACCTGCTGTA CGAATACCCG
    1681  GAAGCGGAAA TCGAAAACCT GGCTATCAAC TTCATGGACA TCCGCAAACG TATCTGCAAC
    1741  TTCCCGAAGC TGGGCACTAA AGCTATTTCC GTTGCCATCC CGACTACCGC GGGCACTGGT
    1801  TCCGAAGCCA CGCCGTTCGC CGTGATCACC AACGATGAAA CCGGTATGAA ATACCCGCTG
    1861  ACCTCTTACG AACTGACCCC GAACATGGCA ATTATCGACA CCGAGCTGAT GCTGAACATG
    1921  CCGCGCAAGC TGACCGCTGC TACCGGCATC GACGCTCTGG TACATGCTAT TGAGGCGTAC
    1981  GTTTCCGTGA TGGCTACCGA TTACACCGAC GAACTGGCCC TGCGTGCGAT CAAAATGATT
    2041  TTCAAGTACC TGCCTCGCGC TTACAAAAAC GGCACGAATG ACATCGAGGC GCGTGAGAAA
    2101  ATGGCCCATG CAAGCAACAT CGCGGGCATG GCCTTCGCCA ACGCGTTCCT GGGCGTGTGC
    2161  CACTCTATGG CTCACAAACT GGGTGCTATG CACCACGTGC CGCACGGTAT CGCGTGTGCT
    2221  GTCCTGATCG AAGAAGTAAT TAAGTACAAC GCTACTGATT GCCCGACTAA ACAGACCGCC
    2281  TTCCCACAGT ACAAATCTCC TAACGCTAAA CGTAAGTACG CTGAGATCGC CGAATACCTG
    2341  AACCTGAAGG GTACGAGCGA CACTGAGAAA GTTACTGCGC TGATCGAAGC TATCTCTAAA
    2401  CTGAAAATTG ACCTGTCCAT CCCGCAGAAC ATCAGCGCCG CAGGCATCAA CAAAAAGGAC
    2461  TTTTACAACA CGCTGGACAA AATGAGCGAA CTGGCTTTTG ACGACCAGTG CACCACTGCA
2521  AACCCGCGTT ACCCGCTGAT CTCTGAGCTG AAAGACATTT ATATCAAATC TTTTTAA
```

**AdhE2.15 (76944)**

```
       1  ATGAAAGTGA CGAACCAAAA AGAACTGAAA CAAAAACTGA ATGAACTGCG TGAGGCCCAG
      61  AAAAAGTTCG CTACGTATAC CCAGGAGCAA GTTGACAAGT TCTTCAAACA GTGTGCCATT
```

121 GCGGCTGCCA AATACCGTAT TAACTTGGCC AAACTGGCGG TGGAAGAAAC CGGCATCGGT

181 CTGGTCGAGG ACAAGATTAT CAAGAATCAC TTTGCCGCGG AGTATATCTA CAACAAAGAG

```
     241  AAAAACGAAA AGACTTGCGG CATCATTGAT CATGATGACA GCTTGGGTAT CACGAAAGTC
     301  GCAGAACCAA TCGGTATTGT CGCGGCAATT GTCCCGACCA CTAATCCGAC CAGCACCGCG
     361  ATCTTCAAGA GCTTGATCAG CCTGAAAACC CGCAACGCGA TTTTCTTCAG CCCGCACCCG
     421  CGTGCGAAGA AGAGCACCAT CGCAGCGGCG AAGCTGATTT TGGATGCCGC GGTTAAGGCC
     481  GGTGCACCGA AAAACATTAT TGGCTGGATT GACGAGCCGA GCATTGAGCT GTCTCAGGAC
     541  CTGATGAGCG AAGCGGACAT TATTCTGGCC ACCGGTGGTC CGTCTATGGT TAAGGCAGCG
     601  TACTCCAGCG GTAAGCCTGC TATCGGCGTT GGTGCGGGCA ATACGCCGGC TATCATCGAT
     661  GAAAGCGCGG ATATTGATAT GGCAGTTAGC TCCATCATTC TGTCCAAAAC CTACGACAAT
     721  GGTGTTATCT GTGCAAGCGA GCAAAGCATC CTGGTGATGA ATAGCATTTA CGAGAAGGTC
     781  AAAGAAGAGT TTGTTAAGCG TGGTTCCTAT ATTCTGAACC AGAACGAGAT TGCAAAAATC
     841  AAAGAAACGA TGTTTAAGAA TGGCGCGATT AATGCAGATA TTGTTGGTAA GTCCGCATAC
     901  ATTATCGCGA AGATGGCGGG TATCGAAGTG CCGCAAACCA CCAAAATCTT GATCGGCGAA
     961  GTGCAGAGCG TGGAGAAGAG CGAACTGTTC AGCCATGAGA AACTGAGCCC GGTTCTGGCT
    1021  ATGTACAAAG TGAAGGACTT TGACGAAGCC CTGAAGAAAG CTCAGCGTCT GATTGAGCTG
    1081  GGCGGCAGCG GTCACACGAG CAGCCTGTAT ATCGATAGCC AAAACAATAA GGACAAGGTG
    1141  AAAGAGTTCG GTCTGGCGAT GAAAACCTCT CGCACGTTTA TCAACATGCC GAGCAGCCAA
    1201  GGTGCGAGCG GCGATCTGTA TAACTTCGCA ATCGCGCCTT CTTTTACGCT GGGTTGCGGT
    1261  ACCTGGGGCG GTAATTCGGT GAGCCAGAAT GTCGAGCCGA AGCACCTGCT GAACATCAAA
    1321  TCGGTCGCGG AGCGCCGTGA GAAGAGACCG TGGTTCAAAG TCCCACAGAA AATTTACTTC
    1381  AAATACGGCT GCCTGCGTTT CGCGCTGAAA GAACTGAAAG ACATGAACAA AAAGCGTGCG
    1441  TTCATTGTTA CCGACAAAGA CCTGTTCAAA CTGGGTTACG TGAACAAAAT CACCAAAGTT
    1501  CTGGATGAAA TTGACATCAA GTACTCCATC TTCACTGATA TCAAATCCGA CCCAACGATT
    1561  GATAGCGTGA AAAAGGGCGC TAAAGAAATG CTGAACTTTG AACCGGACAC CATCATCAGC
    1621  ATCGGTGGTG GCTCTCCTAT GGATGCTGCG AAGGTCATGC ACCTGCTGTA CGAATACCCG
    1681  GAAGCGGAAA TCGAAAACCT GGCTATCAAC TTCATGGACA TCCGCAAACG TATCTGCAAC
    1741  TTCCCGAAGC TGGGCACTAA AGCTATTTCC GTTGCCATCC CGACTACCGC GGGCACTGGT
    1801  TCCGAAGCCA CGCCGTTCGC CGTGATCACC AACGATGAAA CCGGTATGAA ATACCCGCTG
    1861  ACCTCTTACG AACTGACCCC GAACATGGCA ATTATCGACA CCGAGCTGAT GCTGAACATG
    1921  CCGCGCAAGC TGACCGCTGC TACCGGCATC GACGCTCTGG TACATGCTAT TGAGGCGTAC
    1981  GTTTCCGTGA TGGCTACCGA TTACACCGAC GAACTGGCCC TGCGTGCGAT CAAAATGATT
    2041  TTCAAGTACC TGCCTCGCGC TTACAAAAAC GGCACGAATG ACATCGAGGC GCGTGAGAAA
    2101  ATGGCCCATG CAAGCAACAT CGCGGGCATG GCCTTCGCCA ACGCGTTCCT GGGCGTGTGC
    2161  CACTCTATGG CTCACAAACT GGGTGCTATG CACCACGTGC CGCACGGTAT CGCGTGTGCT
    2221  GTCCTGATCG AAGAAGTAAT TAAGTACAAC GCTACTGATT GCCCGACTAA ACAGACCGCC
    2281  TTCCCACAGT ACAAATCTCC TAACGCTAAA CGTAAGTACG CTGAGATCGC CGAATACCTG
    2341  AACCTGAAGG GTACGAGCGA CACTGAGAAA GTTACTGCGC TGATCGAAGC TATCTCTAAA
    2401  CTGAAAATTG ACCTGTCCAT CCCGCAGAAC ATCAGCGCCG CAGGCATCAA CAAAAAGGAC
    2461  TTTTACAACA CGCTGGACAA AATGAGCGAA CTGGCTTTTG ACGACCAGTG CACCACTGCA
    2521  AACCCGCGTT ACCCGCTGAT CTCTGAGCTG AAAGACATTT ATATCAAATC TTTTTAA
```

**AdhE2.16 (76945)**

```
       1  ATGAAAGTGA CGAACCAAAA AGAACTGAAA CAAAAACTGA ATGAACTGCG TGAGGCCCAG
      61  AAAAAGTTCG CTACGTATAC CCAGGAGCAA GTTGAGAAGA TCTTCAAACA GTGTGCCATT
```

121 GCGGCTGCCA AAGAGCGTAT TAACTTGGCC AAACTGGCGG TGGAAGAAAC CGGCATCGGT

181 CTGGTCGAGG ACAAGATTAT CAAGAATCAC TTTGCCGCGG AGTATATCTA CAACAAATAC

```
     241  AAAAACGAAA AGACTTGCGG CATCATTGAT CATGATGACA GCTTGGGTAT CACGAAAGTC
     301  GCAGAACCAA TCGGTATTGT CGCGGCAATT GTCCCGACCA CTAATCCGAC CAGCACCGCG
     361  ATCTTCAAGA GCTTGATCAG CCTGAAAACC CGCAACGCGA TTTTCTTCAG CCCGCACCCG
     421  CGTGCGAAGA AGAGCACCAT CGCAGCGGCG AAGCTGATTT TGGATGCCGC GGTTAAGGCC
     481  GGTGCACCGA AAAACATTAT TGGCTGGATT GACGAGCCGA GCATTGAGCT GTCTCAGGAC
     541  CTGATGAGCG AACCGGACAT TATTCTGGCC ACCGGTGGTC CGTCTATGGT TAAGGCAGCG
     601  TACTCCAGCG GTAAGCCTGC TATCGGCGTT GGTGCGGGCA ATACGCCGGC TATCATCGAT
     661  GAAAGCGCGG ATATTGATAT GGCAGTTAGC TCCATCATTC TGTCCAAAAC CTACGACAAT
     721  GGTGTTATCT GTGCAAGCGA GCAAAGCATC CTGGTGATGA ATAGCATTTA CGAGAAGGTC
     781  AAAGAAGAGT TTGTTAAGCG TGGTTCCTAT ATTCTGAACC AGAACGAGAT TGCAAAAATC
     841  AAAGAAACGA TGTTTAAGAA TGGCGCGATT AATGCAGATA TTGTTGGTAA GTCCGCATAC
     901  ATTATCGCGA AGATGGCGGG TATCGAAGTG CCGCAAACCA CCAAAATCTT GATCGGCGAA
     961  GTGCAGAGCG TGGAGAAGAG CGAACTGTTC AGCAAAGAGA AACTGAGCCC GGTTCTGGCT
    1021  ATGTACAAAG TGAAGGACTT TGACGAAGCC CTGAAGAAAG CTCAGCGTCT GATTGAGCTG
    1081  GGCGGCAGCG GTCACACGAG CAGCCTGTAT ATCGATAGCC AAAACAATAA GGACAAGGTG
    1141  AAAGAGTTCG GTCTGGCGAT GAAAACCTCT CGCACGTTTA TCAACATGCC GAGCAGCCAA
    1201  GGTGCGAGCG GCGATCTGTA TAACTTCGCA ATCGCGCCTT CTTTTACGCT GGGTTGCGGT
    1261  ACCTGGGGCG GTAATTCGGT GAGCCAGAAT GTCGAGCCGA AGCACCTGCT GAACATCAAA
    1321  TCGGTCGCGG AGCGCCGTGA GAAGAGACCG TGGTTCAAAG TCCCACAGAA AATTTACTTC
    1381  AAATACGGCT GCCTGCGTTT CGCGCTGAAA GAACTGAAAG ACATGAACAA AAAGCGTGCG
    1441  TTCATTGTTA CCGACAAAGA CCTGTTCAAA CTGGGTTACG TGAACAAAAT CACCAAAGTT
    1501  CTGGATGAAA TTGACATCAA GTACTCCATC TTCACTGATA TCAAATCCGA CCCAACGATT
    1561  GATAGCGTGA AAAAGGGCGC TAAAGAAATG CTGAACTTTG AACCGGACAC CATCATCAGC
    1621  ATCGGTGGTG GCTCTCCTAT GGATGCTGCG AAGGTCATGC ACCTGCTGTA CGAATACCCG
    1681  GAAGCGGAAA TCGAAAACCT GGCTATCAAC TTCATGGACA TCCGCAAACG TATCTGCAAC
    1741  TTCCCGAAGC TGGGCACTAA AGCTATTTCC GTTGCCATCC CGACTACCGC GGGCACTGGT
    1801  TCCGAAGCCA CGCCGTTCGC CGTGATCACC AACGATGAAA CCGGTATGAA ATACCCGCTG
    1861  ACCTCTTACG AACTGACCCC GAACATGGCA ATTATCGACA CCGAGCTGAT GCTGAACATG
    1921  CCGCGCAAGC TGACCGCTGC TACCGGCATC GACGCTCTGG TACATGCTAT TGAGGCGTAC
    1981  GTTTCCGTGA TGGCTACCGA TTACACCGAC GAACTGGCCC TGCGTGCGAT CAAAATGATT
    2041  TTCAAGTACC TGCCTCGCGC TTACAAAAAC GGCACGAATG ACATCGAGGC GCGTGAGAAA
    2101  ATGGCCCATG CAAGCAACAT CGCGGGCATG GCCTTCGCCA ACGCGTTCCT GGGCGTGTGC
    2161  CACTCTATGG CTCACAAACT GGGTGCTATG CACCACGTGC CGCACGGTAT CGCGTGTGCT
    2221  GTCCTGATCG AAGAAGTAAT TAAGTACAAC GCTACTGATT GCCCGACTAA ACAGACCGCC
    2281  TTCCCACAGT ACAAATCTCC TAACGCTAAA CGTAAGTACG CTGAGATCGC CGAATACCTG
    2341  AACCTGAAGG GTACGAGCGA CACTGAGAAA GTTACTGCGC TGATCGAAGC TATCTCTAAA
    2401  CTGAAAATTG ACCTGTCCAT CCCGCAGAAC ATCAGCGCCG CAGGCATCAA CAAAAAGGAC
    2461  TTTTACAACA CGCTGGACAA AATGAGCGAA CTGGCTTTTG ACGACCAGTG CACCACTGCA
    2521  AACCCGCGTT ACCCGCTGAT CTCTGAGCTG AAAGACATTT ATATCAAATC TTTTTAA
```

**AdhE2.17 (76946)**

```
       1  ATGAAAGTGA CGAACCAAAA AGAACTGAAA CAAAAACTGA ATGAACTGCG TGAGGCCCAG
      61  AAAAAGTTCG CTACGTATAC CCAGGAGCAA GTTGACAAGA TCTTCAAACA GTGTGCCATT
```

121 GCGGCTGCCA AACTGCGTAT TAACTTGGCC AAACTGGCGG TGGAAGAAAC CGGCATCGGT

181 CTGGTCGAGG ACAAGATTAT CAAGAATCAC TTTGCCGCGG AGTATATCTA CAACAAATAC

```
     241  AAAAACGAAA AGACTTGCGG CATCATTGAT CATGATGACA GCTTGGGTAT CACGAAAGTC
     301  GCAGAACCAA TCGGTATTGT CGCGGCAATT GTCCCGACCA CTAATCCGAC CAGCACCGCG
     361  ATCTTCAAGA GCTTGATCAG CCTGAAAACC CGCAACGCGA TTTTCTTCAG CCCGCACCCG
     421  CGTGCGAAGA AGAGCACCAT CGCAGCGGCG AAGCTGATTT TGGATGCCGC GGTTAAGGCC
     481  GGTGCACCGA AAAACATTAT TGGCTGGATT GACGAGCCGA GCATTGAGCT GTCTCAGGAC
     541  CTGATGAGCG AAGCGGACAT TATTCTGGCC ACCGGTGGTC CGTCTATGGT TAAGGCAGCG
     601  TACTCCAGCG GTAAGCCTGC TATCGGCGTT GGTGCGGGCA ATACGCCGGC TATCATCGAT
     661  GAAAGCGCGG ATATTGATAT GGCAGTTAGC TCCATCATTC TGTCCAAAAC CTACGACAAT
     721  GGTGTTATCT GTGCAAGCGA GCAAAGCATC CTGGTGATGA ATAGCATTTA CGAGAAGGTC
     781  AAAGAAGAGT TTGTTAAGCG TGGTTCCTAT ATTCTGAACC AGAACGAGAT TGCAAAAATC
     841  AAAGAAACGA TGTTTAAGAA TGGCGCGATT AATGCAGATA TTGTTGGTAA GTCCGCATAC
     901  ATTATCGCGA AGATGGCGGG TATCGAAGTG CCGCAAACCA CCAAAATCTT GATCGGCGAA
     961  GTGCAGAGCG TGGAGAAGAG CGAACTGTTC AGCGAGGAGC TGCTGAGCCC GGTTCTGGCT
    1021  ATGTACAAAG TGAAGGACTT TGACGAAGCC CTGAAGAAAG CTCAGCGTCT GATTGAGCTG
    1081  GGCGGCAGCG GTCACACGAG CAGCCTGTAT ATCGATAGCC AAAACAATAA GGACAAGGTG
    1141  AAAGAGTTCG GTCTGGCGAT GAAAACCTCT CGCACGTTTA TCAACATGCC GAGCAGCCAA
    1201  GGTGCGAGCG GCGATCTGTA TAACTTCGCA ATCGCGCCTT CTTTTACGCT GGGTTGCGGT
    1261  ACCTGGGGCG GTAATTCGGT GAGCCAGAAT GTCGAGCCGA AGCACCTGCT GAACATCAAA
    1321  TCGGTCGCGG AGCGCCGTGA GAAGAGACCG TGGTTCAAAG TCCCACAGAA AATTTACTTC
    1381  AAATACGGCT GCCTGCGTTT CGCGCTGAAA GAACTGAAAG ACATGAACAA AAAGCGTGCG
    1441  TTCATTGTTA CCGACAAAGA CCTGTTCAAA CTGGGTTACG TGAACAAAAT CACCAAAGTT
    1501  CTGGATGAAA TTGACATCAA GTACTCCATC TTCACTGATA TCAAATCCGA CCCAACGATT
    1561  GATAGCGTGA AAAAGGGCGC TAAAGAAATG CTGAACTTTG AACCGGACAC CATCATCAGC
    1621  ATCGGTGGTG GCTCTCCTAT GGATGCTGCG AAGGTCATGC ACCTGCTGTA CGAATACCCG
    1681  GAAGCGGAAA TCGAAAACCT GGCTATCAAC TTCATGGACA TCCGCAAACG TATCTGCAAC
    1741  TTCCCGAAGC TGGGCACTAA AGCTATTTCC GTTGCCATCC CGACTACCGC GGGCACTGGT
    1801  TCCGAAGCCA CGCCGTTCGC CGTGATCACC AACGATGAAA CCGGTATGAA ATACCCGCTG
    1861  ACCTCTTACG AACTGACCCC GAACATGGCA ATTATCGACA CCGAGCTGAT GCTGAACATG
    1921  CCGCGCAAGC TGACCGCTGC TACCGGCATC GACGCTCTGG TACATGCTAT TGAGGCGTAC
    1981  GTTTCCGTGA TGGCTACCGA TTACACCGAC GAACTGGCCC TGCGTGCGAT CAAAATGATT
    2041  TTCAAGTACC TGCCTCGCGC TTACAAAAAC GGCACGAATG ACATCGAGGC GCGTGAGAAA
    2101  ATGGCCCATG CAAGCAACAT CGCGGGCATG GCCTTCGCCA ACGCGTTCCT GGGCGTGTGC
    2161  CACTCTATGG CTCACAAACT GGGTGCTATG CACCACGTGC CGCACGGTAT CGCGTGTGCT
    2221  GTCCTGATCG AAGAAGTAAT TAAGTACAAC GCTACTGATT GCCCGACTAA ACAGACCGCC
    2281  TTCCCACAGT ACAAATCTCC TAACGCTAAA CGTAAGTACG CTGAGATCGC CGAATACCTG
    2341  AACCTGAAGG GTACGAGCGA CACTGAGAAA GTTACTGCGC TGATCGAAGC TATCTCTAAA
    2401  CTGAAAATTG ACCTGTCCAT CCCGCAGAAC ATCAGCGCCG CAGGCATCAA CAAAAAGGAC
    2461  TTTTACAACA CGCTGGACAA AATGAGCGAA CTGGCTTTTG ACGACCAGTG CACCACTGCA
    2521  AACCCGCGTT ACCCGCTGAT CTCTGAGCTG AAAGACATTT ATATCAAATC TTTTTAA
```

**AdhE2.18 (76947)**

```
       1  ATGAAAGTGA CGAACCAAAA AGAACTGAAA CAAAAACTGA ATGAACTGCG TGAGGCCCAG
      61  AAAAAGTTCG CTACGTATAC CCAGGAGCAA GTTGACAAGA TCTTCAAACA GTGTGCCATT
```

121 GCGGCTGCCA AAGAGCGTAT TAACTTGGCC AAACTGGCGG TGGAAGAAAC CGGCATCGGT

181 CTGGTCGAGG ACAAGATTAT CAAGAATCAC TTTGCCGCGG AGTATATCTA CAACAAATAC

```
     241  AAAAACCACA AGACTTGCGG CATCATTGAT CATGATGACA GCTTGGGTAT CACGAAAGTC
     301  GCAGAACCAA TCGGTATTGT CGCGGCAATT GTCCCGACCA CTAATCCGAC CAGCACCGCG
     361  ATCTTCAAGA GCTTGATCAG CCTGGCGACC CGCAACGCGA TTTTCTTCAG CCCGCACCCG
     421  CGTGCGAAGA AGAGCACCAT CGCAGCGGCG AAGCTGATTT TGGATGCCGC GGTTAAGGCC
     481  GGTGCACCGA AAAACATTAT TGGCTGGATT GACGAGCCGA GCATTGAGCT GTCTCAGGAC
     541  CTGATGAGCG AAGCGGACAT TATTCTGGCC ACCGGTGGTC CGTCTATGGT TAAGGCAGCG
     601  TACTCCAGCG GTAAGCCTGC TATCGGCGTT GGTGCGGGCA ATACGCCGGC TATCATCGAT
     661  GAAAGCGCGG ATATTGATAT GGCAGTTAGC TCCATCATTC TGTCCAAAAC CTACGACAAT
     721  GGTGTTATCT GTGCAAGCGA GCAAAGCATC CTGGTGATGA ATAGCATTTA CGAGAAGGTC
     781  AAAGAAGAGT TTGTTAAGCG TGGTTCCTAT ATTCTGAACC AGAACGAGAT TGCAAAAATC
     841  AAAGAAACGA TGTTTAAGAA TGGCGCGATT AATGCAGATA TTGTTGGTAA GTCCGCATAC
     901  ATTATCGCGA AGATGGCGGG TATCGAAGTG CCGCAAACCA CCAAAATCTT GATCGGCGAA
     961  GTGCAGAGCG TGGAGAAGAG CGAACTGTTC AGCCATGAGA AACTGAGCCC GGTTCTGGCT
    1021  ATGTACAAAG TGAAGGACTT TGACGAAGCC CTGGACAAAG CTCAGCGTCT GATTGAGCTG
    1081  GGCGGCAGCG GTCACACGAG CAGCCTGTAT ATCGATAGCC AAAACAATAA GGACAAGGTG
    1141  AAAGAGTTCG GTCTGGCGAT GAAAACCTCT CGCACGTTTA TCAACATGCC GAGCAGCCAA
    1201  GGTGCGAGCG GCGATCTGTA TAACTTCGCA ATCGCGCCTT CTTTTACGCT GGGTTGCGGT
    1261  ACCTGGGGCG GTAATTCGGT GAGCCAGAAT GTCGAGCCGA AGCACCTGCT GAACATCAAA
    1321  TCGGTCGCGG AGCGCCGTGA GAAGAGACCG TGGTTCAAAG TCCCACAGAA AATTTACTTC
    1381  AAATACGGCT GCCTGCGTTT CGCGCTGAAA GAACTGAAAG ACATGAACAA AAAGCGTGCG
    1441  TTCATTGTTA CCGACAAAGA CCTGTTCAAA CTGGGTTACG TGAACAAAAT CACCAAAGTT
    1501  CTGGATGAAA TTGACATCAA GTACTCCATC TTCACTGATA TCAAATCCGA CCCAACGATT
    1561  GATAGCGTGA AAAAGGGCGC TAAAGAAATG CTGAACTTTG AACCGGACAC CATCATCAGC
    1621  ATCGGTGGTG GCTCTCCTAT GGATGCTGCG AAGGTCATGC ACCTGCTGTA CGAATACCCG
    1681  GAAGCGGAAA TCGAAAACCT GGCTATCAAC TTCATGGACA TCCGCAAACG TATCTGCAAC
    1741  TTCCCGAAGC TGGGCACTAA AGCTATTTCC GTTGCCATCC CGACTACCGC GGGCACTGGT
    1801  TCCGAAGCCA CGCCGTTCGC CGTGATCACC AACGATGAAA CCGGTATGAA ATACCCGCTG
    1861  ACCTCTTACG AACTGACCCC GAACATGGCA ATTATCGACA CCGAGCTGAT GCTGAACATG
    1921  CCGCGCAAGC TGACCGCTGC TACCGGCATC GACGCTCTGG TACATGCTAT TGAGGCGTAC
    1981  GTTTCCGTGA TGGCTACCGA TTACACCGAC GAACTGGCCC TGCGTGCGAT CAAAATGATT
    2041  TTCAAGTACC TGCCTCGCGC TTACAAAAAC GGCACGAATG ACATCGAGGC GCGTGAGAAA
    2101  ATGGCCCATG CAAGCAACAT CGCGGGCATG GCCTTCGCCA ACGCGTTCCT GGGCGTGTGC
    2161  CACTCTATGG CTCACAAACT GGGTGCTATG CACCACGTGC CGCACGGTAT CGCGTGTGCT
    2221  GTCCTGATCG AAGAAGTAAT TAAGTACAAC GCTACTGATT GCCCGACTAA ACAGACCGCC
    2281  TTCCCACAGT ACAAATCTCC TAACGCTAAA CGTAAGTACG CTGAGATCGC CGAATACCTG
    2341  AACCTGAAGG GTACGAGCGA CACTGAGAAA GTTACTGCGC TGATCGAAGC TATCTCTAAA
    2401  CTGAAAATTG ACCTGTCCAT CCCGCAGAAC ATCAGCGCCG CAGGCATCAA CAAAAAGGAC
    2461  TTTTACAACA CGCTGGACAA AATGAGCGAA CTGGCTTTTG ACGACCAGTG CACCACTGCA
    2521  AACCCGCGTT ACCCGCTGAT CTCTGAGCTG AAAGACATTT ATATCAAATC TTTTTAA
```

**AdhE2.19 (76948)**

```
       1  ATGAAAGTGA CGAACCAAAA AGAACTGAAA CAAAAACTGA ATGAACTGCG TGAGGCCCAG
      61  AAAAAGTTCG CTACGTATAC CCAGGAGCAA GTTGACAAGA TCTTCAAACA GTGTGCCATT
```

121 GCGGCTGCCA AAGAGCGTAT TAACTTGGCC AAACTGGCGG TGGAAGAAAC CGGCATCGGT

181 CTGGTCGAGG ACAAGATTAT CAAGAATCAC TTTGCCGCGG AGTATATCTA CAACAAAGAG

```
     241  AAAAACGAAA AGACTTGCGG CATCATTGAT CATGATGACA GCTTGGGTAT CACGAAAGTC
     301  GCAGAACCAA TCGGTATTGT CGCGGCAATT GTCCCGACCA CTAATCCGAC CAGCACCGCG
     361  ATCTTCAAGA GCTTGATCAG CCTGAAAACC CGCAACGCGA TTTTCTTCAG CCCGCACCCG
     421  CGTGCGAAGA AGAGCACCAT CGCAGCGGCG AAGCTGATTT TGGATGCCGC GGTTAAGGCC
     481  GGTGCACCGA AAAACATTAT TGGCTGGATT GACGAGCCGA GCATTGAGCT GTCTCAGGAC
     541  CTGATGAGCG AAGCGGACAT TATTCTGGCC ACCGGTGGTC CGTCTATGGT TAAGGCAGCG
     601  TACTCCAGCG GTAAGCCTGC TATCGGCGTT GGTGCGGGCA ATACGCCGGC TATCATCGAT
     661  GAAAGCGCGG ATATTGATAT GGCAGTTAGC TCCATCATTC TGTCCAAAAC CTACGACAAT
     721  GGTGTTATCT GTGCAAGCGA GCAAAGCATC CTGGTGATGA ATAGCATTTA CGAGAAGGTC
     781  AAAGAAGAGT TTGTTAAGCG TGGTTCCTAT ATTCTGAACC AGAACGAGAT TGCAAAAATC
     841  AAAGAAACGA TGTTTAAGAA TGGCGCGATT AATGCAGATA TTGTTGGTAA GTCCGCATAC
     901  ATTATCGCGA AGATGGCGGG TATCGAAGTG CCGCAAACCA CCAAAATCTT GATCGGCGAA
     961  GTGCAGAGCG TGGAGAAGAG CGAACTGTTC AGCCATGAGA AACTGAGCCC GGTTCTGGCT
    1021  ATGTACAAAG TGAAGGACTT TGACGAAGCC CTGAAGAAAG CTCAGCGTCT GATTGAGCTG
    1081  GGCGGCAGCG GTCACACGAG CAGCCTGTAT ATCGATAGCC AAAACAATAA GGACAAGGTG
    1141  AAAGAGTTCG GTCTGGCGAT GAAAACCTCT CGCACGTTTA TCAACATGCC GAGCAGCCAA
    1201  GGTGCGAGCG GCGATCTGTA TAACTTCGCA ATCGCGCCTT CTTTTACGCT GGGTTGCGGT
    1261  ACCTGGGGCG GTAATTCGGT GAGCCAGAAT GTCGAGCCGA AGCACCTGCT GAACATCGTT
    1321  TCGGTCGCGC GTCGCCGTGA GAAGAGACCG TGGTTCAAAG TCCCACAGAA AATTTACTTC
    1381  AAATACGGCT GCCTGCGTTT CGCGCTGAAA GAACTGAAAG ACATGAACAA AAAGCGTGCG
    1441  TTCATTGTTA CCGACAAAGA CCTGTTCAAA CTGGGTTACG TGAACAAAAT CACCAAAGTT
    1501  CTGGATGAAA TTGACATCAA GTACTCCATC TTCACTGATA TCAAATCCGA CCCAACGATT
    1561  GATAGCGTGA AAAAGGGCGC TAAAGAAATG CTGAACTTTG AACCGGACAC CATCATCAGC
    1621  ATCGGTGGTG GCTCTCCTAT GGATGCTGCG AAGGTCATGC ACCTGCTGTA CGAATACCCG
    1681  GAAGCGGAAA TCGAAAACCT GGCTATCAAC TTCATGGACA TCCGCAAACG TATCTGCAAC
    1741  TTCCCGAAGC TGGGCACTAA AGCTATTTCC GTTGCCATCC CGACTACCGC GGGCACTGGT
    1801  TCCGAAGCCA CGCCGTTCGC CGTGATCACC AACGATGAAA CCGGTATGAA ATACCCGCTG
    1861  ACCTCTTACG AACTGACCCC GAACATGGCA ATTATCGACA CCGAGCTGAT GCTGAACATG
    1921  CCGCGCAAGC TGACCGCTGC TACCGGCATC GACGCTCTGG TACATGCTAT TGAGGCGTAC
    1981  GTTTCCGTGA TGGCTACCGA TTACACCGAC GAACTGGCCC TGCGTGCGAT CAAAATGATT
    2041  TTCAAGTACC TGCCTCGCGC TTACAAAAAC GGCACGAATG ACATCGAGGC GCGTGAGAAA
    2101  ATGGCCCATG CAAGCAACAT CGCGGGCATG GCCTTCGCCA ACGCGTTCCT GGGCGTGTGC
    2161  CACTCTATGG CTCACAAACT GGGTGCTATG CACCACGTGC CGCACGGTAT CGCGTGTGCT
    2221  GTCCTGATCG AAGAAGTAAT TAAGTACAAC GCTACTGATT GCCCGACTAA ACAGACCGCC
    2281  TTCCCACAGT ACAAATCTCC TAACGCTAAA CGTAAGTACG CTGAGATCGC CGAATACCTG
    2341  AACCTGAAGG GTACGAGCGA CACTGAGAAA GTTACTGCGC TGATCGAAGC TATCTCTAAA
    2401  CTGAAAATTG ACCTGTCCAT CCCGCAGAAC ATCAGCGCCG CAGGCATCAA CAAAAAGGAC
    2461  TTTTACAACA CGCTGGACAA AATGAGCGAA CTGGCTTTTG ACGACCAGTG CACCACTGCA
    2521  AACCCGCGTT ACCCGCTGAT CTCTGAGCTG AAAGACATTT ATATCAAATC TTTTTAA
```

**AdhE2.20 (76949)**

```
       1  ATGAAAGTGA CGAACCAAAA AGAACTGAAA CAAAAACTGA ATGAACTGCG TGAGGCCCAG
      61  AAAAAGTTCG CTACGTATAC CCAGGAGCAA GTTGACAAGA TCTTCAAACA GTGTGCCATT
```

121 GCGGCTGCCA AAGAGCGTAT TAACTTGGCC AAACTGGCGG TGGAAGAAAC CGGCATCGGT

181 CTGGTCGAGG ACAAGATTAT CAAGAATCAC TTTGCCGCGG AGTATATCTA CAACAAATAC

```
     241  AAAAACGAAA AGACTTGCGG CATCATTGAT CATGATGACA GCTTGGGTAT CACGAAAGTC
     301  GCAGAACCAA TCGGTATTGT CGCGGCAATT GTCCCGACCA CTAATCCGAC CAGCACCGCG
     361  ATCTTCAAGA GCTTGATCAG CCTGAAAACC CGCAACGCGA TTTTCTTCAG CCCGCACCCG
     421  CGTGCGAAGA AGAGCACCAT CGCAGCGGCG AAGCTGATTT TGGATGCCGC GGTTAAGGCC
     481  GGTGCACCGA AAAACATTAT TGGCTGGATT GACGAGCCGA GCATTGAGCT GTCTCAGGAC
     541  CTGATGAGCG AAGCGGACAT TATTCTGGCC ACCGGTGGTC CGTCTATGGT TAAGGCAGCG
     601  TACTCCAGCG GTAAGCCTGC TATCGGCGCG GGTGCGGGCA ATACGCCGGC TATCATCGAT
     661  GAAAGCGCGG ATATTGATAT GGCAGTTAGC TCCATCATTC TGTCCAAAAC CTACGACAAT
     721  GGTGTTATCT GTGCAAGCGA GCAAAGCATC CTGGTGATGA ATAGCATTTA CGAGAAGGTC
     781  AAAGAAGAGT TTGTTAAGCG TGGTTCCTAT ATTCTGAACC AGAACGAGAT TGCAAAAATC
     841  AAAGAAACGA TGTTTAAGAA TGGCGCGATT AATGCAGATA TTGTTGGTAA GTCCGCATAC
     901  ATTATCGCGA AGATGGCGGG TATCGAAGTG CCGCAAACCA CCAAAATCTT GATCGGCGAA
     961  GTGCAGGACG TGGAGAAGAG CGAACTGTTC AGCGTTGAGA AACTGAGCCC GGTTCTGGCT
    1021  ATGTACAAAG TGAAGGACTT TGACGAAGCC CTGAAGAAAG CTCAGCGTCT GATTGAGCTG
    1081  GGCGGCAGCG GTCACACGAG CAGCCTGTAT ATCGATAGCC AAAACAATAA GGACAAGGTG
    1141  AAAGAGTTCG GTCTGGCGAT GAAAACCTCT CGCACGTTTA TCAACATGCC GAGCAGCCAA
    1201  GGTGCGAGCG GCGATCTGTA TAACTTCGCA ATCGCGCCTT CTTTTACGCT GGGTTGCGGT
    1261  ACCTGGGGCG GTAATTCGGT GAGCCAGAAT GTCGAGCCGA AGCACCTGCT GAACATCAAA
    1321  TCGGTCGCGG AGCGCCGTGA GAAGAGACCG TGGTTCAAAG TCCCACAGAA AATTTACTTC
    1381  AAATACGGCT GCCTGCGTTT CGCGCTGAAA GAACTGAAAG ACATGAACAA AAAGCGTGCG
    1441  TTCATTGTTA CCGACAAAGA CCTGTTCAAA CTGGGTTACG TGAACAAAAT CACCAAAGTT
    1501  CTGGATGAAA TTGACATCAA GTACTCCATC TTCACTGATA TCAAATCCGA CCCAACGATT
    1561  GATAGCGTGA AAAAGGGCGC TAAAGAAATG CTGAACTTTG AACCGGACAC CATCATCAGC
    1621  ATCGGTGGTG GCTCTCCTAT GGATGCTGCG AAGGTCATGC ACCTGCTGTA CGAATACCCG
    1681  GAAGCGGAAA TCGAAAACCT GGCTATCAAC TTCATGGACA TCCGCAAACG TATCTGCAAC
    1741  TTCCCGAAGC TGGGCACTAA AGCTATTTCC GTTGCCATCC CGACTACCGC GGGCACTGGT
    1801  TCCGAAGCCA CGCCGTTCGC CGTGATCACC AACGATGAAA CCGGTATGAA ATACCCGCTG
    1861  ACCTCTTACG AACTGACCCC GAACATGGCA ATTATCGACA CCGAGCTGAT GCTGAACATG
    1921  CCGCGCAAGC TGACCGCTGC TACCGGCATC GACGCTCTGG TACATGCTAT TGAGGCGTAC
    1981  GTTTCCGTGA TGGCTACCGA TTACACCGAC GAACTGGCCC TGCGTGCGAT CAAAATGATT
    2041  TTCAAGTACC TGCCTCGCGC TTACAAAAAC GGCACGAATG ACATCGAGGC GCGTGAGAAA
    2101  ATGGCCCATG CAAGCAACAT CGCGGGCATG GCCTTCGCCA ACGCGTTCCT GGGCGTGTGC
    2161  CACTCTATGG CTCACAAACT GGGTGCTATG CACCACGTGC CGCACGGTAT CGCGTGTGCT
    2221  GTCCTGATCG AAGAAGTAAT TAAGTACAAC GCTACTGATT GCCCGACTAA ACAGACCGCC
    2281  TTCCCACAGT ACAAATCTCC TAACGCTAAA CGTAAGTACG CTGAGATCGC CGAATACCTG
    2341  AACCTGAAGG GTACGAGCGA CACTGAGAAA GTTACTGCGC TGATCGAAGC TATCTCTAAA
    2401  CTGAAAATTG ACCTGTCCAT CCCGCAGAAC ATCAGCGCCG CAGGCATCAA CAAAAAGGAC
    2461  TTTTACAACA CGCTGGACAA AATGAGCGAA CTGGCTTTTG ACGACCAGTG CACCACTGCA
    2521  AACCCGCGTT ACCCGCTGAT CTCTGAGCTG AAAGACATTT ATATCAAATC TTTTTAA
```

**AdhE2.21 (76950)**

```
       1  ATGAAAGTGA CGAACCAAAA AGAACTGAAA CAAAAACTGA ATGAACTGCG TGAGGCCCAG
      61  AAAAAGTTCG CTACGTATAC CCAGGACCAA GTTGACAAGA TCTTCAAACA GTGTGCCATT
```

121 GCGGCTGCCA AAGAGCGTAT TAACTTGGCC AAACTGGCGG TGGAAGAAAC CGGCATCGGT

181 CTGGTCGAGG ACAAGATTAT CAAGAATCAC TTTGCCGCGG AGTATATCTA CAACAAATAC

```
     241  AAAAACGAAA AGACTTGCGG CATCATTGAT CATGATGACA GCTTGGGTAT CACGAAAGTC
     301  GCAGAACCAA TCGGTATTGT CGCGGCAATT GTCCCGACCA CTAATCCGAC CAGCACCGCG
     361  ATCTTCAAGA GCTTGATCAG CCTGAAAACC CGCAACGCGA TTTTCTTCAG CCCGCACCCG
     421  CGTGCGAAGA AGAGCACCAT CGCAGCGGCG AAGCTGATTT TGGATGCCGC GGTTAAGGCC
     481  GGTGCACCGA AAAACATTAT TGGCTGGATT GACGAGCCGA GCATTGAGCT GTCTCAGGAC
     541  CTGATGAAAG AAGCGGACAT TATTCTGGCC ACCGGTGGTC CGTCTATGGT TAAGGCAGCG
     601  TACTCCAGCG GTAAGCCTGC TATCGGCGTT GGTGCGGGCA ATACGCCGGC TATCATCGAT
     661  GAAAGCGCGG ATATTGATAT GGCAGTTAGC TCCATCATTC TGTCCAAAAC CTACGACAAT
     721  GGTGTTATCT GTGCAAGCGA GCAAAGCATC CTGGTGATGA ATAGCATTTA CGAGAAGGTC
     781  AAAAAAGAGT TTGTTAAGCG TGGTTCCTAT ATTCTGAACC AGAACGAGAT TGCAAAAATC
     841  AAAGAAACGA TGTTTAAGAA TGGCGCGATT AATGCAGATA TTGTTGGTAA GTCCGCATAC
     901  ATTATCGCGA AGATGGCGGG TATCGAAGTG CCGCAAACCA CCAAAATCTT GATCGGCGAA
     961  GTGCAGAGCG TGGAGAAGAG CGAACTGTTC AGCCATGAGA AACTGAGCCC GGTTCTGGCT
    1021  ATGTACAAAG TGAAGGACTT TGACGAAGCC CTGAAGAAAG CTCAGCGTCT GATTGAGCTG
    1081  GGCGGCAGCG GTCACACGAG CAGCCTGTAT ATCGATAGCC AAAACAATAA GGACAAGGTG
    1141  AAAGAGTTCG GTCTGGCGAT GAAAACCTCT CGCACGTTTA TCAACATGCC GAGCAGCCAA
    1201  GGTGCGAGCG GCGATCTGTA TAACTTCGCA ATCGCGCCTT CTTTTACGCT GGGTTGCGGT
    1261  ACCTGGGGCG GTAATTCGGT GAGCCAGAAT GTCGAGCCGA AGCACCTGCT GAACATCAAA
    1321  TCGGTCGCGG AGCGCCGTGA GAAGAGACCG TGGTTCAAAG TCCCACAGAA AATTTACTTC
    1381  AAATACGGCT GCCTGCGTTT CGCGCTGAAA GAACTGAAAG ACATGAACAA AAAGCGTGCG
    1441  TTCATTGTTA CCGACAAAGA CCTGTTCAAA CTGGGTTACG TGAACAAAAT CACCAAAGTT
    1501  CTGGATGAAA TTGACATCAA GTACTCCATC TTCACTGATA TCAAATCCGA CCCAACGATT
    1561  GATAGCGTGA AAAAGGGCGC TAAAGAAATG CTGAACTTTG AACCGGACAC CATCATCAGC
    1621  ATCGGTGGTG GCTCTCCTAT GGATGCTGCG AAGGTCATGC ACCTGCTGTA CGAATACCCG
    1681  GAAGCGGAAA TCGAAAACCT GGCTATCAAC TTCATGGACA TCCGCAAACG TATCTGCAAC
    1741  TTCCCGAAGC TGGGCACTAA AGCTATTTCC GTTGCCATCC CGACTACCGC GGGCACTGGT
    1801  TCCGAAGCCA CGCCGTTCGC CGTGATCACC AACGATGAAA CCGGTATGAA ATACCCGCTG
    1861  ACCTCTTACG AACTGACCCC GAACATGGCA ATTATCGACA CCGAGCTGAT GCTGAACATG
    1921  CCGCGCAAGC TGACCGCTGC TACCGGCATC GACGCTCTGG TACATGCTAT TGAGGCGTAC
    1981  GTTTCCGTGA TGGCTACCGA TTACACCGAC GAACTGGCCC TGCGTGCGAT CAAAATGATT
    2041  TTCAAGTACC TGCCTCGCGC TTACAAAAAC GGCACGAATG ACATCGAGGC GCGTGAGAAA
    2101  ATGGCCCATG CAAGCAACAT CGCGGGCATG GCCTTCGCCA ACGCGTTCCT GGGCGTGTGC
    2161  CACTCTATGG CTCACAAACT GGGTGCTATG CACCACGTGC CGCACGGTAT CGCGTGTGCT
    2221  GTCCTGATCG AAGAAGTAAT TAAGTACAAC GCTACTGATT GCCCGACTAA ACAGACCGCC
    2281  TTCCCACAGT ACAAATCTCC TAACGCTAAA CGTAAGTACG CTGAGATCGC CGAATACCTG
    2341  AACCTGAAGG GTACGAGCGA CACTGAGAAA GTTACTGCGC TGATCGAAGC TATCTCTAAA
    2401  CTGAAAATTG ACCTGTCCAT CCCGCAGAAC ATCAGCGCCG CAGGCATCAA CAAAAAGGAC
    2461  TTTTACAACA CGCTGGACAA AATGAGCGAA CTGGCTTTTG ACGACCAGTG CACCACTGCA
2521  AACCCGCGTT ACCCGCTGAT CTCTGAGCTG AAAGACATTT ATATCAAATC TTTTTAA
```

**AdhE2.22 (76951)**

```
       1  ATGAAAGTGA CGAACCAAAA AGAACTGAAA CAAAAACTGA ATGAACTGCG TGAGGCCCAG
      61  AAAAAGTTCG CTACGTATAC CCAGGAGCAA GTTGACAAGT TCTTCAAACA GTGTGCCATT
```

121 GCGGCTGCCA AAGAGCGTAT TAACTTGGCC CGTCTGGCGG TGGAAGAAAC CGGCATCGGT

181 CTGGTCGAGG ACAAGATTAT CAAGAATCAC TTTGCCGCGG AGTATATCTA CAACAAATAC

```
     241  AAAAACGAAA AGACTTGCGG CATCATTGAT CATGATGACA GCTTGGGTAT CACGAAAGTC
     301  GCAGAACCAA TCGGTATTGT CGCGGCAATT GTCCCGACCA CTAATCCGAC CAGCACCGCG
     361  ATCTTCAAGA GCTTGATCAG CCTGAAAACC CGCAACGCGA TTTTCTTCAG CCCGCACCCG
     421  CGTGCGAAGA AGAGCACCAT CGCAGCGGCG AAGCTGATTT TGGATGCCGC GGTTAAGGCC
     481  GGTGCACCGA AAAACATTAT TGGCTGGATT GACGAGCCGA GCATTGAGCT GTCTCAGGAC
     541  CTGATGAGCG AAGCGGACAT TATTCTGGCC ACCGGTGGTC CGTCTATGGT TAAGGCAGCG
     601  TACTCCAGCG GTAAGCCTGC TATCGGCGTT GGTGCGGGCA ATACGCCGGC TATCATCGAT
     661  GAAAGCGCGG ATATTGATAT GGCAGTTAGC TCCATCATTC TGTCCAAAAC CTACGACAAT
     721  GGTGTTATCT GTGCAAGCGA GCAAAGCATC CTGGTGATGA ATAGCATTTA CGAGAAGGTC
     781  AAAGAAGAGT TTGTTAAGCG TGGTTCCTAT ATTCTGAACC AGAACGAGAT TGCAAAAATC
     841  AAAGAAACGA TGTTTAAGAA TGGCGCGATT AATGCAGATA TTGTTGGTAA GTCCGCATAC
     901  ATTATCGCGA AGATGGCGGG TATCGAAGTG CCGCAAACCA CCAAAATCTT GATCGGCGAA
     961  GTGCAGAGCG TGGAGGCGAG CGAACTGTTC AGCCATGAGA AACTGAGCCC GGTTCTGGCT
    1021  ATGTACAAAG TGAAGGACTT TGACGAAGCC CTGAAGAAAG CTCAGCGTCT GATTGAGCTG
    1081  GGCGGCAGCG GTCACACGAG CAGCCTGTAT ATCGATAGCC AAAACAATAA GGACAAGGTG
    1141  AAAGAGTTCG GTCTGGCGAT GAAAACCTCT CGCACGTTTA TCAACATGCC GAGCAGCCAA
    1201  GGTGCGAGCG GCGATCTGTA TAACTTCGCA ATCGCGCCTT CTTTTACGCT GGGTTGCGGT
    1261  ACCTGGGGCG GTAATTCGGT GAGCCAGAAT GTCGAGCCGA AGCACCTGCT GAACATCAAA
    1321  TCGGTCGCGG AGCGCCGTGA GAAGAGACCG TGGTTCAAAG TCCCACAGAA AATTTACTTC
    1381  AAATACGGCT GCCTGCGTTT CGCGCTGAAA GAACTGAAAG ACATGAACAA AAAGCGTGCG
    1441  TTCATTGTTA CCGACAAAGA CCTGTTCAAA CTGGGTTACG TGAACAAAAT CACCAAAGTT
    1501  CTGGATGAAA TTGACATCAA GTACTCCATC TTCACTGATA TCAAATCCGA CCCAACGATT
    1561  GATAGCGTGA AAAAGGGCGC TAAAGAAATG CTGAACTTTG AACCGGACAC CATCATCAGC
    1621  ATCGGTGGTG GCTCTCCTAT GGATGCTGCG AAGGTCATGC ACCTGCTGTA CGAATACCCG
    1681  GAAGCGGAAA TCGAAAACCT GGCTATCAAC TTCATGGACA TCCGCAAACG TATCTGCAAC
    1741  TTCCCGAAGC TGGGCACTAA AGCTATTTCC GTTGCCATCC CGACTACCGC GGGCACTGGT
    1801  TCCGAAGCCA CGCCGTTCGC CGTGATCACC AACGATGAAA CCGGTATGAA ATACCCGCTG
    1861  ACCTCTTACG AACTGACCCC GAACATGGCA ATTATCGACA CCGAGCTGAT GCTGAACATG
    1921  CCGCGCAAGC TGACCGCTGC TACCGGCATC GACGCTCTGG TACATGCTAT TGAGGCGTAC
    1981  GTTTCCGTGA TGGCTACCGA TTACACCGAC GAACTGGCCC TGCGTGCGAT CAAAATGATT
    2041  TTCAAGTACC TGCCTCGCGC TTACAAAAAC GGCACGAATG ACATCGAGGC GCGTGAGAAA
    2101  ATGGCCCATG CAAGCAACAT CGCGGGCATG GCCTTCGCCA ACGCGTTCCT GGGCGTGTGC
    2161  CACTCTATGG CTCACAAACT GGGTGCTATG CACCACGTGC CGCACGGTAT CGCGTGTGCT
    2221  GTCCTGATCG AAGAAGTAAT TAAGTACAAC GCTACTGATT GCCCGACTAA ACAGACCGCC
    2281  TTCCCACAGT ACAAATCTCC TAACGCTAAA CGTAAGTACG CTGAGATCGC CGAATACCTG
    2341  AACCTGAAGG GTACGAGCGA CACTGAGAAA GTTACTGCGC TGATCGAAGC TATCTCTAAA
    2401  CTGAAAATTG ACCTGTCCAT CCCGCAGAAC ATCAGCGCCG CAGGCATCAA CAAAAAGGAC
    2461  TTTTACAACA CGCTGGACAA AATGAGCGAA CTGGCTTTTG ACGACCAGTG CACCACTGCA
    2521  AACCCGCGTT ACCCGCTGAT CTCTGAGCTG AAAGACATTT ATATCAAATC TTTTTAA
```

**AdhE2.23 (76952)**

```
       1  ATGAAAGTGA CGAACCAAAA AGAACTGAAA CAAAAACTGA ATGAACTGCG TGAGGCCCAG
      61  AAAAAGTTCG CTACGTGCAC CCAGGAGCAA GTTGACAAGA TCTTCAAACA GTGTGCCATT
```

121 GCGGCTGCCA AAGAGCGTAT TAACTTGGCC AAACTGGCGG TGGAAGAAAC CGGCATCGGT

181 CTGGTCGAGG ACAAGATTAT CAAGAATCAC TTTGCCGCGG AGTATATCTA CAACAAATAC

```
     241  AAAAACGAAA AGACTTGCGG CATCATTGAT CATGATGACA GCTTGGGTAT CACGAAAGTC
     301  GCAGAACCAA TCGGTATTGT CGCGGCAATT GTCCCGACCA CTAATCCGAC CAGCACCGCG
     361  ATCTTCAAGA GCTTGATCAG CCTGAAAACC CGCAACGCGA TTTTCTTCAG CCCGCACCCG
     421  CGTGCGAAGA AGAGCACCAT CGCAGCGGCG AAGCTGATTT TGGATGCCGC GGTTAAGGCC
     481  GGTGCACCGA AAAACATTAT TGGCTGGATT GACGAGCCGA GCATTGAGCT GTCTCAGGAC
     541  CTGATGAGCG AAGCGGACAT TATTCTGGCC ACCGGTGGTC CGTCTATGGT TAAGGCAGCG
     601  TACTCCAGCG GTAAGCCTGC TATCGGCGTT GGTGCGGGCA ATACGCCGGC TATCATCGAT
     661  GAAAGCGCGG ATATTGATAT GGCAGTTAGC TCCATCATTC TGTCCGCGAC CTACGACAAT
     721  GGTGTTATCT GTGCAAGCGA GCAAAGCATC CTGGTGATGA ATAGCATTTA CGAGAAGGTC
     781  AAAGAAGAGT TTGTTAAGCG TGGTTCCTAT ATTCTGAACC AGAACGAGAT TGCAAAAATC
     841  AAAGAAACGA TGTTTAAGAA TGGCGCGATT AATCGTGATA TTGTTGGTAA GTCCGCATAC
     901  ATTATCGCGA AGATGGCGGG TATCGAAGTG CCGCAAACCA CCAAAATCTT GATCGGCGAA
     961  GTGCAGAGCG TGGAGAAGAG CGAACTGTTC AGCCATGAGA AACTGAGCCC GGTTCTGGCT
    1021  ATGTACAAAG TGAAGGACTT TGACGAAGCC CTGAAGAAAG CTCAGCGTCT GATTGAGCTG
    1081  GGCGGCAGCG GTCACACGAG CAGCCTGTAT ATCGATAGCC AAAACAATAA GGACAAGGTG
    1141  AAAGAGTTCG GTCTGGCGAT GAAAACCTCT CGCACGTTTA TCAACATGCC GAGCAGCCAA
    1201  GGTGCGAGCG GCGATCTGTA TAACTTCGCA ATCGCGCCTT CTTTTACGCT GGGTTGCGGT
    1261  ACCTGGGGCG GTAATTCGGT GAGCCAGAAT GTCGAGCCGA AGCACCTGCT GAACATCAAA
    1321  TCGGTCGCGG AGCGCCGTGA GAAGAGACCG TGGTTCAAAG TCCCACAGAA AATTTACTTC
    1381  AAATACGGCT GCCTGCGTTT CGCGCTGAAA GAACTGAAAG ACATGAACAA AAAGCGTGCG
    1441  TTCATTGTTA CCGACAAAGA CCTGTTCAAA CTGGGTTACG TGAACAAAAT CACCAAAGTT
    1501  CTGGATGAAA TTGACATCAA GTACTCCATC TTCACTGATA TCAAATCCGA CCCAACGATT
    1561  GATAGCGTGA AAAAGGGCGC TAAAGAAATG CTGAACTTTG AACCGGACAC CATCATCAGC
    1621  ATCGGTGGTG GCTCTCCTAT GGATGCTGCG AAGGTCATGC ACCTGCTGTA CGAATACCCG
    1681  GAAGCGGAAA TCGAAAACCT GGCTATCAAC TTCATGGACA TCCGCAAACG TATCTGCAAC
    1741  TTCCCGAAGC TGGGCACTAA AGCTATTTCC GTTGCCATCC CGACTACCGC GGGCACTGGT
    1801  TCCGAAGCCA CGCCGTTCGC CGTGATCACC AACGATGAAA CCGGTATGAA ATACCCGCTG
    1861  ACCTCTTACG AACTGACCCC GAACATGGCA ATTATCGACA CCGAGCTGAT GCTGAACATG
    1921  CCGCGCAAGC TGACCGCTGC TACCGGCATC GACGCTCTGG TACATGCTAT TGAGGCGTAC
    1981  GTTTCCGTGA TGGCTACCGA TTACACCGAC GAACTGGCCC TGCGTGCGAT CAAAATGATT
    2041  TTCAAGTACC TGCCTCGCGC TTACAAAAAC GGCACGAATG ACATCGAGGC GCGTGAGAAA
    2101  ATGGCCCATG CAAGCAACAT CGCGGGCATG GCCTTCGCCA ACGCGTTCCT GGGCGTGTGC
    2161  CACTCTATGG CTCACAAACT GGGTGCTATG CACCACGTGC CGCACGGTAT CGCGTGTGCT
    2221  GTCCTGATCG AAGAAGTAAT TAAGTACAAC GCTACTGATT GCCCGACTAA ACAGACCGCC
    2281  TTCCCACAGT ACAAATCTCC TAACGCTAAA CGTAAGTACG CTGAGATCGC CGAATACCTG
    2341  AACCTGAAGG GTACGAGCGA CACTGAGAAA GTTACTGCGC TGATCGAAGC TATCTCTAAA
    2401  CTGAAAATTG ACCTGTCCAT CCCGCAGAAC ATCAGCGCCG CAGGCATCAA CAAAAAGGAC
    2461  TTTTACAACA CGCTGGACAA AATGAGCGAA CTGGCTTTTG ACGACCAGTG CACCACTGCA
    2521  AACCCGCGTT ACCCGCTGAT CTCTGAGCTG AAAGACATTT ATATCAAATC TTTTTAA
```

**AdhE2.24 (76953)**

```
       1  ATGAAAGTGA CGAACCAAAA AGAACTGAAA CAAAAACTGA ATGAACTGCG TGAGGCCCAG
      61  AAAAAGTTCA ACACGTATAC CCAGGAGCAA GTTGACAAGA TCTTCAAACA GTGTGCCATT
```

121 GCGGCTGCCA AAGAGCGTAT TAACTTGGCC AAACTGGCGG TGGAAGAAAC CGGCATCGGT

181 CTGGTCGAGG ACAAGATTAT CAAGAATCAC TTTGCCGCGG AGTATATCTA CAACAAATAC

```
     241  AAAAACGAAA AGACTTGCGG CATCATTGAT CATGATGACA GCTTGGGTAT CACGAAAGTC
     301  GCAGAACCAA TCGGTATTGT CGCGGCAATT GTCCCGACCA CTAATCCGAC CAGCACCGCG
     361  ATCTTCAAGA GCTTGATCAG CCTGAAAACC CGCAACGCGA TTTTCTTCAG CCCGCACCCG
     421  CGTGCGAAGA AGAGCACCAT CGCAGCGGCG AAGCTGATTT TGGATGCCGC GGTTAAGGCC
     481  GGTGCACCGA AAAACATTAT TGGCTGGATT GACGAGCCGA GCATTGAGCT GTCTCAGGAC
     541  CTGATGAGCG AAGCGGACAT TATTCTGGCC ACCGGTGGTC CGTCTATGGT TAAGGCAGCG
     601  TACTCCAGCG GTAAGCCTGC TATCGGCGTT GGTGCGGGCA ATACGCCGGC TATCATCGAT
     661  GAAAGCGCGG ATATTGATAT GGCAGTTAGC TCCATCATTC TGTCCGCGAC CTACGACAAT
     721  GGTGTTATCT GTGCAAGCGA GCAAAGCATC CTGGTGATGA ATAGCATTTA CGAGAAGGTC
     781  AAAGAAGAGT TTGTTAAGCG TGGTTCCTAT ATTCTGAACC AGAACGAGAT TGCAAAAATC
     841  AAAGAAACGA TGTTTAAGAA TGGCGCGATT AATGCAGATA TTGTTGGTAA GTCCGCATAC
     901  ATTATCGCGA AGATGGCGGG TATCGAAGTG CCGCAAACCA CCAAAATCTT GATCGGCGAA
     961  GTGCAGAGCG TGGAGAAGAG CGAACTGTTC AGCCATGAGA AACTGAGCCC GGTTCTGGCT
    1021  ATGTACAAAG TGAAGGACTT TGACGAAGCC CTGAAGAAAG CTGAGCGTCT GATTGAGCTG
    1081  GGCGGCAGCG GTCACACGAG CAGCCTGTAT ATCGATAGCC AAAACAATAA GGACAAGGTG
    1141  AAAGAGTTCG GTCTGGCGAT GAAAACCTCT CGCACGTTTA TCAACATGCC GAGCAGCCAA
    1201  GGTGCGAGCG GCGATCTGTA TAACTTCGCA ATCGCGCCTT CTTTTACGCT GGGTTGCGGT
    1261  ACCTGGGGCG GTAATTCGGT GAGCCAGAAT GTCGAGCCGA AGCACCTGCT GAACATCAAA
    1321  TCGGTCGCGG AGCGCCGTGA GAAGAGACCG TGGTTCAAAG TCCCACAGAA AATTTACTTC
    1381  AAATACGGCT GCCTGCGTTT CGCGCTGAAA GAACTGAAAG ACATGAACAA AAAGCGTGCG
    1441  TTCATTGTTA CCGACAAAGA CCTGTTCAAA CTGGGTTACG TGAACAAAAT CACCAAAGTT
    1501  CTGGATGAAA TTGACATCAA GTACTCCATC TTCACTGATA TCAAATCCGA CCCAACGATT
    1561  GATAGCGTGA AAAAGGGCGC TAAAGAAATG CTGAACTTTG AACCGGACAC CATCATCAGC
    1621  ATCGGTGGTG GCTCTCCTAT GGATGCTGCG AAGGTCATGC ACCTGCTGTA CGAATACCCG
    1681  GAAGCGGAAA TCGAAAACCT GGCTATCAAC TTCATGGACA TCCGCAAACG TATCTGCAAC
    1741  TTCCCGAAGC TGGGCACTAA AGCTATTTCC GTTGCCATCC CGACTACCGC GGGCACTGGT
    1801  TCCGAAGCCA CGCCGTTCGC CGTGATCACC AACGATGAAA CCGGTATGAA ATACCCGCTG
    1861  ACCTCTTACG AACTGACCCC GAACATGGCA ATTATCGACA CCGAGCTGAT GCTGAACATG
    1921  CCGCGCAAGC TGACCGCTGC TACCGGCATC GACGCTCTGG TACATGCTAT TGAGGCGTAC
    1981  GTTTCCGTGA TGGCTACCGA TTACACCGAC GAACTGGCCC TGCGTGCGAT CAAAATGATT
    2041  TTCAAGTACC TGCCTCGCGC TTACAAAAAC GGCACGAATG ACATCGAGGC GCGTGAGAAA
    2101  ATGGCCCATG CAAGCAACAT CGCGGGCATG GCCTTCGCCA ACGCGTTCCT GGGCGTGTGC
    2161  CACTCTATGG CTCACAAACT GGGTGCTATG CACCACGTGC CGCACGGTAT CGCGTGTGCT
    2221  GTCCTGATCG AAGAAGTAAT TAAGTACAAC GCTACTGATT GCCCGACTAA ACAGACCGCC
    2281  TTCCCACAGT ACAAATCTCC TAACGCTAAA CGTAAGTACG CTGAGATCGC CGAATACCTG
    2341  AACCTGAAGG GTACGAGCGA CACTGAGAAA GTTACTGCGC TGATCGAAGC TATCTCTAAA
    2401  CTGAAAATTG ACCTGTCCAT CCCGCAGAAC ATCAGCGCCG CAGGCATCAA CAAAAAGGAC
    2461  TTTTACAACA CGCTGGACAA AATGAGCGAA CTGGCTTTTG ACGACCAGTG CACCACTGCA
    2521  AACCCGCGTT ACCCGCTGAT CTCTGAGCTG AAAGACATTT ATATCAAATC TTTTTAA
```

**AdhE2.25 (76954)**

```
       1  ATGAAAGTGA CGAACCAAAA AGAACTGAAA CAAAAACTGA ATGAACTGCG TGAGGCCCAG
      61  AAAAAGTTCG CTACGTATAC CCAGGAGCAA GTTGACAAGA TCTTCAAACA GTGTGCCATT
```

121 GCGGCTGCCA AACTGCGTAT TAACTTGGCC AAACTGGCGG TGGAAGAAAC CGGCATCGGT

181 CTGGTCGAGG ACAAGATTAT CAAGAATCAC TTTGCCGCGG AGTATATCTA CAACAAATAC

```
     241  AAAAACGAAA AGACTTGCGG CATCATTGAT CATGATGACA GCTTGGGTAT CACGAAAGTC
     301  GCAGAACCAA TCGGTATTGT CGCGGCAATT GTCCCGACCA CTAATCCGAC CAGCACCGCG
     361  ATCTTCAAGA GCTTGATCAG CCTGAAAACC CGCAACGCGA TTTTCTTCAG CCCGCACCCG
     421  CGTGCGAAGA AGAGCACCAT CGCAGCGGCG ACCCTGATTT TGGATGCCGC GGTTAAGGCC
     481  GGTGCACCGA AAAACATTAT TGGCTGGATT GACGAGCCGA GCATTGAGCT GTCTCAGGAC
     541  CTGATGAGCG AAGCGGACAT TATTCTGGCC ACCGGTGGTC CGTCTATGGT TAAGGCAGCG
     601  TACTCCAGCG GTAAGCCTGC TATCGGCGTT GGTGCGGGCA ATACGCCGGC TATCATCGAT
     661  GAAAGCGCGG ATATTGATAT GGCAGTTAGC TCCATCATTC TGTCCAAAAC CTACGACAAT
     721  GGTGTTATCT GTGCAAGCGA GCAAAGCATC CTGGTGATGA ATAGCATTTA CGAGAAGGTC
     781  AAAGAAGAGT TTGTTAAGCG TGGTTCCTAT ATTCTGAACC AGAACGAGAT TGCAAAAATC
     841  AAAGAAACGA TGTTTAAGAA TGGCGCGATT AATGCAGATA TTGTTGGTAA GTCCGCATAC
     901  ATTATCGCGA AGATGGCGGG TATCGAAGTG CCGCAAACCA CCAAAATCTT GATCGGCGAA
     961  GTGCAGAGCG TGGAGAAGAG CGAACTGTTC AGCCATGAGA AACTGAGCCC GGTTCTGGCT
    1021  ATGTACAAAG TGAAGGACTT TGACGAAGCC CTGAAGCTGG CTCAGCGTCT GATTGAGCTG
    1081  GGCGGCAGCG GTCACACGAG CAGCCTGTAT ATCGATAGCC AAAACAATAA GGACAAGGTG
    1141  AAAGAGTTCG GTCTGGCGAT GAAAACCTCT CGCACGTTTA TCAACATGCC GAGCAGCCAA
    1201  GGTGCGAGCG GCGATCTGTA TAACTTCGCA ATCGCGCCTT CTTTTACGCT GGGTTGCGGT
    1261  ACCTGGGGCG GTAATTCGGT GAGCCAGAAT GTCGAGCCGA AGCACCTGCT GAACATCAAA
    1321  TCGGTCGCGG AGCGCCGTGA GAAGAGACCG TGGTTCAAAG TCCCACAGAA AATTTACTTC
    1381  AAATACGGCT GCCTGCGTTT CGCGCTGAAA GAACTGAAAG ACATGAACAA AAAGCGTGCG
    1441  TTCATTGTTA CCGACAAAGA CCTGTTCAAA CTGGGTTACG TGAACAAAAT CACCAAAGTT
    1501  CTGGATGAAA TTGACATCAA GTACTCCATC TTCACTGATA TCAAATCCGA CCCAACGATT
    1561  GATAGCGTGA AAAAGGGCGC TAAAGAAATG CTGAACTTTG AACCGGACAC CATCATCAGC
    1621  ATCGGTGGTG GCTCTCCTAT GGATGCTGCG AAGGTCATGC ACCTGCTGTA CGAATACCCG
    1681  GAAGCGGAAA TCGAAAACCT GGCTATCAAC TTCATGGACA TCCGCAAACG TATCTGCAAC
    1741  TTCCCGAAGC TGGGCACTAA AGCTATTTCC GTTGCCATCC CGACTACCGC GGGCACTGGT
    1801  TCCGAAGCCA CGCCGTTCGC CGTGATCACC AACGATGAAA CCGGTATGAA ATACCCGCTG
    1861  ACCTCTTACG AACTGACCCC GAACATGGCA ATTATCGACA CCGAGCTGAT GCTGAACATG
    1921  CCGCGCAAGC TGACCGCTGC TACCGGCATC GACGCTCTGG TACATGCTAT TGAGGCGTAC
    1981  GTTTCCGTGA TGGCTACCGA TTACACCGAC GAACTGGCCC TGCGTGCGAT CAAAATGATT
    2041  TTCAAGTACC TGCCTCGCGC TTACAAAAAC GGCACGAATG ACATCGAGGC GCGTGAGAAA
    2101  ATGGCCCATG CAAGCAACAT CGCGGGCATG GCCTTCGCCA ACGCGTTCCT GGGCGTGTGC
    2161  CACTCTATGG CTCACAAACT GGGTGCTATG CACCACGTGC CGCACGGTAT CGCGTGTGCT
    2221  GTCCTGATCG AAGAAGTAAT TAAGTACAAC GCTACTGATT GCCCGACTAA ACAGACCGCC
    2281  TTCCCACAGT ACAAATCTCC TAACGCTAAA CGTAAGTACG CTGAGATCGC CGAATACCTG
    2341  AACCTGAAGG GTACGAGCGA CACTGAGAAA GTTACTGCGC TGATCGAAGC TATCTCTAAA
    2401  CTGAAAATTG ACCTGTCCAT CCCGCAGAAC ATCAGCGCCG CAGGCATCAA CAAAAAGGAC
    2461  TTTTACAACA CGCTGGACAA AATGAGCGAA CTGGCTTTTG ACGACCAGTG CACCACTGCA
    2521  AACCCGCGTT ACCCGCTGAT CTCTGAGCTG AAAGACATTT ATATCAAATC TTTTTAA
```

**AdhE2.26 (76955)**

```
       1  ATGAAAGTGA CGAACCAAAA AGAACTGAAA CAAAAACTGA ATGAACTGCG TGAGGCCCAG
      61  AAAAAGTTCC TGACGATGAC CCAGGAGCAA GTTGACAAGA TCTTCAAACA GTGTGCCATT
```

121 GCGGCTGCCA AAGAGCGTAT TAACTTGGCC AAACTGGCGG TGGAAGAAAC CGGCATCGGT

181 CTGGTCGAGG ACAAGATTAT CAAGAATCAC TTTGCCGCGG AGTATATCTA CAACAAATAC

```
     241  AAAAACGAAA AGACTTGCGG CATCATTGAT CATGATGACA GCTTGGGTAT CACGAAAGTC
     301  GCAGAACCAA TCGGTATTGT CGCGGCAATT GTCCCGACCA CTAATCCGAC CAGCACCGCG
     361  ATCTTCAAGA GCTTGATCAG CCTGAAAACC CGCAACGCGA TTTTCTTCAG CCCGCACCCG
     421  CGTGCGAAGA AGAGCACCAT CGCAGCGGCG AAGCTGATTT TGGATGCCGC GGTTAAGGCC
     481  GGTGCACCGA AAAACATTAT TGGCTGGATT GACGAGCCGA GCATTGAGCT GTCTCAGGAC
     541  CTGATGAGCG AAGCGGACAT TATTCTGGCC ACCGGTGGTC CGTCTATGGT TAAGGCAGCG
     601  TACTCCAGCG GTAAGCCTGC TATCGGCGTT GGTGCGGGCA ATACGCCGGC TATCATCGAT
     661  GAAAGCGCGG ATATTGATAT GGCAGTTAGC TCCATCATTC TGTCCAAAAC CTACGACAAT
     721  GGTGTTATCT GTGCAAGCGA GCAAAGCATC CTGGTGATGA ATAGCATTTA CGAGAAGGTC
     781  AAAGAAGAGT TTGTTAAGCG TGGTTCCTAT ATTCTGAACC AGAACGAGAT TGCAAAAATC
     841  AAAGAAACGA TGTTTAAGAA TGGCGCGATT AATGCAGATA TTGTTGGTAA GTCCGCATAC
     901  ATTATCGCGA AGATGGCGGG TATCGAAGTG CCGCAAACCA CCAAAATCTT GATCGGCGAA
     961  GTGCAGAGCG TGGAGAAGAG CGAACTGTTC AGCCATGAGA AACTGAGCCC GGTTCTGGCT
    1021  ATGTACAAAG TGAAGGACTT TGACGAAGCC CTGAAGAAAG CTCAGCGTCT GATTCACCTG
    1081  GGCGGCAGCG GTCACACGAG CAGCCTGTAT ATCGATAGCC AAAACAATAA GGACAAGGTG
    1141  AAAGAGTTCG GTCTGGCGAT GAAAACCTCT CGCACGTTTA TCAACATGCC GAGCAGCCAA
    1201  GGTGCGAGCG GCGATCTGTA TAACTTCGCA ATCGCGCCTT CTTTTACGCT GGGTTGCGGT
    1261  ACCTGGGGCG GTAATTCGGT GAGCCAGAAT GTCGAGCCGA AGCACCTGCT GAACATCAAA
    1321  TCGGTCGCGG AGCGCCGTGA GAAGAGACCG TGGTTCAAAG TCCCACAGAA AATTTACTTC
    1381  AAATACGGCT GCCTGCGTTT CGCGCTGAAA GAACTGAAAG ACATGAACAA AAAGCGTGCG
    1441  TTCATTGTTA CCGACAAAGA CCTGTTCAAA CTGGGTTACG TGAACAAAAT CACCAAAGTT
    1501  CTGGATGAAA TTGACATCAA GTACTCCATC TTCACTGATA TCAAATCCGA CCCAACGATT
    1561  GATAGCGTGA AAAAGGGCGC TAAAGAAATG CTGAACTTTG AACCGGACAC CATCATCAGC
    1621  ATCGGTGGTG GCTCTCCTAT GGATGCTGCG AAGGTCATGC ACCTGCTGTA CGAATACCCG
    1681  GAAGCGGAAA TCGAAAACCT GGCTATCAAC TTCATGGACA TCCGCAAACG TATCTGCAAC
    1741  TTCCCGAAGC TGGGCACTAA AGCTATTTCC GTTGCCATCC CGACTACCGC GGGCACTGGT
    1801  TCCGAAGCCA CGCCGTTCGC CGTGATCACC AACGATGAAA CCGGTATGAA ATACCCGCTG
    1861  ACCTCTTACG AACTGACCCC GAACATGGCA ATTATCGACA CCGAGCTGAT GCTGAACATG
    1921  CCGCGCAAGC TGACCGCTGC TACCGGCATC GACGCTCTGG TACATGCTAT TGAGGCGTAC
    1981  GTTTCCGTGA TGGCTACCGA TTACACCGAC GAACTGGCCC TGCGTGCGAT CAAAATGATT
    2041  TTCAAGTACC TGCCTCGCGC TTACAAAAAC GGCACGAATG ACATCGAGGC GCGTGAGAAA
    2101  ATGGCCCATG CAAGCAACAT CGCGGGCATG GCCTTCGCCA ACGCGTTCCT GGGCGTGTGC
    2161  CACTCTATGG CTCACAAACT GGGTGCTATG CACCACGTGC CGCACGGTAT CGCGTGTGCT
    2221  GTCCTGATCG AAGAAGTAAT TAAGTACAAC GCTACTGATT GCCCGACTAA ACAGACCGCC
    2281  TTCCCACAGT ACAAATCTCC TAACGCTAAA CGTAAGTACG CTGAGATCGC CGAATACCTG
    2341  AACCTGAAGG GTACGAGCGA CACTGAGAAA GTTACTGCGC TGATCGAAGC TATCTCTAAA
    2401  CTGAAAATTG ACCTGTCCAT CCCGCAGAAC ATCAGCGCCG CAGGCATCAA CAAAAAGGAC
    2461  TTTTACAACA CGCTGGACAA AATGAGCGAA CTGGCTTTTG ACGACCAGTG CACCACTGCA
    2521  AACCCGCGTT ACCCGCTGAT CTCTGAGCTG AAAGACATTT ATATCAAATC TTTTTAA
```

**AdhE2.27 (76956)**

```
       1  ATGAAAGTGA CGAACCAAAA AGAACTGAAA CAAAAACTGA ATGAACTGCG TGAGGCCCAG
      61  AAAAAGTTCG CTTGCTATAC CCAGGAGCAA GTTGAGAAGA TCTTCAAACA GTGTGCCATT
```

121 GCGGCTGCCA AAGAGCGTAT TAACTTGGCC AAACTGGCGG TGGAAGAAAC CGGCATCGGT

181 CTGGTCGAGG ACAAGATTAT CAAGAATCAC TTTGCCGCGG AGTATATCTA CAACAAATAC

```
     241  AAAAACGAAA AGACTTGCGG CATCATTGAT CATGATGACA GCTTGGGTAT CACGAAAGTC
     301  GCAGAACCAA TCGGTATTGT CGCGGCAATT GTCCCGACCA CTAATCCGAC CAGCACCGCG
     361  ATCTTCAAGA GCTTGATCAG CCTGAAAACC CGCAACGCGA TTTTCTTCAG CCCGCACCCG
     421  CGTGCGAAGA AGAGCACCAT CGCAGCGGCG AAGCTGATTT TGGATGCCGC GGTTAAGGCC
     481  GGTGCACCGA AAAACATTAT TGGCTGGATT GACGAGCCGA GCATTGAGCT GTCTCAGGAC
     541  CTGATGAGCG AAGCGGACAT TATTCTGGCC ACCGGTGGTC CGTCTATGGT TAAGGCAGCG
     601  TACTCCAGCG GTAAGCCTGC TATCGGCGTT GGTGCGGGCA ATACGCCGGC TATCATCGAT
     661  GAAAGCGCGG ATATTGATAT GGCAGTTAGC TCCATCATTC TGTCCAAAAC CTACGACAAT
     721  GGTGTTATCT GTGCAAGCGA GCAAAGCATC CTGGTGATGA ATAGCATTTA CGAGAAGGTC
     781  AAAGAAGAGT TTGTTAAGCG TGGTTCCTAT ATTCTGAACC AGAACGAGAT TGCAAAAATC
     841  AAAGAAACGA TGTTTAAGAA TGGCGCGATT AATGCAGATA TTGTTGGTAA GTCCGCATAC
     901  ATTATCGCGA AGATGGCGGG TATCGAAGTG CCGCAAACCA CCAAAATCTT GATCGGCGAA
     961  GTGCAGAGCG TGGAGAAGAG CGAACTGTTC AGCCATGAGA AACTGAGCCC GGTTCTGGCT
    1021  ATGTACAAAG TGAAGGACTT TGACGAAGCC CTGAAGAAAG CTCAGCGTCT GATTGAGCTG
    1081  GGCGGCAGCC ACCACACGAG CAGCCTGTAT ATCGATAGCC AAAACAATAA GGACAAGGTG
    1141  AAAGAGTTCG GTCTGGCGAT GAAAACCTCT CGCACGTTTA TCAACATGCC GAGCAGCCAA
    1201  GGTGCGAGCG GCGATCTGTA TAACTTCGCA ATCGCGCCTT CTTTTACGCT GGGTTGCGGT
    1261  ACCTGGGGCG GTAATTCGGT GAGCCAGAAT GTCGAGCCGA AGCACCTGCT GAACATCAAA
    1321  TCGGTCGCGG AGCGCCGTGA GAAGAGACCG TGGTTCAAAG TCCCACAGAA AATTTACTTC
    1381  AAATACGGCT GCCTGCGTTT CGCGCTGAAA GAACTGAAAG ACATGAACAA AAAGCGTGCG
    1441  TTCATTGTTA CCGACAAAGA CCTGTTCAAA CTGGGTTACG TGAACAAAAT CACCAAAGTT
    1501  CTGGATGAAA TTGACATCAA GTACTCCATC TTCACTGATA TCAAATCCGA CCCAACGATT
    1561  GATAGCGTGA AAAAGGGCGC TAAAGAAATG CTGAACTTTG AACCGGACAC CATCATCAGC
    1621  ATCGGTGGTG GCTCTCCTAT GGATGCTGCG AAGGTCATGC ACCTGCTGTA CGAATACCCG
    1681  GAAGCGGAAA TCGAAAACCT GGCTATCAAC TTCATGGACA TCCGCAAACG TATCTGCAAC
    1741  TTCCCGAAGC TGGGCACTAA AGCTATTTCC GTTGCCATCC CGACTACCGC GGGCACTGGT
    1801  TCCGAAGCCA CGCCGTTCGC CGTGATCACC AACGATGAAA CCGGTATGAA ATACCCGCTG
    1861  ACCTCTTACG AACTGACCCC GAACATGGCA ATTATCGACA CCGAGCTGAT GCTGAACATG
    1921  CCGCGCAAGC TGACCGCTGC TACCGGCATC GACGCTCTGG TACATGCTAT TGAGGCGTAC
    1981  GTTTCCGTGA TGGCTACCGA TTACACCGAC GAACTGGCCC TGCGTGCGAT CAAAATGATT
    2041  TTCAAGTACC TGCCTCGCGC TTACAAAAAC GGCACGAATG ACATCGAGGC GCGTGAGAAA
    2101  ATGGCCCATG CAAGCAACAT CGCGGGCATG GCCTTCGCCA ACGCGTTCCT GGGCGTGTGC
    2161  CACTCTATGG CTCACAAACT GGGTGCTATG CACCACGTGC CGCACGGTAT CGCGTGTGCT
    2221  GTCCTGATCG AAGAAGTAAT TAAGTACAAC GCTACTGATT GCCCGACTAA ACAGACCGCC
    2281  TTCCCACAGT ACAAATCTCC TAACGCTAAA CGTAAGTACG CTGAGATCGC CGAATACCTG
    2341  AACCTGAAGG GTACGAGCGA CACTGAGAAA GTTACTGCGC TGATCGAAGC TATCTCTAAA
    2401  CTGAAAATTG ACCTGTCCAT CCCGCAGAAC ATCAGCGCCG CAGGCATCAA CAAAAAGGAC
    2461  TTTTACAACA CGCTGGACAA AATGAGCGAA CTGGCTTTTG ACGACCAGTG CACCACTGCA
    2521  AACCCGCGTT ACCCGCTGAT CTCTGAGCTG AAAGACATTT ATATCAAATC TTTTTAA
```

**AdhE2.28 (76957)**

```
       1  ATGAAAGTGA CGAACCAAAA AGAACTGAAA CAAAAACTGA ATGAACTGCG TGAGGCCCAG
      61  AAAAAGTTCG CTACGTGCAC CCAGGAGCAA GTTGACAAGA TCTTCAAACA GTGTGCCATT
```

121 GCGGCTGCCA AAGAGCGTAT TAACTTGGCC AAACTGGCGG TGGAAGAAAC CGGCATCGGT

181 CTGGTCGAGG ACAAGATTAT CAAGAATCAC TTTGCCGCGG AGTATATCTA CAACAAATAC

```
     241  AAAAACGAAA AGACTTGCGG CATCATTGAT CATGATGACA GCTTGGGTAT CACGAAAGTC
     301  GCAGAACCAA TCGGTATTGT CGCGGCAATT GTCCCGACCA CTAATCCGAC CAGCACCGCG
     361  ATCTTCAAGA GCTTGATCAG CCTGAAAACC CGCAACGCGA TTTTCTTCAG CCCGCACCCG
     421  CGTGCGAAGA AGAGCACCAT CGCAGCGGCG AAGCTGATTT TGGATGCCGC GGTTAAGGCC
     481  GGTGCACCGA AAAACATTAT TGGCTGGATT GACGAGCCGA GCATTGAGCT GTCTCAGGAC
     541  CTGATGAGCG AAGCGGACAT TATTCTGGCC ACCGGTGGTC CGTCTATGGT TAAGGCAGCG
     601  TACTCCAGCG GTAAGCCTGC TATCGGCGTT GGTGCGGGCA ATACGCCGGC TATCATCGAT
     661  GAAAGCGCGG ATATTGATAT GGCAGTTAGC TCCATCATTC TGTCCAAAAC CTACGACAAT
     721  GGTGTTATCT GTGCAAGCGA GCAAAGCATC CTGGTGATGA ATAGCATTTA CGAGAAGGTC
     781  AAAGAAGAGT TTGTTAAGCG TGGTTCCTAT ATTCTGAACC AGAACGAGAT TGCAAAAATC
     841  AAAGAAACGA TGTTTAAGAA TGGCGCGATT AATGCAGATA TTGTTGGTAA GTCCGCATAC
     901  ATTATCGCGA AGATGGCGGG TATCGAAGTG CCGCAATTCA CCAAAATCTT GATCGGCGAA
     961  GTGCAGAGCG TGGAGAAGAG CGAACTGTTC AGCCATGAGA AACTGAGCCC GGTTCTGGCT
    1021  ATGTACAAAG TGAAGGACTT TGACGAAGCC CTGAAGAAAG CTCAGCGTCT GATTGAGCTG
    1081  GGCGGCAGCG GTCACACGAG CAGCCTGTAT ATCGATAGCC AAAACAATAA GGACAAGGTG
    1141  AAAGAGTTCG GTCTGGCGAT GAAAACCTCT CGCACGTTTA TCAACATGCC GAGCAGCCAA
    1201  GGTGCGAGCG GCGATCTGTA TAACTTCGCA ATCGCGCCTT CTTTTACGCT GGCGTGCGGT
    1261  ACCTGGGGCG GTAATTCGGT GAGCCAGAAT GTCGAGCCGA AGCACCTGCT GAACATCAAA
    1321  TCGGTCGCGG AGCGCCGTGA GAAGAGACCG TGGTTCAAAG TCCCACAGAA AATTTACTTC
    1381  AAATACGGCT GCCTGCGTTT CGCGCTGAAA GAACTGAAAG ACATGAACAA AAAGCGTGCG
    1441  TTCATTGTTA CCGACAAAGA CCTGTTCAAA CTGGGTTACG TGAACAAAAT CACCAAAGTT
    1501  CTGGATGAAA TTGACATCAA GTACTCCATC TTCACTGATA TCAAATCCGA CCCAACGATT
    1561  GATAGCGTGA AAAAGGGCGC TAAAGAAATG CTGAACTTTG AACCGGACAC CATCATCAGC
    1621  ATCGGTGGTG GCTCTCCTAT GGATGCTGCG AAGGTCATGC ACCTGCTGTA CGAATACCCG
    1681  GAAGCGGAAA TCGAAAACCT GGCTATCAAC TTCATGGACA TCCGCAAACG TATCTGCAAC
    1741  TTCCCGAAGC TGGGCACTAA AGCTATTTCC GTTGCCATCC CGACTACCGC GGGCACTGGT
    1801  TCCGAAGCCA CGCCGTTCGC CGTGATCACC AACGATGAAA CCGGTATGAA ATACCCGCTG
    1861  ACCTCTTACG AACTGACCCC GAACATGGCA ATTATCGACA CCGAGCTGAT GCTGAACATG
    1921  CCGCGCAAGC TGACCGCTGC TACCGGCATC GACGCTCTGG TACATGCTAT TGAGGCGTAC
    1981  GTTTCCGTGA TGGCTACCGA TTACACCGAC GAACTGGCCC TGCGTGCGAT CAAAATGATT
    2041  TTCAAGTACC TGCCTCGCGC TTACAAAAAC GGCACGAATG ACATCGAGGC GCGTGAGAAA
    2101  ATGGCCCATG CAAGCAACAT CGCGGGCATG GCCTTCGCCA ACGCGTTCCT GGGCGTGTGC
    2161  CACTCTATGG CTCACAAACT GGGTGCTATG CACCACGTGC CGCACGGTAT CGCGTGTGCT
    2221  GTCCTGATCG AAGAAGTAAT TAAGTACAAC GCTACTGATT GCCCGACTAA ACAGACCGCC
    2281  TTCCCACAGT ACAAATCTCC TAACGCTAAA CGTAAGTACG CTGAGATCGC CGAATACCTG
    2341  AACCTGAAGG GTACGAGCGA CACTGAGAAA GTTACTGCGC TGATCGAAGC TATCTCTAAA
    2401  CTGAAAATTG ACCTGTCCAT CCCGCAGAAC ATCAGCGCCG CAGGCATCAA CAAAAAGGAC
    2461  TTTTACAACA CGCTGGACAA AATGAGCGAA CTGGCTTTTG ACGACCAGTG CACCACTGCA
    2521  AACCCGCGTT ACCCGCTGAT CTCTGAGCTG AAAGACATTT ATATCAAATC TTTTTAA
```

**AdhE2.29 (76958)**

```
       1  ATGAAAGTGA CGAACCAAAA AGAACTGAAA CAAAAACTGA ATGAACTGCG TGAGGCCCAG
      61  AAAAAGTTCG CTACGTATAC CCAGGAGCAA GTTGACAAGA TCTTCAAACA GTGTGCCATT
```

121 GCGGCTGCCA AAGAGCGTAT TAACTTGGCC AAACTGGCGG TGGAAGAAAC CGGCATCGGT

181 CTGGTCGAGG ACAAGATTAT CAAGAATCAC TTTGCCGCGG AGTATATCTA CAACAAATAC

```
     241  AAAAACCACA AGACTTGCGG CATCATTGAT CATGATGACA GCTTGGGTAT CACGAAAGTC
     301  GCAGAACCAA TCGGTATTGT CGCGGCAATT GTCCCGACCA CTAATCCGAC CAGCACCGCG
     361  ATCTTCAAGA GCTTGATCAG CCTGAAAACC CGCAACGCGA TTTTCTTCAG CCCGCACCCG
     421  CGTGCGAAGA AGAGCACCAT CGCAGCGGCG AAGCTGATTT TGGATGCCGC GGTTAAGGCC
     481  GGTGCACCGA AAAACATTAT TGGCTGGATT GACGAGCCGA GCATTGAGCT GTCTCAGGAC
     541  CTGATGAGCG AACCGGACAT TATTCTGGCC ACCGGTGGTC CGTCTATGGT TAAGGCAGCG
     601  TACTCCAGCG GTAAGCCTGC TATCGGCGTT GGTGCGGGCA ATACGCCGGC TATCATCGAT
     661  GAAAGCGCGG ATATTGATAT GGCAGTTAGC TCCATCATTC TGTCCAAAAC CTACGACAAT
     721  GGTGTTATCT GTGCAAGCGA GCAAAGCATC CTGGTGATGA ATAGCATTTA CGAGAAGGTC
     781  AAAGAAGAGT TTGTTAAGCG TGGTTCCTAT ATTCTGAACC AGAACGAGAT TGCAAAAATC
     841  AAAGAAACGA TGTTTAAGAA TGGCGCGATT AATGCAGATA TTGTTGGTAA GTCCGCATAC
     901  ATTATCGCGA AGATGGCGGG TATCGAAGTG CCGCAAACCA CCAAAATCTT GATCGGCGAA
     961  GTGCAGAGCG TGGAGAAGAG CGAACTGTTC AGCCATGAGA AACTGAGCCC GGTTCTGGCT
    1021  ATGTACAAAG TGAAGGACTT TGACGAAGCC CTGAAGAAAG CTCAGCGTCT GATTAACCTG
    1081  GGCGGCAGCG GTCACACGAG CAGCCTGTAT ATCGATAGCC AAAACAATAA GGACAAGGTG
    1141  AAAGAGTTCG GTCTGGCGAT GAAAACCTCT CGCACGTTTA TCAACATGCC GAGCAGCCAA
    1201  GGTGCGAGCG GCGATCTGTA TAACTTCGCA ATCGCGCCTT CTTTTACGCT GGGTTGCGGT
    1261  ACCTGGGGCG GTAATTCGGT GAGCCAGAAT GTCGAGCCGA AGCACCTGCT GAACATCAAA
    1321  TCGGTCGCGG AGCGCCGTGA GAAGAGACCG TGGTTCAAAG TCCCACAGAA AATTTACTTC
    1381  AAATACGGCT GCCTGCGTTT CGCGCTGAAA GAACTGAAAG ACATGAACAA AAAGCGTGCG
    1441  TTCATTGTTA CCGACAAAGA CCTGTTCAAA CTGGGTTACG TGAACAAAAT CACCAAAGTT
    1501  CTGGATGAAA TTGACATCAA GTACTCCATC TTCACTGATA TCAAATCCGA CCCAACGATT
    1561  GATAGCGTGA AAAAGGGCGC TAAAGAAATG CTGAACTTTG AACCGGACAC CATCATCAGC
    1621  ATCGGTGGTG GCTCTCCTAT GGATGCTGCG AAGGTCATGC ACCTGCTGTA CGAATACCCG
    1681  GAAGCGGAAA TCGAAAACCT GGCTATCAAC TTCATGGACA TCCGCAAACG TATCTGCAAC
    1741  TTCCCGAAGC TGGGCACTAA AGCTATTTCC GTTGCCATCC CGACTACCGC GGGCACTGGT
    1801  TCCGAAGCCA CGCCGTTCGC CGTGATCACC AACGATGAAA CCGGTATGAA ATACCCGCTG
    1861  ACCTCTTACG AACTGACCCC GAACATGGCA ATTATCGACA CCGAGCTGAT GCTGAACATG
    1921  CCGCGCAAGC TGACCGCTGC TACCGGCATC GACGCTCTGG TACATGCTAT TGAGGCGTAC
    1981  GTTTCCGTGA TGGCTACCGA TTACACCGAC GAACTGGCCC TGCGTGCGAT CAAAATGATT
    2041  TTCAAGTACC TGCCTCGCGC TTACAAAAAC GGCACGAATG ACATCGAGGC GCGTGAGAAA
    2101  ATGGCCCATG CAAGCAACAT CGCGGGCATG GCCTTCGCCA ACGCGTTCCT GGGCGTGTGC
    2161  CACTCTATGG CTCACAAACT GGGTGCTATG CACCACGTGC CGCACGGTAT CGCGTGTGCT
    2221  GTCCTGATCG AAGAAGTAAT TAAGTACAAC GCTACTGATT GCCCGACTAA ACAGACCGCC
    2281  TTCCCACAGT ACAAATCTCC TAACGCTAAA CGTAAGTACG CTGAGATCGC CGAATACCTG
    2341  AACCTGAAGG GTACGAGCGA CACTGAGAAA GTTACTGCGC TGATCGAAGC TATCTCTAAA
    2401  CTGAAAATTG ACCTGTCCAT CCCGCAGAAC ATCAGCGCCG CAGGCATCAA CAAAAAGGAC
    2461  TTTTACAACA CGCTGGACAA AATGAGCGAA CTGGCTTTTG ACGACCAGTG CACCACTGCA
    2521  AACCCGCGTT ACCCGCTGAT CTCTGAGCTG AAAGACATTT ATATCAAATC TTTTTAA
```

**AdhE2.30 (76959)**

```
       1  ATGAAAGTGA CGAACCAAAA AGAACTGAAA CAAAAACTGA ATGAACTGCG TGAGGCCCAG
      61  AAAAAGTTCG CTTGCTATAC CCAGGAGCAA GTTGACAAGA TCTTCAAACA GTGTGCCATT
```

121 GCGGCTGCCA AAGAGCGTAT TAACTTGGCC AAACTGGCGG TGGAAGAAAC CGGCATCGGT

181 CTGGTCGAGG ACAAGATTAT CAAGAATCAC TTTGCCGCGG AGTATATCTA CAACAAAGAG

```
     241  AAAAACGAAA AGACTTGCGG CATCATTGAT CATGATGACA GCTTGGGTAT CACGAAAGTC
     301  GCAGAACCAA TCGGTATTGT CGCGGCAATT GTCCCGACCA CTAATCCGAC CAGCACCGCG
     361  ATCTTCAAGA GCTTGATCAG CCTGAAAACC CGCAACGCGA TTTTCTTCAG CCCGCACCCG
     421  CGTGCGAAGA AGAGCACCAT CGCAGCGGCG AAGCTGATTT TGGATGCCGC GGTTAAGGCC
     481  GGTGCACCGA AAAACATTAT TGGCTGGATT GACGAGCCGA GCATTGAGCT GTCTCAGGAC
     541  ATGATGAGCG AAGCGGACAT TATTCTGGCC ACCGGTGGTC CGTCTATGGT TAAGGCAGCG
     601  TACTCCAGCG GTAAGCCTGC TATCGGCGTT GGTGCGGGCA ATACGCCGGC TATCATCGAT
     661  GAAAGCGCGG ATATTGATAT GGCAGTTAGC TCCATCATTC TGTCCAAAAC CTACGACAAT
     721  GGTGTTATCT GTGCAAGCGA GCAAAGCATC CTGGTGATGA ATAGCATTTA CGAGAAGGTC
     781  AAAGAAGAGT TTGTTAAGCG TGGTTCCTAT ATTCTGAACC AGAACGAGAT TGCAAAAATC
     841  AAAGAAACGA TGTTTAAGAA TGGCGCGATT AATGCAGATA TTGTTGGTAA GTCCGCATAC
     901  ATTATCGCGA AGATGGCGGG TATCGAAGTG CCGCAAACCA CCAAAATCTT GATCGGCGAA
     961  GTGCAGAGCG TGGAGAAGAG CGAACTGTTC AGCCATGAGA AACTGAGCCC GGTTCTGGCT
    1021  ATGTACAAAG TGAAGGACTT TGACGAAGCC CTGAAGAAAG CTCAGCGTCT GATTGAGCTG
    1081  GGCGGCAGCG GTCACACGAG CAGCCTGTAT ATCGATAGCC AAAACAATAA GGACAAGGTG
    1141  AAAGAGTTCG GTCTGGCGAT GAAAACCTCT CGCACGTTTA TCAACATGCC GAGCAGCCAA
    1201  GGTGCGAGCG GCGATCTGTA TAACTTCGCA ATCGCGCCTT CTTTTACGCT GGGTTGCGGT
    1261  ACCTGGGGCG GTAATTCGGT GAGCCAGAAT GTCGAGCCGA AGCACCTGCT GAACATCAAA
    1321  TCGGTCGCGG AGCGCCGTGA GAAGAGACCG TGGTTCAAAG TCCCACAGAA AATTTACTTC
    1381  AAATACGGCT GCCTGCGTTT CGCGCTGAAA GAACTGAAAG ACATGAACAA AAAGCGTGCG
    1441  TTCATTGTTA CCGACAAAGA CCTGTTCAAA CTGGGTTACG TGAACAAAAT CACCAAAGTT
    1501  CTGGATGAAA TTGACATCAA GTACTCCATC TTCACTGATA TCAAATCCGA CCCAACGATT
    1561  GATAGCGTGA AAAAGGGCGC TAAAGAAATG CTGAACTTTG AACCGGACAC CATCATCAGC
    1621  ATCGGTGGTG GCTCTCCTAT GGATGCTGCG AAGGTCATGC ACCTGCTGTA CGAATACCCG
    1681  GAAGCGGAAA TCGAAAACCT GGCTATCAAC TTCATGGACA TCCGCAAACG TATCTGCAAC
    1741  TTCCCGAAGC TGGGCACTAA AGCTATTTCC GTTGCCATCC CGACTACCGC GGGCACTGGT
    1801  TCCGAAGCCA CGCCGTTCGC CGTGATCACC AACGATGAAA CCGGTATGAA ATACCCGCTG
    1861  ACCTCTTACG AACTGACCCC GAACATGGCA ATTATCGACA CCGAGCTGAT GCTGAACATG
    1921  CCGCGCAAGC TGACCGCTGC TACCGGCATC GACGCTCTGG TACATGCTAT TGAGGCGTAC
    1981  GTTTCCGTGA TGGCTACCGA TTACACCGAC GAACTGGCCC TGCGTGCGAT CAAAATGATT
    2041  TTCAAGTACC TGCCTCGCGC TTACAAAAAC GGCACGAATG ACATCGAGGC GCGTGAGAAA
    2101  ATGGCCCATG CAAGCAACAT CGCGGGCATG GCCTTCGCCA ACGCGTTCCT GGGCGTGTGC
    2161  CACTCTATGG CTCACAAACT GGGTGCTATG CACCACGTGC CGCACGGTAT CGCGTGTGCT
    2221  GTCCTGATCG AAGAAGTAAT TAAGTACAAC GCTACTGATT GCCCGACTAA ACAGACCGCC
    2281  TTCCCACAGT ACAAATCTCC TAACGCTAAA CGTAAGTACG CTGAGATCGC CGAATACCTG
    2341  AACCTGAAGG GTACGAGCGA CACTGAGAAA GTTACTGCGC TGATCGAAGC TATCTCTAAA
    2401  CTGAAAATTG ACCTGTCCAT CCCGCAGAAC ATCAGCGCCG CAGGCATCAA CAAAAAGGAC
    2461  TTTTACAACA CGCTGGACAA AATGAGCGAA CTGGCTTTTG ACGACCAGTG CACCACTGCA
    2521  AACCCGCGTT ACCCGCTGAT CTCTGAGCTG AAAGACATTT ATATCAAATC TTTTTAA
```

**AdhE2.31 (76960)**

```
       1  ATGAAAGTGA CGAACCAAAA AGAACTGAAA CAAAAACTGA ATGAACTGCG TGAGGCCCAG
      61  AAAAAGTTCC TGACGTATAC CCAGGAGCAA GTTGACAAGA TCTTCAAACA GTGTGCCATT
```

121 GCGGCTGCCA AAGAGCGTAT TAACTTGGCC CGTCTGGCGG TGGAAGAAAC CGGCATCGGT

181 CTGGTCGAGG ACAAGATTAT CAAGAATCAC TTTGCCGCGG AGTATATCTA CAACAAATAC

```
     241  AAAAACGAAA AGACTTGCGG CATCATTGAT CATGATGACA GCTTGGGTAT CACGAAAGTC
     301  GCAGAACCAA TCGGTATTGT CGCGGCAATT GTCCCGACCA CTAATCCGAC CAGCACCGCG
     361  ATCTTCAAGA GCTTGATCAG CCTGAAAACC CGCAACGCGA TTTTCTTCAG CCCGCACCCG
     421  CGTGCGAAGA AGAGCACCAT CGCAGCGGCG AAGCTGATTT TGGATGCCGC GGTTAAGGCC
     481  GGTGCACCGA AAAACATTAT TGGCTGGATT GACGAGCCGA GCATTGAGCT GTCTCAGGAC
     541  CTGATGAGCG AAGCGGACAT TATTCTGGCC ACCGGTGGTC CGTCTATGGT TAAGGCAGCG
     601  TACTCCAGCG GTAAGCCTGC TATCGGCGTT GGTGCGGGCA ATACGCCGGC TATCATCGAT
     661  GAAAGCGCGG ATATTGATAT GGCAGTTAGC TCCATCATTC TGTCCAAAAC CTACGACAAT
     721  GGTGTTATCT GTGCAAGCGA GCAAAGCATC CTGGTGATGA ATAGCATTTA CGAGAAGGTC
     781  AAAGAAGAGT TTGTTAAGCG TGGTTCCTAT ATTCTGAACC AGGAGGAGAT TGCAAAAATC
     841  AAAGAAACGA TGTTTAAGAA TGGCGCGATT AATGCAGATA TTGTTGGTAA GTCCGCATAC
     901  ATTATCGCGA AGATGGCGGG TATCGAAGTG CCGCAAACCA CCAAAATCTT GATCGGCGAA
     961  GTGCAGAGCG TGGAGAAGAG CGAACTGTTC AGCCATGAGA AACTGAGCCC GGTTCTGGCT
    1021  ATGTACAAAG TGAAGGACTT TGACGAAGCC CTGAAGAAAG CTCAGCGTCT GATTGAGCTG
    1081  GGCGGCAGCG GTCACACGAG CAGCCTGTAT ATCGATAGCC AAAACAATAA GGACAAGGTG
    1141  AAAGAGTTCG GTCTGGCGAT GAAAACCTCT CGCACGTTTA TCAACATGCC GAGCAGCCAA
    1201  GGTGCGAGCG GCGATCTGTA TAACTTCGCA ATCGCGCCTT CTTTTACGCT GGGTTGCGGT
    1261  ACCTGGGGCG GTAATTCGGT GAGCCAGAAT GTCGAGCCGA AGCACCTGCT GAACATCAAA
    1321  TCGGTCGCGG AGCGCCGTGA GAAGAGACCG TGGTTCAAAG TCCCACAGAA AATTTACTTC
    1381  AAATACGGCT GCCTGCGTTT CGCGCTGAAA GAACTGAAAG ACATGAACAA AAAGCGTGCG
    1441  TTCATTGTTA CCGACAAAGA CCTGTTCAAA CTGGGTTACG TGAACAAAAT CACCAAAGTT
    1501  CTGGATGAAA TTGACATCAA GTACTCCATC TTCACTGATA TCAAATCCGA CCCAACGATT
    1561  GATAGCGTGA AAAAGGGCGC TAAAGAAATG CTGAACTTTG AACCGGACAC CATCATCAGC
    1621  ATCGGTGGTG GCTCTCCTAT GGATGCTGCG AAGGTCATGC ACCTGCTGTA CGAATACCCG
    1681  GAAGCGGAAA TCGAAAACCT GGCTATCAAC TTCATGGACA TCCGCAAACG TATCTGCAAC
    1741  TTCCCGAAGC TGGGCACTAA AGCTATTTCC GTTGCCATCC CGACTACCGC GGGCACTGGT
    1801  TCCGAAGCCA CGCCGTTCGC CGTGATCACC AACGATGAAA CCGGTATGAA ATACCCGCTG
    1861  ACCTCTTACG AACTGACCCC GAACATGGCA ATTATCGACA CCGAGCTGAT GCTGAACATG
    1921  CCGCGCAAGC TGACCGCTGC TACCGGCATC GACGCTCTGG TACATGCTAT TGAGGCGTAC
    1981  GTTTCCGTGA TGGCTACCGA TTACACCGAC GAACTGGCCC TGCGTGCGAT CAAAATGATT
    2041  TTCAAGTACC TGCCTCGCGC TTACAAAAAC GGCACGAATG ACATCGAGGC GCGTGAGAAA
    2101  ATGGCCCATG CAAGCAACAT CGCGGGCATG GCCTTCGCCA ACGCGTTCCT GGGCGTGTGC
    2161  CACTCTATGG CTCACAAACT GGGTGCTATG CACCACGTGC CGCACGGTAT CGCGTGTGCT
    2221  GTCCTGATCG AAGAAGTAAT TAAGTACAAC GCTACTGATT GCCCGACTAA ACAGACCGCC
    2281  TTCCCACAGT ACAAATCTCC TAACGCTAAA CGTAAGTACG CTGAGATCGC CGAATACCTG
    2341  AACCTGAAGG GTACGAGCGA CACTGAGAAA GTTACTGCGC TGATCGAAGC TATCTCTAAA
    2401  CTGAAAATTG ACCTGTCCAT CCCGCAGAAC ATCAGCGCCG CAGGCATCAA CAAAAAGGAC
    2461  TTTTACAACA CGCTGGACAA AATGAGCGAA CTGGCTTTTG ACGACCAGTG CACCACTGCA
    2521  AACCCGCGTT ACCCGCTGAT CTCTGAGCTG AAAGACATTT ATATCAAATC TTTTTAA
```

**AdhE2.32 (76961)**

```
       1  ATGAAAGTGA CGAACCAAAA AGAACTGAAA CAAAAACTGA ATGAACTGCG TGAGGCCCAG
      61  AAAAAGTTCG CTACGATGAC CCAGGAGCAA GTTGACAAGA TCTTCAAACA GTGTGCCATT
```

121 GCGGCTGCCA AAGAGCGTAT TAACTTGGCC AAACTGGCGG TGGAAGAAAC CGGCATCGGT

181 CTGGTCGAGG ACAAGATTAT CAAGAATCAC TTTGCCGCGG AGTATATCTA CAACAAATAC

```
     241  AAAAACGAAA AGACTTGCGG CATCATTGAT CATGATGACA GCTTGGGTAT CACGAAAGTC
     301  GCAGAACCAA TCGGTATTGT CGCGGCAATT GTCCCGACCA CTAATCCGAC CAGCACCGCG
     361  ATCTTCAAGA GCTTGATCAG CCTGAAAACC CGCAACGCGA TTTTCTTCAG CCCGCACCCG
     421  CGTGCGAAGA AGAGCACCAT CGCAGCGGCG AAGCTGATTT TGGATGCCGC GGTTAAGGCC
     481  GGTGCACCGA AAAACATTAT TGGCTGGATT GACGAGCCGA GCATTGAGCT GTCTCAGGAC
     541  ATGATGAGCG AAGCGGACAT TATTCTGGCC ACCGGTGGTC CGTCTATGGT TAAGGCAGCG
     601  TACTCCAGCG GTAAGCCTGC TATCGGCGTT GGTGCGGGCA ATACGCCGGC TATCATCGAT
     661  GAAAGCGCGG ATATTGATAT GGCAGTTAGC TCCATCATTC TGTCCAAAAC CTACGACAAT
     721  GGTGTTATCT GTGCAAGCGA GCAAAGCATC CTGGTGATGA ATAGCATTTA CGAGAAGGTC
     781  AAAGAAGAGT TTGTTAAGCG TGGTTCCTAT ATTCTGAACC AGAACGAGAT TGCAAAAATC
     841  AAAGAAACGA TGTTTAAGAA TGGCGCGATT AATGCAGATA TTGTTGGTAA GTCCGCATAC
     901  ATTATCGCGA AGATGGCGGG TATCGAAGTG CCGCAAACCA CCAAAATCTT GATCGGCGAA
     961  GTGCAGAGCG TGGAGAAGAG CGAACTGTTC AGCCATGAGA AACTGAGCCC GGTTCTGGCT
    1021  ATGTACAAAG TGAAGGACTT TGACGAAGCC CTGAAGAAAG CTCAGCGTCT GATTGAGCTG
    1081  GGCGGCAGCG GTCACACGAG CAGCCTGTAT AGCGATAGCC AAAACAATAA GGACAAGGTG
    1141  AAAGAGTTCG GTCTGGCGAT GAAAACCTCT CGCACGTTTA TCAACATGCC GAGCAGCCAA
    1201  GGTGCGAGCG GCGATCTGTA TAACTTCGCA ATCGCGCCTT CTTTTACGCT GGGTTGCGGT
    1261  ACCTGGGGCG GTAATTCGGT GAGCCAGAAT GTCGAGCCGA AGCACCTGCT GAACATCAAA
    1321  TCGGTCGCGG AGCGCCGTGA GAAGAGACCG TGGTTCAAAG TCCCACAGAA AATTTACTTC
    1381  AAATACGGCT GCCTGCGTTT CGCGCTGAAA GAACTGAAAG ACATGAACAA AAAGCGTGCG
    1441  TTCATTGTTA CCGACAAAGA CCTGTTCAAA CTGGGTTACG TGAACAAAAT CACCAAAGTT
    1501  CTGGATGAAA TTGACATCAA GTACTCCATC TTCACTGATA TCAAATCCGA CCCAACGATT
    1561  GATAGCGTGA AAAAGGGCGC TAAAGAAATG CTGAACTTTG AACCGGACAC CATCATCAGC
    1621  ATCGGTGGTG GCTCTCCTAT GGATGCTGCG AAGGTCATGC ACCTGCTGTA CGAATACCCG
    1681  GAAGCGGAAA TCGAAAACCT GGCTATCAAC TTCATGGACA TCCGCAAACG TATCTGCAAC
    1741  TTCCCGAAGC TGGGCACTAA AGCTATTTCC GTTGCCATCC CGACTACCGC GGGCACTGGT
    1801  TCCGAAGCCA CGCCGTTCGC CGTGATCACC AACGATGAAA CCGGTATGAA ATACCCGCTG
    1861  ACCTCTTACG AACTGACCCC GAACATGGCA ATTATCGACA CCGAGCTGAT GCTGAACATG
    1921  CCGCGCAAGC TGACCGCTGC TACCGGCATC GACGCTCTGG TACATGCTAT TGAGGCGTAC
    1981  GTTTCCGTGA TGGCTACCGA TTACACCGAC GAACTGGCCC TGCGTGCGAT CAAAATGATT
    2041  TTCAAGTACC TGCCTCGCGC TTACAAAAAC GGCACGAATG ACATCGAGGC GCGTGAGAAA
    2101  ATGGCCCATG CAAGCAACAT CGCGGGCATG GCCTTCGCCA ACGCGTTCCT GGGCGTGTGC
    2161  CACTCTATGG CTCACAAACT GGGTGCTATG CACCACGTGC CGCACGGTAT CGCGTGTGCT
    2221  GTCCTGATCG AAGAAGTAAT TAAGTACAAC GCTACTGATT GCCCGACTAA ACAGACCGCC
    2281  TTCCCACAGT ACAAATCTCC TAACGCTAAA CGTAAGTACG CTGAGATCGC CGAATACCTG
    2341  AACCTGAAGG GTACGAGCGA CACTGAGAAA GTTACTGCGC TGATCGAAGC TATCTCTAAA
    2401  CTGAAAATTG ACCTGTCCAT CCCGCAGAAC ATCAGCGCCG CAGGCATCAA CAAAAAGGAC
    2461  TTTTACAACA CGCTGGACAA AATGAGCGAA CTGGCTTTTG ACGACCAGTG CACCACTGCA
    2521  AACCCGCGTT ACCCGCTGAT CTCTGAGCTG AAAGACATTT ATATCAAATC TTTTTAA
```

**AdhE2.33 (76962)**

```
       1  ATGAAAGTGA CGAACCAAAA AGAACTGAAA CAAAAACTGA ATGAACTGCG TGAGGCCCAG
      61  AAAAAGTTCG CTACGTATAC CCAGGAGCAA GTTGACAAGA TCTTCAAACA GTGTGCCATT
```

121 GCGGCTGCCA AAGAGCGTAT TAACTTGGCC AAACTGGCGG TGGAAGAAAC CGGCATCGGT

181 CTGGTCGAGG ACAAGATTAT CAAGAATCAC TTTGCCGCGG AGTATATCTA CAACAAATAC

```
     241  AAAAACGAAA AGACTTGCGG CATCATTGAT CATGATGACA GCTTGGGTAT CACGAAAGTC
     301  GCAGAACCAA TCGGTATTGT CGCGAGCATT GTCCCGACCA CTAATCCGAC CAGCACCGCG
     361  ATCTTCAAGA GCTTGATCAG CCTGAAAACC CGCAACGCGA TTTTCTTCAG CCCGCACCCG
     421  CGTGCGAAGA AGAGCACCAT CGCAGCGGCG AAGCTGATTT TGGATGCCGC GGTTAAGGCC
     481  GGTGCACCGA AAAACATTAT TGGCTGGATT GACGAGCCGA GCATTGAGCT GTCTCAGGAC
     541  CTGATGAGCG AAGCGGACAT TATTCTGGCC ACCGGTGGTC CGTCTATGGT TAAGGCAGCG
     601  TACTCCAGCG GTAAGCCTGC TATCGGCGTT GGTGCGGGCA ATACGCCGGC TATCATCGAT
     661  GAAAGCGCGG ATATTGATAT GGCAGTTAGC TCCATCATTC TGTCCAAAAC CTACGACAAT
     721  GGTGTTATCT GTGCAAGCGA GCAAAGCATC CTGGTGATGA ATAGCATTTA CGAGAAGGTC
     781  AAAGAAGAGT TTGTTAAGCG TGGTTCCTAT ATTCTGAACC AGAACGAGAT TGCAAAAATC
     841  AAAGAAACGA TGTTTAAGAA TGGCGCGATT AATGCAGATA TTGTTGGTAA GTCCGCATAC
     901  ATTATCGCGA AGATGGCGGG TATCGAAGTG CCGCAATTCA CCAAAATCTT GATCGGCGAA
     961  GTGCAGAGCG TGGAGAAGAG CGAACTGTTC AGCCATGAGA AACTGAGCCC GGTTCTGGCT
    1021  ATGTACAAAG TGAAGGACTT TGACGAAGCC CTGAAGAAAG CTCAGCGTCT GATTGAGCTG
    1081  GGCGGCAGCG GTCACACGAG CAGCCTGTAT ATCGATAGCC AAAACAATAA GGACAAGGTG
    1141  AAAGAGTTCG GTCTGCGTAT GAAAACCTCT CGCACGTTTA TCAACATGCC GAGCAGCCAA
    1201  GGTGCGAGCG GCGATCTGTA TAACTTCGCA ATCGCGCCTT CTTTTACGCT GGGTTGCGGT
    1261  ACCTGGGGCG GTAATTCGGT GAGCCAGAAT GTCGAGCCGA AGCACCTGCT GAACATCAAA
    1321  TCGGTCGCGG AGCGCCGTGA GAAGAGACCG TGGTTCAAAG TCCCACAGAA AATTTACTTC
    1381  AAATACGGCT GCCTGCGTTT CGCGCTGAAA GAACTGAAAG ACATGAACAA AAAGCGTGCG
    1441  TTCATTGTTA CCGACAAAGA CCTGTTCAAA CTGGGTTACG TGAACAAAAT CACCAAAGTT
    1501  CTGGATGAAA TTGACATCAA GTACTCCATC TTCACTGATA TCAAATCCGA CCCAACGATT
    1561  GATAGCGTGA AAAAGGGCGC TAAAGAAATG CTGAACTTTG AACCGGACAC CATCATCAGC
    1621  ATCGGTGGTG GCTCTCCTAT GGATGCTGCG AAGGTCATGC ACCTGCTGTA CGAATACCCG
    1681  GAAGCGGAAA TCGAAAACCT GGCTATCAAC TTCATGGACA TCCGCAAACG TATCTGCAAC
    1741  TTCCCGAAGC TGGGCACTAA AGCTATTTCC GTTGCCATCC CGACTACCGC GGGCACTGGT
    1801  TCCGAAGCCA CGCCGTTCGC CGTGATCACC AACGATGAAA CCGGTATGAA ATACCCGCTG
    1861  ACCTCTTACG AACTGACCCC GAACATGGCA ATTATCGACA CCGAGCTGAT GCTGAACATG
    1921  CCGCGCAAGC TGACCGCTGC TACCGGCATC GACGCTCTGG TACATGCTAT TGAGGCGTAC
    1981  GTTTCCGTGA TGGCTACCGA TTACACCGAC GAACTGGCCC TGCGTGCGAT CAAAATGATT
    2041  TTCAAGTACC TGCCTCGCGC TTACAAAAAC GGCACGAATG ACATCGAGGC GCGTGAGAAA
    2101  ATGGCCCATG CAAGCAACAT CGCGGGCATG GCCTTCGCCA ACGCGTTCCT GGGCGTGTGC
    2161  CACTCTATGG CTCACAAACT GGGTGCTATG CACCACGTGC CGCACGGTAT CGCGTGTGCT
    2221  GTCCTGATCG AAGAAGTAAT TAAGTACAAC GCTACTGATT GCCCGACTAA ACAGACCGCC
    2281  TTCCCACAGT ACAAATCTCC TAACGCTAAA CGTAAGTACG CTGAGATCGC CGAATACCTG
    2341  AACCTGAAGG GTACGAGCGA CACTGAGAAA GTTACTGCGC TGATCGAAGC TATCTCTAAA
    2401  CTGAAAATTG ACCTGTCCAT CCCGCAGAAC ATCAGCGCCG CAGGCATCAA CAAAAAGGAC
    2461  TTTTACAACA CGCTGGACAA AATGAGCGAA CTGGCTTTTG ACGACCAGTG CACCACTGCA
    2521  AACCCGCGTT ACCCGCTGAT CTCTGAGCTG AAAGACATTT ATATCAAATC TTTTTAA
```

**AdhE2.34 (76963)**

```
       1  ATGAAAGTGA CGAACCAAAA AGAACTGAAA CAAAAACTGA ATGAACTGCG TGAGGCCCAG
      61  AAAAAGTTCG CTACGTATAC CCAGGAGCAA GTTGACAAGA TCTTCAAACA GTGTGCCATT
```

121 GCGGCTGCCA AAGAGCGTAT TAACTTGGCC AAACTGGCGG TGGAAGAAAC CGGCATCGGT

181 CTGGTCGAGG ACAAGATTAT CAAGAATCAC TTTGCCGCGG AGTATATCTA CAACAAATAC

```
     241  AAAAACGAAA AGACTTGCGG CATCATTGAT CATGATGACA GCTTGGGTAT CACGAAAGTC
     301  GCAGAACCAA TCGGTATTGT CGCGGCAATT GTCCCGACCA CTAATCCGAC CAGCACCGCG
     361  ATCTTCAAGA GCTTGATCAG CCTGAAAACC CGCAACGCGA TTTTCTTCAG CCCGCACCCG
     421  CGTGCGAAGA AGAGCACCAT CGCAGCGGCG AAGCTGATTT TGGATGCCGC GGTTAAGGCC
     481  GGTGCACCGA AAAACATTAT TGGCTGGATT GACGAGCCGA GCATTGAGCT GTCTCAGGAC
     541  CTGATGAGCG AAGCGGACAT TATTCTGGCC ACCGGTGGTC CGTCTATGGT TAAGGCAGCG
     601  TACTCCAGCG GTAAGCCTGC TATCGGCGTT GGTGCGGGCA ATACGCCGGC TATCATCGAT
     661  GAAAGCGCGG ATATTGATAT GGCAGTTAGC TCCATCATTC TGTCCAAAAC CTACGACAAT
     721  GGTGTTATCT GTGCAAGCGA GCAAAGCATC CTGGTGATGA ATAGCATTTA CGAGAAGGTC
     781  AAAGAAGAGT TTGTTAAGCG TGGTTCCTAT ATTCTGAACC AGAACGAGAT TGCAAAAATC
     841  AAAGAAACGA TGTTTAAGAA TGGCGCGATT AATGCAGATA TTGTTGGTAA GTCCGCATAC
     901  ATTATCGCGA AGATGGCGGG TATCGAAGTG CCGCAAACCA CCAAAATCTT GATCGGCGAA
     961  GTGCAGAGCG TGGAGAAGAG CGAACTGTTC AGCGAGGAGA AACTGAGCCC GGTTCTGGCT
    1021  ATGTACAAAG TGAAGGACTT TGACGAAGCC CTGAAGAAAG CTCAGCGTCT GATTGAGCTG
    1081  GGCGGCAGCG GTCACACGAG CAGCCTGTAT ATCGATAGCC AAAACAATAA GGACAAGGTG
    1141  AAAGAGTTCG GTCTGGCGAT GAAAACCTCT ATCACGTTTA TCAACATGCC GAGCAGCCAA
    1201  GGTGCGAGCG GCGATCTGTA TAACTTCGCA ATCGCGCCTT CTTTTACGCT GGGTTGCGGT
    1261  ACCTGGGGCG GTAATTCGGT GAGCCAGAAT GTCGAGCCGA AGCACCTGCT GAACATCGTT
    1321  TCGGTCGCGG AGCGCCGTGA GAAGAGACCG TGGTTCAAAG TCCCACAGAA AATTTACTTC
    1381  AAATACGGCT GCCTGCGTTT CGCGCTGAAA GAACTGAAAG ACATGAACAA AAAGCGTGCG
    1441  TTCATTGTTA CCGACAAAGA CCTGTTCAAA CTGGGTTACG TGAACAAAAT CACCAAAGTT
    1501  CTGGATGAAA TTGACATCAA GTACTCCATC TTCACTGATA TCAAATCCGA CCCAACGATT
    1561  GATAGCGTGA AAAAGGGCGC TAAAGAAATG CTGAACTTTG AACCGGACAC CATCATCAGC
    1621  ATCGGTGGTG GCTCTCCTAT GGATGCTGCG AAGGTCATGC ACCTGCTGTA CGAATACCCG
    1681  GAAGCGGAAA TCGAAAACCT GGCTATCAAC TTCATGGACA TCCGCAAACG TATCTGCAAC
    1741  TTCCCGAAGC TGGGCACTAA AGCTATTTCC GTTGCCATCC CGACTACCGC GGGCACTGGT
    1801  TCCGAAGCCA CGCCGTTCGC CGTGATCACC AACGATGAAA CCGGTATGAA ATACCCGCTG
    1861  ACCTCTTACG AACTGACCCC GAACATGGCA ATTATCGACA CCGAGCTGAT GCTGAACATG
    1921  CCGCGCAAGC TGACCGCTGC TACCGGCATC GACGCTCTGG TACATGCTAT TGAGGCGTAC
    1981  GTTTCCGTGA TGGCTACCGA TTACACCGAC GAACTGGCCC TGCGTGCGAT CAAAATGATT
    2041  TTCAAGTACC TGCCTCGCGC TTACAAAAAC GGCACGAATG ACATCGAGGC GCGTGAGAAA
    2101  ATGGCCCATG CAAGCAACAT CGCGGGCATG GCCTTCGCCA ACGCGTTCCT GGGCGTGTGC
    2161  CACTCTATGG CTCACAAACT GGGTGCTATG CACCACGTGC CGCACGGTAT CGCGTGTGCT
    2221  GTCCTGATCG AAGAAGTAAT TAAGTACAAC GCTACTGATT GCCCGACTAA ACAGACCGCC
    2281  TTCCCACAGT ACAAATCTCC TAACGCTAAA CGTAAGTACG CTGAGATCGC CGAATACCTG
    2341  AACCTGAAGG GTACGAGCGA CACTGAGAAA GTTACTGCGC TGATCGAAGC TATCTCTAAA
    2401  CTGAAAATTG ACCTGTCCAT CCCGCAGAAC ATCAGCGCCG CAGGCATCAA CAAAAAGGAC
    2461  TTTTACAACA CGCTGGACAA AATGAGCGAA CTGGCTTTTG ACGACCAGTG CACCACTGCA
2521  AACCCGCGTT ACCCGCTGAT CTCTGAGCTG AAAGACATTT ATATCAAATC TTTTTAA
```

**AdhE2.35 (76964)**

```
       1  ATGAAAGTGA CGAACCAAAA AGAACTGAAA CAAAAACTGA ATGAACTGCG TGAGGCCCAG
      61  AAAAAGTTCG CTACGTATAC CCAGGAGCAA GTTGACAAGA TCTTCAAACA GTGTGCCATT
```

121 GCGGCTGCCA AAGAGCGTAT TAACTTGGCC AAACTGGCGG TGGAAGAAAC CGGCATCGGT

181 CTGGTCGAGG ACAAGATTAT CAAGAATCAC TTTGCCGCGG AGTATATCTA CAACAAATAC

```
     241  AAAAACGAAA AGACTTGCGG CATCATTGAT CATGATGACA GCTTGGGTAT CACGAAAGTC
     301  GCAGAACCAA TCGGTATTGT CGCGGCAATT GTCCCGACCA CTAATCCGAC CAGCACCGCG
     361  ATCTTCAAGA GCTTGATCAG CCTGAAAACC CGCAACGCGA TTTTCTTCAG CCCGCACCCG
     421  CGTGCGAAGA AGAGCACCAT CGCAGCGGCG AAGCTGATTT TGGATGCCGC GGTTAAGGCC
     481  GGTGCACCGA AAAACATTAT TGGCTGGATT GACGAGCCGA GCATTGAGCT GTCTCAGGAC
     541  CTGATGAAAG AAGCGGACAT TATTCTGGCC ACCGGTGGTC CGTCTATGGT TAAGGCAGCG
     601  TACTCCAGCG GTAAGAAAGC TATCGGCGTT GGTGCGGGCA ATACGCCGGC TATCATCGAT
     661  GAAAGCGCGG ATATTGATAT GGCAGTTAGC TCCATCATTC TGTCCAAAAC CTACGACAAT
     721  GGTGTTATCT GTGCAAGCGA GCAAAGCATC CTGGTGATGA ATAGCATTTA CGAGAAGGTC
     781  AAAGAAGAGT TTGTTAAGCG TGGTTCCTAT ATTCTGAACC AGAACGAGAT TGCAAAAATC
     841  AAAGAAACGA TGTTTAAGAA TGGCGCGATT AATGCAGATA TTGTTGGTAA GTCCGCATAC
     901  ATTATCGCGA AGATGGCGGG TATCGAAGTG CCGCAAACCA CCAAAATCTT GATCGGCGAA
     961  GTGCAGAGCG TGGAGAAGAG CGAACTGTTC AGCCATGAGA AACTGAGCCC GGTTCTGGCT
    1021  ATGTACAAAG TGAAGGACTT TGACGAAGCC CTGAAGAAAG CTCAGCGTCT GATTGAGCTG
    1081  GGCGGCAGCC ACCACACGAG CAGCCTGTAT ATCGATAGCC AAAACAATAA GGACAAGGTG
    1141  AAAGAGTTCG GTCTGGCGAT GAAAACCTCT CGCACGTTTA TCAACATGCC GAGCAGCCAA
    1201  GGTGCGAGCG GCGATCTGTA TAACTTCGCA ATCGCGCCTT CTTTTACGCT GGGTTGCGGT
    1261  ACCTGGGGCG GTAATTCGGT GAGCCAGAAT GTCGAGCCGA AGCACCTGCT GAACATCAAA
    1321  TCGGTCGCGG AGCGCCGTGA GAAGAGACCG TGGTTCAAAG TCCCACAGAA AATTTACTTC
    1381  AAATACGGCT GCCTGCGTTT CGCGCTGAAA GAACTGAAAG ACATGAACAA AAAGCGTGCG
    1441  TTCATTGTTA CCGACAAAGA CCTGTTCAAA CTGGGTTACG TGAACAAAAT CACCAAAGTT
    1501  CTGGATGAAA TTGACATCAA GTACTCCATC TTCACTGATA TCAAATCCGA CCCAACGATT
    1561  GATAGCGTGA AAAAGGGCGC TAAAGAAATG CTGAACTTTG AACCGGACAC CATCATCAGC
    1621  ATCGGTGGTG GCTCTCCTAT GGATGCTGCG AAGGTCATGC ACCTGCTGTA CGAATACCCG
    1681  GAAGCGGAAA TCGAAAACCT GGCTATCAAC TTCATGGACA TCCGCAAACG TATCTGCAAC
    1741  TTCCCGAAGC TGGGCACTAA AGCTATTTCC GTTGCCATCC CGACTACCGC GGGCACTGGT
    1801  TCCGAAGCCA CGCCGTTCGC CGTGATCACC AACGATGAAA CCGGTATGAA ATACCCGCTG
    1861  ACCTCTTACG AACTGACCCC GAACATGGCA ATTATCGACA CCGAGCTGAT GCTGAACATG
    1921  CCGCGCAAGC TGACCGCTGC TACCGGCATC GACGCTCTGG TACATGCTAT TGAGGCGTAC
    1981  GTTTCCGTGA TGGCTACCGA TTACACCGAC GAACTGGCCC TGCGTGCGAT CAAAATGATT
    2041  TTCAAGTACC TGCCTCGCGC TTACAAAAAC GGCACGAATG ACATCGAGGC GCGTGAGAAA
    2101  ATGGCCCATG CAAGCAACAT CGCGGGCATG GCCTTCGCCA ACGCGTTCCT GGGCGTGTGC
    2161  CACTCTATGG CTCACAAACT GGGTGCTATG CACCACGTGC CGCACGGTAT CGCGTGTGCT
    2221  GTCCTGATCG AAGAAGTAAT TAAGTACAAC GCTACTGATT GCCCGACTAA ACAGACCGCC
    2281  TTCCCACAGT ACAAATCTCC TAACGCTAAA CGTAAGTACG CTGAGATCGC CGAATACCTG
    2341  AACCTGAAGG GTACGAGCGA CACTGAGAAA GTTACTGCGC TGATCGAAGC TATCTCTAAA
    2401  CTGAAAATTG ACCTGTCCAT CCCGCAGAAC ATCAGCGCCG CAGGCATCAA CAAAAAGGAC
    2461  TTTTACAACA CGCTGGACAA AATGAGCGAA CTGGCTTTTG ACGACCAGTG CACCACTGCA
    2521  AACCCGCGTT ACCCGCTGAT CTCTGAGCTG AAAGACATTT ATATCAAATC TTTTTAA
```

**AdhE2.36 (76965)**

```
       1  ATGAAAGTGA CGAACCAAAA AGAACTGAAA CAAAAACTGA ATGAACTGCG TGAGGCCCAG
      61  AAAAAGTTCG CTACGTATAC CCAGGAGCAA GTTGACAAGA TCTTCAAACA GTGTGCCATT
```

121 GCGGCTGCCA AATACCGTAT TAACTTGGCC AAACTGGCGG TGGAAGAAAC CGGCATCGGT

181 CTGGTCGAGG ACAAGATTAT CAAGAATCAC TTTGCCGCGG AGTATATCTA CAACAAATAC

```
     241  AAAAACCACA AGACTTGCGG CATCATTGAT CATGATGACA GCTTGGGTAT CACGAAAGTC
     301  GCAGAACCAA TCGGTATTGT CGCGGCAATT GTCCCGACCA CTAATCCGAC CAGCACCGCG
     361  ATCTTCAAGA GCTTGATCAG CCTGAAAACC CGCAACGCGA TTTTCTTCAG CCCGCACCCG
     421  CGTGCGAAGA AGAGCACCAT CGCAGCGGCG AAGCTGATTT TGGATGCCGC GGTTAAGGCC
     481  GGTGCACCGA AAAACATTAT TGGCTGGATT GACGAGCCGA GCATTGAGCT GTCTCAGGAC
     541  CTGATGAAAG AAGCGGACAT TATTCTGGCC ACCGGTGGTC CGTCTATGGT TAAGGCAGCG
     601  TACTCCAGCG GTAAGCCTGC TATCGGCGTT GGTGCGGGCA ATACGCCGGC TATCATCGAT
     661  GAAAGCGCGG ATATTGATAT GGCAGTTAGC TCCATCATTC TGTCCAAAAC CTACGACAAT
     721  GGTGTTATCT GTGCAAGCGA GCAAAGCATC CTGGTGATGA ATAGCATTTA CGAGAAGGTC
     781  AAAGAAGAGT TTGTTAAGCG TGGTTCCTAT ATTCTGAACC AGAACGAGAT TGCAAAAATC
     841  AAAGAAACGA TGTTTAAGAA TGGCGCGATT AATGCAGATA TTGTTGGTAA GTCCGCATAC
     901  ATTATCGCGA AGATGGCGGG TATCGAAGTG CCGCAAACCA CCAAAATCTT GATCGGCGAA
     961  GTGCAGAGCG TGGAGAAGAG CGAACTGTTC AGCCATGAGA AACTGAGCCC GGTTCTGGCT
    1021  ATGTACAAAG TGAAGGACTT TGACGAAGCC CTGAAGAAAG CTCAGCGTCT GATTGAGCTG
    1081  GGCGGCAGCG GTCACACGAG CAGCCTGTAT ATCGATAGCC AAAACAATAA GGACAAGGTG
    1141  AAAGAGTTCG GTCTGGCGAT GAAAACCTCT CGCACGTTTA TCAACATGCC GAGCAGCCAA
    1201  GGTGCGAGCG GCGATCTGTA TAACTTCGCA ATCGCGCCTT CTTTTACGCT GGGTTGCGGT
    1261  ACCTGGGGCG GTAATTCGGT GAGCCAGAAT GTCGAGCCGA AGCACCTGCT GAACATCAAA
    1321  TCGGTCGCGG AGCGCCGTGA GAAGAGACCG TGGTTCAAAG TCCCACAGAA AATTTACTTC
    1381  AAATACGGCT GCCTGCGTTT CGCGCTGAAA GAACTGAAAG ACATGAACAA AAAGCGTGCG
    1441  TTCATTGTTA CCGACAAAGA CCTGTTCAAA CTGGGTTACG TGAACAAAAT CACCAAAGTT
    1501  CTGGATGAAA TTGACATCAA GTACTCCATC TTCACTGATA TCAAATCCGA CCCAACGATT
    1561  GATAGCGTGA AAAAGGGCGC TAAAGAAATG CTGAACTTTG AACCGGACAC CATCATCAGC
    1621  ATCGGTGGTG GCTCTCCTAT GGATGCTGCG AAGGTCATGC ACCTGCTGTA CGAATACCCG
    1681  GAAGCGGAAA TCGAAAACCT GGCTATCAAC TTCATGGACA TCCGCAAACG TATCTGCAAC
    1741  TTCCCGAAGC TGGGCACTAA AGCTATTTCC GTTGCCATCC CGACTACCGC GGGCACTGGT
    1801  TCCGAAGCCA CGCCGTTCGC CGTGATCACC AACGATGAAA CCGGTATGAA ATACCCGCTG
    1861  ACCTCTTACG AACTGACCCC GAACATGGCA ATTATCGACA CCGAGCTGAT GCTGAACATG
    1921  CCGCGCAAGC TGACCGCTGC TACCGGCATC GACGCTCTGG TACATGCTAT TGAGGCGTAC
    1981  GTTTCCGTGA TGGCTACCGA TTACACCGAC GAACTGGCCC TGCGTGCGAT CAAAATGATT
    2041  TTCAAGTACC TGCCTCGCGC TTACAAAAAC GGCACGAATG ACATCGAGGC GCGTGAGAAA
    2101  ATGGCCCATG CAAGCAACAT CGCGGGCATG GCCTTCGCCA ACGCGTTCCT GGGCGTGTGC
    2161  CACTCTATGG CTCACAAACT GGGTGCTATG CACCACGTGC CGCACGGTAT CGCGTGTGCT
    2221  GTCCTGATCG AAGAAGTAAT TAAGTACAAC GCTACTGATT GCCCGACTAA ACAGACCGCC
    2281  TTCCCACAGT ACAAATCTCC TAACGCTAAA CGTAAGTACG CTGAGATCGC CGAATACCTG
    2341  AACCTGAAGG GTACGAGCGA CACTGAGAAA GTTACTGCGC TGATCGAAGC TATCTCTAAA
    2401  CTGAAAATTG ACCTGTCCAT CCCGCAGAAC ATCAGCGCCG CAGGCATCAA CAAAAAGGAC
    2461  TTTTACAACA CGCTGGACAA AATGAGCGAA CTGGCTTTTG ACGACCAGTG CACCACTGCA
    2521  AACCCGCGTT ACCCGCTGAT CTCTGAGCTG AAAGACATTT ATATCAAATC TTTTTAA
```

**AdhE2.37 (76966)**

```
       1  ATGAAAGTGA CGAACCAAAA AGAACTGAAA CAAAAACTGA ATGAACTGCG TGAGGCCCAG
      61  AAAAAGTTCG CTACGTATAC CCAGGCGCAA GTTGACAAGA TCTTCAAACA GTGTGCCATT
```

121 GCGGCTGCCA AAGAGCGTAT TAACTTGGCC AAACTGGCGG TGGAAGAAAC CGGCATCGGT

181 CTGGTCGAGG ACAAGATTAT CAAGAATCAC TTTGCCGCGG AGTATATCTA CAACAAATAC

```
     241  AAAAACGAAA AGACTTGCGG CATCATTGAT CATGATGACA GCTTGGGTAT CACGAAAGTC
     301  GCAGAACCAA TCGGTATTGT CGCGGCAATT GTCCCGACCA CTAATCCGAC CAGCACCGCG
     361  ATCTTCAAGA GCTTGATCAG CCTGAAAACC CGCAACGCGA TTTTCTTCAG CCCGCACCCG
     421  CGTGCGAAGA AGAGCACCAT CGCAGCGGCG AAGCTGATTT TGGATGCCGC GGTTAAGGCC
     481  GGTGCACCGA AAAACATTAT TGGCTGGATT GACGAGCCGA GCATTGAGCT GTCTCAGGAC
     541  CTGATGAGCG AAGCGGACAT TATTCTGGCC ACCGGTGGTC CGTCTATGGT TAAGGCAGCG
     601  TACTCCAGCG GTAAGCCTGC TATCGGCGTT GGTGCGGGCA ATACGCCGGC TATCATCGAT
     661  GAAAGCGCGG ATATTGATAT GGCAGTTAGC TCCATCATTC TGTCCAAAAC CTACGACAAT
     721  GGTGTTATCT GTGCAAGCGA GCAAAGCATC CTGGTGATGA ATAGCATTTA CGAGAAGGTC
     781  AAAGAAGAGT TTGTTAAGCG TGGTTCCTAT ATTCTGAACC AGAACGAGAT TGCAAAAATC
     841  AAAGAAACGA TGTTTAAGAA TGGCGCGATT AATGCAGATA TTGTTGGTAA GTCCGCATAC
     901  ATTATCGCGA AGATGGCGGG TATCGAAGTG CCGCAAACCA CCAAAATCTT GATCGGCGAA
     961  GTGCAGGACG TGGAGAAGAG CGAACTGTTC AGCCATGAGA AACTGAGCCC GGTTCTGGCT
    1021  ATGTACAAAG TGAAGGACTT TGACGAAGCC CTGAAGAAAG CTCAGCGTCT GATTGAGCTG
    1081  GGCGGCAGCG GTCACACGAG CAGCCTGTAT ATCGATAGCC AAAACAATAA GGACAAGGTG
    1141  AAAGAGTTCG GTCTGGCGAT GAAAACCTCT CGCACGTTTA TCAACATGCC GAGCAGCCAA
    1201  GGTGCGAGCG GCGATCTGTA TAACTTCGCA ATCGCGCCTT CTTTTACGCT GGGTTGCCCG
    1261  ACCTGGGGCG GTAATTCGGT GAGCCAGAAT GTCGAGCCGA AGCACCTGCT GAACATCAAA
    1321  TCGGTCGCGG AGCGCCGTGA GAAGAGACCG TGGTTCAAAG TCCCACAGAA AATTTACTTC
    1381  AAATACGGCT GCCTGCGTTT CGCGCTGAAA GAACTGAAAG ACATGAACAA AAAGCGTGCG
    1441  TTCATTGTTA CCGACAAAGA CCTGTTCAAA CTGGGTTACG TGAACAAAAT CACCAAAGTT
    1501  CTGGATGAAA TTGACATCAA GTACTCCATC TTCACTGATA TCAAATCCGA CCCAACGATT
    1561  GATAGCGTGA AAAAGGGCGC TAAAGAAATG CTGAACTTTG AACCGGACAC CATCATCAGC
    1621  ATCGGTGGTG GCTCTCCTAT GGATGCTGCG AAGGTCATGC ACCTGCTGTA CGAATACCCG
    1681  GAAGCGGAAA TCGAAAACCT GGCTATCAAC TTCATGGACA TCCGCAAACG TATCTGCAAC
    1741  TTCCCGAAGC TGGGCACTAA AGCTATTTCC GTTGCCATCC CGACTACCGC GGGCACTGGT
    1801  TCCGAAGCCA CGCCGTTCGC CGTGATCACC AACGATGAAA CCGGTATGAA ATACCCGCTG
    1861  ACCTCTTACG AACTGACCCC GAACATGGCA ATTATCGACA CCGAGCTGAT GCTGAACATG
    1921  CCGCGCAAGC TGACCGCTGC TACCGGCATC GACGCTCTGG TACATGCTAT TGAGGCGTAC
    1981  GTTTCCGTGA TGGCTACCGA TTACACCGAC GAACTGGCCC TGCGTGCGAT CAAAATGATT
    2041  TTCAAGTACC TGCCTCGCGC TTACAAAAAC GGCACGAATG ACATCGAGGC GCGTGAGAAA
    2101  ATGGCCCATG CAAGCAACAT CGCGGGCATG GCCTTCGCCA ACGCGTTCCT GGGCGTGTGC
    2161  CACTCTATGG CTCACAAACT GGGTGCTATG CACCACGTGC CGCACGGTAT CGCGTGTGCT
    2221  GTCCTGATCG AAGAAGTAAT TAAGTACAAC GCTACTGATT GCCCGACTAA ACAGACCGCC
    2281  TTCCCACAGT ACAAATCTCC TAACGCTAAA CGTAAGTACG CTGAGATCGC CGAATACCTG
    2341  AACCTGAAGG GTACGAGCGA CACTGAGAAA GTTACTGCGC TGATCGAAGC TATCTCTAAA
    2401  CTGAAAATTG ACCTGTCCAT CCCGCAGAAC ATCAGCGCCG CAGGCATCAA CAAAAAGGAC
    2461  TTTTACAACA CGCTGGACAA AATGAGCGAA CTGGCTTTTG ACGACCAGTG CACCACTGCA
    2521  AACCCGCGTT ACCCGCTGAT CTCTGAGCTG AAAGACATTT ATATCAAATC TTTTTAA
```

**AdhE2.38 (76967)**

```
       1  ATGAAAGTGA CGAACCAAAA AGAACTGAAA CAAAAACTGA ATGAACTGCG TGAGGCCCAG
      61  AAAAAGTTCG CTACGATGAC CCAGGAGCAA GTTGACAAGA TCTTCAAACA GTGTGCCATT
```

121 GCGGCTGCCA AAGAGCGTAT TAACTTGGCC AAACTGGCGG TGGAAGAAAC CGGCATCGGT

181 CTGGTCGAGG ACAAGATTAT CAAGAATCAC TTTGCCGCGG AGTATATCTA CAACAAATAC

```
     241  AAAAACGAAA AGACTTGCGG CATCATTGAT CATGATGACA GCTTGGGTAT CACGAAAGTC
     301  GCAGAACCAA TCGGTATTGT CGCGGCAATT GTCCCGACCA CTAATCCGAC CAGCACCGCG
     361  ATCTTCAAGA GCTTGATCAG CCTGAAAACC CGCAACGCGA TTTTCTTCAG CCCGCACCCG
     421  CGTGCGAAGA AGAGCACCAT CGCAGCGGCG AAGCTGATTT TGGATGCCGC GGTTAAGGCC
     481  GGTGCACCGA AAAACATTAT TGGCTGGATT GACGAGCCGA GCATTGAGCT GTCTCAGGAC
     541  CTGATGAGCG AAGCGGACAT TATTCTGGCC ACCGGTGGTC CGTCTATGGT TAAGGCAGCG
     601  TACTCCAGCG GTAAGCCTGC TATCGGCGTT GGTGCGGGCA ATACGCCGGC TATCATCGAT
     661  GAAAGCGCGG ATATTGATAT GGCAGTTAGC TCCATCATTC TGTCCAAAAC CTACGACAAT
     721  GGTGTTATCT GTGCAAGCGA GCAAAGCATC CTGGTGATGA ATAGCATTTA CGAGAAGGTC
     781  AAAGAAGAGT TTGTTAAGCG TGGTTCCTAT ATTCTGAACC AGCAGGAGAT TGCAAAAATC
     841  AAAGAAACGA TGTTTAAGAA TGGCGCGATT AATGCAGATA TTGTTGGTAA GTCCGCATAC
     901  ATTATCGCGA AGATGGCGGG TATCGAAGTG CCGCAAACCA CCAAAATCTT GATCGGCGAA
     961  GTGCAGAGCG TGGAGAAGAG CGAACTGTTC AGCCATGAGA AACTGAGCCC GGTTCTGGCT
    1021  ATGTACAAAG TGAAGGACTT TGACGAAGCC CTGAAGAAAG CTCAGCGTCT GATTGAGCTG
    1081  GGCGGCAGCG GTCACACGAG CAGCCTGTAT ATCGATAGCC AAAACAATAA GGACAAGGTG
    1141  AAAGAGTTCG GTCTGGCGAT GAAAACCTCT CGCACGTTTA TCAACATGCC GAGCAGCCAA
    1201  GGTGCGAGCG GCGATCTGTA TAACTTCGCA ATCGCGCCTT CTTTTACGCT GGGTTGCGGT
    1261  ACCTGGGAGG GTAATTCGGT GAGCCAGAAT GTCGAGCCGA AGCACCTGCT GAACATCAAA
    1321  TCGGTCGCGG AGCGCCGTGA GAAGAGACCG TGGTTCAAAG TCCCACAGAA AATTTACTTC
    1381  AAATACGGCT GCCTGCGTTT CGCGCTGAAA GAACTGAAAG ACATGAACAA AAAGCGTGCG
    1441  TTCATTGTTA CCGACAAAGA CCTGTTCAAA CTGGGTTACG TGAACAAAAT CACCAAAGTT
    1501  CTGGATGAAA TTGACATCAA GTACTCCATC TTCACTGATA TCAAATCCGA CCCAACGATT
    1561  GATAGCGTGA AAAAGGGCGC TAAAGAAATG CTGAACTTTG AACCGGACAC CATCATCAGC
    1621  ATCGGTGGTG GCTCTCCTAT GGATGCTGCG AAGGTCATGC ACCTGCTGTA CGAATACCCG
    1681  GAAGCGGAAA TCGAAAACCT GGCTATCAAC TTCATGGACA TCCGCAAACG TATCTGCAAC
    1741  TTCCCGAAGC TGGGCACTAA AGCTATTTCC GTTGCCATCC CGACTACCGC GGGCACTGGT
    1801  TCCGAAGCCA CGCCGTTCGC CGTGATCACC AACGATGAAA CCGGTATGAA ATACCCGCTG
    1861  ACCTCTTACG AACTGACCCC GAACATGGCA ATTATCGACA CCGAGCTGAT GCTGAACATG
    1921  CCGCGCAAGC TGACCGCTGC TACCGGCATC GACGCTCTGG TACATGCTAT TGAGGCGTAC
    1981  GTTTCCGTGA TGGCTACCGA TTACACCGAC GAACTGGCCC TGCGTGCGAT CAAAATGATT
    2041  TTCAAGTACC TGCCTCGCGC TTACAAAAAC GGCACGAATG ACATCGAGGC GCGTGAGAAA
    2101  ATGGCCCATG CAAGCAACAT CGCGGGCATG GCCTTCGCCA ACGCGTTCCT GGGCGTGTGC
    2161  CACTCTATGG CTCACAAACT GGGTGCTATG CACCACGTGC CGCACGGTAT CGCGTGTGCT
    2221  GTCCTGATCG AAGAAGTAAT TAAGTACAAC GCTACTGATT GCCCGACTAA ACAGACCGCC
    2281  TTCCCACAGT ACAAATCTCC TAACGCTAAA CGTAAGTACG CTGAGATCGC CGAATACCTG
    2341  AACCTGAAGG GTACGAGCGA CACTGAGAAA GTTACTGCGC TGATCGAAGC TATCTCTAAA
    2401  CTGAAAATTG ACCTGTCCAT CCCGCAGAAC ATCAGCGCCG CAGGCATCAA CAAAAAGGAC
    2461  TTTTACAACA CGCTGGACAA AATGAGCGAA CTGGCTTTTG ACGACCAGTG CACCACTGCA
    2521  AACCCGCGTT ACCCGCTGAT CTCTGAGCTG AAAGACATTT ATATCAAATC TTTTTAA
```

**AdhE2.39 (76968)**

```
       1  ATGAAAGTGA CGAACCAAAA AGAACTGAAA CAAAAACTGA ATGAACTGCG TGAGGCCCAG
      61  AAAAAGTTCG CTACGTATAC CCAGGAGCAA GTTGACAAGA TCTTCAAACA GTGTGCCATT
```

121 GCGGCTGCCA AAGAGCGTAT TAACTTGGCC AAACTGGCGG TGGAAGAAAC CGGCATCGGT

181 CTGGTCGAGG ACAAGATTAT CAAGAATCAC TTTGCCGCGG AGTATATCTA CAACAAATAC

```
     241  AAAAACGAAA AGACTTGCGG CATCATTGAT CATGATGACA GCTTGGGTAT CACGAAAGTC
     301  GCAGAACCAA TCGGTATTGT CGCGGCAATT GTCCCGACCA CTAATCCGAC CAGCACCGCG
     361  ATCTTCAAGA GCTTGATCAG CCTGAAAACC CGCAACGCGA TTTTCTTCAG CCCGCACCCG
     421  CGTGCGAAGA AGAGCACCAT CGCAGCGGCG ACCCTGATTT TGGATGCCGC GGTTAAGGCC
     481  GGTGCACCGA AAAACATTAT TGGCTGGATT GACGAGCCGA GCATTGAGCT GTCTCAGGAC
     541  CTGATGAGCG AAGCGGACAT TATTCTGGCC ACCGGTGGTC CGTCTATGGT TAAGGCAGCG
     601  TACTCCAGCG GTAAGCCTGC TATCGGCGTT GGTGCGGGCA ATACGCCGGC TATCATCGAT
     661  GAAAGCGCGG ATATTGATAT GGCAGTTAGC TCCATCATTC TGTCCAAAAC CTACGACAAT
     721  GGTGTTATCT GTGCAAGCGA GCAAAGCATC CTGGTGATGA ATAGCATTTA CGAGAAGGTC
     781  AAAGAAGAGT TTGTTAAGCG TGGTTCCTAT ATTCTGAACC AGAACGAGAT TGCAAAAATC
     841  AAAGAAACGA TGTTTAAGAA TGGCGCGATT AATGCAGATA TTGTTGGTAA GTCCGCATAC
     901  ATTATCGCGA AGATGGCGGG TATCGAAGTG CCGCAAACCA CCAAAATCTT GATCGGCGAA
     961  GTGCAGAGCG TGGAGAAGAG CGAACTGTTC AGCCATGAGA AACTGAGCCC GGTTCTGGCT
    1021  ATGTACAAAG TGAAGGACTT TGACGAAGCC CTGAAGAAAG CTGAGCGTCT GATTGAGCTG
    1081  GGCGGCAGCG GTCACACGAG CAGCCTGTAT ATCGATAGCC AAAACAATAA GGACAAGGTG
    1141  AAAGAGTTCG GTCTGGCGAT GAAAACCTCT ATCACGTTTA TCAACATGCC GAGCAGCCAA
    1201  GGTGCGAGCG GCGATCTGTA TAACTTCGCA ATCGCGCCTT CTTTTACGCT GGGTTGCGGT
    1261  ACCTGGGGCG GTAATTCGGT GAGCCAGAAT GTCGAGCCGA AGCACCTGCT GAACATCAAA
    1321  TCGGTCGCGG AGCGCCGTGA GAAGAGACCG TGGTTCAAAG TCCCACAGAA AATTTACTTC
    1381  AAATACGGCT GCCTGCGTTT CGCGCTGAAA GAACTGAAAG ACATGAACAA AAAGCGTGCG
    1441  TTCATTGTTA CCGACAAAGA CCTGTTCAAA CTGGGTTACG TGAACAAAAT CACCAAAGTT
    1501  CTGGATGAAA TTGACATCAA GTACTCCATC TTCACTGATA TCAAATCCGA CCCAACGATT
    1561  GATAGCGTGA AAAAGGGCGC TAAAGAAATG CTGAACTTTG AACCGGACAC CATCATCAGC
    1621  ATCGGTGGTG GCTCTCCTAT GGATGCTGCG AAGGTCATGC ACCTGCTGTA CGAATACCCG
    1681  GAAGCGGAAA TCGAAAACCT GGCTATCAAC TTCATGGACA TCCGCAAACG TATCTGCAAC
    1741  TTCCCGAAGC TGGGCACTAA AGCTATTTCC GTTGCCATCC CGACTACCGC GGGCACTGGT
    1801  TCCGAAGCCA CGCCGTTCGC CGTGATCACC AACGATGAAA CCGGTATGAA ATACCCGCTG
    1861  ACCTCTTACG AACTGACCCC GAACATGGCA ATTATCGACA CCGAGCTGAT GCTGAACATG
    1921  CCGCGCAAGC TGACCGCTGC TACCGGCATC GACGCTCTGG TACATGCTAT TGAGGCGTAC
    1981  GTTTCCGTGA TGGCTACCGA TTACACCGAC GAACTGGCCC TGCGTGCGAT CAAAATGATT
    2041  TTCAAGTACC TGCCTCGCGC TTACAAAAAC GGCACGAATG ACATCGAGGC GCGTGAGAAA
    2101  ATGGCCCATG CAAGCAACAT CGCGGGCATG GCCTTCGCCA ACGCGTTCCT GGGCGTGTGC
    2161  CACTCTATGG CTCACAAACT GGGTGCTATG CACCACGTGC CGCACGGTAT CGCGTGTGCT
    2221  GTCCTGATCG AAGAAGTAAT TAAGTACAAC GCTACTGATT GCCCGACTAA ACAGACCGCC
    2281  TTCCCACAGT ACAAATCTCC TAACGCTAAA CGTAAGTACG CTGAGATCGC CGAATACCTG
    2341  AACCTGAAGG GTACGAGCGA CACTGAGAAA GTTACTGCGC TGATCGAAGC TATCTCTAAA
    2401  CTGAAAATTG ACCTGTCCAT CCCGCAGAAC ATCAGCGCCG CAGGCATCAA CAAAAAGGAC
    2461  TTTTACAACA CGCTGGACAA AATGAGCGAA CTGGCTTTTG ACGACCAGTG CACCACTGCA
    2521  AACCCGCGTT ACCCGCTGAT CTCTGAGCTG AAAGACATTT ATATCAAATC TTTTTAA
```

**AdhE2.40 (76969)**

```
       1  ATGAAAGTGA CGAACCAAAA AGAACTGAAA CAAAAACTGA ATGAACTGCG TGAGGCCCAG
      61  AAAAAGTTCG CTACGTATAC CCAGGCGCAA GTTGAGAAGA TCTTCAAACA GTGTGCCATT
```

121 GCGGCTGCCA AAGAGCGTAT TAACTTGGCC AAACTGGCGG TGGAAGAAAC CGGCATCGGT

181 CTGGTCGAGG ACAAGATTAT CAAGAATCAC TTTGCCGCGG AGTATATCTA CAACAAATAC

```
     241  AAAAACGAAA AGACTTGCGG CATCATTGAT CATGATGACA GCTTGGGTAT CACGAAAGTC
     301  GCAGAACCAA TCGGTATTGT CGCGGCAATT GTCCCGACCA CTAATCCGAC CAGCACCGCG
     361  ATCTTCAAGA GCTTGATCAG CCTGAAAACC CGCAACGCGA TTTTCTTCAG CCCGCACCCG
     421  CGTGCGAAGA AGAGCACCAT CGCAGCGGCG AAGCTGATTT TGGATGCCGC GGTTAAGGCC
     481  GGTGCACCGA AAAACATTAT TGGCTGGATT GACGAGCCGA GCATTGAGCT GTCTCAGGAC
     541  CTGATGAGCG AAGCGGACAT TATTCTGGCC ACCGGTGGTC CGTCTATGGT TAAGGCAGCG
     601  TACTCCAGCG GTAAGCCTGC TATCGGCGTT GGTGCGGGCA ATACGCCGGC TATCATCGAT
     661  GAAAGCGCGG ATATTGATAT GGCAGTTAGC TCCATCATTC TGTCCAAAAC CTACGACAAT
     721  GGTGTTATCT GTGCAAGCGA GCAAAGCATC CTGGTGATGA ATAGCATTTA CGAGAAGGTC
     781  AAAGAAGAGT TTGTTAAGCG TGGTTCCTAT ATTCTGAACC AGAACGAGAT TGCAAAAATC
     841  AAAGAAACGA TGTTTAAGAA TGGCGCGATT AATGCAGATA TTGTTGGTAA GTCCGCATAC
     901  ATTATCGCGA AGATGGCGGG TATCGAAGTG CCGCAAACCA CCAAAATCTT GATCGGCGAA
     961  GTGCAGAGCG TGGAGAAGAG CGAACTGTTC AGCCATGAGA AACTGAGCCC GGTTCTGGCT
    1021  ATGTACAAAG TGAAGGACTT TGACGAAGCC CTGAAGAAAG CTCAGCGTCT GATTGAGCTG
    1081  GGCGGCAGCG GTCACACGAG CAGCCTGTAT ATCGATAGCC AAAACAATAA GGACAAGGTG
    1141  AAAGAGTTCG GTCTGGCGAT GAAAACCTCT CGCACGTTTA TCAACATGCC GAGCAGCCAA
    1201  GGTGCGAGCG GCGATCTGTA TAACTTCGCA ATCGCGCCTT CTTTTACGCT GGCGTGCGGT
    1261  ACCTGGGGCG GTAATTCGGT GAGCCAGAAT GTCGAGCCGA AGCACCTGCT GAACATCAAA
    1321  TCGGTCGCGG AGCGCCGTGA GAAGAGACCG TGGTTCAAAG TCCCACAGAA AATTTACTTC
    1381  AAATACGGCT GCCTGCGTTT CGCGCTGAAA GAACTGAAAG ACATGAACAA AAAGCGTGCG
    1441  TTCATTGTTA CCGACAAAGA CCTGTTCAAA CTGGGTTACG TGAACAAAAT CACCAAAGTT
    1501  CTGGATGAAA TTGACATCAA GTACTCCATC TTCACTGATA TCAAATCCGA CCCAACGATT
    1561  GATAGCGTGA AAAAGGGCGC TAAAGAAATG CTGAACTTTG AACCGGACAC CATCATCAGC
    1621  ATCGGTGGTG GCTCTCCTAT GGATGCTGCG AAGGTCATGC ACCTGCTGTA CGAATACCCG
    1681  GAAGCGGAAA TCGAAAACCT GGCTATCAAC TTCATGGACA TCCGCAAACG TATCTGCAAC
    1741  TTCCCGAAGC TGGGCACTAA AGCTATTTCC GTTGCCATCC CGACTACCGC GGGCACTGGT
    1801  TCCGAAGCCA CGCCGTTCGC CGTGATCACC AACGATGAAA CCGGTATGAA ATACCCGCTG
    1861  ACCTCTTACG AACTGACCCC GAACATGGCA ATTATCGACA CCGAGCTGAT GCTGAACATG
    1921  CCGCGCAAGC TGACCGCTGC TACCGGCATC GACGCTCTGG TACATGCTAT TGAGGCGTAC
    1981  GTTTCCGTGA TGGCTACCGA TTACACCGAC GAACTGGCCC TGCGTGCGAT CAAAATGATT
    2041  TTCAAGTACC TGCCTCGCGC TTACAAAAAC GGCACGAATG ACATCGAGGC GCGTGAGAAA
    2101  ATGGCCCATG CAAGCAACAT CGCGGGCATG GCCTTCGCCA ACGCGTTCCT GGGCGTGTGC
    2161  CACTCTATGG CTCACAAACT GGGTGCTATG CACCACGTGC CGCACGGTAT CGCGTGTGCT
    2221  GTCCTGATCG AAGAAGTAAT TAAGTACAAC GCTACTGATT GCCCGACTAA ACAGACCGCC
    2281  TTCCCACAGT ACAAATCTCC TAACGCTAAA CGTAAGTACG CTGAGATCGC CGAATACCTG
    2341  AACCTGAAGG GTACGAGCGA CACTGAGAAA GTTACTGCGC TGATCGAAGC TATCTCTAAA
    2401  CTGAAAATTG ACCTGTCCAT CCCGCAGAAC ATCAGCGCCG CAGGCATCAA CAAAAAGGAC
    2461  TTTTACAACA CGCTGGACAA AATGAGCGAA CTGGCTTTTG ACGACCAGTG CACCACTGCA
    2521  AACCCGCGTT ACCCGCTGAT CTCTGAGCTG AAAGACATTT ATATCAAATC TTTTTAA
```

**AdhE2.41 (76970)**

```
       1  ATGAAAGTGA CGAACCAAAA AGAACTGAAA CAAAAACTGA ATGAACTGCG TGAGGCCCAG
      61  AAAAAGTTCG CTACGTATAC CCAGGAGCAA GTTGACAAGA TCTTCAAACA GTGTGCCATT
```

121 GCGGCTGCCA AAGAGCGTAT TAACTTGGCC GAGCTGGCGG TGGAAGAAAC CGGCATCGGT

181 CTGGTCGAGG ACAAGATTAT CAAGAATCAC TTTGCCGCGG AGTATATCTA CAACAAATAC

```
     241  AAAAACGAAA AGACTTGCGG CATCATTGAT CATGATGACA GCTTGGGTAT CACGAAAGTC
     301  GCAGAACCAA TCGGTATTGT CGCGAGCATT GTCCCGACCA CTAATCCGAC CAGCACCGCG
     361  ATCTTCAAGA GCTTGATCAG CCTGAAAACC CGCAACGCGA TTTTCTTCAG CCCGCACCCG
     421  CGTGCGAAGA AGAGCACCAT CGCAGCGGCG AAGCTGATTT TGGATGCCGC GGTTAAGGCC
     481  GGTGCACCGA AAAACATTAT TGGCTGGATT GACGAGCCGA GCATTGAGCT GTCTCAGGAC
     541  CTGATGAGCG AACCGGACAT TATTCTGGCC ACCGGTGGTC CGTCTATGGT TAAGGCAGCG
     601  TACTCCAGCG GTAAGCCTGC TATCGGCGTT GGTGCGGGCA ATACGCCGGC TATCATCGAT
     661  GAAAGCGCGG ATATTGATAT GGCAGTTAGC TCCATCATTC TGTCCAAAAC CTACGACAAT
     721  GGTGTTATCT GTGCAAGCGA GCAAAGCATC CTGGTGATGA ATAGCATTTA CGAGAAGGTC
     781  AAAGAAGAGT TTGTTAAGCG TGGTTCCTAT ATTCTGAACC AGAACGAGAT TGCAAAAATC
     841  AAAGAAACGA TGTTTAAGAA TGGCGCGATT AATGCAGATA TTGTTGGTAA GTCCGCATAC
     901  ATTATCGCGA AGATGGCGGG TATCGAAGTG CCGCAAACCA CCAAAATCTT GATCGGCGAA
     961  GTGCAGAGCG TGGAGAAGAG CGAACTGTTC AGCCATGAGA AACTGAGCCC GGTTCTGGCT
    1021  ATGTACAAAG TGAAGGACTT TGACGAAGCC CTGAAGAAAG CTCAGCGTCT GATTGAGCTG
    1081  GGCGGCAGCG GTCACACGAG CAGCCTGTAT ATCGATAGCC AAAACAATAA GGACAAGGTG
    1141  AAAGAGTTCG GTCTGGCGAT GAAAACCTCT CGCACGTTTA TCAACATGCC GAGCAGCCAA
    1201  GGTGCGAGCG GCGATCTGTA TAACTTCGCA ATCGCGCCTT CTTTTACGCT GGGTTGCGGT
    1261  ACCTGGGGCG GTAATTCGGT GAGCCAGAAT GTCGAGCCGA AGCACCTGCT GAACATCAAA
    1321  TCGGTCGCGG AGCGCCGTGA GAAGAGACCG TGGTTCAAAG TCCCACAGAA AATTTACTTC
    1381  AAATACGGCT GCCTGCGTTT CGCGCTGAAA GAACTGAAAG ACATGAACAA AAAGCGTGCG
    1441  TTCATTGTTA CCGACAAAGA CCTGTTCAAA CTGGGTTACG TGAACAAAAT CACCAAAGTT
    1501  CTGGATGAAA TTGACATCAA GTACTCCATC TTCACTGATA TCAAATCCGA CCCAACGATT
    1561  GATAGCGTGA AAAAGGGCGC TAAAGAAATG CTGAACTTTG AACCGGACAC CATCATCAGC
    1621  ATCGGTGGTG GCTCTCCTAT GGATGCTGCG AAGGTCATGC ACCTGCTGTA CGAATACCCG
    1681  GAAGCGGAAA TCGAAAACCT GGCTATCAAC TTCATGGACA TCCGCAAACG TATCTGCAAC
    1741  TTCCCGAAGC TGGGCACTAA AGCTATTTCC GTTGCCATCC CGACTACCGC GGGCACTGGT
    1801  TCCGAAGCCA CGCCGTTCGC CGTGATCACC AACGATGAAA CCGGTATGAA ATACCCGCTG
    1861  ACCTCTTACG AACTGACCCC GAACATGGCA ATTATCGACA CCGAGCTGAT GCTGAACATG
    1921  CCGCGCAAGC TGACCGCTGC TACCGGCATC GACGCTCTGG TACATGCTAT TGAGGCGTAC
    1981  GTTTCCGTGA TGGCTACCGA TTACACCGAC GAACTGGCCC TGCGTGCGAT CAAAATGATT
    2041  TTCAAGTACC TGCCTCGCGC TTACAAAAAC GGCACGAATG ACATCGAGGC GCGTGAGAAA
    2101  ATGGCCCATG CAAGCAACAT CGCGGGCATG GCCTTCGCCA ACGCGTTCCT GGGCGTGTGC
    2161  CACTCTATGG CTCACAAACT GGGTGCTATG CACCACGTGC CGCACGGTAT CGCGTGTGCT
    2221  GTCCTGATCG AAGAAGTAAT TAAGTACAAC GCTACTGATT GCCCGACTAA ACAGACCGCC
    2281  TTCCCACAGT ACAAATCTCC TAACGCTAAA CGTAAGTACG CTGAGATCGC CGAATACCTG
    2341  AACCTGAAGG GTACGAGCGA CACTGAGAAA GTTACTGCGC TGATCGAAGC TATCTCTAAA
    2401  CTGAAAATTG ACCTGTCCAT CCCGCAGAAC ATCAGCGCCG CAGGCATCAA CAAAAAGGAC
    2461  TTTTACAACA CGCTGGACAA AATGAGCGAA CTGGCTTTTG ACGACCAGTG CACCACTGCA
    2521  AACCCGCGTT ACCCGCTGAT CTCTGAGCTG AAAGACATTT ATATCAAATC TTTTTAA
```

**AdhE2.42 (76971)**

```
       1  ATGAAAGTGA CGAACCAAAA AGAACTGAAA CAAAAACTGA ATGAACTGCG TGAGGCCCAG
      61  AAAAAGTTCG CTACGTATAC CCAGGAGCAA GTTGACAAGA TCTTCAAACA GTGTGCCATT
```

121 GCGGCTGCCA AAGAGCGTAT TAACTTGGCC AAACTGGCGG TGGAAGAAAC CGGCATCGGT

181 CTGGTCGAGG ACAAGATTAT CAAGAATCAC TTTGCCGCGG AGTATATCTA CAACAAATAC

```
     241  AAAAACGAAA AGACTTGCGG CATCATTGAT CATGATGACA GCTTGGGTAT CACGAAAGTC
     301  GCAGAACCAA TCGGTATTGT CGCGGCAATT GTCCCGACCA CTAATCCGAC CAGCACCGCG
     361  ATCTTCAAGA GCTTGATCAG CCTGGCGACC CGCAACGCGA TTTTCTTCAG CCCGCACCCG
     421  CGTGCGAAGA AGAGCACCAT CGCAGCGGCG AAGCTGATTT TGGATGCCGC GGTTAAGGCC
     481  GGTGCACCGA AAAACATTAT TGGCTGGATT GACGAGCCGA GCATTGAGCT GTCTCAGGAC
     541  CTGATGAGCG AAGCGGACAT TATTCTGGCC ACCGGTGGTC CGTCTATGGT TAAGGCAGCG
     601  TACTCCAGCG GTAAGCCTGC TATCGGCGTT GGTGCGGGCA ATACGCCGGC TATCATCGAT
     661  GAAAGCGCGG ATATTGATAT GGCAGTTAGC TCCATCATTC TGTCCAAAAC CTACGACAAT
     721  AACGTTATCT GTGCAAGCGA GCAAAGCATC CTGGTGATGA ATAGCATTTA CGAGAAGGTC
     781  AAAGAAGAGT TTGTTAAGCG TGGTTCCTAT ATTCTGAACC AGAACGAGAT TGCAAAAATC
     841  AAAGAAACGA TGTTTAAGAA TGGCGCGATT AATGCAGATA TTGTTGGTAA GTCCGCATAC
     901  ATTATCGCGA AGATGGCGGG TATCGAAGTG CCGCAAACCA CCAAAATCTT GATCGGCGAA
     961  GTGCAGAGCG TGGAGAAGAG CGAACTGTTC AGCCATGAGA AACTGAGCCC GGTTCTGGCT
    1021  ATGTACAAAG TGAAGGACTT TGACGAAGCC CTGAAGAAAG CTCAGCGTCT GATTGAGCTG
    1081  GGCGGCAGCG GTCACACGAG CAGCCTGTAT ATCGATAGCC AAAACAATAA GGACAAGGTG
    1141  AAAGAGTTCG GTCTGCGTAT GAAAACCTCT CGCACGTTTA TCAACATGCC GAGCAGCCAA
    1201  GGTGCGAGCG GCGATCTGTA TAACTTCGCA ATCGCGCCTT CTTTTACGCT GGGTTGCGGT
    1261  ACCTGGGGCG GTAATTCGGT GAGCCAGAAT GTCGAGCCGA AGCACCTGCT GAACATCAAA
    1321  TCGGTCGCGG AGCGCCGTGA GAAGAGACCG TGGTTCAAAG TCCCACAGAA AATTTACTTC
    1381  AAATACGGCT GCCTGCGTTT CGCGCTGAAA GAACTGAAAG ACATGAACAA AAAGCGTGCG
    1441  TTCATTGTTA CCGACAAAGA CCTGTTCAAA CTGGGTTACG TGAACAAAAT CACCAAAGTT
    1501  CTGGATGAAA TTGACATCAA GTACTCCATC TTCACTGATA TCAAATCCGA CCCAACGATT
    1561  GATAGCGTGA AAAAGGGCGC TAAAGAAATG CTGAACTTTG AACCGGACAC CATCATCAGC
    1621  ATCGGTGGTG GCTCTCCTAT GGATGCTGCG AAGGTCATGC ACCTGCTGTA CGAATACCCG
    1681  GAAGCGGAAA TCGAAAACCT GGCTATCAAC TTCATGGACA TCCGCAAACG TATCTGCAAC
    1741  TTCCCGAAGC TGGGCACTAA AGCTATTTCC GTTGCCATCC CGACTACCGC GGGCACTGGT
    1801  TCCGAAGCCA CGCCGTTCGC CGTGATCACC AACGATGAAA CCGGTATGAA ATACCCGCTG
    1861  ACCTCTTACG AACTGACCCC GAACATGGCA ATTATCGACA CCGAGCTGAT GCTGAACATG
    1921  CCGCGCAAGC TGACCGCTGC TACCGGCATC GACGCTCTGG TACATGCTAT TGAGGCGTAC
    1981  GTTTCCGTGA TGGCTACCGA TTACACCGAC GAACTGGCCC TGCGTGCGAT CAAAATGATT
    2041  TTCAAGTACC TGCCTCGCGC TTACAAAAAC GGCACGAATG ACATCGAGGC GCGTGAGAAA
    2101  ATGGCCCATG CAAGCAACAT CGCGGGCATG GCCTTCGCCA ACGCGTTCCT GGGCGTGTGC
    2161  CACTCTATGG CTCACAAACT GGGTGCTATG CACCACGTGC CGCACGGTAT CGCGTGTGCT
    2221  GTCCTGATCG AAGAAGTAAT TAAGTACAAC GCTACTGATT GCCCGACTAA ACAGACCGCC
    2281  TTCCCACAGT ACAAATCTCC TAACGCTAAA CGTAAGTACG CTGAGATCGC CGAATACCTG
    2341  AACCTGAAGG GTACGAGCGA CACTGAGAAA GTTACTGCGC TGATCGAAGC TATCTCTAAA
    2401  CTGAAAATTG ACCTGTCCAT CCCGCAGAAC ATCAGCGCCG CAGGCATCAA CAAAAAGGAC
    2461  TTTTACAACA CGCTGGACAA AATGAGCGAA CTGGCTTTTG ACGACCAGTG CACCACTGCA
    2521  AACCCGCGTT ACCCGCTGAT CTCTGAGCTG AAAGACATTT ATATCAAATC TTTTTAA
```

**AdhE2.43 (76972)**

```
       1  ATGAAAGTGA CGAACCAAAA AGAACTGAAA CAAAAACTGA ATGAACTGCG TGAGGCCCAG
      61  AAAAAGTTCG CTACGTATAC CCAGGAGCAA GTTGACAAGA TCTTCAAACA GTGTGCCATT
```

121 GCGGCTGCCA AAGAGCGTAT TAACTTGGCC CGTCTGGCGG TGGAAGAAAC CGGCATCGGT

181 CTGGTCGAGG ACAAGATTAT CAAGAATCAC TTTGCCGCGG AGTATATCTA CAACAAATAC

```
     241  AAAAACGAAA AGACTTGCGG CATCATTGAT CATGATGACA GCTTGGGTAT CACGAAAGTC
     301  GCAGAACCAA TCGGTATTGT CGCGGCAATT GTCCCGACCA CTAATCCGAC CAGCACCGCG
     361  ATCTTCAAGA GCTTGATCAG CCTGAAAACC CGCAACGCGA TTTTCTTCAG CCCGCACCCG
     421  CGTGCGAAGA AGAGCACCAT CGCAGCGGCG AAGCTGATTT TGGATGCCGC GGTTAAGGCC
     481  GGTGCACCGA AAAACATTAT TGGCTGGATT GACGAGCCGA GCATTGAGCT GTCTCAGGAC
     541  CTGATGAGCG AAGCGGACAT TATTCTGGCC ACCGGTGGTC CGTCTATGGT TAAGGCAGCG
     601  TACTCCAGCG GTAAGCCTGC TATCGGCGTT GGTGCGGGCA ATACGCCGGC TATCATCGAT
     661  GAAAGCGCGG ATATTGATAT GGCAGTTAGC TCCATCATTC TGTCCAAAAC CTACGACAAT
     721  GGTGTTATCT GTGCAAGCGA GCAAAGCATC CTGGTGATGA ATAGCATTTA CGAGAAGGTC
     781  AAAAAAGAGT TTGTTAAGCG TGGTTCCTAT ATTCTGAACC AGAACGAGAT TGCAAAAATC
     841  AAAGAAACGA TGTTTAAGAA TGGCGCGATT AATGCAGATA TTGTTGGTAA GTCCGCATAC
     901  ATTATCGCGA AGATGGCGGG TATCGAAGTG CCGCAAACCA CCAAAATCTT GATCGGCGAA
     961  GTGCAGAGCG TGGAGAAGAG CGAACTGTTC AGCCATGAGA AACTGAGCCC GGTTCTGGCT
    1021  ATGTACAAAG TGAAGGACTT TGACGAAGCC CTGAAGAAAG CTCAGCGTCT GATTGAGCTG
    1081  GGCGGCAGCG GTCACACGAG CAGCCTGTAT ATCGATAGCC AAAACAATAA GGACAAGGTG
    1141  AAAGAGTTCG GTCTGGCGAT GAAAACCTCT CGCTTCTTTA TCAACATGCC GAGCAGCCAA
    1201  GGTGCGAGCG GCGATCTGTA TAACTTCGCA ATCGCGCCTT CTTTTACGCT GGGTTGCGGT
    1261  ACCTGGGGCG GTAATTCGGT GAGCCAGAAT GTCGAGCCGA AGCACCTGCT GAACATCAAA
    1321  TCGGTCGCGG AGCGCCGTGA GAAGAGACCG TGGTTCAAAG TCCCACAGAA AATTTACTTC
    1381  AAATACGGCT GCCTGCGTTT CGCGCTGAAA GAACTGAAAG ACATGAACAA AAAGCGTGCG
    1441  TTCATTGTTA CCGACAAAGA CCTGTTCAAA CTGGGTTACG TGAACAAAAT CACCAAAGTT
    1501  CTGGATGAAA TTGACATCAA GTACTCCATC TTCACTGATA TCAAATCCGA CCCAACGATT
    1561  GATAGCGTGA AAAAGGGCGC TAAAGAAATG CTGAACTTTG AACCGGACAC CATCATCAGC
    1621  ATCGGTGGTG GCTCTCCTAT GGATGCTGCG AAGGTCATGC ACCTGCTGTA CGAATACCCG
    1681  GAAGCGGAAA TCGAAAACCT GGCTATCAAC TTCATGGACA TCCGCAAACG TATCTGCAAC
    1741  TTCCCGAAGC TGGGCACTAA AGCTATTTCC GTTGCCATCC CGACTACCGC GGGCACTGGT
    1801  TCCGAAGCCA CGCCGTTCGC CGTGATCACC AACGATGAAA CCGGTATGAA ATACCCGCTG
    1861  ACCTCTTACG AACTGACCCC GAACATGGCA ATTATCGACA CCGAGCTGAT GCTGAACATG
    1921  CCGCGCAAGC TGACCGCTGC TACCGGCATC GACGCTCTGG TACATGCTAT TGAGGCGTAC
    1981  GTTTCCGTGA TGGCTACCGA TTACACCGAC GAACTGGCCC TGCGTGCGAT CAAAATGATT
    2041  TTCAAGTACC TGCCTCGCGC TTACAAAAAC GGCACGAATG ACATCGAGGC GCGTGAGAAA
    2101  ATGGCCCATG CAAGCAACAT CGCGGGCATG GCCTTCGCCA ACGCGTTCCT GGGCGTGTGC
    2161  CACTCTATGG CTCACAAACT GGGTGCTATG CACCACGTGC CGCACGGTAT CGCGTGTGCT
    2221  GTCCTGATCG AAGAAGTAAT TAAGTACAAC GCTACTGATT GCCCGACTAA ACAGACCGCC
    2281  TTCCCACAGT ACAAATCTCC TAACGCTAAA CGTAAGTACG CTGAGATCGC CGAATACCTG
    2341  AACCTGAAGG GTACGAGCGA CACTGAGAAA GTTACTGCGC TGATCGAAGC TATCTCTAAA
    2401  CTGAAAATTG ACCTGTCCAT CCCGCAGAAC ATCAGCGCCG CAGGCATCAA CAAAAAGGAC
    2461  TTTTACAACA CGCTGGACAA AATGAGCGAA CTGGCTTTTG ACGACCAGTG CACCACTGCA
    2521  AACCCGCGTT ACCCGCTGAT CTCTGAGCTG AAAGACATTT ATATCAAATC TTTTTAA
```

**AdhE2.44 (76973)**

```
       1  ATGAAAGTGA CGAACCAAAA AGAACTGAAA CAAAAACTGA ATGAACTGCG TGAGGCCCAG
      61  AAAAAGTTCA ACACGTATAC CCAGGAGCAA GTTGACAAGA TCTTCAAACA GTGTGCCATT
```

121 GCGGCTGCCA AAGAGCGTAT TAACTTGGCC AAACTGGCGG TGGAAGAAAC CGGCATCGGT

181 CTGGTCGAGG ACAAGATTAT CAAGAATCAC TTTGCCGCGG AGTATATCTA CAACAAATAC

```
     241  AAAAACGAAA AGACTTGCGG CATCATTGAT CATGATGACA GCTTGGGTAT CACGAAAGTC
     301  GCAGAACCAA TCGGTATTGT CGCGGCAATT GTCCCGACCA CTAATCCGAC CAGCACCGCG
     361  ATCTTCAAGA GCTTGATCAG CCTGAAAACC CGCAACGCGA TTTTCTTCAG CCCGCACCCG
     421  CGTGCGAAGA AGAGCACCAT CGCAGCGGCG AAGCTGATTT TGGATGCCGC GGTTAAGGCC
     481  GGTGCACCGA AAAACATTAT TGGCTGGATT GACGAGCCGA GCATTGAGCT GTCTCAGGAC
     541  CTGATGAGCG AAGCGGACAT TATTCTGGCC ACCGGTGGTC CGTCTATGGT TAAGGCAGCG
     601  TACTCCAGCG GTAAGCCTGC TATCGGCGTT GGTGCGGGCA ATACGCCGGC TATCATCGAT
     661  GAAAGCGCGG ATATTGATAT GGCAGTTAGC TCCATCATTC TGTCCAAAAC CTACGACAAT
     721  GGTGTTATCT GTGCAAGCGA GCAAAGCATC CTGGTGATGA ATAGCATTTA CGAGAAGGTC
     781  AAAGAAGAGT TTGTTAAGCG TGGTTCCTAT ATTCTGAACC AGAACGAGAT TGCAAAAATC
     841  AAAGAAACGA TGTTTAAGAA TGGCGCGATT AATGCAGATA TTGTTGGTAA GTCCGCATAC
     901  ATTATCGCGA AGATGGCGGG TATCGAAGTG CCGCAAACCA CCAAAATCTT GATCGGCGAA
     961  GTGCAGCACG TGGAGAAGAG CGAACTGTTC AGCCATGAGA AACTGAGCCC GGTTCTGGCT
    1021  ATGTACAAAG TGAAGGACTT TGACGAAGCC CTGAAGAAAG CTCAGCGTCT GATTGAGCTG
    1081  GGCGGCAGCG GTCACACGAG CAGCCTGTAT ATCGATAGCC AAAACAATAA GGACAAGGTG
    1141  AAAGAGTTCG GTCTGGCGAT GAAAACCTCT CGCACGTTTA TCAACATGCC GAGCAGCCAA
    1201  GGTGCGAGCG GCGATCTGTA TAACTTCGCA ATCGCGCCTT CTTTTACGCT GGGTTGCGGT
    1261  ACCTGGGGCG GTAATTCGGT GAGCCAGAAT GTCGAGCCGA AGCACCTGCT GAACATCAAA
    1321  TCGGTCGCGC GTCGCCGTGA GAAGAGACCG TGGTTCAAAG TCCCACAGAA AATTTACTTC
    1381  AAATACGGCT GCCTGCGTTT CGCGCTGAAA GAACTGAAAG ACATGAACAA AAAGCGTGCG
    1441  TTCATTGTTA CCGACAAAGA CCTGTTCAAA CTGGGTTACG TGAACAAAAT CACCAAAGTT
    1501  CTGGATGAAA TTGACATCAA GTACTCCATC TTCACTGATA TCAAATCCGA CCCAACGATT
    1561  GATAGCGTGA AAAAGGGCGC TAAAGAAATG CTGAACTTTG AACCGGACAC CATCATCAGC
    1621  ATCGGTGGTG GCTCTCCTAT GGATGCTGCG AAGGTCATGC ACCTGCTGTA CGAATACCCG
    1681  GAAGCGGAAA TCGAAAACCT GGCTATCAAC TTCATGGACA TCCGCAAACG TATCTGCAAC
    1741  TTCCCGAAGC TGGGCACTAA AGCTATTTCC GTTGCCATCC CGACTACCGC GGGCACTGGT
    1801  TCCGAAGCCA CGCCGTTCGC CGTGATCACC AACGATGAAA CCGGTATGAA ATACCCGCTG
    1861  ACCTCTTACG AACTGACCCC GAACATGGCA ATTATCGACA CCGAGCTGAT GCTGAACATG
    1921  CCGCGCAAGC TGACCGCTGC TACCGGCATC GACGCTCTGG TACATGCTAT TGAGGCGTAC
    1981  GTTTCCGTGA TGGCTACCGA TTACACCGAC GAACTGGCCC TGCGTGCGAT CAAAATGATT
    2041  TTCAAGTACC TGCCTCGCGC TTACAAAAAC GGCACGAATG ACATCGAGGC GCGTGAGAAA
    2101  ATGGCCCATG CAAGCAACAT CGCGGGCATG GCCTTCGCCA ACGCGTTCCT GGGCGTGTGC
    2161  CACTCTATGG CTCACAAACT GGGTGCTATG CACCACGTGC CGCACGGTAT CGCGTGTGCT
    2221  GTCCTGATCG AAGAAGTAAT TAAGTACAAC GCTACTGATT GCCCGACTAA ACAGACCGCC
    2281  TTCCCACAGT ACAAATCTCC TAACGCTAAA CGTAAGTACG CTGAGATCGC CGAATACCTG
    2341  AACCTGAAGG GTACGAGCGA CACTGAGAAA GTTACTGCGC TGATCGAAGC TATCTCTAAA
    2401  CTGAAAATTG ACCTGTCCAT CCCGCAGAAC ATCAGCGCCG CAGGCATCAA CAAAAAGGAC
    2461  TTTTACAACA CGCTGGACAA AATGAGCGAA CTGGCTTTTG ACGACCAGTG CACCACTGCA
    2521  AACCCGCGTT ACCCGCTGAT CTCTGAGCTG AAAGACATTT ATATCAAATC TTTTTAA
```

**AdhE2.45 (76974)**

```
       1  ATGAAAGTGA CGAACCAAAA AGAACTGAAA CAAAAAGACA ATGAACTGCG TGAGGCCCAG
      61  AAAAAGTTCG CTACGTATAC CCAGGAGCAA GTTGACAAGA TCTTCAAACA GTGTGCCATT
```

121 GCGGCTGCCA AAGAGCGTAT TAACTTGGCC AAACTGGCGG TGGAAGAAAC CGGCATCGGT

181 CTGGTCGAGG ACAAGATTAT CAAGAATCAC TTTGCCGCGG AGTATATCTA CAACAAATAC

```
     241  AAAAACGAAA AGACTTGCGG CATCATTGAT CATGATGACA GCTTGGGTAT CACGAAAGTC
     301  GCAGAACCAA TCGGTATTGT CGCGGCAATT GTCCCGACCA CTAATCCGAC CAGCACCGCG
     361  ATCTTCAAGA GCTTGATCAG CCTGAAAACC CGCAACGCGA TTTTCTTCAG CCCGCACCCG
     421  CGTGCGAAGA AGAGCACCAT CGCAGCGGCG AAGCTGATTT TGGATGCCGC GGTTAAGGCC
     481  GGTGCACCGA AAAACATTAT TGGCTGGATT GACGAGCCGA GCATTGAGCT GTCTCAGGAC
     541  ATGATGAGCG AAGCGGACAT TATTCTGGCC ACCGGTGGTC CGTCTATGGT TAAGGCAGCG
     601  TACTCCAGCG GTAAGCCTGC TATCGGCGTT GGTGCGGGCA ATACGCCGGC TATCATCGAT
     661  GAAAGCGCGG ATATTGATAT GGCAGTTAGC TCCATCATTC TGTCCAAAAC CTACGACAAT
     721  GGTGTTATCT GTGCAAGCGA GCAAAGCATC CTGGTGATGA ATAGCATTTA CGAGAAGGTC
     781  AAAGAAGAGT TTGTTAAGCG TGGTTCCTAT ATTCTGAACC AGAACGCGAT TGCAAAAATC
     841  AAAGAAACGA TGTTTAAGAA TGGCGCGATT AATGCAGATA TTGTTGGTAA GTCCGCATAC
     901  ATTATCGCGA AGATGGCGGG TATCGAAGTG CCGCAAACCA CCAAAATCTT GATCGGCGAA
     961  GTGCAGAGCG TGGAGAAGAG CGAACTGTTC AGCCATGAGA AACTGAGCCC GGTTCTGGCT
    1021  ATGTACAAAG TGAAGGACTT TGACGAAGCC CTGAAGAAAG CTCAGCGTCT GATTGAGCTG
    1081  GGCGGCAGCG GTCACACGAG CAGCCTGTAT ATCGATAGCC AAAACAATAA GGACAAGGTG
    1141  AAAGAGTTCG GTCTGGCGAT GAAAACCTCT CGCACGTTTA TCAACATGCC GAGCAGCCAA
    1201  GGTGCGAGCG GCGATCTGTA TAACTTCGCA ATCGCGCCTT CTTTTACGCT GGGTTGCGGT
    1261  ACCTGGGGCG GTAATTCGGT GAGCCAGAAT GTCGAGCCGA AGCACCTGCT GAACATCAAA
    1321  TCGGTCGCGG AGCGCCGTGA GAAGAGACCG TGGTTCAAAG TCCCACAGAA AATTTACTTC
    1381  AAATACGGCT GCCTGCGTTT CGCGCTGAAA GAACTGAAAG ACATGAACAA AAAGCGTGCG
    1441  TTCATTGTTA CCGACAAAGA CCTGTTCAAA CTGGGTTACG TGAACAAAAT CACCAAAGTT
    1501  CTGGATGAAA TTGACATCAA GTACTCCATC TTCACTGATA TCAAATCCGA CCCAACGATT
    1561  GATAGCGTGA AAAAGGGCGC TAAAGAAATG CTGAACTTTG AACCGGACAC CATCATCAGC
    1621  ATCGGTGGTG GCTCTCCTAT GGATGCTGCG AAGGTCATGC ACCTGCTGTA CGAATACCCG
    1681  GAAGCGGAAA TCGAAAACCT GGCTATCAAC TTCATGGACA TCCGCAAACG TATCTGCAAC
    1741  TTCCCGAAGC TGGGCACTAA AGCTATTTCC GTTGCCATCC CGACTACCGC GGGCACTGGT
    1801  TCCGAAGCCA CGCCGTTCGC CGTGATCACC AACGATGAAA CCGGTATGAA ATACCCGCTG
    1861  ACCTCTTACG AACTGACCCC GAACATGGCA ATTATCGACA CCGAGCTGAT GCTGAACATG
    1921  CCGCGCAAGC TGACCGCTGC TACCGGCATC GACGCTCTGG TACATGCTAT TGAGGCGTAC
    1981  GTTTCCGTGA TGGCTACCGA TTACACCGAC GAACTGGCCC TGCGTGCGAT CAAAATGATT
    2041  TTCAAGTACC TGCCTCGCGC TTACAAAAAC GGCACGAATG ACATCGAGGC GCGTGAGAAA
    2101  ATGGCCCATG CAAGCAACAT CGCGGGCATG GCCTTCGCCA ACGCGTTCCT GGGCGTGTGC
    2161  CACTCTATGG CTCACAAACT GGGTGCTATG CACCACGTGC CGCACGGTAT CGCGTGTGCT
    2221  GTCCTGATCG AAGAAGTAAT TAAGTACAAC GCTACTGATT GCCCGACTAA ACAGACCGCC
    2281  TTCCCACAGT ACAAATCTCC TAACGCTAAA CGTAAGTACG CTGAGATCGC CGAATACCTG
    2341  AACCTGAAGG GTACGAGCGA CACTGAGAAA GTTACTGCGC TGATCGAAGC TATCTCTAAA
    2401  CTGAAAATTG ACCTGTCCAT CCCGCAGAAC ATCAGCGCCG CAGGCATCAA CAAAAAGGAC
    2461  TTTTACAACA CGCTGGACAA AATGAGCGAA CTGGCTTTTG ACGACCAGTG CACCACTGCA
    2521  AACCCGCGTT ACCCGCTGAT CTCTGAGCTG AAAGACATTT ATATCAAATC TTTTTAA
```

**AdhE2.46 (76975)**

```
       1  ATGAAAGTGA CGAACCAAAA AGAACTGAAA CAAAAACTGA ATGAACTGCG TGAGGCCCAG
      61  AAAAAGTTCG CTACGTATAC CCAGGCGCAA GTTGACAAGA TCTTCAAACA GTGTGCCATT
```

121 GCGGCTGCCA AATACCGTAT TAACTTGGCC AAACTGGCGG TGGAAGAAAC CGGCATCGGT

181 CTGGTCGAGG ACAAGATTAT CAAGAATCAC TTTGCCGCGG AGTATATCTA CAACAAATAC

```
     241  AAAAACGAAA AGACTTGCGG CATCATTGAT CATGATGACA GCTTGGGTAT CACGAAAGTC
     301  GCAGAACCAA TCGGTATTGT CGCGGCAATT GTCCCGACCA CTAATCCGAC CAGCACCGCG
     361  ATCTTCAAGA GCTTGATCAG CCTGAAAACC CGCAACGCGA TTTTCTTCAG CCCGCACCCG
     421  CGTGCGAAGA AGAGCACCAT CGCAGCGGCG AAGCTGATTT TGGATGCCGC GGTTAAGGCC
     481  GGTGCACCGA AAAACATTAT TGGCTGGATT GACGAGCCGA GCATTGAGCT GTCTCAGGAC
     541  CTGATGAGCG AAGCGGACAT TATTCTGGCC ACCGGTGGTC CGTCTATGGT TAAGGCAGCG
     601  TACTCCAGCG GTAAGCCTGC TATCGGCGTT GGTGCGGGCA ATACGCCGGC TATCATCGAT
     661  GAAAGCGCGG ATATTGATAT GGCAGTTAGC TCCATCATTC TGTCCAAAAC CTACGACAAT
     721  GGTGTTATCT GTGCAAGCGA GCAAAGCATC CTGGTGATGA ATAGCATTTA CGAGAAGGTC
     781  AAAGAAGAGT TTGTTAAGCG TGGTTCCTAT ATTCTGAACC AGAACGAGAT TGCAAAAATC
     841  AAAGAAACGA TGTTTAAGAA TGGCGCGATT AATGCAGATA TTGTTGGTAA GTCCGCATAC
     901  ATTATCGCGA AGATGGCGGG TATCGAAGTG CCGCAAACCA CCAAAATCTT GATCGGCGAA
     961  GTGCAGAGCG TGGAGAAGAG CGAACTGTTC AGCCATGAGA AACTGAGCCC GGTTCTGGCT
    1021  ATGTACAAAG TGAAGGACTT TGACGAAGCC CTGAAGAAAG CTCAGCGTCT GATTCACCTG
    1081  GGCGGCAGCG GTCACACGAG CAGCCTGTAT ATCGATAGCC AAAACAATAA GGACAAGGTG
    1141  AAAGAGTTCG GTCTGGCGAT GAAAACCTCT CGCACGTTTA TCAACATGCC GAGCAGCCAA
    1201  GGTGCGAGCG GCGATCTGTA TAACTTCGCA ATCGCGCCTT CTTTTACGCT GGGTTGCGGT
    1261  ACCTGGGGCG GTAATTCGGT GAGCCAGAAT GTCGAGCCGA AGCACCTGCT GAACATCAAA
    1321  TCGGTCGCGG AGCGCCGTGA GAAGAGACCG TGGTTCAAAG TCCCACAGAA AATTTACTTC
    1381  AAATACGGCT GCCTGCGTTT CGCGCTGAAA GAACTGAAAG ACATGAACAA AAAGCGTGCG
    1441  TTCATTGTTA CCGACAAAGA CCTGTTCAAA CTGGGTTACG TGAACAAAAT CACCAAAGTT
    1501  CTGGATGAAA TTGACATCAA GTACTCCATC TTCACTGATA TCAAATCCGA CCCAACGATT
    1561  GATAGCGTGA AAAAGGGCGC TAAAGAAATG CTGAACTTTG AACCGGACAC CATCATCAGC
    1621  ATCGGTGGTG GCTCTCCTAT GGATGCTGCG AAGGTCATGC ACCTGCTGTA CGAATACCCG
    1681  GAAGCGGAAA TCGAAAACCT GGCTATCAAC TTCATGGACA TCCGCAAACG TATCTGCAAC
    1741  TTCCCGAAGC TGGGCACTAA AGCTATTTCC GTTGCCATCC CGACTACCGC GGGCACTGGT
    1801  TCCGAAGCCA CGCCGTTCGC CGTGATCACC AACGATGAAA CCGGTATGAA ATACCCGCTG
    1861  ACCTCTTACG AACTGACCCC GAACATGGCA ATTATCGACA CCGAGCTGAT GCTGAACATG
    1921  CCGCGCAAGC TGACCGCTGC TACCGGCATC GACGCTCTGG TACATGCTAT TGAGGCGTAC
    1981  GTTTCCGTGA TGGCTACCGA TTACACCGAC GAACTGGCCC TGCGTGCGAT CAAAATGATT
    2041  TTCAAGTACC TGCCTCGCGC TTACAAAAAC GGCACGAATG ACATCGAGGC GCGTGAGAAA
    2101  ATGGCCCATG CAAGCAACAT CGCGGGCATG GCCTTCGCCA ACGCGTTCCT GGGCGTGTGC
    2161  CACTCTATGG CTCACAAACT GGGTGCTATG CACCACGTGC CGCACGGTAT CGCGTGTGCT
    2221  GTCCTGATCG AAGAAGTAAT TAAGTACAAC GCTACTGATT GCCCGACTAA ACAGACCGCC
    2281  TTCCCACAGT ACAAATCTCC TAACGCTAAA CGTAAGTACG CTGAGATCGC CGAATACCTG
    2341  AACCTGAAGG GTACGAGCGA CACTGAGAAA GTTACTGCGC TGATCGAAGC TATCTCTAAA
    2401  CTGAAAATTG ACCTGTCCAT CCCGCAGAAC ATCAGCGCCG CAGGCATCAA CAAAAAGGAC
    2461  TTTTACAACA CGCTGGACAA AATGAGCGAA CTGGCTTTTG ACGACCAGTG CACCACTGCA
    2521  AACCCGCGTT ACCCGCTGAT CTCTGAGCTG AAAGACATTT ATATCAAATC TTTTTAA
```

**AdhE2.47 (76976)**

```
       1  ATGAAAGTGA CGAACCAAAA AGAACTGAAA CAAAAACTGA ATGAACTGCG TGAGGCCCAG
      61  AAAAAGTTCG CTACGTATAC CCAGGAGCAA GTTGACAAGA TCTTCAAACA GTGTGCCATT
```

121 GCGGCTGCCA AAGAGCGTAT TAACTTGGCC AAACTGGCGG TGGAAGAAAC CGGCATCGGT

181 CTGGTCGAGG ACAAGATTAT CAAGAATCAC TTTGCCGCGG AGTATATCTA CAACAAATAC

```
     241  AAAAACGGTA AGACTTGCGG CATCATTGAT CATGATGACA GCTTGGGTAT CACGAAAGTC
     301  GCAGAACCAA TCGGTATTGT CGCGGCAATT GTCCCGACCA CTAATCCGAC CAGCACCGCG
     361  ATCTTCAAGA GCTTGATCAG CCTGAAAACC CGCAACGCGA TTTTCTTCAG CCCGCACCCG
     421  CGTGCGAAGA AGAGCACCAT CGCAGCGGCG AAGCTGATTT TGGATGCCGC GGTTAAGGCC
     481  GGTGCACCGA AAAACATTAT TGGCTGGATT GACGAGCCGA GCATTGAGCT GTCTCAGGAC
     541  CTGATGAGCG AAGCGGACAT TATTCTGGCC ACCGGTGGTC CGTCTATGGT TAAGGCAGCG
     601  TACTCCAGCG GTAAGCCTGC TATCGGCGTT GGTGCGGGCA ATACGCCGGC TATCATCGAT
     661  GAAAGCGCGG ATATTGATAT GGCAGTTAGC TCCATCATTC TGTCCAAAAC CTACGACAAT
     721  GGTGTTATCT GTGCAAGCGA GCAAAGCATC CTGGTGATGA ATAGCATTTA CGAGAAGGTC
     781  AAAGCGGAGT TTGTTAAGCG TGGTTCCTAT ATTCTGAACC AGAACGAGAT TGCAAAAATC
     841  AAAGAAACGA TGTTTAAGAA TGGCGCGATT AATGCAGATA TTGTTGGTAA GTCCGCATAC
     901  ATTATCGCGA AGATGGCGGG TATCGAAGTG CCGCAAACCA CCAAAATCTT GATCGGCGAA
     961  GTGCAGAGCG TGGAGAAGAG CGAACTGTTC AGCCATGAGA AACTGAGCCC GGTTCTGGCT
    1021  ATGTACAAAG TGAAGGACTT TGACGAAGCC CTGGACAAAG CTCAGCGTCT GATTGAGCTG
    1081  GGCGGCAGCG GTCACACGAG CAGCCTGTAT ATCGATAGCC AAAACAATAA GGACAAGGTG
    1141  AAAGAGTTCG GTCTGGCGAT GAAAACCTCT CGCACGTTTA TCAACATGCC GAGCAGCCAA
    1201  GGTGCGAGCG GCGATCTGTA TAACTTCGCA ATCGCGCCTT CTTTTACGCT GGGTTGCGGT
    1261  ACCTGGGGCG GTAATTCGGT GAGCCAGAAT GTCGAGCCGA AGCACCTGCT GAACATCAAA
    1321  TCGGTCGCGG AGCGCCGTGA GAAGAGACCG TGGTTCAAAG TCCCACAGAA AATTTACTTC
    1381  AAATACGGCT GCCTGCGTTT CGCGCTGAAA GAACTGAAAG ACATGAACAA AAAGCGTGCG
    1441  TTCATTGTTA CCGACAAAGA CCTGTTCAAA CTGGGTTACG TGAACAAAAT CACCAAAGTT
    1501  CTGGATGAAA TTGACATCAA GTACTCCATC TTCACTGATA TCAAATCCGA CCCAACGATT
    1561  GATAGCGTGA AAAAGGGCGC TAAAGAAATG CTGAACTTTG AACCGGACAC CATCATCAGC
    1621  ATCGGTGGTG GCTCTCCTAT GGATGCTGCG AAGGTCATGC ACCTGCTGTA CGAATACCCG
    1681  GAAGCGGAAA TCGAAAACCT GGCTATCAAC TTCATGGACA TCCGCAAACG TATCTGCAAC
    1741  TTCCCGAAGC TGGGCACTAA AGCTATTTCC GTTGCCATCC CGACTACCGC GGGCACTGGT
    1801  TCCGAAGCCA CGCCGTTCGC CGTGATCACC AACGATGAAA CCGGTATGAA ATACCCGCTG
    1861  ACCTCTTACG AACTGACCCC GAACATGGCA ATTATCGACA CCGAGCTGAT GCTGAACATG
    1921  CCGCGCAAGC TGACCGCTGC TACCGGCATC GACGCTCTGG TACATGCTAT TGAGGCGTAC
    1981  GTTTCCGTGA TGGCTACCGA TTACACCGAC GAACTGGCCC TGCGTGCGAT CAAAATGATT
    2041  TTCAAGTACC TGCCTCGCGC TTACAAAAAC GGCACGAATG ACATCGAGGC GCGTGAGAAA
    2101  ATGGCCCATG CAAGCAACAT CGCGGGCATG GCCTTCGCCA ACGCGTTCCT GGGCGTGTGC
    2161  CACTCTATGG CTCACAAACT GGGTGCTATG CACCACGTGC CGCACGGTAT CGCGTGTGCT
    2221  GTCCTGATCG AAGAAGTAAT TAAGTACAAC GCTACTGATT GCCCGACTAA ACAGACCGCC
    2281  TTCCCACAGT ACAAATCTCC TAACGCTAAA CGTAAGTACG CTGAGATCGC CGAATACCTG
    2341  AACCTGAAGG GTACGAGCGA CACTGAGAAA GTTACTGCGC TGATCGAAGC TATCTCTAAA
    2401  CTGAAAATTG ACCTGTCCAT CCCGCAGAAC ATCAGCGCCG CAGGCATCAA CAAAAAGGAC
    2461  TTTTACAACA CGCTGGACAA AATGAGCGAA CTGGCTTTTG ACGACCAGTG CACCACTGCA
    2521  AACCCGCGTT ACCCGCTGAT CTCTGAGCTG AAAGACATTT ATATCAAATC TTTTTAA
```

**AdhE2.48 (76977)**

```
       1  ATGAAAGTGA CGAACCAAAA AGAACTGAAA CAAAAACTGA ATGAACTGCG TGAGGCCCAG
      61  AAAAAGTTCG CTACGTATAC CCAGGAGCAA GTTGACAAGA TCTTCAAACA GTGTGCCATT
```

121 GCGGCTGCCA AACTGCGTAT TAACTTGGCC GAGCTGGCGG TGGAAGAAAC CGGCATCGGT

181 CTGGTCGAGG ACAAGATTAT CAAGAATCAC TTTGCCGCGG AGTATATCTA CAACAAATAC

```
     241  AAAAACGAAA AGACTTGCGG CATCATTGAT CATGATGACA GCTTGGGTAT CACGAAAGTC
     301  GCAGAACCAA TCGGTATTGT CGCGGCAATT GTCCCGACCA CTAATCCGAC CAGCACCGCG
     361  ATCTTCAAGA GCTTGATCAG CCTGAAAACC CGCAACGCGA TTTTCTTCAG CCCGCACCCG
     421  CGTGCGAAGA AGAGCACCAT CGCAGCGGCG AAGCTGATTT TGGATGCCGC GGTTAAGGCC
     481  GGTGCACCGA AAAACATTAT TGGCTGGATT GACGAGCCGA GCATTGAGCT GTCTCAGGAC
     541  CTGATGAGCG AAGCGGACAT TATTCTGGCC ACCGGTGGTC CGTCTATGGT TAAGGCAGCG
     601  TACTCCAGCG GTAAGCCTGC TATCGGCGTT GGTGCGGGCA ATACGCCGGC TATCATCGAT
     661  GAAAGCGCGG ATATTGATAT GGCAGTTAGC TCCATCATTC TGTCCAAAAC CTACGACAAT
     721  GGTGTTATCT GTGCAAGCGA GCAAAGCATC CTGGTGATGA ATAGCATTTA CGAGAAGGTC
     781  AAAGAAGAGT TTGTTAAGCG TGGTTCCTAT ATTCTGAACC AGAACGAGAT TGCAAAAATC
     841  AAAGAAACGA TGTTTAAGAA TGGCGCGATT AATGCAGATA TTGTTGGTAA GTCCGCATAC
     901  ATTATCGCGA AGATGGCGGG TATCGAAGTG CCGCAAACCA CCAAAATCTT GATCGGCGAA
     961  GTGCAGAGCG TGGAGAAGAG CGAACTGTTC AGCGTTGAGA AACTGAGCCC GGTTCTGGCT
    1021  ATGTACAAAG TGAAGGACTT TGACGAAGCC CTGAAGAAAG CTCAGCGTCT GATTGAGCTG
    1081  GGCGGCAGCG GTCACACGAG CAGCCTGTAT ATCGATAGCC AAAACAATAA GGACAAGGTG
    1141  AAAGAGTTCG GTCTGGCGAT GAAAACCTCT CGCACGTTTA TCAACATGCC GAGCAGCCAA
    1201  GGTGCGAGCG GCGATCTGTA TAACTTCGCA ATCGCGCCTT CTTTTACGCT GGGTTGCGGT
    1261  ACCTGGGGCG GTAATTCGGT GAGCCAGAAT GTCGAGCCGA AGCACCTGCT GAACATCAAA
    1321  TCGGTCGCGG AGCGCCGTGA GAAGAGACCG TGGTTCAAAG TCCCACAGAA AATTTACTTC
    1381  AAATACGGCT GCCTGCGTTT CGCGCTGAAA GAACTGAAAG ACATGAACAA AAAGCGTGCG
    1441  TTCATTGTTA CCGACAAAGA CCTGTTCAAA CTGGGTTACG TGAACAAAAT CACCAAAGTT
    1501  CTGGATGAAA TTGACATCAA GTACTCCATC TTCACTGATA TCAAATCCGA CCCAACGATT
    1561  GATAGCGTGA AAAAGGGCGC TAAAGAAATG CTGAACTTTG AACCGGACAC CATCATCAGC
    1621  ATCGGTGGTG GCTCTCCTAT GGATGCTGCG AAGGTCATGC ACCTGCTGTA CGAATACCCG
    1681  GAAGCGGAAA TCGAAAACCT GGCTATCAAC TTCATGGACA TCCGCAAACG TATCTGCAAC
    1741  TTCCCGAAGC TGGGCACTAA AGCTATTTCC GTTGCCATCC CGACTACCGC GGGCACTGGT
    1801  TCCGAAGCCA CGCCGTTCGC CGTGATCACC AACGATGAAA CCGGTATGAA ATACCCGCTG
    1861  ACCTCTTACG AACTGACCCC GAACATGGCA ATTATCGACA CCGAGCTGAT GCTGAACATG
    1921  CCGCGCAAGC TGACCGCTGC TACCGGCATC GACGCTCTGG TACATGCTAT TGAGGCGTAC
    1981  GTTTCCGTGA TGGCTACCGA TTACACCGAC GAACTGGCCC TGCGTGCGAT CAAAATGATT
    2041  TTCAAGTACC TGCCTCGCGC TTACAAAAAC GGCACGAATG ACATCGAGGC GCGTGAGAAA
    2101  ATGGCCCATG CAAGCAACAT CGCGGGCATG GCCTTCGCCA ACGCGTTCCT GGGCGTGTGC
    2161  CACTCTATGG CTCACAAACT GGGTGCTATG CACCACGTGC CGCACGGTAT CGCGTGTGCT
    2221  GTCCTGATCG AAGAAGTAAT TAAGTACAAC GCTACTGATT GCCCGACTAA ACAGACCGCC
    2281  TTCCCACAGT ACAAATCTCC TAACGCTAAA CGTAAGTACG CTGAGATCGC CGAATACCTG
    2341  AACCTGAAGG GTACGAGCGA CACTGAGAAA GTTACTGCGC TGATCGAAGC TATCTCTAAA
    2401  CTGAAAATTG ACCTGTCCAT CCCGCAGAAC ATCAGCGCCG CAGGCATCAA CAAAAAGGAC
    2461  TTTTACAACA CGCTGGACAA AATGAGCGAA CTGGCTTTTG ACGACCAGTG CACCACTGCA
    2521  AACCCGCGTT ACCCGCTGAT CTCTGAGCTG AAAGACATTT ATATCAAATC TTTTTAA
```

**AdhE2.49 (76978)**

```
       1  ATGAAAGTGA CGAACCAAAA AGAACTGAAA CAAAAACTGA ATGAACTGCG TGAGGCCCAG
      61  AAAAAGTTCG CTACGTGCAC CCAGGAGCAA GTTGACAAGA TCTTCAAACA GTGTGCCATT
```

121 GCGGCTGCCA AAGAGCGTAT TAACTTGGCC AAACTGGCGG TGGAAGAAAC CGGCATCGGT

181 CTGGTCGAGG ACAAGATTAT CAAGAATCAC TTTGCCGCGG AGTATATCTA CAACAAATAC

```
     241  AAAAACGGTA AGACTTGCGG CATCATTGAT CATGATGACA GCTTGGGTAT CACGAAAGTC
     301  GCAGAACCAA TCGGTATTGT CGCGGCAATT GTCCCGACCA CTAATCCGAC CAGCACCGCG
     361  ATCTTCAAGA GCTTGATCAG CCTGAAAACC CGCAACGCGA TTTTCTTCAG CCCGCACCCG
     421  CGTGCGAAGA AGAGCACCAT CGCAGCGGCG AAGCTGATTT TGGATGCCGC GGTTAAGGCC
     481  GGTGCACCGA AAAACATTAT TGGCTGGATT GACGAGCCGA GCATTGAGCT GTCTCAGGAC
     541  CTGATGAGCG AAGCGGACAT TATTCTGGCC ACCGGTGGTC CGTCTATGGT TAAGGCAGCG
     601  TACTCCAGCG GTAAGCCTGC TATCGGCGTT GGTGCGGGCA ATACGCCGGC TATCATCGAT
     661  GAAAGCGCGG ATATTGATAT GGCAGTTAGC TCCATCATTC TGTCCAAAAC CTACGACAAT
     721  GGTGTTATCT GTGCAAGCGA GCAAAGCATC CTGGTGATGA ATAGCATTTA CGAGAAGGTC
     781  AAAGAAGAGT TTGTTAAGCG TGGTTCCTAT ATTCTGAACC AGAACGAGAT TGCAAAAATC
     841  AAAGAAACGA TGTTTAAGAA TGGCGCGATT AATGCAGATA TTGTTGGTAA GTCCGCATAC
     901  ATTATCGCGA AGATGGCGGG TATCGAAGTG CCGCAAACCA CCAAAATCTT GATCGGCGAA
     961  GTGCAGAGCG TGGAGGCGAG CGAACTGTTC AGCCATGAGA AACTGAGCCC GGTTCTGGCT
    1021  ATGTACAAAG TGAAGGACTT TGACGAAGCC CTGAAGAAAG CTCAGCGTCT GATTGAGCTG
    1081  GGCGGCAGCG GTCACACGAG CAGCCTGTAT ATCGATAGCC AAAACAATAA GGACAAGGTG
    1141  AAAGAGTTCG GTCTGGCGAT GAAAACCTCT CGCACGTTTA TCAACATGCC GAGCAGCCAA
    1201  GGTGCGAGCG GCGATCTGTA TAACTTCGCA ATCGCGCCTT CTTTTACGCT GGGTTGCGGT
    1261  ACCTGGGGCG GTAATTCGGT GAGCCAGAAT GTCGAGCCGA AGCACCTGCT GAACATCAAA
    1321  TCGGTCGCGG AGCGCCGTGA GAAGAGACCG TGGTTCAAAG TCCCACAGAA AATTTACTTC
    1381  AAATACGGCT GCCTGCGTTT CGCGCTGAAA GAACTGAAAG ACATGAACAA AAAGCGTGCG
    1441  TTCATTGTTA CCGACAAAGA CCTGTTCAAA CTGGGTTACG TGAACAAAAT CACCAAAGTT
    1501  CTGGATGAAA TTGACATCAA GTACTCCATC TTCACTGATA TCAAATCCGA CCCAACGATT
    1561  GATAGCGTGA AAAAGGGCGC TAAAGAAATG CTGAACTTTG AACCGGACAC CATCATCAGC
    1621  ATCGGTGGTG GCTCTCCTAT GGATGCTGCG AAGGTCATGC ACCTGCTGTA CGAATACCCG
    1681  GAAGCGGAAA TCGAAAACCT GGCTATCAAC TTCATGGACA TCCGCAAACG TATCTGCAAC
    1741  TTCCCGAAGC TGGGCACTAA AGCTATTTCC GTTGCCATCC CGACTACCGC GGGCACTGGT
    1801  TCCGAAGCCA CGCCGTTCGC CGTGATCACC AACGATGAAA CCGGTATGAA ATACCCGCTG
    1861  ACCTCTTACG AACTGACCCC GAACATGGCA ATTATCGACA CCGAGCTGAT GCTGAACATG
    1921  CCGCGCAAGC TGACCGCTGC TACCGGCATC GACGCTCTGG TACATGCTAT TGAGGCGTAC
    1981  GTTTCCGTGA TGGCTACCGA TTACACCGAC GAACTGGCCC TGCGTGCGAT CAAAATGATT
    2041  TTCAAGTACC TGCCTCGCGC TTACAAAAAC GGCACGAATG ACATCGAGGC GCGTGAGAAA
    2101  ATGGCCCATG CAAGCAACAT CGCGGGCATG GCCTTCGCCA ACGCGTTCCT GGGCGTGTGC
    2161  CACTCTATGG CTCACAAACT GGGTGCTATG CACCACGTGC CGCACGGTAT CGCGTGTGCT
    2221  GTCCTGATCG AAGAAGTAAT TAAGTACAAC GCTACTGATT GCCCGACTAA ACAGACCGCC
    2281  TTCCCACAGT ACAAATCTCC TAACGCTAAA CGTAAGTACG CTGAGATCGC CGAATACCTG
    2341  AACCTGAAGG GTACGAGCGA CACTGAGAAA GTTACTGCGC TGATCGAAGC TATCTCTAAA
    2401  CTGAAAATTG ACCTGTCCAT CCCGCAGAAC ATCAGCGCCG CAGGCATCAA CAAAAAGGAC
    2461  TTTTACAACA CGCTGGACAA AATGAGCGAA CTGGCTTTTG ACGACCAGTG CACCACTGCA
    2521  AACCCGCGTT ACCCGCTGAT CTCTGAGCTG AAAGACATTT ATATCAAATC TTTTTAA
```

**AdhE2.50 (76979)**

```
       1  ATGAAAGTGA CGAACCAAAA AGAACTGAAA CAAAAAGACA ATGAACTGCG TGAGGCCCAG
      61  AAAAAGTTCG CTACGTATAC CCAGGAGCAA GTTGACAAGA TCTTCAAACA GTGTGCCATT
```

121 GCGGCTGCCA AAGAGCGTAT TAACTTGGCC GAGCTGGCGG TGGAAGAAAC CGGCATCGGT

181 CTGGTCGAGG ACAAGATTAT CAAGAATCAC TTTGCCGCGG AGTATATCTA CAACAAATAC

```
     241  AAAAACGAAA AGACTTGCGG CATCATTGAT CATGATGACA GCTTGGGTAT CACGAAAGTC
     301  GCAGAACCAA TCGGTATTGT CGCGGCAATT GTCCCGACCA CTAATCCGAC CAGCACCGCG
     361  ATCTTCAAGA GCTTGATCAG CCTGAAAACC CGCAACGCGA TTTTCTTCAG CCCGCACCCG
     421  CGTGCGAAGA AGAGCACCAT CGCAGCGGCG AAGCTGATTT TGGATGCCGC GGTTAAGGCC
     481  GGTGCACCGA AAAACATTAT TGGCTGGATT GACGAGCCGA GCATTGAGCT GTCTCAGGAC
     541  CTGATGAGCG AAGCGGACAT TATTCTGGCC ACCGGTGGTC CGTCTATGGT TAAGGCAGCG
     601  TACTCCAGCG GTAAGCCTGC TATCGGCGTT GGTGCGGGCA ATACGCCGGC TATCATCGAT
     661  GAAAGCGCGG ATATTGATAT GGCAGTTAGC TCCATCATTC TGTCCAAAAC CTACGACAAT
     721  GGTGTTATCT GTGCAAGCGA GCAAAGCATC CTGGTGATGA ATAGCATTTA CGAGAAGGTC
     781  AAAGAAGAGT TTGTTAAGCG TGGTTCCTAT ATTCTGAACC AGAACGAGAT TGCAAAAATC
     841  AAAGAAACGA TGTTTAAGAA TGGCGCGATT AATGCAGATA TTGTTGGTAA GTCCGCATAC
     901  ATTATCGCGA AGATGGCGGG TATCGAAGTG CCGCAAACCA CCAAAATCTT GATCGGCGAA
     961  GTGCAGAGCG TGGAGAAGAG CGAACTGTTC AGCCATGAGA AACTGAGCCC GGTTCTGGCT
    1021  ATGTACAAAG TGAAGGACTT TGACGAAGCC CTGAAGAAAG CTGAGCGTCT GATTGAGCTG
    1081  GGCGGCAGCG GTCACACGAG CAGCCTGTAT ATCGATAGCC AAAACAATAA GGACAAGGTG
    1141  AAAGAGTTCG GTCTGGCGAT GAAAACCTCT CGCACGTTTA TCAACATGCC GAGCAGCCAA
    1201  GGTGCGAGCG GCGATCTGTA TAACTTCGCA ATCGCGCCTT CTTTTACGCT GGGTTGCGGT
    1261  ACCTGGGGCG GTAATTCGGT GAGCCAGAAT GTCGAGCCGA AGCACCTGCT GAACATCGTT
    1321  TCGGTCGCGG AGCGCCGTGA GAAGAGACCG TGGTTCAAAG TCCCACAGAA AATTTACTTC
    1381  AAATACGGCT GCCTGCGTTT CGCGCTGAAA GAACTGAAAG ACATGAACAA AAAGCGTGCG
    1441  TTCATTGTTA CCGACAAAGA CCTGTTCAAA CTGGGTTACG TGAACAAAAT CACCAAAGTT
    1501  CTGGATGAAA TTGACATCAA GTACTCCATC TTCACTGATA TCAAATCCGA CCCAACGATT
    1561  GATAGCGTGA AAAAGGGCGC TAAAGAAATG CTGAACTTTG AACCGGACAC CATCATCAGC
    1621  ATCGGTGGTG GCTCTCCTAT GGATGCTGCG AAGGTCATGC ACCTGCTGTA CGAATACCCG
    1681  GAAGCGGAAA TCGAAAACCT GGCTATCAAC TTCATGGACA TCCGCAAACG TATCTGCAAC
    1741  TTCCCGAAGC TGGGCACTAA AGCTATTTCC GTTGCCATCC CGACTACCGC GGGCACTGGT
    1801  TCCGAAGCCA CGCCGTTCGC CGTGATCACC AACGATGAAA CCGGTATGAA ATACCCGCTG
    1861  ACCTCTTACG AACTGACCCC GAACATGGCA ATTATCGACA CCGAGCTGAT GCTGAACATG
    1921  CCGCGCAAGC TGACCGCTGC TACCGGCATC GACGCTCTGG TACATGCTAT TGAGGCGTAC
    1981  GTTTCCGTGA TGGCTACCGA TTACACCGAC GAACTGGCCC TGCGTGCGAT CAAAATGATT
    2041  TTCAAGTACC TGCCTCGCGC TTACAAAAAC GGCACGAATG ACATCGAGGC GCGTGAGAAA
    2101  ATGGCCCATG CAAGCAACAT CGCGGGCATG GCCTTCGCCA ACGCGTTCCT GGGCGTGTGC
    2161  CACTCTATGG CTCACAAACT GGGTGCTATG CACCACGTGC CGCACGGTAT CGCGTGTGCT
    2221  GTCCTGATCG AAGAAGTAAT TAAGTACAAC GCTACTGATT GCCCGACTAA ACAGACCGCC
    2281  TTCCCACAGT ACAAATCTCC TAACGCTAAA CGTAAGTACG CTGAGATCGC CGAATACCTG
    2341  AACCTGAAGG GTACGAGCGA CACTGAGAAA GTTACTGCGC TGATCGAAGC TATCTCTAAA
    2401  CTGAAAATTG ACCTGTCCAT CCCGCAGAAC ATCAGCGCCG CAGGCATCAA CAAAAAGGAC
    2461  TTTTACAACA CGCTGGACAA AATGAGCGAA CTGGCTTTTG ACGACCAGTG CACCACTGCA
    2521  AACCCGCGTT ACCCGCTGAT CTCTGAGCTG AAAGACATTT ATATCAAATC TTTTTAA
```

**AdhE2.51 (76980)**

```
       1  ATGAAAGTGA CGAACCAAAA AGAACTGAAA CAAAAACTGA ATGAACTGCG TGAGGCCCAG
      61  AAAAAGTTCG CTACGTATAC CCAGGCGCAA GTTGACAAGT TCTTCAAACA GTGTGCCATT
```

121 GCGGCTGCCA AAGAGCGTAT TAACTTGGCC AAACTGGCGG TGGAAGAAAC CGGCATCGGT

181 CTGGTCGAGG ACAAGATTAT CAAGAATCAC TTTGCCGCGG AGTATATCTA CAACAAATAC

```
     241  AAAAACGAAA AGACTTGCGG CATCATTGAT CATGATGACA GCTTGGGTAT CACGAAAGTC
     301  GCAGAACCAA TCGGTATTGT CGCGGCAATT GTCCCGACCA CTAATCCGAC CAGCACCGCG
     361  ATCTTCAAGA GCTTGATCAG CCTGAAAACC CGCAACGCGA TTTTCTTCAG CCCGCACCCG
     421  CGTGCGAAGA AGAGCACCAT CGCAGCGGCG AAGCTGATTT TGGATGCCGC GGTTAAGGCC
     481  GGTGCACCGA AAAACATTAT TGGCTGGATT GACGAGCCGA GCATTGAGCT GTCTCAGGAC
     541  CTGATGAGCG AAGCGGACAT TATTCTGGCC ACCGGTGGTC CGTCTATGGT TAAGGCAGCG
     601  TACTCCAGCG GTAAGCCTGC TATCGGCGTT GGTGCGGGCA ATACGCCGGC TATCATCGAT
     661  GAAAGCGCGG ATATTGATAT GGCAGTTAGC TCCATCATTC TGTCCAAAAC CTACGACAAT
     721  GGTGTTATCT GTGCAAGCGA GCAAAGCATC CTGGTGATGA ATAGCATTTA CGAGAAGGTC
     781  AAAGAAGAGT TTGTTAAGCG TGGTTCCTAT ATTCTGAACC AGGAGGAGAT TGCAAAAATC
     841  AAAGAAACGA TGTTTAAGAA TGGCGCGATT AATGCAGATA TTGTTGGTAA GTCCGCATAC
     901  ATTATCGCGA AGATGGCGGG TATCGAAGTG CCGCAATTCA CCAAAATCTT GATCGGCGAA
     961  GTGCAGAGCG TGGAGAAGAG CGAACTGTTC AGCCATGAGA AACTGAGCCC GGTTCTGGCT
    1021  ATGTACAAAG TGAAGGACTT TGACGAAGCC CTGAAGAAAG CTCAGCGTCT GATTGAGCTG
    1081  GGCGGCAGCG GTCACACGAG CAGCCTGTAT ATCGATAGCC AAAACAATAA GGACAAGGTG
    1141  AAAGAGTTCG GTCTGGCGAT GAAAACCTCT CGCACGTTTA TCAACATGCC GAGCAGCCAA
    1201  GGTGCGAGCG GCGATCTGTA TAACTTCGCA ATCGCGCCTT CTTTTACGCT GGGTTGCGGT
    1261  ACCTGGGGCG GTAATTCGGT GAGCCAGAAT GTCGAGCCGA AGCACCTGCT GAACATCAAA
    1321  TCGGTCGCGG AGCGCCGTGA GAAGAGACCG TGGTTCAAAG TCCCACAGAA AATTTACTTC
    1381  AAATACGGCT GCCTGCGTTT CGCGCTGAAA GAACTGAAAG ACATGAACAA AAAGCGTGCG
    1441  TTCATTGTTA CCGACAAAGA CCTGTTCAAA CTGGGTTACG TGAACAAAAT CACCAAAGTT
    1501  CTGGATGAAA TTGACATCAA GTACTCCATC TTCACTGATA TCAAATCCGA CCCAACGATT
    1561  GATAGCGTGA AAAAGGGCGC TAAAGAAATG CTGAACTTTG AACCGGACAC CATCATCAGC
    1621  ATCGGTGGTG GCTCTCCTAT GGATGCTGCG AAGGTCATGC ACCTGCTGTA CGAATACCCG
    1681  GAAGCGGAAA TCGAAAACCT GGCTATCAAC TTCATGGACA TCCGCAAACG TATCTGCAAC
    1741  TTCCCGAAGC TGGGCACTAA AGCTATTTCC GTTGCCATCC CGACTACCGC GGGCACTGGT
    1801  TCCGAAGCCA CGCCGTTCGC CGTGATCACC AACGATGAAA CCGGTATGAA ATACCCGCTG
    1861  ACCTCTTACG AACTGACCCC GAACATGGCA ATTATCGACA CCGAGCTGAT GCTGAACATG
    1921  CCGCGCAAGC TGACCGCTGC TACCGGCATC GACGCTCTGG TACATGCTAT TGAGGCGTAC
    1981  GTTTCCGTGA TGGCTACCGA TTACACCGAC GAACTGGCCC TGCGTGCGAT CAAAATGATT
    2041  TTCAAGTACC TGCCTCGCGC TTACAAAAAC GGCACGAATG ACATCGAGGC GCGTGAGAAA
    2101  ATGGCCCATG CAAGCAACAT CGCGGGCATG GCCTTCGCCA ACGCGTTCCT GGGCGTGTGC
    2161  CACTCTATGG CTCACAAACT GGGTGCTATG CACCACGTGC CGCACGGTAT CGCGTGTGCT
    2221  GTCCTGATCG AAGAAGTAAT TAAGTACAAC GCTACTGATT GCCCGACTAA ACAGACCGCC
    2281  TTCCCACAGT ACAAATCTCC TAACGCTAAA CGTAAGTACG CTGAGATCGC CGAATACCTG
    2341  AACCTGAAGG GTACGAGCGA CACTGAGAAA GTTACTGCGC TGATCGAAGC TATCTCTAAA
    2401  CTGAAAATTG ACCTGTCCAT CCCGCAGAAC ATCAGCGCCG CAGGCATCAA CAAAAAGGAC
    2461  TTTTACAACA CGCTGGACAA AATGAGCGAA CTGGCTTTTG ACGACCAGTG CACCACTGCA
    2521  AACCCGCGTT ACCCGCTGAT CTCTGAGCTG AAAGACATTT ATATCAAATC TTTTTAA
```

**AdhE2.52 (76981)**

```
       1  ATGAAAGTGA CGAACCAAAA AGAACTGAAA CAAAAACTGA ATGAACTGCG TGAGGCCCAG
      61  AAAAAGTTCG CTACGTATAC CCAGGAGCAA GTTGAGAAGA TCTTCAAACA GTGTGCCATT
```

121 GCGGCTGCCA AAGAGCGTAT TAACTTGGCC AAACTGGCGG TGGAAGAAAC CGGCATCGGT

181 CTGGTCGAGG ACAAGATTAT CAAGAATCAC TTTGCCGCGG AGTATATCTA CAACAAATAC

```
     241  AAAAACGAAA AGACTTGCGG CATCATTGAT CATGATGACA GCTTGGGTAT CACGAAAGTC
     301  GCAGAACCAA TCGGTATTGT CGCGGCAATT GTCCCGACCA CTAATCCGAC CAGCACCGCG
     361  ATCTTCAAGA GCTTGATCAG CCTGAAAACC CGCAACGCGA TTTTCTTCAG CCCGCACCCG
     421  CGTGCGAAGA AGAGCACCAT CGCAGCGGCG AAGCTGATTT TGGATGCCGC GGTTAAGGCC
     481  GGTGCACCGA AAAACATTAT TGGCTGGATT GACGAGCCGA GCATTGAGCT GTCTCAGGAC
     541  CTGATGAGCG AAGCGGACAT TATTCTGGCC ACCGGTGGTC CGTCTATGGT TAAGGCAGCG
     601  TACTCCAGCG GTAAGCCTGC TATCGGCGTT GGTGCGGGCA ATACGCCGGC TATCATCGAT
     661  GAAAGCGCGG ATATTGATAT GGCAGTTAGC TCCATCATTC TGTCCAAAAC CTACGACAAT
     721  GGTGTTATCT GTGCAAGCGA GCAAAGCATC CTGGTGATGA ATAGCATTTA CGAGAAGGTC
     781  AAAGAAGAGT TTGTTAAGCG TGGTTCCTAT ATTCTGAACC AGCAGGAGAT TGCAAAAATC
     841  AAAGAAACGA TGTTTAAGAA TGGCGCGATT AATGCAGATA TTGTTGGTAA GTCCGCATAC
     901  ATTATCGCGA AGATGGCGGG TATCGAAGTG CCGCAAACCA CCAAAATCTT GATCGGCGAA
     961  GTGCAGGACG TGGAGAAGAG CGAACTGTTC AGCCATGAGA AACTGAGCCC GGTTCTGGCT
    1021  ATGTACAAAG TGAAGGACTT TGACGAAGCC CTGAAGAAAG CTCAGCGTCT GATTGAGCTG
    1081  GGCGGCAGCG GTCACACGAG CAGCCTGTAT ATCGATAGCC AAAACAATAA GGACAAGGTG
    1141  AAAGAGTTCG GTCTGGAGAT GAAAACCTCT CGCACGTTTA TCAACATGCC GAGCAGCCAA
    1201  GGTGCGAGCG GCGATCTGTA TAACTTCGCA ATCGCGCCTT CTTTTACGCT GGGTTGCGGT
    1261  ACCTGGGGCG GTAATTCGGT GAGCCAGAAT GTCGAGCCGA AGCACCTGCT GAACATCAAA
    1321  TCGGTCGCGG AGCGCCGTGA GAAGAGACCG TGGTTCAAAG TCCCACAGAA AATTTACTTC
    1381  AAATACGGCT GCCTGCGTTT CGCGCTGAAA GAACTGAAAG ACATGAACAA AAAGCGTGCG
    1441  TTCATTGTTA CCGACAAAGA CCTGTTCAAA CTGGGTTACG TGAACAAAAT CACCAAAGTT
    1501  CTGGATGAAA TTGACATCAA GTACTCCATC TTCACTGATA TCAAATCCGA CCCAACGATT
    1561  GATAGCGTGA AAAAGGGCGC TAAAGAAATG CTGAACTTTG AACCGGACAC CATCATCAGC
    1621  ATCGGTGGTG GCTCTCCTAT GGATGCTGCG AAGGTCATGC ACCTGCTGTA CGAATACCCG
    1681  GAAGCGGAAA TCGAAAACCT GGCTATCAAC TTCATGGACA TCCGCAAACG TATCTGCAAC
    1741  TTCCCGAAGC TGGGCACTAA AGCTATTTCC GTTGCCATCC CGACTACCGC GGGCACTGGT
    1801  TCCGAAGCCA CGCCGTTCGC CGTGATCACC AACGATGAAA CCGGTATGAA ATACCCGCTG
    1861  ACCTCTTACG AACTGACCCC GAACATGGCA ATTATCGACA CCGAGCTGAT GCTGAACATG
    1921  CCGCGCAAGC TGACCGCTGC TACCGGCATC GACGCTCTGG TACATGCTAT TGAGGCGTAC
    1981  GTTTCCGTGA TGGCTACCGA TTACACCGAC GAACTGGCCC TGCGTGCGAT CAAAATGATT
    2041  TTCAAGTACC TGCCTCGCGC TTACAAAAAC GGCACGAATG ACATCGAGGC GCGTGAGAAA
    2101  ATGGCCCATG CAAGCAACAT CGCGGGCATG GCCTTCGCCA ACGCGTTCCT GGGCGTGTGC
    2161  CACTCTATGG CTCACAAACT GGGTGCTATG CACCACGTGC CGCACGGTAT CGCGTGTGCT
    2221  GTCCTGATCG AAGAAGTAAT TAAGTACAAC GCTACTGATT GCCCGACTAA ACAGACCGCC
    2281  TTCCCACAGT ACAAATCTCC TAACGCTAAA CGTAAGTACG CTGAGATCGC CGAATACCTG
    2341  AACCTGAAGG GTACGAGCGA CACTGAGAAA GTTACTGCGC TGATCGAAGC TATCTCTAAA
    2401  CTGAAAATTG ACCTGTCCAT CCCGCAGAAC ATCAGCGCCG CAGGCATCAA CAAAAAGGAC
    2461  TTTTACAACA CGCTGGACAA AATGAGCGAA CTGGCTTTTG ACGACCAGTG CACCACTGCA
    2521  AACCCGCGTT ACCCGCTGAT CTCTGAGCTG AAAGACATTT ATATCAAATC TTTTTAA
```

**AdhE2.53 (76982)**

```
       1  ATGAAAGTGA CGAACCAAAA AGAACTGAAA CAAAAACTGA ATGAACTGCG TGAGGCCCAG
      61  AAAAAGTTCA ACACGTATAC CCAGGAGCAA GTTGACAAGA TCTTCAAACA GTGTGCCATT
```

121 GCGGCTGCCA AATACCGTAT TAACTTGGCC AAACTGGCGG TGGAAGAAAC CGGCATCGGT

181 CTGGTCGAGG ACAAGATTAT CAAGAATCAC TTTGCCGCGG AGTATATCTA CAACAAATAC

```
     241  AAAAACGAAA AGACTTGCGG CATCATTGAT CATGATGACA GCTTGGGTAT CACGAAAGTC
     301  GCAGAACCAA TCGGTATTGT CGCGGCAATT GTCCCGACCA CTAATCCGAC CAGCACCGCG
     361  ATCTTCAAGA GCTTGATCAG CCTGAAAACC CGCAACGCGA TTTTCTTCAG CCCGCACCCG
     421  CGTGCGAAGA AGAGCACCAT CGCAGCGGCG AAGCTGATTT TGGATGCCGC GGTTAAGGCC
     481  GGTGCACCGA AAAACATTAT TGGCTGGATT GACGAGCCGA GCATTGAGCT GTCTCAGGAC
     541  CTGATGAGCG AAGCGGACAT TATTCTGGCC ACCGGTGGTC CGTCTATGGT TAAGGCAGCG
     601  TACTCCAGCG GTAAGCCTGC TATCGGCGTT GGTGCGGGCA ATACGCCGGC TATCATCGAT
     661  GAAAGCGCGG ATATTGATAT GGCAGTTAGC TCCATCATTC TGTCCAAAAC CTACGACAAT
     721  GGTGTTATCT GTGCAAGCGA GCAAAGCATC CTGGTGATGA ATAGCATTTA CGAGAAGGTC
     781  AAAGAAGAGT TTGTTAAGCG TGGTTCCTAT ATTCTGAACC AGAACGCGAT TGCAAAAATC
     841  AAAGAAACGA TGTTTAAGAA TGGCGCGATT AATGCAGATA TTGTTGGTAA GTCCGCATAC
     901  ATTATCGCGA AGATGGCGGG TATCGAAGTG CCGCAAACCA CCAAAATCTT GATCGGCGAA
     961  GTGCAGAGCG TGGAGAAGAG CGAACTGTTC AGCCATGAGA AACTGAGCCC GGTTCTGGCT
    1021  ATGTACAAAG TGAAGGACTT TGACGAAGCC CTGAAGAAAG CTCAGCGTCT GATTGAGCTG
    1081  GGCGGCAGCG GTCACACGAG CAGCCTGTAT ATCGATAGCC AAAACAATAA GGACAAGGTG
    1141  AAAGAGTTCG GTCTGGCGAT GAAAACCTCT CGCACGTTTA TCAACATGCC GAGCAGCCAA
    1201  GGTGCGAGCG GCGATCTGTA TAACTTCGCA ATCGCGCCTT CTTTTACGCT GGGTTGCCCG
    1261  ACCTGGGGCG GTAATTCGGT GAGCCAGAAT GTCGAGCCGA AGCACCTGCT GAACATCAAA
    1321  TCGGTCGCGG AGCGCCGTGA GAAGAGACCG TGGTTCAAAG TCCCACAGAA AATTTACTTC
    1381  AAATACGGCT GCCTGCGTTT CGCGCTGAAA GAACTGAAAG ACATGAACAA AAAGCGTGCG
    1441  TTCATTGTTA CCGACAAAGA CCTGTTCAAA CTGGGTTACG TGAACAAAAT CACCAAAGTT
    1501  CTGGATGAAA TTGACATCAA GTACTCCATC TTCACTGATA TCAAATCCGA CCCAACGATT
    1561  GATAGCGTGA AAAAGGGCGC TAAAGAAATG CTGAACTTTG AACCGGACAC CATCATCAGC
    1621  ATCGGTGGTG GCTCTCCTAT GGATGCTGCG AAGGTCATGC ACCTGCTGTA CGAATACCCG
    1681  GAAGCGGAAA TCGAAAACCT GGCTATCAAC TTCATGGACA TCCGCAAACG TATCTGCAAC
    1741  TTCCCGAAGC TGGGCACTAA AGCTATTTCC GTTGCCATCC CGACTACCGC GGGCACTGGT
    1801  TCCGAAGCCA CGCCGTTCGC CGTGATCACC AACGATGAAA CCGGTATGAA ATACCCGCTG
    1861  ACCTCTTACG AACTGACCCC GAACATGGCA ATTATCGACA CCGAGCTGAT GCTGAACATG
    1921  CCGCGCAAGC TGACCGCTGC TACCGGCATC GACGCTCTGG TACATGCTAT TGAGGCGTAC
    1981  GTTTCCGTGA TGGCTACCGA TTACACCGAC GAACTGGCCC TGCGTGCGAT CAAAATGATT
    2041  TTCAAGTACC TGCCTCGCGC TTACAAAAAC GGCACGAATG ACATCGAGGC GCGTGAGAAA
    2101  ATGGCCCATG CAAGCAACAT CGCGGGCATG GCCTTCGCCA ACGCGTTCCT GGGCGTGTGC
    2161  CACTCTATGG CTCACAAACT GGGTGCTATG CACCACGTGC CGCACGGTAT CGCGTGTGCT
    2221  GTCCTGATCG AAGAAGTAAT TAAGTACAAC GCTACTGATT GCCCGACTAA ACAGACCGCC
    2281  TTCCCACAGT ACAAATCTCC TAACGCTAAA CGTAAGTACG CTGAGATCGC CGAATACCTG
    2341  AACCTGAAGG GTACGAGCGA CACTGAGAAA GTTACTGCGC TGATCGAAGC TATCTCTAAA
    2401  CTGAAAATTG ACCTGTCCAT CCCGCAGAAC ATCAGCGCCG CAGGCATCAA CAAAAAGGAC
    2461  TTTTACAACA CGCTGGACAA AATGAGCGAA CTGGCTTTTG ACGACCAGTG CACCACTGCA
    2521  AACCCGCGTT ACCCGCTGAT CTCTGAGCTG AAAGACATTT ATATCAAATC TTTTTAA
```

**AdhE2.54 (76983)**

```
       1  ATGAAAGTGA CGAACCAAAA AGAACTGAAA CAAAAACTGA ATGAACTGCG TGAGGCCCAG
      61  AAAAAGTTCG CTACGTATAC CCAGGAGCAA GTTGACAAGA TCTTCAAACA GTGTGCCATT
```

121 GCGGCTGCCA AAGAGCGTAT TAACTTGGCC AAACTGGCGG TGGAAGAAAC CGGCATCGGT

181 CTGGTCGAGG ACAAGATTAT CAAGAATCAC TTTGCCGCGG AGTATATCTA CAACAAATAC

```
     241  AAAAACGAAA AGACTTGCGG CATCATTGAT CATGATGACA GCTTGGGTAT CACGAAAGTC
     301  GCAGAACCAA TCGGTATTGT CGCGGCAATT GTCCCGACCA CTAATCCGAC CAGCACCGCG
     361  ATCTTCAAGA GCTTGATCAG CCTGAAAACC CGCAACGCGA TTTTCTTCAG CCCGCACCCG
     421  CGTGCGAAGA AGAGCACCAT CGCAGCGGCG AAGCTGATTT TGGATGCCGC GGTTAAGGCC
     481  GGTGCACCGA AAAACATTAT TGGCTGGATT GACGAGCCGA GCATTGAGCT GTCTCAGGAC
     541  CTGATGAGCG AAGCGGACAT TATTCTGGCC ACCGGTGGTC CGTCTATGGT TAAGGCAGCG
     601  TACTCCAGCG GTAAGCCTGC TATCGGCGCG GGTGCGGGCA ATACGCCGGC TATCATCGAT
     661  GAAAGCGCGG ATATTGATAT GGCAGTTAGC TCCATCATTC TGTCCAAAAC CTACGACAAT
     721  GGTGTTATCT GTGCAAGCGA GCAAAGCATC CTGGTGATGA ATAGCATTTA CGAGAAGGTC
     781  AAAGAAGAGT TTGTTAAGCG TGGTTCCTAT ATTCTGAACC AGCAGGAGAT TGCAAAAATC
     841  AAAGAAACGA TGTTTAAGAA TGGCGCGATT AATGCAGATA TTGTTGGTAA GTCCGCATAC
     901  ATTATCGCGA AGATGGCGGG TATCGAAGTG CCGCAAACCA CCAAAATCTT GATCGGCGAA
     961  GTGCAGAGCG TGGAGAAGAG CGAACTGTTC AGCGAGGAGA AACTGAGCCC GGTTCTGGCT
    1021  ATGTACAAAG TGAAGGACTT TGACGAAGCC CTGAAGAAAG CTCAGCGTCT GATTGAGCTG
    1081  GGCGGCAGCG GTCACACGAG CAGCCTGTAT ATCGATAGCC AAAACAATAA GGACAAGGTG
    1141  AAAGAGTTCG GTCTGGCGAT GAAAACCTCT CGCACGTTTA TCAACATGCC GAGCAGCCAA
    1201  GGTGCGAGCG GCGATCTGTA TAACTTCGCA ATCGCGCCTT CTTTTACGCT GGGTTGCGGT
    1261  ACCTGGGGCG GTAATTCGGT GAGCCAGAAT GTCGAGCCGA AGCACCTGCT GAACATCAAA
    1321  TCGGTCGCGC GTCGCCGTGA GAAGAGACCG TGGTTCAAAG TCCCACAGAA AATTTACTTC
    1381  AAATACGGCT GCCTGCGTTT CGCGCTGAAA GAACTGAAAG ACATGAACAA AAAGCGTGCG
    1441  TTCATTGTTA CCGACAAAGA CCTGTTCAAA CTGGGTTACG TGAACAAAAT CACCAAAGTT
    1501  CTGGATGAAA TTGACATCAA GTACTCCATC TTCACTGATA TCAAATCCGA CCCAACGATT
    1561  GATAGCGTGA AAAAGGGCGC TAAAGAAATG CTGAACTTTG AACCGGACAC CATCATCAGC
    1621  ATCGGTGGTG GCTCTCCTAT GGATGCTGCG AAGGTCATGC ACCTGCTGTA CGAATACCCG
    1681  GAAGCGGAAA TCGAAAACCT GGCTATCAAC TTCATGGACA TCCGCAAACG TATCTGCAAC
    1741  TTCCCGAAGC TGGGCACTAA AGCTATTTCC GTTGCCATCC CGACTACCGC GGGCACTGGT
    1801  TCCGAAGCCA CGCCGTTCGC CGTGATCACC AACGATGAAA CCGGTATGAA ATACCCGCTG
    1861  ACCTCTTACG AACTGACCCC GAACATGGCA ATTATCGACA CCGAGCTGAT GCTGAACATG
    1921  CCGCGCAAGC TGACCGCTGC TACCGGCATC GACGCTCTGG TACATGCTAT TGAGGCGTAC
    1981  GTTTCCGTGA TGGCTACCGA TTACACCGAC GAACTGGCCC TGCGTGCGAT CAAAATGATT
    2041  TTCAAGTACC TGCCTCGCGC TTACAAAAAC GGCACGAATG ACATCGAGGC GCGTGAGAAA
    2101  ATGGCCCATG CAAGCAACAT CGCGGGCATG GCCTTCGCCA ACGCGTTCCT GGGCGTGTGC
    2161  CACTCTATGG CTCACAAACT GGGTGCTATG CACCACGTGC CGCACGGTAT CGCGTGTGCT
    2221  GTCCTGATCG AAGAAGTAAT TAAGTACAAC GCTACTGATT GCCCGACTAA ACAGACCGCC
    2281  TTCCCACAGT ACAAATCTCC TAACGCTAAA CGTAAGTACG CTGAGATCGC CGAATACCTG
    2341  AACCTGAAGG GTACGAGCGA CACTGAGAAA GTTACTGCGC TGATCGAAGC TATCTCTAAA
    2401  CTGAAAATTG ACCTGTCCAT CCCGCAGAAC ATCAGCGCCG CAGGCATCAA CAAAAAGGAC
    2461  TTTTACAACA CGCTGGACAA AATGAGCGAA CTGGCTTTTG ACGACCAGTG CACCACTGCA
    2521  AACCCGCGTT ACCCGCTGAT CTCTGAGCTG AAAGACATTT ATATCAAATC TTTTTAA
```

**AdhE2.55 (76984)**

```
       1  ATGAAAGTGA CGAACCAAAA AGAACTGAAA CAAAAACTGA ATGAACTGCG TGAGGCCCAG
      61  AAAAAGTTCG CTACGTATAC CCAGGAGCAA GTTGACAAGA TCTTCAAACA GTGTGCCATT
```

121 GCGGCTGCCA AAGAGCGTAT TAACTTGGCC AAACTGGCGG TGGAAGAAAC CGGCATCGGT

181 CTGGTCGAGG ACAAGATTAT CAAGAATCAC TTTGCCGCGG AGTATATCTA CAACAAATAC

```
     241  AAAAACGAAA AGACTTGCGG CATCATTGAT CATGATGACA GCTTGGGTAT CACGAAAGTC
     301  GCAGAACCAA TCGGTATTGT CGCGGCAATT GTCCCGACCA CTAATCCGAC CAGCACCGCG
     361  ATCTTCAAGA GCTTGATCAG CCTGAAAACC CGCAACGCGA TTTTCTTCAG CCCGCACCCG
     421  CGTGCGAAGA AGAGCACCAT CGCAGCGGCG ACCCTGATTT TGGATGCCGC GGTTAAGGCC
     481  GGTGCACCGA AAAACATTAT TGGCTGGATT GACGAGCCGA GCATTGAGCT GTCTCAGGAC
     541  CTGATGAGCG AAGCGGACAT TATTCTGGCC ACCGGTGGTC CGTCTATGGT TAAGGCAGCG
     601  TACTCCAGCG GTAAGCCTGC TATCGGCGTT GGTGCGGGCA ATACGCCGGC TATCATCGAT
     661  GAAAGCGCGG ATATTGATAT GGCAGTTAGC TCCATCATTC TGTCCAAAAC CTACGACAAT
     721  GGTGTTATCT GTGCAAGCGA GCAAAGCATC CTGGTGATGA ATAGCATTTA CGAGAAGGTC
     781  AAAGCGGAGT TTGTTAAGCG TGGTTCCTAT ATTCTGAACC AGAACGAGAT TGCAAAAATC
     841  AAAGAAACGA TGTTTAAGAA TGGCGCGATT AATGCAGATA TTGTTGGTAA GTCCGCATAC
     901  ATTATCGCGA AGATGGCGGG TATCGAAGTG CCGCAAACCA CCAAAATCTT GATCGGCGAA
     961  GTGCAGAGCG TGGAGAAGAG CGAACTGTTC AGCAAAGAGA AACTGAGCCC GGTTCTGGCT
    1021  ATGTACAAAG TGAAGGACTT TGACGAAGCC CTGAAGAAAG CTCAGCGTCT GATTAACCTG
    1081  GGCGGCAGCG GTCACACGAG CAGCCTGTAT ATCGATAGCC AAAACAATAA GGACAAGGTG
    1141  AAAGAGTTCG GTCTGGCGAT GAAAACCTCT CGCACGTTTA TCAACATGCC GAGCAGCCAA
    1201  GGTGCGAGCG GCGATCTGTA TAACTTCGCA ATCGCGCCTT CTTTTACGCT GGGTTGCGGT
    1261  ACCTGGGGCG GTAATTCGGT GAGCCAGAAT GTCGAGCCGA AGCACCTGCT GAACATCAAA
    1321  TCGGTCGCGG AGCGCCGTGA GAAGAGACCG TGGTTCAAAG TCCCACAGAA AATTTACTTC
    1381  AAATACGGCT GCCTGCGTTT CGCGCTGAAA GAACTGAAAG ACATGAACAA AAAGCGTGCG
    1441  TTCATTGTTA CCGACAAAGA CCTGTTCAAA CTGGGTTACG TGAACAAAAT CACCAAAGTT
    1501  CTGGATGAAA TTGACATCAA GTACTCCATC TTCACTGATA TCAAATCCGA CCCAACGATT
    1561  GATAGCGTGA AAAAGGGCGC TAAAGAAATG CTGAACTTTG AACCGGACAC CATCATCAGC
    1621  ATCGGTGGTG GCTCTCCTAT GGATGCTGCG AAGGTCATGC ACCTGCTGTA CGAATACCCG
    1681  GAAGCGGAAA TCGAAAACCT GGCTATCAAC TTCATGGACA TCCGCAAACG TATCTGCAAC
    1741  TTCCCGAAGC TGGGCACTAA AGCTATTTCC GTTGCCATCC CGACTACCGC GGGCACTGGT
    1801  TCCGAAGCCA CGCCGTTCGC CGTGATCACC AACGATGAAA CCGGTATGAA ATACCCGCTG
    1861  ACCTCTTACG AACTGACCCC GAACATGGCA ATTATCGACA CCGAGCTGAT GCTGAACATG
    1921  CCGCGCAAGC TGACCGCTGC TACCGGCATC GACGCTCTGG TACATGCTAT TGAGGCGTAC
    1981  GTTTCCGTGA TGGCTACCGA TTACACCGAC GAACTGGCCC TGCGTGCGAT CAAAATGATT
    2041  TTCAAGTACC TGCCTCGCGC TTACAAAAAC GGCACGAATG ACATCGAGGC GCGTGAGAAA
    2101  ATGGCCCATG CAAGCAACAT CGCGGGCATG GCCTTCGCCA ACGCGTTCCT GGGCGTGTGC
    2161  CACTCTATGG CTCACAAACT GGGTGCTATG CACCACGTGC CGCACGGTAT CGCGTGTGCT
    2221  GTCCTGATCG AAGAAGTAAT TAAGTACAAC GCTACTGATT GCCCGACTAA ACAGACCGCC
    2281  TTCCCACAGT ACAAATCTCC TAACGCTAAA CGTAAGTACG CTGAGATCGC CGAATACCTG
    2341  AACCTGAAGG GTACGAGCGA CACTGAGAAA GTTACTGCGC TGATCGAAGC TATCTCTAAA
    2401  CTGAAAATTG ACCTGTCCAT CCCGCAGAAC ATCAGCGCCG CAGGCATCAA CAAAAAGGAC
    2461  TTTTACAACA CGCTGGACAA AATGAGCGAA CTGGCTTTTG ACGACCAGTG CACCACTGCA
    2521  AACCCGCGTT ACCCGCTGAT CTCTGAGCTG AAAGACATTT ATATCAAATC TTTTTAA
```

**AdhE2.56 (76985)**

```
       1  ATGAAAGTGA CGAACCAAAA AGAACTGAAA CAAAAACTGA ATGAACTGCG TGAGGCCCAG
      61  AAAAAGTTCG CTACGTATAC CCAGGAGCAA GTTGACAAGA TCTTCAAACA GTGTGCCATT
```

121 GCGGCTGCCA AAGAGCGTAT TAACTTGGCC AAACTGGCGG TGGAAGAAAC CGGCATCGGT

181 CTGGTCGAGG ACAAGATTAT CAAGAATCAC TTTGCCGCGG AGTATATCTA CAACAAATAC

```
     241  AAAAACGAAA AGACTTGCGG CATCATTGAT CATGATGACA GCTTGGGTAT CACGAAAGTC
     301  GCAGAACCAA TCGGTATTGT CGCGGCAATT GTCCCGACCA CTAATCCGAC CAGCACCGCG
     361  ATCTTCAAGA GCTTGATCAG CCTGAAAACC CGCAACGCGA TTTTCTTCAG CCCGCACCCG
     421  CGTGCGAAGA AGAGCACCAT CGCAGCGGCG AAGCTGATTT TGGATGCCGC GGTTAAGGCC
     481  GGTGCACCGA AAAACATTAT TGGCTGGATT GACGAGCCGA GCATTGAGCT GTCTCAGGAC
     541  CTGATGAGCG AAGCGGACAT TATTCTGGCC ACCGGTGGTC CGTCTATGGT TAAGGCAGCG
     601  TACTCCAGCG GTAAGCCTGC TATCGGCGTT GGTGCGGGCA ATACGCCGGC TATCATCGAT
     661  GAAAGCGCGG ATATTGATAT GGCAGTTAGC TCCATCATTC TGTCCAAAAC CTACGACAAT
     721  GGTGTTATCT GTGCAAGCGA GCAAAGCATC CTGGTGATGA ATAGCATTTA CGAGAAGGTC
     781  AAAGAAGAGT TTGTTAAGCG TGGTTCCTAT ATTCTGAACC AGAACGAGAT TGCAAAAATC
     841  AAAGAAACGA TGTTTAAGAA TGGCGCGATT AATCGTGATA TTGTTGGTAA GTCCGCATAC
     901  ATTATCGCGA AGATGGCGGG TATCGAAGTG CCGCAATTCA CCAAAATCTT GATCGGCGAA
     961  GTGCAGGACG TGGAGAAGAG CGAACTGTTC AGCCATGAGA AACTGAGCCC GGTTCTGGCT
    1021  ATGTACAAAG TGAAGGACTT TGACGAAGCC CTGAAGAAAG CTCAGCGTCT GATTGAGCTG
    1081  GGCGGCAGCG GTCACACGAG CAGCCTGTAT ATCGATAGCC AAAACAATAA GGACAAGGTG
    1141  AAAGAGTTCG GTCTGGCGAT GAAAACCTCT CGCACGTTTA TCAACATGCC GAGCAGCCAA
    1201  GGTGCGAGCG GCGATCTGTA TAACTTCGCA ATCGCGCCTT CTTTTACGCT GGGTTGCGGT
    1261  ACCTGGGGCG GTAATTCGGT GAGCCAGAAT GTCGAGCCGA AGCACCTGCT GAACATCAAA
    1321  TCGGTCGCGC GTCGCCGTGA GAAGAGACCG TGGTTCAAAG TCCCACAGAA AATTTACTTC
    1381  AAATACGGCT GCCTGCGTTT CGCGCTGAAA GAACTGAAAG ACATGAACAA AAAGCGTGCG
    1441  TTCATTGTTA CCGACAAAGA CCTGTTCAAA CTGGGTTACG TGAACAAAAT CACCAAAGTT
    1501  CTGGATGAAA TTGACATCAA GTACTCCATC TTCACTGATA TCAAATCCGA CCCAACGATT
    1561  GATAGCGTGA AAAAGGGCGC TAAAGAAATG CTGAACTTTG AACCGGACAC CATCATCAGC
    1621  ATCGGTGGTG GCTCTCCTAT GGATGCTGCG AAGGTCATGC ACCTGCTGTA CGAATACCCG
    1681  GAAGCGGAAA TCGAAAACCT GGCTATCAAC TTCATGGACA TCCGCAAACG TATCTGCAAC
    1741  TTCCCGAAGC TGGGCACTAA AGCTATTTCC GTTGCCATCC CGACTACCGC GGGCACTGGT
    1801  TCCGAAGCCA CGCCGTTCGC CGTGATCACC AACGATGAAA CCGGTATGAA ATACCCGCTG
    1861  ACCTCTTACG AACTGACCCC GAACATGGCA ATTATCGACA CCGAGCTGAT GCTGAACATG
    1921  CCGCGCAAGC TGACCGCTGC TACCGGCATC GACGCTCTGG TACATGCTAT TGAGGCGTAC
    1981  GTTTCCGTGA TGGCTACCGA TTACACCGAC GAACTGGCCC TGCGTGCGAT CAAAATGATT
    2041  TTCAAGTACC TGCCTCGCGC TTACAAAAAC GGCACGAATG ACATCGAGGC GCGTGAGAAA
    2101  ATGGCCCATG CAAGCAACAT CGCGGGCATG GCCTTCGCCA ACGCGTTCCT GGGCGTGTGC
    2161  CACTCTATGG CTCACAAACT GGGTGCTATG CACCACGTGC CGCACGGTAT CGCGTGTGCT
    2221  GTCCTGATCG AAGAAGTAAT TAAGTACAAC GCTACTGATT GCCCGACTAA ACAGACCGCC
    2281  TTCCCACAGT ACAAATCTCC TAACGCTAAA CGTAAGTACG CTGAGATCGC CGAATACCTG
    2341  AACCTGAAGG GTACGAGCGA CACTGAGAAA GTTACTGCGC TGATCGAAGC TATCTCTAAA
    2401  CTGAAAATTG ACCTGTCCAT CCCGCAGAAC ATCAGCGCCG CAGGCATCAA CAAAAAGGAC
    2461  TTTTACAACA CGCTGGACAA AATGAGCGAA CTGGCTTTTG ACGACCAGTG CACCACTGCA
    2521  AACCCGCGTT ACCCGCTGAT CTCTGAGCTG AAAGACATTT ATATCAAATC TTTTTAA
```

**AdhE2.57 (76986)**

```
       1  ATGAAAGTGA CGAACCAAAA AGAACTGAAA CAAAAACTGA ATGAACTGCG TGAGGCCCAG
      61  AAAAAGTTCG CTACGATGAC CCAGGAGCAA GTTGACAAGA TCTTCAAACA GTGTGCCATT
```

121 GCGGCTGCCA AAGAGCGTAT TAACTTGGCC AAACTGGCGG TGGAAGAAAC CGGCATCGGT

181 CTGGTCGAGG ACAAGATTAT CAAGAATCAC TTTGCCGCGG AGTATATCTA CAACAAATAC

```
     241  AAAAACGAAA AGACTTGCGG CATCATTGAT CATGATGACA GCTTGGGTAT CACGAAAGTC
     301  GCAGAACCAA TCGGTATTGT CGCGGCAATT GTCCCGACCA CTAATCCGAC CAGCACCGCG
     361  ATCTTCAAGA GCTTGATCAG CCTGAAAACC CGCAACGCGA TTTTCTTCAG CCCGCACCCG
     421  CGTGCGAAGA AGAGCACCAT CGCAGCGGCG AAGCTGATTT TGGATGCCGC GGTTAAGGCC
     481  GGTGCACCGA AAAACATTAT TGGCTGGATT GACGAGCCGA GCATTGAGCT GTCTCAGGAC
     541  CTGATGAGCG AAGCGGACAT TATTCTGGCC ACCGGTGGTC CGTCTATGGT TAAGGCAGCG
     601  TACTCCAGCG GTAAGCCTGC TATCGGCGTT GGTGCGGGCA ATACGCCGGC TATCATCGAT
     661  GAAAGCGCGG ATATTGATAT GGCAGTTAGC TCCATCATTC TGTCCAAAAC CTACGACAAT
     721  AACGTTATCT GTGCAAGCGA GCAAAGCATC CTGGTGATGA ATAGCATTTA CGAGAAGGTC
     781  AAAGAAGAGT TTGTTAAGCG TGGTTCCTAT ATTCTGAACC AGAACGAGAT TGCAAAAATC
     841  AAAGAAACGA TGTTTAAGAA TGGCGCGATT AATCGTGATA TTGTTGGTAA GTCCGCATAC
     901  ATTATCGCGA AGATGGCGGG TATCGAAGTG CCGCAAACCA CCAAAATCTT GATCGGCGAA
     961  GTGCAGAGCG TGGAGAAGAG CGAACTGTTC AGCGTTGAGA AACTGAGCCC GGTTCTGGCT
    1021  ATGTACAAAG TGAAGGACTT TGACGAAGCC CTGAAGAAAG CTCAGCGTCT GATTGAGCTG
    1081  GGCGGCAGCG GTCACACGAG CAGCCTGTAT ATCGATAGCC AAAACAATAA GGACAAGGTG
    1141  AAAGAGTTCG GTCTGGCGAT GAAAACCTCT CGCACGTTTA TCAACATGCC GAGCAGCCAA
    1201  GGTGCGAGCG GCGATCTGTA TAACTTCGCA ATCGCGCCTT CTTTTACGCT GGGTTGCGGT
    1261  ACCTGGGGCG GTAATTCGGT GAGCCAGAAT GTCGAGCCGA AGCACCTGCT GAACATCAAA
    1321  TCGGTCGCGG AGCGCCGTGA GAAGAGACCG TGGTTCAAAG TCCCACAGAA AATTTACTTC
    1381  AAATACGGCT GCCTGCGTTT CGCGCTGAAA GAACTGAAAG ACATGAACAA AAAGCGTGCG
    1441  TTCATTGTTA CCGACAAAGA CCTGTTCAAA CTGGGTTACG TGAACAAAAT CACCAAAGTT
    1501  CTGGATGAAA TTGACATCAA GTACTCCATC TTCACTGATA TCAAATCCGA CCCAACGATT
    1561  GATAGCGTGA AAAAGGGCGC TAAAGAAATG CTGAACTTTG AACCGGACAC CATCATCAGC
    1621  ATCGGTGGTG GCTCTCCTAT GGATGCTGCG AAGGTCATGC ACCTGCTGTA CGAATACCCG
    1681  GAAGCGGAAA TCGAAAACCT GGCTATCAAC TTCATGGACA TCCGCAAACG TATCTGCAAC
    1741  TTCCCGAAGC TGGGCACTAA AGCTATTTCC GTTGCCATCC CGACTACCGC GGGCACTGGT
    1801  TCCGAAGCCA CGCCGTTCGC CGTGATCACC AACGATGAAA CCGGTATGAA ATACCCGCTG
    1861  ACCTCTTACG AACTGACCCC GAACATGGCA ATTATCGACA CCGAGCTGAT GCTGAACATG
    1921  CCGCGCAAGC TGACCGCTGC TACCGGCATC GACGCTCTGG TACATGCTAT TGAGGCGTAC
    1981  GTTTCCGTGA TGGCTACCGA TTACACCGAC GAACTGGCCC TGCGTGCGAT CAAAATGATT
    2041  TTCAAGTACC TGCCTCGCGC TTACAAAAAC GGCACGAATG ACATCGAGGC GCGTGAGAAA
    2101  ATGGCCCATG CAAGCAACAT CGCGGGCATG GCCTTCGCCA ACGCGTTCCT GGGCGTGTGC
    2161  CACTCTATGG CTCACAAACT GGGTGCTATG CACCACGTGC CGCACGGTAT CGCGTGTGCT
    2221  GTCCTGATCG AAGAAGTAAT TAAGTACAAC GCTACTGATT GCCCGACTAA ACAGACCGCC
    2281  TTCCCACAGT ACAAATCTCC TAACGCTAAA CGTAAGTACG CTGAGATCGC CGAATACCTG
    2341  AACCTGAAGG GTACGAGCGA CACTGAGAAA GTTACTGCGC TGATCGAAGC TATCTCTAAA
    2401  CTGAAAATTG ACCTGTCCAT CCCGCAGAAC ATCAGCGCCG CAGGCATCAA CAAAAAGGAC
    2461  TTTTACAACA CGCTGGACAA AATGAGCGAA CTGGCTTTTG ACGACCAGTG CACCACTGCA
    2521  AACCCGCGTT ACCCGCTGAT CTCTGAGCTG AAAGACATTT ATATCAAATC TTTTTAA
```

**AdhE2.58 (76987)**

```
       1  ATGAAAGTGA CGAACCAAAA AGAACTGAAA CAAAAACTGA ATGAACTGCG TGAGGCCCAG
      61  AAAAAGTTCG CTACGTATAC CCAGGAGCAA GTTGACAAGA TCTTCAAACA GTGTGCCATT
```

121 GCGGCTGCCA AAGAGCGTAT TAACTTGGCC AAACTGGCGG TGGAAGAAAC CGGCATCGGT

181 CTGGTCGAGG ACAAGATTAT CAAGAATCAC TTTGCCGCGG AGTATATCTA CAACAAATAC

```
     241  AAAAACGAAA AGACTTGCGG CATCATTGAT CATGATGACA GCTTGGGTAT CACGAAAGTC
     301  GCAGAACCAA TCGGTATTGT CGCGAGCATT GTCCCGACCA CTAATCCGAC CAGCACCGCG
     361  ATCTTCAAGA GCTTGATCAG CCTGAAAACC CGCAACGCGA TTTTCTTCAG CCCGCACCCG
     421  CGTGCGAAGA AGAGCACCAT CGCAGCGGCG AAGCTGATTT TGGATGCCGC GGTTAAGGCC
     481  GGTGCACCGA AAAACATTAT TGGCTGGATT GACGAGCCGA GCATTGAGCT GTCTCAGGAC
     541  ATGATGAGCG AAGCGGACAT TATTCTGGCC ACCGGTGGTC CGTCTATGGT TAAGGCAGCG
     601  TACTCCAGCG GTAAGCCTGC TATCGGCGTT GGTGCGGGCA ATACGCCGGC TATCATCGAT
     661  GAAAGCGCGG ATATTGATAT GGCAGTTAGC TCCATCATTC TGTCCAAAAC CTACGACAAT
     721  GGTGTTATCT GTGCAAGCGA GCAAAGCATC CTGGTGATGA ATAGCATTTA CGAGAAGGTC
     781  AAAAAAGAGT TTGTTAAGCG TGGTTCCTAT ATTCTGAACC AGAACGAGAT TGCAAAAATC
     841  AAAGAAACGA TGTTTAAGAA TGGCGCGATT AATGCAGATA TTGTTGGTAA GTCCGCATAC
     901  ATTATCGCGA AGATGGCGGG TATCGAAGTG CCGCAAACCA CCAAAATCTT GATCGGCGAA
     961  GTGCAGAGCG TGGAGAAGAG CGAACTGTTC AGCCATGAGA AACTGAGCCC GGTTCTGGCT
    1021  ATGTACAAAG TGAAGGACTT TGACGAAGCC CTGAAGAAAG CTCAGCGTCT GATTGAGCTG
    1081  GGCGGCAGCC ACCACACGAG CAGCCTGTAT ATCGATAGCC AAAACAATAA GGACAAGGTG
    1141  AAAGAGTTCG GTCTGGCGAT GAAAACCTCT CGCACGTTTA TCAACATGCC GAGCAGCCAA
    1201  GGTGCGAGCG GCGATCTGTA TAACTTCGCA ATCGCGCCTT CTTTTACGCT GGGTTGCGGT
    1261  ACCTGGGGCG GTAATTCGGT GAGCCAGAAT GTCGAGCCGA AGCACCTGCT GAACATCAAA
    1321  TCGGTCGCGG AGCGCCGTGA GAAGAGACCG TGGTTCAAAG TCCCACAGAA AATTTACTTC
    1381  AAATACGGCT GCCTGCGTTT CGCGCTGAAA GAACTGAAAG ACATGAACAA AAAGCGTGCG
    1441  TTCATTGTTA CCGACAAAGA CCTGTTCAAA CTGGGTTACG TGAACAAAAT CACCAAAGTT
    1501  CTGGATGAAA TTGACATCAA GTACTCCATC TTCACTGATA TCAAATCCGA CCCAACGATT
    1561  GATAGCGTGA AAAAGGGCGC TAAAGAAATG CTGAACTTTG AACCGGACAC CATCATCAGC
    1621  ATCGGTGGTG GCTCTCCTAT GGATGCTGCG AAGGTCATGC ACCTGCTGTA CGAATACCCG
    1681  GAAGCGGAAA TCGAAAACCT GGCTATCAAC TTCATGGACA TCCGCAAACG TATCTGCAAC
    1741  TTCCCGAAGC TGGGCACTAA AGCTATTTCC GTTGCCATCC CGACTACCGC GGGCACTGGT
    1801  TCCGAAGCCA CGCCGTTCGC CGTGATCACC AACGATGAAA CCGGTATGAA ATACCCGCTG
    1861  ACCTCTTACG AACTGACCCC GAACATGGCA ATTATCGACA CCGAGCTGAT GCTGAACATG
    1921  CCGCGCAAGC TGACCGCTGC TACCGGCATC GACGCTCTGG TACATGCTAT TGAGGCGTAC
    1981  GTTTCCGTGA TGGCTACCGA TTACACCGAC GAACTGGCCC TGCGTGCGAT CAAAATGATT
    2041  TTCAAGTACC TGCCTCGCGC TTACAAAAAC GGCACGAATG ACATCGAGGC GCGTGAGAAA
    2101  ATGGCCCATG CAAGCAACAT CGCGGGCATG GCCTTCGCCA ACGCGTTCCT GGGCGTGTGC
    2161  CACTCTATGG CTCACAAACT GGGTGCTATG CACCACGTGC CGCACGGTAT CGCGTGTGCT
    2221  GTCCTGATCG AAGAAGTAAT TAAGTACAAC GCTACTGATT GCCCGACTAA ACAGACCGCC
    2281  TTCCCACAGT ACAAATCTCC TAACGCTAAA CGTAAGTACG CTGAGATCGC CGAATACCTG
    2341  AACCTGAAGG GTACGAGCGA CACTGAGAAA GTTACTGCGC TGATCGAAGC TATCTCTAAA
    2401  CTGAAAATTG ACCTGTCCAT CCCGCAGAAC ATCAGCGCCG CAGGCATCAA CAAAAAGGAC
    2461  TTTTACAACA CGCTGGACAA AATGAGCGAA CTGGCTTTTG ACGACCAGTG CACCACTGCA
    2521  AACCCGCGTT ACCCGCTGAT CTCTGAGCTG AAAGACATTT ATATCAAATC TTTTTAA
```

**AdhE2.59 (76988)**

```
       1  ATGAAAGTGA CGAACCAAAA AGAACTGAAA CAAAAACTGA ATGAACTGCG TGAGGCCCAG
      61  AAAAAGTTCG CTACGTATAC CCAGGAGCAA GTTGACAAGA TCTTCAAACA GTGTGCCATT
```

121 GCGGCTGCCA AAGAGCGTAT TAACTTGGCC AAACTGGCGG TGGAAGAAAC CGGCATCGGT

181 CTGGTCGAGG ACAAGATTAT CAAGAATCAC TTTGCCGCGG AGTATATCTA CAACAAATAC

```
     241  AAAAACGAAA AGACTTGCGG CATCATTGAT CATGATGACA GCTTGGGTAT CACGAAAGTC
     301  GCAGAACCAA TCGGTATTGT CGCGGCAATT GTCCCGACCA CTAATCCGAC CAGCACCGCG
     361  ATCTTCAAGA GCTTGATCAG CCTGAAAACC CGCAACGCGA TTTTCTTCAG CCCGCACCCG
     421  CGTGCGAAGA AGAGCACCAT CGCAGCGGCG AAGCTGATTT TGGATGCCGC GGTTAAGGCC
     481  GGTGCACCGA AAAACATTAT TGGCTGGATT GACGAGCCGA GCATTGAGCT GTCTCAGGAC
     541  CTGATGAGCG AAGCGGACAT TATTCTGGCC ACCGGTGGTC CGTCTATGGT TAAGGCAGCG
     601  TACTCCAGCG GTAAGCCTGC TATCGGCGTT GGTGCGGGCA ATACGCCGGC TATCATCGAT
     661  GAAAGCGCGG ATATTGATAT GGCAGTTAGC TCCATCATTC TGTCCGCGAC CTACGACAAT
     721  GGTGTTATCT GTGCAAGCGA GCAAAGCATC CTGGTGATGA ATAGCATTTA CGAGAAGGTC
     781  AAAGAAGAGT TTGTTAAGCG TGGTTCCTAT ATTCTGAACC AGAACGAGAT TGCAAAAATC
     841  AAAGAAACGA TGTTTAAGAA TGGCGCGATT AATGCAGATA TTGTTGGTAA GTCCGCATAC
     901  ATTATCGCGA AGATGGCGGG TATCGAAGTG CCGCAAACCA CCAAAATCTT GATCGGCGAA
     961  GTGCAGAGCG TGGAGGCGAG CGAACTGTTC AGCAAAGAGA AACTGAGCCC GGTTCTGGCT
    1021  ATGTACAAAG TGAAGGACTT TGACGAAGCC CTGAAGCTGG CTCAGCGTCT GATTGAGCTG
    1081  GGCGGCAGCG GTCACACGAG CAGCCTGTAT ATCGATAGCC AAAACAATAA GGACAAGGTG
    1141  AAAGAGTTCG GTCTGGCGAT GAAAACCTCT CGCACGTTTA TCAACATGCC GAGCAGCCAA
    1201  GGTGCGAGCG GCGATCTGTA TAACTTCGCA ATCGCGCCTT CTTTTACGCT GGGTTGCGGT
    1261  ACCTGGGGCG GTAATTCGGT GAGCCAGAAT GTCGAGCCGA AGCACCTGCT GAACATCAAA
    1321  TCGGTCGCGG AGCGCCGTGA GAAGAGACCG TGGTTCAAAG TCCCACAGAA AATTTACTTC
    1381  AAATACGGCT GCCTGCGTTT CGCGCTGAAA GAACTGAAAG ACATGAACAA AAAGCGTGCG
    1441  TTCATTGTTA CCGACAAAGA CCTGTTCAAA CTGGGTTACG TGAACAAAAT CACCAAAGTT
    1501  CTGGATGAAA TTGACATCAA GTACTCCATC TTCACTGATA TCAAATCCGA CCCAACGATT
    1561  GATAGCGTGA AAAAGGGCGC TAAAGAAATG CTGAACTTTG AACCGGACAC CATCATCAGC
    1621  ATCGGTGGTG GCTCTCCTAT GGATGCTGCG AAGGTCATGC ACCTGCTGTA CGAATACCCG
    1681  GAAGCGGAAA TCGAAAACCT GGCTATCAAC TTCATGGACA TCCGCAAACG TATCTGCAAC
    1741  TTCCCGAAGC TGGGCACTAA AGCTATTTCC GTTGCCATCC CGACTACCGC GGGCACTGGT
    1801  TCCGAAGCCA CGCCGTTCGC CGTGATCACC AACGATGAAA CCGGTATGAA ATACCCGCTG
    1861  ACCTCTTACG AACTGACCCC GAACATGGCA ATTATCGACA CCGAGCTGAT GCTGAACATG
    1921  CCGCGCAAGC TGACCGCTGC TACCGGCATC GACGCTCTGG TACATGCTAT TGAGGCGTAC
    1981  GTTTCCGTGA TGGCTACCGA TTACACCGAC GAACTGGCCC TGCGTGCGAT CAAAATGATT
    2041  TTCAAGTACC TGCCTCGCGC TTACAAAAAC GGCACGAATG ACATCGAGGC GCGTGAGAAA
    2101  ATGGCCCATG CAAGCAACAT CGCGGGCATG GCCTTCGCCA ACGCGTTCCT GGGCGTGTGC
    2161  CACTCTATGG CTCACAAACT GGGTGCTATG CACCACGTGC CGCACGGTAT CGCGTGTGCT
    2221  GTCCTGATCG AAGAAGTAAT TAAGTACAAC GCTACTGATT GCCCGACTAA ACAGACCGCC
    2281  TTCCCACAGT ACAAATCTCC TAACGCTAAA CGTAAGTACG CTGAGATCGC CGAATACCTG
    2341  AACCTGAAGG GTACGAGCGA CACTGAGAAA GTTACTGCGC TGATCGAAGC TATCTCTAAA
    2401  CTGAAAATTG ACCTGTCCAT CCCGCAGAAC ATCAGCGCCG CAGGCATCAA CAAAAAGGAC
    2461  TTTTACAACA CGCTGGACAA AATGAGCGAA CTGGCTTTTG ACGACCAGTG CACCACTGCA
    2521  AACCCGCGTT ACCCGCTGAT CTCTGAGCTG AAAGACATTT ATATCAAATC TTTTTAA
```

**AdhE2.60 (76989)**

```
       1  ATGAAAGTGA CGAACCAAAA AGAACTGAAA CAAAAACTGA ATGAACTGCG TGAGGCCCAG
      61  AAAAAGTTCG CTACGTATAC CCAGGAGCAA GTTGACAAGA TCTTCAAACA GTGTGCCATT
```

121 GCGGCTGCCA AAGAGCGTAT TAACTTGGCC AAACTGGCGG TGGAAGAAAC CGGCATCGGT

181 CTGGTCGAGG ACAAGATTAT CAAGAATCAC TTTGCCGCGG AGTATATCTA CAACAAATAC

```
     241  AAAAACGAAA AGACTTGCGG CATCATTGAT CATGATGACA GCTTGGGTAT CACGAAAGTC
     301  GCAGAACCAA TCGGTATTGT CGCGGCAATT GTCCCGACCA CTAATCCGAC CAGCACCGCG
     361  ATCTTCAAGA GCTTGATCAG CCTGAAAACC CGCAACGCGA TTTTCTTCAG CCCGCACCCG
     421  CGTGCGAAGA AGAGCACCAT CGCAGCGGCG AAGCTGATTT TGGATGCCGC GGTTAAGGCC
     481  GGTGCACCGA AAAACATTAT TGGCTGGATT GACGAGCCGA GCATTGAGCT GTCTCAGGAC
     541  CTGATGAGCG AAGCGGACAT TATTCTGGCC ACCGGTGGTC CGTCTATGGT TAAGGCAGCG
     601  TACTCCAGCG GTAAGCCTGC TATCGGCGTT GGTGCGGGCA ATACGCCGGC TATCATCGAT
     661  GAAAGCGCGG ATATTGATAT GGCAGTTAGC TCCATCATTC TGTCCAAAAC CTACGACAAT
     721  GGTGTTATCT GTGCAAGCGA GCAAAGCATC CTGGTGATGA ATAGCATTTA CGAGAAGGTC
     781  AAAGAAGAGT TTGTTAAGCG TGGTTCCTAT ATTCTGAACC AGAACGAGAT TGCAAAAATC
     841  AAAGAAACGA TGTTTAAGAA TGGCGCGATT AATGCAGATA TTGTTGGTAA GTCCGCATAC
     901  ATTATCGCGA AGATGGCGGG TATCGAAGTG CCGCAAACCA CCAAAATCTT GATCGGCGAA
     961  GTGCAGAGCG TGGAGGCGAG CGAACTGTTC AGCCATGAGA AACTGAGCCC GGTTCTGGCT
    1021  ATGTACAAAG TGAAGGACTT TGACGAAGCC CTGAAGAAAG CTCAGCGTCT GATTAACCTG
    1081  GGCGGCAGCC ACCACACGAG CAGCCTGTAT ATCGATAGCC AAAACAATAA GGACAAGGTG
    1141  AAAGAGTTCG GTCTGGCGAT GAAAACCTCT CGCACGTTTA TCAACATGCC GAGCAGCCAA
    1201  GGTGCGAGCG GCGATCTGTA TAACTTCGCA ATCGCGCCTT CTTTTACGCT GGGTTGCGGT
    1261  ACCTGGGGCG GTAATTCGGT GAGCCAGAAT GTCGAGCCGA AGCACCTGCT GAACATCGTT
    1321  TCGGTCGCGG AGCGCCGTGA GAAGAGACCG TGGTTCAAAG TCCCACAGAA AATTTACTTC
    1381  AAATACGGCT GCCTGCGTTT CGCGCTGAAA GAACTGAAAG ACATGAACAA AAAGCGTGCG
    1441  TTCATTGTTA CCGACAAAGA CCTGTTCAAA CTGGGTTACG TGAACAAAAT CACCAAAGTT
    1501  CTGGATGAAA TTGACATCAA GTACTCCATC TTCACTGATA TCAAATCCGA CCCAACGATT
    1561  GATAGCGTGA AAAAGGGCGC TAAAGAAATG CTGAACTTTG AACCGGACAC CATCATCAGC
    1621  ATCGGTGGTG GCTCTCCTAT GGATGCTGCG AAGGTCATGC ACCTGCTGTA CGAATACCCG
    1681  GAAGCGGAAA TCGAAAACCT GGCTATCAAC TTCATGGACA TCCGCAAACG TATCTGCAAC
    1741  TTCCCGAAGC TGGGCACTAA AGCTATTTCC GTTGCCATCC CGACTACCGC GGGCACTGGT
    1801  TCCGAAGCCA CGCCGTTCGC CGTGATCACC AACGATGAAA CCGGTATGAA ATACCCGCTG
    1861  ACCTCTTACG AACTGACCCC GAACATGGCA ATTATCGACA CCGAGCTGAT GCTGAACATG
    1921  CCGCGCAAGC TGACCGCTGC TACCGGCATC GACGCTCTGG TACATGCTAT TGAGGCGTAC
    1981  GTTTCCGTGA TGGCTACCGA TTACACCGAC GAACTGGCCC TGCGTGCGAT CAAAATGATT
    2041  TTCAAGTACC TGCCTCGCGC TTACAAAAAC GGCACGAATG ACATCGAGGC GCGTGAGAAA
    2101  ATGGCCCATG CAAGCAACAT CGCGGGCATG GCCTTCGCCA ACGCGTTCCT GGGCGTGTGC
    2161  CACTCTATGG CTCACAAACT GGGTGCTATG CACCACGTGC CGCACGGTAT CGCGTGTGCT
    2221  GTCCTGATCG AAGAAGTAAT TAAGTACAAC GCTACTGATT GCCCGACTAA ACAGACCGCC
    2281  TTCCCACAGT ACAAATCTCC TAACGCTAAA CGTAAGTACG CTGAGATCGC CGAATACCTG
    2341  AACCTGAAGG GTACGAGCGA CACTGAGAAA GTTACTGCGC TGATCGAAGC TATCTCTAAA
    2401  CTGAAAATTG ACCTGTCCAT CCCGCAGAAC ATCAGCGCCG CAGGCATCAA CAAAAAGGAC
    2461  TTTTACAACA CGCTGGACAA AATGAGCGAA CTGGCTTTTG ACGACCAGTG CACCACTGCA
    2521  AACCCGCGTT ACCCGCTGAT CTCTGAGCTG AAAGACATTT ATATCAAATC TTTTTAA
```

**AdhE2.61 (76990)**

```
       1  ATGAAAGTGA CGAACCAAAA AGAACTGAAA CAAAAACTGA ATGAACTGCG TGAGGCCCAG
      61  AAAAAGTTCG CTAGCTATAC CCAGGAGCAA GTTGACAAGA TCTTCAAACA GTGTGCCATT
```

121 GCGGCTGCCA AAGAGCGTAT TAACTTGGCC AAACTGGCGG TGGAAGAAAC CGGCATCGGT

181 CTGGTCGAGG ACAAGATTAT CAAGAATCAC TTTGCCGCGG AGTATATCTA CAACAAATAC

```
     241  AAAAACGAAA AGACTTGCGG CATCATTGAT CATGATGACA GCTTGGGTAT CACGAAAGTC
     301  GCAGAACCAA TCGGTATTGT CGCGGCAATT GTCCCGACCA CTAATCCGAC CAGCACCGCG
     361  ATCTTCAAGA GCTTGATCAG CCTGAAAACC CGCAACGCGA TTTTCTTCAG CCCGCACCCG
     421  CGTGCGAAGA AGAGCACCAT CGCAGCGGCG AAGCTGATTT TGGATGCCGC GGTTAAGGCC
     481  GGTGCACCGA AAAACATTAT TGGCTGGATT GACGAGCCGA GCATTGAGCT GTCTCAGGAC
     541  CTGATGAGCG AAGCGGACAT TATTCTGGCC ACCGGTGGTC CGTCTATGGT TAAGGCAGCG
     601  TACTCCAGCG GTAAGAAAGC TATCGGCGTT GGTGCGGGCA ATACGCCGGC TATCATCGAT
     661  GAAAGCGCGG ATATTGATAT GGCAGTTAGC TCCATCATTC TGTCCAAAAC CTACGACAAT
     721  GGTGTTATCT GTGCAAGCGA GCAAAGCATC CTGGTGATGA ATAGCATTTA CGAGAAGGTC
     781  AAAGAAGAGT TTGTTAAGCG TGGTTCCTAT ATTCTGAACC AGAACGAGAT TGCAAAAATC
     841  AAAGAAACGA TGTTTAAGAA TGGCGCGATT AATGCAGATA TTGTTGGTAA GTCCGCATAC
     901  ATTATCGCGA AGATGGCGGG TATCGAAGTG CCGCAATACA CCAAAATCTT GATCGGCGAA
     961  GTGCAGAGCG TGGAGAAGAG CGAACTGTTC AGCCATGAGA AACTGAGCCC GGTTCTGGCT
    1021  ATGTACAAAG TGAAGGACTT TGACGAAGCC CTGAAGAAAG CTCAGCGTCT GATTGAGCTG
    1081  GGCGGCAGCG GTCACACGAG CAGCCTGTAT ATCGATAGCC AAAACAATAA GGACAAGGTG
    1141  AAAGAGTTCG GTCTGGCGAT GAAAACCTCT CGCTTCTTTA TCAACATGCC GAGCAGCCAA
    1201  GGTGCGAGCG GCGATCTGTA TAACTTCGCA ATCGCGCCTT CTTTTACGCT GGGTTGCGGT
    1261  ACCTGGGGCG GTAATTCGGT GAGCCAGAAT GTCGAGCCGA AGCACCTGCT GAACATCAAA
    1321  TCGGTCGCGG AGCGCCGTGA GAAGAGACCG TGGTTCAAAG TCCCACAGAA AATTTACTTC
    1381  AAATACGGCT GCCTGCGTTT CGCGCTGAAA GAACTGAAAG ACATGAACAA AAAGCGTGCG
    1441  TTCATTGTTA CCGACAAAGA CCTGTTCAAA CTGGGTTACG TGAACAAAAT CACCAAAGTT
    1501  CTGGATGAAA TTGACATCAA GTACTCCATC TTCACTGATA TCAAATCCGA CCCAACGATT
    1561  GATAGCGTGA AAAAGGGCGC TAAAGAAATG CTGAACTTTG AACCGGACAC CATCATCAGC
    1621  ATCGGTGGTG GCTCTCCTAT GGATGCTGCG AAGGTCATGC ACCTGCTGTA CGAATACCCG
    1681  GAAGCGGAAA TCGAAAACCT GGCTATCAAC TTCATGGACA TCCGCAAACG TATCTGCAAC
    1741  TTCCCGAAGC TGGGCACTAA AGCTATTTCC GTTGCCATCC CGACTACCGC GGGCACTGGT
    1801  TCCGAAGCCA CGCCGTTCGC CGTGATCACC AACGATGAAA CCGGTATGAA ATACCCGCTG
    1861  ACCTCTTACG AACTGACCCC GAACATGGCA ATTATCGACA CCGAGCTGAT GCTGAACATG
    1921  CCGCGCAAGC TGACCGCTGC TACCGGCATC GACGCTCTGG TACATGCTAT TGAGGCGTAC
    1981  GTTTCCGTGA TGGCTACCGA TTACACCGAC GAACTGGCCC TGCGTGCGAT CAAAATGATT
    2041  TTCAAGTACC TGCCTCGCGC TTACAAAAAC GGCACGAATG ACATCGAGGC GCGTGAGAAA
    2101  ATGGCCCATG CAAGCAACAT CGCGGGCATG GCCTTCGCCA ACGCGTTCCT GGGCGTGTGC
    2161  CACTCTATGG CTCACAAACT GGGTGCTATG CACCACGTGC CGCACGGTAT CGCGTGTGCT
    2221  GTCCTGATCG AAGAAGTAAT TAAGTACAAC GCTACTGATT GCCCGACTAA ACAGACCGCC
    2281  TTCCCACAGT ACAAATCTCC TAACGCTAAA CGTAAGTACG CTGAGATCGC CGAATACCTG
    2341  AACCTGAAGG GTACGAGCGA CACTGAGAAA GTTACTGCGC TGATCGAAGC TATCTCTAAA
    2401  CTGAAAATTG ACCTGTCCAT CCCGCAGAAC ATCAGCGCCG CAGGCATCAA CAAAAAGGAC
    2461  TTTTACAACA CGCTGGACAA AATGAGCGAA CTGGCTTTTG ACGACCAGTG CACCACTGCA
    2521  AACCCGCGTT ACCCGCTGAT CTCTGAGCTG AAAGACATTT ATATCAAATC TTTTTAA
```

**AdhE2.62 (76991)**

```
       1  ATGAAAGTGA CGAACCAAAA AGAACTGAAA CAAAAACTGA ATGAACTGCG TGAGGCCCAG
      61  AAAAAGTTCG CTACGTATAC CCAGGAGCAA GTTGACAAGA TCTTCAAACA GTGTGCCATT
```

121 GCGGCTGCCA AAGAGCGTAT TAACTTGGCC AAACTGGCGG TGGAAGAAAC CGGCATCGGT

181 CTGGTCGAGG ACAAGATTAT CAAGAATCAC TTTGCCGCGG AGTATATCTA CAACAAATAC

```
     241  AAAAACGAAA AGACTTGCGG CATCATTGAT CATGATGACA GCTTGGGTAT CACGAAAGTC
     301  GCAGAACCAA TCGGTATTGT CGCGGCAATT GTCCCGACCA CTAATCCGAC CAGCACCGCG
     361  ATCTTCAAGA GCTTGATCAG CCTGAAAACC CGCAACGCGA TTTTCTTCAG CCCGCACCCG
     421  CGTGCGAAGA AGAGCACCAT CGCAGCGGCG AAGCTGATTT TGGATGCCGC GGTTAAGGCC
     481  GGTGCACCGA AAAACATTAT TGGCTGGATT GACGAGCCGA GCATTGAGCT GTCTCAGGAC
     541  CTGATGAGCG AAGCGGACAT TATTCTGGCC ACCGGTGGTC CGTCTATGGT TAAGGCAGCG
     601  TACTCCAGCG GTAAGCCTGC TATCGGCGTT GGTGCGGGCA ATACGCCGGC TATCATCGAT
     661  GAAAGCGCGG ATATTGATAT GGCAGTTAGC TCCATCATTC TGTCCAAAAC CTACGACAAT
     721  GGTGTTATCT GTGCAAGCGA GCAAAGCATC CTGGTGATGA ATAGCATTTA CGAGAAGGTC
     781  AAAGAAGAGT TTGTTAAGCG TGGTTCCTAT ATTCTGAACC AGAACGAGAT TGCAAAAATC
     841  AAAGAAACGA TGTTTAAGAA TGGCGCGATT AATGCAGATA TTGTTGGTAA GTCCGCATAC
     901  ATTATCGCGA AGATGGCGGG TATCGAAGTG CCGCAATACA CCAAAATCTT GATCGGCGAA
     961  GTGCAGAGCG TGGAGAAGAG CGAACTGTTC AGCGTTGAGA AACTGAGCCC GGTTCTGGCT
    1021  ATGTACAAAG TGAAGGACTT TGACGAAGCC CTGAAGCTGG CTCAGCGTCT GATTCACCTG
    1081  GGCGGCAGCG GTCACACGAG CAGCCTGTAT ATCGATAGCC AAAACAATAA GGACAAGGTG
    1141  AAAGAGTTCG GTCTGGCGAT GAAAACCTCT CGCACGTTTA TCAACATGCC GAGCAGCCAA
    1201  GGTGCGAGCG GCGATCTGTA TAACTTCGCA ATCGCGCCTT CTTTTACGCT GGGTTGCGGT
    1261  ACCTGGGGCG GTAATTCGGT GAGCCAGAAT GTCGAGCCGA AGCACCTGCT GAACATCAAA
    1321  TCGGTCGCGG AGCGCCGTGA GAAGAGACCG TGGTTCAAAG TCCCACAGAA AATTTACTTC
    1381  AAATACGGCT GCCTGCGTTT CGCGCTGAAA GAACTGAAAG ACATGAACAA AAAGCGTGCG
    1441  TTCATTGTTA CCGACAAAGA CCTGTTCAAA CTGGGTTACG TGAACAAAAT CACCAAAGTT
    1501  CTGGATGAAA TTGACATCAA GTACTCCATC TTCACTGATA TCAAATCCGA CCCAACGATT
    1561  GATAGCGTGA AAAAGGGCGC TAAAGAAATG CTGAACTTTG AACCGGACAC CATCATCAGC
    1621  ATCGGTGGTG GCTCTCCTAT GGATGCTGCG AAGGTCATGC ACCTGCTGTA CGAATACCCG
    1681  GAAGCGGAAA TCGAAAACCT GGCTATCAAC TTCATGGACA TCCGCAAACG TATCTGCAAC
    1741  TTCCCGAAGC TGGGCACTAA AGCTATTTCC GTTGCCATCC CGACTACCGC GGGCACTGGT
    1801  TCCGAAGCCA CGCCGTTCGC CGTGATCACC AACGATGAAA CCGGTATGAA ATACCCGCTG
    1861  ACCTCTTACG AACTGACCCC GAACATGGCA ATTATCGACA CCGAGCTGAT GCTGAACATG
    1921  CCGCGCAAGC TGACCGCTGC TACCGGCATC GACGCTCTGG TACATGCTAT TGAGGCGTAC
    1981  GTTTCCGTGA TGGCTACCGA TTACACCGAC GAACTGGCCC TGCGTGCGAT CAAAATGATT
    2041  TTCAAGTACC TGCCTCGCGC TTACAAAAAC GGCACGAATG ACATCGAGGC GCGTGAGAAA
    2101  ATGGCCCATG CAAGCAACAT CGCGGGCATG GCCTTCGCCA ACGCGTTCCT GGGCGTGTGC
    2161  CACTCTATGG CTCACAAACT GGGTGCTATG CACCACGTGC CGCACGGTAT CGCGTGTGCT
    2221  GTCCTGATCG AAGAAGTAAT TAAGTACAAC GCTACTGATT GCCCGACTAA ACAGACCGCC
    2281  TTCCCACAGT ACAAATCTCC TAACGCTAAA CGTAAGTACG CTGAGATCGC CGAATACCTG
    2341  AACCTGAAGG GTACGAGCGA CACTGAGAAA GTTACTGCGC TGATCGAAGC TATCTCTAAA
    2401  CTGAAAATTG ACCTGTCCAT CCCGCAGAAC ATCAGCGCCG CAGGCATCAA CAAAAAGGAC
    2461  TTTTACAACA CGCTGGACAA AATGAGCGAA CTGGCTTTTG ACGACCAGTG CACCACTGCA
    2521  AACCCGCGTT ACCCGCTGAT CTCTGAGCTG AAAGACATTT ATATCAAATC TTTTTAA
```

**AdhE2.63 (76992)**

```
       1  ATGAAAGTGA CGAACCAAAA AGAACTGAAA CAAAAACTGA ATGAACTGCG TGAGGCCCAG
      61  AAAAAGTTCG CTACGTGCAC CCAGGAGCAA GTTGACAAGA TCTTCAAACA GTGTGCCATT
```

121 GCGGCTGCCA AAGAGCGTAT TAACTTGGCC AAACTGGCGG TGGAAGAAAC CGGCATCGGT

181 CTGGTCGAGG ACAAGATTAT CAAGAATCAC TTTGCCGCGG AGTATATCTA CAACAAAGAG

```
     241  AAAAACGAAA AGACTTGCGG CATCATTGAT CATGATGACA GCTTGGGTAT CACGAAAGTC
     301  GCAGAACCAA TCGGTATTGT CGCGGCAATT GTCCCGACCA CTAATCCGAC CAGCACCGCG
     361  ATCTTCAAGA GCTTGATCAG CCTGAAAACC CGCAACGCGA TTTTCTTCAG CCCGCACCCG
     421  CGTGCGAAGA AGAGCACCAT CGCAGCGGCG AAGCTGATTT TGGATGCCGC GGTTAAGGCC
     481  GGTGCACCGA AAAACATTAT TGGCTGGATT GACGAGCCGA GCATTGAGCT GTCTCAGGAC
     541  CTGATGAGCG AAGCGGACAT TATTCTGGCC ACCGGTGGTC CGTCTATGGT TAAGGCAGCG
     601  TACTCCAGCG GTAAGCCTGC TATCGGCGTT GGTGCGGGCA ATACGCCGGC TATCATCGAT
     661  GAAAGCGCGG ATATTGATAT GGCAGTTAGC TCCATCATTC TGTCCAAAAC CTACGACAAT
     721  GGTGTTATCT GTGCAAGCGA GCAAAGCATC CTGGTGATGA ATAGCATTTA CGAGAAGGTC
     781  AAAGAAGAGT TTGTTAAGCG TGGTTCCTAT ATTCTGAACC AGAACGAGAT TGCAAAAATC
     841  AAAGAAACGA TGTTTAAGAA TGGCGCGATT AATGCAGATA TTGTTGGTAA GTCCGCATAC
     901  ATTATCGCGA AGATGGCGGG TATCGAAGTG CCGCAAACCA CCAAAATCTT GATCGGCGAA
     961  GTGCAGAGCG TGGAGAAGAG CGAACTGTTC AGCCATGAGA AACTGAGCCC GGTTCTGGCT
    1021  ATGTACAAAG TGAAGGACTT TGACGAAGCC CTGGACAAAG CTCAGCGTCT GATTGAGCTG
    1081  GGCGGCAGCG GTCACACGAG CAGCCTGTAT ATCGATAGCC AAAACAATAA GGACAAGGTG
    1141  AAAGAGTTCG GTCTGCGTAT GAAAACCTCT CGCACGTTTA TCAACATGCC GAGCAGCCAA
    1201  GGTGCGAGCG GCGATCTGTA TAACTTCGCA ATCGCGCCTT CTTTTACGCT GGGTTGCGGT
    1261  ACCTGGGGCG GTAATTCGGT GAGCCAGAAT GTCGAGCCGA AGCACCTGCT GAACATCAAA
    1321  TCGGTCGCGG AGCGCCGTGA GAAGAGACCG TGGTTCAAAG TCCCACAGAA AATTTACTTC
    1381  AAATACGGCT GCCTGCGTTT CGCGCTGAAA GAACTGAAAG ACATGAACAA AAAGCGTGCG
    1441  TTCATTGTTA CCGACAAAGA CCTGTTCAAA CTGGGTTACG TGAACAAAAT CACCAAAGTT
    1501  CTGGATGAAA TTGACATCAA GTACTCCATC TTCACTGATA TCAAATCCGA CCCAACGATT
    1561  GATAGCGTGA AAAAGGGCGC TAAAGAAATG CTGAACTTTG AACCGGACAC CATCATCAGC
    1621  ATCGGTGGTG GCTCTCCTAT GGATGCTGCG AAGGTCATGC ACCTGCTGTA CGAATACCCG
    1681  GAAGCGGAAA TCGAAAACCT GGCTATCAAC TTCATGGACA TCCGCAAACG TATCTGCAAC
    1741  TTCCCGAAGC TGGGCACTAA AGCTATTTCC GTTGCCATCC CGACTACCGC GGGCACTGGT
    1801  TCCGAAGCCA CGCCGTTCGC CGTGATCACC AACGATGAAA CCGGTATGAA ATACCCGCTG
    1861  ACCTCTTACG AACTGACCCC GAACATGGCA ATTATCGACA CCGAGCTGAT GCTGAACATG
    1921  CCGCGCAAGC TGACCGCTGC TACCGGCATC GACGCTCTGG TACATGCTAT TGAGGCGTAC
    1981  GTTTCCGTGA TGGCTACCGA TTACACCGAC GAACTGGCCC TGCGTGCGAT CAAAATGATT
    2041  TTCAAGTACC TGCCTCGCGC TTACAAAAAC GGCACGAATG ACATCGAGGC GCGTGAGAAA
    2101  ATGGCCCATG CAAGCAACAT CGCGGGCATG GCCTTCGCCA ACGCGTTCCT GGGCGTGTGC
    2161  CACTCTATGG CTCACAAACT GGGTGCTATG CACCACGTGC CGCACGGTAT CGCGTGTGCT
    2221  GTCCTGATCG AAGAAGTAAT TAAGTACAAC GCTACTGATT GCCCGACTAA ACAGACCGCC
    2281  TTCCCACAGT ACAAATCTCC TAACGCTAAA CGTAAGTACG CTGAGATCGC CGAATACCTG
    2341  AACCTGAAGG GTACGAGCGA CACTGAGAAA GTTACTGCGC TGATCGAAGC TATCTCTAAA
    2401  CTGAAAATTG ACCTGTCCAT CCCGCAGAAC ATCAGCGCCG CAGGCATCAA CAAAAAGGAC
    2461  TTTTACAACA CGCTGGACAA AATGAGCGAA CTGGCTTTTG ACGACCAGTG CACCACTGCA
    2521  AACCCGCGTT ACCCGCTGAT CTCTGAGCTG AAAGACATTT ATATCAAATC TTTTTAA
```

**AdhE2.64 (76993)**

```
       1  ATGAAAGTGA CGAACCAAAA AGAACTGAAA CAAAAACTGA ATGAACTGCG TGAGGCCCAG
      61  AAAAAGTTCG CTACGTATAC CCAGGAGCAA GTTGACAAGA TCTTCAAACA GTGTGCCATT
```

121 GCGGCTGCCA AACTGCGTAT TAACTTGGCC AAACTGGCGG TGGAAGAAAC CGGCATCGGT

181 CTGGTCGAGG ACAAGATTAT CAAGAATCAC TTTGCCGCGG AGTATATCTA CAACAAATAC

```
     241  AAAAACGAAA AGACTTGCGG CATCATTGAT CATGATGACA GCTTGGGTAT CACGAAAGTC
     301  GCAGAACCAA TCGGTATTGT CGCGGCAATT GTCCCGACCA CTAATCCGAC CAGCACCGCG
     361  ATCTTCAAGA GCTTGATCAG CCTGAAAACC CGCAACGCGA TTTTCTTCAG CCCGCACCCG
     421  CGTGCGAAGA AGAGCACCAT CGCAGCGGCG AAGCTGATTT TGGATGCCGC GGTTAAGGCC
     481  GGTGCACCGA AAAACATTAT TGGCTGGATT GACGAGCCGA GCATTGAGCT GTCTCAGGAC
     541  CTGATGAGCG AAGCGGACAT TATTCTGGCC ACCGGTGGTC CGTCTATGGT TAAGGCAGCG
     601  TACTCCAGCG GTAAGCCTGC TATCGGCGTT GGTGCGGGCA ATACGCCGGC TATCATCGAT
     661  GAAAGCGCGG ATATTGATAT GGCAGTTAGC TCCATCATTC TGTCCAAAAC CTACGACAAT
     721  GGTGTTATCT GTGCAAGCGA GCAAAGCATC CTGGTGATGA ATAGCATTTA CGAGAAGGTC
     781  AAAGAAGAGT TTGTTAAGCG TGGTTCCTAT ATTCTGAACC AGAACGAGAT TGCAAAAATC
     841  AAAGAAACGA TGTTTAAGAA TGGCGCGATT AATGCAGATA TTGTTGGTAA GTCCGCATAC
     901  ATTATCGCGA AGATGGCGGG TATCGAAGTG CCGCAAACCA CCAAAATCTT GATCGGCGAA
     961  GTGCAGAGCG TGGAGAAGAG CGAACTGTTC AGCCATGAGA AACTGAGCCC GGTTCTGGCT
    1021  ATGTACAAAG TGAAGGACTT TGACGAAGCC CTGAAGAAAG CTGAGCGTCT GATTGAGCTG
    1081  GGCGGCAGCG GTCACACGAG CAGCCTGTAT ATCGATAGCC AAAACAATAA GGACAAGGTG
    1141  AAAGAGTTCG GTCTGCGTAT GAAAACCTCT CGCACGTTTA TCAACATGCC GAGCAGCCAA
    1201  GGTGCGAGCG GCGATCTGTA TAACTTCGCA ATCGCGCCTT CTTTTACGCT GGCGTGCGGT
    1261  ACCTGGGGCG GTAATTCGGT GAGCCAGAAT GTCGAGCCGA AGCACCTGCT GAACATCAAA
    1321  TCGGTCGCGG AGCGCCGTGA GAAGAGACCG TGGTTCAAAG TCCCACAGAA AATTTACTTC
    1381  AAATACGGCT GCCTGCGTTT CGCGCTGAAA GAACTGAAAG ACATGAACAA AAAGCGTGCG
    1441  TTCATTGTTA CCGACAAAGA CCTGTTCAAA CTGGGTTACG TGAACAAAAT CACCAAAGTT
    1501  CTGGATGAAA TTGACATCAA GTACTCCATC TTCACTGATA TCAAATCCGA CCCAACGATT
    1561  GATAGCGTGA AAAAGGGCGC TAAAGAAATG CTGAACTTTG AACCGGACAC CATCATCAGC
    1621  ATCGGTGGTG GCTCTCCTAT GGATGCTGCG AAGGTCATGC ACCTGCTGTA CGAATACCCG
    1681  GAAGCGGAAA TCGAAAACCT GGCTATCAAC TTCATGGACA TCCGCAAACG TATCTGCAAC
    1741  TTCCCGAAGC TGGGCACTAA AGCTATTTCC GTTGCCATCC CGACTACCGC GGGCACTGGT
    1801  TCCGAAGCCA CGCCGTTCGC CGTGATCACC AACGATGAAA CCGGTATGAA ATACCCGCTG
    1861  ACCTCTTACG AACTGACCCC GAACATGGCA ATTATCGACA CCGAGCTGAT GCTGAACATG
    1921  CCGCGCAAGC TGACCGCTGC TACCGGCATC GACGCTCTGG TACATGCTAT TGAGGCGTAC
    1981  GTTTCCGTGA TGGCTACCGA TTACACCGAC GAACTGGCCC TGCGTGCGAT CAAAATGATT
    2041  TTCAAGTACC TGCCTCGCGC TTACAAAAAC GGCACGAATG ACATCGAGGC GCGTGAGAAA
    2101  ATGGCCCATG CAAGCAACAT CGCGGGCATG GCCTTCGCCA ACGCGTTCCT GGGCGTGTGC
    2161  CACTCTATGG CTCACAAACT GGGTGCTATG CACCACGTGC CGCACGGTAT CGCGTGTGCT
    2221  GTCCTGATCG AAGAAGTAAT TAAGTACAAC GCTACTGATT GCCCGACTAA ACAGACCGCC
    2281  TTCCCACAGT ACAAATCTCC TAACGCTAAA CGTAAGTACG CTGAGATCGC CGAATACCTG
    2341  AACCTGAAGG GTACGAGCGA CACTGAGAAA GTTACTGCGC TGATCGAAGC TATCTCTAAA
    2401  CTGAAAATTG ACCTGTCCAT CCCGCAGAAC ATCAGCGCCG CAGGCATCAA CAAAAAGGAC
    2461  TTTTACAACA CGCTGGACAA AATGAGCGAA CTGGCTTTTG ACGACCAGTG CACCACTGCA
    2521  AACCCGCGTT ACCCGCTGAT CTCTGAGCTG AAAGACATTT ATATCAAATC TTTTTAA
```

**AdhE2.65 (76994)**

```
       1  ATGAAAGTGA CGAACCAAAA AGAACTGAAA CAAAAACTGA ATGAACTGCG TGAGGCCCAG
      61  AAAAAGTTCG CTACGTATAC CCAGGAGCAA GTTGACAAGA TCTTCAAACA GTGTGCCATT
```

121 GCGGCTGCCA AAGAGCGTAT TAACTTGGCC AAACTGGCGG TGGAAGAAAC CGGCATCGGT

181 CTGGTCGAGG ACAAGATTAT CAAGAATCAC TTTGCCGCGG AGTATATCTA CAACAAATAC

```
     241  AAAAACGAAA AGACTTGCGG CATCATTGAT CATGATGACA GCTTGGGTAT CACGAAAGTC
     301  GCAGAACCAA TCGGTATTGT CGCGGCAATT GTCCCGACCA CTAATCCGAC CAGCACCGCG
     361  ATCTTCAAGA GCTTGATCAG CCTGAAAACC CGCAACGCGA TTTTCTTCAG CCCGCACCCG
     421  CGTGCGAAGA AGAGCACCAT CGCAGCGGCG AAGCTGATTT TGGATGCCGC GGTTAAGGCC
     481  GGTGCACCGA AAAACATTAT TGGCTGGATT GACGAGCCGA GCATTGAGCT GTCTCAGGAC
     541  CTGATGAGCG AAGCGGACAT TATTCTGGCC ACCGGTGGTC CGTCTATGGT TAAGGCAGCG
     601  TACTCCAGCG GTAAGCCTGC TATCGGCGTT GGTGCGGGCA ATACGCCGGC TATCATCGAT
     661  GAAAGCGCGG ATATTGATAT GGCAGTTAGC TCCATCATTC TGGGTAAAAC CTACGACAAT
     721  GGTGTTATCT GTGCAAGCGA GCAAAGCATC CTGGTGATGA ATAGCATTTA CGAGAAGGTC
     781  AAAGCGGAGT TTGTTAAGCG TGGTTCCTAT ATTCTGAACC AGAACGAGAT TGCAAAAATC
     841  AAAGAAACGA TGTTTAAGAA TGGCGCGATT AATGCAGATA TTGTTGGTAA GTCCGCATAC
     901  ATTATCGCGA AGATGGCGGG TATCGAAGTG CCGCAAACCA CCAAAATCTT GATCGGCGAA
     961  GTGCAGCACG TGGAGAAGAG CGAACTGTTC AGCCATGAGA AACTGAGCCC GGTTCTGGCT
    1021  ATGTACAAAG TGAAGGACTT TGACGAAGCC CTGAAGAAAG CTCAGCGTCT GATTGAGCTG
    1081  GGCGGCAGCG GTCACACGAG CAGCCTGTAT ATCGATAGCC AAAACAATAA GGACAAGGTG
    1141  AAAGAGTTCG GTCTGGCGAT GAAAACCTCT ATCACGTTTA TCAACATGCC GAGCAGCCAA
    1201  GGTGCGAGCG GCGATCTGTA TAACTTCGCA ATCGCGCCTT CTTTTACGCT GGGTTGCGGT
    1261  ACCTGGGGCG GTAATTCGGT GAGCCAGAAT GTCGAGCCGA AGCACCTGCT GAACATCAAA
    1321  TCGGTCGCGG AGCGCCGTGA GAAGAGACCG TGGTTCAAAG TCCCACAGAA AATTTACTTC
    1381  AAATACGGCT GCCTGCGTTT CGCGCTGAAA GAACTGAAAG ACATGAACAA AAAGCGTGCG
    1441  TTCATTGTTA CCGACAAAGA CCTGTTCAAA CTGGGTTACG TGAACAAAAT CACCAAAGTT
    1501  CTGGATGAAA TTGACATCAA GTACTCCATC TTCACTGATA TCAAATCCGA CCCAACGATT
    1561  GATAGCGTGA AAAAGGGCGC TAAAGAAATG CTGAACTTTG AACCGGACAC CATCATCAGC
    1621  ATCGGTGGTG GCTCTCCTAT GGATGCTGCG AAGGTCATGC ACCTGCTGTA CGAATACCCG
    1681  GAAGCGGAAA TCGAAAACCT GGCTATCAAC TTCATGGACA TCCGCAAACG TATCTGCAAC
    1741  TTCCCGAAGC TGGGCACTAA AGCTATTTCC GTTGCCATCC CGACTACCGC GGGCACTGGT
    1801  TCCGAAGCCA CGCCGTTCGC CGTGATCACC AACGATGAAA CCGGTATGAA ATACCCGCTG
    1861  ACCTCTTACG AACTGACCCC GAACATGGCA ATTATCGACA CCGAGCTGAT GCTGAACATG
    1921  CCGCGCAAGC TGACCGCTGC TACCGGCATC GACGCTCTGG TACATGCTAT TGAGGCGTAC
    1981  GTTTCCGTGA TGGCTACCGA TTACACCGAC GAACTGGCCC TGCGTGCGAT CAAAATGATT
    2041  TTCAAGTACC TGCCTCGCGC TTACAAAAAC GGCACGAATG ACATCGAGGC GCGTGAGAAA
    2101  ATGGCCCATG CAAGCAACAT CGCGGGCATG GCCTTCGCCA ACGCGTTCCT GGGCGTGTGC
    2161  CACTCTATGG CTCACAAACT GGGTGCTATG CACCACGTGC CGCACGGTAT CGCGTGTGCT
    2221  GTCCTGATCG AAGAAGTAAT TAAGTACAAC GCTACTGATT GCCCGACTAA ACAGACCGCC
    2281  TTCCCACAGT ACAAATCTCC TAACGCTAAA CGTAAGTACG CTGAGATCGC CGAATACCTG
    2341  AACCTGAAGG GTACGAGCGA CACTGAGAAA GTTACTGCGC TGATCGAAGC TATCTCTAAA
    2401  CTGAAAATTG ACCTGTCCAT CCCGCAGAAC ATCAGCGCCG CAGGCATCAA CAAAAAGGAC
    2461  TTTTACAACA CGCTGGACAA AATGAGCGAA CTGGCTTTTG ACGACCAGTG CACCACTGCA
    2521  AACCCGCGTT ACCCGCTGAT CTCTGAGCTG AAAGACATTT ATATCAAATC TTTTTAA
```

**AdhE2.66 (76995	)**

```
       1  ATGAAAGTGA CGAACCAAAA AGAACTGAAA CAAAAACTGA ATGAACTGCG TGAGGCCCAG
      61  AAAAAGTTCG CTACGTATAC CCAGGAGCAA GTTGACAAGA TCTTCAAACA GTGTGCCATT
```

121 GCGGCTGCCA AAGAGCGTAT TAACTTGGCC AAACTGGCGG TGGAAGAAAC CGGCATCGGT

181 CTGGTCGAGG ACAAGATTAT CAAGAATCAC TTTGCCGCGG AGTATATCTA CAACAAATAC

```
     241  AAAAACGAAA AGACTTGCGG CATCATTGAT CATGATGACA GCTTGGGTAT CACGAAAGTC
     301  GCAGAACCAA TCGGTATTGT CGCGGCAATT GTCCCGACCA CTAATCCGAC CAGCACCGCG
     361  ATCTTCAAGA GCTTGATCAG CCTGAAAACC CGCAACGCGA TTTTCTTCAG CCCGCACCCG
     421  CGTGCGAAGA AGAGCACCAT CGCAGCGGCG AAGCTGATTT TGGATGCCGC GGTTAAGGCC
     481  GGTGCACCGA AAAACATTAT TGGCTGGATT GACGAGCCGA GCATTGAGCT GTCTCAGGAC
     541  CTGATGAGCG AAGCGGACAT TATTCTGGCC ACCGGTGGTC CGTCTATGGT TAAGGCAGCG
     601  TACTCCAGCG GTAAGCCTGC TATCGGCGTT GGTGCGGGCA ATACGCCGGC TATCATCGAT
     661  GAAAGCGCGG ATATTGATAT GGCAGTTAGC TCCATCATTC TGTCCGCGAC CTACGACAAT
     721  GGTGTTATCT GTGCAAGCGA GCAAAGCATC CTGGTGATGA ATAGCATTTA CGAGAAGGTC
     781  AAAGAAGAGT TTGTTAAGCG TGGTTCCTAT ATTCTGAACC AGAACGAGAT TGCAAAAATC
     841  AAAGAAACGA TGTTTAAGAA TGGCGCGATT AATGCAGATA TTGTTGGTAA GTCCGCATAC
     901  ATTATCGCGA AGATGGCGGG TATCGAAGTG CCGCAAACCA CCAAAATCTT GATCGGCGAA
     961  GTGCAGAGCG TGGAGAAGAG CGAACTGTTC AGCCATGAGC TGCTGAGCCC GGTTCTGGCT
    1021  ATGTACAAAG TGAAGGACTT TGACGAAGCC CTGAAGAAAG CTCAGCGTCT GATTCACCTG
    1081  GGCGGCAGCG GTCACACGAG CAGCCTGTAT ATCGATAGCC AAAACAATAA GGACAAGGTG
    1141  AAAGAGTTCG GTCTGGCGAT GAAAACCTCT ATCACGTTTA TCAACATGCC GAGCAGCCAA
    1201  GGTGCGAGCG GCGATCTGTA TAACTTCGCA ATCGCGCCTT CTTTTACGCT GGGTTGCGGT
    1261  ACCTGGGGCG GTAATTCGGT GAGCCAGAAT GTCGAGCCGA AGCACCTGCT GAACATCAAA
    1321  TCGGTCGCGG AGCGCCGTGA GAAGAGACCG TGGTTCAAAG TCCCACAGAA AATTTACTTC
    1381  AAATACGGCT GCCTGCGTTT CGCGCTGAAA GAACTGAAAG ACATGAACAA AAAGCGTGCG
    1441  TTCATTGTTA CCGACAAAGA CCTGTTCAAA CTGGGTTACG TGAACAAAAT CACCAAAGTT
    1501  CTGGATGAAA TTGACATCAA GTACTCCATC TTCACTGATA TCAAATCCGA CCCAACGATT
    1561  GATAGCGTGA AAAAGGGCGC TAAAGAAATG CTGAACTTTG AACCGGACAC CATCATCAGC
    1621  ATCGGTGGTG GCTCTCCTAT GGATGCTGCG AAGGTCATGC ACCTGCTGTA CGAATACCCG
    1681  GAAGCGGAAA TCGAAAACCT GGCTATCAAC TTCATGGACA TCCGCAAACG TATCTGCAAC
    1741  TTCCCGAAGC TGGGCACTAA AGCTATTTCC GTTGCCATCC CGACTACCGC GGGCACTGGT
    1801  TCCGAAGCCA CGCCGTTCGC CGTGATCACC AACGATGAAA CCGGTATGAA ATACCCGCTG
    1861  ACCTCTTACG AACTGACCCC GAACATGGCA ATTATCGACA CCGAGCTGAT GCTGAACATG
    1921  CCGCGCAAGC TGACCGCTGC TACCGGCATC GACGCTCTGG TACATGCTAT TGAGGCGTAC
    1981  GTTTCCGTGA TGGCTACCGA TTACACCGAC GAACTGGCCC TGCGTGCGAT CAAAATGATT
    2041  TTCAAGTACC TGCCTCGCGC TTACAAAAAC GGCACGAATG ACATCGAGGC GCGTGAGAAA
    2101  ATGGCCCATG CAAGCAACAT CGCGGGCATG GCCTTCGCCA ACGCGTTCCT GGGCGTGTGC
    2161  CACTCTATGG CTCACAAACT GGGTGCTATG CACCACGTGC CGCACGGTAT CGCGTGTGCT
    2221  GTCCTGATCG AAGAAGTAAT TAAGTACAAC GCTACTGATT GCCCGACTAA ACAGACCGCC
    2281  TTCCCACAGT ACAAATCTCC TAACGCTAAA CGTAAGTACG CTGAGATCGC CGAATACCTG
    2341  AACCTGAAGG GTACGAGCGA CACTGAGAAA GTTACTGCGC TGATCGAAGC TATCTCTAAA
    2401  CTGAAAATTG ACCTGTCCAT CCCGCAGAAC ATCAGCGCCG CAGGCATCAA CAAAAAGGAC
    2461  TTTTACAACA CGCTGGACAA AATGAGCGAA CTGGCTTTTG ACGACCAGTG CACCACTGCA
    2521  AACCCGCGTT ACCCGCTGAT CTCTGAGCTG AAAGACATTT ATATCAAATC TTTTTAA
```

**AdhE2.67 (76996)**

```
       1  ATGAAAGTGA CGAACCAAAA AGAACTGAAA CAAAAACTGA ATGAACTGCG TGAGGCCCAG
      61  AAAAAGTTCG CTACGTATAC CCAGGAGCAA GTTGACAAGA TCTTCAAACA GTGTGCCATT
```

121 GCGGCTGCCA AAGAGCGTAT TAACTTGGCC AAACTGGCGG TGGAAGAAAC CGGCATCGGT

181 CTGGTCGAGG ACAAGATTAT CAAGAATCAC TTTGCCGCGG AGTATATCTA CAACAAATAC

```
     241  AAAAACGAAA AGACTTGCGG CATCATTGAT CATGATGACA GCTTGGGTAT CACGAAAGTC
     301  GCAGAACCAA TCGGTATTGT CGCGGCAATT GTCCCGACCA CTAATCCGAC CAGCACCGCG
     361  ATCTTCAAGA GCTTGATCAG CCTGAAAACC CGCAACGCGA TTTTCTTCAG CCCGCACCCG
     421  CGTGCGAAGA AGAGCACCAT CGCAGCGGCG AAGCTGATTT TGGATGCCGC GGTTAAGGCC
     481  GGTGCACCGA AAAACATTAT TGGCTGGATT GACGAGCCGA GCATTGAGCT GTCTCAGGAC
     541  CTGATGAGCG AAGCGGACAT TATTCTGGCC ACCGGTGGTC CGTCTATGGT TAAGGCAGCG
     601  TACTCCAGCG GTAAGAAAGC TATCGGCGTT GGTGCGGGCA ATACGCCGGC TATCATCGAT
     661  GAAAGCGCGG ATATTGATAT GGCAGTTAGC TCCATCATTC TGTCCAAAAC CTACGACAAT
     721  GGTGTTATCT GTGCAAGCGA GCAAAGCATC CTGGTGATGA ATAGCATTTA CGAGAAGGTC
     781  AAAAAAGAGT TTGTTAAGCG TGGTTCCTAT ATTCTGAACC AGAACGCGAT TGCAAAAATC
     841  AAAGAAACGA TGTTTAAGAA TGGCGCGATT AATGCAGATA TTGTTGGTAA GTCCGCATAC
     901  ATTATCGCGA AGATGGCGGG TATCGAAGTG CCGCAAACCA CCAAAATCTT GATCGGCGAA
     961  GTGCAGAGCG TGGAGAAGAG CGAACTGTTC AGCCATGAGC TGCTGAGCCC GGTTCTGGCT
    1021  ATGTACAAAG TGAAGGACTT TGACGAAGCC CTGAAGAAAG CTCAGCGTCT GATTGAGCTG
    1081  GGCGGCAGCG GTCACACGAG CAGCCTGTAT ATCGATAGCC AAAACAATAA GGACAAGGTG
    1141  AAAGAGTTCG GTCTGGCGAT GAAAACCTCT CGCACGTTTA TCAACATGCC GAGCAGCCAA
    1201  GGTGCGAGCG GCGATCTGTA TAACTTCGCA ATCGCGCCTT CTTTTACGCT GGGTTGCGGT
    1261  ACCTGGGGCG GTAATTCGGT GAGCCAGAAT GTCGAGCCGA AGCACCTGCT GAACATCAAA
    1321  TCGGTCGCGG AGCGCCGTGA GAAGAGACCG TGGTTCAAAG TCCCACAGAA AATTTACTTC
    1381  AAATACGGCT GCCTGCGTTT CGCGCTGAAA GAACTGAAAG ACATGAACAA AAAGCGTGCG
    1441  TTCATTGTTA CCGACAAAGA CCTGTTCAAA CTGGGTTACG TGAACAAAAT CACCAAAGTT
    1501  CTGGATGAAA TTGACATCAA GTACTCCATC TTCACTGATA TCAAATCCGA CCCAACGATT
    1561  GATAGCGTGA AAAAGGGCGC TAAAGAAATG CTGAACTTTG AACCGGACAC CATCATCAGC
    1621  ATCGGTGGTG GCTCTCCTAT GGATGCTGCG AAGGTCATGC ACCTGCTGTA CGAATACCCG
    1681  GAAGCGGAAA TCGAAAACCT GGCTATCAAC TTCATGGACA TCCGCAAACG TATCTGCAAC
    1741  TTCCCGAAGC TGGGCACTAA AGCTATTTCC GTTGCCATCC CGACTACCGC GGGCACTGGT
    1801  TCCGAAGCCA CGCCGTTCGC CGTGATCACC AACGATGAAA CCGGTATGAA ATACCCGCTG
    1861  ACCTCTTACG AACTGACCCC GAACATGGCA ATTATCGACA CCGAGCTGAT GCTGAACATG
    1921  CCGCGCAAGC TGACCGCTGC TACCGGCATC GACGCTCTGG TACATGCTAT TGAGGCGTAC
    1981  GTTTCCGTGA TGGCTACCGA TTACACCGAC GAACTGGCCC TGCGTGCGAT CAAAATGATT
    2041  TTCAAGTACC TGCCTCGCGC TTACAAAAAC GGCACGAATG ACATCGAGGC GCGTGAGAAA
    2101  ATGGCCCATG CAAGCAACAT CGCGGGCATG GCCTTCGCCA ACGCGTTCCT GGGCGTGTGC
    2161  CACTCTATGG CTCACAAACT GGGTGCTATG CACCACGTGC CGCACGGTAT CGCGTGTGCT
    2221  GTCCTGATCG AAGAAGTAAT TAAGTACAAC GCTACTGATT GCCCGACTAA ACAGACCGCC
    2281  TTCCCACAGT ACAAATCTCC TAACGCTAAA CGTAAGTACG CTGAGATCGC CGAATACCTG
    2341  AACCTGAAGG GTACGAGCGA CACTGAGAAA GTTACTGCGC TGATCGAAGC TATCTCTAAA
    2401  CTGAAAATTG ACCTGTCCAT CCCGCAGAAC ATCAGCGCCG CAGGCATCAA CAAAAAGGAC
    2461  TTTTACAACA CGCTGGACAA AATGAGCGAA CTGGCTTTTG ACGACCAGTG CACCACTGCA
    2521  AACCCGCGTT ACCCGCTGAT CTCTGAGCTG AAAGACATTT ATATCAAATC TTTTTAA
```

**AdhE2.68 (76997)**

```
       1  ATGAAAGTGA CGAACCAAAA AGAACTGAAA CAAAAACTGA ATGAACTGCG TGAGGCCCAG
      61  AAAAAGTTCG CTACGTATAC CCAGGAGCAA GTTGACAAGA TCTTCAAACA GTGTGCCATT
```

121 GCGGCTGCCA AAGAGCGTAT TAACTTGGCC AAACTGGCGG TGGAAGAAAC CGGCATCGGT

181 CTGGTCGAGG ACAAGATTAT CAAGAATCAC TTTGCCGCGG AGTATATCTA CAACAAATAC

```
     241  AAAAACGAAA AGACTTGCGG CATCATTGAT CATGATGACA GCTTGGGTAT CACGAAAGTC
     301  GCAGAACCAA TCGGTATTGT CGCGGCAATT GTCCCGACCA CTAATCCGAC CAGCACCGCG
     361  ATCTTCAAGA GCTTGATCAG CCTGAAAACC CGCAACGCGA TTTTCTTCAG CCCGCACCCG
     421  CGTGCGAAGA AGAGCACCAT CGCAGCGGCG AAGCTGATTT TGGATGCCGC GGTTAAGGCC
     481  GGTGCACCGA AAAACATTAT TGGCTGGATT GACGAGCCGA GCATTGAGCT GTCTCAGGAC
     541  CTGATGAAAG AAGCGGACAT TATTCTGGCC ACCGGTGGTC CGTCTATGGT TAAGGCAGCG
     601  TACTCCAGCG GTAAGCCTGC TATCGGCGTT GGTGCGGGCA ATACGCCGGC TATCATCGAT
     661  GAAAGCGCGG ATATTGATAT GGCAGTTAGC TCCATCATTC TGTCCAAAAC CTACGACAAT
     721  GGTGTTATCT GTGCAAGCGA GCAAAGCATC CTGGTGATGA ATAGCATTTA CGAGAAGGTC
     781  AAAGAAGAGT TTGTTAAGCG TGGTTCCTAT ATTCTGAACC AGAACGAGAT TGCAAAAATC
     841  AAAGAAACGA TGTTTAAGAA TGGCGCGATT AATGCAGATA TTGTTGGTAA GTCCGCATAC
     901  ATTATCGCGA AGATGGCGGG TATCGAAGTG CCGCAAACCA CCAAAATCTT GATCGGCGAA
     961  GTGCAGAGCG TGGAGAAGAG CGAACTGTTC AGCGAGGAGA AACTGAGCCC GGTTCTGGCT
    1021  ATGTACAAAG TGAAGGACTT TGACGAAGCC CTGAAGAAAG CTCAGCGTCT GATTGAGCTG
    1081  GGCGGCAGCG GTCACACGAG CAGCCTGTAT ATCGATAGCC AAAACAATAA GGACAAGGTG
    1141  AAAGAGTTCG GTCTGGAGAT GAAAACCTCT CGCACGTTTA TCAACATGCC GAGCAGCCAA
    1201  GGTGCGAGCG GCGATCTGTA TAACTTCGCA ATCGCGCCTT CTTTTACGCT GGGTTGCGGT
    1261  ACCTGGGAGG GTAATTCGGT GAGCCAGAAT GTCGAGCCGA AGCACCTGCT GAACATCAAA
    1321  TCGGTCGCGG AGCGCCGTGA GAAGAGACCG TGGTTCAAAG TCCCACAGAA AATTTACTTC
    1381  AAATACGGCT GCCTGCGTTT CGCGCTGAAA GAACTGAAAG ACATGAACAA AAAGCGTGCG
    1441  TTCATTGTTA CCGACAAAGA CCTGTTCAAA CTGGGTTACG TGAACAAAAT CACCAAAGTT
    1501  CTGGATGAAA TTGACATCAA GTACTCCATC TTCACTGATA TCAAATCCGA CCCAACGATT
    1561  GATAGCGTGA AAAAGGGCGC TAAAGAAATG CTGAACTTTG AACCGGACAC CATCATCAGC
    1621  ATCGGTGGTG GCTCTCCTAT GGATGCTGCG AAGGTCATGC ACCTGCTGTA CGAATACCCG
    1681  GAAGCGGAAA TCGAAAACCT GGCTATCAAC TTCATGGACA TCCGCAAACG TATCTGCAAC
    1741  TTCCCGAAGC TGGGCACTAA AGCTATTTCC GTTGCCATCC CGACTACCGC GGGCACTGGT
    1801  TCCGAAGCCA CGCCGTTCGC CGTGATCACC AACGATGAAA CCGGTATGAA ATACCCGCTG
    1861  ACCTCTTACG AACTGACCCC GAACATGGCA ATTATCGACA CCGAGCTGAT GCTGAACATG
    1921  CCGCGCAAGC TGACCGCTGC TACCGGCATC GACGCTCTGG TACATGCTAT TGAGGCGTAC
    1981  GTTTCCGTGA TGGCTACCGA TTACACCGAC GAACTGGCCC TGCGTGCGAT CAAAATGATT
    2041  TTCAAGTACC TGCCTCGCGC TTACAAAAAC GGCACGAATG ACATCGAGGC GCGTGAGAAA
    2101  ATGGCCCATG CAAGCAACAT CGCGGGCATG GCCTTCGCCA ACGCGTTCCT GGGCGTGTGC
    2161  CACTCTATGG CTCACAAACT GGGTGCTATG CACCACGTGC CGCACGGTAT CGCGTGTGCT
    2221  GTCCTGATCG AAGAAGTAAT TAAGTACAAC GCTACTGATT GCCCGACTAA ACAGACCGCC
    2281  TTCCCACAGT ACAAATCTCC TAACGCTAAA CGTAAGTACG CTGAGATCGC CGAATACCTG
    2341  AACCTGAAGG GTACGAGCGA CACTGAGAAA GTTACTGCGC TGATCGAAGC TATCTCTAAA
    2401  CTGAAAATTG ACCTGTCCAT CCCGCAGAAC ATCAGCGCCG CAGGCATCAA CAAAAAGGAC
    2461  TTTTACAACA CGCTGGACAA AATGAGCGAA CTGGCTTTTG ACGACCAGTG CACCACTGCA
    2521  AACCCGCGTT ACCCGCTGAT CTCTGAGCTG AAAGACATTT ATATCAAATC TTTTTAA
```

**AdhE2.69 (76998)**

```
       1  ATGAAAGTGA CGAACCAAAA AGAACTGAAA CAAAAACTGA ATGAACTGCG TGAGGCCCAG
      61  AAAAAGTTCG CTACGTATAC CCAGGACCAA GTTGACAAGA TCTTCAAACA GTGTGCCATT
```

121 GCGGCTGCCA AAGAGCGTAT TAACTTGGCC AAACTGGCGG TGGAAGAAAC CGGCATCGGT

181 CTGGTCGAGG ACAAGATTAT CAAGAATCAC TTTGCCGCGG AGTATATCTA CAACAAATAC

```
     241  AAAAACGAAA AGACTTGCGG CATCATTGAT CATGATGACA GCTTGGGTAT CACGAAAGTC
     301  GCAGAACCAA TCGGTATTGT CGCGGCAATT GTCCCGACCA CTAATCCGAC CAGCACCGCG
     361  ATCTTCAAGA GCTTGATCAG CCTGAAAACC CGCAACGCGA TTTTCTTCAG CCCGCACCCG
     421  CGTGCGAAGA AGAGCACCAT CGCAGCGGCG AAGCTGATTT TGGATGCCGC GGTTAAGGCC
     481  GGTGCACCGA AAAACATTAT TGGCTGGATT GACGAGCCGA GCATTGAGCT GTCTCAGGAC
     541  CTGATGAGCG AAGCGGACAT TATTCTGGCC ACCGGTGGTC CGTCTATGGT TAAGGCAGCG
     601  TACTCCAGCG GTAAGCCTGC TATCGGCGTT GGTGCGGGCA ATACGCCGGC TATCATCGAT
     661  GAAAGCGCGG ATATTGATAT GGCAGTTAGC TCCATCATTC TGTCCAAAAC CTACGACAAT
     721  GGTGTTATCT GTGCAAGCGA GCAAAGCATC CTGGTGATGA ATAGCATTTA CGAGAAGGTC
     781  AAAGAAGAGT TTGTTAAGCG TGGTTCCTAT ATTCTGAACC AGAACGAGAT TGCAAAAATC
     841  AAAGAAACGA TGTTTAAGAA TGGCGCGATT AATGCAGATA TTGTTGGTAA GTCCGCATAC
     901  ATTATCGCGA AGATGGCGGG TATCGAAGTG CCGCAAACCA CCAAAATCTT GATCGGCGAA
     961  GTGCAGCACG TGGAGAAGAG CGAACTGTTC AGCCATGAGA AACTGAGCCC GGTTCTGGCT
    1021  ATGTACAAAG TGAAGGACTT TGACGAAGCC CTGGACAAAG CTCAGCGTCT GATTGAGCTG
    1081  GGCGGCAGCG GTCACACGAG CAGCCTGTAT AGCGATAGCC AAAACAATAA GGACAAGGTG
    1141  AAAGAGTTCG GTCTGGCGAT GAAAACCTCT CGCACGTTTA TCAACATGCC GAGCAGCCAA
    1201  GGTGCGAGCG GCGATCTGTA TAACTTCGCA ATCGCGCCTT CTTTTACGCT GGGTTGCGGT
    1261  ACCTGGGGCG GTAATTCGGT GAGCCAGAAT GTCGAGCCGA AGCACCTGCT GAACATCAAA
    1321  TCGGTCGCGG AGCGCCGTGA GAAGAGACCG TGGTTCAAAG TCCCACAGAA AATTTACTTC
    1381  AAATACGGCT GCCTGCGTTT CGCGCTGAAA GAACTGAAAG ACATGAACAA AAAGCGTGCG
    1441  TTCATTGTTA CCGACAAAGA CCTGTTCAAA CTGGGTTACG TGAACAAAAT CACCAAAGTT
    1501  CTGGATGAAA TTGACATCAA GTACTCCATC TTCACTGATA TCAAATCCGA CCCAACGATT
    1561  GATAGCGTGA AAAAGGGCGC TAAAGAAATG CTGAACTTTG AACCGGACAC CATCATCAGC
    1621  ATCGGTGGTG GCTCTCCTAT GGATGCTGCG AAGGTCATGC ACCTGCTGTA CGAATACCCG
    1681  GAAGCGGAAA TCGAAAACCT GGCTATCAAC TTCATGGACA TCCGCAAACG TATCTGCAAC
    1741  TTCCCGAAGC TGGGCACTAA AGCTATTTCC GTTGCCATCC CGACTACCGC GGGCACTGGT
    1801  TCCGAAGCCA CGCCGTTCGC CGTGATCACC AACGATGAAA CCGGTATGAA ATACCCGCTG
    1861  ACCTCTTACG AACTGACCCC GAACATGGCA ATTATCGACA CCGAGCTGAT GCTGAACATG
    1921  CCGCGCAAGC TGACCGCTGC TACCGGCATC GACGCTCTGG TACATGCTAT TGAGGCGTAC
    1981  GTTTCCGTGA TGGCTACCGA TTACACCGAC GAACTGGCCC TGCGTGCGAT CAAAATGATT
    2041  TTCAAGTACC TGCCTCGCGC TTACAAAAAC GGCACGAATG ACATCGAGGC GCGTGAGAAA
    2101  ATGGCCCATG CAAGCAACAT CGCGGGCATG GCCTTCGCCA ACGCGTTCCT GGGCGTGTGC
    2161  CACTCTATGG CTCACAAACT GGGTGCTATG CACCACGTGC CGCACGGTAT CGCGTGTGCT
    2221  GTCCTGATCG AAGAAGTAAT TAAGTACAAC GCTACTGATT GCCCGACTAA ACAGACCGCC
    2281  TTCCCACAGT ACAAATCTCC TAACGCTAAA CGTAAGTACG CTGAGATCGC CGAATACCTG
    2341  AACCTGAAGG GTACGAGCGA CACTGAGAAA GTTACTGCGC TGATCGAAGC TATCTCTAAA
    2401  CTGAAAATTG ACCTGTCCAT CCCGCAGAAC ATCAGCGCCG CAGGCATCAA CAAAAAGGAC
    2461  TTTTACAACA CGCTGGACAA AATGAGCGAA CTGGCTTTTG ACGACCAGTG CACCACTGCA
    2521  AACCCGCGTT ACCCGCTGAT CTCTGAGCTG AAAGACATTT ATATCAAATC TTTTTAA
```

**AdhE2.70 (76999)**

```
       1  ATGAAAGTGA CGAACCAAAA AGAACTGAAA CAAAAACTGA ATGAACTGCG TGAGGCCCAG
      61  AAAAAGTTCC TGACGTATAC CCAGGAGCAA GTTGACAAGA TCTTCAAACA GTGTGCCATT
```

121 GCGGCTGCCA AAGAGCGTAT TAACTTGGCC AAACTGGCGG TGGAAGAAAC CGGCATCGGT

181 CTGGTCGAGG ACAAGATTAT CAAGAATCAC TTTGCCGCGG AGTATATCTA CAACAAATAC

```
     241  AAAAACGAAA AGACTTGCGG CATCATTGAT CATGATGACA GCTTGGGTAT CACGAAAGTC
     301  GCAGAACCAA TCGGTATTGT CGCGGCAATT GTCCCGACCA CTAATCCGAC CAGCACCGCG
     361  ATCTTCAAGA GCTTGATCAG CCTGGCGACC CGCAACGCGA TTTTCTTCAG CCCGCACCCG
     421  CGTGCGAAGA AGAGCACCAT CGCAGCGGCG AAGCTGATTT TGGATGCCGC GGTTAAGGCC
     481  GGTGCACCGA AAAACATTAT TGGCTGGATT GACGAGCCGA GCATTGAGCT GTCTCAGGAC
     541  CTGATGAGCG AAGCGGACAT TATTCTGGCC ACCGGTGGTC CGTCTATGGT TAAGGCAGCG
     601  TACTCCAGCG GTAAGCCTGC TATCGGCGTT GGTGCGGGCA ATACGCCGGC TATCATCGAT
     661  GAAAGCGCGG ATATTGATAT GGCAGTTAGC TCCATCATTC TGGGTAAAAC CTACGACAAT
     721  GGTGTTATCT GTGCAAGCGA GCAAAGCATC CTGGTGATGA ATAGCATTTA CGAGAAGGTC
     781  AAAGAAGAGT TTGTTAAGCG TGGTTCCTAT ATTCTGAACC AGAACGAGAT TGCAAAAATC
     841  AAAGAAACGA TGTTTAAGAA TGGCGCGATT AATGCAGATA TTGTTGGTAA GTCCGCATAC
     901  ATTATCGCGA AGATGGCGGG TATCGAAGTG CCGCAAACCA CCAAAATCTT GATCGGCGAA
     961  GTGCAGAGCG TGGAGAAGAG CGAACTGTTC AGCCATGAGA AACTGAGCCC GGTTCTGGCT
    1021  ATGTACAAAG TGAAGGACTT TGACGAAGCC CTGAAGAAAG CTCAGCGTCT GATTGAGCTG
    1081  GGCGGCAGCG GTCACACGAG CAGCCTGTAT ATCGATAGCC AAAACAATAA GGACAAGGTG
    1141  AAAGAGTTCG GTCTGGCGAT GAAAACCTCT CGCACGTTTA TCAACATGCC GAGCAGCCAA
    1201  GGTGCGAGCG GCGATCTGTA TAACTTCGCA ATCGCGCCTT CTTTTACGCT GGCGTGCGGT
    1261  ACCTGGGGCG GTAATTCGGT GAGCCAGAAT GTCGAGCCGA AGCACCTGCT GAACATCAAA
    1321  TCGGTCGCGG AGCGCCGTGA GAAGAGACCG TGGTTCAAAG TCCCACAGAA AATTTACTTC
    1381  AAATACGGCT GCCTGCGTTT CGCGCTGAAA GAACTGAAAG ACATGAACAA AAAGCGTGCG
    1441  TTCATTGTTA CCGACAAAGA CCTGTTCAAA CTGGGTTACG TGAACAAAAT CACCAAAGTT
    1501  CTGGATGAAA TTGACATCAA GTACTCCATC TTCACTGATA TCAAATCCGA CCCAACGATT
    1561  GATAGCGTGA AAAAGGGCGC TAAAGAAATG CTGAACTTTG AACCGGACAC CATCATCAGC
    1621  ATCGGTGGTG GCTCTCCTAT GGATGCTGCG AAGGTCATGC ACCTGCTGTA CGAATACCCG
    1681  GAAGCGGAAA TCGAAAACCT GGCTATCAAC TTCATGGACA TCCGCAAACG TATCTGCAAC
    1741  TTCCCGAAGC TGGGCACTAA AGCTATTTCC GTTGCCATCC CGACTACCGC GGGCACTGGT
    1801  TCCGAAGCCA CGCCGTTCGC CGTGATCACC AACGATGAAA CCGGTATGAA ATACCCGCTG
    1861  ACCTCTTACG AACTGACCCC GAACATGGCA ATTATCGACA CCGAGCTGAT GCTGAACATG
    1921  CCGCGCAAGC TGACCGCTGC TACCGGCATC GACGCTCTGG TACATGCTAT TGAGGCGTAC
    1981  GTTTCCGTGA TGGCTACCGA TTACACCGAC GAACTGGCCC TGCGTGCGAT CAAAATGATT
    2041  TTCAAGTACC TGCCTCGCGC TTACAAAAAC GGCACGAATG ACATCGAGGC GCGTGAGAAA
    2101  ATGGCCCATG CAAGCAACAT CGCGGGCATG GCCTTCGCCA ACGCGTTCCT GGGCGTGTGC
    2161  CACTCTATGG CTCACAAACT GGGTGCTATG CACCACGTGC CGCACGGTAT CGCGTGTGCT
    2221  GTCCTGATCG AAGAAGTAAT TAAGTACAAC GCTACTGATT GCCCGACTAA ACAGACCGCC
    2281  TTCCCACAGT ACAAATCTCC TAACGCTAAA CGTAAGTACG CTGAGATCGC CGAATACCTG
    2341  AACCTGAAGG GTACGAGCGA CACTGAGAAA GTTACTGCGC TGATCGAAGC TATCTCTAAA
    2401  CTGAAAATTG ACCTGTCCAT CCCGCAGAAC ATCAGCGCCG CAGGCATCAA CAAAAAGGAC
    2461  TTTTACAACA CGCTGGACAA AATGAGCGAA CTGGCTTTTG ACGACCAGTG CACCACTGCA
    2521  AACCCGCGTT ACCCGCTGAT CTCTGAGCTG AAAGACATTT ATATCAAATC TTTTTAA
```

**AdhE2.71 (77000)**

```
       1  ATGAAAGTGA CGAACCAAAA AGAACTGAAA CAAAAACTGA ATGAACTGCG TGAGGCCCAG
      61  AAAAAGTTCG CTACGTATAC CCAGGAGCAA GTTGACAAGA TCTTCAAACA GTGTGCCATT
```

121 GCGGCTGCCA AAGAGCGTAT TAACTTGGCC AAACTGGCGG TGGAAGAAAC CGGCATCGGT

181 CTGGTCGAGG ACAAGATTAT CAAGAATCAC TTTGCCGCGG AGTATATCTA CAACAAATAC

```
     241  AAAAACCACA AGACTTGCGG CATCATTGAT CATGATGACA GCTTGGGTAT CACGAAAGTC
     301  GCAGAACCAA TCGGTATTGT CGCGGCAATT GTCCCGACCA CTAATCCGAC CAGCACCGCG
     361  ATCTTCAAGA GCTTGATCAG CCTGAAAACC CGCAACGCGA TTTTCTTCAG CCCGCACCCG
     421  CGTGCGAAGA AGAGCACCAT CGCAGCGGCG AAGCTGATTT TGGATGCCGC GGTTAAGGCC
     481  GGTGCACCGA AAAACATTAT TGGCTGGATT GACGAGCCGA GCATTGAGCT GTCTCAGGAC
     541  CTGATGAGCG AAGCGGACAT TATTCTGGCC ACCGGTGGTC CGTCTATGGT TAAGGCAGCG
     601  TACTCCAGCG GTAAGCCTGC TATCGGCGTT GGTGCGGGCA ATACGCCGGC TATCATCGAT
     661  GAAAGCGCGG ATATTGATAT GGCAGTTAGC TCCATCATTC TGTCCAAAAC CTACGACAAT
     721  AACGTTATCT GTGCAAGCGA GCAAAGCATC CTGGTGATGA ATAGCATTTA CGAGAAGGTC
     781  AAAGAAGAGT TTGTTAAGCG TGGTTCCTAT ATTCTGAACC AGAACGAGAT TGCAAAAATC
     841  AAAGAAACGA TGTTTAAGAA TGGCGCGATT AATGCAGATA TTGTTGGTAA GTCCGCATAC
     901  ATTATCGCGA AGATGGCGGG TATCGAAGTG CCGCAAACCA CCAAAATCTT GATCGGCGAA
     961  GTGCAGAGCG TGGAGAAGAG CGAACTGTTC AGCCATGAGA AACTGAGCCC GGTTCTGGCT
    1021  ATGTACAAAG TGAAGGACTT TGACGAAGCC CTGAAGAAAG CTCAGCGTCT GATTGAGCTG
    1081  GGCGGCAGCG GTCACACGAG CAGCCTGTAT ATCGATAGCC AAAACAATAA GGACAAGGTG
    1141  AAAGAGTTCG GTCTGGCGAT GAAAACCTCT CGCTTCTTTA TCAACATGCC GAGCAGCCAA
    1201  GGTGCGAGCG GCGATCTGTA TAACTTCGCA ATCGCGCCTT CTTTTACGCT GGGTTGCCCG
    1261  ACCTGGGGCG GTAATTCGGT GAGCCAGAAT GTCGAGCCGA AGCACCTGCT GAACATCAAA
    1321  TCGGTCGCGG AGCGCCGTGA GAAGAGACCG TGGTTCAAAG TCCCACAGAA AATTTACTTC
    1381  AAATACGGCT GCCTGCGTTT CGCGCTGAAA GAACTGAAAG ACATGAACAA AAAGCGTGCG
    1441  TTCATTGTTA CCGACAAAGA CCTGTTCAAA CTGGGTTACG TGAACAAAAT CACCAAAGTT
    1501  CTGGATGAAA TTGACATCAA GTACTCCATC TTCACTGATA TCAAATCCGA CCCAACGATT
    1561  GATAGCGTGA AAAAGGGCGC TAAAGAAATG CTGAACTTTG AACCGGACAC CATCATCAGC
    1621  ATCGGTGGTG GCTCTCCTAT GGATGCTGCG AAGGTCATGC ACCTGCTGTA CGAATACCCG
    1681  GAAGCGGAAA TCGAAAACCT GGCTATCAAC TTCATGGACA TCCGCAAACG TATCTGCAAC
    1741  TTCCCGAAGC TGGGCACTAA AGCTATTTCC GTTGCCATCC CGACTACCGC GGGCACTGGT
    1801  TCCGAAGCCA CGCCGTTCGC CGTGATCACC AACGATGAAA CCGGTATGAA ATACCCGCTG
    1861  ACCTCTTACG AACTGACCCC GAACATGGCA ATTATCGACA CCGAGCTGAT GCTGAACATG
    1921  CCGCGCAAGC TGACCGCTGC TACCGGCATC GACGCTCTGG TACATGCTAT TGAGGCGTAC
    1981  GTTTCCGTGA TGGCTACCGA TTACACCGAC GAACTGGCCC TGCGTGCGAT CAAAATGATT
    2041  TTCAAGTACC TGCCTCGCGC TTACAAAAAC GGCACGAATG ACATCGAGGC GCGTGAGAAA
    2101  ATGGCCCATG CAAGCAACAT CGCGGGCATG GCCTTCGCCA ACGCGTTCCT GGGCGTGTGC
    2161  CACTCTATGG CTCACAAACT GGGTGCTATG CACCACGTGC CGCACGGTAT CGCGTGTGCT
    2221  GTCCTGATCG AAGAAGTAAT TAAGTACAAC GCTACTGATT GCCCGACTAA ACAGACCGCC
    2281  TTCCCACAGT ACAAATCTCC TAACGCTAAA CGTAAGTACG CTGAGATCGC CGAATACCTG
    2341  AACCTGAAGG GTACGAGCGA CACTGAGAAA GTTACTGCGC TGATCGAAGC TATCTCTAAA
    2401  CTGAAAATTG ACCTGTCCAT CCCGCAGAAC ATCAGCGCCG CAGGCATCAA CAAAAAGGAC
    2461  TTTTACAACA CGCTGGACAA AATGAGCGAA CTGGCTTTTG ACGACCAGTG CACCACTGCA
    2521  AACCCGCGTT ACCCGCTGAT CTCTGAGCTG AAAGACATTT ATATCAAATC TTTTTAA
```

**AdhE2.72 (77001)**

```
       1  ATGAAAGTGA CGAACCAAAA AGAACTGAAA CAAAAACTGA ATGAACTGCG TGAGGCCCAG
      61  AAAAAGTTCG CTACGTATAC CCAGGAGCAA GTTGACAAGA TCTTCAAACA GTGTGCCATT
```

121 GCGGCTGCCA AAGAGCGTAT TAACTTGGCC AAACTGGCGG TGGAAGAAAC CGGCATCGGT

181 CTGGTCGAGG ACAAGATTAT CAAGAATCAC TTTGCCGCGG AGTATATCTA CAACAAATAC

```
     241  AAAAACGAAA AGACTTGCGG CATCATTGAT CATGATGACA GCTTGGGTAT CACGAAAGTC
     301  GCAGAACCAA TCGGTATTGT CGCGGCAATT GTCCCGACCA CTAATCCGAC CAGCACCGCG
     361  ATCTTCAAGA GCTTGATCAG CCTGAAAACC CGCAACGCGA TTTTCTTCAG CCCGCACCCG
     421  CGTGCGAAGA AGAGCACCAT CGCAGCGGCG AAGCTGATTT TGGATGCCGC GGTTAAGGCC
     481  GGTGCACCGA AAAACATTAT TGGCTGGATT GACGAGCCGA GCATTGAGCT GTCTCAGGAC
     541  CTGATGAGCG AACCGGACAT TATTCTGGCC ACCGGTGGTC CGTCTATGGT TAAGGCAGCG
     601  TACTCCAGCG GTAAGCCTGC TATCGGCGCG GGTGCGGGCA ATACGCCGGC TATCATCGAT
     661  GAAAGCGCGG ATATTGATAT GGCAGTTAGC TCCATCATTC TGTCCAAAAC CTACGACAAT
     721  GGTGTTATCT GTGCAAGCGA GCAAAGCATC CTGGTGATGA ATAGCATTTA CGAGAAGGTC
     781  AAAGAAGAGT TTGTTAAGCG TGGTTCCTAT ATTCTGAACC AGGAGGAGAT TGCAAAAATC
     841  AAAGAAACGA TGTTTAAGAA TGGCGCGATT AATGCAGATA TTGTTGGTAA GTCCGCATAC
     901  ATTATCGCGA AGATGGCGGG TATCGAAGTG CCGCAAACCA CCAAAATCTT GATCGGCGAA
     961  GTGCAGAGCG TGGAGAAGAG CGAACTGTTC AGCCATGAGA AACTGAGCCC GGTTCTGGCT
    1021  ATGTACAAAG TGAAGGACTT TGACGAAGCC CTGAAGAAAG CTCAGCGTCT GATTGAGCTG
    1081  GGCGGCAGCG GTCACACGAG CAGCCTGTAT AGCGATAGCC AAAACAATAA GGACAAGGTG
    1141  AAAGAGTTCG GTCTGGCGAT GAAAACCTCT CGCACGTTTA TCAACATGCC GAGCAGCCAA
    1201  GGTGCGAGCG GCGATCTGTA TAACTTCGCA ATCGCGCCTT CTTTTACGCT GGGTTGCGGT
    1261  ACCTGGGGCG GTAATTCGGT GAGCCAGAAT GTCGAGCCGA AGCACCTGCT GAACATCAAA
    1321  TCGGTCGCGG AGCGCCGTGA GAAGAGACCG TGGTTCAAAG TCCCACAGAA AATTTACTTC
    1381  AAATACGGCT GCCTGCGTTT CGCGCTGAAA GAACTGAAAG ACATGAACAA AAAGCGTGCG
    1441  TTCATTGTTA CCGACAAAGA CCTGTTCAAA CTGGGTTACG TGAACAAAAT CACCAAAGTT
    1501  CTGGATGAAA TTGACATCAA GTACTCCATC TTCACTGATA TCAAATCCGA CCCAACGATT
    1561  GATAGCGTGA AAAAGGGCGC TAAAGAAATG CTGAACTTTG AACCGGACAC CATCATCAGC
    1621  ATCGGTGGTG GCTCTCCTAT GGATGCTGCG AAGGTCATGC ACCTGCTGTA CGAATACCCG
    1681  GAAGCGGAAA TCGAAAACCT GGCTATCAAC TTCATGGACA TCCGCAAACG TATCTGCAAC
    1741  TTCCCGAAGC TGGGCACTAA AGCTATTTCC GTTGCCATCC CGACTACCGC GGGCACTGGT
    1801  TCCGAAGCCA CGCCGTTCGC CGTGATCACC AACGATGAAA CCGGTATGAA ATACCCGCTG
    1861  ACCTCTTACG AACTGACCCC GAACATGGCA ATTATCGACA CCGAGCTGAT GCTGAACATG
    1921  CCGCGCAAGC TGACCGCTGC TACCGGCATC GACGCTCTGG TACATGCTAT TGAGGCGTAC
    1981  GTTTCCGTGA TGGCTACCGA TTACACCGAC GAACTGGCCC TGCGTGCGAT CAAAATGATT
    2041  TTCAAGTACC TGCCTCGCGC TTACAAAAAC GGCACGAATG ACATCGAGGC GCGTGAGAAA
    2101  ATGGCCCATG CAAGCAACAT CGCGGGCATG GCCTTCGCCA ACGCGTTCCT GGGCGTGTGC
    2161  CACTCTATGG CTCACAAACT GGGTGCTATG CACCACGTGC CGCACGGTAT CGCGTGTGCT
    2221  GTCCTGATCG AAGAAGTAAT TAAGTACAAC GCTACTGATT GCCCGACTAA ACAGACCGCC
    2281  TTCCCACAGT ACAAATCTCC TAACGCTAAA CGTAAGTACG CTGAGATCGC CGAATACCTG
    2341  AACCTGAAGG GTACGAGCGA CACTGAGAAA GTTACTGCGC TGATCGAAGC TATCTCTAAA
    2401  CTGAAAATTG ACCTGTCCAT CCCGCAGAAC ATCAGCGCCG CAGGCATCAA CAAAAAGGAC
    2461  TTTTACAACA CGCTGGACAA AATGAGCGAA CTGGCTTTTG ACGACCAGTG CACCACTGCA
    2521  AACCCGCGTT ACCCGCTGAT CTCTGAGCTG AAAGACATTT ATATCAAATC TTTTTAA
```

**AdhE2.73 (77002)**

```
       1  ATGAAAGTGA CGAACCAAAA AGAACTGAAA CAAAAACTGA ATGAACTGCG TGAGGCCCAG
      61  AAAAAGTTCG CTTGCTATAC CCAGGAGCAA GTTGACAAGA TCTTCAAACA GTGTGCCATT
```

121 GCGGCTGCCA AAGAGCGTAT TAACTTGGCC CGTCTGGCGG TGGAAGAAAC CGGCATCGGT

181 CTGGTCGAGG ACAAGATTAT CAAGAATCAC TTTGCCGCGG AGTATATCTA CAACAAATAC

```
     241  AAAAACGGTA AGACTTGCGG CATCATTGAT CATGATGACA GCTTGGGTAT CACGAAAGTC
     301  GCAGAACCAA TCGGTATTGT CGCGGCAATT GTCCCGACCA CTAATCCGAC CAGCACCGCG
     361  ATCTTCAAGA GCTTGATCAG CCTGAAAACC CGCAACGCGA TTTTCTTCAG CCCGCACCCG
     421  CGTGCGAAGA AGAGCACCAT CGCAGCGGCG AAGCTGATTT TGGATGCCGC GGTTAAGGCC
     481  GGTGCACCGA AAAACATTAT TGGCTGGATT GACGAGCCGA GCATTGAGCT GTCTCAGGAC
     541  CTGATGAGCG AAGCGGACAT TATTCTGGCC ACCGGTGGTC CGTCTATGGT TAAGGCAGCG
     601  TACTCCAGCG GTAAGCCTGC TATCGGCGTT GGTGCGGGCA ATACGCCGGC TATCATCGAT
     661  GAAAGCGCGG ATATTGATAT GGCAGTTAGC TCCATCATTC TGTCCAAAAC CTACGACAAT
     721  GGTGTTATCT GTGCAAGCGA GCAAAGCATC CTGGTGATGA ATAGCATTTA CGAGAAGGTC
     781  AAAGAAGAGT TTGTTAAGCG TGGTTCCTAT ATTCTGAACC AGAACGAGAT TGCAAAAATC
     841  AAAGAAACGA TGTTTAAGAA TGGCGCGATT AATGCAGATA TTGTTGGTAA GTCCGCATAC
     901  ATTATCGCGA AGATGGCGGG TATCGAAGTG CCGCAAACCA CCAAAATCTT GATCGGCGAA
     961  GTGCAGAGCG TGGAGAAGAG CGAACTGTTC AGCCATGAGA AACTGAGCCC GGTTCTGGCT
    1021  ATGTACAAAG TGAAGGACTT TGACGAAGCC CTGAAGAAAG CTCAGCGTCT GATTGAGCTG
    1081  GGCGGCAGCG GTCACACGAG CAGCCTGTAT ATCGATAGCC AAAACAATAA GGACAAGGTG
    1141  AAAGAGTTCG GTCTGGCGAT GAAAACCTCT CGCACGTTTA TCAACATGCC GAGCAGCCAA
    1201  GGTGCGAGCG GCGATCTGTA TAACTTCGCA ATCGCGCCTT CTTTTACGCT GGGTTGCGGT
    1261  ACCTGGGAGG GTAATTCGGT GAGCCAGAAT GTCGAGCCGA AGCACCTGCT GAACATCAAA
    1321  TCGGTCGCGG AGCGCCGTGA GAAGAGACCG TGGTTCAAAG TCCCACAGAA AATTTACTTC
    1381  AAATACGGCT GCCTGCGTTT CGCGCTGAAA GAACTGAAAG ACATGAACAA AAAGCGTGCG
    1441  TTCATTGTTA CCGACAAAGA CCTGTTCAAA CTGGGTTACG TGAACAAAAT CACCAAAGTT
    1501  CTGGATGAAA TTGACATCAA GTACTCCATC TTCACTGATA TCAAATCCGA CCCAACGATT
    1561  GATAGCGTGA AAAAGGGCGC TAAAGAAATG CTGAACTTTG AACCGGACAC CATCATCAGC
    1621  ATCGGTGGTG GCTCTCCTAT GGATGCTGCG AAGGTCATGC ACCTGCTGTA CGAATACCCG
    1681  GAAGCGGAAA TCGAAAACCT GGCTATCAAC TTCATGGACA TCCGCAAACG TATCTGCAAC
    1741  TTCCCGAAGC TGGGCACTAA AGCTATTTCC GTTGCCATCC CGACTACCGC GGGCACTGGT
    1801  TCCGAAGCCA CGCCGTTCGC CGTGATCACC AACGATGAAA CCGGTATGAA ATACCCGCTG
    1861  ACCTCTTACG AACTGACCCC GAACATGGCA ATTATCGACA CCGAGCTGAT GCTGAACATG
    1921  CCGCGCAAGC TGACCGCTGC TACCGGCATC GACGCTCTGG TACATGCTAT TGAGGCGTAC
    1981  GTTTCCGTGA TGGCTACCGA TTACACCGAC GAACTGGCCC TGCGTGCGAT CAAAATGATT
    2041  TTCAAGTACC TGCCTCGCGC TTACAAAAAC GGCACGAATG ACATCGAGGC GCGTGAGAAA
    2101  ATGGCCCATG CAAGCAACAT CGCGGGCATG GCCTTCGCCA ACGCGTTCCT GGGCGTGTGC
    2161  CACTCTATGG CTCACAAACT GGGTGCTATG CACCACGTGC CGCACGGTAT CGCGTGTGCT
    2221  GTCCTGATCG AAGAAGTAAT TAAGTACAAC GCTACTGATT GCCCGACTAA ACAGACCGCC
    2281  TTCCCACAGT ACAAATCTCC TAACGCTAAA CGTAAGTACG CTGAGATCGC CGAATACCTG
    2341  AACCTGAAGG GTACGAGCGA CACTGAGAAA GTTACTGCGC TGATCGAAGC TATCTCTAAA
    2401  CTGAAAATTG ACCTGTCCAT CCCGCAGAAC ATCAGCGCCG CAGGCATCAA CAAAAAGGAC
    2461  TTTTACAACA CGCTGGACAA AATGAGCGAA CTGGCTTTTG ACGACCAGTG CACCACTGCA
    2521  AACCCGCGTT ACCCGCTGAT CTCTGAGCTG AAAGACATTT ATATCAAATC TTTTTAA
```

**AdhE2.74 (77003)**

```
       1  ATGAAAGTGA CGAACCAAAA AGAACTGAAA CAAAAAGACA ATGAACTGCG TGAGGCCCAG
      61  AAAAAGTTCG CTACGTATAC CCAGGAGCAA GTTGACAAGA TCTTCAAACA GTGTGCCATT
```

121 GCGGCTGCCA AAGAGCGTAT TAACTTGGCC CGTCTGGCGG TGGAAGAAAC CGGCATCGGT

181 CTGGTCGAGG ACAAGATTAT CAAGAATCAC TTTGCCGCGG AGTATATCTA CAACAAATAC

```
     241  AAAAACGAAA AGACTTGCGG CATCATTGAT CATGATGACA GCTTGGGTAT CACGAAAGTC
     301  GCAGAACCAA TCGGTATTGT CGCGGCAATT GTCCCGACCA CTAATCCGAC CAGCACCGCG
     361  ATCTTCAAGA GCTTGATCAG CCTGAAAACC CGCAACGCGA TTTTCTTCAG CCCGCACCCG
     421  CGTGCGAAGA AGAGCACCAT CGCAGCGGCG AAGCTGATTT TGGATGCCGC GGTTAAGGCC
     481  GGTGCACCGA AAAACATTAT TGGCTGGATT GACGAGCCGA GCATTGAGCT GTCTCAGGAC
     541  CTGATGAGCG AAGCGGACAT TATTCTGGCC ACCGGTGGTC CGTCTATGGT TAAGGCAGCG
     601  TACTCCAGCG GTAAGCCTGC TATCGGCGTT GGTGCGGGCA ATACGCCGGC TATCATCGAT
     661  GAAAGCGCGG ATATTGATAT GGCAGTTAGC TCCATCATTC TGTCCAAAAC CTACGACAAT
     721  GGTGTTATCT GTGCAAGCGA GCAAAGCATC CTGGTGATGA ATAGCATTTA CGAGAAGGTC
     781  AAAGCGGAGT TTGTTAAGCG TGGTTCCTAT ATTCTGAACC AGAACGAGAT TGCAAAAATC
     841  AAAGAAACGA TGTTTAAGAA TGGCGCGATT AATGCAGATA TTGTTGGTAA GTCCGCATAC
     901  ATTATCGCGA AGATGGCGGG TATCGAAGTG CCGCAAACCA CCAAAATCTT GATCGGCGAA
     961  GTGCAGAGCG TGGAGAAGAG CGAACTGTTC AGCGAGGAGA AACTGAGCCC GGTTCTGGCT
    1021  ATGTACAAAG TGAAGGACTT TGACGAAGCC CTGAAGAAAG CTCAGCGTCT GATTCACCTG
    1081  GGCGGCAGCG GTCACACGAG CAGCCTGTAT ATCGATAGCC AAAACAATAA GGACAAGGTG
    1141  AAAGAGTTCG GTCTGGCGAT GAAAACCTCT CGCACGTTTA TCAACATGCC GAGCAGCCAA
    1201  GGTGCGAGCG GCGATCTGTA TAACTTCGCA ATCGCGCCTT CTTTTACGCT GGGTTGCGGT
    1261  ACCTGGGGCG GTAATTCGGT GAGCCAGAAT GTCGAGCCGA AGCACCTGCT GAACATCAAA
    1321  TCGGTCGCGG AGCGCCGTGA GAAGAGACCG TGGTTCAAAG TCCCACAGAA AATTTACTTC
    1381  AAATACGGCT GCCTGCGTTT CGCGCTGAAA GAACTGAAAG ACATGAACAA AAAGCGTGCG
    1441  TTCATTGTTA CCGACAAAGA CCTGTTCAAA CTGGGTTACG TGAACAAAAT CACCAAAGTT
    1501  CTGGATGAAA TTGACATCAA GTACTCCATC TTCACTGATA TCAAATCCGA CCCAACGATT
    1561  GATAGCGTGA AAAAGGGCGC TAAAGAAATG CTGAACTTTG AACCGGACAC CATCATCAGC
    1621  ATCGGTGGTG GCTCTCCTAT GGATGCTGCG AAGGTCATGC ACCTGCTGTA CGAATACCCG
    1681  GAAGCGGAAA TCGAAAACCT GGCTATCAAC TTCATGGACA TCCGCAAACG TATCTGCAAC
    1741  TTCCCGAAGC TGGGCACTAA AGCTATTTCC GTTGCCATCC CGACTACCGC GGGCACTGGT
    1801  TCCGAAGCCA CGCCGTTCGC CGTGATCACC AACGATGAAA CCGGTATGAA ATACCCGCTG
    1861  ACCTCTTACG AACTGACCCC GAACATGGCA ATTATCGACA CCGAGCTGAT GCTGAACATG
    1921  CCGCGCAAGC TGACCGCTGC TACCGGCATC GACGCTCTGG TACATGCTAT TGAGGCGTAC
    1981  GTTTCCGTGA TGGCTACCGA TTACACCGAC GAACTGGCCC TGCGTGCGAT CAAAATGATT
    2041  TTCAAGTACC TGCCTCGCGC TTACAAAAAC GGCACGAATG ACATCGAGGC GCGTGAGAAA
    2101  ATGGCCCATG CAAGCAACAT CGCGGGCATG GCCTTCGCCA ACGCGTTCCT GGGCGTGTGC
    2161  CACTCTATGG CTCACAAACT GGGTGCTATG CACCACGTGC CGCACGGTAT CGCGTGTGCT
    2221  GTCCTGATCG AAGAAGTAAT TAAGTACAAC GCTACTGATT GCCCGACTAA ACAGACCGCC
    2281  TTCCCACAGT ACAAATCTCC TAACGCTAAA CGTAAGTACG CTGAGATCGC CGAATACCTG
    2341  AACCTGAAGG GTACGAGCGA CACTGAGAAA GTTACTGCGC TGATCGAAGC TATCTCTAAA
    2401  CTGAAAATTG ACCTGTCCAT CCCGCAGAAC ATCAGCGCCG CAGGCATCAA CAAAAAGGAC
    2461  TTTTACAACA CGCTGGACAA AATGAGCGAA CTGGCTTTTG ACGACCAGTG CACCACTGCA
    2521  AACCCGCGTT ACCCGCTGAT CTCTGAGCTG AAAGACATTT ATATCAAATC TTTTTAA
```

**AdhE2.75 (77004)**

```
       1  ATGAAAGTGA CGAACCAAAA AGAACTGAAA CAAAAACTGA ATGAACTGCG TGAGGCCCAG
      61  AAAAAGTTCG CTACGTATAC CCAGGAGCAA GTTGACAAGA TCTTCAAACA GTGTGCCATT
```

121 GCGGCTGCCA AAGAGCGTAT TAACTTGGCC AAACTGGCGG TGGAAGAAAC CGGCATCGGT

181 CTGGTCGAGG ACAAGATTAT CAAGAATCAC TTTGCCGCGG AGTATATCTA CAACAAATAC

```
     241  AAAAACGAAA AGACTTGCGG CATCATTGAT CATGATGACA GCTTGGGTAT CACGAAAGTC
     301  GCAGAACCAA TCGGTATTGT CGCGGCAATT GTCCCGACCA CTAATCCGAC CAGCACCGCG
     361  ATCTTCAAGA GCTTGATCAG CCTGAAAACC CGCAACGCGA TTTTCTTCAG CCCGCACCCG
     421  CGTGCGAAGA AGAGCACCAT CGCAGCGGCG AAGCTGATTT TGGATGCCGC GGTTAAGGCC
     481  GGTGCACCGA AAAACATTAT TGGCTGGATT GACGAGCCGA GCATTGAGCT GTCTCAGGAC
     541  ATGATGAGCG AAGCGGACAT TATTCTGGCC ACCGGTGGTC CGTCTATGGT TAAGGCAGCG
     601  TACTCCAGCG GTAAGCCTGC TATCGGCGTT GGTGCGGGCA ATACGCCGGC TATCATCGAT
     661  GAAAGCGCGG ATATTGATAT GGCAGTTAGC TCCATCATTC TGTCCAAAAC CTACGACAAT
     721  GGTGTTATCT GTGCAAGCGA GCAAAGCATC CTGGTGATGA ATAGCATTTA CGAGAAGGTC
     781  AAAGAAGAGT TTGTTAAGCG TGGTTCCTAT ATTCTGAACC AGAACGAGAT TGCAAAAATC
     841  AAAGAAACGA TGTTTAAGAA TGGCGCGATT AATGCAGATA TTGTTGGTAA GTCCGCATAC
     901  ATTATCGCGA AGATGGCGGG TATCGAAGTG CCGCAAACCA CCAAAATCTT GATCGGCGAA
     961  GTGCAGAGCG TGGAGAAGAG CGAACTGTTC AGCGAGGAGA AACTGAGCCC GGTTCTGGCT
    1021  ATGTACAAAG TGAAGGACTT TGACGAAGCC CTGGACAAAG CTCAGCGTCT GATTAACCTG
    1081  GGCGGCAGCG GTCACACGAG CAGCCTGTAT ATCGATAGCC AAAACAATAA GGACAAGGTG
    1141  AAAGAGTTCG GTCTGGCGAT GAAAACCTCT CGCACGTTTA TCAACATGCC GAGCAGCCAA
    1201  GGTGCGAGCG GCGATCTGTA TAACTTCGCA ATCGCGCCTT CTTTTACGCT GGCGTGCGGT
    1261  ACCTGGGGCG GTAATTCGGT GAGCCAGAAT GTCGAGCCGA AGCACCTGCT GAACATCAAA
    1321  TCGGTCGCGG AGCGCCGTGA GAAGAGACCG TGGTTCAAAG TCCCACAGAA AATTTACTTC
    1381  AAATACGGCT GCCTGCGTTT CGCGCTGAAA GAACTGAAAG ACATGAACAA AAAGCGTGCG
    1441  TTCATTGTTA CCGACAAAGA CCTGTTCAAA CTGGGTTACG TGAACAAAAT CACCAAAGTT
    1501  CTGGATGAAA TTGACATCAA GTACTCCATC TTCACTGATA TCAAATCCGA CCCAACGATT
    1561  GATAGCGTGA AAAAGGGCGC TAAAGAAATG CTGAACTTTG AACCGGACAC CATCATCAGC
    1621  ATCGGTGGTG GCTCTCCTAT GGATGCTGCG AAGGTCATGC ACCTGCTGTA CGAATACCCG
    1681  GAAGCGGAAA TCGAAAACCT GGCTATCAAC TTCATGGACA TCCGCAAACG TATCTGCAAC
    1741  TTCCCGAAGC TGGGCACTAA AGCTATTTCC GTTGCCATCC CGACTACCGC GGGCACTGGT
    1801  TCCGAAGCCA CGCCGTTCGC CGTGATCACC AACGATGAAA CCGGTATGAA ATACCCGCTG
    1861  ACCTCTTACG AACTGACCCC GAACATGGCA ATTATCGACA CCGAGCTGAT GCTGAACATG
    1921  CCGCGCAAGC TGACCGCTGC TACCGGCATC GACGCTCTGG TACATGCTAT TGAGGCGTAC
    1981  GTTTCCGTGA TGGCTACCGA TTACACCGAC GAACTGGCCC TGCGTGCGAT CAAAATGATT
    2041  TTCAAGTACC TGCCTCGCGC TTACAAAAAC GGCACGAATG ACATCGAGGC GCGTGAGAAA
    2101  ATGGCCCATG CAAGCAACAT CGCGGGCATG GCCTTCGCCA ACGCGTTCCT GGGCGTGTGC
    2161  CACTCTATGG CTCACAAACT GGGTGCTATG CACCACGTGC CGCACGGTAT CGCGTGTGCT
    2221  GTCCTGATCG AAGAAGTAAT TAAGTACAAC GCTACTGATT GCCCGACTAA ACAGACCGCC
    2281  TTCCCACAGT ACAAATCTCC TAACGCTAAA CGTAAGTACG CTGAGATCGC CGAATACCTG
    2341  AACCTGAAGG GTACGAGCGA CACTGAGAAA GTTACTGCGC TGATCGAAGC TATCTCTAAA
    2401  CTGAAAATTG ACCTGTCCAT CCCGCAGAAC ATCAGCGCCG CAGGCATCAA CAAAAAGGAC
    2461  TTTTACAACA CGCTGGACAA AATGAGCGAA CTGGCTTTTG ACGACCAGTG CACCACTGCA
    2521  AACCCGCGTT ACCCGCTGAT CTCTGAGCTG AAAGACATTT ATATCAAATC TTTTTAA
```

**AdhE2.76 (77005)**

```
       1  ATGAAAGTGA CGAACCAAAA AGAACTGAAA CAAAAACTGA ATGAACTGCG TGAGGCCCAG
      61  AAAAAGTTCG CTACGTATAC CCAGGAGCAA GTTGACAAGA TCTTCAAACA GTGTGCCATT
```

121 GCGGCTGCCA AAGAGCGTAT TAACTTGGCC AAACTGGCGG TGGAAGAAAC CGGCATCGGT

181 CTGGTCGAGG ACAAGATTAT CAAGAATCAC TTTGCCGCGG AGTATATCTA CAACAAATAC

```
     241  AAAAACGGTA AGACTTGCGG CATCATTGAT CATGATGACA GCTTGGGTAT CACGAAAGTC
     301  GCAGAACCAA TCGGTATTGT CGCGGCAATT GTCCCGACCA CTAATCCGAC CAGCACCGCG
     361  ATCTTCAAGA GCTTGATCAG CCTGAAAACC CGCAACGCGA TTTTCTTCAG CCCGCACCCG
     421  CGTGCGAAGA AGAGCACCAT CGCAGCGGCG ACCCTGATTT TGGATGCCGC GGTTAAGGCC
     481  GGTGCACCGA AAAACATTAT TGGCTGGATT GACGAGCCGA GCATTGAGCT GTCTCAGGAC
     541  CTGATGAGCG AAGCGGACAT TATTCTGGCC ACCGGTGGTC CGTCTATGGT TAAGGCAGCG
     601  TACTCCAGCG GTAAGCCTGC TATCGGCGTT GGTGCGGGCA ATACGCCGGC TATCATCGAT
     661  GAAAGCGCGG ATATTGATAT GGCAGTTAGC TCCATCATTC TGTCCAAAAC CTACGACAAT
     721  GGTGTTATCT GTGCAAGCGA GCAAAGCATC CTGGTGATGA ATAGCATTTA CGAGAAGGTC
     781  AAAGAAGAGT TTGTTAAGCG TGGTTCCTAT ATTCTGAACC AGAACGCGAT TGCAAAAATC
     841  AAAGAAACGA TGTTTAAGAA TGGCGCGATT AATGCAGATA TTGTTGGTAA GTCCGCATAC
     901  ATTATCGCGA AGATGGCGGG TATCGAAGTG CCGCAAACCA CCAAAATCTT GATCGGCGAA
     961  GTGCAGCACG TGGAGAAGAG CGAACTGTTC AGCCATGAGA AACTGAGCCC GGTTCTGGCT
    1021  ATGTACAAAG TGAAGGACTT TGACGAAGCC CTGAAGAAAG CTCAGCGTCT GATTCACCTG
    1081  GGCGGCAGCG GTCACACGAG CAGCCTGTAT ATCGATAGCC AAAACAATAA GGACAAGGTG
    1141  AAAGAGTTCG GTCTGGCGAT GAAAACCTCT CGCACGTTTA TCAACATGCC GAGCAGCCAA
    1201  GGTGCGAGCG GCGATCTGTA TAACTTCGCA ATCGCGCCTT CTTTTACGCT GGGTTGCGGT
    1261  ACCTGGGGCG GTAATTCGGT GAGCCAGAAT GTCGAGCCGA AGCACCTGCT GAACATCAAA
    1321  TCGGTCGCGG AGCGCCGTGA GAAGAGACCG TGGTTCAAAG TCCCACAGAA AATTTACTTC
    1381  AAATACGGCT GCCTGCGTTT CGCGCTGAAA GAACTGAAAG ACATGAACAA AAAGCGTGCG
    1441  TTCATTGTTA CCGACAAAGA CCTGTTCAAA CTGGGTTACG TGAACAAAAT CACCAAAGTT
    1501  CTGGATGAAA TTGACATCAA GTACTCCATC TTCACTGATA TCAAATCCGA CCCAACGATT
    1561  GATAGCGTGA AAAAGGGCGC TAAAGAAATG CTGAACTTTG AACCGGACAC CATCATCAGC
    1621  ATCGGTGGTG GCTCTCCTAT GGATGCTGCG AAGGTCATGC ACCTGCTGTA CGAATACCCG
    1681  GAAGCGGAAA TCGAAAACCT GGCTATCAAC TTCATGGACA TCCGCAAACG TATCTGCAAC
    1741  TTCCCGAAGC TGGGCACTAA AGCTATTTCC GTTGCCATCC CGACTACCGC GGGCACTGGT
    1801  TCCGAAGCCA CGCCGTTCGC CGTGATCACC AACGATGAAA CCGGTATGAA ATACCCGCTG
    1861  ACCTCTTACG AACTGACCCC GAACATGGCA ATTATCGACA CCGAGCTGAT GCTGAACATG
    1921  CCGCGCAAGC TGACCGCTGC TACCGGCATC GACGCTCTGG TACATGCTAT TGAGGCGTAC
    1981  GTTTCCGTGA TGGCTACCGA TTACACCGAC GAACTGGCCC TGCGTGCGAT CAAAATGATT
    2041  TTCAAGTACC TGCCTCGCGC TTACAAAAAC GGCACGAATG ACATCGAGGC GCGTGAGAAA
    2101  ATGGCCCATG CAAGCAACAT CGCGGGCATG GCCTTCGCCA ACGCGTTCCT GGGCGTGTGC
    2161  CACTCTATGG CTCACAAACT GGGTGCTATG CACCACGTGC CGCACGGTAT CGCGTGTGCT
    2221  GTCCTGATCG AAGAAGTAAT TAAGTACAAC GCTACTGATT GCCCGACTAA ACAGACCGCC
    2281  TTCCCACAGT ACAAATCTCC TAACGCTAAA CGTAAGTACG CTGAGATCGC CGAATACCTG
    2341  AACCTGAAGG GTACGAGCGA CACTGAGAAA GTTACTGCGC TGATCGAAGC TATCTCTAAA
    2401  CTGAAAATTG ACCTGTCCAT CCCGCAGAAC ATCAGCGCCG CAGGCATCAA CAAAAAGGAC
    2461  TTTTACAACA CGCTGGACAA AATGAGCGAA CTGGCTTTTG ACGACCAGTG CACCACTGCA
    2521  AACCCGCGTT ACCCGCTGAT CTCTGAGCTG AAAGACATTT ATATCAAATC TTTTTAA
```

**AdhE2.77 (77006)**

```
       1  ATGAAAGTGA CGAACCAAAA AGAACTGAAA CAAAAACTGA ATGAACTGCG TGAGGCCCAG
      61  AAAAAGTTCG CTACGATGAC CCAGGAGCAA GTTGAGAAGA TCTTCAAACA GTGTGCCATT
```

121 GCGGCTGCCA AACTGCGTAT TAACTTGGCC AAACTGGCGG TGGAAGAAAC CGGCATCGGT

181 CTGGTCGAGG ACAAGATTAT CAAGAATCAC TTTGCCGCGG AGTATATCTA CAACAAATAC

```
     241  AAAAACGAAA AGACTTGCGG CATCATTGAT CATGATGACA GCTTGGGTAT CACGAAAGTC
     301  GCAGAACCAA TCGGTATTGT CGCGGCAATT GTCCCGACCA CTAATCCGAC CAGCACCGCG
     361  ATCTTCAAGA GCTTGATCAG CCTGAAAACC CGCAACGCGA TTTTCTTCAG CCCGCACCCG
     421  CGTGCGAAGA AGAGCACCAT CGCAGCGGCG AAGCTGATTT TGGATGCCGC GGTTAAGGCC
     481  GGTGCACCGA AAAACATTAT TGGCTGGATT GACGAGCCGA GCATTGAGCT GTCTCAGGAC
     541  CTGATGAGCG AAGCGGACAT TATTCTGGCC ACCGGTGGTC CGTCTATGGT TAAGGCAGCG
     601  TACTCCAGCG GTAAGCCTGC TATCGGCGTT GGTGCGGGCA ATACGCCGGC TATCATCGAT
     661  GAAAGCGCGG ATATTGATAT GGCAGTTAGC TCCATCATTC TGTCCAAAAC CTACGACAAT
     721  GGTGTTATCT GTGCAAGCGA GCAAAGCATC CTGGTGATGA ATAGCATTTA CGAGAAGGTC
     781  AAAAAAGAGT TTGTTAAGCG TGGTTCCTAT ATTCTGAACC AGAACGAGAT TGCAAAAATC
     841  AAAGAAACGA TGTTTAAGAA TGGCGCGATT AATGCAGATA TTGTTGGTAA GTCCGCATAC
     901  ATTATCGCGA AGATGGCGGG TATCGAAGTG CCGCAATACA CCAAAATCTT GATCGGCGAA
     961  GTGCAGAGCG TGGAGAAGAG CGAACTGTTC AGCCATGAGA AACTGAGCCC GGTTCTGGCT
    1021  ATGTACAAAG TGAAGGACTT TGACGAAGCC CTGAAGAAAG CTCAGCGTCT GATTGAGCTG
    1081  GGCGGCAGCG GTCACACGAG CAGCCTGTAT ATCGATAGCC AAAACAATAA GGACAAGGTG
    1141  AAAGAGTTCG GTCTGGCGAT GAAAACCTCT CGCACGTTTA TCAACATGCC GAGCAGCCAA
    1201  GGTGCGAGCG GCGATCTGTA TAACTTCGCA ATCGCGCCTT CTTTTACGCT GGGTTGCGGT
    1261  ACCTGGGGCG GTAATTCGGT GAGCCAGAAT GTCGAGCCGA AGCACCTGCT GAACATCAAA
    1321  TCGGTCGCGG AGCGCCGTGA GAAGAGACCG TGGTTCAAAG TCCCACAGAA AATTTACTTC
    1381  AAATACGGCT GCCTGCGTTT CGCGCTGAAA GAACTGAAAG ACATGAACAA AAAGCGTGCG
    1441  TTCATTGTTA CCGACAAAGA CCTGTTCAAA CTGGGTTACG TGAACAAAAT CACCAAAGTT
    1501  CTGGATGAAA TTGACATCAA GTACTCCATC TTCACTGATA TCAAATCCGA CCCAACGATT
    1561  GATAGCGTGA AAAAGGGCGC TAAAGAAATG CTGAACTTTG AACCGGACAC CATCATCAGC
    1621  ATCGGTGGTG GCTCTCCTAT GGATGCTGCG AAGGTCATGC ACCTGCTGTA CGAATACCCG
    1681  GAAGCGGAAA TCGAAAACCT GGCTATCAAC TTCATGGACA TCCGCAAACG TATCTGCAAC
    1741  TTCCCGAAGC TGGGCACTAA AGCTATTTCC GTTGCCATCC CGACTACCGC GGGCACTGGT
    1801  TCCGAAGCCA CGCCGTTCGC CGTGATCACC AACGATGAAA CCGGTATGAA ATACCCGCTG
    1861  ACCTCTTACG AACTGACCCC GAACATGGCA ATTATCGACA CCGAGCTGAT GCTGAACATG
    1921  CCGCGCAAGC TGACCGCTGC TACCGGCATC GACGCTCTGG TACATGCTAT TGAGGCGTAC
    1981  GTTTCCGTGA TGGCTACCGA TTACACCGAC GAACTGGCCC TGCGTGCGAT CAAAATGATT
    2041  TTCAAGTACC TGCCTCGCGC TTACAAAAAC GGCACGAATG ACATCGAGGC GCGTGAGAAA
    2101  ATGGCCCATG CAAGCAACAT CGCGGGCATG GCCTTCGCCA ACGCGTTCCT GGGCGTGTGC
    2161  CACTCTATGG CTCACAAACT GGGTGCTATG CACCACGTGC CGCACGGTAT CGCGTGTGCT
    2221  GTCCTGATCG AAGAAGTAAT TAAGTACAAC GCTACTGATT GCCCGACTAA ACAGACCGCC
    2281  TTCCCACAGT ACAAATCTCC TAACGCTAAA CGTAAGTACG CTGAGATCGC CGAATACCTG
    2341  AACCTGAAGG GTACGAGCGA CACTGAGAAA GTTACTGCGC TGATCGAAGC TATCTCTAAA
    2401  CTGAAAATTG ACCTGTCCAT CCCGCAGAAC ATCAGCGCCG CAGGCATCAA CAAAAAGGAC
    2461  TTTTACAACA CGCTGGACAA AATGAGCGAA CTGGCTTTTG ACGACCAGTG CACCACTGCA
    2521  AACCCGCGTT ACCCGCTGAT CTCTGAGCTG AAAGACATTT ATATCAAATC TTTTTAA
```

**AdhE2.78 (77007)**

```
       1  ATGAAAGTGA CGAACCAAAA AGAACTGAAA CAAAAACTGA ATGAACTGCG TGAGGCCCAG
      61  AAAAAGTTCG CTTGCATGAC CCAGGAGCAA GTTGACAAGA TCTTCAAACA GTGTGCCATT
```

121 GCGGCTGCCA AAGAGCGTAT TAACTTGGCC AAACTGGCGG TGGAAGAAAC CGGCATCGGT

181 CTGGTCGAGG ACAAGATTAT CAAGAATCAC TTTGCCGCGG AGTATATCTA CAACAAATAC

```
     241  AAAAACGAAA AGACTTGCGG CATCATTGAT CATGATGACA GCTTGGGTAT CACGAAAGTC
     301  GCAGAACCAA TCGGTATTGT CGCGGCAATT GTCCCGACCA CTAATCCGAC CAGCACCGCG
     361  ATCTTCAAGA GCTTGATCAG CCTGAAAACC CGCAACGCGA TTTTCTTCAG CCCGCACCCG
     421  CGTGCGAAGA AGAGCACCAT CGCAGCGGCG AAGCTGATTT TGGATGCCGC GGTTAAGGCC
     481  GGTGCACCGA AAAACATTAT TGGCTGGATT GACGAGCCGA GCATTGAGCT GTCTCAGGAC
     541  CTGATGAGCG AAGCGGACAT TATTCTGGCC ACCGGTGGTC CGTCTATGGT TAAGGCAGCG
     601  TACTCCAGCG GTAAGAAAGC TATCGGCGTT GGTGCGGGCA ATACGCCGGC TATCATCGAT
     661  GAAAGCGCGG ATATTGATAT GGCAGTTAGC TCCATCATTC TGTCCAAAAC CTACGACAAT
     721  GGTGTTATCT GTGCAAGCGA GCAAAGCATC CTGGTGATGA ATAGCATTTA CGAGAAGGTC
     781  AAAGAAGAGT TTGTTAAGCG TGGTTCCTAT ATTCTGAACC AGAACGAGAT TGCAAAAATC
     841  AAAGAAACGA TGTTTAAGAA TGGCGCGATT AATGCAGATA TTGTTGGTAA GTCCGCATAC
     901  ATTATCGCGA AGATGGCGGG TATCGAAGTG CCGCAAACCA CCAAAATCTT GATCGGCGAA
     961  GTGCAGAGCG TGGAGAAGAG CGAACTGTTC AGCCATGAGA AACTGAGCCC GGTTCTGGCT
    1021  ATGTACAAAG TGAAGGACTT TGACGAAGCC CTGAAGCTGG CTCAGCGTCT GATTGAGCTG
    1081  GGCGGCAGCG GTCACACGAG CAGCCTGTAT ATCGATAGCC AAAACAATAA GGACAAGGTG
    1141  AAAGAGTTCG GTCTGGCGAT GAAAACCTCT ATCACGTTTA TCAACATGCC GAGCAGCCAA
    1201  GGTGCGAGCG GCGATCTGTA TAACTTCGCA ATCGCGCCTT CTTTTACGCT GGGTTGCGGT
    1261  ACCTGGGGCG GTAATTCGGT GAGCCAGAAT GTCGAGCCGA AGCACCTGCT GAACATCAAA
    1321  TCGGTCGCGG AGCGCCGTGA GAAGAGACCG TGGTTCAAAG TCCCACAGAA AATTTACTTC
    1381  AAATACGGCT GCCTGCGTTT CGCGCTGAAA GAACTGAAAG ACATGAACAA AAAGCGTGCG
    1441  TTCATTGTTA CCGACAAAGA CCTGTTCAAA CTGGGTTACG TGAACAAAAT CACCAAAGTT
    1501  CTGGATGAAA TTGACATCAA GTACTCCATC TTCACTGATA TCAAATCCGA CCCAACGATT
    1561  GATAGCGTGA AAAAGGGCGC TAAAGAAATG CTGAACTTTG AACCGGACAC CATCATCAGC
    1621  ATCGGTGGTG GCTCTCCTAT GGATGCTGCG AAGGTCATGC ACCTGCTGTA CGAATACCCG
    1681  GAAGCGGAAA TCGAAAACCT GGCTATCAAC TTCATGGACA TCCGCAAACG TATCTGCAAC
    1741  TTCCCGAAGC TGGGCACTAA AGCTATTTCC GTTGCCATCC CGACTACCGC GGGCACTGGT
    1801  TCCGAAGCCA CGCCGTTCGC CGTGATCACC AACGATGAAA CCGGTATGAA ATACCCGCTG
    1861  ACCTCTTACG AACTGACCCC GAACATGGCA ATTATCGACA CCGAGCTGAT GCTGAACATG
    1921  CCGCGCAAGC TGACCGCTGC TACCGGCATC GACGCTCTGG TACATGCTAT TGAGGCGTAC
    1981  GTTTCCGTGA TGGCTACCGA TTACACCGAC GAACTGGCCC TGCGTGCGAT CAAAATGATT
    2041  TTCAAGTACC TGCCTCGCGC TTACAAAAAC GGCACGAATG ACATCGAGGC GCGTGAGAAA
    2101  ATGGCCCATG CAAGCAACAT CGCGGGCATG GCCTTCGCCA ACGCGTTCCT GGGCGTGTGC
    2161  CACTCTATGG CTCACAAACT GGGTGCTATG CACCACGTGC CGCACGGTAT CGCGTGTGCT
    2221  GTCCTGATCG AAGAAGTAAT TAAGTACAAC GCTACTGATT GCCCGACTAA ACAGACCGCC
    2281  TTCCCACAGT ACAAATCTCC TAACGCTAAA CGTAAGTACG CTGAGATCGC CGAATACCTG
    2341  AACCTGAAGG GTACGAGCGA CACTGAGAAA GTTACTGCGC TGATCGAAGC TATCTCTAAA
    2401  CTGAAAATTG ACCTGTCCAT CCCGCAGAAC ATCAGCGCCG CAGGCATCAA CAAAAAGGAC
    2461  TTTTACAACA CGCTGGACAA AATGAGCGAA CTGGCTTTTG ACGACCAGTG CACCACTGCA
    2521  AACCCGCGTT ACCCGCTGAT CTCTGAGCTG AAAGACATTT ATATCAAATC TTTTTAA
```

**AdhE2.79 (77008)**

```
       1  ATGAAAGTGA CGAACCAAAA AGAACTGAAA CAAAAACTGA ATGAACTGCG TGAGGCCCAG
      61  AAAAAGTTCG CTACGTATAC CCAGGAGCAA GTTGACAAGA TCTTCAAACA GTGTGCCATT
```

121 GCGGCTGCCA AAGAGCGTAT TAACTTGGCC AAACTGGCGG TGGAAGAAAC CGGCATCGGT

181 CTGGTCGAGG ACAAGATTAT CAAGAATCAC TTTGCCGCGG AGTATATCTA CAACAAATAC

```
     241  AAAAACGAAA AGACTTGCGG CATCATTGAT CATGATGACA GCTTGGGTAT CACGAAAGTC
     301  GCAGAACCAA TCGGTATTGT CGCGGCAATT GTCCCGACCA CTAATCCGAC CAGCACCGCG
     361  ATCTTCAAGA GCTTGATCAG CCTGAAAACC CGCAACGCGA TTTTCTTCAG CCCGCACCCG
     421  CGTGCGAAGA AGAGCACCAT CGCAGCGGCG AAGCTGATTT TGGATGCCGC GGTTAAGGCC
     481  GGTGCACCGA AAAACATTAT TGGCTGGATT GACGAGCCGA GCATTGAGCT GTCTCAGGAC
     541  CTGATGAGCG AAGCGGACAT TATTCTGGCC ACCGGTGGTC CGTCTATGGT TAAGGCAGCG
     601  TACTCCAGCG GTAAGCCTGC TATCGGCGTT GGTGCGGGCA ATACGCCGGC TATCATCGAT
     661  GAAAGCGCGG ATATTGATAT GGCAGTTAGC TCCATCATTC TGTCCAAAAC CTACGACAAT
     721  GGTGTTATCT GTGCAAGCGA GCAAAGCATC CTGGTGATGA ATAGCATTTA CGAGAAGGTC
     781  AAAGAAGAGT TTGTTAAGCG TGGTTCCTAT ATTCTGAACC AGCAGGCGAT TGCAAAAATC
     841  AAAGAAACGA TGTTTAAGAA TGGCGCGATT AATGCAGATA TTGTTGGTAA GTCCGCATAC
     901  ATTATCGCGA AGATGGCGGG TATCGAAGTG CCGCAATACA CCAAAATCTT GATCGGCGAA
     961  GTGCAGAGCG TGGAGAAGAG CGAACTGTTC AGCCATGAGA AACTGAGCCC GGTTCTGGCT
    1021  ATGTACAAAG TGAAGGACTT TGACGAAGCC CTGAAGAAAG CTGAGCGTCT GATTGAGCTG
    1081  GGCGGCAGCG GTCACACGAG CAGCCTGTAT AGCGATAGCC AAAACAATAA GGACAAGGTG
    1141  AAAGAGTTCG GTCTGGCGAT GAAAACCTCT CGCACGTTTA TCAACATGCC GAGCAGCCAA
    1201  GGTGCGAGCG GCGATCTGTA TAACTTCGCA ATCGCGCCTT CTTTTACGCT GGGTTGCGGT
    1261  ACCTGGGGCG GTAATTCGGT GAGCCAGAAT GTCGAGCCGA AGCACCTGCT GAACATCAAA
    1321  TCGGTCGCGG AGCGCCGTGA GAAGAGACCG TGGTTCAAAG TCCCACAGAA AATTTACTTC
    1381  AAATACGGCT GCCTGCGTTT CGCGCTGAAA GAACTGAAAG ACATGAACAA AAAGCGTGCG
    1441  TTCATTGTTA CCGACAAAGA CCTGTTCAAA CTGGGTTACG TGAACAAAAT CACCAAAGTT
    1501  CTGGATGAAA TTGACATCAA GTACTCCATC TTCACTGATA TCAAATCCGA CCCAACGATT
    1561  GATAGCGTGA AAAAGGGCGC TAAAGAAATG CTGAACTTTG AACCGGACAC CATCATCAGC
    1621  ATCGGTGGTG GCTCTCCTAT GGATGCTGCG AAGGTCATGC ACCTGCTGTA CGAATACCCG
    1681  GAAGCGGAAA TCGAAAACCT GGCTATCAAC TTCATGGACA TCCGCAAACG TATCTGCAAC
    1741  TTCCCGAAGC TGGGCACTAA AGCTATTTCC GTTGCCATCC CGACTACCGC GGGCACTGGT
    1801  TCCGAAGCCA CGCCGTTCGC CGTGATCACC AACGATGAAA CCGGTATGAA ATACCCGCTG
    1861  ACCTCTTACG AACTGACCCC GAACATGGCA ATTATCGACA CCGAGCTGAT GCTGAACATG
    1921  CCGCGCAAGC TGACCGCTGC TACCGGCATC GACGCTCTGG TACATGCTAT TGAGGCGTAC
    1981  GTTTCCGTGA TGGCTACCGA TTACACCGAC GAACTGGCCC TGCGTGCGAT CAAAATGATT
    2041  TTCAAGTACC TGCCTCGCGC TTACAAAAAC GGCACGAATG ACATCGAGGC GCGTGAGAAA
    2101  ATGGCCCATG CAAGCAACAT CGCGGGCATG GCCTTCGCCA ACGCGTTCCT GGGCGTGTGC
    2161  CACTCTATGG CTCACAAACT GGGTGCTATG CACCACGTGC CGCACGGTAT CGCGTGTGCT
    2221  GTCCTGATCG AAGAAGTAAT TAAGTACAAC GCTACTGATT GCCCGACTAA ACAGACCGCC
    2281  TTCCCACAGT ACAAATCTCC TAACGCTAAA CGTAAGTACG CTGAGATCGC CGAATACCTG
    2341  AACCTGAAGG GTACGAGCGA CACTGAGAAA GTTACTGCGC TGATCGAAGC TATCTCTAAA
    2401  CTGAAAATTG ACCTGTCCAT CCCGCAGAAC ATCAGCGCCG CAGGCATCAA CAAAAAGGAC
    2461  TTTTACAACA CGCTGGACAA AATGAGCGAA CTGGCTTTTG ACGACCAGTG CACCACTGCA
    2521  AACCCGCGTT ACCCGCTGAT CTCTGAGCTG AAAGACATTT ATATCAAATC TTTTTAA
```

**AdhE2.80 (77009	)**

```
       1  ATGAAAGTGA CGAACCAAAA AGAACTGAAA CAAAAACTGA ATGAACTGCG TGAGGCCCAG
      61  AAAAAGTTCG CTACGTATAC CCAGGCGCAA GTTGACAAGA TCTTCAAACA GTGTGCCATT
```

121 GCGGCTGCCA AAGAGCGTAT TAACTTGGCC AAACTGGCGG TGGAAGAAAC CGGCATCGGT

181 CTGGTCGAGG ACAAGATTAT CAAGAATCAC TTTGCCGCGG AGTATATCTA CAACAAATAC

```
     241  AAAAACGAAA AGACTTGCGG CATCATTGAT CATGATGACA GCTTGGGTAT CACGAAAGTC
     301  GCAGAACCAA TCGGTATTGT CGCGGCAATT GTCCCGACCA CTAATCCGAC CAGCACCGCG
     361  ATCTTCAAGA GCTTGATCAG CCTGAAAACC CGCAACGCGA TTTTCTTCAG CCCGCACCCG
     421  CGTGCGAAGA AGAGCACCAT CGCAGCGGCG AAGCTGATTT TGGATGCCGC GGTTAAGGCC
     481  GGTGCACCGA AAAACATTAT TGGCTGGATT GACGAGCCGA GCATTGAGCT GTCTCAGGAC
     541  CTGATGAGCG AAGCGGACAT TATTCTGGCC ACCGGTGGTC CGTCTATGGT TAAGGCAGCG
     601  TACTCCAGCG GTAAGCCTGC TATCGGCGCG GGTGCGGGCA ATACGCCGGC TATCATCGAT
     661  GAAAGCGCGG ATATTGATAT GGCAGTTAGC TCCATCATTC TGTCCAAAAC CTACGACAAT
     721  AACGTTATCT GTGCAAGCGA GCAAAGCATC CTGGTGATGA ATAGCATTTA CGAGAAGGTC
     781  AAAAAAGAGT TTGTTAAGCG TGGTTCCTAT ATTCTGAACC AGAACGAGAT TGCAAAAATC
     841  AAAGAAACGA TGTTTAAGAA TGGCGCGATT AATGCAGATA TTGTTGGTAA GTCCGCATAC
     901  ATTATCGCGA AGATGGCGGG TATCGAAGTG CCGCAAACCA CCAAAATCTT GATCGGCGAA
     961  GTGCAGCACG TGGAGAAGAG CGAACTGTTC AGCCATGAGA AACTGAGCCC GGTTCTGGCT
    1021  ATGTACAAAG TGAAGGACTT TGACGAAGCC CTGAAGAAAG CTCAGCGTCT GATTGAGCTG
    1081  GGCGGCAGCG GTCACACGAG CAGCCTGTAT ATCGATAGCC AAAACAATAA GGACAAGGTG
    1141  AAAGAGTTCG GTCTGGCGAT GAAAACCTCT CGCACGTTTA TCAACATGCC GAGCAGCCAA
    1201  GGTGCGAGCG GCGATCTGTA TAACTTCGCA ATCGCGCCTT CTTTTACGCT GGGTTGCGGT
    1261  ACCTGGGGCG GTAATTCGGT GAGCCAGAAT GTCGAGCCGA AGCACCTGCT GAACATCAAA
    1321  TCGGTCGCGG AGCGCCGTGA GAAGAGACCG TGGTTCAAAG TCCCACAGAA AATTTACTTC
    1381  AAATACGGCT GCCTGCGTTT CGCGCTGAAA GAACTGAAAG ACATGAACAA AAAGCGTGCG
    1441  TTCATTGTTA CCGACAAAGA CCTGTTCAAA CTGGGTTACG TGAACAAAAT CACCAAAGTT
    1501  CTGGATGAAA TTGACATCAA GTACTCCATC TTCACTGATA TCAAATCCGA CCCAACGATT
    1561  GATAGCGTGA AAAAGGGCGC TAAAGAAATG CTGAACTTTG AACCGGACAC CATCATCAGC
    1621  ATCGGTGGTG GCTCTCCTAT GGATGCTGCG AAGGTCATGC ACCTGCTGTA CGAATACCCG
    1681  GAAGCGGAAA TCGAAAACCT GGCTATCAAC TTCATGGACA TCCGCAAACG TATCTGCAAC
    1741  TTCCCGAAGC TGGGCACTAA AGCTATTTCC GTTGCCATCC CGACTACCGC GGGCACTGGT
    1801  TCCGAAGCCA CGCCGTTCGC CGTGATCACC AACGATGAAA CCGGTATGAA ATACCCGCTG
    1861  ACCTCTTACG AACTGACCCC GAACATGGCA ATTATCGACA CCGAGCTGAT GCTGAACATG
    1921  CCGCGCAAGC TGACCGCTGC TACCGGCATC GACGCTCTGG TACATGCTAT TGAGGCGTAC
    1981  GTTTCCGTGA TGGCTACCGA TTACACCGAC GAACTGGCCC TGCGTGCGAT CAAAATGATT
    2041  TTCAAGTACC TGCCTCGCGC TTACAAAAAC GGCACGAATG ACATCGAGGC GCGTGAGAAA
    2101  ATGGCCCATG CAAGCAACAT CGCGGGCATG GCCTTCGCCA ACGCGTTCCT GGGCGTGTGC
    2161  CACTCTATGG CTCACAAACT GGGTGCTATG CACCACGTGC CGCACGGTAT CGCGTGTGCT
    2221  GTCCTGATCG AAGAAGTAAT TAAGTACAAC GCTACTGATT GCCCGACTAA ACAGACCGCC
    2281  TTCCCACAGT ACAAATCTCC TAACGCTAAA CGTAAGTACG CTGAGATCGC CGAATACCTG
    2341  AACCTGAAGG GTACGAGCGA CACTGAGAAA GTTACTGCGC TGATCGAAGC TATCTCTAAA
    2401  CTGAAAATTG ACCTGTCCAT CCCGCAGAAC ATCAGCGCCG CAGGCATCAA CAAAAAGGAC
    2461  TTTTACAACA CGCTGGACAA AATGAGCGAA CTGGCTTTTG ACGACCAGTG CACCACTGCA
    2521  AACCCGCGTT ACCCGCTGAT CTCTGAGCTG AAAGACATTT ATATCAAATC TTTTTAA
```

**AdhE2.81 (77010)**

```
       1  ATGAAAGTGA CGAACCAAAA AGAACTGAAA CAAAAACTGA ATGAACTGCG TGAGGCCCAG
      61  AAAAAGTTCG CTAGCTATAC CCAGGCGCAA GTTGACAAGA TCTTCAAACA GTGTGCCATT
```

121 GCGGCTGCCA AAGAGCGTAT TAACTTGGCC AAACTGGCGG TGGAAGAAAC CGGCATCGGT

181 CTGGTCGAGG ACAAGATTAT CAAGAATCAC TTTGCCGCGG AGTATATCTA CAACAAATAC

```
     241  AAAAACCACA AGACTTGCGG CATCATTGAT CATGATGACA GCTTGGGTAT CACGAAAGTC
     301  GCAGAACCAA TCGGTATTGT CGCGGCAATT GTCCCGACCA CTAATCCGAC CAGCACCGCG
     361  ATCTTCAAGA GCTTGATCAG CCTGAAAACC CGCAACGCGA TTTTCTTCAG CCCGCACCCG
     421  CGTGCGAAGA AGAGCACCAT CGCAGCGGCG AAGCTGATTT TGGATGCCGC GGTTAAGGCC
     481  GGTGCACCGA AAAACATTAT TGGCTGGATT GACGAGCCGA GCATTGAGCT GTCTCAGGAC
     541  CTGATGAGCG AAGCGGACAT TATTCTGGCC ACCGGTGGTC CGTCTATGGT TAAGGCAGCG
     601  TACTCCAGCG GTAAGCCTGC TATCGGCGTT GGTGCGGGCA ATACGCCGGC TATCATCGAT
     661  GAAAGCGCGG ATATTGATAT GGCAGTTAGC TCCATCATTC TGTCCAAAAC CTACGACAAT
     721  GGTGTTATCT GTGCAAGCGA GCAAAGCATC CTGGTGATGA ATAGCATTTA CGAGAAGGTC
     781  AAAGAAGAGT TTGTTAAGCG TGGTTCCTAT ATTCTGAACC AGAACGAGAT TGCAAAAATC
     841  AAAGAAACGA TGTTTAAGAA TGGCGCGATT AATGCAGATA TTGTTGGTAA GTCCGCATAC
     901  ATTATCGCGA AGATGGCGGG TATCGAAGTG CCGCAAACCA CCAAAATCTT GATCGGCGAA
     961  GTGCAGAGCG TGGAGAAGAG CGAACTGTTC AGCGTTGAGA AACTGAGCCC GGTTCTGGCT
    1021  ATGTACAAAG TGAAGGACTT TGACGAAGCC CTGAAGAAAG CTCAGCGTCT GATTGAGCTG
    1081  GGCGGCAGCG GTCACACGAG CAGCCTGTAT ATCGATAGCC AAAACAATAA GGACAAGGTG
    1141  AAAGAGTTCG GTCTGCGTAT GAAAACCTCT CGCACGTTTA TCAACATGCC GAGCAGCCAA
    1201  GGTGCGAGCG GCGATCTGTA TAACTTCGCA ATCGCGCCTT CTTTTACGCT GGGTTGCGGT
    1261  ACCTGGGGCG GTAATTCGGT GAGCCAGAAT GTCGAGCCGA AGCACCTGCT GAACATCAAA
    1321  TCGGTCGCGG AGCGCCGTGA GAAGAGACCG TGGTTCAAAG TCCCACAGAA AATTTACTTC
    1381  AAATACGGCT GCCTGCGTTT CGCGCTGAAA GAACTGAAAG ACATGAACAA AAAGCGTGCG
    1441  TTCATTGTTA CCGACAAAGA CCTGTTCAAA CTGGGTTACG TGAACAAAAT CACCAAAGTT
    1501  CTGGATGAAA TTGACATCAA GTACTCCATC TTCACTGATA TCAAATCCGA CCCAACGATT
    1561  GATAGCGTGA AAAAGGGCGC TAAAGAAATG CTGAACTTTG AACCGGACAC CATCATCAGC
    1621  ATCGGTGGTG GCTCTCCTAT GGATGCTGCG AAGGTCATGC ACCTGCTGTA CGAATACCCG
    1681  GAAGCGGAAA TCGAAAACCT GGCTATCAAC TTCATGGACA TCCGCAAACG TATCTGCAAC
    1741  TTCCCGAAGC TGGGCACTAA AGCTATTTCC GTTGCCATCC CGACTACCGC GGGCACTGGT
    1801  TCCGAAGCCA CGCCGTTCGC CGTGATCACC AACGATGAAA CCGGTATGAA ATACCCGCTG
    1861  ACCTCTTACG AACTGACCCC GAACATGGCA ATTATCGACA CCGAGCTGAT GCTGAACATG
    1921  CCGCGCAAGC TGACCGCTGC TACCGGCATC GACGCTCTGG TACATGCTAT TGAGGCGTAC
    1981  GTTTCCGTGA TGGCTACCGA TTACACCGAC GAACTGGCCC TGCGTGCGAT CAAAATGATT
    2041  TTCAAGTACC TGCCTCGCGC TTACAAAAAC GGCACGAATG ACATCGAGGC GCGTGAGAAA
    2101  ATGGCCCATG CAAGCAACAT CGCGGGCATG GCCTTCGCCA ACGCGTTCCT GGGCGTGTGC
    2161  CACTCTATGG CTCACAAACT GGGTGCTATG CACCACGTGC CGCACGGTAT CGCGTGTGCT
    2221  GTCCTGATCG AAGAAGTAAT TAAGTACAAC GCTACTGATT GCCCGACTAA ACAGACCGCC
    2281  TTCCCACAGT ACAAATCTCC TAACGCTAAA CGTAAGTACG CTGAGATCGC CGAATACCTG
    2341  AACCTGAAGG GTACGAGCGA CACTGAGAAA GTTACTGCGC TGATCGAAGC TATCTCTAAA
    2401  CTGAAAATTG ACCTGTCCAT CCCGCAGAAC ATCAGCGCCG CAGGCATCAA CAAAAAGGAC
    2461  TTTTACAACA CGCTGGACAA AATGAGCGAA CTGGCTTTTG ACGACCAGTG CACCACTGCA
    2521  AACCCGCGTT ACCCGCTGAT CTCTGAGCTG AAAGACATTT ATATCAAATC TTTTTAA
```

**AdhE2.82 (77011)**

```
       1  ATGAAAGTGA CGAACCAAAA AGAACTGAAA CAAAAACTGA ATGAACTGCG TGAGGCCCAG
      61  AAAAAGTTCG CTACGTATAC CCAGGACCAA GTTGACAAGA TCTTCAAACA GTGTGCCATT
```

121 GCGGCTGCCA AAGAGCGTAT TAACTTGGCC AAACTGGCGG TGGAAGAAAC CGGCATCGGT

181 CTGGTCGAGG ACAAGATTAT CAAGAATCAC TTTGCCGCGG AGTATATCTA CAACAAATAC

```
     241  AAAAACGAAA AGACTTGCGG CATCATTGAT CATGATGACA GCTTGGGTAT CACGAAAGTC
     301  GCAGAACCAA TCGGTATTGT CGCGGCAATT GTCCCGACCA CTAATCCGAC CAGCACCGCG
     361  ATCTTCAAGA GCTTGATCAG CCTGAAAACC CGCAACGCGA TTTTCTTCAG CCCGCACCCG
     421  CGTGCGAAGA AGAGCACCAT CGCAGCGGCG AAGCTGATTT TGGATGCCGC GGTTAAGGCC
     481  GGTGCACCGA AAAACATTAT TGGCTGGATT GACGAGCCGA GCATTGAGCT GTCTCAGGAC
     541  CTGATGAGCG AAGCGGACAT TATTCTGGCC ACCGGTGGTC CGTCTATGGT TAAGGCAGCG
     601  TACTCCAGCG GTAAGCCTGC TATCGGCGTT GGTGCGGGCA ATACGCCGGC TATCATCGAT
     661  GAAAGCGCGG ATATTGATAT GGCAGTTAGC TCCATCATTC TGTCCAAAAC CTACGACAAT
     721  AACGTTATCT GTGCAAGCGA GCAAAGCATC CTGGTGATGA ATAGCATTTA CGAGAAGGTC
     781  AAAGAAGAGT TTGTTAAGCG TGGTTCCTAT ATTCTGAACC AGAACGAGAT TGCAAAAATC
     841  AAAGAAACGA TGTTTAAGAA TGGCGCGATT AATGCAGATA TTGTTGGTAA GTCCGCATAC
     901  ATTATCGCGA AGATGGCGGG TATCGAAGTG CCGCAAACCA CCAAAATCTT GATCGGCGAA
     961  GTGCAGAGCG TGGAGGCGAG CGAACTGTTC AGCCATGAGA AACTGAGCCC GGTTCTGGCT
    1021  ATGTACAAAG TGAAGGACTT TGACGAAGCC CTGAAGAAAG CTCAGCGTCT GATTGAGCTG
    1081  GGCGGCAGCG GTCACACGAG CAGCCTGTAT ATCGATAGCC AAAACAATAA GGACAAGGTG
    1141  AAAGAGTTCG GTCTGGCGAT GAAAACCTCT ATCACGTTTA TCAACATGCC GAGCAGCCAA
    1201  GGTGCGAGCG GCGATCTGTA TAACTTCGCA ATCGCGCCTT CTTTTACGCT GGGTTGCGGT
    1261  ACCTGGGAGG GTAATTCGGT GAGCCAGAAT GTCGAGCCGA AGCACCTGCT GAACATCAAA
    1321  TCGGTCGCGG AGCGCCGTGA GAAGAGACCG TGGTTCAAAG TCCCACAGAA AATTTACTTC
    1381  AAATACGGCT GCCTGCGTTT CGCGCTGAAA GAACTGAAAG ACATGAACAA AAAGCGTGCG
    1441  TTCATTGTTA CCGACAAAGA CCTGTTCAAA CTGGGTTACG TGAACAAAAT CACCAAAGTT
    1501  CTGGATGAAA TTGACATCAA GTACTCCATC TTCACTGATA TCAAATCCGA CCCAACGATT
    1561  GATAGCGTGA AAAAGGGCGC TAAAGAAATG CTGAACTTTG AACCGGACAC CATCATCAGC
    1621  ATCGGTGGTG GCTCTCCTAT GGATGCTGCG AAGGTCATGC ACCTGCTGTA CGAATACCCG
    1681  GAAGCGGAAA TCGAAAACCT GGCTATCAAC TTCATGGACA TCCGCAAACG TATCTGCAAC
    1741  TTCCCGAAGC TGGGCACTAA AGCTATTTCC GTTGCCATCC CGACTACCGC GGGCACTGGT
    1801  TCCGAAGCCA CGCCGTTCGC CGTGATCACC AACGATGAAA CCGGTATGAA ATACCCGCTG
    1861  ACCTCTTACG AACTGACCCC GAACATGGCA ATTATCGACA CCGAGCTGAT GCTGAACATG
    1921  CCGCGCAAGC TGACCGCTGC TACCGGCATC GACGCTCTGG TACATGCTAT TGAGGCGTAC
    1981  GTTTCCGTGA TGGCTACCGA TTACACCGAC GAACTGGCCC TGCGTGCGAT CAAAATGATT
    2041  TTCAAGTACC TGCCTCGCGC TTACAAAAAC GGCACGAATG ACATCGAGGC GCGTGAGAAA
    2101  ATGGCCCATG CAAGCAACAT CGCGGGCATG GCCTTCGCCA ACGCGTTCCT GGGCGTGTGC
    2161  CACTCTATGG CTCACAAACT GGGTGCTATG CACCACGTGC CGCACGGTAT CGCGTGTGCT
    2221  GTCCTGATCG AAGAAGTAAT TAAGTACAAC GCTACTGATT GCCCGACTAA ACAGACCGCC
    2281  TTCCCACAGT ACAAATCTCC TAACGCTAAA CGTAAGTACG CTGAGATCGC CGAATACCTG
    2341  AACCTGAAGG GTACGAGCGA CACTGAGAAA GTTACTGCGC TGATCGAAGC TATCTCTAAA
    2401  CTGAAAATTG ACCTGTCCAT CCCGCAGAAC ATCAGCGCCG CAGGCATCAA CAAAAAGGAC
    2461  TTTTACAACA CGCTGGACAA AATGAGCGAA CTGGCTTTTG ACGACCAGTG CACCACTGCA
    2521  AACCCGCGTT ACCCGCTGAT CTCTGAGCTG AAAGACATTT ATATCAAATC TTTTTAA
```

**AdhE2.83 (77012)**

```
       1  ATGAAAGTGA CGAACCAAAA AGAACTGAAA CAAAAACTGA ATGAACTGCG TGAGGCCCAG
      61  AAAAAGTTCG CTACGTATAC CCAGGAGCAA GTTGACAAGA TCTTCAAACA GTGTGCCATT
```

121 GCGGCTGCCA AAGAGCGTAT TAACTTGGCC AAACTGGCGG TGGAAGAAAC CGGCATCGGT

181 CTGGTCGAGG ACAAGATTAT CAAGAATCAC TTTGCCGCGG AGTATATCTA CAACAAATAC

```
     241  AAAAACGAAA AGACTTGCGG CATCATTGAT CATGATGACA GCTTGGGTAT CACGAAAGTC
     301  GCAGAACCAA TCGGTATTGT CGCGGCAATT GTCCCGACCA CTAATCCGAC CAGCACCGCG
     361  ATCTTCAAGA GCTTGATCAG CCTGAAAACC CGCAACGCGA TTTTCTTCAG CCCGCACCCG
     421  CGTGCGAAGA AGAGCACCAT CGCAGCGGCG ACCCTGATTT TGGATGCCGC GGTTAAGGCC
     481  GGTGCACCGA AAAACATTAT TGGCTGGATT GACGAGCCGA GCATTGAGCT GTCTCAGGAC
     541  CTGATGAGCG AAGCGGACAT TATTCTGGCC ACCGGTGGTC CGTCTATGGT TAAGGCAGCG
     601  TACTCCAGCG GTAAGCCTGC TATCGGCGTT GGTGCGGGCA ATACGCCGGC TATCATCGAT
     661  GAAAGCGCGG ATATTGATAT GGCAGTTAGC TCCATCATTC TGGGTAAAAC CTACGACAAT
     721  GGTGTTATCT GTGCAAGCGA GCAAAGCATC CTGGTGATGA ATAGCATTTA CGAGAAGGTC
     781  AAAGAAGAGT TTGTTAAGCG TGGTTCCTAT ATTCTGAACC AGAACGAGAT TGCAAAAATC
     841  AAAGAAACGA TGTTTAAGAA TGGCGCGATT AATGCAGATA TTGTTGGTAA GTCCGCATAC
     901  ATTATCGCGA AGATGGCGGG TATCGAAGTG CCGCAAACCA CCAAAATCTT GATCGGCGAA
     961  GTGCAGAGCG TGGAGAAGAG CGAACTGTTC AGCCATGAGC TGCTGAGCCC GGTTCTGGCT
    1021  ATGTACAAAG TGAAGGACTT TGACGAAGCC CTGAAGAAAG CTCAGCGTCT GATTGAGCTG
    1081  GGCGGCAGCC ACCACACGAG CAGCCTGTAT ATCGATAGCC AAAACAATAA GGACAAGGTG
    1141  AAAGAGTTCG GTCTGGAGAT GAAAACCTCT CGCACGTTTA TCAACATGCC GAGCAGCCAA
    1201  GGTGCGAGCG GCGATCTGTA TAACTTCGCA ATCGCGCCTT CTTTTACGCT GGGTTGCGGT
    1261  ACCTGGGGCG GTAATTCGGT GAGCCAGAAT GTCGAGCCGA AGCACCTGCT GAACATCAAA
    1321  TCGGTCGCGG AGCGCCGTGA GAAGAGACCG TGGTTCAAAG TCCCACAGAA AATTTACTTC
    1381  AAATACGGCT GCCTGCGTTT CGCGCTGAAA GAACTGAAAG ACATGAACAA AAAGCGTGCG
    1441  TTCATTGTTA CCGACAAAGA CCTGTTCAAA CTGGGTTACG TGAACAAAAT CACCAAAGTT
    1501  CTGGATGAAA TTGACATCAA GTACTCCATC TTCACTGATA TCAAATCCGA CCCAACGATT
    1561  GATAGCGTGA AAAAGGGCGC TAAAGAAATG CTGAACTTTG AACCGGACAC CATCATCAGC
    1621  ATCGGTGGTG GCTCTCCTAT GGATGCTGCG AAGGTCATGC ACCTGCTGTA CGAATACCCG
    1681  GAAGCGGAAA TCGAAAACCT GGCTATCAAC TTCATGGACA TCCGCAAACG TATCTGCAAC
    1741  TTCCCGAAGC TGGGCACTAA AGCTATTTCC GTTGCCATCC CGACTACCGC GGGCACTGGT
    1801  TCCGAAGCCA CGCCGTTCGC CGTGATCACC AACGATGAAA CCGGTATGAA ATACCCGCTG
    1861  ACCTCTTACG AACTGACCCC GAACATGGCA ATTATCGACA CCGAGCTGAT GCTGAACATG
    1921  CCGCGCAAGC TGACCGCTGC TACCGGCATC GACGCTCTGG TACATGCTAT TGAGGCGTAC
    1981  GTTTCCGTGA TGGCTACCGA TTACACCGAC GAACTGGCCC TGCGTGCGAT CAAAATGATT
    2041  TTCAAGTACC TGCCTCGCGC TTACAAAAAC GGCACGAATG ACATCGAGGC GCGTGAGAAA
    2101  ATGGCCCATG CAAGCAACAT CGCGGGCATG GCCTTCGCCA ACGCGTTCCT GGGCGTGTGC
    2161  CACTCTATGG CTCACAAACT GGGTGCTATG CACCACGTGC CGCACGGTAT CGCGTGTGCT
    2221  GTCCTGATCG AAGAAGTAAT TAAGTACAAC GCTACTGATT GCCCGACTAA ACAGACCGCC
    2281  TTCCCACAGT ACAAATCTCC TAACGCTAAA CGTAAGTACG CTGAGATCGC CGAATACCTG
    2341  AACCTGAAGG GTACGAGCGA CACTGAGAAA GTTACTGCGC TGATCGAAGC TATCTCTAAA
    2401  CTGAAAATTG ACCTGTCCAT CCCGCAGAAC ATCAGCGCCG CAGGCATCAA CAAAAAGGAC
    2461  TTTTACAACA CGCTGGACAA AATGAGCGAA CTGGCTTTTG ACGACCAGTG CACCACTGCA
    2521  AACCCGCGTT ACCCGCTGAT CTCTGAGCTG AAAGACATTT ATATCAAATC TTTTTAA
```

**AdhE2.84 (77013)**

```
       1  ATGAAAGTGA CGAACCAAAA AGAACTGAAA CAAAAACTGA ATGAACTGCG TGAGGCCCAG
      61  AAAAAGTTCG CTACGTATAC CCAGGAGCAA GTTGAGAAGT TCTTCAAACA GTGTGCCATT
```

121 GCGGCTGCCA AAGAGCGTAT TAACTTGGCC AAACTGGCGG TGGAAGAAAC CGGCATCGGT

181 CTGGTCGAGG ACAAGATTAT CAAGAATCAC TTTGCCGCGG AGTATATCTA CAACAAATAC

```
     241  AAAAACGAAA AGACTTGCGG CATCATTGAT CATGATGACA GCTTGGGTAT CACGAAAGTC
     301  GCAGAACCAA TCGGTATTGT CGCGAGCATT GTCCCGACCA CTAATCCGAC CAGCACCGCG
     361  ATCTTCAAGA GCTTGATCAG CCTGAAAACC CGCAACGCGA TTTTCTTCAG CCCGCACCCG
     421  CGTGCGAAGA AGAGCACCAT CGCAGCGGCG AAGCTGATTT TGGATGCCGC GGTTAAGGCC
     481  GGTGCACCGA AAAACATTAT TGGCTGGATT GACGAGCCGA GCATTGAGCT GTCTCAGGAC
     541  CTGATGAGCG AAGCGGACAT TATTCTGGCC ACCGGTGGTC CGTCTATGGT TAAGGCAGCG
     601  TACTCCAGCG GTAAGCCTGC TATCGGCGTT GGTGCGGGCA ATACGCCGGC TATCATCGAT
     661  GAAAGCGCGG ATATTGATAT GGCAGTTAGC TCCATCATTC TGTCCAAAAC CTACGACAAT
     721  GGTGTTATCT GTGCAAGCGA GCAAAGCATC CTGGTGATGA ATAGCATTTA CGAGAAGGTC
     781  AAAGAAGAGT TTGTTAAGCG TGGTTCCTAT ATTCTGAACC AGAACGAGAT TGCAAAAATC
     841  AAAGAAACGA TGTTTAAGAA TGGCGCGATT AATGCAGATA TTGTTGGTAA GTCCGCATAC
     901  ATTATCGCGA AGATGGCGGG TATCGAAGTG CCGCAAACCA CCAAAATCTT GATCGGCGAA
     961  GTGCAGAGCG TGGAGAAGAG CGAACTGTTC AGCCATGAGA AACTGAGCCC GGTTCTGGCT
    1021  ATGTACAAAG TGAAGGACTT TGACGAAGCC CTGAAGAAAG CTCAGCGTCT GATTGAGCTG
    1081  GGCGGCAGCG GTCACACGAG CAGCCTGTAT ATCGATAGCC AAAACAATAA GGACAAGGTG
    1141  AAAGAGTTCG GTCTGGCGAT GAAAACCTCT CGCACGTTTA TCAACATGCC GAGCAGCCAA
    1201  GGTGCGAGCG GCGATCTGTA TAACTTCGCA ATCGCGCCTT CTTTTACGCT GGGTTGCCCG
    1261  ACCTGGGGCG GTAATTCGGT GAGCCAGAAT GTCGAGCCGA AGCACCTGCT GAACATCAAA
    1321  TCGGTCGCGC GTCGCCGTGA GAAGAGACCG TGGTTCAAAG TCCCACAGAA AATTTACTTC
    1381  AAATACGGCT GCCTGCGTTT CGCGCTGAAA GAACTGAAAG ACATGAACAA AAAGCGTGCG
    1441  TTCATTGTTA CCGACAAAGA CCTGTTCAAA CTGGGTTACG TGAACAAAAT CACCAAAGTT
    1501  CTGGATGAAA TTGACATCAA GTACTCCATC TTCACTGATA TCAAATCCGA CCCAACGATT
    1561  GATAGCGTGA AAAAGGGCGC TAAAGAAATG CTGAACTTTG AACCGGACAC CATCATCAGC
    1621  ATCGGTGGTG GCTCTCCTAT GGATGCTGCG AAGGTCATGC ACCTGCTGTA CGAATACCCG
    1681  GAAGCGGAAA TCGAAAACCT GGCTATCAAC TTCATGGACA TCCGCAAACG TATCTGCAAC
    1741  TTCCCGAAGC TGGGCACTAA AGCTATTTCC GTTGCCATCC CGACTACCGC GGGCACTGGT
    1801  TCCGAAGCCA CGCCGTTCGC CGTGATCACC AACGATGAAA CCGGTATGAA ATACCCGCTG
    1861  ACCTCTTACG AACTGACCCC GAACATGGCA ATTATCGACA CCGAGCTGAT GCTGAACATG
    1921  CCGCGCAAGC TGACCGCTGC TACCGGCATC GACGCTCTGG TACATGCTAT TGAGGCGTAC
    1981  GTTTCCGTGA TGGCTACCGA TTACACCGAC GAACTGGCCC TGCGTGCGAT CAAAATGATT
    2041  TTCAAGTACC TGCCTCGCGC TTACAAAAAC GGCACGAATG ACATCGAGGC GCGTGAGAAA
    2101  ATGGCCCATG CAAGCAACAT CGCGGGCATG GCCTTCGCCA ACGCGTTCCT GGGCGTGTGC
    2161  CACTCTATGG CTCACAAACT GGGTGCTATG CACCACGTGC CGCACGGTAT CGCGTGTGCT
    2221  GTCCTGATCG AAGAAGTAAT TAAGTACAAC GCTACTGATT GCCCGACTAA ACAGACCGCC
    2281  TTCCCACAGT ACAAATCTCC TAACGCTAAA CGTAAGTACG CTGAGATCGC CGAATACCTG
    2341  AACCTGAAGG GTACGAGCGA CACTGAGAAA GTTACTGCGC TGATCGAAGC TATCTCTAAA
    2401  CTGAAAATTG ACCTGTCCAT CCCGCAGAAC ATCAGCGCCG CAGGCATCAA CAAAAAGGAC
    2461  TTTTACAACA CGCTGGACAA AATGAGCGAA CTGGCTTTTG ACGACCAGTG CACCACTGCA
    2521  AACCCGCGTT ACCCGCTGAT CTCTGAGCTG AAAGACATTT ATATCAAATC TTTTTAA
```

**AdhE2.85 (77014)**

```
       1  ATGAAAGTGA CGAACCAAAA AGAACTGAAA CAAAAACTGA ATGAACTGCG TGAGGCCCAG
      61  AAAAAGTTCG CTACGTATAC CCAGGAGCAA GTTGACAAGA TCTTCAAACA GTGTGCCATT
```

121 GCGGCTGCCA AAGAGCGTAT TAACTTGGCC AAACTGGCGG TGGAAGAAAC CGGCATCGGT

181 CTGGTCGAGG ACAAGATTAT CAAGAATCAC TTTGCCGCGG AGTATATCTA CAACAAATAC

```
     241  AAAAACGAAA AGACTTGCGG CATCATTGAT CATGATGACA GCTTGGGTAT CACGAAAGTC
     301  GCAGAACCAA TCGGTATTGT CGCGGCAATT GTCCCGACCA CTAATCCGAC CAGCACCGCG
     361  ATCTTCAAGA GCTTGATCAG CCTGAAAACC CGCAACGCGA TTTTCTTCAG CCCGCACCCG
     421  CGTGCGAAGA AGAGCACCAT CGCAGCGGCG AAGCTGATTT TGGATGCCGC GGTTAAGGCC
     481  GGTGCACCGA AAAACATTAT TGGCTGGATT GACGAGCCGA GCATTGAGCT GTCTCAGGAC
     541  CTGATGAGCG AAGCGGACAT TATTCTGGCC ACCGGTGGTC CGTCTATGGT TAAGGCAGCG
     601  TACTCCAGCG GTAAGCCTGC TATCGGCGTT GGTGCGGGCA ATACGCCGGC TATCATCGAT
     661  GAAAGCGCGG ATATTGATAT GGCAGTTAGC TCCATCATTC TGTCCAAAAC CTACGACAAT
     721  GGTGTTATCT GTGCAAGCGA GCAAAGCATC CTGGTGATGA ATAGCATTTA CGAGAAGGTC
     781  AAAGAAGAGT TTGTTAAGCG TGGTTCCTAT ATTCTGAACC AGCAGGAGAT TGCAAAAATC
     841  AAAGAAACGA TGTTTAAGAA TGGCGCGATT AATGCAGATA TTGTTGGTAA GTCCGCATAC
     901  ATTATCGCGA AGATGGCGGG TATCGAAGTG CCGCAAACCA CCAAAATCTT GATCGGCGAA
     961  GTGCAGAGCG TGGAGAAGAG CGAACTGTTC AGCAAAGAGA AACTGAGCCC GGTTCTGGCT
    1021  ATGTACAAAG TGAAGGACTT TGACGAAGCC CTGAAGAAAG CTCAGCGTCT GATTGAGCTG
    1081  GGCGGCAGCC ACCACACGAG CAGCCTGTAT ATCGATAGCC AAAACAATAA GGACAAGGTG
    1141  AAAGAGTTCG GTCTGCGTAT GAAAACCTCT CGCTTCTTTA TCAACATGCC GAGCAGCCAA
    1201  GGTGCGAGCG GCGATCTGTA TAACTTCGCA ATCGCGCCTT CTTTTACGCT GGGTTGCGGT
    1261  ACCTGGGGCG GTAATTCGGT GAGCCAGAAT GTCGAGCCGA AGCACCTGCT GAACATCAAA
    1321  TCGGTCGCGG AGCGCCGTGA GAAGAGACCG TGGTTCAAAG TCCCACAGAA AATTTACTTC
    1381  AAATACGGCT GCCTGCGTTT CGCGCTGAAA GAACTGAAAG ACATGAACAA AAAGCGTGCG
    1441  TTCATTGTTA CCGACAAAGA CCTGTTCAAA CTGGGTTACG TGAACAAAAT CACCAAAGTT
    1501  CTGGATGAAA TTGACATCAA GTACTCCATC TTCACTGATA TCAAATCCGA CCCAACGATT
    1561  GATAGCGTGA AAAAGGGCGC TAAAGAAATG CTGAACTTTG AACCGGACAC CATCATCAGC
    1621  ATCGGTGGTG GCTCTCCTAT GGATGCTGCG AAGGTCATGC ACCTGCTGTA CGAATACCCG
    1681  GAAGCGGAAA TCGAAAACCT GGCTATCAAC TTCATGGACA TCCGCAAACG TATCTGCAAC
    1741  TTCCCGAAGC TGGGCACTAA AGCTATTTCC GTTGCCATCC CGACTACCGC GGGCACTGGT
    1801  TCCGAAGCCA CGCCGTTCGC CGTGATCACC AACGATGAAA CCGGTATGAA ATACCCGCTG
    1861  ACCTCTTACG AACTGACCCC GAACATGGCA ATTATCGACA CCGAGCTGAT GCTGAACATG
    1921  CCGCGCAAGC TGACCGCTGC TACCGGCATC GACGCTCTGG TACATGCTAT TGAGGCGTAC
    1981  GTTTCCGTGA TGGCTACCGA TTACACCGAC GAACTGGCCC TGCGTGCGAT CAAAATGATT
    2041  TTCAAGTACC TGCCTCGCGC TTACAAAAAC GGCACGAATG ACATCGAGGC GCGTGAGAAA
    2101  ATGGCCCATG CAAGCAACAT CGCGGGCATG GCCTTCGCCA ACGCGTTCCT GGGCGTGTGC
    2161  CACTCTATGG CTCACAAACT GGGTGCTATG CACCACGTGC CGCACGGTAT CGCGTGTGCT
    2221  GTCCTGATCG AAGAAGTAAT TAAGTACAAC GCTACTGATT GCCCGACTAA ACAGACCGCC
    2281  TTCCCACAGT ACAAATCTCC TAACGCTAAA CGTAAGTACG CTGAGATCGC CGAATACCTG
    2341  AACCTGAAGG GTACGAGCGA CACTGAGAAA GTTACTGCGC TGATCGAAGC TATCTCTAAA
    2401  CTGAAAATTG ACCTGTCCAT CCCGCAGAAC ATCAGCGCCG CAGGCATCAA CAAAAAGGAC
    2461  TTTTACAACA CGCTGGACAA AATGAGCGAA CTGGCTTTTG ACGACCAGTG CACCACTGCA
    2521  AACCCGCGTT ACCCGCTGAT CTCTGAGCTG AAAGACATTT ATATCAAATC TTTTTAA
```

**AdhE2.86 (77015)**

```
       1  ATGAAAGTGA CGAACCAAAA AGAACTGAAA CAAAAACTGA ATGAACTGCG TGAGGCCCAG
      61  AAAAAGTTCA ACACGTATAC CCAGGAGCAA GTTGACAAGA TCTTCAAACA GTGTGCCATT
```

121 GCGGCTGCCA AAGAGCGTAT TAACTTGGCC AAACTGGCGG TGGAAGAAAC CGGCATCGGT

181 CTGGTCGAGG ACAAGATTAT CAAGAATCAC TTTGCCGCGG AGTATATCTA CAACAAATAC

```
     241  AAAAACGAAA AGACTTGCGG CATCATTGAT CATGATGACA GCTTGGGTAT CACGAAAGTC
     301  GCAGAACCAA TCGGTATTGT CGCGGCAATT GTCCCGACCA CTAATCCGAC CAGCACCGCG
     361  ATCTTCAAGA GCTTGATCAG CCTGAAAACC CGCAACGCGA TTTTCTTCAG CCCGCACCCG
     421  CGTGCGAAGA AGAGCACCAT CGCAGCGGCG AAGCTGATTT TGGATGCCGC GGTTAAGGCC
     481  GGTGCACCGA AAAACATTAT TGGCTGGATT GACGAGCCGA GCATTGAGCT GTCTCAGGAC
     541  CTGATGAAAG AAGCGGACAT TATTCTGGCC ACCGGTGGTC CGTCTATGGT TAAGGCAGCG
     601  TACTCCAGCG GTAAGCCTGC TATCGGCGTT GGTGCGGGCA ATACGCCGGC TATCATCGAT
     661  GAAAGCGCGG ATATTGATAT GGCAGTTAGC TCCATCATTC TGTCCAAAAC CTACGACAAT
     721  GGTGTTATCT GTGCAAGCGA GCAAAGCATC CTGGTGATGA ATAGCATTTA CGAGAAGGTC
     781  AAAGAAGAGT TTGTTAAGCG TGGTTCCTAT ATTCTGAACC AGAACGAGAT TGCAAAAATC
     841  AAAGAAACGA TGTTTAAGAA TGGCGCGATT AATGCAGATA TTGTTGGTAA GTCCGCATAC
     901  ATTATCGCGA AGATGGCGGG TATCGAAGTG CCGCAATTCA CCAAAATCTT GATCGGCGAA
     961  GTGCAGAGCG TGGAGAAGAG CGAACTGTTC AGCCATGAGA AACTGAGCCC GGTTCTGGCT
    1021  ATGTACAAAG TGAAGGACTT TGACGAAGCC CTGAAGAAAG CTCAGCGTCT GATTAACCTG
    1081  GGCGGCAGCG GTCACACGAG CAGCCTGTAT AGCGATAGCC AAAACAATAA GGACAAGGTG
    1141  AAAGAGTTCG GTCTGGCGAT GAAAACCTCT CGCACGTTTA TCAACATGCC GAGCAGCCAA
    1201  GGTGCGAGCG GCGATCTGTA TAACTTCGCA ATCGCGCCTT CTTTTACGCT GGGTTGCGGT
    1261  ACCTGGGGCG GTAATTCGGT GAGCCAGAAT GTCGAGCCGA AGCACCTGCT GAACATCAAA
    1321  TCGGTCGCGG AGCGCCGTGA GAAGAGACCG TGGTTCAAAG TCCCACAGAA AATTTACTTC
    1381  AAATACGGCT GCCTGCGTTT CGCGCTGAAA GAACTGAAAG ACATGAACAA AAAGCGTGCG
    1441  TTCATTGTTA CCGACAAAGA CCTGTTCAAA CTGGGTTACG TGAACAAAAT CACCAAAGTT
    1501  CTGGATGAAA TTGACATCAA GTACTCCATC TTCACTGATA TCAAATCCGA CCCAACGATT
    1561  GATAGCGTGA AAAAGGGCGC TAAAGAAATG CTGAACTTTG AACCGGACAC CATCATCAGC
    1621  ATCGGTGGTG GCTCTCCTAT GGATGCTGCG AAGGTCATGC ACCTGCTGTA CGAATACCCG
    1681  GAAGCGGAAA TCGAAAACCT GGCTATCAAC TTCATGGACA TCCGCAAACG TATCTGCAAC
    1741  TTCCCGAAGC TGGGCACTAA AGCTATTTCC GTTGCCATCC CGACTACCGC GGGCACTGGT
    1801  TCCGAAGCCA CGCCGTTCGC CGTGATCACC AACGATGAAA CCGGTATGAA ATACCCGCTG
    1861  ACCTCTTACG AACTGACCCC GAACATGGCA ATTATCGACA CCGAGCTGAT GCTGAACATG
    1921  CCGCGCAAGC TGACCGCTGC TACCGGCATC GACGCTCTGG TACATGCTAT TGAGGCGTAC
    1981  GTTTCCGTGA TGGCTACCGA TTACACCGAC GAACTGGCCC TGCGTGCGAT CAAAATGATT
    2041  TTCAAGTACC TGCCTCGCGC TTACAAAAAC GGCACGAATG ACATCGAGGC GCGTGAGAAA
    2101  ATGGCCCATG CAAGCAACAT CGCGGGCATG GCCTTCGCCA ACGCGTTCCT GGGCGTGTGC
    2161  CACTCTATGG CTCACAAACT GGGTGCTATG CACCACGTGC CGCACGGTAT CGCGTGTGCT
    2221  GTCCTGATCG AAGAAGTAAT TAAGTACAAC GCTACTGATT GCCCGACTAA ACAGACCGCC
    2281  TTCCCACAGT ACAAATCTCC TAACGCTAAA CGTAAGTACG CTGAGATCGC CGAATACCTG
    2341  AACCTGAAGG GTACGAGCGA CACTGAGAAA GTTACTGCGC TGATCGAAGC TATCTCTAAA
    2401  CTGAAAATTG ACCTGTCCAT CCCGCAGAAC ATCAGCGCCG CAGGCATCAA CAAAAAGGAC
    2461  TTTTACAACA CGCTGGACAA AATGAGCGAA CTGGCTTTTG ACGACCAGTG CACCACTGCA
    2521  AACCCGCGTT ACCCGCTGAT CTCTGAGCTG AAAGACATTT ATATCAAATC TTTTTAA
```

**AdhE2.87 (77016)**

```
       1  ATGAAAGTGA CGAACCAAAA AGAACTGAAA CAAAAAGACA ATGAACTGCG TGAGGCCCAG
      61  AAAAAGTTCG CTACGTGCAC CCAGGACCAA GTTGACAAGA TCTTCAAACA GTGTGCCATT
```

121 GCGGCTGCCA AAGAGCGTAT TAACTTGGCC AAACTGGCGG TGGAAGAAAC CGGCATCGGT

181 CTGGTCGAGG ACAAGATTAT CAAGAATCAC TTTGCCGCGG AGTATATCTA CAACAAATAC

```
     241  AAAAACGAAA AGACTTGCGG CATCATTGAT CATGATGACA GCTTGGGTAT CACGAAAGTC
     301  GCAGAACCAA TCGGTATTGT CGCGGCAATT GTCCCGACCA CTAATCCGAC CAGCACCGCG
     361  ATCTTCAAGA GCTTGATCAG CCTGAAAACC CGCAACGCGA TTTTCTTCAG CCCGCACCCG
     421  CGTGCGAAGA AGAGCACCAT CGCAGCGGCG AAGCTGATTT TGGATGCCGC GGTTAAGGCC
     481  GGTGCACCGA AAAACATTAT TGGCTGGATT GACGAGCCGA GCATTGAGCT GTCTCAGGAC
     541  CTGATGAGCG AAGCGGACAT TATTCTGGCC ACCGGTGGTC CGTCTATGGT TAAGGCAGCG
     601  TACTCCAGCG GTAAGCCTGC TATCGGCGTT GGTGCGGGCA ATACGCCGGC TATCATCGAT
     661  GAAAGCGCGG ATATTGATAT GGCAGTTAGC TCCATCATTC TGGGTAAAAC CTACGACAAT
     721  GGTGTTATCT GTGCAAGCGA GCAAAGCATC CTGGTGATGA ATAGCATTTA CGAGAAGGTC
     781  AAAGAAGAGT TTGTTAAGCG TGGTTCCTAT ATTCTGAACC AGAACGAGAT TGCAAAAATC
     841  AAAGAAACGA TGTTTAAGAA TGGCGCGATT AATGCAGATA TTGTTGGTAA GTCCGCATAC
     901  ATTATCGCGA AGATGGCGGG TATCGAAGTG CCGCAAACCA CCAAAATCTT GATCGGCGAA
     961  GTGCAGAGCG TGGAGAAGAG CGAACTGTTC AGCAAAGAGA AACTGAGCCC GGTTCTGGCT
    1021  ATGTACAAAG TGAAGGACTT TGACGAAGCC CTGAAGAAAG CTCAGCGTCT GATTGAGCTG
    1081  GGCGGCAGCG GTCACACGAG CAGCCTGTAT ATCGATAGCC AAAACAATAA GGACAAGGTG
    1141  AAAGAGTTCG GTCTGGCGAT GAAAACCTCT CGCACGTTTA TCAACATGCC GAGCAGCCAA
    1201  GGTGCGAGCG GCGATCTGTA TAACTTCGCA ATCGCGCCTT CTTTTACGCT GGGTTGCGGT
    1261  ACCTGGGGCG GTAATTCGGT GAGCCAGAAT GTCGAGCCGA AGCACCTGCT GAACATCAAA
    1321  TCGGTCGCGG AGCGCCGTGA GAAGAGACCG TGGTTCAAAG TCCCACAGAA AATTTACTTC
    1381  AAATACGGCT GCCTGCGTTT CGCGCTGAAA GAACTGAAAG ACATGAACAA AAAGCGTGCG
    1441  TTCATTGTTA CCGACAAAGA CCTGTTCAAA CTGGGTTACG TGAACAAAAT CACCAAAGTT
    1501  CTGGATGAAA TTGACATCAA GTACTCCATC TTCACTGATA TCAAATCCGA CCCAACGATT
    1561  GATAGCGTGA AAAAGGGCGC TAAAGAAATG CTGAACTTTG AACCGGACAC CATCATCAGC
    1621  ATCGGTGGTG GCTCTCCTAT GGATGCTGCG AAGGTCATGC ACCTGCTGTA CGAATACCCG
    1681  GAAGCGGAAA TCGAAAACCT GGCTATCAAC TTCATGGACA TCCGCAAACG TATCTGCAAC
    1741  TTCCCGAAGC TGGGCACTAA AGCTATTTCC GTTGCCATCC CGACTACCGC GGGCACTGGT
    1801  TCCGAAGCCA CGCCGTTCGC CGTGATCACC AACGATGAAA CCGGTATGAA ATACCCGCTG
    1861  ACCTCTTACG AACTGACCCC GAACATGGCA ATTATCGACA CCGAGCTGAT GCTGAACATG
    1921  CCGCGCAAGC TGACCGCTGC TACCGGCATC GACGCTCTGG TACATGCTAT TGAGGCGTAC
    1981  GTTTCCGTGA TGGCTACCGA TTACACCGAC GAACTGGCCC TGCGTGCGAT CAAAATGATT
    2041  TTCAAGTACC TGCCTCGCGC TTACAAAAAC GGCACGAATG ACATCGAGGC GCGTGAGAAA
    2101  ATGGCCCATG CAAGCAACAT CGCGGGCATG GCCTTCGCCA ACGCGTTCCT GGGCGTGTGC
    2161  CACTCTATGG CTCACAAACT GGGTGCTATG CACCACGTGC CGCACGGTAT CGCGTGTGCT
    2221  GTCCTGATCG AAGAAGTAAT TAAGTACAAC GCTACTGATT GCCCGACTAA ACAGACCGCC
    2281  TTCCCACAGT ACAAATCTCC TAACGCTAAA CGTAAGTACG CTGAGATCGC CGAATACCTG
    2341  AACCTGAAGG GTACGAGCGA CACTGAGAAA GTTACTGCGC TGATCGAAGC TATCTCTAAA
    2401  CTGAAAATTG ACCTGTCCAT CCCGCAGAAC ATCAGCGCCG CAGGCATCAA CAAAAAGGAC
    2461  TTTTACAACA CGCTGGACAA AATGAGCGAA CTGGCTTTTG ACGACCAGTG CACCACTGCA
    2521  AACCCGCGTT ACCCGCTGAT CTCTGAGCTG AAAGACATTT ATATCAAATC TTTTTAA
```

**AdhE2.88 (77017)**

```
       1  ATGAAAGTGA CGAACCAAAA AGAACTGAAA CAAAAACTGA ATGAACTGCG TGAGGCCCAG
      61  AAAAAGTTCA ACACGTATAC CCAGGAGCAA GTTGACAAGA TCTTCAAACA GTGTGCCATT
```

121 GCGGCTGCCA AAGAGCGTAT TAACTTGGCC AAACTGGCGG TGGAAGAAAC CGGCATCGGT

181 CTGGTCGAGG ACAAGATTAT CAAGAATCAC TTTGCCGCGG AGTATATCTA CAACAAAGAG

```
     241  AAAAACGAAA AGACTTGCGG CATCATTGAT CATGATGACA GCTTGGGTAT CACGAAAGTC
     301  GCAGAACCAA TCGGTATTGT CGCGAGCATT GTCCCGACCA CTAATCCGAC CAGCACCGCG
     361  ATCTTCAAGA GCTTGATCAG CCTGGCGACC CGCAACGCGA TTTTCTTCAG CCCGCACCCG
     421  CGTGCGAAGA AGAGCACCAT CGCAGCGGCG AAGCTGATTT TGGATGCCGC GGTTAAGGCC
     481  GGTGCACCGA AAAACATTAT TGGCTGGATT GACGAGCCGA GCATTGAGCT GTCTCAGGAC
     541  CTGATGAGCG AAGCGGACAT TATTCTGGCC ACCGGTGGTC CGTCTATGGT TAAGGCAGCG
     601  TACTCCAGCG GTAAGCCTGC TATCGGCGTT GGTGCGGGCA ATACGCCGGC TATCATCGAT
     661  GAAAGCGCGG ATATTGATAT GGCAGTTAGC TCCATCATTC TGTCCAAAAC CTACGACAAT
     721  GGTGTTATCT GTGCAAGCGA GCAAAGCATC CTGGTGATGA ATAGCATTTA CGAGAAGGTC
     781  AAAGAAGAGT TTGTTAAGCG TGGTTCCTAT ATTCTGAACC AGGAGGAGAT TGCAAAAATC
     841  AAAGAAACGA TGTTTAAGAA TGGCGCGATT AATGCAGATA TTGTTGGTAA GTCCGCATAC
     901  ATTATCGCGA AGATGGCGGG TATCGAAGTG CCGCAAACCA CCAAAATCTT GATCGGCGAA
     961  GTGCAGAGCG TGGAGAAGAG CGAACTGTTC AGCCATGAGA AACTGAGCCC GGTTCTGGCT
    1021  ATGTACAAAG TGAAGGACTT TGACGAAGCC CTGAAGAAAG CTCAGCGTCT GATTGAGCTG
    1081  GGCGGCAGCG GTCACACGAG CAGCCTGTAT ATCGATAGCC AAAACAATAA GGACAAGGTG
    1141  AAAGAGTTCG GTCTGGCGAT GAAAACCTCT CGCACGTTTA TCAACATGCC GAGCAGCCAA
    1201  GGTGCGAGCG GCGATCTGTA TAACTTCGCA ATCGCGCCTT CTTTTACGCT GGGTTGCGGT
    1261  ACCTGGGGCG GTAATTCGGT GAGCCAGAAT GTCGAGCCGA AGCACCTGCT GAACATCAAA
    1321  TCGGTCGCGG AGCGCCGTGA GAAGAGACCG TGGTTCAAAG TCCCACAGAA AATTTACTTC
    1381  AAATACGGCT GCCTGCGTTT CGCGCTGAAA GAACTGAAAG ACATGAACAA AAAGCGTGCG
    1441  TTCATTGTTA CCGACAAAGA CCTGTTCAAA CTGGGTTACG TGAACAAAAT CACCAAAGTT
    1501  CTGGATGAAA TTGACATCAA GTACTCCATC TTCACTGATA TCAAATCCGA CCCAACGATT
    1561  GATAGCGTGA AAAAGGGCGC TAAAGAAATG CTGAACTTTG AACCGGACAC CATCATCAGC
    1621  ATCGGTGGTG GCTCTCCTAT GGATGCTGCG AAGGTCATGC ACCTGCTGTA CGAATACCCG
    1681  GAAGCGGAAA TCGAAAACCT GGCTATCAAC TTCATGGACA TCCGCAAACG TATCTGCAAC
    1741  TTCCCGAAGC TGGGCACTAA AGCTATTTCC GTTGCCATCC CGACTACCGC GGGCACTGGT
    1801  TCCGAAGCCA CGCCGTTCGC CGTGATCACC AACGATGAAA CCGGTATGAA ATACCCGCTG
    1861  ACCTCTTACG AACTGACCCC GAACATGGCA ATTATCGACA CCGAGCTGAT GCTGAACATG
    1921  CCGCGCAAGC TGACCGCTGC TACCGGCATC GACGCTCTGG TACATGCTAT TGAGGCGTAC
    1981  GTTTCCGTGA TGGCTACCGA TTACACCGAC GAACTGGCCC TGCGTGCGAT CAAAATGATT
    2041  TTCAAGTACC TGCCTCGCGC TTACAAAAAC GGCACGAATG ACATCGAGGC GCGTGAGAAA
    2101  ATGGCCCATG CAAGCAACAT CGCGGGCATG GCCTTCGCCA ACGCGTTCCT GGGCGTGTGC
    2161  CACTCTATGG CTCACAAACT GGGTGCTATG CACCACGTGC CGCACGGTAT CGCGTGTGCT
    2221  GTCCTGATCG AAGAAGTAAT TAAGTACAAC GCTACTGATT GCCCGACTAA ACAGACCGCC
    2281  TTCCCACAGT ACAAATCTCC TAACGCTAAA CGTAAGTACG CTGAGATCGC CGAATACCTG
    2341  AACCTGAAGG GTACGAGCGA CACTGAGAAA GTTACTGCGC TGATCGAAGC TATCTCTAAA
    2401  CTGAAAATTG ACCTGTCCAT CCCGCAGAAC ATCAGCGCCG CAGGCATCAA CAAAAAGGAC
    2461  TTTTACAACA CGCTGGACAA AATGAGCGAA CTGGCTTTTG ACGACCAGTG CACCACTGCA
    2521  AACCCGCGTT ACCCGCTGAT CTCTGAGCTG AAAGACATTT ATATCAAATC TTTTTAA
```

**AdhE2.89 (77018)**

```
       1  ATGAAAGTGA CGAACCAAAA AGAACTGAAA CAAAAACTGA ATGAACTGCG TGAGGCCCAG
      61  AAAAAGTTCG CTAGCTATAC CCAGGAGCAA GTTGACAAGA TCTTCAAACA GTGTGCCATT
```

121 GCGGCTGCCA AAGAGCGTAT TAACTTGGCC GAGCTGGCGG TGGAAGAAAC CGGCATCGGT

181 CTGGTCGAGG ACAAGATTAT CAAGAATCAC TTTGCCGCGG AGTATATCTA CAACAAAGAG

```
     241  AAAAACGAAA AGACTTGCGG CATCATTGAT CATGATGACA GCTTGGGTAT CACGAAAGTC
     301  GCAGAACCAA TCGGTATTGT CGCGGCAATT GTCCCGACCA CTAATCCGAC CAGCACCGCG
     361  ATCTTCAAGA GCTTGATCAG CCTGAAAACC CGCAACGCGA TTTTCTTCAG CCCGCACCCG
     421  CGTGCGAAGA AGAGCACCAT CGCAGCGGCG AAGCTGATTT TGGATGCCGC GGTTAAGGCC
     481  GGTGCACCGA AAAACATTAT TGGCTGGATT GACGAGCCGA GCATTGAGCT GTCTCAGGAC
     541  CTGATGAGCG AAGCGGACAT TATTCTGGCC ACCGGTGGTC CGTCTATGGT TAAGGCAGCG
     601  TACTCCAGCG GTAAGCCTGC TATCGGCGTT GGTGCGGGCA ATACGCCGGC TATCATCGAT
     661  GAAAGCGCGG ATATTGATAT GGCAGTTAGC TCCATCATTC TGTCCAAAAC CTACGACAAT
     721  GGTGTTATCT GTGCAAGCGA GCAAAGCATC CTGGTGATGA ATAGCATTTA CGAGAAGGTC
     781  AAAGAAGAGT TTGTTAAGCG TGGTTCCTAT ATTCTGAACC AGAACGAGAT TGCAAAAATC
     841  AAAGAAACGA TGTTTAAGAA TGGCGCGATT AATGCAGATA TTGTTGGTAA GTCCGCATAC
     901  ATTATCGCGA AGATGGCGGG TATCGAAGTG CCGCAATTCA CCAAAATCTT GATCGGCGAA
     961  GTGCAGAGCG TGGAGAAGAG CGAACTGTTC AGCCATGAGA AACTGAGCCC GGTTCTGGCT
    1021  ATGTACAAAG TGAAGGACTT TGACGAAGCC CTGAAGAAAG CTCAGCGTCT GATTGAGCTG
    1081  GGCGGCAGCG GTCACACGAG CAGCCTGTAT ATCGATAGCC AAAACAATAA GGACAAGGTG
    1141  AAAGAGTTCG GTCTGGAGAT GAAAACCTCT CGCACGTTTA TCAACATGCC GAGCAGCCAA
    1201  GGTGCGAGCG GCGATCTGTA TAACTTCGCA ATCGCGCCTT CTTTTACGCT GGGTTGCGGT
    1261  ACCTGGGGCG GTAATTCGGT GAGCCAGAAT GTCGAGCCGA AGCACCTGCT GAACATCAAA
    1321  TCGGTCGCGG AGCGCCGTGA GAAGAGACCG TGGTTCAAAG TCCCACAGAA AATTTACTTC
    1381  AAATACGGCT GCCTGCGTTT CGCGCTGAAA GAACTGAAAG ACATGAACAA AAAGCGTGCG
    1441  TTCATTGTTA CCGACAAAGA CCTGTTCAAA CTGGGTTACG TGAACAAAAT CACCAAAGTT
    1501  CTGGATGAAA TTGACATCAA GTACTCCATC TTCACTGATA TCAAATCCGA CCCAACGATT
    1561  GATAGCGTGA AAAAGGGCGC TAAAGAAATG CTGAACTTTG AACCGGACAC CATCATCAGC
    1621  ATCGGTGGTG GCTCTCCTAT GGATGCTGCG AAGGTCATGC ACCTGCTGTA CGAATACCCG
    1681  GAAGCGGAAA TCGAAAACCT GGCTATCAAC TTCATGGACA TCCGCAAACG TATCTGCAAC
    1741  TTCCCGAAGC TGGGCACTAA AGCTATTTCC GTTGCCATCC CGACTACCGC GGGCACTGGT
    1801  TCCGAAGCCA CGCCGTTCGC CGTGATCACC AACGATGAAA CCGGTATGAA ATACCCGCTG
    1861  ACCTCTTACG AACTGACCCC GAACATGGCA ATTATCGACA CCGAGCTGAT GCTGAACATG
    1921  CCGCGCAAGC TGACCGCTGC TACCGGCATC GACGCTCTGG TACATGCTAT TGAGGCGTAC
    1981  GTTTCCGTGA TGGCTACCGA TTACACCGAC GAACTGGCCC TGCGTGCGAT CAAAATGATT
    2041  TTCAAGTACC TGCCTCGCGC TTACAAAAAC GGCACGAATG ACATCGAGGC GCGTGAGAAA
    2101  ATGGCCCATG CAAGCAACAT CGCGGGCATG GCCTTCGCCA ACGCGTTCCT GGGCGTGTGC
    2161  CACTCTATGG CTCACAAACT GGGTGCTATG CACCACGTGC CGCACGGTAT CGCGTGTGCT
    2221  GTCCTGATCG AAGAAGTAAT TAAGTACAAC GCTACTGATT GCCCGACTAA ACAGACCGCC
    2281  TTCCCACAGT ACAAATCTCC TAACGCTAAA CGTAAGTACG CTGAGATCGC CGAATACCTG
    2341  AACCTGAAGG GTACGAGCGA CACTGAGAAA GTTACTGCGC TGATCGAAGC TATCTCTAAA
    2401  CTGAAAATTG ACCTGTCCAT CCCGCAGAAC ATCAGCGCCG CAGGCATCAA CAAAAAGGAC
    2461  TTTTACAACA CGCTGGACAA AATGAGCGAA CTGGCTTTTG ACGACCAGTG CACCACTGCA
    2521  AACCCGCGTT ACCCGCTGAT CTCTGAGCTG AAAGACATTT ATATCAAATC TTTTTAA
```

**AdhE2.90 (77019)**

```
       1  ATGAAAGTGA CGAACCAAAA AGAACTGAAA CAAAAACTGA ATGAACTGCG TGAGGCCCAG
      61  AAAAAGTTCG CTACGTATAC CCAGGAGCAA GTTGACAAGA TCTTCAAACA GTGTGCCATT
```

121 GCGGCTGCCA AACTGCGTAT TAACTTGGCC CGTCTGGCGG TGGAAGAAAC CGGCATCGGT

181 CTGGTCGAGG ACAAGATTAT CAAGAATCAC TTTGCCGCGG AGTATATCTA CAACAAATAC

```
     241  AAAAACGAAA AGACTTGCGG CATCATTGAT CATGATGACA GCTTGGGTAT CACGAAAGTC
     301  GCAGAACCAA TCGGTATTGT CGCGGCAATT GTCCCGACCA CTAATCCGAC CAGCACCGCG
     361  ATCTTCAAGA GCTTGATCAG CCTGAAAACC CGCAACGCGA TTTTCTTCAG CCCGCACCCG
     421  CGTGCGAAGA AGAGCACCAT CGCAGCGGCG AAGCTGATTT TGGATGCCGC GGTTAAGGCC
     481  GGTGCACCGA AAAACATTAT TGGCTGGATT GACGAGCCGA GCATTGAGCT GTCTCAGGAC
     541  ATGATGAGCG AAGCGGACAT TATTCTGGCC ACCGGTGGTC CGTCTATGGT TAAGGCAGCG
     601  TACTCCAGCG GTAAGCCTGC TATCGGCGTT GGTGCGGGCA ATACGCCGGC TATCATCGAT
     661  GAAAGCGCGG ATATTGATAT GGCAGTTAGC TCCATCATTC TGTCCGCGAC CTACGACAAT
     721  GGTGTTATCT GTGCAAGCGA GCAAAGCATC CTGGTGATGA ATAGCATTTA CGAGAAGGTC
     781  AAAGAAGAGT TTGTTAAGCG TGGTTCCTAT ATTCTGAACC AGAACGAGAT TGCAAAAATC
     841  AAAGAAACGA TGTTTAAGAA TGGCGCGATT AATGCAGATA TTGTTGGTAA GTCCGCATAC
     901  ATTATCGCGA AGATGGCGGG TATCGAAGTG CCGCAAACCA CCAAAATCTT GATCGGCGAA
     961  GTGCAGAGCG TGGAGAAGAG CGAACTGTTC AGCCATGAGA AACTGAGCCC GGTTCTGGCT
    1021  ATGTACAAAG TGAAGGACTT TGACGAAGCC CTGAAGAAAG CTCAGCGTCT GATTGAGCTG
    1081  GGCGGCAGCG GTCACACGAG CAGCCTGTAT ATCGATAGCC AAAACAATAA GGACAAGGTG
    1141  AAAGAGTTCG GTCTGGCGAT GAAAACCTCT CGCACGTTTA TCAACATGCC GAGCAGCCAA
    1201  GGTGCGAGCG GCGATCTGTA TAACTTCGCA ATCGCGCCTT CTTTTACGCT GGGTTGCGGT
    1261  ACCTGGGGCG GTAATTCGGT GAGCCAGAAT GTCGAGCCGA AGCACCTGCT GAACATCGTT
    1321  TCGGTCGCGG AGCGCCGTGA GAAGAGACCG TGGTTCAAAG TCCCACAGAA AATTTACTTC
    1381  AAATACGGCT GCCTGCGTTT CGCGCTGAAA GAACTGAAAG ACATGAACAA AAAGCGTGCG
    1441  TTCATTGTTA CCGACAAAGA CCTGTTCAAA CTGGGTTACG TGAACAAAAT CACCAAAGTT
    1501  CTGGATGAAA TTGACATCAA GTACTCCATC TTCACTGATA TCAAATCCGA CCCAACGATT
    1561  GATAGCGTGA AAAAGGGCGC TAAAGAAATG CTGAACTTTG AACCGGACAC CATCATCAGC
    1621  ATCGGTGGTG GCTCTCCTAT GGATGCTGCG AAGGTCATGC ACCTGCTGTA CGAATACCCG
    1681  GAAGCGGAAA TCGAAAACCT GGCTATCAAC TTCATGGACA TCCGCAAACG TATCTGCAAC
    1741  TTCCCGAAGC TGGGCACTAA AGCTATTTCC GTTGCCATCC CGACTACCGC GGGCACTGGT
    1801  TCCGAAGCCA CGCCGTTCGC CGTGATCACC AACGATGAAA CCGGTATGAA ATACCCGCTG
    1861  ACCTCTTACG AACTGACCCC GAACATGGCA ATTATCGACA CCGAGCTGAT GCTGAACATG
    1921  CCGCGCAAGC TGACCGCTGC TACCGGCATC GACGCTCTGG TACATGCTAT TGAGGCGTAC
    1981  GTTTCCGTGA TGGCTACCGA TTACACCGAC GAACTGGCCC TGCGTGCGAT CAAAATGATT
    2041  TTCAAGTACC TGCCTCGCGC TTACAAAAAC GGCACGAATG ACATCGAGGC GCGTGAGAAA
    2101  ATGGCCCATG CAAGCAACAT CGCGGGCATG GCCTTCGCCA ACGCGTTCCT GGGCGTGTGC
    2161  CACTCTATGG CTCACAAACT GGGTGCTATG CACCACGTGC CGCACGGTAT CGCGTGTGCT
    2221  GTCCTGATCG AAGAAGTAAT TAAGTACAAC GCTACTGATT GCCCGACTAA ACAGACCGCC
    2281  TTCCCACAGT ACAAATCTCC TAACGCTAAA CGTAAGTACG CTGAGATCGC CGAATACCTG
    2341  AACCTGAAGG GTACGAGCGA CACTGAGAAA GTTACTGCGC TGATCGAAGC TATCTCTAAA
    2401  CTGAAAATTG ACCTGTCCAT CCCGCAGAAC ATCAGCGCCG CAGGCATCAA CAAAAAGGAC
    2461  TTTTACAACA CGCTGGACAA AATGAGCGAA CTGGCTTTTG ACGACCAGTG CACCACTGCA
    2521  AACCCGCGTT ACCCGCTGAT CTCTGAGCTG AAAGACATTT ATATCAAATC TTTTTAA
```

**AdhE2.91 (77020)**

```
       1  ATGAAAGTGA CGAACCAAAA AGAACTGAAA CAAAAACTGA ATGAACTGCG TGAGGCCCAG
      61  AAAAAGTTCG CTACGTGCAC CCAGGAGCAA GTTGACAAGT TCTTCAAACA GTGTGCCATT
```

121 GCGGCTGCCA AAGAGCGTAT TAACTTGGCC AAACTGGCGG TGGAAGAAAC CGGCATCGGT

181 CTGGTCGAGG ACAAGATTAT CAAGAATCAC TTTGCCGCGG AGTATATCTA CAACAAATAC

```
     241  AAAAACGAAA AGACTTGCGG CATCATTGAT CATGATGACA GCTTGGGTAT CACGAAAGTC
     301  GCAGAACCAA TCGGTATTGT CGCGGCAATT GTCCCGACCA CTAATCCGAC CAGCACCGCG
     361  ATCTTCAAGA GCTTGATCAG CCTGGCGACC CGCAACGCGA TTTTCTTCAG CCCGCACCCG
     421  CGTGCGAAGA AGAGCACCAT CGCAGCGGCG AAGCTGATTT TGGATGCCGC GGTTAAGGCC
     481  GGTGCACCGA AAAACATTAT TGGCTGGATT GACGAGCCGA GCATTGAGCT GTCTCAGGAC
     541  CTGATGAGCG AAGCGGACAT TATTCTGGCC ACCGGTGGTC CGTCTATGGT TAAGGCAGCG
     601  TACTCCAGCG GTAAGCCTGC TATCGGCGTT GGTGCGGGCA ATACGCCGGC TATCATCGAT
     661  GAAAGCGCGG ATATTGATAT GGCAGTTAGC TCCATCATTC TGTCCAAAAC CTACGACAAT
     721  GGTGTTATCT GTGCAAGCGA GCAAAGCATC CTGGTGATGA ATAGCATTTA CGAGAAGGTC
     781  AAAGAAGAGT TTGTTAAGCG TGGTTCCTAT ATTCTGAACC AGAACGAGAT TGCAAAAATC
     841  AAAGAAACGA TGTTTAAGAA TGGCGCGATT AATGCAGATA TTGTTGGTAA GTCCGCATAC
     901  ATTATCGCGA AGATGGCGGG TATCGAAGTG CCGCAAACCA CCAAAATCTT GATCGGCGAA
     961  GTGCAGGACG TGGAGAAGAG CGAACTGTTC AGCCATGAGA AACTGAGCCC GGTTCTGGCT
    1021  ATGTACAAAG TGAAGGACTT TGACGAAGCC CTGAAGCTGG CTCAGCGTCT GATTGAGCTG
    1081  GGCGGCAGCG GTCACACGAG CAGCCTGTAT ATCGATAGCC AAAACAATAA GGACAAGGTG
    1141  AAAGAGTTCG GTCTGGCGAT GAAAACCTCT CGCACGTTTA TCAACATGCC GAGCAGCCAA
    1201  GGTGCGAGCG GCGATCTGTA TAACTTCGCA ATCGCGCCTT CTTTTACGCT GGGTTGCGGT
    1261  ACCTGGGGCG GTAATTCGGT GAGCCAGAAT GTCGAGCCGA AGCACCTGCT GAACATCAAA
    1321  TCGGTCGCGG AGCGCCGTGA GAAGAGACCG TGGTTCAAAG TCCCACAGAA AATTTACTTC
    1381  AAATACGGCT GCCTGCGTTT CGCGCTGAAA GAACTGAAAG ACATGAACAA AAAGCGTGCG
    1441  TTCATTGTTA CCGACAAAGA CCTGTTCAAA CTGGGTTACG TGAACAAAAT CACCAAAGTT
    1501  CTGGATGAAA TTGACATCAA GTACTCCATC TTCACTGATA TCAAATCCGA CCCAACGATT
    1561  GATAGCGTGA AAAAGGGCGC TAAAGAAATG CTGAACTTTG AACCGGACAC CATCATCAGC
    1621  ATCGGTGGTG GCTCTCCTAT GGATGCTGCG AAGGTCATGC ACCTGCTGTA CGAATACCCG
    1681  GAAGCGGAAA TCGAAAACCT GGCTATCAAC TTCATGGACA TCCGCAAACG TATCTGCAAC
    1741  TTCCCGAAGC TGGGCACTAA AGCTATTTCC GTTGCCATCC CGACTACCGC GGGCACTGGT
    1801  TCCGAAGCCA CGCCGTTCGC CGTGATCACC AACGATGAAA CCGGTATGAA ATACCCGCTG
    1861  ACCTCTTACG AACTGACCCC GAACATGGCA ATTATCGACA CCGAGCTGAT GCTGAACATG
    1921  CCGCGCAAGC TGACCGCTGC TACCGGCATC GACGCTCTGG TACATGCTAT TGAGGCGTAC
    1981  GTTTCCGTGA TGGCTACCGA TTACACCGAC GAACTGGCCC TGCGTGCGAT CAAAATGATT
    2041  TTCAAGTACC TGCCTCGCGC TTACAAAAAC GGCACGAATG ACATCGAGGC GCGTGAGAAA
    2101  ATGGCCCATG CAAGCAACAT CGCGGGCATG GCCTTCGCCA ACGCGTTCCT GGGCGTGTGC
    2161  CACTCTATGG CTCACAAACT GGGTGCTATG CACCACGTGC CGCACGGTAT CGCGTGTGCT
    2221  GTCCTGATCG AAGAAGTAAT TAAGTACAAC GCTACTGATT GCCCGACTAA ACAGACCGCC
    2281  TTCCCACAGT ACAAATCTCC TAACGCTAAA CGTAAGTACG CTGAGATCGC CGAATACCTG
    2341  AACCTGAAGG GTACGAGCGA CACTGAGAAA GTTACTGCGC TGATCGAAGC TATCTCTAAA
    2401  CTGAAAATTG ACCTGTCCAT CCCGCAGAAC ATCAGCGCCG CAGGCATCAA CAAAAAGGAC
    2461  TTTTACAACA CGCTGGACAA AATGAGCGAA CTGGCTTTTG ACGACCAGTG CACCACTGCA
    2521  AACCCGCGTT ACCCGCTGAT CTCTGAGCTG AAAGACATTT ATATCAAATC TTTTTAA
```

**AdhE2.92 (77021)**

```
       1  ATGAAAGTGA CGAACCAAAA AGAACTGAAA CAAAAACTGA ATGAACTGCG TGAGGCCCAG
      61  AAAAAGTTCC TGACGTATAC CCAGGAGCAA GTTGACAAGA TCTTCAAACA GTGTGCCATT
```

121 GCGGCTGCCA AAGAGCGTAT TAACTTGGCC AAACTGGCGG TGGAAGAAAC CGGCATCGGT

181 CTGGTCGAGG ACAAGATTAT CAAGAATCAC TTTGCCGCGG AGTATATCTA CAACAAATAC

```
     241  AAAAACGAAA AGACTTGCGG CATCATTGAT CATGATGACA GCTTGGGTAT CACGAAAGTC
     301  GCAGAACCAA TCGGTATTGT CGCGGCAATT GTCCCGACCA CTAATCCGAC CAGCACCGCG
     361  ATCTTCAAGA GCTTGATCAG CCTGAAAACC CGCAACGCGA TTTTCTTCAG CCCGCACCCG
     421  CGTGCGAAGA AGAGCACCAT CGCAGCGGCG AAGCTGATTT TGGATGCCGC GGTTAAGGCC
     481  GGTGCACCGA AAAACATTAT TGGCTGGATT GACGAGCCGA GCATTGAGCT GTCTCAGGAC
     541  CTGATGAAAG AACCGGACAT TATTCTGGCC ACCGGTGGTC CGTCTATGGT TAAGGCAGCG
     601  TACTCCAGCG GTAAGCCTGC TATCGGCGTT GGTGCGGGCA ATACGCCGGC TATCATCGAT
     661  GAAAGCGCGG ATATTGATAT GGCAGTTAGC TCCATCATTC TGTCCAAAAC CTACGACAAT
     721  GGTGTTATCT GTGCAAGCGA GCAAAGCATC CTGGTGATGA ATAGCATTTA CGAGAAGGTC
     781  AAAGAAGAGT TTGTTAAGCG TGGTTCCTAT ATTCTGAACC AGAACGAGAT TGCAAAAATC
     841  AAAGAAACGA TGTTTAAGAA TGGCGCGATT AATGCAGATA TTGTTGGTAA GTCCGCATAC
     901  ATTATCGCGA AGATGGCGGG TATCGAAGTG CCGCAAACCA CCAAAATCTT GATCGGCGAA
     961  GTGCAGGACG TGGAGGCGAG CGAACTGTTC AGCCATGAGA AACTGAGCCC GGTTCTGGCT
    1021  ATGTACAAAG TGAAGGACTT TGACGAAGCC CTGAAGAAAG CTCAGCGTCT GATTGAGCTG
    1081  GGCGGCAGCG GTCACACGAG CAGCCTGTAT ATCGATAGCC AAAACAATAA GGACAAGGTG
    1141  AAAGAGTTCG GTCTGGCGAT GAAAACCTCT CGCACGTTTA TCAACATGCC GAGCAGCCAA
    1201  GGTGCGAGCG GCGATCTGTA TAACTTCGCA ATCGCGCCTT CTTTTACGCT GGGTTGCGGT
    1261  ACCTGGGGCG GTAATTCGGT GAGCCAGAAT GTCGAGCCGA AGCACCTGCT GAACATCAAA
    1321  TCGGTCGCGG AGCGCCGTGA GAAGAGACCG TGGTTCAAAG TCCCACAGAA AATTTACTTC
    1381  AAATACGGCT GCCTGCGTTT CGCGCTGAAA GAACTGAAAG ACATGAACAA AAAGCGTGCG
    1441  TTCATTGTTA CCGACAAAGA CCTGTTCAAA CTGGGTTACG TGAACAAAAT CACCAAAGTT
    1501  CTGGATGAAA TTGACATCAA GTACTCCATC TTCACTGATA TCAAATCCGA CCCAACGATT
    1561  GATAGCGTGA AAAAGGGCGC TAAAGAAATG CTGAACTTTG AACCGGACAC CATCATCAGC
    1621  ATCGGTGGTG GCTCTCCTAT GGATGCTGCG AAGGTCATGC ACCTGCTGTA CGAATACCCG
    1681  GAAGCGGAAA TCGAAAACCT GGCTATCAAC TTCATGGACA TCCGCAAACG TATCTGCAAC
    1741  TTCCCGAAGC TGGGCACTAA AGCTATTTCC GTTGCCATCC CGACTACCGC GGGCACTGGT
    1801  TCCGAAGCCA CGCCGTTCGC CGTGATCACC AACGATGAAA CCGGTATGAA ATACCCGCTG
    1861  ACCTCTTACG AACTGACCCC GAACATGGCA ATTATCGACA CCGAGCTGAT GCTGAACATG
    1921  CCGCGCAAGC TGACCGCTGC TACCGGCATC GACGCTCTGG TACATGCTAT TGAGGCGTAC
    1981  GTTTCCGTGA TGGCTACCGA TTACACCGAC GAACTGGCCC TGCGTGCGAT CAAAATGATT
    2041  TTCAAGTACC TGCCTCGCGC TTACAAAAAC GGCACGAATG ACATCGAGGC GCGTGAGAAA
    2101  ATGGCCCATG CAAGCAACAT CGCGGGCATG GCCTTCGCCA ACGCGTTCCT GGGCGTGTGC
    2161  CACTCTATGG CTCACAAACT GGGTGCTATG CACCACGTGC CGCACGGTAT CGCGTGTGCT
    2221  GTCCTGATCG AAGAAGTAAT TAAGTACAAC GCTACTGATT GCCCGACTAA ACAGACCGCC
    2281  TTCCCACAGT ACAAATCTCC TAACGCTAAA CGTAAGTACG CTGAGATCGC CGAATACCTG
    2341  AACCTGAAGG GTACGAGCGA CACTGAGAAA GTTACTGCGC TGATCGAAGC TATCTCTAAA
    2401  CTGAAAATTG ACCTGTCCAT CCCGCAGAAC ATCAGCGCCG CAGGCATCAA CAAAAAGGAC
    2461  TTTTACAACA CGCTGGACAA AATGAGCGAA CTGGCTTTTG ACGACCAGTG CACCACTGCA
    2521  AACCCGCGTT ACCCGCTGAT CTCTGAGCTG AAAGACATTT ATATCAAATC TTTTTAA
```

**AdhE2.93 (77022)**

```
       1  ATGAAAGTGA CGAACCAAAA AGAACTGAAA CAAAAACTGA ATGAACTGCG TGAGGCCCAG
      61  AAAAAGTTCC TGTGCTATAC CCAGGAGCAA GTTGACAAGA TCTTCAAACA GTGTGCCATT
```

121 GCGGCTGCCA AAGAGCGTAT TAACTTGGCC AAACTGGCGG TGGAAGAAAC CGGCATCGGT

181 CTGGTCGAGG ACAAGATTAT CAAGAATCAC TTTGCCGCGG AGTATATCTA CAACAAATAC

```
     241  AAAAACCACA AGACTTGCGG CATCATTGAT CATGATGACA GCTTGGGTAT CACGAAAGTC
     301  GCAGAACCAA TCGGTATTGT CGCGGCAATT GTCCCGACCA CTAATCCGAC CAGCACCGCG
     361  ATCTTCAAGA GCTTGATCAG CCTGAAAACC CGCAACGCGA TTTTCTTCAG CCCGCACCCG
     421  CGTGCGAAGA AGAGCACCAT CGCAGCGGCG AAGCTGATTT TGGATGCCGC GGTTAAGGCC
     481  GGTGCACCGA AAAACATTAT TGGCTGGATT GACGAGCCGA GCATTGAGCT GTCTCAGGAC
     541  CTGATGAGCG AAGCGGACAT TATTCTGGCC ACCGGTGGTC CGTCTATGGT TAAGGCAGCG
     601  TACTCCAGCG GTAAGCCTGC TATCGGCGTT GGTGCGGGCA ATACGCCGGC TATCATCGAT
     661  GAAAGCGCGG ATATTGATAT GGCAGTTAGC TCCATCATTC TGTCCAAAAC CTACGACAAT
     721  GGTGTTATCT GTGCAAGCGA GCAAAGCATC CTGGTGATGA ATAGCATTTA CGAGAAGGTC
     781  AAAGCGGAGT TTGTTAAGCG TGGTTCCTAT ATTCTGAACC AGAACGAGAT TGCAAAAATC
     841  AAAGAAACGA TGTTTAAGAA TGGCGCGATT AATCGTGATA TTGTTGGTAA GTCCGCATAC
     901  ATTATCGCGA AGATGGCGGG TATCGAAGTG CCGCAAACCA CCAAAATCTT GATCGGCGAA
     961  GTGCAGAGCG TGGAGAAGAG CGAACTGTTC AGCCATGAGA AACTGAGCCC GGTTCTGGCT
    1021  ATGTACAAAG TGAAGGACTT TGACGAAGCC CTGAAGAAAG CTCAGCGTCT GATTGAGCTG
    1081  GGCGGCAGCG GTCACACGAG CAGCCTGTAT ATCGATAGCC AAAACAATAA GGACAAGGTG
    1141  AAAGAGTTCG GTCTGGCGAT GAAAACCTCT CGCACGTTTA TCAACATGCC GAGCAGCCAA
    1201  GGTGCGAGCG GCGATCTGTA TAACTTCGCA ATCGCGCCTT CTTTTACGCT GGGTTGCGGT
    1261  ACCTGGGGCG GTAATTCGGT GAGCCAGAAT GTCGAGCCGA AGCACCTGCT GAACATCAAA
    1321  TCGGTCGCGG AGCGCCGTGA GAAGAGACCG TGGTTCAAAG TCCCACAGAA AATTTACTTC
    1381  AAATACGGCT GCCTGCGTTT CGCGCTGAAA GAACTGAAAG ACATGAACAA AAAGCGTGCG
    1441  TTCATTGTTA CCGACAAAGA CCTGTTCAAA CTGGGTTACG TGAACAAAAT CACCAAAGTT
    1501  CTGGATGAAA TTGACATCAA GTACTCCATC TTCACTGATA TCAAATCCGA CCCAACGATT
    1561  GATAGCGTGA AAAAGGGCGC TAAAGAAATG CTGAACTTTG AACCGGACAC CATCATCAGC
    1621  ATCGGTGGTG GCTCTCCTAT GGATGCTGCG AAGGTCATGC ACCTGCTGTA CGAATACCCG
    1681  GAAGCGGAAA TCGAAAACCT GGCTATCAAC TTCATGGACA TCCGCAAACG TATCTGCAAC
    1741  TTCCCGAAGC TGGGCACTAA AGCTATTTCC GTTGCCATCC CGACTACCGC GGGCACTGGT
    1801  TCCGAAGCCA CGCCGTTCGC CGTGATCACC AACGATGAAA CCGGTATGAA ATACCCGCTG
    1861  ACCTCTTACG AACTGACCCC GAACATGGCA ATTATCGACA CCGAGCTGAT GCTGAACATG
    1921  CCGCGCAAGC TGACCGCTGC TACCGGCATC GACGCTCTGG TACATGCTAT TGAGGCGTAC
    1981  GTTTCCGTGA TGGCTACCGA TTACACCGAC GAACTGGCCC TGCGTGCGAT CAAAATGATT
    2041  TTCAAGTACC TGCCTCGCGC TTACAAAAAC GGCACGAATG ACATCGAGGC GCGTGAGAAA
    2101  ATGGCCCATG CAAGCAACAT CGCGGGCATG GCCTTCGCCA ACGCGTTCCT GGGCGTGTGC
    2161  CACTCTATGG CTCACAAACT GGGTGCTATG CACCACGTGC CGCACGGTAT CGCGTGTGCT
    2221  GTCCTGATCG AAGAAGTAAT TAAGTACAAC GCTACTGATT GCCCGACTAA ACAGACCGCC
    2281  TTCCCACAGT ACAAATCTCC TAACGCTAAA CGTAAGTACG CTGAGATCGC CGAATACCTG
    2341  AACCTGAAGG GTACGAGCGA CACTGAGAAA GTTACTGCGC TGATCGAAGC TATCTCTAAA
    2401  CTGAAAATTG ACCTGTCCAT CCCGCAGAAC ATCAGCGCCG CAGGCATCAA CAAAAAGGAC
    2461  TTTTACAACA CGCTGGACAA AATGAGCGAA CTGGCTTTTG ACGACCAGTG CACCACTGCA
    2521  AACCCGCGTT ACCCGCTGAT CTCTGAGCTG AAAGACATTT ATATCAAATC TTTTTAA
```

**AdhE2.94 (77023)**

```
       1  ATGAAAGTGA CGAACCAAAA AGAACTGAAA CAAAAACTGA ATGAACTGCG TGAGGCCCAG
      61  AAAAAGTTCG CTACGTATAC CCAGGAGCAA GTTGACAAGA TCTTCAAACA GTGTGCCATT
```

121 GCGGCTGCCA AATACCGTAT TAACTTGGCC AAACTGGCGG TGGAAGAAAC CGGCATCGGT

181 CTGGTCGAGG ACAAGATTAT CAAGAATCAC TTTGCCGCGG AGTATATCTA CAACAAATAC

```
     241  AAAAACGAAA AGACTTGCGG CATCATTGAT CATGATGACA GCTTGGGTAT CACGAAAGTC
     301  GCAGAACCAA TCGGTATTGT CGCGGCAATT GTCCCGACCA CTAATCCGAC CAGCACCGCG
     361  ATCTTCAAGA GCTTGATCAG CCTGAAAACC CGCAACGCGA TTTTCTTCAG CCCGCACCCG
     421  CGTGCGAAGA AGAGCACCAT CGCAGCGGCG AAGCTGATTT TGGATGCCGC GGTTAAGGCC
     481  GGTGCACCGA AAAACATTAT TGGCTGGATT GACGAGCCGA GCATTGAGCT GTCTCAGGAC
     541  CTGATGAGCG AACCGGACAT TATTCTGGCC ACCGGTGGTC CGTCTATGGT TAAGGCAGCG
     601  TACTCCAGCG GTAAGCCTGC TATCGGCGTT GGTGCGGGCA ATACGCCGGC TATCATCGAT
     661  GAAAGCGCGG ATATTGATAT GGCAGTTAGC TCCATCATTC TGTCCAAAAC CTACGACAAT
     721  GGTGTTATCT GTGCAAGCGA GCAAAGCATC CTGGTGATGA ATAGCATTTA CGAGAAGGTC
     781  AAAGAAGAGT TTGTTAAGCG TGGTTCCTAT ATTCTGAACC AGAACGAGAT TGCAAAAATC
     841  AAAGAAACGA TGTTTAAGAA TGGCGCGATT AATGCAGATA TTGTTGGTAA GTCCGCATAC
     901  ATTATCGCGA AGATGGCGGG TATCGAAGTG CCGCAAACCA CCAAAATCTT GATCGGCGAA
     961  GTGCAGAGCG TGGAGAAGAG CGAACTGTTC AGCGTTGAGC TGCTGAGCCC GGTTCTGGCT
    1021  ATGTACAAAG TGAAGGACTT TGACGAAGCC CTGAAGAAAG CTGAGCGTCT GATTGAGCTG
    1081  GGCGGCAGCG GTCACACGAG CAGCCTGTAT ATCGATAGCC AAAACAATAA GGACAAGGTG
    1141  AAAGAGTTCG GTCTGGCGAT GAAAACCTCT CGCACGTTTA TCAACATGCC GAGCAGCCAA
    1201  GGTGCGAGCG GCGATCTGTA TAACTTCGCA ATCGCGCCTT CTTTTACGCT GGGTTGCGGT
    1261  ACCTGGGGCG GTAATTCGGT GAGCCAGAAT GTCGAGCCGA AGCACCTGCT GAACATCAAA
    1321  TCGGTCGCGG AGCGCCGTGA GAAGAGACCG TGGTTCAAAG TCCCACAGAA AATTTACTTC
    1381  AAATACGGCT GCCTGCGTTT CGCGCTGAAA GAACTGAAAG ACATGAACAA AAAGCGTGCG
    1441  TTCATTGTTA CCGACAAAGA CCTGTTCAAA CTGGGTTACG TGAACAAAAT CACCAAAGTT
    1501  CTGGATGAAA TTGACATCAA GTACTCCATC TTCACTGATA TCAAATCCGA CCCAACGATT
    1561  GATAGCGTGA AAAAGGGCGC TAAAGAAATG CTGAACTTTG AACCGGACAC CATCATCAGC
    1621  ATCGGTGGTG GCTCTCCTAT GGATGCTGCG AAGGTCATGC ACCTGCTGTA CGAATACCCG
    1681  GAAGCGGAAA TCGAAAACCT GGCTATCAAC TTCATGGACA TCCGCAAACG TATCTGCAAC
    1741  TTCCCGAAGC TGGGCACTAA AGCTATTTCC GTTGCCATCC CGACTACCGC GGGCACTGGT
    1801  TCCGAAGCCA CGCCGTTCGC CGTGATCACC AACGATGAAA CCGGTATGAA ATACCCGCTG
    1861  ACCTCTTACG AACTGACCCC GAACATGGCA ATTATCGACA CCGAGCTGAT GCTGAACATG
    1921  CCGCGCAAGC TGACCGCTGC TACCGGCATC GACGCTCTGG TACATGCTAT TGAGGCGTAC
    1981  GTTTCCGTGA TGGCTACCGA TTACACCGAC GAACTGGCCC TGCGTGCGAT CAAAATGATT
    2041  TTCAAGTACC TGCCTCGCGC TTACAAAAAC GGCACGAATG ACATCGAGGC GCGTGAGAAA
    2101  ATGGCCCATG CAAGCAACAT CGCGGGCATG GCCTTCGCCA ACGCGTTCCT GGGCGTGTGC
    2161  CACTCTATGG CTCACAAACT GGGTGCTATG CACCACGTGC CGCACGGTAT CGCGTGTGCT
    2221  GTCCTGATCG AAGAAGTAAT TAAGTACAAC GCTACTGATT GCCCGACTAA ACAGACCGCC
    2281  TTCCCACAGT ACAAATCTCC TAACGCTAAA CGTAAGTACG CTGAGATCGC CGAATACCTG
    2341  AACCTGAAGG GTACGAGCGA CACTGAGAAA GTTACTGCGC TGATCGAAGC TATCTCTAAA
    2401  CTGAAAATTG ACCTGTCCAT CCCGCAGAAC ATCAGCGCCG CAGGCATCAA CAAAAAGGAC
    2461  TTTTACAACA CGCTGGACAA AATGAGCGAA CTGGCTTTTG ACGACCAGTG CACCACTGCA
    2521  AACCCGCGTT ACCCGCTGAT CTCTGAGCTG AAAGACATTT ATATCAAATC TTTTTAA
```

**AdhE2.95 (77024)**

```
       1  ATGAAAGTGA CGAACCAAAA AGAACTGAAA CAAAAACTGA ATGAACTGCG TGAGGCCCAG
      61  AAAAAGTTCG CTACGTATAC CCAGGAGCAA GTTGACAAGA TCTTCAAACA GTGTGCCATT
```

121 GCGGCTGCCA AATACCGTAT TAACTTGGCC AAACTGGCGG TGGAAGAAAC CGGCATCGGT

181 CTGGTCGAGG ACAAGATTAT CAAGAATCAC TTTGCCGCGG AGTATATCTA CAACAAATAC

```
     241  AAAAACGAAA AGACTTGCGG CATCATTGAT CATGATGACA GCTTGGGTAT CACGAAAGTC
     301  GCAGAACCAA TCGGTATTGT CGCGGCAATT GTCCCGACCA CTAATCCGAC CAGCACCGCG
     361  ATCTTCAAGA GCTTGATCAG CCTGAAAACC CGCAACGCGA TTTTCTTCAG CCCGCACCCG
     421  CGTGCGAAGA AGAGCACCAT CGCAGCGGCG AAGCTGATTT TGGATGCCGC GGTTAAGGCC
     481  GGTGCACCGA AAAACATTAT TGGCTGGATT GACGAGCCGA GCATTGAGCT GTCTCAGGAC
     541  CTGATGAGCG AAGCGGACAT TATTCTGGCC ACCGGTGGTC CGTCTATGGT TAAGGCAGCG
     601  TACTCCAGCG GTAAGCCTGC TATCGGCGTT GGTGCGGGCA ATACGCCGGC TATCATCGAT
     661  GAAAGCGCGG ATATTGATAT GGCAGTTAGC TCCATCATTC TGTCCAAAAC CTACGACAAT
     721  GGTGTTATCT GTGCAAGCGA GCAAAGCATC CTGGTGATGA ATAGCATTTA CGAGAAGGTC
     781  AAAGAAGAGT TTGTTAAGCG TGGTTCCTAT ATTCTGAACC AGGAGGAGAT TGCAAAAATC
     841  AAAGAAACGA TGTTTAAGAA TGGCGCGATT AATCGTGATA TTGTTGGTAA GTCCGCATAC
     901  ATTATCGCGA AGATGGCGGG TATCGAAGTG CCGCAAACCA CCAAAATCTT GATCGGCGAA
     961  GTGCAGAGCG TGGAGAAGAG CGAACTGTTC AGCCATGAGA AACTGAGCCC GGTTCTGGCT
    1021  ATGTACAAAG TGAAGGACTT TGACGAAGCC CTGGACAAAG CTCAGCGTCT GATTGAGCTG
    1081  GGCGGCAGCG GTCACACGAG CAGCCTGTAT ATCGATAGCC AAAACAATAA GGACAAGGTG
    1141  AAAGAGTTCG GTCTGGCGAT GAAAACCTCT CGCTTCTTTA TCAACATGCC GAGCAGCCAA
    1201  GGTGCGAGCG GCGATCTGTA TAACTTCGCA ATCGCGCCTT CTTTTACGCT GGGTTGCGGT
    1261  ACCTGGGGCG GTAATTCGGT GAGCCAGAAT GTCGAGCCGA AGCACCTGCT GAACATCAAA
    1321  TCGGTCGCGG AGCGCCGTGA GAAGAGACCG TGGTTCAAAG TCCCACAGAA AATTTACTTC
    1381  AAATACGGCT GCCTGCGTTT CGCGCTGAAA GAACTGAAAG ACATGAACAA AAAGCGTGCG
    1441  TTCATTGTTA CCGACAAAGA CCTGTTCAAA CTGGGTTACG TGAACAAAAT CACCAAAGTT
    1501  CTGGATGAAA TTGACATCAA GTACTCCATC TTCACTGATA TCAAATCCGA CCCAACGATT
    1561  GATAGCGTGA AAAAGGGCGC TAAAGAAATG CTGAACTTTG AACCGGACAC CATCATCAGC
    1621  ATCGGTGGTG GCTCTCCTAT GGATGCTGCG AAGGTCATGC ACCTGCTGTA CGAATACCCG
    1681  GAAGCGGAAA TCGAAAACCT GGCTATCAAC TTCATGGACA TCCGCAAACG TATCTGCAAC
    1741  TTCCCGAAGC TGGGCACTAA AGCTATTTCC GTTGCCATCC CGACTACCGC GGGCACTGGT
    1801  TCCGAAGCCA CGCCGTTCGC CGTGATCACC AACGATGAAA CCGGTATGAA ATACCCGCTG
    1861  ACCTCTTACG AACTGACCCC GAACATGGCA ATTATCGACA CCGAGCTGAT GCTGAACATG
    1921  CCGCGCAAGC TGACCGCTGC TACCGGCATC GACGCTCTGG TACATGCTAT TGAGGCGTAC
    1981  GTTTCCGTGA TGGCTACCGA TTACACCGAC GAACTGGCCC TGCGTGCGAT CAAAATGATT
    2041  TTCAAGTACC TGCCTCGCGC TTACAAAAAC GGCACGAATG ACATCGAGGC GCGTGAGAAA
    2101  ATGGCCCATG CAAGCAACAT CGCGGGCATG GCCTTCGCCA ACGCGTTCCT GGGCGTGTGC
    2161  CACTCTATGG CTCACAAACT GGGTGCTATG CACCACGTGC CGCACGGTAT CGCGTGTGCT
    2221  GTCCTGATCG AAGAAGTAAT TAAGTACAAC GCTACTGATT GCCCGACTAA ACAGACCGCC
    2281  TTCCCACAGT ACAAATCTCC TAACGCTAAA CGTAAGTACG CTGAGATCGC CGAATACCTG
    2341  AACCTGAAGG GTACGAGCGA CACTGAGAAA GTTACTGCGC TGATCGAAGC TATCTCTAAA
    2401  CTGAAAATTG ACCTGTCCAT CCCGCAGAAC ATCAGCGCCG CAGGCATCAA CAAAAAGGAC
    2461  TTTTACAACA CGCTGGACAA AATGAGCGAA CTGGCTTTTG ACGACCAGTG CACCACTGCA
    2521  AACCCGCGTT ACCCGCTGAT CTCTGAGCTG AAAGACATTT ATATCAAATC TTTTTAA
```

**AdhE2.96 (77025)**

```
       1  ATGAAAGTGA CGAACCAAAA AGAACTGAAA CAAAAACTGA ATGAACTGCG TGAGGCCCAG
      61  AAAAAGTTCG CTACGTATAC CCAGGAGCAA GTTGACAAGA TCTTCAAACA GTGTGCCATT
```

121 GCGGCTGCCA AAGAGCGTAT TAACTTGGCC GAGCTGGCGG TGGAAGAAAC CGGCATCGGT

181 CTGGTCGAGG ACAAGATTAT CAAGAATCAC TTTGCCGCGG AGTATATCTA CAACAAATAC

```
     241  AAAAACGGTA AGACTTGCGG CATCATTGAT CATGATGACA GCTTGGGTAT CACGAAAGTC
     301  GCAGAACCAA TCGGTATTGT CGCGGCAATT GTCCCGACCA CTAATCCGAC CAGCACCGCG
     361  ATCTTCAAGA GCTTGATCAG CCTGAAAACC CGCAACGCGA TTTTCTTCAG CCCGCACCCG
     421  CGTGCGAAGA AGAGCACCAT CGCAGCGGCG AAGCTGATTT TGGATGCCGC GGTTAAGGCC
     481  GGTGCACCGA AAAACATTAT TGGCTGGATT GACGAGCCGA GCATTGAGCT GTCTCAGGAC
     541  CTGATGAGCG AAGCGGACAT TATTCTGGCC ACCGGTGGTC CGTCTATGGT TAAGGCAGCG
     601  TACTCCAGCG GTAAGAAAGC TATCGGCGCG GGTGCGGGCA ATACGCCGGC TATCATCGAT
     661  GAAAGCGCGG ATATTGATAT GGCAGTTAGC TCCATCATTC TGTCCGCGAC CTACGACAAT
     721  GGTGTTATCT GTGCAAGCGA GCAAAGCATC CTGGTGATGA ATAGCATTTA CGAGAAGGTC
     781  AAAGAAGAGT TTGTTAAGCG TGGTTCCTAT ATTCTGAACC AGAACGAGAT TGCAAAAATC
     841  AAAGAAACGA TGTTTAAGAA TGGCGCGATT AATGCAGATA TTGTTGGTAA GTCCGCATAC
     901  ATTATCGCGA AGATGGCGGG TATCGAAGTG CCGCAAACCA CCAAAATCTT GATCGGCGAA
     961  GTGCAGAGCG TGGAGAAGAG CGAACTGTTC AGCCATGAGA AACTGAGCCC GGTTCTGGCT
    1021  ATGTACAAAG TGAAGGACTT TGACGAAGCC CTGAAGAAAG CTCAGCGTCT GATTGAGCTG
    1081  GGCGGCAGCG GTCACACGAG CAGCCTGTAT ATCGATAGCC AAAACAATAA GGACAAGGTG
    1141  AAAGAGTTCG GTCTGGCGAT GAAAACCTCT CGCACGTTTA TCAACATGCC GAGCAGCCAA
    1201  GGTGCGAGCG GCGATCTGTA TAACTTCGCA ATCGCGCCTT CTTTTACGCT GGGTTGCGGT
    1261  ACCTGGGGCG GTAATTCGGT GAGCCAGAAT GTCGAGCCGA AGCACCTGCT GAACATCAAA
    1321  TCGGTCGCGG AGCGCCGTGA GAAGAGACCG TGGTTCAAAG TCCCACAGAA AATTTACTTC
    1381  AAATACGGCT GCCTGCGTTT CGCGCTGAAA GAACTGAAAG ACATGAACAA AAAGCGTGCG
    1441  TTCATTGTTA CCGACAAAGA CCTGTTCAAA CTGGGTTACG TGAACAAAAT CACCAAAGTT
    1501  CTGGATGAAA TTGACATCAA GTACTCCATC TTCACTGATA TCAAATCCGA CCCAACGATT
    1561  GATAGCGTGA AAAAGGGCGC TAAAGAAATG CTGAACTTTG AACCGGACAC CATCATCAGC
    1621  ATCGGTGGTG GCTCTCCTAT GGATGCTGCG AAGGTCATGC ACCTGCTGTA CGAATACCCG
    1681  GAAGCGGAAA TCGAAAACCT GGCTATCAAC TTCATGGACA TCCGCAAACG TATCTGCAAC
    1741  TTCCCGAAGC TGGGCACTAA AGCTATTTCC GTTGCCATCC CGACTACCGC GGGCACTGGT
    1801  TCCGAAGCCA CGCCGTTCGC CGTGATCACC AACGATGAAA CCGGTATGAA ATACCCGCTG
    1861  ACCTCTTACG AACTGACCCC GAACATGGCA ATTATCGACA CCGAGCTGAT GCTGAACATG
    1921  CCGCGCAAGC TGACCGCTGC TACCGGCATC GACGCTCTGG TACATGCTAT TGAGGCGTAC
    1981  GTTTCCGTGA TGGCTACCGA TTACACCGAC GAACTGGCCC TGCGTGCGAT CAAAATGATT
    2041  TTCAAGTACC TGCCTCGCGC TTACAAAAAC GGCACGAATG ACATCGAGGC GCGTGAGAAA
    2101  ATGGCCCATG CAAGCAACAT CGCGGGCATG GCCTTCGCCA ACGCGTTCCT GGGCGTGTGC
    2161  CACTCTATGG CTCACAAACT GGGTGCTATG CACCACGTGC CGCACGGTAT CGCGTGTGCT
    2221  GTCCTGATCG AAGAAGTAAT TAAGTACAAC GCTACTGATT GCCCGACTAA ACAGACCGCC
    2281  TTCCCACAGT ACAAATCTCC TAACGCTAAA CGTAAGTACG CTGAGATCGC CGAATACCTG
    2341  AACCTGAAGG GTACGAGCGA CACTGAGAAA GTTACTGCGC TGATCGAAGC TATCTCTAAA
    2401  CTGAAAATTG ACCTGTCCAT CCCGCAGAAC ATCAGCGCCG CAGGCATCAA CAAAAAGGAC
    2461  TTTTACAACA CGCTGGACAA AATGAGCGAA CTGGCTTTTG ACGACCAGTG CACCACTGCA
    2521  AACCCGCGTT ACCCGCTGAT CTCTGAGCTG AAAGACATTT ATATCAAATC TTTTTAA
```

*D2. AdhE2 homologs in the pCDF3-TdTer.AdhE29-45 plasmids*

| **Name** | **Accession No**. |
| --- | --- |
| AdhE30 | 317057809 |
| AdhE31 | 221194444 |
| AdhE32 | 302875249 |
| AdhE33 | 149276422 |
| AdhE34 | 15004865 |
| AdhE35 | 260891427 |
| AdhE36 | 126697906 |
| AdhE37 | 188590416 |
| AdhE38 | 300854834 |
| AdhE39 | 110802422 |
| AdhE40 | 210633271 |
| AdhE41 | 220930559 |
| AdhE42 | 168184834 |
| AdhE43 | 317133832 |
| AdhE44 | 307687897 |
| AdhE45 | 225571592 |

*D3. Monofunctional ALDH variants in the pCDF3-TdTer.ALDH1-15 and ALDH46 plasmids*

**ALDH1**

atgaataaag acaccctgat tccaactacc aaagatctga agctgaaaac taatgtcgaa

aacatcaatt tgaagaacta caaagataac agctcgtgtt ttggcgtgtt cgaaaacgtt

gagaacgcga tcaattccgc cgttcacgca cagaagattc tgagcctgca ctacaccaaa

gagcagcgtg agaagatcat tacggaaatc cgcaaagcgg cgctggagaa taaagaggtg

ctggctacca tgattctgga agaaacccac atgggtcgtt atgaggacaa aatcctgaag

cacgagctgg tcgctaagta cacccctggc accgaggacc tgacgaccac ggcatggagc

ggtgataacg gtttgacggt cgtcgaaatg agcccgtatg gcgtcattgg tgcaattacc

cctagcacca atccgaccga aactgtgatc tgcaactcta tcggtatgat tgctgcgggc

aatgccgtcg ttttcaatgg ccatccgggt gcgaagaagt gcgttgcgtt cgctattgaa

atgatcaaca aggcgatcat ctcctgtggt ggtccggaga acttggtgac cacgatcaaa

aatccgacga tggagagcct ggatgcgatc attaaacatc cgctgattaa actgttgtgc

ggtacgggcg gtccgggtat ggttaaaacg ctgctgaata gcggcaaaaa ggcaatcggt

gccggtgcgg gcaacccgcc agttatcgta gacgacacgg cggacattga aaaggccggt

aagagcatta ttgagggttg ttcttttgac aacaatctgc cgtgtattgc ggagaaagaa

gtgtttgttt tcgagaatgt ggcggatgac ctgattagca acatgctgaa aaacaatgca

gttatcatca acgaggatca agtctccaag ctgatcgatc tggtgttgca gaaaaacaac

gaaacccaag agtacttcat taacaagaag tgggttggta aggatgcaaa gctgtttagc

gacgagattg acgtggaaag cccgagcaat atcaaatgca tcgtgtgcga ggtcaatgca

aatcacccgt tcgttatgac cgaactgatg atgccgatcc tgccgattgt tcgcgtgaaa

gatatcgatg aggcggtcaa atacactaag atcgcggagc agaatcgtaa acatagcgcg

tacatctata gcaagaacat cgacaacttg aatcgtttcg aacgtgagat cgacaccacg

atttttgtga aaaacgcaaa gagcttcgcc ggtgtgggct atgaagccga aggctttacc

acctttacca ttgcgggcag cacgggcgag ggtattacct ctgcacgtaa tttcacccgt

caacgccgct gcgttctggc cggttaa

**ALDH2**

atgaatgaca tcgaaatcgc ccaagccgta agcactattc tgagcaagtt cactaaagca

acgcctgacg aggctccggc gacctcggaa gccgcacgtg tcgatggtct ggatgagatt

gtggcaaaag ccttggccca gcacagcagc gtgcgcgatg cttctgcgat tagccaagtt

gcgaaagttg ccaacgcttc taccggtgcg ttcgatacga tggacgaggc gatctccgca

gcggttttgg cacaggtcca atatcgtcat tgttctatgc aggatcgcgc aagctttatc

aatggtattc gcgacgtgtt cctgcaagag gacgtgctgt gtgccctgag ccgcatggcg

gtggaagaaa ccggtatggg taactacgaa gataagctga tcaaaaatcg cgtggccgca

ctgaaaacgc cgggtattga ggatctgacg accagcgcgg ttagcggcga cggtggcctg

acgctgattg aatacagcgc gttcggcgtc attggcagca tcaccccaac cacgaacccg

acggaaacga tcatcaacaa ttctatcggc atgctggcag cgggcaatac cgtcgtcttt

agcccgcacc cgcgttcccg caaggtttcc ctgtacgcgg tggaattgat caacaataaa

ctggcgcagc tgggtgcacc ggccaacatg gtagtgaccg tgaccaagcc gagcatcgac

aacaccaatg ttctgattaa tgatccgcgt attaacatgc tggtagcaac cggcggtccg

gcgattgtta agaccgttat gagcagcggt aaaaaggcga tcggtgcggg tgctggtaac

ccgcctgcgg ttgtggatga aacggcggac attgagaagg ctgcgcgtga tatcattaaa

ggttgcagct tcgacaacaa tctgccatgt gtcgcagaaa aagaggtcat cgttgtcaat

caggttgctg attacctgat ccattgcatg aagaaaagcg gtgcctatct gctgtgcgac

aagaaactga gccagcaact gcagagcctg gtcttgaacg agaagggtac tggcccgaat

accgcgttcg tgggcaaaga cgcacgttac atcctgcagc aactgggcat ccaggttggc

gacgacatta aggtcatttt gatcgaagcg gagaaaaccc acccgtttgt tgttcacgag

ctgatgatgc cggtcttgcc ggttgtgcgt gtggacaatg tggatgaggc gattgagctg

gcagtgaagg tggagcatgg taaccgccac acggcggtca tgcactccac caacgttgag

aagttgacca agatggcgcg tctgattcaa acgaccatct ttgtcaaaaa tggtccgtcg

tatgcgggcc tgggcgttgg tggtgagggt catgcgacct ttaccattgc tggcccgacg

ggtgaaggtc tgaccagcgc ccgtagcttc gcacgtcgtc gtcgttgcgt gatggtcgag

gcgctgaaca ttcgctaa

**ALDH3**

atgattaagg acactctcgt aagcatcacc aaggatctga aattgaaaac gaatgtagag

aacgccaatc tgaagaacta caaggacgat tcgagctgct tcggtgtttt tgaaaatgtg

gagaatgcta ttagcaatgc ggtgcatgcg cagaaaatcc tgtccctgca ttacaccaaa

gagcaacgcg aaaagatcat cactgagatt cgtaaggccg cactggagaa taaagagatc

ctggcgacca tgattctgga agaaacccac atgggtcgtt acgaggataa gattctgaag

cacgaattgg ttgccaagta cactccgggt accgaagatc tgaccaccac ggcgtggagc

ggtgataacg gtctgaccgt tgtcgagatg agcccgtatg gtgttatcgg tgccattacc

ccttctacga atccgacgga aaccgtgatc tgcaacagca tcggcatgat tgcggcaggc

aataccgtgg tgttcaatgg ccatccgggt gccaagaagt gtgtcgcgtt tgcagttgag

atgattaaca aagcaatcat ttcttgtggt ggcccggaaa acctggttac caccatcaag

aacccgacga tggacagctt ggacgcaatt atcaaacacc cgtccattaa actgctgtgc

ggtacgggtg gcccaggcat ggtcaagacg ttgctgaaca gcggtaaaaa ggcgattggt

gcgggtgccg gcaatccgcc ggtcattgtg gacgacacgg ctgacatcga gaaagcgggc

aaaagcatca ttgaaggctg cagcttcgac aacaatctgc cgtgcatcgc ggagaaagag

gtttttgttt ttgagaacgt cgcagacgat ctgatttcga acatgctgaa gaataatgcg

gtcattatca atgaggacca ggttagcaaa ttgatcgatc tggtcctgca gaagaacaac

gagactcaag aatatagcat taacaaaaag tgggtgggta aagatgcgaa gctgtttctg

gacgagattg atgtggagtc tccgagcagc gttaagtgta tcatctgcga agtgtccgct

cgccacccgt tcgtcatgac cgagctgatg atgccgatcc tgccaattgt gcgtgtgaaa

gatattgacg aagcaatcga gtacgctaaa atcgcagaac aaaatcgcaa acacagcgca

tatatctata gcaaaaacat cgacaacctg aaccgtttcg aacgcgaaat tgataccacc

attttcgtca agaacgctaa aagctttgcg ggtgttggtt acgaggccga aggctttacc

acgttcacca ttgcgggcag cacgggcgag ggtatcacgt ccgcgcgtaa tttcacccgt

cagcgtcgtt gtgttctggc gggttaa

**ALDH4**

atgtcatttg atatcaacaa tgcacaaggc gtatttgaaa cggtagaagc agcaattgaa

gccacccaca aagcccaggt ggagttctat gcgaactcca ctaaagaggg ccgtgaggcg

atcctgaccg ctatccgtgg cgccgtgttg gcgaaagcgg aagatttcgc caaaatggtt

cgcgaagaaa ccaagctggg ccgtgtcgag gataagatcg cgaaacatca actgaccgca

gccaagaccc cgggtaccga ggtcctggaa acgaaggttt ggagcggtga caacggtatc

agcctggaag agcgtgcgcc gtacggtgtc atcggcgctg tcaccccggt tacgaatccg

acggaaacga tcgtcaacaa cgcaattagc atgctggcga gcggcaacgc ggtgacgttc

aatgtgcatc catcctcgaa agttgtgagc gcagttatga tcgacatgat taacaaaacg

attgttgctg cgggtggtcc ggcgaacctg gtgactatgg ttaaagaacc aacgctggaa

acgctgaacg aaatcgcgaa aagcccgctg gtgaatatgt tggtcggtac gggcggtccg

ggcctggtga aggcgattct gcaatctggc aagaaaggtg tcggtgcggg tgcgggtaat

ccgccggtga ttgtcgatgc atctgctaat ctggacctgg ctgcagcggg tgtatacggc

ggtgccagct tcgacaataa cctgttgtgt attggcgaga aagaggtgtt cgttgaggat

agcgtcgcgg acgagtttct ggctaagctg gaagcgaccg gtgcctatgt tctgagcgca

gaagaggcgg agaagttgac cgctcagatc ctgacgatgg acgagatcga cggtgcgaaa

ccgtgtaccg cacaggaaat tgcgcgtgtg tggcacccgg tcaagcagca cgttggtcaa

gatgcgggtg agatcctgaa gtccatcggt gtcgagagcg aaacccgtct ggcggtgatg

gttgtggaga atgatcatcc tctggttcac gtcgagcaga tgatgccggt gctgccggtt

gtgcgttgcg cgaatattga cgaggcgatc gagcgcgcag ttgcggccga gcgtggcaac

aagcacagcg cgtgcatcta cagcggcaac attgagaatg ttaccaagtt cggtcgtgca

attaacacca ccatctttgc ccacaacggt ccgaccttga gcggtgtcgg ctacaatgca

gaaggtacca gcacctttac cattgcaggc ccgactggtg agggtattac caatgcgtat

agcttcaccc gcgcacgtcg ctttgccatt gcccagggcg gtctgcgcat tgtttaa

**ALDH5**

atgtccgtaa acgagaagat ggtccaagat attgtacaag aagtcgtagc taaaatgcaa

attagctccg acgtcagcgg caagaagggc gtttttagcg atatgaatga agcaatcgag

gcgagcaaaa aggcacagaa aatcgtggct aaaatgagca tggaccaacg cgaagccatt

atcagcaaga tccgtgagaa gattaaagag aatgcggaaa ttctggcgcg tatgggtgtt

gaagaaaccg gcatgggtaa tgttggccac aaaattctga agcatcagct ggttgcggaa

aagaccccgg gtaccgagga catcacgacg acggcttggt ctggtgatcg tggtttgact

ttgatcgaaa tgggcccgtt cggcgttatc ggcgcgatca ccccgtgcac taacccgtct

gaaaccgtgc tgtgtaatac gatcggtatg ctggcgggtg gtaacaccgt tgtctttaac

ccacatccag ccgccatcaa gaccagcatc tatgcggtga atctgctgaa cgaggcatcc

gtcgaggttg gtggtccgga gaatattgcg gtgaccgtcg agcacccgac gatggaaacc

tcggatatca tgatgaagca caaggacatc catctgattg cggctacggg cggtccgggc

gttgtgaccg ccgtcctgag cagcggtaaa cgcggtattg gtgcgggtgc tggcaacccg

cctgcgttgg tcgacgaaac ggccgacatt cgcaaggccg cagaggatat tgtgaacggt

tgtaccttcg acaataatct gccgtgcatt gcggagaaag aaattgtggc agtggattcg

atcgcagatg agctgttgca ctacatggtg agcgagcagg gctgttacat gatcagcaaa

gaagagcagg acgcgctgac cgaagttgtt ctgaaaggcg gtcgtctgaa tcgcaaatgc

gtgggccgtg acgcgaaaac gttgctgggt atgattggta tcacggttcc ggacaatatt

cgttgcatca cgtttgaggg tccgaaagag catccgctga tcgcggaaga actgatgatg

ccgattctgg gcgtggttcg tgcgaaagat tttgatgatg cagtggagca ggcagtgtgg

ctggagcacg gtaaccgcca cagcgcgcac attcatagca agaacgttga caacatcacc

aaatacgcaa aagccattga caccgcgatt ctggtcaaga acggtccgag ctatgcagca

ctgggcttcg gtggtgaggg ctattgcacc ttcaccatcg ccagccgtac cggcgagggt

ctgactagcg cgagcacgtt caccaagcgt cgccgttgtg tcatgaccga ttctctgtgc

attcgttaa

**ALDH6**

atgaaagagg gtgtaattcg cttggacatg gacattaagg taattgaaca gttggtagaa

caagcgctga aagagattaa ggctgagcaa cctctgaaat tcaccgctcc gaaactggaa

cgttacggcg tgttcaagac gatggacgag gcgatcgctg cgtctgaaga ggcacagaaa

aagctgctgt tctccaaaat cagcgatcgt cagaagtacg ttgatgtgat tcgtagcacc

atcattaagc gcgagaacct ggaactgatc agccgcctgt ctgttgaaga gactgaaatt

ggtgactacg aacacaaatt gatcaaaaat cgtctggcag cggaaaagac gccaggcacg

gaagatctgc tgaccgaggc cattacgggt gataacggct tgaccctggt tgagtattgc

ccgttcggtg tgattggtgc gattaccccg accaccaatc caaccgaaac gatcatcaat

aacagcatca gcatgattgc gggtggcaac acggtcgtct ttagcccgca tccgcgtgca

aagaaggtga gccagatgac cgtcaagatg ctgaacaaag cactgattga caacggcgca

ccgccgaatc tgatcactat ggtggaagag ccgtctattg agaacacgaa caaaatgatc

gacaatccgt ccgttcgcct gctggttgct accggtggcc cgagcatcgt caagaaagtc

ctgtccagcg gcaagaaagc catcggtgcc ggtgcgggta atccgccagt cgttgtcgac

gagactgccg acattgataa ggcggccaaa gatattgtgg atggttgtag ctttgacaac

aatgtgccgt gcattgcaga gaaagaagtc tttgcggttg actcgatttg cgactacctg

atccaccaca tgaaagagaa tggcgcgtat cagatcacgg accctatgtt gctggagcaa

ctggttgcgc tggttacgac cgaaaagggc ggtccgaaaa ccagcttcgt gggcaagagc

gctcgttata tcctggataa gctgggtatc acggtcgatg cgtccgtccg tgtgattatc

atggaagtgc cgaaggatca cctgttggtg caagaagaga tgatgatgcc gatcctgccg

gtggtccgtg ttagcgatgt ggataccgca atcgagtacg cacaccaggc ggagcatggt

aatcgccata ccgcgatgat gcacagcaaa aacgttgaga aactgagcaa aatggccaag

attatggaaa ccacgatctt tgttaagaac gcgccgagct atgcgggcat tggtgttggt

ggtgagggct acaccacctt cactatcgca ggcccgaccg gtgagggtct gaccagcccg

cgtaccttct gtcgtaagcg caaatgtgtt atgacggacg cctttagcat tcgttaa

**ALDH7**

atggaacgca acttgtcggt actctcgcaa actaatgact tgaaaatcac taaacgcacg

gaaggtgata aaagcaataa caaagaaagc tatctgggtg tgtttaagaa ggtcgaaaat

gcgatcacca aagccattta cgcgcagaag aaactgtctc tgtattacac caaagaggac

cgcgagcgta tcattaagag cattcgtaag gccaccttgg aaaacaaaga gatcctggcc

aagatgatcg tggatgaaac gcacatgggc cgttatgagg acaagatcct gaagcacgag

ttggtggcga aatacacgcc tggtaccgag gacctgatca cgaccgcgtg gagcggcgat

caaggtctga cgctggtcga aatgagcccg tacggcgtta ttggcgcgat tacgccgagc

accaatccta ctgaaaccgt gatctgcaac agcattggta tgattgcagc tggcgattcc

gtcgtgttta atggtcatcc gggtgccaag aaatgtgttg cgtttgcagt cgacatgatt

aacaaagctg ttatccgtga gggcggtccg gagaacctgg tgaccacggt ggagaacccg

acgatggaga gcctgaatgt cattatgaag cacccgtaca tcaagctgct gtgtggcacc

ggtggtccgg gtttgattaa gaccctgctg aactccggta agaaagcgat tggcgcaggc

gctggtaatc cgccggttat tgttgatgat tccgccgaca tcgacaaagc ggcaaagaac

atcattgagg gttgcagctt cgacaacaat ctgccgtgta tcgcggaaaa agaggttttt

gtgttcgaga atgtcgcgaa tgatctgatt cagaacatga tcaagaataa cgcagtgctg

attaatgaaa accaagtcag caaactgctg gatctggttc tgctggagcg caaggatgaa

accctggagt atgcgattaa caagaaatgg gtgggtaagg atgcgaaact gtttctggac

aaaatcggca ttaaggctag cgataacgtt cgttgcatca tctgcgaagt tgacgcgaac

cacccgttcg ttatgaccga attgatgatg ccgattctgc caattgtccg tgttaaggac

gtcgacgagg cgattgaatg tgcgaaaacc gcagagcagc gtaaacgcca ttctgcatat

atgtacagca agaatattga caatctgaat cgttttgaaa aagagatcga tacgacgatc

ttcgtgaaga atgccaaaag cttcgcgggt gtgggtttcg gtgcagaagg ctttacgacc

ttcaccatcg ctggcccgac cggtgagggc atcaccagcg cacgtaactt cacccgtcag

cgccgttgcg ttctggccgg ttaa

**ALDH8**

atgaataaca atctgtttgt aagccctgaa acgaaagact tgaaactgcg cactaatgtt

```
      gagaacttga aattcaaagg ttgtgagggt ggctccacct acatcggcgt gtttgagaat
      gcagaaaccg cgatcgacga ggcggttaac gcgcaaaagc gtctgagcct gtactacacc
      aaagaacagc gtgagaagat tatcacggaa attcgtaaag ttaccctgaa gaataaagag
      attctggcac aaatgattct ggaagaaacg cacatgggtc gttatgaaga taagatcctg
      aagcacgagc tggtcgcgaa gtatacgccg ggtaccgagg acctggcaac caccgcgtgg
      tccggtgaca acggcctgac tgtcgtggag atgtctccgt acggtgttat tggtgcgatc
      accccgtcga ccaatccgac cgaaacgatc atctgcaaca gcatcggtat gatcgcgagc
      ggtaacgcgg ttgttttcaa cggccatcca ggcgcaaaga aatgtgtggc gttcgcggtt
      gatatgatta accgcgcgat tatcagctgc ggtggtccgc gcaatctggt gaccgcgatc
      aagaacccga cgatggagag cttggatgcc atcatcaagc acccggcgat taagctgttg
      tgcggtacgg gcggtccggg tatggttaaa accctgctga gcagcggtaa gaagagcatt
      ggtgcaggcg ctggtaatcc accggtcatt gtggatgaca ccgccgacat cgagaaggct
      ggtaagagca ttatcgaggg ttgtagcttc gacaataatc tgccgtgcat cgcggaaaaa
     gaggtgttcg tttttgaaaa cgtcgcagac gacctgatta agaatatgct gaaaaacaat
     gcagtcatta tcaataaaga ccaagttagc cgcctggtga atctggtcct gcagaagaac
     aatgaaacca gcgaatatac catcaacaag aaatgggtcg gcaaagacgc aaagctgttc
     ttggatgaga ttgatgtcga gtctagctcc gatgttcgct gcattatctg cgaagtggat
     gccgaccacc cgttcgtcat gaccgaactg atgatgccga tcctgccgat tgtgcgtgtg
```

aaagatattg atgaggccat caaatatgcc aaaattgccg agcagaaccg taaacatagc

gcgtacatct atagcaaaaa cattgagaac ctgaatcgtt ttgaaaaaga gattgatacg

accatttttg tgaagaacgc gaagtcgttt gcaggcgtcg gctacggcgc tgagggtttc

acgactttta ccattgctgg ctgtacgggc gagggcatca cgagcgcccg taacttcacc

cgtcagcgtc gctgtgtgtt tgttggttaa

**ALDH9**

atgaatgact ttaacatgat cgatatcgag agcattgtca aaaacattgt aaaagaattg

accggtaacg agaagggcca gggtgcgatc acgaccgcga ccgctccgaa agaagccaat

ccgctggttg acattgagaa aaagattatg ggttttatga ataccccgac catgcctgtg

ggtgagtacg gcgtgttcga ggacatcaac gacgcgatcg aacaagcatg gctggccgag

caggagtatc gtaaagttgg cctggataag cgtacggaga ttatcgaggc tttcaaggca

gaagtgcgca aaaatgtcga agagatctcc cgtcgtacct ttgaagaaac gggtatgggc

cgttatgagg ataagatcct gaaaaacaac ctggccttgg ataagacgcc gggtgtggaa

gatctggaag cgggtgtgaa aacgggcgat ggtggtctga ccctgtatga gatgtcgccg

ttcggtgtca ttggtgcgat cgctccgagc accaatccga cggaaactat tatcaataat

ggcattagca tgctggcggg tggtaacacc gtcgtgttca gcccgcatcc aggtgcgaaa

gacgtcagcg tgtttatcgt tcaactgatt aacaaagcga tcgagcgtat caacggtccg

aagaacctga tcgttacggt gaagaacccg aacatcgaaa gcaccaacat tatgttggcg

catccgaagg tgaatatgat ttgcgcgacc ggcggtccgg gcatcgttaa ggttgctctg

agctctggca agaaggcgat tggtgccggt gcgggcaatc cgccggtggt ggtggacgaa

accgcagaca tcgagaaagc ggcagttgac attatcgacg gctgtagctt cgacaataat

ctgccgtgta tctgcgagaa agaggtcatt gttgttgaca aggttgcgga ctacctgaaa

acgtgtatga gcaagtattg cgcactggag attacggaca agaacatgtt ggcacagctg

gagaagctgg tgctgaccga aaatggcacg atcaacaaac aatttgtcgg caagaacgca

gattacatta tgagcaaatt gggtgtcaat atcgatccga gcattcgcgt catctttgca

gaggtggaag cgaatcaccc gttcgccgtc gaagagctga tgatgcctat tctgccggtc

atccgtgttc gcaacgttga tgaggccatc gatctgggtg tagagctgga acatggtaat

cgtcacaccg cgatcatgca cagcaaacac attgataatc tgtccaagtt tgccaaagcg

gttcagacca cgattttcgt caaaaacgcg ccatcctacg caggcattgg ttacggcgca

gaaggccacg gtaccttcac cattgccggt ccgactggtg agggcctgac cagcgctcgc

accttcactc gcaaacgtcg ttgcgttatg gttgacaact tttctattaa gtaa

**ALDH10**

atggaattgg aaagcaacga attgagcgtg attattgaga aggtactgaa agaaatgaac

aagaaagagt ttggtaagaa agagagcgac ggtattttcg atacgatgga cgaggccgtt

gaggcgtctt acgaggcaca gaagaaatac agctcgtact ccctggagca gcgcgagaag

ctgattcaag caatgcgtaa agcgatcatg gataatgcga tggaagtcgc taatctgtgt

gtgaaagaaa gcggtatggg tcgtgtcgac cacaaatact tgaaactgaa attgattgtt

gaaaagacgc aaggtacgga aatcctgcgt ccggaagttt acaccggtga caacggcctg

accctgattg aacatggtgc tttcggtgtt atcggtgcca ttacgccgag caccaatccg

gcagcgaccg tcgcgtgcaa ctccatctgc atgctggcgg gtggtaatac tgtggttttt

agcccgcacc caggtgcgct gaatagctgc ttgaccatga tccgcattct gaataaagca

atcaaagagg ccggtggtcc ggagaacctg attaccagcg tgaaagcacc tagcattgag

aataccaata tcatgattaa ccacaagcgt attcgcctgg tcgtggctac cggcggtccg

ggcattgtga aactggtgct gtccagcggc aagaaggcga tcggtgcggg tgccggcaat

ccgccggttg ttgtggatga aaccgccgac attccgaagg cggcacgtga catcattgcc

ggctgcagct ttgacaataa tctgccgtgc atcgcagaga aagaagcaat tgtcgtcgag

agcgtttacg aagaattgat taaagagttc aagaaaaacc gcgtcgttta cgagctgacg

gacgaagagg ccgaaaaact ggttggcaag gtcctgaact atgatgagaa gaacaagaag

tatagcatca acaaaaagtt cgtcggtaaa gatgcgaaat atctgctgga gagcatcggc

aaggatgcgg gcacgggtgt tgagtgtctg atttatcgtg cggagaatag ccacccgttc

gtccaagaag agctgatgat gccgatcctg ccgatcgtca aggttaagaa cgtggacgaa

gcgatcgaaa ccgcagtgga agatgagcat ggcaatcgtc atacggcgat gatgcacagc

aaaaacgttg tgaacctgac gaagatggcg cgtgcgatcg ataccactat cttcgtgaaa

aacgcaccgt cttatgcggg tatcggcttt ggtggcgagg gtcacaccac ctttaccatt

gctggcccaa ccggtgaggg catcaccaac gccgttacct tcacgcgcca gcgtcgttgt

acgatggtgg actcttttcg catcgtgtaa

**ALDH11**

atggagatcg gcgcaaaaga aattgagtta atcgtaagag aagttttggc aggcattgaa

tctcgtggcc cgaagctgag ctacattccg gcccaaagcg acaacggtgt ttttgagcgc

gtggaagatg ccattggtgc ggcgcatacc gcgcaacgcg aatgggtcga gcattaccgt

gttgaggatc gccgtcgcat catcgaggca atccgtatga cggcaaagag ccacgcgaaa

accttggcga agctggtgtg ggaagaaacg ggcatgggtc gctttgagga taagattcag

aagcacatgg cagtcatcga gaaaacgcca ggcgttgagt gcctgaccac ggacgcaatt

tccggcgacg agggtctgat gatcgaagag tacgctccgt ttggtgttat tggtgcgatc

accccgtcca cgaacccaac cgaaaccatc attaacaata ctatcagcat gattgcgggt

ggcaatgcgg tggtgttcaa cgttcaccct ggtgcgaaga aatgttgcgc gcactgtctg

aagctgctgc atcaagctat cgtcgagaac ggtggccctg ccaacctgat taccatgcag

aaagagccga ctatggaagc tgtgaccaag atgacctctg acccgcgtat ccgtctgatg

gtcggtacgg gtggtatgcc gatggtcaat gcgttgctgc gttcgggcaa gaaaacgatc

ggtgcaggcg ctggtaatcc gccggttatt gtggatgatt ccgcggacgt gagcctggca

gcgcgtgaga tttatcgcgg tgccagcttc gataacaata ttctgtgcct ggcggaaaaa

gaggtttttg tgatggagaa agctgcggat gaactggtta acaacctggt gaaagaaggc

gcatatctgc tgaatccgat ggagctgaat gagattttga aattcgcaat gatcgaaaag

aacggcagct gcgaggtcaa caagaagtgg gtcggcaagg acgccggtct gtttctggaa

gccattggcg tcagcggcca caaagacgtt cgtctgctga tttgtgaaac cgaccgcaat

cacccgttcg tcatggttga gcagctgatg ccgattctgc cgatcgtccg tctgcgcacc

ttcgaagagt gcgtggagag cgcggtggca gcggaaagcg gcaatcgtca cacggcgagc

atgttcagcc gcaatgtgga gaatatgacc cgtttcggta aagttatcga gactaccatt

ttcaccaaaa acggtagcac gttgaaaggt gttggtatcg gtggtgaggg tcataccacc

atgaccatcg cgggtccgac gggtgaaggt ctgacctgtg cccgtagctt tacgcgtcgt

cgtcgctgca tgctggccga gggcggtttg cgtatcattt aa

**ALDH12**

atggacgcac aaaagattga aaaactggta cgcaagattt tggaagagat ggaagagaaa

aagaaaccgg ccgagactga gtgtgaatgg ggtatctttg accacatgaa ccaggcgatt

gaagcggcgg aaattgcgca aaaagagctg gttcaactga gcctgggtca gcgtggcaaa

ctgattgaag caattcgtaa ggctgcgaaa gagaacgcgg agaagttcgc gcgcatggca

gtcgatgaga ctggtatggg caaatacgag gacaaaatcg tcaaaaatct gctggctgcc

gaaaagaccc cgggtatcga agatctgcgc accgaggtgt ttagcggtga cgacggcttg

acgttggtgg agctgagccc gtacggcgtg atcggcgcta tcaccccgac caccaacccg

accgaaacca tcatttgtaa ttccattggt atgatcgcgg caggcaacgc agtcgtcttt

tccccgcacc cgcgtgcgaa gaacacctct ctgtacgcaa ttaagatttt caatcaggcg

atcgttgagg cgggtggtcc gaagaacctg attaccacgg tagcaaaccc gagcattgaa

caagccgaga tcatgatgaa gcacaaaacg atcaaaatgc tggttgctac cggtggtccg

ggtgtggtga aggcggttct gagcagcggt aagaaggcca tcggcgctgg tgcgggtaat

ccgcctgtgg ttgttgacga aactgcggat attgagaagg cagccaaaga catcatcgca

ggctgctcgt tcgataacaa tttgccgtgc gttgccgaga aagaggtgat tgcagtggaa

agcatcgcag atcgtctgat cgactatatg aaaaagcacg gtgcgtatga gattaccaat

aaagagcaga tccagcaact gaccgatctg gttgtcgaga acggccatgc caacaaagag

ttcgtcggta aagacgccgc gtacatcctg aagcatatcg gtatcaatgt tccgccggat

acccgtgtgg ccattatgga agtggatggc aaacacccac tggttacggt tgagctgatg

atgccgatcc tgccaattgt gcgtgtcaaa aatgttgacc aggcaatcga actggcggtc

gaagttgagc acggcttccg tcatacggcg attatgcata gcaagaacgt tgatcacctg

acgaaattcg caaaggcgat ccagacgacc atttttgtga agaatgctcc tagctatgcg

ggcattggtg tgggcggtga aggttacgct acctttacca tcgcgggtcc gacgggtgag

ggcctgacga gcgcgaagga tttcgcgcgt aagcgcaaat gcgtcctggt cgacgccttg

tctattcgct aa

**ALDH13**

atgaacaagg atacgacgat tagcgaaacc gagaacttga aatttaaaac gaacattaag

aatgctgacc tgaagaatta cgagaatagc acgagctatt ccggcgtttt tgaagatgtc

gaggtggcga tcaacaaggc catcaccgcg cagaaagagt tcagcctgta ctatacgaaa

gagcagcgcg agaaaatcct gactgagatt cgtaaagcga ccctgaaaaa caaaaagatt

ctggcgaaga tgattctgga cgaaacccac atgggccgct atgaggataa gatcttgaag

catgaactgg ttgcaaaata caccccgggt attgaggatc tgactaccac cgcttggtcc

ggcgacaatg gcctgaccgt tgttgaaatg gcgccgtacg gtgtgattgg tgcaattacg

cctagcacca acccgacgga aaccgttatc tgcaatagca tcggtatgat cgcagcgggc

aatgcagtgg ttttcaatgg tcacccgagc gcaaagaagt gtgtggcctt tgctgtcgat

atgatcaata aagcaatcgt cagctgtggt ggcccgaaaa acctgattac cgcggtgaaa

aacccgacga tggagagctt ggatgcgatt atcaagcatc cggaaatcaa actgctgtgt

ggtaccggtg gcccaggtat ggtgaaaacc ctgttgaaca gcggcaagaa agccatcggt

gcgggtgccg gtaatccgcc ggtgattgtc gacgataccg cggatatcga aaaggcgggt

aaaaacatca ttgagggttg ctcgttcgac aataatctgc catgcatcgc cgaaaaagag

gtttttgtct ttgacaacgt tgccgacaat ctgattgata acatgttgaa gaataacgct

gtgatcatca ataaggacaa aatcaccaag ctgctgaatc tgatcctgca gaaaaacaat

gaaacgcagg agtataacat taacaagaag tgggtcggca aagacgcgaa gctgttcctg

aatgagattg acgttgaggc gccgagcagc gttcgttgca ttatctgtga ggtggaaccg

gatcacccgt tcgtgatgac cgagctgatg atgccgatcc tgccgattgt ccgtgttaag

aacattgacg acgcgatcca atacgcaaag atcgcggaac aatctcgcaa acacagcgcg

tacatttact ccaaaaacat cgataatctg aatcgttttg aaaaagagat tgacaccacg

attttcgtca agaacgcaaa gtctttcgcg ggtgtgggct ataacgcaga aggtttcacg

accttcacta ttgcgggttg cacgggcgag ggtattacca gcgctcgtaa ctttacccgt

caacgccgtt gcgtcctggc cggctaa

**ALDH14**

atggaatttg aggtaaacaa cattgaagaa attgtggaac tgattatgaa gaagatggca

gagtctaaca tcagcacggc gggtaattcc aaaaatggtg tgttcgacaa tgtggacgag

gcgattgaag aagcgaagaa agcgcaggca attctgttca gcagcaagtt ggagctgcgt

gagaagatca tcgctagcat tcgcgacacc ctgaagaatc acgttaccga gctggcagag

ttggcagtta aagaaaccgg tatgggtcgt gtcgcggaca aagagttgaa aaacaaaatc

gctattgaaa agaccccggg tttggaagat ctgaaggcat tcgcattcag cggtgatgat

ggcctgacgg ttatggaact gtccccgtat ggtgtgattg gcgcaattac gccgagcacc

aacccgagcg aaacggtgat ctgtaacagc atcggcatga tcgccgctgg taatgcggtg

attttcgcac cgcatccggg tgccaagcgc accagcatcc gcaccgtcga gctgatcaat

gaggcgatcc gtaaggttgg tggccctgat aatctggttg ttaccatccg tgagcctagc

attgagaata ccgagaaaat cattgccaat ccaaatatca aaatgctggt tgctaccggc

ggtccgggcg ttgtcaaaac cgttatgagc agcggtaaga aggcgattgg tgccggtgcg

ggcaatccac cggtcctggt cgatgaaacc gcggacatcg agaaagccgc gaaagacatt

attgcgggct gtagctttga caacaatctg ccgtgcactg ccgagaaaga ggtcgttgca

gttgattcta tcgtgaacta cctgatcttt gagatgcaaa agaacggcgc gtatctgctg

aaggacaaag aactgattga aaagctgctg agcctggtgc tgaagaacaa cagcccggat

cgtaagtacg tcggtcgtga cgccaagtat ttgctgaaac agatcggtat cgaggtgggt

gatgaaatca aggtcattat cgtcgaaacg gacaagaacc acccgttcgc tgtggaagag

ttgctgatgc cgattctgcc gatcgtcaaa gttaaagacg ccctggaagg tatcaaagtc

gcgaaagagc tggagcgtgg cctgcgtcat actgcggtga tccactctaa gaatattgat

attctgacca aatacgcgcg tgagatggaa acgacgatcc tggtgaaaaa cggtccgagc

tacgcgggta ttggtatcgg cggtgagggc cacgttacgt ttaccattgc aggcccgacg

ggcgagggtc tgacctcggc gaaatccttc gcgcgcaacc gccgttgcgt attggtgggc

ggttttagca ttaaataa

**ALDH15**

atgaatttgg aagcaaacaa catggacgaa attgtggcac tgattatgaa agaactgaag

aaaaccgaca ttaaggcggg ttgtcaatct tgtgagagct tgaaaaacgg cgttttcagc

agcatggatg aggccattgc tgcagcgaag aaggcgcagg agatcctgtt cagctcccgt

ctggagatgc gtgagaagat tgtcgcgagc attcgcgaag tgatgaagga ctatgttgtg

gagctggccg agctgggtgt gaaagaaacc ggtatgggtc gtgccgcaga caaagcgctg

aaacaccagg tgacgatcga gaaaaccccg ggtgttgagg acttgcgcgc ctttgcgttt

agcggcgatg atggtctgac cgtcatggag ctgagcccgt atggcgtgat tggcgcgatc

accccaagca ccaatccgtc cgaaacgatc atctgcaata gcattggcat gatctccgct

ggcaattctg ttgttttcgc gccacatccg ggtgcgaaac gcacgtcgat taagactgtc

gaaatcatta acgaggccgt tcgccgtgca ggcggtccgg agaacctggt ggtcacgatc

gcggagccga gcatcgaaaa caccaatcgt atgatggaga atccggatat caagatgctg

gtcgccacgg gtggtccggg tgtggttaaa agcgtcatga gcagcggtaa gaaagcgatt

ggcgcaggcg caggcaatcc gccggtgctg gttgatgaaa ccgctgatat cgagaaggcg

gcacgtgaca tcgtcgccgg ctgtagcttt gacaataatc tgccgtgcat tgctgagaaa

gaagtcgttg cggttgattc tatcaccgac tacctgattt ttgagatgca aaagaacggc

gcgtatctga ttaaagacaa atccgtgatt gaccgcctgg tggcgatggt tctgaagaac

ggtagcccga accgcgcgta cgttggcaaa gatgcgagct acatcctgaa agacctgggt

attaacgttg gcgacgagat tcgtgtgatc atcaccgaaa ccgacaagga tcacccgttt

gcagttgaag agctgctgat gcctatcctg ccgatcatcc gtgtcaagaa cgcgctggaa

ggtattgagg taagcaagaa attggaacac ggtctgcgcc ataccgcgat gattcatagc

aaaaacattg atatcttgac gaagtacgcg cgtgatatgg aaacgaccat cctggtcaag

aatggcccga gcttcgcagg catcggtgtg ggtggtgagg gtcacacgac tttcaccatt

gccggtccta cgggtgaagg tctgaccagc gcaaagtctt tcgctcgtaa tcgtcgttgc

gtgttggtcg gtggtctgag cattaaataa

**ALDH46**

atgaataaag acaccctgat tccgaccacg aaagatctga aagttaagac taacggcgag

aacattaacc tgaagaatta caaagacaat agcagctgtt ttggcgtctt tgaaaatgtg

gagaatgcga tttcttctgc ggtgcacgcg caaaagattc tgtccctgca ctatacgaag

gagcagcgcg agaaaatcat tactgaaatc cgtaaagcgg ccctgcagaa taaagaggtg

ctggcaacca tgattttgga agaaacgcac atgggtcgct acgaagataa gattctgaaa

catgagctgg tcgcgaaata caccccgggt accgaggact tgaccactac cgcgtggagc

ggcgacaacg gtctgaccgt cgtcgagatg agcccgtacg gtgtcattgg tgcaatcacg

ccgagcacca acccgacgga aacggtgatc tgcaacagca ttggtatgat cgctgcaggc

aacgcggtcg ttttcaatgg ccacccgtgt gcgaagaagt gtgttgcctt tgctgttgag

atgatcaaca aagcgattat cagctgtggc ggtccggaga atctggtcac gaccattaag

aatccgacca tggaatccct ggacgcaatc attaagcacc cgtcgattaa actgctgtgc

ggcaccggtg gtccaggtat ggttaagacg ctgctgaaca gcggtaagaa agcaatcggt

gctggcgctg gtaacccgcc tgtcatcgtt gacgatacgg cagacattga aaaggcgggt

cgttccatca ttgagggctg cagcttcgat aacaacctgc cgtgcattgc ggagaaagag

gttttcgtgt ttgagaatgt ggcagacgat ctgatcagca acatgctgaa gaataacgcg

gtaatcatta acgaggacca agttagcaag ctgatcgacc tggttttgca gaaaaacaac

gaaacccaag agtacttcat caataagaaa tgggtgggta aggatgcgaa gttgttcctg

gatgagatcg atgtggaaag cccaagcaat gtgaaatgca tcatctgcga agttaatgcc

aatcatccgt tcgttatgac cgaactgatg atgccgatct tgccgatcgt gcgtgtcaaa

gatatcgatg aggccattaa gtatgcgaag atcgccgaac agaatcgtaa acatagcgct

tatatctaca gcaaaaacat tgacaatctg aatcgcttcg aacgtgagat tgacaccacg

atttttgtga aaaacgcaaa aagctttgcg ggtgtgggct atgaggcgga aggcttcacc

acctttacca ttgcaggttc taccggtgaa ggtatcacga gcgcccgtaa cttcacgcgc

caacgtcgtt gtgttctggc cggctaa

**E. gBlock sequences for ADH screening**

*E1. Primary ADHs*

**aldh46.x G1**

ATCGCTTCGA ACGTGAGATT GACACCACGA TTTTTGTGAA AAACGCAAAA AGCTTTGCGG

GTGTGGGCTA TGAGGCGGAA GGCTTCACCA CCTTTACCAT TGCAGGTTCT ACCGGTGAAG

GTATCACGAG CGCCCGTAAC TTCACGCGCC AACGTCGTTG TGTTCTGGCC GGCTAATCTA

GATGAACAAC TTCCGTTTCT GCACCCTACC GAATTACTAG TATCGATGAT AAGCTGTCAA

ACATGAGCAG ATCTGAGCCC GCCTAATGAG C

**ADH1 G1 (Accession No. B6YQP9_AZOPC)**

CGAGCGCCCG TAACTTCACG CGCCAACGTC GTTGTGTTCT GGCCGGCTAA TCTAGACTCG

CCATATCCGA CCCACCCAAG GACAACTCAT ATGAACAACT TCCGTTTCTG CAGCCCTACC

GAATTCATTT TTGGTAAAAA CACCATCTGT AAAGTGGCTC AGCTGGTTAA ACAGTATGGT

GGCTCTAAAG TTCTGATCCA TTACGGCAAT AAATCTGCGA AAAAATCTGG TCTGCTGACC

CAGATCGAGA ACTGCTTCCA GAACGAATTT ATCGAATATG TCAAACTGGG TGGTGTTCAG

CCGAACCCGA TCGACGAACT GGTCTACAAG GGTATCGAAC TGGGCCGTAA AGAAAAAGTT

AACTTCATCC TGGCTATCGG TGGCGGTAGC GTTATCGACT CTGCTAAAGC AATCGCTGCG

GGCATTCTGT ACAACGGTGA TTTCTGGAAC TTTTTCGAAG GCATCGTTAC CATTAACCAC

GCCCTGCCAA TTGCAACTGT TCTGACCCTG CCTGCTGCGG GCTCTGAGGG TTCTCCGAAC

ACTGTCATCA CGAAAACCGA CGGTATGCTG AAACGTGGCA TCGGTTCTTC CTTCATCCGC

CCAGTCTTCT CTATCATGGA TCCAGTGCTG ACGTTCACCC TGCCGACCTG TCAGACCGTT

TATGGCATCG CAGATATGAT GGCCCACGTT ATGGA

**ADH1 G2 (Accession No. B6YQP9_AZOPC)**

ACCTGTCAGA CCGTTTATGG CATCGCAGAT ATGATGGCCC ACGTTATGGA ACGCTACTTC

ACCCAGACCC AGGGTGTGGA TATTACTGAC CGCATGTGCG AGTCTATCCT GCTGTCTATT

ATCCACAGCG CGAAAACTCT GATTCGCGAA CCGGAAAACT ACGACGCTCG TGCCAACATC

ATGTGGGCCT CCACGATCGC GCACAACGGT ATCTGCGGCG TGGGTCGTGA AGAAGACTGG

GCGACCCATG CTCTGGAACA TGAACTGTCC GCGCTGTATA ACATCGCACA CGGCGCCGGC

CTGGCTGTGA TGTTTCCGGC GTGGATGCAA TACGTATACA CCGCGGGTAT CGACCGTTTC

GTGCAATTTG CTACCCGCGT TTGGAACATC GAAAACATCG GCTCTAAAAA AGAGATTGCC

CTGAAAGGTA TCCACGCTCT GAAAGACTTT TTCTCCTCCA TCAAACTGCC AATCAACTTT

GAACAGCTGG GCGCACAGAA AAGCGATATT GACAAACTGA TTGACACCCT GAAAATTAAC

ACCAAAGGTA AACTGGGTAA CTTCCTGCTG CTGGACATGA ACGATGCTCG TGCAATCTAC

GAAATTGCTG CTAAGCGTTA AACTAGTATC GATGATAAGC TGTCAAACAT GAGCAGATCT

GAGCCCGCCT AATGAGC

**ADH2 G1 (Accession No. A0RQF7_CAMFF)**

CGAGCGCCCG TAACTTCACG CGCCAACGTC GTTGTGTTCT GGCCGGCTAA TCTAGAAGAT

TAAACTCTAA GCGAGGAATA CATGGTCAAC TTTTCCTACT GCAATCCAAC CCGTATCGAA

TTCGGCAAAG GTAAAGAAAA CTCCATCGGT GAATACCTGA ACGAATATGG CGCAAAAAAC

GTGCTGATTC TGTTCGGCTC CGACCGCGTT AAAAAAGACG GTCTGTTTGA CAAAGCGACT

GCGTCCCTGA CCAAATTCGG CATCAAATTC TCCGAACTGG GTGACATTGT GAGCAATCCA

GTACTGTCCA AAGTTTATGA AGCTATCAAC CTGGCCCGCA AAAACGGCGT GGATAGCGTT

CTGGCGATCG GCGGTGGTTC TGTCCTGGAT ACTGCCAAAT CCGTAGCAGC CGGTGCAAAA

TACGACGGTG ACGTTTGGGA TCTGTTCCTG GCCAAAGCTC CGATTAAAGA TGCTCTGATG

GTTTTCGATA TTATGACCCT GGCTGCAACT GGTAGCGAAA TGAACAGCTT CGCCGTTGTC

ACCAACGAAG ACACTAAAGA GAAAATCTCT ATCACCTCTT CCCTGGTGAA CCCAAAAGTA

AGCGTAATCA ATCCGGAACT GATGAAATCC ATTTCTAAAA ACTACCTGGT GTACTCCGCG

GCCGACATCA TCGCGCATTC TATCGAAGGC TACCTGACCG CAACTCATCA CCCGGAAATT

ATCTCCAAAC TGGTTGAAGC GAATATCTCC

**ADH2 G2 (Accession No. A0RQF7_CAMFF)**

CAACTCATCA CCCGGAAATT ATCTCCAAAC TGGTTGAAGC GAATATCTCC ACTATTATTA

AAACGACCGA AATCCTGCTG GCTGACCCAG ACAACTACGA CGCACGTGCG GAATTTGCGT

GGGCAGCAAC TTGTGCTCTG AACGGCACCA CTTACGTTGG CGTTGGTGGT TACTCCTACC

CGAACCACAT GATCGAACAT TCCATCTCTG CACTGTACGG TGTACCGCAT GGTGCGGGTC

TGTCCGTAGT AATGCCGGCA TGGATGAAAT GGTATAAGGA CAAAAATGAA GCCCAGTTCT

CTCGCTTCGC TAAAGTAATC TTCGGTAAAA ACAGCGCTGA TGAAGGTATT GAAGCCCTGA

AGACGTGGTT CAAAAAAATC GGCACCCCGA CCAAACTGCG CGACTTCGGC CTGGACATGT

CCGTATCTGA CATCACCACT GCTGCGCTGC ATCACGCTAA AGCATTTGGT ATCGCTGATA

TCTATACCAA AGACGTTCTG GAAGAAATTC TGAACCTGGC TTACTAAACT AGTATCGATG

ATAAGCTGTC AAACATGAGC AGATCTGAGC CCGCCTAATG AGC

**ADH3 G1 (Accession No. G5F136_9ACTN)**

CGAGCGCCCG TAACTTCACG CGCCAACGTC GTTGTGTTCT GGCCGGCTAA TCTAGAGATA

CCTCTCCCTT AAGAGCGAGG TCATTATGAT TAACTTCGAC TATTGCGTGC CGACTAAAGT

TGTTTTCGGT CATGGTGTTG AATCTAACGT TGGCAAATAC GTAAAAGAGT TCGGTGGTAC

CAAAGCGATG ATTCACTGGG GCGGTGACTA TGTTCGCGAT ACGGGTCTGC TGGACCGTGT

CGAAAAATCT CTGTCCGCGG AAGGTATCGG CTACGTTGAG TTTGAAGGCG TCGTACCGAA

CCCGCGCCTG TCCACCGCTA AAGAGGGCCT GGCTCTGGCG AAACGTGAAG GTGTAGATTT

CCTGCTGGCT ATCGGCGGCG GTTCTGCAAT CGATAGCAGC AAAACCATCG CATACGGTCT

GGCGAACGAT TTCGAGCTGG AAGACCTGTT CCTGGGTAAA GTAAGCACTG ACCGTATCGC

GGGCCTGGGT GCGATCTCTA CCCTGGCCGG CACCGGTTCT GAAACCTCTA ACTCTACTGT

TATCAACATC GATACGATGG GTGACGTCGA GCTGAAACGT AGCTACAACC ACGAATGTGC

CCGTCCGAAA TTCGCGATCA TGGATCCGGA ACTGACCTAT ACCGTTCCGG CATGGCAGAC

GGCCGCCGCT GGCTGCGACA TTATGATGCA CACTA

**ADH3 G2 (Accession No. G5F136_9ACTN)**

TTCCGGCATG GCAGACGGCC GCCGCTGGCT GCGACATTAT GATGCACACT ATGGAACGTT

TCTTCACTAC CGTTTCTCAT ACGGAACTGA TCGATCAAAT GTCCCTGGGT CTGCTGCGTG

CTGTCAAAAC CGCGATTCCA CTGGCTCTGG CTGAGCCGGA TGACTATGAT GCACGCGCCA

CCCTGCTGTG GGCGGGCTCT CTGTCTCACA ACGGTCTGAC CGGCACCGGT CAGCAGGGTG

ACTTCGCATC CCATGCAATT GAACACGAAA TGGGTGCTCT GTACAACTGC ACCCACGGCG

CAGGTCTGTG CGCGATGTGG TCTTCCTGGG CTCGTTATGT CATTGATGTG CGTCCGGAAC

GTTTCGCACA GTTCGGTGTG GAAGTCTTCG GTGTGGTAAA CGACTACTCT GATCCGAAAG

GTACCGGTCT GCGCGGTATC GAGGCTTGGG AAAAATTCTG CAAATCTGTG GGTATGCCGG

TACGTATGAG CGAACTGGCA ATCAACCCGA CTGATGAGGA GATCCGTCAT ATGGCTCAGG

GCGCCATTGA CGCCCGTGGT GGTGATCATT GCGGTTCTTT CATGGAACTG CGTGTTGATG

ACGTCGTAAA AATTCTGGAA ATGGCCCGCT AAACTAGTAT CGATGATAAG CTGTCAAACA

TGAGCAGATC TGAGCCCGCC TAATGAGC

**ADH4 G1 (Accession No. B1C7G7_9FIRM)**

CGAGCGCCCG TAACTTCACG CGCCAACGTC GTTGTGTTCT GGCCGGCTAA TCTAGACCCA

CCTTCCAAAA CTCCCAGAGG TATTCATGCA GAAATTTGAC TACTATACTC CGACCAAAGT

TATCTTTGGC AAAGGCACCG AAAACAAAGT GGGTAAAGAG ATGAAAAAAG ACGGTGCTAA

GAAGGCTTAT ATCGTTTACG GCGGCAAATC CGCGAAAAAA AGCGGTCTGC TGGACAAAGT

GGAGAAATCT CTGAAAGACG AAAACATTGA ATACAAAATG ATCGGTGGCG TGAAACCGAA

CCCTCGCCTG TCTCTGGCTC GCGAAGGTGT GAAGGAAGCG AAGGAATTCG GTGCCGATTT

TATTCTGGCG GTTGGTGGTG GCTCTGTTAT CGATACCGCA AAAGGCATCG CACATGGCGT

AGCAAACCCT GACACTGACA TCTGGGATTT CTGGGAAGGT AAAGCCAAGG TTGAAAAATC

CCTGCCTGTT GGCGTTATCC TGACCATTTC TGCTGCGGGT TCTGAAATGA GCAACTCCGC

GGTGCTGACG AATGAAGAAA CTGGCATGAA GCGTGGCCTG TCCACCGATT TCAACCGTCC

GAAATTCGCC ATCATGGACC CGGAACTGAC CTACACGCTG CCGGATTACC AGGTTGGTTG

CGGTGTGGTA GACATCATGA TGCACACCAT GGATC

**ADH4 G2 (Accession No. B1C7G7_9FIRM)**

ATTACCAGGT TGGTTGCGGT GTGGTAGACA TCATGATGCA CACCATGGAT CGTTATTTCA

```
     CTGACCTGAC TGATTGCCAG AACGATCTGA CCGATGAAAT CGCAGAGTCT CTGCTGCGTA
     TCGTTATCAA AAACGGTCGT GTAGCTTGCA AGAATAAAGA AGACTACCAC GCTATGAGCG
     AAATCATGTG GGCAGGTTCC CTGTCCCATA ACGGCCTGAC CGGTCTGGGC GCCCCGATGG
     ACTTTGCAAC GCACCGCCTG GGTCACTCTC TGTCCGCGAA ATTTGATGTT GCACACGGTG
     CGTCCCTGTC CGCCATGTGG CCGCACTGGG CTAACTACGT AAAACATAAA GACATCGAGC
     GTTTTGCACG CTATGCGCGT AACGTTTGGG GCATTACGGA AGGCACCGAT GAAGAACTGG
     CTGATAAAGG TATTGAAGCG ACCGTGGAAT TCTTCAAATC TATCAACATG CCGACCTGCT
     TTAGCGAACT GGGTATCGGC ATCCAGGATG AGGATGGCCT GCGTGAGCTG ACCAACCGTT
     GCTTCTACGT GAAAGGTACC AAAGTAGGTA AACTGATTCC GCTGACCGAA GAAGATATTT
     ACCCGATCTA TGTATCTGCG AACAAATAAA CTAGTATCGA TGATAAGCTG TCAAACATGA
GCAGATCTGA GCCCGCCTAA TGAGC
```

**ADH5 G1 (Accession No. YUGK_BACSU)**

CGAGCGCCCG TAACTTCACG CGCCAACGTC GTTGTGTTCT GGCCGGCTAA TCTAGATAAC

```
     ACACCTATCA AGAAATAATT CAGAGGTCCC AATGGAAAAC TTCACCTACT ACAACCCGAC
     CAAACTGATC TTCGGCAAAG GCCAGCTGGA ACAGCTGCGC AAAGAATTTA AACGTTATGG
     TAAAAACGTT CTGCTGGTTT ATGGTGGCGG CTCCATCAAA CGCAACGGTC TGTACGACCA
     GGTCACCGGC ATCCTGAAAG AGGAGGGCGC GGTGGTTCAC GAACTGAGCG GTGTTGAACC
     GAACCCGCGC CTGGCTACCG TGGAAAAGGG CATTGGTCTG TGCCGTGAAC ACGATATCGA
     TTTTCTGCTG GCCGTCGGTG GTGGCTCTGT CATTGACTGC ACCAAAGCAA TCGCGGCGGG
     TGTAAAATAC GATGGTGACG CTTGGGATAT CTTTTCCAAA AAGGTTACCG CCGAAGACGC
     TCTGCCGTTT GGCACCGTAC TGACCCTGGC CGCTACCGGT TCCGAGATGA ACCCGGATTC
     CGTTATCACC AACTGGGAAA CTAACGAAAA ATTCGTCTGG GGTTCCAACG TTACCCACCC
     GCGCTTCTCT ATCCTGGACC CGGAAAACAC CTTTACCGTA CCGGAAAACC AGACAGTGTA
TGGCATGGTT GACAT
```

**ADH5 G2 (Accession No. YUGK_BACSU)**

AAACACCTTT ACCGTACCGG AAAACCAGAC AGTGTATGGC ATGGTTGACA TGATGTCTCA

```
     CGTTTTCGAA CAGTATTTCC ATAACGTAGA AAACACTCCG CTGCAGGATC GTATGTGCTT
     TGCTGTGCTG CAGACCGTCA TCGAAACGGC TCCGAAGCTG CTGGAAGACC TGGAAAATTA
     CGAACTGCGT GAAACCATTC TGTACGCGGG TACCATTGCG CTGAACGGTA CTCTGCAGAT
     GGGTTACTTC GGTGATTGGG CGTCTCACAC TATGGAACAC GCAGTGAGCG CAGTGTACGA
     CATTCCGCAC GCGGGCGGTC TGGCGATTCT GTTTCCGAAT TGGATGCGTT ACACGCTGGA
     TACTAACGTG GGTCGTTTCA AAAACCTGAT GCTGAACATG TTCGATATCG ATACGGAAGG
     CAAAACTGAC AAGGAGATCG CCCTGGAAGG TATTGACAAA CTGTCCGCAT TTTGGACGAG
     CCTGGGCGCG CCGTCCCGTC TGGCCGATTA CAACATCGGC GAAGAAAAAC TGGAGCTGAT
     CGCAGACATT GCTGCGAAAG AGATGGAGCA CGGCGGCTTC GGCAACTTTC AGAAGCTGAA
     TAAAGACGAC GTACTGGCGA TCCTGCGTGC ATCTCTGTAA ACTAGTATCG ATGATAAGCT
GTCAAACATG AGCAGATCTG AGCCCGCCTA ATGAGC
```

**ADH6 G1 (Accession No. A8SGI9_9FIRM)**

CGAGCGCCCG TAACTTCACG CGCCAACGTC GTTGTGTTCT GGCCGGCTAA TCTAGACCTC

TCCCGGTACG ATAATAAGGA GGCATCAATG AACAACTTCC TGTTCGAAAA CAAAACCAAA

GTATACTTCG GTAAGGGTGG TGTTAAAGAA TATCTGGGTT GTCTGCTGGA ACATTATGGT

GACACCGTTA TGCTGGCCTA TGGCGGCGGC TCCATCAAAC ATAACGGTGT ATATGATGAA

ATTGTGGGCA TCCTGAACGC CGAAGGCAAA CGCATCGTTG AATTCCCGGG TATCATGCCG

AACCCGACGT ATGCTAAGGT GCAAGAAGGT GCTAAACTGG CGCGTGAAAA CCACGTAGAC

CTGATCCTGG CCGTTGGCGG TGGTAGCGTT TCCGACTGCT GCAAAGTTGT GAGCGCGCAG

GCAAAAGTAG ATGAAGATCT GTGGGAGCTG GAAAACACTA AACACACTCG CCCGACTGCA

TTCATTCCGC TGGGTACCAT TGTGACCGTT TTTGGTACTG GCAGCGAAAT GAACAACGGC

GCTGTAATCA CCCACGAGGA GAAAAAAATT AAAGGTGCTC TGTGGGGCGC ACAGGCGGAC

TTTGCATTCC TGGACCCGAC TTATACTCTG TCCGTGCCGA TGAAACAGGT TATTAGCGGT

GCGTTCGACA CTCTG

**ADH6 G2 (Accession No. A8SGI9_9FIRM)**

ACTCTGTCCG TGCCGATGAA ACAGGTTATT AGCGGTGCGT TCGACACTCT GAGCCACGCT

ATGGAAACTT ATTTCGGCAA ACCGGATGAG AACAATCTGT CCGACGACAT CAACGAAGCG

GTGATGCGTT CCGTTATCCG TAACATTCGT GTGCTGCTGA CCGACAAGGA TAACTACGAA

GCACGCTCCG AACTGACCTG GGCTTCTGCG ATGGCAGAAA ACGGTATTCT GAAAATCGGT

AAAGTAACTG ACTTTCAATG CCACATGATC GAACATCAGC TGGGCGCATA CACTAACTGT

AACCACGGCG CTGGTCTGGC GGTTATCCAC CCGGTTCTGT ATCGTCATCT GCTGCCGGCG

AACACCGCAC GTTTCGCGCG TTTCGCTCAA AACGTTTGGG GCATCGATCC AGCAGGTAAA

TCCGAACTGA AACTGGCGCA GGCGGGTGTG GAAGCTCTGG CGGCGTTTAT CAAGGAAATT

GGCATGCCGA CTACCTTCGC TGAGCTGGGC GTTCCGGCGG ACACCGATCT GAAAGCCGTA

GCTGACTCTA CCGTCCTGAC CGGTGGTTGT TGCAAAAAAC TGTCTCGTGA AGAGCTGCTG

GACATCCTGA ACGAATGTAA ATAAACTAGT ATCGATGATA AGCTGTCAAA CATGAGCAGA

TCTGAGCCCG CCTAATGAGC

**ADH7 G1 (Accession No. E2SQ66_9FIRM)**

CGAGCGCCCG TAACTTCACG CGCCAACGTC GTTGTGTTCT GGCCGGCTAA TCTAGATATC

CAGCCATTCC CCAGGAGAAA CCACTATGCG TAACTTTACC TACCACAACC CGGTCCGTAT

CCTGTTCGGC GATCATGCTC TGGACCAGCT GCCGGATCTG TTCCGTGAAT TCCACGTGTC

TAACCTGCTG CTGGTGTATT CTGGCGATTT TATTAAAGAA CTGGGCATCT GGGATGCCGT

TTACAACGCT TGCGCGGAAA ATGGTATCGC ATTTTACGAA GAAGGTGGTG TAGTCCCGAA

CCCGAAAATT GAACTGGTTC GTGAACTGGT CGCACTGGGC AAAAAAAAAA AGATCGACTT

CATTCTGGCT GTAGGCGGTG GTTCTTCCAT CGACACTGCT AAGGCTGTTG CCGCAGGCAT

CCCGTACGCC CACGACGTGT GGGACTTCTT CGAATACACT GCGGTTCCGG AAACGGCGGT

GCCGATCGGT GTAATCACCA CGATCCCAGC GTCTGGTTCC GAATGTTCTA ATTGCAGCAT

TATCTCCAAC GGTCTGCACA AATGCGGTAT TGAGTACGAT TGCATCATCC CACAGTTTGC

CATCATGAAC CCGGAGTACA CCCGTACCCT GCCTGCGTAC CAGACCTCCG CAGGCATCGC

GGACATTCTG TCCCA

**ADH7 G2 (Accession No. E2SQ66_9FIRM)**

GTACCCTGCC TGCGTACCAG ACCTCCGCAG GCATCGCGGA CATTCTGTCC CACATGCTGG

AACGCTACTT CACGAACACT ACTCACGTTG ACACCACCGA CTACATGCTG GAAGGTACCA

TGCAGGCTCT GATGGTCAAC GCGCGCCGCC TGATGAAACA GCCGGATGAC ATCCACGCGC

GCGCAGAAGT TCAGTGTCTG GCTTTCCTGG CACATAACAA CCTGCTGGAC ATCGGTCGCG

AATCTGACTG GGGCCCGCAT CGTATTGAAC ACGAACTGTC CGCACAGTAC GGCATTACCC

ACGGTGAAGG TATGGCAGTT GTAACCATCG CGTGGGCACG CTACATGGCT GCACACCACC

CGGACAAACT GGCACAGCTG GCCTCCCGTA TCTTCGGTGC TGATCCGTTT GTACATTCCA

AAGAGGATAT GGCACTGCTG CTGGCTGACC ACCTGGAAGA GTTTTTCAAA TCCCTGCACC

TGAAAACCAC CCTGCACGAA ATGGGTATCG ACGATACCCA CTTTGAAGAG ATGGCAAACC

GTGCCACCAA TAACGGTAAG GATTGTGTTG GCCACTACGT GGCTCTGAAC AAACAGATCT

TTATCGACAT TCTGCACATG GCCCTGTAAA CTAGTATCGA TGATAAGCTG TCAAACATGA

GCAGATCTGA GCCCGCCTAA TGAGC

**ADH8 G1 (Accession No. E1QYZ8_OLSUV)**

CGAGCGCCCG TAACTTCACG CGCCAACGTC GTTGTGTTCT GGCCGGCTAA TCTAGAGACT

TAGTAGTCAC ACGCAAGAGG AGGATTCCAG TATGTACGAC TTCATGTTCC ACGTACCGAC

CAAGATCTAC TTCGGCCGCG GCCAGATCTC TCACCTGGCA GAACTGTCTG ATTTTGGCCA

GAAAGCGCTG CTGGTTTACG GTGGCGGCAG CATCAAACGT AACGGCATTT ACGACGAAGC

GATTCGTATT CTGACCCATG CGGGTATCGA AGTTGTAGAA CTGAGCGGCG TTGAACCGAA

CCCGCGTATT GAAACCGTGC GTCGCGGTGT CGGTCTGTGC GCTCGCGAAG GTGTTGACAT

GGTTCTGGCT ATCGGCGGCG GTAGCACCAT CGATTGCGCT AAAGTAGTTG CGGCCGGCGC

GCGTTACGAT GGCGACCCGT GGGACCTGGT ACTGGACGGT TCTAAGGCGG CTTCCGCGCT

GCCAATCTTT TCTGTGCTGA CCCTGTCCGC GACCGGTTCT GAGATGGATG CATTCGCTGT

CATCAGCGAT ATGAGCAAAA ATGAAAAGTG GGGTACCGGC GCAGAGTGTA TGAAACCGAC

CATGTCTGTG CTGGACCCGT CTTACACCTT CAGCGTGAGC CCTAAACAGA CCGCGGCTGG

CACCGCCGAT ATGAT

**ADH8 G2 (Accession No. E1QYZ8_OLSUV)**

ACACCTTCAG CGTGAGCCCT AAACAGACCG CGGCTGGCAC CGCCGATATG ATGAGCCATA

CCTTCGAATC TTATTTTTCC ATGGACGAAG GTGCGTACGT CCAGAAGCGT CTGGCAGAAG

GTCTGCTGGG CACTATGATC CACTTCGGCC CGATTGCCCT GGCACATCCG GACGACTACG

ATGCGCGTGC GAACCTGATG TGGGCGGCTT CTCACGCAAT TAACGGCCTG GTTTCTGATG

GTTGTAGCCC TGCCTGGTGC GTTCACCCGA TGGAACACGA GCTGTCTGCA TTCTACGATA

TCACTCACGG CGAGGGTCTG GCGATCCTGA CGCCGGCATG GATGGAGCAC GTTCTGGATG

CTCAGACTGC TCCTCTGTTT GCTGCATACG GTTGCAACGT ATGGGGTCTG TCCGGCGTAG

ATGACATGAA AGTTGCTCGT GAAGCAATCA GCCGCACTCG TGCGTTTTTT GTTGAAGCTA

TGCATCTGCC GGCAACCCTG CGCGAGGTCG GCATTACCGA TGAAAAAAAC TTCGAAGTTA

TGGCTCGCAA AGCCGCCGAT GGTTGCAAAG GCAGCTTCGT TGCGCTGTCT CAGGACGACA

TCGTAGAAAT CTACCGTGCT GCTCTGTAAA CTAGTATCGA TGATAAGCTG TCAAACATGA

GCAGATCTGA GCCCGCCTAA TGAGC

**ADH9 G1 (Accession No. F5X0G1_STRG1)**

CGAGCGCCCG TAACTTCACG CGCCAACGTC GTTGTGTTCT GGCCGGCTAA TCTAGACTCC

TTCAATAAGC CCAGGGAGGA TTAAAGCATG AATGATTTCC AGTTTCAGAA CACTACCAAA

GTTTATTTCG GTAAACATCA GCTGCAACAC CTGCACCAGG AAGTGCTGAA ATACGGTCAG

AAAGTGCTGA TCGCTGATGG CGGTGAATTC ATCCGTCAGT CTCCGCTGTA TGCTCAAGTT

CTGAAAGAAC TGACGGACAA CGGCATCCAG ATCTTCGAAC TGGGTTCTGT GGAGCCGAAT

CCGCGCCACA CCACCGTTAA CCGCGAAGTA AAACTGTGTA AAGGCAACAA CATCCAGACC

GTACTGGCCG TTGGCGGCGG CTCCACGATT GACTGCTGTA AAGCGATCGC GGCGACCTCT

TGCACCGACG AAGACGACGT TTGGACCCTG ATCGAAAAAC GTGAACCGAT CAACCAAGCG

CTGGCGGTTA TCGCTATGCC GACCATCGCG TCCACGGGCT CTGAAATGGA CAAGAGCTGC

GTGATTGCCA ACGAAGAGCT GCACCTGAAA AAGGGTCTGA ACGGCGAAGC TATCCGTCCG

AAAGCGGCTT TTCTGAACCC GGAAAACACC TTCACCGTTC CGGCACGTCA GACCGCGTGT

GGTGGCTTCG ACATCATGAT GCATCTGCTG GATAT

**ADH9 G2 (Accession No. F5X0G1_STRG1)**

CGTCAGACCG CGTGTGGTGG CTTCGACATC ATGATGCATC TGCTGGATAT GAACTATTTT

GTAGACTCTG ATAAATATCC GCTGCAGTTC AATGTGGTAG AAACCCTGCT GCGCACTATT

CGTGAGCAGC TGCCGATCGC GCTGCGTGAG CCGGAAAACT ACGAGGCTCG TGCGACCCTG

CTGTGGGGTG CTTCCTGGGC GCTGAACTCT TTCTGTACCT CCGGTTTCAA AACCGCACCG

AGCAACCACG GTCTGGAACA ATTCTCTGCG TTCTACGATC AGACGCATGG TCTGGGTCTG

GCTCTGGTGG TTACCAAATG GATGACCTAC CTGCTGGAAA AGGACCCGAC CGTGGCACCA

GATTTCGCTC GTCTGGGCAC CAATGTGCTG GGCTGTCAGC CAGTTGACGA TGTGATCGAG

GGCGCAAAAA ACGCTATCAA AGCCTTTGAC GCATTCATTG TGAATGACCT GGGTCTGCCG

CGTACCATGA CTGAAATCGG TCTGAACGAC TCTAAGCTGA GCGAGATGGC TCATGCTGCG

GTAACCGGTT ATGGCGACGG CACGCTGAAG GGCTACCGTG AACTGACTGA AGCGAACTGC

CTGGCCATTT ATAAAATGTG CCTGTAAACT AGTATCGATG ATAAGCTGTC AAACATGAGC

AGATCTGAGC CCGCCTAATG AGC

**ADH10 G1 (Accession No. E6W4G5_DESIS)**

CGAGCGCCCG TAACTTCACG CGCCAACGTC GTTGTGTTCT GGCCGGCTAA TCTAGAGCCT

TAATCCCCGT AAGCACAGGA GATCCACAAT GCAGAATTTC GTTTTTCACA ACCCGACCCG

TATCGTTTTC GGCCGTGACA AGACGGCGAG CATCGGCAAG GCGACCCTGC CGTATGGTCG

CCGCGTTCTG CTGCTGACGG GTCAGGGTTC CGTCGTGAAA CACGGTATCC TGGCGAAAGT

GACCTCTTCC CTGTCTACTG CGGGTATCTC CTGGGTTGAG TGTAGCGGTG TGCAGCCGAA

CCCGGTTCTG GGCTTCGTGC GTCAGGCCAT CGACACTTTC CGTCGTGAAA ACCTGGACGC

CATTGTAGCG GTTGGCGGTG GCTCCGTGAT CGACACCGCG AAGGCGGTGG CTGCGGGCGT

TCGTTACGAA GGCGATGTTT GGGACTTCTT TACCGGTAAA GCTAACGTCC TGGACGCGGC

CCCGATCACT GTAGTGCTGA CTCTGCCGGC GGCTGCATCC GAGATGAACA GCGGCGGTGT

TATCACTAAT GAACAAACTC GTCAAAAATT CAACCTGGGC GGCGAACCGC TGTCTCCGAA

AGTTTCTATC CTGGACCCGG TCAACAGCTT TAGCGCCCCG GTGAATCACT CCCTGTACGG

TGTTGTTGAC GCGAT

**ADH10 G2 (Accession No. E6W4G5_DESIS)**

ACAGCTTTAG CGCCCCGGTG AATCACTCCC TGTACGGTGT TGTTGACGCG ATGGTTCATC

TGCTGGAGGG CTACTTCAAC GGCTCTGACC CGTGGACTCC ACTGCAGGAC CGTTACGCGG

AAGGTATCAT TCGCACTCTG ATGGAATGCG CTGCCATTAT TCGTGAACAG CCAGACCACT

ACGACGCACG TGCTAACATC ATGTGGGGCG CGACTCTGGC TTTCAACGGC CTGGCACCGT

GCGGTATCGG CCCGGCAGGT TTTCCGATGC ACATGATCGA ACACAGCCTG TCTGCACTGT

ATGATGTATC TCATGGTGCG GGTCTGGCGA TGATCCTGCC GGGTTGGCTG AAGTACCACT

CCGATTCCAG CCCGCGCAAA GTTAACCAGT TTGGCCGTCG TATTTTTGAA CTGGATCACC

AGGATGATCG TCAGGGCGCT CAAGCAGCCA TTGCCGAGCT GGAACGTTGG CTGCGTTCCA

TGGATATCCC GGCATCCCTG CACGAAGGTG GCATCCCGAT CGATGAGATC CCAGCAATTG

CGGAGAACGC TGTGATGCTG GCGCAGAAAT GGGGTCTGAA AGCTTACACT CAGGCCGTTA

TCGAAGACGT TCTGCGTCGC GCTTCTCGCT AAACTAGTAT CGATGATAAG CTGTCAAACA

TGAGCAGATC TGAGCCCGCC TAATGAGC

**ADH11 G1 (Accession No. E6K7W2_9BACT)**

CGAGCGCCCG TAACTTCACG CGCCAACGTC GTTGTGTTCT GGCCGGCTAA TCTAGATGAT

CCCTCCACAA CTAAAGGCGG TATTCAAATG AAAGACTTCA ACTTCTACGC ACCGACCCGT

GTAGTGTTCG GCAAACAGAG CGAAGAGCAG CTGCCGCGCC TGCTGAAAGA AGCGGGTGGT

AAAAAGGTTC TGGTACACTA TGGTGGCGGC TCTGCAAAAC GTTCTGGCCT GCTGGATAAA

GTGTATGGTA TGCTGGACGA CGCGGGCATC GAACATGTAG GTCTGGGCGG TGTAGTACCG

AACCCGCTGC TGTCCAAAGT AAACGAAGGC ATTGACCTGT GCCGTCGTAA AGGTGTAAAC

TTCATTCTGG CTGTAGGCGG CGGCTCCGTA ATCGATAGCG CGAAAGCAAT TGCGTATGGT

GTGCCGTACG AGGGTGACGT TTGGGATTTC TGGAATGGTA AGCCGGCAAC CGCTGCCCTG

CCGGTCGGTG CAATGCTGAC TATCCCGGCT GCTGGCTCTG AAATGAGCAA TTCTTGCGTG

ATTACTAAAG ACGAAGGTGC TGTTAAACGT GGCTTCAACA ACGATCTGTG CCGCTGTAAA

TTCGCGATCA TGAACCCAGA ACGCACTTAC ACGCTGCCGC CGTACCAGAC TGCCGCGGGT

GCGACCGACA TCATG

**ADH11 G2 (Accession No. E6K7W2_9BACT)**

CACTTACACG CTGCCGCCGT ACCAGACTGC CGCGGGTGCG ACCGACATCA TGATGCACAC

CATGGAACGC TACTTTTCCA AACATGAAGA CATGACCCTG ACCGACGCAA TTGCGGAAGC

CCTGCTGCGC ACGGTTAAAG AAAGCACCTT CGAAGTGCTG AAACACCCGG AGGACTACCG

TAACCGCGCT CAGATTATGT GGGCCGGCTC CCTGTCTCAT AACGATCTGA CCGAATGTGG

TCTGGAAAAG GATTTCGCGA CTCACCGCCT GGAACACGAG CTGTCTGCGC TGTTCGGCGT

TACCCATGGC GCCGGCCTGG CAGCCGTGTG GCCTGCATGG GCGCGTTATG TGATGAAGAA

ACACATTTCC CGCTTCGTTC AGTTCGCGGT CAACGTGATG GGCGTTCCGA ACGACTTTTC

TAACCCGGAA GCTACCGCTG AGAAAGGTAT CTGTCGTATG GAACACTTCT TCCACGCGAT

CGGTATGCCG ACCTCCATCA AAGAACTGCT GGGTCATGAT ATCACCGAAG CGCAGATTGA

CGAAATGGTT GACAAATGCT CTCGTGGTGG TACTATCACT GTTGGTGCCA TGGAGGTGAT

TGCCCCAGAC GACATGCGTG CGATCTACCG TATGGCACGC TAAACTAGTA TCGATGATAA

GCTGTCAAAC ATGAGCAGAT CTGAGCCCGC CTAATGAGC

**ADH12 G1 (Accession No. B1C4Z8_9FIRM)**

CGAGCGCCCG TAACTTCACG CGCCAACGTC GTTGTGTTCT GGCCGGCTAA TCTAGATGGT

TCTACAATAA TAGGAGGACT CTACACATGC TGGGCGACTT TACCTACTCC AACCCGACGA

AAATTTATTT CGGCGAGAAC TCTCTGGACA ACCTGTCTAC CGAACTGAAA AACTATGGCA

AGAACGTGCT GCTGGTATAC GGTGGTGGTT CTATCAAAAA AAACGGTATC TACGATAAGG

TTATCGACAT TCTGAAAAAG TGTGATAAGA CTATTATTGA GGATGCGGGC GTAATGCCTA

ATCCGACTGT TGAAAAGCTG TATGAAGGTT GCAAACTGGC TCGTGAAGGT AACGTTGACC

TGATTCTGGC GGTTGGCGGT GGCAGCGTGT GTGACTACGC GAAAGCAGTT AGCGTCAGCA

CGTATTGCAA CGAGGATCCG TGGGAAAAGT ACTACCTGCG TATGGAGGAC GTTGATAACA

AAATTATCCC AGTTGGTTGT ATCCTGACCA TGGTTGGTAC TGGTTCCGAA ATGAATGGCG

GCTCTGTTAT CACCAATCAT GAACAGAAAC TGAAAATTGG TCACGTTTTC GGCGACAATG

TGTTCCCGAA GTTCTCCATT CTGAACCCGA CCTTCACCTA CACGCTGCCG AAATATCAGA

TGATCGCTGG TTTCT

**ADH12 G2 (Accession No. B1C4Z8_9FIRM)**

AACCCGACCT TCACCTACAC GCTGCCGAAA TATCAGATGA TCGCTGGTTT CTACGACATC

ATGTCCCATA TCCTGGAACA GTACTTTAGC GGTGAAGACG ACAACACCTC TGATTATATC

ATGGAAGGTC TGCTGAAATC TCTGATCCAT TCTAGCAAAA TTGCCGTGAA CGATCCTACC

AACTACGAGG CTCGTTCTAA CATCATGTGG ATTGCAACCT GGGCTCTGAA CACCCTGGTG

GCTAAAGGCA AAACCACGGA TTGGATGGTT CACATGATCG GCCAGAGCAT CGGTGCTTAC

ACCGACGCCA CGCATGGTAT GACCCTGGCT GCCGTGTCCA TTCCGTACTA CAAGTACATT

TGTCCATACG GCCTGAACAA ATTCAAACGC TATGCGATTA ACGTTTGGGA TGTTCTGTCT

GAAGGCAAAA CTGACGAGCA GATCGCTAAC GAAGGTCTGG AATGTATGGA AAAATACATG

CGTGACCTGG GTCTGGTAAT GAACATTTCC GATCTGGGCG TCAAAGAAGA GATGCTGGAG

GGTATCGCTG AAGGTACGTT CATCATGAAC GGCGGTTATA AAGTACTGAC CAAAGACGAA

ATTATCACCA TCCTGAAACA ATCCATGAAA TAAACTAGTA TCGATGATAA GCTGTCAAAC

ATGAGCAGAT CTGAGCCCGC CTAATGAGC

**ADH13 G1 (Accession No. G4L3E3_TETHN)**

CGAGCGCCCG TAACTTCACG CGCCAACGTC GTTGTGTTCT GGCCGGCTAA TCTAGAACGT

AAGGCCACTA CATTAACTAA GGAGCAAAAT ATGGAAAATT TCGATTTCCA CGTTACTACT

GATATCCGCT TTGGCAAAGA CCGTCTGGGT GAACTGCCGC AGGTTCTGAA CAACTTCGGC

AAAAACGTGC TGCTGGTTTA CGGTGGTGGC TCCATCAAGC GTAATGGTCT GTACGACAAA

CTGTACGAAC TGTTCAACCA GAACGACAAT AACGTTGTTG AACTGGCGGG TGTAGACCCG

AACCCGCGCA TTGAAACCGT GCAAAAAGGT GTCCAGCTGT GTAAGGAACA CGCGATCGAC

GTCGTGCTGC CGGTAGGTGG CGGCTCTGTG ATTGACTGCT CCAAAGCTGT GGCGGCTTGC

GTCTTTGTTA GCGGTGACCT GTGGGAAAAC TTCGTGCTGC AGAAAAACTA TAAAGGCCCG

GCACTGCCGA TTGTCACCAT TCTGACGCTG GCCGCTACGG GCTCTGAGAT GAACGGTACG

TGCGTAATCT CTAACATGGA TGCGCAGATT AAACTGGGCG TCCACGGTAC CACCAACCTG

CTGCCAAAGG TATCCTTCCT GGATCCGACT AACACCTTCT CTGTTGGTGC ATACCAGACT

GCAGCTGGCT CCGCTGACAT CCTGAGCCAC CTGAT

**ADH13 G2 (Accession No. G4L3E3_TETHN)**

TGGTGCATAC CAGACTGCAG CTGGCTCCGC TGACATCCTG AGCCACCTGA TGGAGAACTA

TTTCAACGCG ACCGAAGGCA CCGAAGTTCA GGATGAAATC GCTGAAGGCC TGATGAAAAC

GGTGATCAAA TATCTGCCGG TGGCGCTGGA CGAACCGGAC AACTATATTG CCCGTGCTAA

CCTGATGTGG GCCTCTACTC TGGCGCTGAA CGGCCTGGTT GGCAAAGGTA AAAAAGGCAG

CTGGTCTTGT CATGCTATGG AACACGAACT GTCCGCTTTC TATGACATCA CTCACGGCGT

CGGCCTGGCT ATGCTGACCC CGCGTTGGAT GGCACACATC CTGGACGAAG ACACCCTGCC

GAAATTTCAA CGTTTTGCTG AAGAGGTCTG GAATGTTAAA GAAAAGGAAC CGAAACGTAC

GGCGGAGATC GGCATTCAGA AACTGTACGA TTTTTTCGTC TCCTGCAACA TCCCTATGAC

CCTGTCCGGT GTGGGCATCC AGACCGAAGA AAATTTTGAA GAAATGGGTC AGCGTGCCGT

TGCTCACTCC TCCATCTCTA ATCAGGGCTT CGTACCGCTG CACGAGGACG ACGTGGTCTC

CATCTATCGC GACTGCATGT CCGAGTCTTC TTTCGTCTAA ACTAGTATCG ATGATAAGCT

GTCAAACATG AGCAGATCTG AGCCCGCCTA ATGAGC

**ADH14 G1 (Accession No. E8LLW8_9GAMM)**

CGAGCGCCCG TAACTTCACG CGCCAACGTC GTTGTGTTCT GGCCGGCTAA TCTAGAAGTA

TATTTCCCGC TCAATATAAG GAGGAGTACA TATGGAATCT TTCGATTTTT TCCGTCGCAC

TCGTATCATC TTTGGCCAGT CTGCGGACAA CGAAGTAGGT CAGATTATCA AATATCAAGG

TGGCACTCGT GTGCTGCTGC TGCACGGTGA AAAAGCAGCG ATCAAGTACG GTATTGTGGA

GCGTATTGGT CGTACTCTGG ACCGTTCCGG TCTGAAATAC TTCTCCAAAG GCGGCATCAA

GAGCAACCCG CATATTGATA AAGTTTACGA ATGCATTGAA TTCTGCCTGT CCAACTCCAT

TAATTATATC CTGGCTGTGG GTGGTGGTTC CGTGATCGAC ACCGCCAAAA TCGTCGCGGC

GGGCGTATTC TTCGACGGCG ACATCTGGGA CATGTTTGAA AAACATCGCG AACCGTACCG

TTCCCTGCCG CTGGGCTGCG TAGTTACCGT TCCTGCAAGC GGTACTGAAT GCAGCAACTC

TTCTTCCCTG ATGCGTGAAA AAGACGGCCG CCGTGAAAAA CTGATCGCGT ATTCTAACAG

CTTCGTACCG GAGTTCGCCA TTCTGAACCC GGACCTGACG CTGTCTCTGT CTCCGCGTGT

GACCGCTAGC GGTTGCGTTG ATATGATTAA CCATG

**ADH14 G2 (Accession No. E8LLW8_9GAMM)**

CTCTGTCTCC GCGTGTGACC GCTAGCGGTT GCGTTGATAT GATTAACCAT GTCCTGGAAG

GTTATTTCTC CAACTCTACC GGTGTACTGC TGAGCGATAA GCTGTGTGAA GCGGTTCTGA

GCTCTATTAT CGAACTGCTG CCGCAGATCT ATGAAGATCC GAATAACATT GATGCGCGCG

CAAACCTGAT GCTGGCAGCA ACCCTGTCTC ACAATGATAT CTGCTGCATG GGCCGCAAGT

CCGACAACGT TATCACGAAA CTGGCCAACC AGCTGGTGGT TGAAAACGAT TGTCCGTTCG

GTGATGCACT GGCTGTTCTG ATCCCGGCTT GGATGGAATA TGTTGTTCAG TTTAACCCGC

TGCGCATCGC ACAATTCTCC AACCGCGTTT TTGGTATCGC AATCAACTTC GAAGATCCGA

AAATTACCGC GTATGACGGT ATCAAAGCCC TGCGCGCTTT TTTCAAAAAT GTAAAACTGC

CGTGCAACTT CGTTGAACTG GGTATCAAGA CCGAAGCAAT CGCGGACATC GTAAACGCTC

TGGACCTGAA AGAAGGTAAA ACTCTGGGTT CTTTTGTGCC GCTGGACGCT GTGGCCTGCG

AAGCAATCCT GTCCCTGGCC GCCAATTACT GCGAAGGTCG CGATATTTTC TAAACTAGTA

TCGATGATAA GCTGTCAAAC ATGAGCAGAT CTGAGCCCGC CTAATGAGC

**DhaT2 G1 (Accession No. E4RKV2_HALSL)**

CGAGCGCCCG TAACTTCACG CGCCAACGTC GTTGTGTTCT GGCCGGCTAA TCTAGATCCT

TACCGCTTAC CACGAAATAA AGAGGGAAAA ATGTCCGACT ACTATGATTA TATGCTGCCG

ACCGTTAACT TCATGGGCCC GGGCTGTGTG GAGGTTGTGG GCGAACGTTG CAAGATCCTG

GGTGCTAAAA AAGTTCTGAT TGTTACCGAT TCTTTCCTGC GTAACATGGA AGGCGGCCCG

GTCGACCAGG TGGTCAAATA CCTGAAAAAG GCGAATCTGA ATTACGCTTT TTACGACGAG

GTTGAACCGA ACCCGAAAGA CGTTAACGTT TACGCCGGTC TGAAAATTTA CGAACGTGAA

AACTGCGATA TGATCGTTAC CATTGGTGGT GGTTCCGCTC ACGATTGTGG TAAAGCTATT

GGTGTTGCTG CTACTCATGA CGGTGATCTG TACAAAGACT ACGCTGGCAT CGAAAAGCTG

GAAAACGAAA CTCCGCCGAT GGTATGTGTG AACACCACTG CTGGCACCGC TTCTGAAGTT

ACTCGTCACA CCGTGATTAC TGACACCAGC CAGACGCCTA ACGTCAAGTT CGTGATTGTA

TCCTGGCGTA ACACTCCAGA TGTGTCTATC AACGATCCGG AACTGATGGT TGGTAAGCCG

CCGGGTCTGA CCGCAGCTAC TGGCATGGAC GCTCTGACTC ACGCGGTTGA AACCTATGTG

AGCACCAATG CGAAC

**DhaT2 G2 (Accession No. E4RKV2_HALSL)**

CATGGACGCT CTGACTCACG CGGTTGAAAC CTATGTGAGC ACCAATGCGA ACGCGCTGAC

TGATGCCGCG GCTATTAAGT CTATTGAACT GGTTGCTAAT AACCTGCGTA AAGTTGTTAA

AGATGGCCAG GACATCAAAG CGCGCGAAAA TATGGCGAAC GCTTCCGTTC TGTCTGGTTT

CGCGTTTAAT AACGGTGGTC TGGGTTACGT TCACGCTATG GCACACCAGC TGGGCGGCTT

CTACGACATG CCTCACGGTA TCGCAAATGC AATTCTGCTG CCGTACGTCG AGAAATTCAA

CCTGGGTACC GACGTTGAAC GTTTCAGCAA CATCACTGAA ATCTTCGGCA AAGAACAGTC

TAAAATCAGC AATAATCCGG AAGCACAGGA AAGCATCAAA GCCATCAAAG ACGAAATCGA

TAAACTGAAA CGCTTTAAAA AAATTGCAGA AGTGTTTGGC GTGGATACCT CTAACATGTC

TACCCGCGAA GCTGCAGAGG CTAGCCTGGA TGCCATCAAA GAACTGGCCC GTGATATCGG

TATTCCGAGC TCTCTGTCTG AGTCTAAATT CGACGTTAAA CGTGACGATT TCGAGGAAAT

GGCTAAACTG GCGCTGGAAG ATGGTAACGC GGGCACCAAC CCGCGTAAAG GTTCTGTAGA

AGATATTGTT CGTATCTTCG AGGATGCGTT CTAAACTAGT ATCGATGATA AGCTGTCAAA

CATGAGCAGA TCTGAGCCCG CCTAATGAGC

**DhaT3 G1 (Accession No. Q15G22_CITFR)**

CGAGCGCCCG TAACTTCACG CGCCAACGTC GTTGTGTTCT GGCCGGCTAA TCTAGATAAT

AGCAATACTA ATAAAGAGGT CTCCTAATGA GCTATCGCAT GTTCGACTAC CTGGTCCCGA

ACGTGAACTT CTTCGGTCCA AACGCAATCA GCGTGGTGGG TGAACGCTGC CAGCTGCTGG

GTGGTAACAA AGCTCTGCTG GTGACCGACA AGGGTCTGCG CGCTATCAAA GACGGTGCGG

TAGACAAGAC CCTGCATTAC CTGCGTGAGG CGGGTATCGA GGTTGCTATC TTCGATGGCG

TTGAACCTAA CCCGAAGGAT ACTAACGTGC GCGACGGTCT GGCAGTTTTC CGCCGTGAAC

AATGCGATAT TATTGTGACG GTTGGTGGCG GTTCCCCGCA TGATTGCGGC AAAGGTATCG

GCATCGCTGC TACGCACGAA GGTGCGATCT GCACTTCTAT GCCGGAATCC CGTCCGTGGA

CCAACCCGCT GCCTCCGATC GTGGCCGTAA ACACCACTGC AGGTACCGCC AGCGAAGTGA

CCCGTCACTG TGTTCTGACC AACACCGAGA GCAAAGTAAA GTTCGTTATT GTCTCCTGGC

GTAACCTGCC GTCTGTGTCC ATCAATGACC CGCTGCTGAT GATCGGTAAA CCGGCTGCCC

TGACGGCAGC CACCG

**DhaT3 G2 (Accession No. Q15G22_CITFR)**

TGACCCGCTG CTGATGATCG GTAAACCGGC TGCCCTGACG GCAGCCACCG GCATGGACGC

TCTGACCCAC GCAGTGGAGG CATACATCTC CAAGGATGCC TCCCCGGTTA CTGATGCAGC

GGCTATGCAG GCAATCCGTC TGATCGCCCG TAACCTGCGC CAGGCCGTTG CTCTGGGCAG

CAACCTGCAG GCTCGTGAAA ACATGGCGTA CGCTAGCCTG CTGGCTGGTA TGGCATTTAA

TAATGCGAAC CTGGGTTACG TGCACGCAAT GGCCCACCAG CTGGGTGGTC TGTACGACAT

GCCACATGGC GTGGCGAACG CAGTTCTGCT GCCGCACGTT GCCCGCTACA ACCTGATCGC

GAATCCGGAA AAATTCGCGG ACATCGCGGA ACTGATGGGC GAAAATATCA CCGGTCTGTC

TACTCTGGAC GCGGCAGAGA AAGCTATCGC AGCGATCACC CGTCTGAGCA TGGATATTGG

TATCCCGCAA CATCTGCGTG ACCTGGGTGT CAAAGAAGCG GACTTCCCGT ACATGGCGGA

AATGGCACTG AAAGACGGTA ATGCGTTCAG CAACCCGCGT AAAGGTAACG AGCAAGAGAT

TGCGGCTATC TTTCGTCAAG CTTTTTAAAC TAGTATCGAT GATAAGCTGT CAAACATGAG

CAGATCTGAG CCCGCCTAAT GAGC

**DhaT4 G1 (Accession No. A0PY50_CLONN)**

CGAGCGCCCG TAACTTCACG CGCCAACGTC GTTGTGTTCT GGCCGGCTAA TCTAGAATAC

GGAAACCTCA CGTATAACCA AGGAGGTTAC AATGCGCATG TATGACTACC TGCTGCCTAA

CGTTAACTTC ATGGGCCCGG GCTGCATCAA AGTTATCGGC GAACGTTGCA AACTGCTGGG

CGCTAAAAAA GCGTTCATCG TAACCGGTAA ACACATTGGT AGCATGGAAA ACGGTCCGCT

GCAGATCGTT GTAAAATACC TGACCGATGA AGGTATCGAC TACGTTCACT TTAGCGGCTC

TGAACCGAAT CCGAAAGACA TTAACGTGCG TAAAGGTGTT GAACTGTTCA AAAAAGAAAA

CTGCGATATG ATCATTACCA TCGGCGGTGG TTCTGCACAT GACTGCGGTA AAGGTATTGG

CATCGGCGCG ACGCACGAAG GTGACCTGTA CGACTACGCA GGCATCGAGA CTCTGACCAA

CCCACTGCCG CCTATTGTAG CGGTGAACAC CACCGCGGGC ACGGGCTCCG AAGTTACCCG

TCACTGTGTG CTGACGAACA CCGAGAAAAA AATCAAATTC GTAATCGTGT CTTGGCGTAA

TCTGCCGCAG GTGAGCATCA ACGATCCGCT GCTGATGGTG GATATGAGCC CGCGTCTGAC

TGCGGCGACC GGTAT

**DhaT4 G2 (Accession No. A0PY50_CLONN)**

TCCGCTGCTG ATGGTGGATA TGAGCCCGCG TCTGACTGCG GCGACCGGTA TGGATGCACT

GACTCACGCG ATTGAGGCCT ACGTGAGCAA AGACGCAAAC GTGGTTACGG ACGCGGCGGC

AATTCAAGCG ATCAAACTGA TCAGCAAAAA CCTGCGTAAA GCCGTCGCAC TGGGCGAAAA

CCTGGAGGCC CGTGACAACA TGGCAAACGC TTCTCTGCTG GCAGGCATGG CCTTTAACAA

TGCGAACCTG GGTTATGTTC ATGCGATGGC TCATCAGCTG GGCGGTCAGT ACGACCTGGC

CCACGGTGTT GCAAACGCCA TGCTGCTGCC GCATGTCGAA CGCTACAACA TTATCTCTAA

CCCAGAAAAA TTCCGTGACA TCGCTGAGTT CATGGGTGAG AATATCGAGG GTCTGTCTGT

CATGGAGGCT GCAGAAAAAG CAATCGATGC GATGTTCAAA CTGGCCGAAG ACATTGGCAT

TCCGCGTCGC CTGCGTGACG TAGGTGTCAA GGAAGAAGAT TTCGAATATA TGGCGGGCAA

CGCGCTGAAA GATGGTAACG CATTTTCTAA CCCTCGTAAG GGCACTGAAG AAGATATTGT

CAACATTTTT AAAGCTGCAT ACTAAACTAG TATCGATGAT AAGCTGTCAA ACATGAGCAG

ATCTGAGCCC GCCTAATGAG C

**DhaT5 G1 (Accession No. Q3A1K9_PELCD)**

CGAGCGCCCG TAACTTCACG CGCCAACGTC GTTGTGTTCT GGCCGGCTAA TCTAGAAATA

TTATCTCTAA TCATAAAAGG AGGTTCAAAC TATGGCGGAT TTCCTGTGCC CTGCTGTTAA

CTTTATTGGC GCGGGTACCG TAGCAGAAAC TGGCCCGCGT GCTGCGATGT TCGGTACGAA

AGCTCTGATC GTGTGCGACG GCTTCCTGGC CAAACTGGAG GGCGGCCCTG TAAGCAAAGT

GAAAGATACG CTGACTTCCT CTGGCGTTGA GTTCGCAGTT TATGACGGCG TGGAACCGAA

TCCAAAAGAT ACGAACGTAG CAGCCGGCCT GAAAATTTAT AAATCTGAAA ACTGTGACCT

GATCGTAACC GTTGGTGGCG GCTCCTCTCA TGATTGCGGT AAAGGCATCG GCATCGCAGC

TACTCACGAG GGCGACCTGT ACGAAGACTA CGCCGGTATC GAAACCCTGA CTAACGAACT

GCCACCGATC ATCGCGGTAA ACACCACTGC TGGCACTGCG AGCGAGGTTA CGCGTCACTG

CGTGATCACC AACACGGCGA AAAAGGTTAA GTTCGTTATC GTTTCTTGGC GTAACCTGCC

AAAAGTGTCT ATCAATGACC CGGAGCTGAT GGTTGCAAAA CCGGCTGGTC TGACCGCCGC

GACCGGTATG GACGC

**DhaT5 G2 (Accession No. Q3A1K9_PELCD)**

GCTGATGGTT GCAAAACCGG CTGGTCTGAC CGCCGCGACC GGTATGGACG CACTGACCCA

CGCAGTTGAG TGTTACGTAA CTAAGGACGC GAACCCGGCG ACTGATGCTG TGGCTATCCA

CGCTATCAAA CTGATCGGCA AGTACCTGCG TCGCGCAGTG GCAAACGGTG AAGACCTGGA

AGCGCGCGAA GGTATGGCTT ACGGCTCTCT GCTGGCTGGT ATGGCTTTCA ACAACGCGGG

CCTGGGCTAT GTTCACGCTA TGGCGCACCA GCTGGGCGGT CTGCTGGATA TGCCGCACGG

CATCGCAAAC GCGGTGCTGC TGCCGCATAT CGAACGTTAC AACCTGATGG TGAACCCGGA

AAAATTCGCC GATATTGCGG AGGCGATGGG TGAAAACATC GATGGTCTGG GTAAAATGGA

AGCTGCTGAA AAGGCTATCG ATGCTATCGT TCGCCTGTCC ATTGATGTTG GTATTCCGCA

GCATCTGGCT GACCTGGGCG TTAAAGAATC CGATCTGGAG CCGATGGCGA AACTGGCCAT

GCAAGATGGC AACGCAGGTA CTAACCCGCG TGTTGGCAAA GTTGAAGATA TCATCCAACT

GTTCAAAAAC GCAATGTAAA CTAGTATCGA TGATAAGCTG TCAAACATGA GCAGATCTGA

GCCCGCCTAA TGAGC

**DhaT6 G1 (Accession No. A5D4X5_PELTS)**

CGAGCGCCCG TAACTTCACG CGCCAACGTC GTTGTGTTCT GGCCGGCTAA TCTAGATTCC

ATTTTTCTTC CACCAAAAGG AGTTACTTCC ATGTATTACG ATTTCCTGAA CCCGTCCGTT

AACTTTTTTG GCCCGGGTTG TGTCTCCGTT GTAGGCGAGC GTTGCAAGAT CCTGGGCGGC

AAAAAAGCTC TGATTGTTTG CGATCCGTTT CTGGCAAAGA TGGAAGGCGG TCCGGTGGAA

CAGGTACTGG GTTACGTTCA GGAAGCGGGC CTGAAAACCG TGGTCTTTGA CGGTGTGGAA

CCAAACCCAA AAGACAAAAA TGTACACGCA GGCCTGAAAG TGTTCAAAGA AGAAAAATGC

GATATGATCA TTACTGTTGG TGGCGGTAGC GCGCACGATT GTGGCAAAGG TATTGGTATC

GCGGCTACCC ACCCGGGTGA TCTGTACAAA GACTACGCGG GTATTGAAAA ACTGACCAAT

CCGCTGCCAC CGATTGTTGC AGTGAACACC ACTGCTGGTA CCGGCTCCGA AGTTACTCGT

CACTGCGTTC TGACTAACAC CTCCACTTCC ATCAAATTCG TCATTGTCTC TTGGCGCAAC

CTGCCACTGG TTTCCATCAA CGATCCGATG CTGATGCTGA AAAAGCCGGC GGGTCTGACT

GCGGCCACTG GCATG

**DhaT6 G2 (Accession No. A5D4X5_PELTS)**

TCCGATGCTG ATGCTGAAAA AGCCGGCGGG TCTGACTGCG GCCACTGGCA TGGATGCTCT

GACGCACGCA GTCGAATGCT ATGTTACCAA AGCTGCAAAC CCGGTGACGG ACGCACTGTG

TGCGCAATCC ATCAAACTGA TCGCGAATAA CCTGCGTCAG GCGGTGGCGA ACGGTGAAAA

CCTGACCGCT CGTGAAAACA TGGCATATGC TAGCATCCTG GCGGGCATGG CCTTCAACAA

CGCCGGTCTG GGCTATGTTC ATGCAATGGC TCACCAACTG GGTGGCTACT ATGACATGGC

GCACGGTGTG GCGAACGCTA TCCTGCTGCC GCACGTGGCT CGCTTTAACC TGATCAGCAA

CCCACAAAAA TTCGCGGATA TCGCAGTGTT TATGGGTGAA AACATCGAAG GTCTGTCCGT

ACGTGCGGCG GCAGAAAAAG CGATTGACGC CATTGTTCAG CTGTCCAAGG ATGTGGGCAT

CCCATCTGGC CTGGCTGAGA TGGGTGTAAA AGAGGAAGAT TTTGACAAAA TGGCGAAAAC

GGCACTGGAG GATGGTAACG CTGGCTGCAA CCCGATTGTG GGTACCCACC AGGATATTGT

GAAGCTGTTC GCGGCTGCTA TGTAAACTAG TATCGATGAT AAGCTGTCAA ACATGAGCAG

ATCTGAGCCC GCCTAATGAG C

**DhaT7 G1 (Accession No. B1V2D9_CLOPF)**

CGAGCGCCCG TAACTTCACG CGCCAACGTC GTTGTGTTCT GGCCGGCTAA TCTAGAAGCC

TAGGCTAAGA AGCATAAGGA CCCCACTATG GAAGAGAACG AAATGCGTAT GTATGACTAC

CTGGTTCCGA GCGTGAACTT CATGGGTGCT AACAGCATTT CCGTTGTAGG CGAGCGTTGC

AAAATCCTGG GTGGCAAGAA AGCTCTGATC GTTACCGACA AATTCCTGCG TGGCCTGAAA

GGTGGTGCAG TTGAACTGAC GGAAAAATAC CTGAAGGAGG CGGGCATTGA AGTAGCATAC

TACGACGGCG TAGAACCGAA CCCGAAAGAC ACTAACGTGA AAGATGGTCT GAAGATCTTT

CAGGACGAGA ACTGTGATAT GATTGTAACC GTTGGTGGTG GTTCCTCCCA CGACTGCGGC

AAGGGCATTG GTATCGCGGC GACTCACGAA GGCGATCTGT ACGACTATGC AGGCATTGAA

ACCCTGACCA ACCCGCTGCC TCCGATCGTG GCTGTAAATA CTACCGCCGG TACGGCAAGC

GAAGTTACCC GCCACTGCGT CATCACTAAC ACCAAAACTA AAGTTAAATT CGTGATTGTT

AGCTGGCGTA ACCTGCCGCT GGTCAGCATT AATGATCCGA TGCTGATGGT TGGTAAGCCG

GCGGGCCTGA CCGCA

**DhaT7 G2 (Accession No. B1V2D9_CLOPF)**

CAGCATTAAT GATCCGATGC TGATGGTTGG TAAGCCGGCG GGCCTGACCG CAGCAACTGG

CATGGATGCT CTGACCCACG CAGTTGAAGC TTACGTTTCT AAAGATGCCA ATCCTGTTAC

CGACGCGGCA GCAATTCAGG CAATTAAACT GATTTCCTCC AATCTGCGTC AGGCGGTAGC

CCTGGGCGAA AACCTGGTGG CTCGTGAAAA CATGGCGTAT GGCAGCCTGC TGGCTGGTAT

GGCCTTCAAC AACGCAAACC TGGGCTATGT GCACGCTATG GCGCACCAGC TGGGCGGCCT

GTACGACATG CCACACGGTG TTGCTAACGC GATGCTGCTG CCTCACGTGT GTAAATATAA

TCTGATCAGC AACCCGCAGA AGTTCGCTGA TATCGCCGAA TTTATGGGCG AAAACATCGA

GGGCCTGTCC GTGATGGACG CGGCACAGAA AGCGATCGAC GCGATGTTCC GTCTGTCTAC

GGATATCGGC ATCCCTGCTA AACTGCGCGA TATGGGCGTT AAGGAGGAAG ACTTCGGTTA

CATGGCCGAA ATGGCGCTGA AAGATGGCAA CGCGTTTTCC AATCCGCGCA AAGGCAACGA

ACGCGATATT GTTGAAATTT TCAAAGCAGC TTTCTAAACT AGTATCGATG ATAAGCTGTC

AAACATGAGC AGATCTGAGC CCGCCTAATG AGC

**DhaT8 G1 (Accession No. E3H9G9_ILYPC)**

CGAGCGCCCG TAACTTCACG CGCCAACGTC GTTGTGTTCT GGCCGGCTAA TCTAGAAGCC

TAACTTAAAT TTAAAAGGAG GCCCTTATGC GTTACTACGA CTATCTGATG CCGTCTGTGA

ACTTTTTCGG TCCGGGTTGC CTGGAAGTGA TCGGCGAACG TGCAAAAATC CTGAATGGCA

CTAAAGCTCT GATCGTTACG GACAAATTCC TGTCTTCCCT GAAAGGCGGT GCTGTTGAAA

AAACCCTGGA GTATCTGAAA TCCGCTGGCG TTGAAGCCGT TGTATTTGAT AACGTGGAAC

CGAACCCAAA GGATACCAAC GTTTACGAGG GTGCAAAAGT ATATAAGGAA AACAATTGCG

ATATGATCAT CACCGTGGGC GGTGGTAGCC CACACGACTG CGGTAAGGGC ATCGGTATTG

CAGCGACCCA TGACGGCGAC ATCTGTGATT ATGCCGGTAT TGAAACCCTG ACCAACGCGC

TGCCGCCGAT CATCGCTGTT AACACCACCG CTGGCACTGC TTCTGAAGTT ACTCGTCACG

CCGTCATTAC TAACACTAAG ACCAAAGTTA AATTTGTGAT CGTGAGCTGG CGTAACCTGC

CACAAGTTAG CATCAACGAC CCGCTGCTGA TGATCGGTAA ACCGGCTGGT CTGACCGCAG

CTACCGGTAT GGACG

**DhaT8 G2 (Accession No. E3H9G9_ILYPC)**

GCTGCTGATG ATCGGTAAAC CGGCTGGTCT GACCGCAGCT ACCGGTATGG ACGCTCTGAC

CCATGCTGTG GAAGCTTACA TTTCCAAGGA CGCGAACCCA ATTACTGATG CTGCGGCAAT

CCAAGCGATC AAACTGATCG CCCAGAACCT GCGTCTGGCC GTGGCCAACG GCGAGAACCT

GAAAGCGCGT GAAAACATGG CGTACGCTTC TGTACTGGCC GGTATGGCAT TTAACAACGG

CAACCTGGGT TACGTTCACG CGATGGCGCA CCAGCTGGGC GGCCTGTATG ATATGCCGCA

CGGCATCGCA AACGCAATGC TGCTGCCGCA CGTGTGTAAA TATAACATGA TCTCTAATCT

GGATAAGTTT GCAGATATCG CAGAATTTAT GGGTGAAAAC GTTGATGGCC TGTCTAAATC

TGAAGCGGCC GAAAAAGCGA TCTCTGCGAT GTTCCGTCTG TCTTCCGATC TGGGCATTCC

GACCTCTCTG GAAGAAGCAG GCATCAAAGA ATCCGACATT GAACTGATGG CTGAGAATGC

CCTGAAGGAT GGTAACGCGT TCTCTAACCC ACGTAAAGGC AACGAGAAAG ATGTTGAAAA

CATTTTTAAA GCGGCGATGT AAACTAGTAT CGATGATAAG CTGTCAAACA TGAGCAGATC

TGAGCCCGCC TAATGAGC

**ADH15 (Accession No. Q1JYE4_DESAC)**

GATACAGGTC TCGTCTAGAA GAGATCATAA TAGAGAGGAT ACC**ATG**CAGC CGTTCGTGTT

TCATAACCCT ACCGAAATCG TTTTCGGCGT TGACACTGCT GATAAAGTAG GCAAATACGC

CGCACGTCAG GGTGGTAAAG CCCTGCTGGT TTATGGCCGC AACTCTATCA AAACCACTGG

CCTGTATGAT CGCGTAACTG CTAGCCTGCA GGCCGCCGGC CTGAGCTGGG TTGACCACGG

CGGTGTCAAA TCCAACCCGG TTCTGAGCCA CGTGCGTGAA GGCGTTGCAG TTGCAAAACG

TGAACAAGTA GACGTTGTTG TCGCAGTAGG CGGCGGTAGC GTTCTGGATG AGTCTAAAGC

GATCGCTGCA GGCGCGCTGT GCGATCACGA TGTTTGGGAG TTCTTCCTGC AGGCTAAAGT

CGAGAAAGCG CTGCCGCTGG TTACTGTTCT GACTCTGGCT GCAACGGGTT CCGAAATGAA

CAGCGGCGGC GTTGTAACCA ACGAGAACAC CGCTCAGAAA TTCAACATCG GCTCCCCGCT

GCTGTTCCCG AAAACGTCTA TTCTGGACCC GGCACTGACC TATACTGTTC CGGCTGATTA

CACCGCATAT AGCGCCGTGG ATGCGATCTC TCACATCATC GAAGGTTATT TCACCTCCAA

CGATCAGGCA ACTCCTCTGC AGGACCGTTT CGTAGAAGGC CTGGTGAAAA CCATCATGGA

GTCCACCGAG CAGATTCTGC AGCAACCGGA TCACGCGGAC GCGCGTGCTA CCATGATGTG

GTCTGCCACT TGGGCTCTGA ACGGCCTGTC CACTGCTGGT ATTGGTCTGT ATCAATTCCC

GAACCACATG ATCGAACACT CTCTGTCCGC TATGTACGAC ATCGCCCATG GTGCCGGTCT

GTCCATTGTA ATCCCAGGTT GGATGGATTA TGCGGCGACC CAAAACCCGG CTAAATTCGC

TCAGTTTGCA CGTCGTGTTT TCGACTGTGA ACTGAGCGAT GACCTGGAAT GCGCTCGCTA

CGGCATCGAA GCGCTGAAAA CGTGGTTTCA TTCTATCGGC TCTCCGGTGT CCCTGGCTCA

GGGTAACATT CCTGATGAAG AGATCGGCGC TATCGCAGAT AACGCTGTAA TGCTGGCCCG

CAAATGGGGT CTGAAAGCGT ACACCGCCGA AGTTATTGCC GACATCCTGA GCCGCTGTCG

TAGCTAAACT AGTAGAGACC GATACA

**ADH16 (Accession No. B5YIE2_THEYD)**

ATGGAAAACT TTGAGTTCTA CAACCCGACG CGCATTATCT TTGGCAAAGA AGCCGAAAAG

AAGATCGGCA AAATCCTGGA GAAAGACGAC GTTGAACGTG TGCTGTTCGT GTATGGTAAA

AGCTCTATCA AAGAGACAGG CCTGTATGAT CGCATTGTTA AGGCTCTGAA AAAAGAAGGC

ATCGAATTTA TTGAGCACGG TGGTGTGAAG CCGAATCCGG TGCTGTCCCA CACCCGTGAA

GGTATTGAAA AAGCGAAGAA GCACAAGGTA GACGCGATTC TGGCTGTAGG CGGCGGTAGC

GTTATCGACG AAGGCAAAAC TATTGCAGTC GGTACCAAAA CCGACAAAGA CGTTTGGGAT

TTCTTCAAAC GTAACAAAGA AATTAAAAAA GCACTGCCGA TTTATGTGAT CCTGACTCTG

GCGGCAACCG GTTCTGAAAT GAACGGTTTC GCTGTAATCA CCAACGAAGA AACCCAGGAA

AAGCTGAGCA TCTCTTCTGA ACACATCTTC CCGCGTGTTT CTATCCTGAA CCCGGAACTG

ACCTTCACCG TATCCGCTAA ATATCAAGCG TACGCGGCAG TAGACGCAAT CGCGCACGTT

ATCGAACACT ATTTCTCCGG TTCTTACTGC CCGAACCTGC AGAACCGTTT CGTTGAGGGT

CTGATCAAAA CCGTAATGGA AACCACCGAA ATCATTCTGA AAGAACAGAA GAACTACAAC

GCTCGTGCAG AATTTATGTG GGCGGCTACC CTGGCCCTGA ACGGCCTGGC AAAGCTGGGC

ATCAAAGGCG GCTCCTTCCC GAACCACATG ATTGCACATT CCCTGGGTGC TATCTACGAT

CTGCCACATG GTGCTTGTCT GAGCATCGTA ATTCCGGCGT GGATGAAATG GTACCAAGAA

AAGAACCTGA TTCAGTTCGA ACGCTTCGCC AAAGAAATCT TCGGCGTCAA CACTGCTACC

GAGGGTGTGT TTCAACTGAA AGAATGGTTT CGCAAAATCG GTGCGCCGGT TTCTCTGAAA

GAGGCAGGCA TCTCCATTGG CGAGATCGAT CGTATTGTAG ACAATGCGTA TAACATCGCG

AAAGTCTGGC AGATGGAAAA AGACTACACT AAGGAGGTAC TGACCGAAAT TATCAAAAAC

GCTAACGATT AA

**ADH17 (Accession No. D2BSS7_DICD5)**

GATACAGGTC TCGTCTAGAT AGCCAGATTA AGGAGCAGCA AAAATGGACA ATTTTAGCTT

TTATAACCCG ACCCGTATCG AATTCGGTGC TGGCAAAGAA CAGCTGATCG GTGAAATCAT

GGCTTCTCAC GGCATTAAGA AAGTCCTGCT GTCCTATGGT AGCGAACGCA TTAAAGATAA

CGGCCTGTTC GTGACCGTTT CCGACTCTCT GAACCGTCAC GGCATCACTC TGATCGAATG

CGGTGGTATC ATCTCTAATC CGCTGATTTC CAAAGTACGT GATGCGATCA CCGTAGCTAA

AGAGCAGAAT GTTGACGCCA TCCTGTCTGT TGGTGGTGGT TCTGTGCTGG ATTCCGCAAA

AGCAATCGCA GCCGGTTCCC TGTATCAGGG TGACGTGTGG GACCTGTTCA TCGGTAAGGG

CCAGATTGAC GCAGCACTGC CAGTTTTCGC CATCCTGACT CTGGCGGCGA CTGGTTCTGA

AATGAACTCT GGTGCGGTGG TGACTAACGA CGACACCAAA GAAAAATTCG CAATCAACAG

CGTACACATC TTCCCGAAAG TGTCCATTGT AAACCCAGCT CTGATGCAAA CCGTTTCCCG

TGACTACCTG GTGTATTCTG CTGCAGACAT CATTGCACAT TCTATTGAAG GTTACTTCAC

TGCGACCATT CAGCCGAAAA TTCAGTCCCG TCTGGTTGAA TCTGTCATTG CTACCGTTCT

GGAAACTACC GAACAACTGC TGGCTGACAG CGCAGACTAT AACGCTCGCG CAGAATTCGC

TTGGGCTGCG ACCCTGGCGC TGAACGGTCT GACCTATGCT GGCACCTCCG GTTTCGGTTA

CCCTAACCAC ATGATTGAAC ACGCGCTGTC TGCACTGTTC AACGTACCGC ATGGTGCTGG

CCTGAGCGTT ATCATGCCTG CATGGATGAA GTGGTTCCAC TCCCAGAACA CTGCTCAGTT

CGAACGCTTC GCCCAGCACC TGTTCGGCCT GAACACCGCG GAAGAGGGCA TCGCTGCTCT

GGAGAAATGG TTCGATAAAA TTGGTACCCC GACTCGTCTG AGCCAGCTGG GCATCAAAGC

CACTGACCTG CCGGCTATCC TGGACAACCT GGAAGGTAAC GCGCGTTGGT TCGGTCTGGC

AGAAACTTAT ACCGAGGAAG TACTGGCGAC CATCCTGCGT CTGGCACTGT AAACTAGTAG

AGACCGATAC A

**ADH18 (Accession No. F0ERB1_HAEPA)**

GATACAGGTC TCGTCTAGAC ACGACAAGAC AGGGGGCCAA TCCATGTATC CATTCTCCTT

TCAGAATCCA ACCCGTATCG AGTTCGGTCT GGATAAAGAA AAGGAAATGG GTAAATACAT

GCACGAATAC GGCGCGAAGA AAGCGCTGAT CATCTATGGC AGCGAGCGTG TGAAACAATC

CGGCCTGTTC GAAGACGTTA CCAAGTCTCT GCGTGAACAC GGCATCGAGT ACATCGAATG

TGGCGGCGTT AAAAGCAACC CGACCATCAG CAAAGTCCGT GAAGCTGTTG CAATGGCCAA

AGCATTCGGC GCAGATAGCG TGCTGTCTAT TGGTGGTGGT AGCTGCCTGG ACAGCGCGAA

AGCGATCGCA GCCGGTGCGT GCTACGATGG CGATACTTGG GACTTCTTCA AGGGCACCCT

GGTTCAAAAG GCGCTGATGA TCTTCGATGT AATCACCCTG GCGGCTACGG GTTCCGAAAT

GAACTGGGGT AGCGTGATCA CCAACGAGGA AACCCAACAG AAATACTCCA TTCACAACAA

CCATCTGTTC CCTAAAGTTA GCGTAATCAA CCCGAAGCTG CAGGCGACTG TGTCTCGTGA

TTACCTGGTT TATTCTGCTG CCGATATCAT CGCACACTCT ATCGAAGCGT ATTTCACCGC

AGAATACCGC CCGGAGATCA TCGACTTTCT GGTAGAGAGC AACATCAAAA CTGTCATCCG

TACCACGGAA ATCCTGCTGA ACGACCCGCA GGATCTGAAC GCGCGCGGCG AATTTGCATG

GGCCGCTACG CTGGCACTGA ATGGCCTGAC CCACCTGGGT ATCAGCCCGT ACGGCTTTCC

GAACCACATG ATCGAACACA GCATGTCTGC GATTAGCGAC GTTCCGCACG GTGCAGGTCT

GAGCGTGATC ATGCCGGCGT GGATGCAGTG GTATCAGAGC CAGCGTCCGG CGCAGTTCAA

ACGTTTTGCC AAAGAAATCT TCGGTCTGGA GAACGCGGAA GAGGGTATCC AGGCGCTGAA

AACTTGGTTC GACAAAATTG GTACTCCAAC GCGTCTGGAA CAGCTGGGTA TTGACGACAA

AACCCTGTTT GAAATTGTTG ATAATGCAGT TCAGACTGCC ATCCGTGCCA AAGTTGAAAA

AACCTACACC AAAGAGGCGA TTAAAGCGAT CTTCGCCCTG GCAAAATAAA CTAGTAGAGA

CCGATACA

**ADH19 (Accession No. G5IQ05_9ENTE)**

GATACAGGTC TCGTCTAGAA TAAGAGCATA CAGGGGGCCT ATTATGCAGA ACTTCCGTTT

TTACGTTCCG ACCGACATTC GTTTTGGCAC GGACCGTCTG AGCGAACTGC CAGAAGCGCT

GTCCAGCATC GGCAAACGCG TTCTGCTGGT ATACGGCGGC GGCTCTATCA AAAAGAGCGG

TCTGTATGAT AAAGTCCAGC AGCAGCTGGT TAAGGGCGGT TTCGAAGTCG TAGAACTGAG

CGGTATCGAA CCGAATCCGA AAATTCACAG CGTTCGTTCC GGTGTTAAAC TGATCCGCAA

ACACCAGCTG GAAGTAATCC TGGCGATCGG TGGTGGCAGC GTGATCGATA CTGCTAAAGT

AATCGCGGCT GGCGTATTCT ATGAAGGCGA CCCGTGGGAC CTGGTTGTCG ACAGCAGCAA

AATCAAACAA GCTCTGCCGA TCGTTGACAT CCTGACCCTG GCGGCAACCG GCACCGAAAT

GAACCGCAAT GCGGTGATTT CTAACGCTGA TACCAAAGAA AAACTGGGTA CCGGCGGCGC

TGAACTGCTG CCGCACGTTT CTTTTCTGGA CCCTACCACC ACTTTCTCCG TTTCCAAATG

GCAGACCGCG GCTGGTGCGG CAGATATCCT GTCCCACCTG TTCGAACAGT ACTTCAACCG

TACCAAAGCT GTCCAAGTTC AAGACAACAT TGCGGAAGCC CTGATGAAGG TAGTAATCGA

TTTCGCTCCG GTGGCCTTTG AAAATCCTGA CGACTATGGC GCACGTGCGA ACCTGCTGTG

GGCTAGCACC CTGGCTCTGA ACGGCCTGGT TGGCAACGGT CGCTCCGGTG GCTGGACTTG

CCACCCGATC GAACATGAGC TGTCTGCTTT CTATGATATC ACTCATGGCA TTGGTCTGGC

TATCCTGACC CCGCGCTGGA TGGCTTACTG TATCACCCAC GACCCAAGCA CCCATGAGAA

ATTCGCTAAG TATGGTGAAG CTGTGTGGGG CCTGACGGGT GACTCCCAGG CAGAAATCGC

TCGTAAGGCG GTACGTACTA CTTACGAATT TTTCGAAGAC CAGCTGGAAA TTCCGATGAC

CCTGCAGAAA GTGGGTATCG AAACGACGGA ACTGGTGGAC GAAATGTCTC AGCAGGCGGT

GGTGCATGGC GAGCTGAACA CTGACGGCCC GTTCGTTCGC CTGGACGAAG AAGCGGTTAA

AAGCATCATT ACCAGCTGTT TCGAAGAAAT GACTCTGAAC TAAACTAGTA GAGACCGATA

CA

**ADH20 (Accession No. B2V5D0_CLOBA)**

GATACAGGTC TCGTCTAGAC ATAAAGCTCA GAGGACATCA CACATGGAAA ATTTCAACTA

CTCTATTCCG ACCGAAATCT ATTTCGGTAA GGGTCAGATC AAAAACCTGG GTAATGCCAT

CAAAAAGTAC GGTAGCAAAG TGCTGGTTGT GTATGGCGGC GGTAGCATCA AACGTATCGG

CCTGTACGAC GACATGATGA AAATCCTGAA AGACAACAAC ATCTCTTACG TTGAACTGTC

CAACATCGCA CCAAACCCGC GCATCGAATC TGTGCGTGAC GGCGTGAAAC TGTGCCGTGA

TAACGACGTT GAAGTCGTTC TGGCAGTCGG TGGCGGCAGC ACGATCGACT GCGCTAAAGT

AATCGCAGCA GGTGTAAACT ATGACAAAGA CCCGTGGGAC CTGGTTCTGG ACAGCTCTAA

AATGACCACG GTTCTGCCAG TTATCACTAT TCTGACCCTG AGCGCAACCG GTTCTGAAAT

GGACCCATTC GCAGTGATCT CTGACATGAG CAAAAACGAA AAGGTAGGCG TGGGTAACGA

TAAAATGAAA CCGAAAGTGT CTATCCTGGA CCCGGAATAC ACCTACAGCG TTCCGAAAAA

CCAAACCGCG GCGGGCACGG CTGATATCAT GTCCCACATC TTCGAAAACT ACTTCAACAA

CACGAAAGGC GCCTTTATCC AGGCACGCAC TGCTGAAGGT CTGCTGAAAG CTTGTATGAA

ATATGGCAAG ATCGCCATCG AGGAACCAAA CAACTACGAA GCTCGTGCAA ACCTGATGTG

GGCATCCAGC CTGGCCATTA ACGGCCTGAT CTCTTACGGC ACCGCGGGTG CGTGGAGCGT

CCACCCGATG GAACACGAAC TGTCTGCATT CTACGACATT ACGCATGGTG TCGGTCTGGC

GATCCTGACC CCGCATTGGA TGCGCTACGT CCTGAACGAA GATACTCTGG ACAAATTCGT

TGAATATGGC ATTAACGTGT GGGAGCTGGA CAAAAACCTG GACAAATATA CCATCGCGAA

CACTGCGATC GATAAAACGG CAGAATTTTT CAAGGAAATG GGTATTCCGA GCACTCTGCG

CGAAGTGGGT ATTGGCGAGG AGAACTTCAA AATCATGGCG CAAAAAGCGG TTAAAAGCGG

CCTGGAATAC GGTTTCAAAC CGCTGGCTCC GGAAGATGTG GTTAACATCT ACAAAGCTGC

TCTGTAAACT AGTAGAGACC GATACA

**ADH21 (Accession No. E2SME8_9FIRM)**

GATACAGGTC TCGTCTAGAA ACACACGACG GGGAGGGGAC GCAATGACCG AAGCGATGGA

AAACTTCATT TACGATATCC CGACCAAAGT TTACTTTGGC AAAGGTCAGC TGAACCAGCT

GGCAGATATC GTGCAGGTCT ATGGCCAGCG TGTTCTGCTG GTTTATGGTG GCGGTTCCAT

TCAACGTAAC GGCATTTATG ACGCAGCAGT TGCACAACTG AAAAAGGCGG GTAAGAAGTA

CACCGAACTG TCTGGTGTTG AACCGAATCC GTCCATCCAC ACGGTCGAAA AAGGTGTTAT

GCTGTGCCAG CGTGAACAGA TCGACATGCT GGTAGCTATT GGTGGCGGCA GCGCTATCGA

TTGTGCCAAG GTGATCAGCG CCGCCGCCTG CTCCACCCGT CGTCCGTGGG AACTGGTTAC

GCACCCAGAG GAAATTCAGC GCGCGCTGCC GGTTATCGCC GTTCTGACCA TCGCTGCAAC

CGGTAGCGAA ATGGATCACA TTGCTGTGAT CACGAATCCG CAAACCAAAG AGAAAATTGG

CACCCGTCAC CCGCTGCTGC GTCCGAAAGC CGCTATCCTG GATCCGAGCT TCACTTTCTC

TGTTAACGCT TACCAAAGCG CCTGTGGCGT GGCTGACATC ATGTCTCATA CGATGGAGTC

CTATTTTGCC CGTAAAGAAG CAGCCCTGCA AGATCGTTTC GCAGAGGGTA TTCTGAAGAT

CTGCCTGACC TACGGCCCGA TCGTGCTGCA GCAGCCGGAT AATTACGAGG CGCGTAGCAA

CCTGATGTGG GCAGCTTCTT GGGCTATCAA CGATCTGCTG AAACTGGGTC ACATGACTCA

GTGGTCCGTG CACCCGATGG AACACCCGCT GAGCGCGTTC TACAGCGTTA CTCACGGCGA

AGGTCTGGCG ATTCTGACCC CGCATTGGAT GGATTACGTC CTGTCTGAAG CTACCGTTGG

TAAATTCGCG TGTTTCGCTC GCGAAGTATG GCAGGTTCGT GAAATGGATC CATGGGACAT

GGCTCGTGAG GGCATCGAAC GCCTGCGTGG TTTCTACAAA CAGCTGCAGC TGCGCAGCAG

CCTGGGCGAA CTGGGCATCG ACGAAACCCA CTTTGACGCT ATGGCAGCAG ACGCAGCCAA

ACAAACCGTT AACGGTTACG TTGCACTGCT GGCAGAAGAT GTCAAGAACA TCTACCGTAA

CTCCCTGTAA ACTAGTAGAG ACCGATACA

**ADH22 (Accession No. B0NYL0_9CLOT)**

GATACAGGTC TCGTCTAGAT TAAAATCCTC CAGAGGTCAA TCCATGAATA ACTTCACCTA

CAGCATCCCG ACTAAGATTC ACTTCGGCAA AGGTCAGATC TCCCACCTGT CTGAGCTGTC

CGAGTCTGGC AACAAAGTTC TGCTGTGCTA TGGCGGCGGC AGCATCAAGA AAGCGGGCAT

CTATGACGAA GCCGTGAAAA TCCTGAAAGA AGAAGATATG GAAATCTTCG AACTGTCTGG

TATTGCACCG AACCCGAAAA TCGAAAGCGT ACGTGAGGGT GTGAAACTGT GCAAGGAGAA

CTCCATTGAT ATGGTTCTGG CTATCGGCGG CGGCTCCGTA ATCGATTGTG CTAAAGTCGT

AGCCGCTGGT GCCTGTTATG ACGGCGACCC GTGGGACCTG GTTATTACCC CGCGTTGGAT

CAAAAAGGCA CTGCCGATTT ACTCCGTACT GACTCTGAGC GCGACCGGCT CCGAGATGGA

TAAATTCGCA GTAATCTCTG ACATGTCTAA AAACGAAAAA TGGGGTACTG CGTCTGATCA

CATGAAGCCG AAAATGTCTA TTCTGGACCC GGAATACACG TATTCTGTTA GCAAAAAACA

AACTGCCGCG GGTACCGCGG ATATCATCTC TCATATCTGC GAAAACTACT TCACGAACGT

TAAAAACGCA GATGTGCAGG CTCGCTTCGC GGAAGGTCTG CTGAAAAATT GCTTTAAGTA

TGGCCCGGTT GCCCTGGAAG AACCGGATAA TTATGACGCT CGTGCGAACC TGATGTGGAC

CGCATCTATG GCGATTAACG GTATGATTCA GTACGGCGCT GAAGTGGCGT GGTGTGTACA

TCCGATGGAA CATGAACTGT CCGCGTTCTA TGATATCACC CACGGTGAAG GTCTGGCAAT

TCTGACCCCG CATTGGATGG AGTTTGCGCT GAACGATGAT ACCGCTTATA AATTCGCTGA

TTACGCACGC AACGTTTGGG ACGTTGTTAA CGACGACGAC ATGGCCGCAG CAAAAGAGGG

TATCGCGTAC ACTCGCGAAT ACTTTAAAAA GATGGGTCTG CCGCAGACCC TGACCGACGT

GGGTATCGAT AAAGAATATT TCGACATCAT GGCTCAGAAA GCTGCGGATG GCTGCAAGGG

CAGCTTCGTG CCACTGAGCA AAGAAGACAT CGTTTCCATC TATGAAGCTG CCCTGTAAAC

TAGTAGAGAC CGATACA

*E2. Secondary ADH sequences*

**sADH1 (Accession No. KGK36767.1)**

ATGTTCATGA AAGGTCTGAC GTATCTGAAG CCGGGCATTG TATCCTGGCA AAGCATCCCG

AAACCGGTTC TGAAAAAACC GACTGACGTG ATTGGTAAAG TTGTCACCAC TACCATCTGC

GGCTCCGACC TGCACATTCT GAAAGGTGAT GTTCCTGAGA CCACCGCGCT GGCGGCCACT

ACGGGTCATG GTGTGGTCCT GGGCCACGAA GCCATCATCG AAATTGAATC TGTAGGCGAC

GCTGTGAAAA ACTTTAGCAA AGGCGACGTT TGTATCGTAT CCTGCATCAC CTCTTGTGGT

AAATGCTACT ACTGCAAACG CAACCTGCAG TCTCACTGTA CCGGCCACAT GGAAGGCACT

TCCGGTTGGG TGTTCGGTCA TGAAATCGAT GGCACCCAGG CCGAATACGT GCGTGTTCCG

TGCGCTGATT ATGGCCTGTA CAAGGTTCCT GAAGGCGTTG CGTACGAAAA ACTGCTGATG

CTGTCCGACG CCATTCCAAC CTCCTACGAA ATCGGCATCC TGAATGGCGA GGTGAAGGAA

GGCGACTCCG TTGCTGTTGT CGGCCTGGGT CCGGTAGGTC TGAGCGCACT GCTGACCGCA

ATTAACAAAA AACCGAAACA ACTGATCGCA ATCGACATGG ACGAGAACCG TCTGGACCTG

GCTAAGCAGC TGGGTGCGAC GCATATCATC AACTCCACCA ACATGCCGAA CGAAGAGGTC

GCGAAGAAAG TGCAGGAGAT CAGCAAAGAC CTGGAACCGG GCCGTGAATC CGGCGTGGAC

GTCGCTATTG AATGTGTGGG CGTCCCTCCG ACCTTCGAGC TGTGTGAAGA CCTGATCGCA

TCTGGTGGCA CTATTGCTAA CGTGGGCGTC CACGGTGCGA AAGTAGACCT GAAACTGCAA

GAACTGTGGA TCAAGAACTG TAAGATTACC ACGGGTCTGG TGAGCACCTA CTCCATCCCG

GACCTGCTGA ACCAGGTTGC AGACGGTTCC CTGGACCCTA GCCCAATCAT TACCCATCAC

TTCAAATTTG ACGAATTCGA GAAAGCTTAC CAGGTTTTCA AAGACGCTAA AAATACCAAA

GCGATGAAAA TTATTCTGAC GCCG

**sADH2 (Accession No. WP_011011186.1)**

ATGTCTAAGA ACTACAAAAA GCACCAGGCA TATATCGCAG GTCGCGACAA AATGAAAGTT

GCTGTTATTA CTGGTGCTTC TCGTGGCATC GGCGAAGCTA TTGCTAAAGC CCTGGCGGAA

GATGGCTACT CTCTGGCGCT GGGTGCGCGC TCCGTTGACC GTCTGGAAAA AATCGCAAAA

GAGCTGAGCG AGAAACACGG TGTGGAAGTC TTTTACGATT ATCTGGACGT GTCCAAACCG

GAATCTGTAG AAGAATTTGC ACGCAAAACG CTGGCACACT TCGGTGACGT TGATGTTGTT

GTGGCGAACG CCGGTCTGGG TTATTTCGGT CGCCTGGAAG AGCTGACCGA AGAACAGTTC

CACGAAATGA TCGAAGTTAA CCTGCTGGGC GTTTGGCGTA CCATCAAAGC ATTCCTGAAC

TCCCTGAAAC GTACTGGTGG CGTTGCGATC GTCGTTACCT CCGATGTCTC CGCTCGTCTG

CTGCCGTACG GCGGCGGTTA TGTCGCAACG AAATGGGCAG CTCGCGCGCT GGTACGCACG

TTCCAGATCG AAAACCCTGA CGTGCGTTTC TTCGAACTGC GTCCGGGTGC TGTTGACACC

TACTTCGGCG GTAGCAAAGC CGGCAAACCG AAGGAGCAGG GCTACCTGAA ACCGGAAGAA

GTTGCGGAAG CAGTTAAATA CCTGCTGCGT CTGCCGAAGG ACGTTCGTGT TGAGGAGCTG

ATGCTGCGCT CTATTTACCA GAAACCGGAA TAC

**sADH3 (Accession No. WP_011614641.1)**

ATGAAAATCT CTCTGACTTC CGCACGTCAG CTGGCCCGTG ATATCCTGGC CGCGCAGCAG

GTGCCGGCCG ATATCGCGGA CGACGTGGCT GAACATCTGG TTGAATCTGA CCGTTGTGGC

TATATCTCCC ACGGCCTGTC TATCCTGCCG AACTATCGTA CTGCGCTGGA CGGCCACTCT

GTTAACCCGC AAGGTCGTGC AAAATGTGTG CTGGATCAGG GTACCCTGAT GGTCTTCGAC

GGCGATGGTG GTTTTGGTCA ACACGTAGGC AAATCTGTTA TGCAGGCTGC GATTGAACGC

GTTCGTCAGC ACGGTCACTG TATCGTTACC CTGCGTCGTT CCCACCACCT GGGTCGCATG

GGCCACTACG GCGAAATGGC AGCTGCTGCG GGCTTTGTTC TGCTGTCCTT CACCAATGTG

ATCAACCGTG CGCCAGTTGT GGCGCCGTTC GGCGGTCGTG TTGCCCGTCT GACTACTAAC

CCGCTGTGCT TTGCTGGCCC GATGCCAAAC GGCCGCCCGC CTCTGGTAGT TGACATCGCA

ACCAGCGCCA TCGCCATCAA CAAAGCTCGC GTTCTGGCCG AAAAAGGCGA ACCAGCACCA

GAAGGCTCTA TCATCGGTGC GGATGGTAAC CCGACCACCG ATGCCTCTAC CATGTTTGGT

GAACACCCGG GTGCTCTGCT GCCATTCGGT GGTCACAAAG GTTATGCACT GGGTGTTGTA

GCTGAACTGC TGGCAGGTGT GCTGAGCGGC GGCGGTACGA TCCAGCCGGA CAACCCGCGT

GGCGGTGTGG CGACTAACAA CCTGTTCGCT GTTCTGCTGA ACCCGGCGCT GGATCTGGGC

CTGGACTGGC AGAGCGCTGA AGTTGAGGCT TTCGTTCGTT ACCTGCATGA TACCCCGCCG

GCCCCGGGTG TTGACCGTGT TCAGTACCCG GGCGAATATG AAGCAGCGAA CCGTGCGCAG

GCTTCCGACA CGCTGAACAT CAACCCGGCG ATCTGGCGCA ACCTGGAGCG CCTGGCGCAG

TCTCTGAACG TCGCAGTACC GACCGCT

**sADH4 (Accession No. P14941.1)**

ATGAAGGGTT TCGCTATGCT GTCCATCGGC AAGGTGGGCT GGATTGAAAA AGAGAAACCG

GCGCCTGGTC CGTTCGACGC CATTGTTCGC CCGCTGGCCG TAGCGCCGTG TACCTCTGAT

ATCCACACTG TGTTCGAGGG TGCGATTGGC GAACGCCACA ACATGATCCT GGGCCACGAG

GCGGTCGGCG AAGTAGTGGA AGTTGGTAGC GAGGTAAAGG ATTTCAAACC GGGCGATCGC

GTCGTTGTGC CGGCAATCAC TCCGGATTGG CGTACTAGCG AAGTACAGCG CGGTTACCAT

CAGCACAGCG GCGGCATGCT GGCGGGCTGG AAATTCAGCA ATGTTAAAGA CGGTGTTTTC

GGTGAATTTT TCCACGTGAA CGACGCCGAT ATGAACCTGG CGCACCTGCC GAAAGAAATT

CCGCTGGAAG CAGCGGTTAT GATCCCGGAC ATGATGACTA CTGGCTTCCA CGGTGCTGAA

CTGGCTGATA TTGAACTGGG TGCGACCGTA GCAGTACTGG GCATCGGTCC GGTGGGCCTG

ATGGCAGTGG CCGGCGCTAA ACTGCGTGGT GCGGGTCGCA TCATCGCCGT TGGCTCCCGC

CCGGTGTGCG TCGACGCTGC GAAATACTAT GGCGCGACGG ATATTGTTAA CTACAAAGAC

GGCCCGATTG AATCTCAGAT CATGAACCTG ACTGAAGGTA AAGGTGTTGA CGCTGCTATC

ATCGCCGGCG GCAATGCAGA CATCATGGCG ACTGCCGTGA AAATTGTGAA ACCGGGTGGT

ACCATCGCGA ACGTTAACTA TTTCGGTGAG GGTGAAGTTC TGCCAGTACC GCGTCTGGAG

TGGGGTTGTG GCATGGCCCA TAAAACCATC AAAGGTGGCC TGTGTCCGGG CGGTCGTCTG

CGCATGGAAC GTCTGATTGA CCTGGTCTTC TATAAACGTG TGGACCCGTC TAAACTGGTC

ACCCACGTAT TCCGTGGCTT CGATAACATT GAAAAAGCTT TCATGCTGAT GAAAGACAAA

CCGAAAGACC TGATTAAGCC GGTAGTCATC CTGGCG

**sADH5 (Accession No. AAA23199.2)**

ATGAAAGGCT TCGCTATGCT GGGTATCAAC AAACTGGGTT GGATCGAAAA GGAACGTCCG

GTGGCAGGCT CTTACGATGC TATTGTTCGT CCACTGGCTG TCTCTCCGTG CACCTCCGAT

ATCCACACTG TCTTTGAGGG CGCGCTGGGT GATCGTAAAA ACATGATCCT GGGCCACGAA

GCAGTCGGCG AGGTCGTTGA AGTTGGTTCT GAGGTGAAAG ACTTCAAACC AGGCGATCGT

GTTATCGTTC CGTGTACCAC CCCGGACTGG CGCTCTCTGG AAGTACAAGC AGGCTTCCAG

CAGCATAGCA ATGGTATGCT GGCAGGCTGG AAGTTCAGCA ACTTTAAGGA CGGTGTGTTC

GGCGAATACT TCCACGTTAA CGACGCAGAT ATGAACCTGG CGATCCTGCC GAAAGATATG

CCGCTGGAAA ACGCGGTTAT GATCACCGAC ATGATGACTA CCGGCTTCCA CGGTGCAGAG

CTGGCTGACA TCCAAATGGG TTCTTCTGTC GTGGTGATCG GCATCGGCGC TGTTGGCCTG

ATGGGCATCG CTGGCGCTAA GCTGCGCGGC GCCGGCCGTA TTATCGGCGT TGGCAGCCGT

CCGATCTGCG TGGAAGCTGC TAAATTCTAT GGCGCCACCG ACATTCTGAA CTACAAAAAT

GGTCACATCG TGGATCAGGT GATGAAACTG ACCAACGGTA AAGGTGTAGA CCGTGTAATC

ATGGCAGGTG GTGGTTCTGA AACCCTGAGC CAGGCGGTAT CTATGGTGAA ACCGGGTGGC

ATCATCTCCA ACATCAACTA TCACGGCTCT GGCGATGCGC TGCTGATCCC GCGTGTGGAG

TGGGGTTGCG GTATGGCTCA TAAAACCATC AAGGGCGGCC TGTGTCCGGG CGGTCGTCTG

CGCGCAGAAA TGCTGCGCGA TATGGTGGTA TATAACCGTG TAGACCTGAG CAAGCTGGTT

ACCCATGTTT ATCACGGTTT CGACCACATC GAGGAAGCCC TGCTGCTGAT GAAAGACAAA

CCTAAAGATC TGATCAAAGC TGTTGTAATT CTG

**sADH6 (Accession No. XP_455102.1)**

ATGCGCGCAC TGGCTTATTT CGGCAAGCAG GATATCCGTT ACACCAAGGA TCTGGAAGAA

CCGGTTATCG AGACCGACGA CGGCATCGAA ATTGAAGTCA GCTGGTGCGG TATCTGTGGC

TCCGACCTGC ACGAATATCT GGATGGTCCG ATTTTTTTCC CTGAAGATGG TAAAGTGCAT

GACGTTAGCG GTCTGGGCCT GCCACAGGCG ATGGGTCACG AGATGTCTGG CATCGTATCT

AAAGTGGGCC CGAAAGTTAC CAACATCAAA GCTGGTGACC ACGTGGTCGT CGAGGCAACT

GGCACTTGTC TGGATCATTA CACCTGGCCG AACGCGGCTC ATGCGAAAGA TGCTGAATGC

GCGGCGTGCC AGCGTGGCTT TTACAACTGC TGCGCCCACC TGGGTTTCAT GGGTCTGGGT

GTTCACAGCG GCGGTTTCGC GGAAAAAGTG GTGGTGAGCG AGAAACATGT TGTTAAGATC

CCGAACACCC TGCCGCTGGA CGTTGCAGCT CTGGTCGAAC CAATTTCTGT CTCCTGGCAC

GCGGTTCGTA TCAGCAAGCT GCAAAAAGGC CAATCTGCTC TGGTGCTGGG CGCTGGCCCA

ATTGGCCTGG CCACCATCCT GGCGCTGCAG GGTCACGGTG CAAGCAAAAT CGTTGTATCC

GAACCAGCGG AAATCCGTCG CAATCAAGCA GCAAAACTGG GCGTTGAAAC GTTCGATCCG

TCCGAACACA AAGAAGACGC GGTTAACATC CTGAAGAAAC TGGCACCGGG TGGTGAGGGT

TTCGATTTCG CCTACGACTG TAGCGGTGTC AAACCTACCT TTGATACTGG TGTACACGCT

ACCACCTTCC GCGGTATGTA CGTGAACATC GCAATTTGGG GTCATAAACC GATCGATTTC

AAACCGATGG ACGTGACTCT GCAGGAGAAG TTCGTCACGG GTTCCATGTG CTACACCATT

AAGGATTTCG AAGATGTGGT TCAGGCTCTG GGTAACGGCT CCATCGCCAT CGACAAGGCG

CGCCACCTGA TTACTGGCCG CCAGAAAATT GAAGATGGCT TCACCAAAGG CTTCGACGAA

CTGATGAACC ATAAAGAAAA AAACATCAAG ATCCTGCTGA CTCCTAATAA CCACGGCGAA

CTGGACGCGA CCAAC

**sADH7 (Accession No. AAP39869.1)**

ATGAAAGCAG CCGTGTTCAA AGGTAAGAAC CGTATCGTAC TGGATGAAAA GCCGGTGCCG

GTGCCAAAAC ATAGCGAAGC GCTGATTAAA ATCACTACCA CTACCATCTG CGGTACCGAC

ATTCACATCC TGAAGGGCGA ATACCCGGTA GCGGAAGGCC TGACCATTGG CCACGAACCG

ATCGGTGTGA TTGAGAGCTT CGGCGACGGT GTTACCGGCT TCAAGAAAGG CCAGCGTGTT

GTTATTGGCG CTATCACGCC GTGCAGCTCT TGTAGCAGCT GTATGGAAGG CATCCGTTCT

CAGTGTGGCA GCAAACTGAT GGGCGGCTGG AAATTCGGTA ATACCATCGA TGGCTCTCAG

GCGGAATATC TGATCGTGCC GGACGCTGCA GCGAACATGT ACCCTATTCC GGATGGTATT

ACCGATGAAC AGGTGCTGAT GTGTCCGGAC ATTATGTCCA CTGGTTTCTC CGGTCCAGAA

TCTGCCGGTG TGAAAGTGGG TGACACCGTC GTTATTTATG CTCAGGGTCC GATCGGTCTG

GGCGCTACCG CGGGTGCCAA AATGATGGGC GCAACGAAAG TAATTGTTGT AGATCGTTTC

CCGGAGCGTC TGGCGCTGGC AAAAAAGCTG GGCGCGGATT ATACGCTGGA TTTCACCAAA

TGTAACCCTA TTGAAGAAGT GATGCGTCTG ACTGGTGGTC GTGGCGTTGA CGTTGCAATT

GAAGCGCTGG GTCTGCAGTC CACCTTTGAG TCTTGCCTGC GTTGCCTGAA ACCGGGCGGT

GTGCTGAGCA GCCTGGGTGT CTATTCTGAT GACCTGCGTC TGCCGAACGA CGCTTTCGCT

GCGGGCCTGG GCGACTTCAA GATCGTTACT ACCCTGTGTC CTGGCGGCAA AGAGCGTATG

CGTCGTCTGC TGTCTGTAAT CGAGTCTGGC CGTGTTGACA TGCGCCCGAT GGTGACTCAC

ACCTTTAAGC TGGATGAAAT CGAAAAAGCC TACGACCTGT TCGGTAACCA GCGCGATGGC

GTTCTGAAAG TAGCCATCAA GCCG

**sADH8 (Accession No. Q0KDL6.1)**

ATGACCGCGA TGATGAAGGC AGCGGTATTT GTTGAACCTG GTCGTATTGA ACTGGCGGAT

AAACCGATTC CGGATATCGG TCCGAACGAC GCACTGGTTC GCATTACCAC CACCACTATT

TGCGGCACCG ACGTTCACAT CCTGAAGGGC GAATATCCGG TTGCTAAGGG CCTGACCGTG

GGCCACGAGC CGGTTGGCAT CATCGAGAAA CTGGGCAGCG CAGTAACGGG CTATCGCGAA

GGTCAGCGTG TAATCGCAGG TGCGATCTGT CCTAACTTCA ACTCTTACGC GGCGCAGGAT

GGTGTTGCGT CTCAGGATGG CAGCTACCTG ATGGCTTCTG GCCAGTGCGG CTGTCATGGT

TACAAAGCAA CCGCCGGCTG GCGTTTCGGT AATATGATTG ACGGTACCCA GGCTGAATAT

GTTCTGGTAC CGGATGCGCA GGCCAACCTG ACCCCGATTC CGGACGGCCT GACGGACGAA

CAGGTCCTGA TGTGTCCGGA TATTATGTCC ACCGGTTTCA AAGGTGCGGA GAACGCAAAC

ATTCGCATTG GTGACACCGT TGCGGTGTTT GCGCAGGGTC CGATCGGTCT GTGCGCGACT

GCTGGTGCGC GTCTGTGCGG CGCAACTACT ATCATCGCTA TCGACGGCAA CGATCATCGT

CTGGAAATCG CGCGCAAAAT GGGTGCGGAC GTTGTGCTGA ACTTCCGTAA CTGCGACGTG

GTCGACGAGG TGATGAAACT GACCGGCGGT CGTGGTGTGG ATGCGTCTAT TGAAGCACTG

GGTACCCAGG CGACCTTCGA ACAATCTCTG CGTGTTCTGA AGCCGGGTGG TACGCTGTCT

TCTCTGGGTG TCTACTCCTC TGATCTGACT ATCCCGCTGA GCGCCTTCGC AGCCGGTCTG

GGCGACCACA AAATTAACAC CGCCCTGTGC CCGGGTGGCA AAGAACGTAT GCGTCGTCTG

ATTAACGTTA TCGAATCCGG TCGCGTGGAC CTGGGTGCGC TGGTAACCCA CCAGTATCGC

CTGGATGATA TCGTTGCGGC GTACGACCTG TTCGCTAACC AGCGTGATGG TGTTCTGAAA

ATCGCCATCA AACCGCAC

**sADH9 (Accession No. XP_001580601.1)**

ATGACCTTCG AACTGCCGAA AACTATGAAA GCGTTTGCGA TGCGCAAGAT CGGTGAAGTC

GGCTGGATTG AAAAGCCGGT GCCAGAATGC GGTCCGAACG ATGCAATCTG CCGTCCGCTG

GCTCTGGCTC CGTGCACCTC TGACATTCAC ACTGTGTGGG CGGGTGCGAT CGGTGAACGC

CATGACATGA TCCTGGGCCA TGAAGCCGTT GGCCAAGTTG TTAAAATCGG TTCCGAAGTT

AAAAACCTGA AAGTTGGCGA TAAGGTTCTG GTTCCGGCAG TAACTCCGGA TTGGGGTAGC

GAAGCAGCAC AGGAAGGCTT CCCGGCACAC TCCGGCGGTA TGCTGGGTGG TTGGAAATTT

TCCAACTTCA AAGATGGCGT TTTTGCTGAA TACTTCCACG TCAACGAAGC CGACGCTAAC

CTGGCCAAAC TGCCGGAAGG TCTGACGCCG CGTGATGTAA TCATGTGCAG CGACATGATG

ACTACCGGTT TCCATGGTGC AGAACTGGCG GAGGTTAAGC TGGGTGATAT CGTAGTAGTT

ATCGGTATCG GTCCGGTTGG TCTGATGTCT GTACGTGGCG CTGCTCTGAT GGGCGCAAGC

CGTATCTTCG CAGTCGGTTC CCGTCCGCAC TGCTGCGACA CCGCAGTCCA GTACGGTGCC

ACTGACATCA TCAACTATAA AAATGGTGAT ATCGTGGAAC AGATCCTGAA AGCTACTGGC

GGTAAAGGCG TTGATCGTGT GATTATCGCG GGTGGCGACG TCCATACTTT CGCGCAAGCT

ATTAAAATGA TCCGTGCTGG CGGCGTTATC GGCAACGTGA ACTACCTGGG CGAAGGTGAA

ATGATCGACG TACCGCGTGT AGAATGGGGT GTTGGCATGG GTCACAAATT CATCCACGGT

GGTCTGACCC CGGGTGGTGC ACTGCGTATG GAAAAAATGG CTAACCTGAT CAAGTATAAA

AAAGTCGATC CGACGAAACT GATTACCCAT GAATTCAAGG GTCTGGAAAA AGTTGAAGAT

GCTCTGATGC TGATGAAAGA CAAACCGGTT GACCTGATCA AACCGGTTGT TCTGATTGAA

TACAACGACA AACTG

**sADH10 (Accession No. AJP52792.1)**

ATGATGGCGA CCATGAAAGC AGCGATCTTT GTTGAAAAAA ACCGTATTGT TCTGGAAGAC

AAACCAATCC CGGAAGTTGG CCCGCTGGAT GCACTGATCC GTATCACCAC CACTACGATT

TGCGGCACCG ATGTTCACAT CCTGCGCGGT GAGTATCCGG TAGCGAAGGG TCTGACGATC

GGTCATGAAC CAGTAGGTAT TATCGAACGT CTGGGTTCCC AGGTTCGTGG TTTCGTAGAA

GGTCAGCGTG TTATTGCAGG CGCGATCACC CCGTCTGGCC AAAGCTACGC ATGCCTGTGT

GGCTGTGCCA GCCAGGACGG TCCGGATACC CGTCACGGTT TTCGTGCGAC CGGCGGCTGG

AAATTCGGCA ACATTATCGA CGGCTGCCAG GCGGAGTATG TACTGGTGCC AGACGCGCTG

GCGAACCTGT GCCCAATTCC GGATGGCCTG AGCGACGAAC AAGTTCTGAT GTGCCCGGAC

ATCATGTCCA CTGGTTTCTC TGGCGCAGAA CGTGGCGAAA TTAACATTGG TGATACTGTT

GCGGTATTCG CACTGGGTCC GATCGGCCTG TGTGCTGTGG CGGGCGCCCG TCTGAAGGGT

GCGACCACCA TCATCGGTGT GGACGCAGTG GCTCAGCGTA TGTCTGTTGC ACGTCAGCTG

GGCGCCACCC ACGTGGTAAA CTTCAAAGAG GCGAACGTTG TGGAACAGAT TATGGCGCTG

ACGGACGGCC GTGGTGTTGA TGTATCTATC GAAGCACTGG GCACCCAGGG CACCTTCGAA

TCTGCTCTGC GTGTCCTGCG CCCGGGCGGT CGCCTGTCCT CCCTGGGTGT TTATTCCAGC

GACCTGCGTA TCCCGCTGGA CGCCTTCGCC GCAGGCCTGG GCGATTACTC CATCGTCACT

ACCCTGTGTC CGGGTGGCAA AGAGCGTATG CGTCGCCTGA TGGCCGTAGT TCAGAGCGGC

GCGGTCGACC TGTCTCCGCT GGTCACTCAC CACTTCAAGC TGGATGACAT CGAAGCGGCA

TATGAACTGT TTGCGAACCA GCGTGATGGT GTAATGAAAG TTGCAATCAC CCCG

**sADH11 (Accession No. WP_011835462.1)**

ATGTCTAAAG TGGCTGCCGT GACCGGTGCA GGTCAGGGCA TTGGTTTTGC AATCGCGAAA

CGCCTGTATA ACGACGGTTT CAAAGTAGCC ATCATCGATT ACAACGAAGA AACCGCTCAG

CAGGCTGCGA AAGAACTGGG TGGTGAATCT TTCGCGCTGA AGGCGGATGT TTCTGACCGT

GACCAGGTAG TCGCCGCGCT GGAAGCTGTT GTTGAGAAAT TCGGTGATCT GAACGTGGTA

GTAAACAACG CGGGTATCGC CCCGACTACC CCGATCGAAA CGATCACCCC GGAACAGTTT

CACCAGGTGT ACAACATCAA TGTTGGTGGT GTGCTGTGGG GTACCCAGGC TGCTACTGCT

CTGTTCCGTA AACTGGGCCA CGGCGGTAAG ATTATCAACG CCACCTCCCA GGCTGGTGTG

GTCGGTAACC CGAACCTGAT GCTGTACAGC AGCTCCAAAT TCGCTGTCCG CGGCATGACC

CAGATCGCAG CACGTGACCT GGCAGAAGAG GGTATCACCG TCAACGCCTA CGCTCCAGGC

ATTGTGAAAA CCCCGATGAT GTTCGATATC GCTCATCAGG TGGGTAAGAA CGCCGGCAAA

GACGACGAGT GGGGTATGCA GACCTTCGCT AAAGACATCG CGATGAAACG TCTGAGCGAG

CCGGAAGATG TAGCAAACGT TGTTTCCTTT CTGGCCGGCC CGGATTCCAA CTACATCACC

GGCCAGACTA TCATTGTAGA CGGTGGCATG CAATTTCAT

**sADH12 (Accession No. AAC04974.1)**

ATGCGTGCGC TGGCCTATTT CAAGAAAGGT GACATTCACT TCACTAATGA CATCCCACGT

CCGGAAATCC AGACGGACGA CGAAGTTATC ATCGATGTGT CTTGGTGCGG CATCTGTGGT

TCTGACCTGC ACGAATACCT GGATGGCCCG ATCTTTATGC CGAAAGACGG TGAGTGCCAC

AAACTGAGCA ACGCCGCTCT GCCTCTGGCT ATGGGCCACG AAATGAGCGG TATCGTTTCC

AAGGTTGGTC CGAAAGTTAC TAAAGTAAAA GTGGGTGATC ACGTAGTTGT TGATGCGGCC

AGCTCCTGCG CCGACCTGCA CTGCTGGCCG CACTCCAAAT TTTATAACTC TAAGCCGTGT

GACGCATGTC AGCGTGGTTC CGAAAACCTG TGCACCCACG CGGGCTTTGT AGGTCTGGGC

GTCATTAGCG GTGGCTTCGC GGAGCAGGTA GTTGTATCTC AACACCACAT CATCCCGGTT

CCGAAGGAGA TCCCGCTGGA CGTTGCAGCT CTGGTTGAAC CGCTGAGCGT TACTTGGCAT

GCAGTGAAAA TTTCTGGTTT CAAAAAAGGT TCTTCCGCAC TGGTTCTGGG CGCAGGTCCG

ATCGGTCTGT GCACCATCCT GGTTCTGAAA GGCATGGGTG CATCCAAGAT TGTCGTATCC

GAAATTGCGG AGCGCCGTAT TGAAATGGCG AAAAAGCTGG GTGTTGAAGT ATTCAACCCG

TCTAAACACG GCCACAAATC CATTGAGATC CTGCGTGGCC TGACCAAAAG CCATGATGGC

TTTGACTACA GCTATGATTG CTCTGGCATC CAGGTGACTT TCGAAACTTC TCTGAAAGCG

CTGACTTTCA AAGGTACCGC TACCAACATT GCAGTGTGGG GCCCGAAACC GGTTCCATTC

CAGCCGATGG ATGTCACCCT GCAGGAAAAA GTAATGACCG GCTCCATTGG TTACGTTGTG

GAAGATTTCG AAGAAGTTGT GCGTGCTATT CATAACGGCG ATATTGCTAT GGAAGATTGT

AAGCAGCTGA TCACCGGTAA ACAGCGTATC GAAGATGGTT GGGAGAAAGG CTTTCAGGAA

CTGATGGATC ACAAAGAATC TAACGTAAAA ATTCTGCTGA CTCCAAACAA CCACGGTGAA

ATGAAA

**sADH13 (Accession No. WP_000374004.1)**

ATGGATCGTA TCATTCAGTC TCCTGGCAAA TACATCCAGG GCGCAGATGT GATCAACCGT

CTGGGTGAAT ATCTGAAACC GCTGGCAGAG CGTTGGCTGG TGGTTGGCGA TAAATTCGTG

CTGGGCTTTG CTCAGTCTAC CGTGGAAAAA TCCTTCAAGG ATGCGGGCCT GGTGGTTGAA

ATTGCACCGT TCGGCGGTGA ATGTAGCCAC AACGAAATTG ACCGTCTGCG TGGCATCGCT

GAAACTGCCC AATGTGGCGC GATTCTGGGC ATCGGTGGCG GCAAGACCCT GGACACTGCC

AAAGCGCTGG CGCATTTCAT GGGTGTCCCA GTTGCGATCG CTCCGACCAT CGCGTCTACC

GCTGCGCCGT GCTCTGCCCT GTCCGTGATC TACACGGACG AAGGCGAGTT CGACCGCTAC

CTGCTGCTGC CAAACAACCC GAATATGGTG ATTGTGGACA CCAAAATCGT TGCAGGTGCG

CCGGCGCGTC TGCTGGCGGC TGGCATTGGT GACGCACTGG CGACCTGGTT CGAAGCACGC

GCGTGCTCCC GTTCTGGTGC GACGACCATG GCTGGCTGCA AATGCACCCA GGCAGCGCTG

GCGCTGGCGG AACTGTGCTA TAACACCCTG CTGGAGGAAG GCGAAAAAGC AATGCTGGCG

GCGGAACAGC ACGTCGTAAC CCCTGCTCTG GAACGTGTAA TTGAAGCAAA TACTTACCTG

AGCGGCGTGG GTTTCGAATC CGGTGGCCTG GCGGCTGCGC ACGCTGTTCA TAACGGCCTG

ACCGCGATCC CTGACGCGCA CCACTACTAT CACGGCGAGA AAGTGGCTTT CGGCACCCTG

ACTCAACTGG TACTGGAAAA CGCGCCGGTC CAGGAAATCG AAACCGTGGC GGCGCTGTCC

CATGCAGTAG GCCTGCCGAT CACCCTGACC CAGCTGGACA TCAAAGAAGA CGTGCCAGCA

AAAATGCGTA TCGTGGCCGA AGCGGCATGC GCAGAAGGTG AGACCATCCA TAACATGCCG

GGCGGCGCTA CCCCGGATCA GGTCTACGCC GCCCTGCTGG TAGCAGACCA ATATGGCCAG

CGTTTTCTGC AGGAATGGGA ATAA

**sADH14 (Accession No. BAD32689.1)**

ATGATGAAAG CGCTGTGTTA CCTGGGTTCT AAAACTATCA AATGGCAGAC CGTAGCCAAA

CCGACCCTGA AATCCCCAAC CGACGTTATC GGTAAACTGT CTGCACTGAC CCTGTGTGGC

TCTGATCTGC ACATTATTGC GGGTCACGTA AAAGAAAGCA CCGATATCGC CGAGTCCCAG

CCGGGCCGTG GCCTGATCCT GGGTCACGAA GGTATCATCA AAGTGGAAGA GGTAGGCGAT

GCTGTCAAAA ACTTCAAACG TGGTGACGTT TGCATCGTCT CTTGCATCAC CAGCTGCGGT

GAATGCTACT ACTGCAAACG TGATCTGCAG TCCCATTGCA ACCGTACCGA AGGTACCTCC

GGCTGGATCC TGGGCCACGA AATTGATGGT ACCCAGGCCG AATACGTCCG TATCCCGTAC

GCCGACCAGT CTCTGTACAA AGCTCCGGAG AACGTGCCGA TCGAGAGCCT GCTGATGCTG

AGCGACATCC TGCCGACCGC GTACGAAGTG GGCGTTGTCT CTGGCAATGT GAAAAAAGGT

GATACCGTCG CCATCGTAGG TCTGGGCCCT GTTGGCCTGT CCGCGCTGCT GTCCGCTAAA

GCCCTGGACC CGGCTAAAAT CATTGCAATT GATATGGACG ACTCTCGTCT GGAGGTTGCA

CGTCGTCTGG GTGCGCACGA AACTATCAAT CCGGGCAAAC AGGATGCTGC AAAGCTGGTT

CATGAACTGA CCGCGAGCGA AGGTAAGTCT TCTGGTGTAG ACGTTGCAAT CGAATGTGTT

GGCGTGCCGG CCACCTTTGA AATGTGCGAA GACCTGCTGT GTCCGGGTGG CCACCTGTCT

AACGTTGGCG TTCACGGTTC TAGCGTCGAA CTGAAACTGC AGGAGCTGTG GATCAAAAAC

ATTTCTATCA GCACCGGTCT GGTATCTGCT TATTCCACCG AAACCCTGCT GCAGAAAGTT

ATCGACAAAA AACTGGACCC GACCCCTCTG GCAACCCACC ATTTCAAGCT GAGCGAAATC

GAAAAGGCGT ACGATGTCTT CTCTCACGCC GCAGATAACC AGGCCATCAA AATGGTGATC

ACCTGTGACG AA

**sADH15 (Accession No. BAA24528.1)**

ATGAGCATTC CGTCCAGCCA GTACGGCTTC GTGTTCAACA AACAGTCCGG CCTGAACCTG

CGCAACGATC TGCCTGTTCA TAAACCGAAG GCGGGTCAGC TGCTGCTGAA AGTTGATGCT

GTTGGTCTGT GCCATTCCGA TCTGCACGTA ATTTATGAGG GCCTGGATTG CGGCGATAAC

TACGTGATGG GCCACGAAAT TGCAGGTACC GTGGCTGCGG TAGGCGACGA CGTTATCAAC

TACAAAGTAG GCGACCGTGT TGCTTGTGTG GGCCCGAACG GTTGCGGTGG TTGCAAGTAT

TGCCGTGGTG CAATCGACAA CGTTTGCAAG AACGCATTTG GTGACTGGTT TGGTCTGGGC

TACGATGGTG GTTATCAGCA GTATCTGCTG GTAACCCGTC CGCGTAACCT GAGCCGTATC

CCAGATAACG TTTCCGCCGA CGTAGCGGCG GCATCTACCG ACGCGGTACT GACTCCGTAT

CATGCTATTA AGATGGCACA GGTGTCTCCT ACCAGCAACA TCCTGCTGAT CGGTGCGGGC

GGCCTGGGTG GTAACGCGAT CCAGGTAGCG AAAGCCTTCG GTGCAAAAGT TACTGTACTG

GATAAAAAAA AAGAGGCACG CGATCAAGCT AAAAAACTGG GTGCAGATGC GGTCTACGAA

ACTCTGCCGG AATCTATCTC TCCTGGTTCC TTCAGCGCTT GCTTCGATTT CGTCTCTGTA

CAGGCGACGT TCGATGTTTG CCAGAAATAT GTTGAGCCAA AAGGTGTAAT TATGCCGGTT

GGTCTGGGCG CGCCTAACCT GTCTTTTAAC CTGGGTGATC TGGCGCTGCG CGAAATTCGT

ATTCTGGGTT CCTTCTGGGG TACCACTAAC GATCTGGACG ACGTCCTGAA ACTGGTTTCC

GAGGGCAAAG TGAAACCGGT TGTTCGCAGC GCAAAACTGA AAGAGCTGCC GGAGTATATC

GAAAAACTGC GTAACAACGC GTATGAGGGT CGTGTTGTCT TTAACCCG

**sADH16 (Accession No. BAN45671.1)**

ATGCTGCGCA CTAATCTGCG CACGTTCGCT CGTCCGCAGT TTATTCGCGG CCTGGCGACT

GCGCCGGTAA TCCCGAAAAC CCAAAAAGGC GTTATTTTCT ACGAAAACGG TGGTGAACTG

CAGTACAAGG ACATCCCGGT ACCGGAGCCA AAACCGAACG AAATCCTGGT TAATGTTAAG

TACAGCGGTG TTTGTCACAC CGATCTGCAC GCCTGGAAGG GTGACTGGCC ACTGCCAGTG

AAACTGCCGC TGGTGGGTGG TCACGAGGGC GCCGGCATCG TGGTAGCTAA GGGCAGCGAA

GTAAAAAACT TTGAGATCGG TGATTATGCG GGCATCAAAT GGCTGAACGG CAGCTGTATG

AGCTGTGAGC TGTGCGAAAA AGGTTACGAA TCCAACTGCC TGCAGGCCGA TCTGAGCGGC

TATACCCATG ACGGTAGCTT TCAGCAATAC GCTACCGCGG ATGCGGTTCA GGCGGCTCAG

ATCCCGAAGA ACGTAGACCT GGCTGAGATC AGCCCTATCC TGTGCGCTGG TGTCACTGTT

TATAAGGCGC TGAAAACCGC GGATCTGGCT CCGGGCCAGT GGGTCGCTAT CTCTGGCGCG

GCTGGTGGTC TGGGTTCTCT GGCGGTGCAA TACGCTAAAG CTATGGGTCT GCGTGTCCTG

GGCATCGACG GTGGTGCAGA AAAAGAAAAG CTGTTTAAAA GCCTGGGTGG CGAGATCTTC

ATCGATTTCA CCAAAGAAAA AAACATCGTA GAAGCCATTC AGGAGGCAAC TAATGGCGGC

CCGCATGGCG TCATCAACGT ATCCGTTGCG GAAGCAGCGA TCTCTCAGTC CACCGAGTAT

GTTCGTCCGA CCGGTACCGT AGTACTGGTG GGTCTGCCAG CAGGTGCTGT CTGCAAAAGC

GAAGTGTTCT CTCACGTGGT AAAATCTATC TCTATCAAAG GCAGCTACGT GGGTAATCGT

GCGGACACCC GTGAGGCGAT CGACTTCTTC GAACGTGGTC TGGTGCGTAG CCCGATCAAG

ATCGTGGGTC TGAGCGAACT GCCGGAAGTG TACAAACTGA TGGAGCAGGG CAAAATCCTG

GGCCGTTACG TCGTGGATAC TACCAAA

**sADH17 (Accession No. CAD36475.1)**

ATGAAAGCGC TGCAATACAC CGAAATCGGT AGCGAACCGG TGGTCGTTGA TGTCCCGACC

CCTGCGCCGG GTCCGGGTGA AATTCTGCTG AAAGTAACCG CTGCGGGTCT GTGTCACAGC

GACATCTTCG TTATGGACAT GCCGGCTGAA CAGTATATCT ACGGCCTGCC GCTGACCCTG

GGCCATGAAG GCGTGGGCAC CGTTGCGGAA CTGGGTGCGG GCGTGACCGG TTTCGAAACT

GGCGACGCTG TTGCAGTTTA CGGCCCTTGG GGCTGTGGCG CTTGCCACGC ATGCGCACGT

GGCCGTGAGA ACTACTGCAC CCGTGCGGCA GAGCTGGGCA TCACTCCTCC GGGCCTGGGC

TCTCCGGGTA GCATGGCTGA GTACATGATC GTTGACTCCG CTCGTCACCT GGTCCCGATC

GGTGACCTGG ACCCGGTGGC AGCAGTGCCG CTGACCGATG CTGGCCTGAC CCCGTACCAC

GCCATCAGCC GTGTACTGCC GCTGCTGGGC CCAGGTTCTA CGGCTGTTGT AATTGGCGTC

GGTGGTCTGG GTCACGTTGG CATTCAGATT CTGCGCGCCG TTAGCGCCGC GCGTGTTATC

GCGGTTGATC TGGACGACGA CCGCCTGGCG CTGGCGCGTG AAGTTGGTGC AGATGCGGCT

GTTAAATCTG GTGCGGGTGC GGCGGATGCT ATTCGTGAAC TGACTGGTGG CGAAGGTGCT

ACTGCTGTTT TCGATTTTGT GGGTGCGCAG TCTACCATCG ACACCGCACA GCAAGTCGTG

GCAATCGATG GCCACATCAG CGTTGTCGGC ATCCACGCCG GTGCACACGC CAAAGTTGGT

TTCTTCATGA TCCCATTCGG TGCTAGCGTT GTTACCCCGT ATTGGGGTAC TCGCTCCGAG

CTGATGGACG TTGTTGATCT GGCTCGTGCA GGCCGCCTGG ATATTCATAC CGAAACCTTC

ACCCTGGACG AAGGCCCTAC TGCATACCGC CGTCTGCGCG AGGGCTCTAT CCGTGGCCGT

GGCGTGGTTG TACCGGGTTA A

**F. DNA probes for rRNA depletion.**

| **Name** | **Sequence** |
| --- | --- |
| 23S-3 | CACTTATCTCTTCCGCATTTAGCTACCGGGCAGTGCCATTGGCATGACAACCCGAACACCAGTGATGCGTCCACTCCGGT |
| 23S-4 | CCTCTCGTACTAGGAGCAGCCCCCCTCAGTTCTCCAGCGCCCACGGCAGATAGGGACCGAACTGTCTCACGACGTTCTAA |
| 23S-5 | ACCCAGCTCGCGTACCACTTTAAATGGCGAACAGCCATACCCTTGGGACCTACTTCAGCCCCAGGATGTGATGAGCCGAC |
| 23S-6 | ATCGAGGTGCCAAACACCGCCGTCGATATGAACTCTTGGGCGGTATCAGCCTGTTATCCCCGGAGTACCTTTTATCCGTT |
| 23S-7 | GAGCGATGGCCCTTCCATTCAGAACCACCGGATCACTATGACCTGCTTTCGCACCTGCTCGCGCCGTCACGCTCGCAGTC |
| 23S-8 | AAGCTGGCTTATGCCATTGCACTAACCTCCTGATGTCCGACCAGGATTAGCCAACCTTCGTGCTCCTCCGTTACTCTTTA |
| 23S-9 | GGAGGAGACCGCCCCAGTCAAACTACCCACCAGACACTGTCCGCAACCCGGATTACGGGTCAACGTTAGAACATCAAACA |
| 23S-10 | TTAAAGGGTGGTATTTCAAGGTCGGCTCCATGCAGACTGGCGTCCACACTTCAAAGCCTCCCACCTATCCTACACATCAA |
| 23S-11 | GGCTCAATGTTCAGTGTCAAGCTATAGTAAAGGTTCACGGGGTCTTTCCGTCTTGCCGCGGGTACACTGCATCTTCACAG |
| 23S-12 | CGAGTTCAATTTCACTGAGTCTCGGGTGGAGACAGCCTGGCCATCATTACGCCATTCGTGCAGGTCGGAACTTACCCGAC |
| 23S-13 | AAGGAATTTCGCTACCTTAGGACCGTTATAGTTACGGCCGCCGTTTACCGGGGCTTCGATCAAGAGCTTCGCTTGCGCTA |
| 23S-14 | ACCCCATCAATTAACCTTCCGGCACCGGGCAGGCGTCACACCGTATACGTCCACTTTCGTGTTTGCACAGTGCTGTGTTT |
| 23S-15 | TTAATAAACAGTTGCAGCCAGCTGGTATCTTCGACTGATTTCAGCTCCACGAGCAAGTCGCTTCACCTACATATCAGCGT |
| 23S-16 | GCCTTCTCCCGAAGTTACGGCACCATTTTGCCTAGTTCCTTCACCCGAGTTCTCTCAAGCGCCTTGGTATTCTCTACCTG |
| 23S-17 | ACCACCTGTGTCGGTTTGGGGTACGATTTGATGTTACCTGATGCTTAGAGGCTTTTCCTGGAAGCAGGGCATTTGTTGCT |
| 23S-18 | TCAGCACCGTAGTGCCTCGTCATCACGCCTCAGCCTTGATTTTCCGGATTTGCCTGGAAAATCAGCCTACACGCTTAAAC |
| 23S-19 | CGGGACAACCGTCGCCCGGCCAACATAGCCTTCTCCGTCCCCCCTTCGCAGTAACACCAAGTACAGGAATATTAACCTGT |
| 23S-20 | TTCCCATCGACTACGCCTTTCGGCCTCGCCTTAGGGGTCGACTCACCCTGCCCCGATTAACGTTGGACAGGAACCCTTGG |
| 23S-21 | TCTTCCGGCGAGCGGGCTTTTCACCCGCTTTATCGTTACTTATGTCAGCATTCGCACTTCTGATACCTCCAGCATACCTC |
| 23S-22 | ACAGTACACCTTCACAGGCTTACAGAACGCTCCCCTACCCAACAACGCATAAGCGTCGCTGCCGCAGCTTCGGTGCATGG |
| 23S-23 | TTTAGCCCCGTTACATCTTCCGCGCAGGCCGACTCGACCAGTGAGCTATTACGCTTTCTTTAAATGATGGCTGCTTCTAA |
| 23S-24 | GCCAACATCCTGGCTGTCTGGGCCTTCCCACATCGTTTCCCACTTAACCATGACTTTGGGACCTTAGCTGGCGGTCTGGG |
| 23S-25 | TTGTTTCCCTCTTCACGACGGACGTTAGCACCCGCCGTGTGTCTCCCGTGATAACATTCTCCGGTATTCGCAGTTTGCAT |
| 23S-26 | CGGGTTGGTAAGTCGGGATGACCCCCTTGCCGAAACAGTGCTCTACCCCCGGAGATGAGTTCACGAGGCGCTACCTAAAT |
| 23S-27 | AGCTTTCGGGGAGAACCAGCTATCTCCCGGTTTGATTGGCCTTTCACCCCCAGCCACAAGTCATCCGCTAATTTTTCAAC |
| 23S-28 | ATTAGTCGGTTCGGTCCTCCAGTTAGTGTTACCCAACCTTCAACCTGCCCATGGCTAGATCACCGGGTTTCGGGTCTATA |
| 23S-29 | CCCTGCAACTTAACGCCCAGTTAAGACTCGGTTTCCCTTCGGCTCCCCTATTCGGTTAACCTTGCTACAGAATATAAGTC |
| 23S-30 | GCTGACCCATTATACAAAAGGTACGCAGTCACACGCCTAAGCATGCTCCCACTGCTTGTACGTACACGGTTTCAGGTTCT |
| 23S-31 | TTTTCACTCCCCTCGCCGGGGTTCTTTTCGCCTTTCCCTCACGGTACTGGTTCACTATCGGTCAGTCAGGAGTATTTAGC |
| 23S-32 | CTTGGAGGATGGTCCCCCCATATTCAGACAGGATACCACGTGTCCCGCCCTACTCATCGAGCTCACAGCATGTGCATTTT |
| 23S-33 | TGTGTACGGGGCTGTCACCCTGTATCGCGCGCCTTTCCAGACGCTTCCACTAACACACACACTGATTCAGGCTCTGGGCT |
| 23S-34 | CCTCCCCGTTCGCTCGCCGCTACTGGGGGAATCTCGGTTGATTTCTTTTCCTCGGGGTACTTAGATGTTTCAGTTCCCCC |
| 23S-35 | GGTTCGCCTCATTAACCTATGGATTCAGTTAATGATAGTGTGTCGAAACACACTGGGTTTCCCCATTCGGAAATCGCCGG |
| 23S-36 | TTATAACGGTTCATATCACCTTACCGACGCTTATCGCAGATTAGCACGTCCTTCATCGCCTCTGACTGCCAGGGCATCCA |
| 23S-37 | CCGTGTACGCTTAGTCGCTTAA |
| 16S-1 | TAAGGAGGTGATCCAACCGCAGGTTCCCCTACGGTTACCTTGTTACGACTTCACCCCAGTCATGAATCACAAAGTGGTAA |
| 16S-2 | GCGCCCTCCCGAAGGTTAAGCTACCTACTTCTTTTGCAACCCACTCCCATGGTGTGACGGGCGGTGTGTACAAGGCCCGG |
| 16S-3 | GAACGTATTCACCGTGGCATTCTGATCCACGATTACTAGCGATTCCGACTTCATGGAGTCGAGTTGCAGACTCCAATCCG |
| 16S-4 | GACTACGACGCACTTTATGAGGTCCGCTTGCTCTCGCGAGGTCGCTTCTCTTTGTATGCGCCATTGTAGCACGTGTGTAG |
| 16S-5 | CCCTGGTCGTAAGGGCCATGATGACTTGACGTCATCCCCACCTTCCTCCAGTTTATCACTGGCAGTCTCCTTTGAGTTCC |
| 16S-6 | CGGCCGGACCGCTGGCAACAAAAGATAAGGGTTGCGCTCGTTGCGGGACTTAACCCAACATTTCACAACACGAGCTGACG |
| 16S-7 | ACAGCCATGCAGCACCTGTCTCACAGTTCCCGAAGGCACCAATCCATCTCTGGAAAGTTCTGTGGATGTCAAGACCAGGT |
| 16S-8 | AAGGTTCTTCGCGTTGCATCGAATTAAACCACATGCTCCACCGCTTGTGCGGGCCCCCGTCAATTCATTTGAGTTTTAAC |
| 16S-9 | CTTGCGGCCGTACTCCCCAGGCGGTCGACTTAACGCGTTAGCTCCGGTAGCCACGCCTCAAGGGCACAACCTCCAAGTCG |
| 16S-10 | ACATCGTTTACGGCGTGGACTACCAGGGTATCTAATCCTGTTTGCTCCCCACGCTTTCGCACCTGAGCGTCAGTCTTCGT |
| 16S-11 | CCAGGGGGCCGCCTTCGCCACCGGTATTCCTCCAGATCTCTACGCATTTCACCGCTACACCTGGAATTCTACCCCCCTCT |
| 16S-12 | ACGAGACTCAAGCTTGCCAGTATCAGATGCAGTTCCCAGGTTGAGCCCGGGGATTTCACATCTGACTTAACAAACCGCCT |
| 16S-13 | GCGTGCGCTTTACGCCCAGTAATTCCGATTAACGCTTGCACCCTCCGTATTACCGCGGCTGCTGGCACGGAGTTAGCCGG |
| 16S-14 | TGCTTCTTCTGCGGGTAACGTCAATGAGCAAAGGTATTAACTTTACTCCCTTCCTCCCCGCTGAAAGTACTTTACAACCC |
| 16S-15 | GAAGGCCTTCTTCATACACGCGGCATGGCTGCATCAGGCTTGCGCCCATTGTGCAATATTCCCCACTGCTGCCTCCCGTA |
| 16S-16 | GGAGTCTGGACCGTGTCTCAGTTCCAGTGTGGCTGGTCATCCTCTCAGACCAGCTAGGGATCGTCGCCTAGGTGAGCCGT |
| 16S-17 | TACCCCACCTACTAGCTAATCCCATCTGGGCACATCCGATGGCAAGAGGCCCGAAGGTCCCCCTCTTTGGTCTTGCGACG |
| 16S-18 | TTATGCGGTATTAGCTACCGTTTCCAGTAGTTATCCCCCTCCATCAGGCAGTTTCCCAGACATTACTCACCCGTCCGCCA |
| 16S-19 | CTCGTCAGCAAAGAAGCAAGCTTCTTCCTGTTACCGTTCGACTTGCATGTGTTAGGCCTGCCGCCAGCGTTCAATCTGAG |
| 16S-20 | CCATGATCAAACTCTTCAATTTAAA |
| 5S-1 | ATGCCTGGCAGTTCCCTACTCTCGCATGGGGAGACCCCACACTACCATCGGCGCTACGGCGTTTCACTTCTGAGTTCGGC |
| 5S-2 | ATGGGGTCAGGTGGGACCACCGCGCTACGGCCGCCAGGCA |
| 23S-1 | AAGGTTAAGCCTCACGGTTCATTAGTACCGGTTAGCTCAACGCATCGCTGCGCTTACACACCCGGCCTATCAACGTCGTC |
| 23S-2 | GTCTTCAACGTTCCTTCAGGACTCTCAAGGAGTCAGGGAGAACTCATCTCGGGGCAAGTTTCGTGCTTAGATGCTTTCAG |

**G. Open reading frame for *pncB* and *rpoC***

***pcnB***

```
       1  GTGCTAAGCC GCGAGGAAAG CGAGGCTGAA CAGGCAGTCG CCCGTCCACA GGTGACGGTG
      61  ATCCCGCGTG AGCAGCATGC TATTTCCCGC AAAGATATCA GTGAAAATGC CCTGAAGGTA
```

121 ATGTACAGGC TCAATAAAGC GGGATACGAA GCCTGGCTGG TTGGCGGCGG CGTGCGCGAC

181 CTGTTACTTG GCAAAAAGCC GAAAGATTTT GACGTAACCA CTAACGCCAC GCCTGAGCAG

241 GTGCGCAAAC TGTTCCGTAA CTGCCGCCTG GTGGGTCGCC GTTTCCGTCT GGCTCATGTA

301 ATGTTTGGCC CGGAGATTAT CGAAGTTGCG ACCTTCCGTG GACACCACGA AGGTAACGTC

361 AGCGACCGCA CGACCTCCCA ACGCGGGCAA AACGGCATGT TGCTGCGCGA CAACATTTTC

421 GGCTCCATCG AAGAAGACGC CCAGCGCCGC GATTTCACTA TCAACAGCCT GTATTACAGC

481 GTAGCGGATT TTACCGTCCG TGATTACGTT GGCGGCATGA AGGATCTGAA GGACGGCGTT

541 ATCCGTCTGA TTGGTAACCC GGAAACGCGC TACCGTGAAG ATCCGGTACG TATGCTGCGC

601 GCGGTACGTT TTGCCGCCAA ATTGGGTATG CGCATCAGCC CGGAAACCGC AGAACCGATC

661 CCTCGCCTCG CTACCCTGCT GAACGATATC CCACCGGCAC GCCTGTTTGA AGAATCGCTT

721 AAACTGCTAC AAGCGGGCTA CGGTTACGAA ACCTATAAGC TGTTGTGTGA ATATCATCTG

781 TTCCAGCCGC TGTTCCCGAC CATTACCCGC TACTTCACGG AAAATGGCGA CAGCCCGATG

841 GAGCGGATCA TTGAACAGGT GCTGAAGAAT ACCGATACGC GTATCCATAA CGATATGCGC

901 GTGAACCCGG CGTTCCTGTT TGCCGCCATG TTCTGGTACC CACTGCTGGA GACGGCACAG

961 AAGATCGCCC AGGAAAGCGG CCTGACCTAT CACGACGCTT TCGCGCTGGC GATGAACGAC

1021 GTGCTGGACG AAGCCTGCCG TTCACTGGCA ATCCCGAAAC GTCTGACGAC ATTAACCCGC

1081 GATATCTGGC AGTTGCAGTT GCGTATGTCC CGTCGTCAGG GTAAACGCGC ATGGAAACTG

1141 CTGGAGCATC CTAAGTTCCG TGCGGCTTAT GACCTGTTGG CCTTGCGAGC TGAAGTTGAG

1201 CGTAACGCTG AACTGCAGCG TCTGGTGAAA TGGTGGGGTG AGTTCCAGGT TTCCGCGCCA

```
    1261  CCAGACCAAA AAGGGATGCT CAACGAGCTG GATGAAGAAC CGTCACCGCG TCGTCGTACT
    1321  CGTCGTCCAC GCAAACGCGC ACCACGTCGT GAGGGTACCG CATGA
```

***rpoC***

```
       1  gtgAAAGATT TATTAAAGTT TCTGAAAGCG CAGACTAAAA CCGAAGAGTT TGATGCGATC
      61  AAAATTGCTC TGGCTTCGCC AGACATGATC CGTTCATGGT CTTTCGGTGA AGTTAAAAAG
```

121 CCGGAAACCA TCAACTACCG TACGTTCAAA CCAGAACGTG ACGGCCTTTT CTGCGCCCGT

181 ATCTTTGGGC CGGTAAAAGA TTACGAGTGC CTGTGCGGTA AGTACAAGCG CCTGAAACAC

241 CGTGGCGTCA TCTGTGAGAA GTGCGGCGTT GAAGTGACCC AGACTAAAGT ACGCCGTGAG

301 CGTATGGGCC ACATCGAACT GGCTTCCCCG ACTGCGCACA TCTGGTTCCT GAAATCGCTG

361 CCGTCCCGTA TCGGTCTGCT GCTCGATATG CCGCTGCGCG ATATCGAACG CGTACTGTAC

421 TTTGAATTCT ATGTGGTTAT CGAAGGCGGT ATGACCAACC TGGAACGTCA GCAGATCCTG

481 ACTGAAGAGC AGTATCTGGA CGCGCTGGAA GAGTTCGGTG ACGAATTCGA CGCGAAGATG

541 GGGGCGGAAG CAATCCAGGC TCTGCTGAAG AGCATGGATC TGGAGCAAGA GTGCGAACAG

601 CTGCGTGAAG AGCTGAACGA AACCAACTCC GAAACCAAGC GTAAAAAGCT GACCAAGCGT

661 ATCAAACTGC TGGAAGCGTT CGTTCAGTCT GGTAACAAAC CAGAGTGGAT GATCCTGACC

721 GTTCTGCCGG TACTGCCGCC AGATCTGCGT CCGCTGGTTC CGCTGGATGG TGGTCGTTTC

781 GCGACTTCTG ACCTGAACGA TCTGTATCGT CGCGTCATTA ACCGTAACAA CCGTCTGAAA

841 CGTCTGCTGG ATCTGGCTGC GCCGGACATC ATCGTACGTA ACGAAAAACG TATGCTGCAG

901 GAAGCGGTAG ACGCCCTGCT GGATAACGGT CGTCGCGGTC GTGCGATCAC CGGTTCTAAC

961 AAGCGTCCTC TGAAATCTTT GGCCGACATG ATCAAAGGTA AACAGGGTCG TTTCCGTCAG

1021 AACCTGCTCG GTAAGCGTGT TGACTACTCC GGTCGTTCTG TAATCACCGT AGGTCCATAC

1081 CTGCGTCTGC ATCAGTGCGG TCTGCCGAAG AAAATGGCAC TGGAGCTGTT CAAACCGTTC

1141 ATCTACGGCA AGCTGGAACT GCGTGGTCTT GCTACCACCA TTAAAGCTGC GAAGAAAATG

1201 GTTGAGCGCG AAGAAGCTGT CGTTTGGGAT ATCCTGGACG AAGTTATCCG CGAACACCCG

1261 GTACTGCTGA ACCGTGCACC GACTCTGCAC CGTCTGGGTA TCCAGGCATT TGAACCGGTA

1321 CTGATCGAAG GTAAAGCTAT CCAGCTGCAC CCGCTGGTTT GTGCGGCATA TAACGCCGAC

1381 TTCGATGGTG ACCAGATGGC TGTTCACGTA CCGCTGACGC TGGAAGCCCA GCTGGAAGCG

1441 CGTGCGCTGA TGATGTCTAC CAACAACATC CTGTCCCCGG CGAACGGCGA ACCAATCATC

1501 GTTCCGTCTC AGGACGTTGT ACTGGGTCTG TACTACATGA CCCGTGACTG TGTTAACGCC

1561 AAAGGCGAAG GCATGGTGCT GACTGGCCCG AAAGAAGCAG AACGTCTGTA TCGCTCTGGT

1621 CTGGCTTCTC TGCATGCGCG CGTTAAAGTG CGTATCACCG AGTATGAAAA AGATGCTAAC

1681 GGTGAATTAG TAGCGAAAAC CAGCCTGAAA GACACGACTG TTGGCCGTGC CATTCTGTGG

1741 ATGATTGTAC CGAAAGGTCT GCCTTACTCC ATCGTCAACC AGGCGCTGGG TAAAAAAGCA

1801 ATCTCCAAAA TGCTGAACAC CTGCTACCGC ATTCTCGGTC TGAAACCGAC CGTTATTTTT

1861 GCGGACCAGA TCATGTACAC CGGCTTCGCC TATGCAGCGC GTTCTGGTGC ATCTGTTGGT

1921 ATCGATGACA TGGTCATCCC GGAGAAGAAA CACGAAATCA TCTCCGAGGC AGAAGCAGAA

1981 GTTGCTGAAA TTCAGGAGCA GTTCCAGTCT GGTCTGGTAA CTGCGGGCGA ACGCTACAAC

2041 AAAGTTATCG ATATCTGGGC TGCGGCGAAC GATCGTGTAT CCAAAGCGAT GATGGATAAC

2101 CTGCAAACTG AAACCGTGAT TAACCGTGAC GGTCAGGAAG AGAAGCAGGT TTCCTTCAAC

2161 AGCATCTACA TGATGGCCGA CTCCGGTGCG CGTGGTTCTG CGGCACAGAT TCGTCAGCTT

2221 GCTGGTATGC GTGGTCTGAT GGCGAAGCCG GATGGCTCCA TCATCGAAAC GCCAATCACC

2281 GCGAACTTCC GTGAAGGTCT GAACGTACTC CAGTACTTCA TCTCCACCCA CGGTGCTCGT

2341 AAAGGTCTGG CGGATACCGC ACTGAAAACT GCGAACTCCG GTTACCTGAC TCGTCGTCTG

2401 GTTGACGTGG CGCAGGACCT GGTGGTTACC GAAGACGATT GTGGTACCCA TGAAGGTATC

2461 ATGATGACTC CGGTTATCGA GGGTGGTGAC GTTAAAGAGC CGCTGCGCGA TCGCGTACTG

2521 GGTCGTGTAA CTGCTGAAGA CGTTCTGAAG CCGGGTACTG CTGATATCCT CGTTCCGCGC

2581 AACACGCTGC TGCACGAACA GTGGTGTGAC CTGCTGGAAG AGAACTCTGT CGACGCGGTT

2641 AAAGTACGTT CTGTTGTATC TTGTGACACC GACTTTGGTG TATGTGCGCA CTGCTACGGT

2701 CGTGACCTGG CGCGTGGCCA CATCATCAAC AAGGGTGAAG CAATCGGTGT TATCGCGGCA

2761 CAGTCCATCG GTGAACCGGG TACACAGCTG ACCATGCGTA CGTTCCACAT CGGTGGTGCG

2821 GCATCTCGTG CGGCTGCTGA ATCCAGCATC CAAGTGAAAA ACAAAGGTAG CATCAAGCTC

2881 AGCAACGTGA AGTCGGTTGT GAACTCCAGC GGTAAACTGG TTATCACTTC CCGTAATACT

2941 GAACTGAAAC TGATCGACGA ATTCGGTCGT ACTAAAGAAA GCTACAAAGT ACCTTACGGT

3001 GCGGTACTGG CGAAAGGCGA TGGCGAACAG GTTGCTGGCG GCGAAACCGT TGCAAACTGG

3061 GACCCGCACA CCATGCCGGT TATCACCGAA GTAAGCGGTT TTGTACGCTT TACTGACATG

3121 ATCGACGGCC AGACCATTAC GCGTCAGACC GACGAACTGA CCGGTCTGTC TTCGCTGGTG

3181 GTTCTGGATT CCGCAGAACG TACCGCAGGT GGTAAAGATC TGCGTCCGGC ACTGAAAATC

3241 GTTGATGCTC AGGGTAACGA CGTTCTGATC CCAGGTACCG ATATGCCAGC GCAGTACTTC

3301 CTGCCGGGTA AAGCGATTGT TCAGCTGGAA GATGGCGTAC AGATCAGCTC TGGTGACACC

3361 CTGGCGCGTA TTCCGCAGGA ATCCGGCGGT ACCAAGGACA TCACCGGTGG TCTGCCGCGC

3421 GTTGCGGACC TGTTCGAAGC ACGTCGTCCG AAAGAGCCGG CAATCCTGGC TGAAATCAGC

3481 GGTATCGTTT CCTTCGGTAA AGAAACCAAA GGTAAACGTC GTCTGGTTAT CACCCCGGTA

3541 GACGGTAGCG ATCCGTACGA AGAGATGATT CCGAAATGGC GTCAGCTCAA CGTGTTCGAA

3601 GGTGAACGTG TAGAACGTGG TGACGTAATT TCCGACGGTC CGGAAGCGCC GCACGACATT

3661 CTGCGTCTGC GTGGTGTTCA TGCTGTTACT CGTTACATCG TTAACGAAGT ACAGGACGTA

3721 TACCGTCTGC AGGGCGTTAA GATTAACGAT AAACACATCG AAGTTATCGT TCGTCAGATG

3781 CTGCGTAAAG CTACCATCGT TAACGCGGGT AGCTCCGACT TCCTGGAAGG CGAACAGGTT

3841 GAATACTCTC GCGTCAAGAT CGCAAACCGC GAACTGGAAG CGAACGGCAA AGTGGGTGCA

3901 ACTTACTCCC GCGATCTGCT GGGTATCACC AAAGCGTCTC TGGCAACCGA GTCCTTCATC

3961 TCCGCGGCAT CGTTCCAGGA GACCACTCGC GTGCTGACCG AAGCAGCCGT TGCGGGCAAA

4021 CGCGACGAAC TGCGCGGCCT GAAAGAGAAC GTTATCGTGG GTCGTCTGAT CCCGGCAGGT

4081 ACCGGTTACG CGTACCACCA GGATCGTATG CGTCGCCGTG CTGCGGGTGA AGCTCCGGCT

```
    4141  GCACCGCAGG TGACTGCAGA AGACGCATCT GCCAGCCTGG CAGAACTGCT GAACGCAGGT
    4201  CTGGGCGGTT CTGATAACGA Gtaa
```

**Figure S1.** **Fermentation pathways of *E. coli* and gene knockouts.** Major fermentation pathways of *E. coli* and the five gene loci deleted in the DH1Δ*5* strain (Δ*ackA-pta* Δ*adhE* Δ*ldhA* Δ*poxB* Δ*frdBC*).

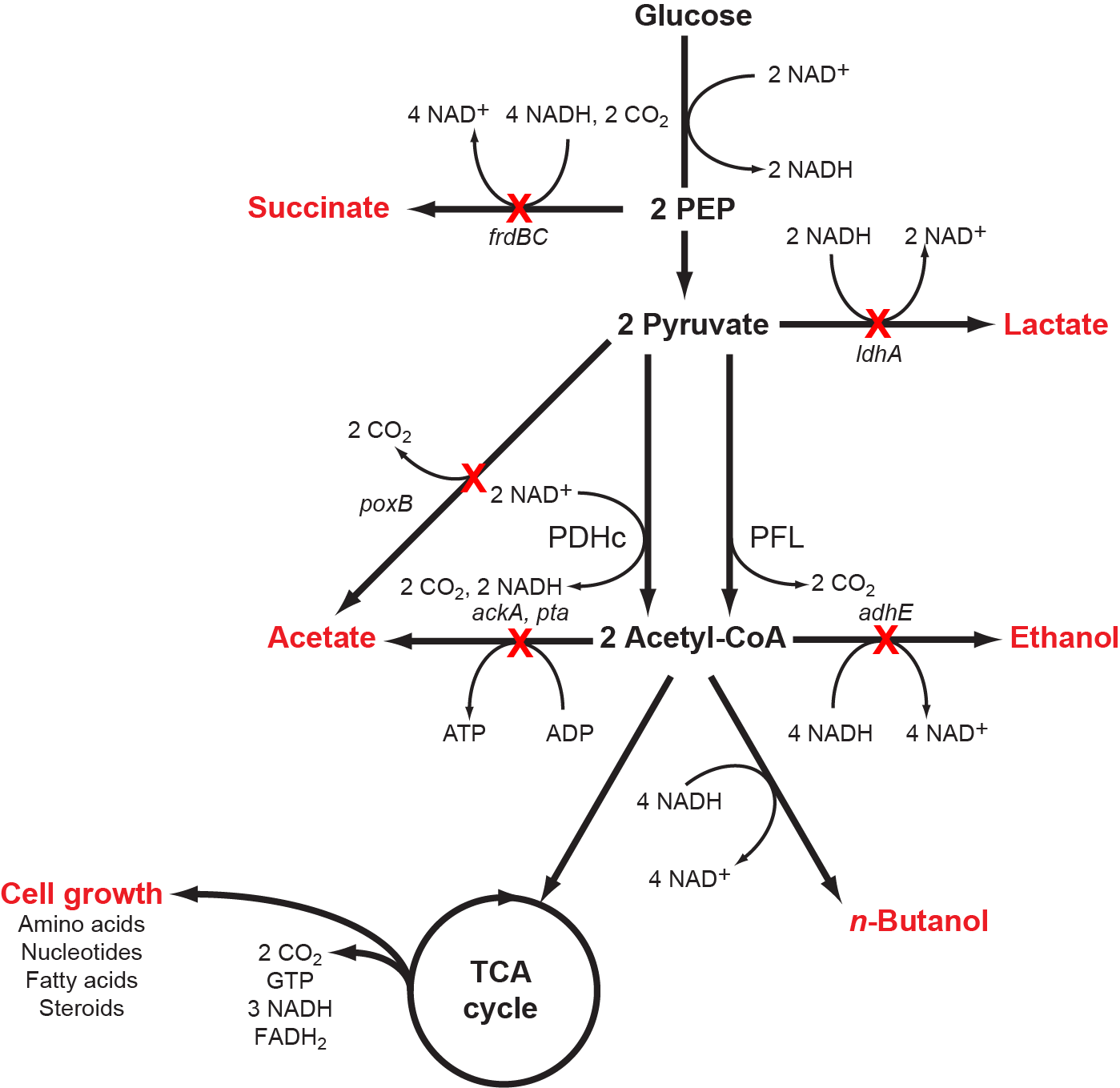

**Figure S2.** **Dependence of the *n*-butanol:ethanol ratio on oxygen.** When DH1 transformed with the first generation *n*-butanol pathway (pBT33-Bu2 pCWori-ter.adhE2 pBBR1-aceEF.lpd) is cultured under anaerobic conditions, the level of *n*-butanol drops but the level of ethanol remains similar. This result suggests that ethanol produced via AdhE2 can support cell growth under anaerobic conditions. Cultures were grown in TB with 2.5% (*w*/*v)* glucose media. Titers from aerobic cultures were analyzed via GC-FID after 3 d of growth and after 5 d for anaerobic cultures. Data are mean ± s.d. of biological replicates (n = 3).

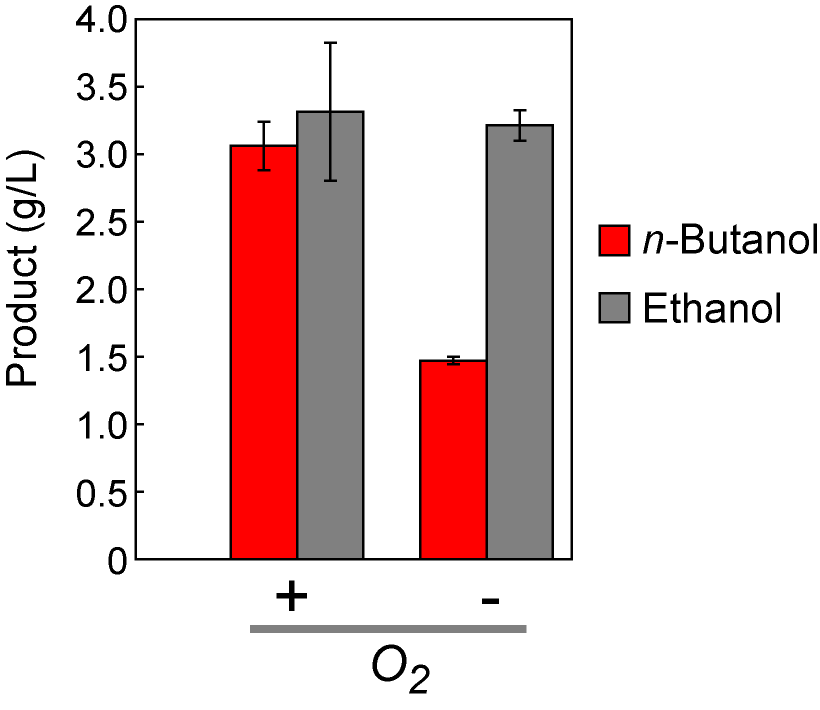

**Figure S3.** **Biochemical characterization of AdhE2 constructs.** The different domains of the bifunctional AdhE2 from *Clostridium acetobutylicum* were characterized biochemically using 5 different constructs and 3 different assays. (A) AdhE2 constructs consisted of domain-inactivated mutants of either the ALDH or ADH domain. Single domain constructs were also generated by truncation. (*, inactivated). (B) Representative SDS-PAGE of Strep tag affinity purification of full-length AdhE2s: Pre-induction (lane 1), post-induction (lane 2), soluble lysate (lane 3), insoluble lysate (lane 4), flow-through (lane 5), eluted AdhE2 (lane 6). (C) Wt AdhE2 was characterized by monitoring NADH oxidation in the presence of either an acyl-CoA (ALDH and ADH activity measured) or aldehyde (ADH activity measured). Activity was also measured by CoA release (ALDH activity measured). Domain inactivated and single domain constructs could be characterized directly by the NADH oxidation assay with addition of the appropriate substrate. *k*_cat_ and *K*_M_ were obtained by curve-fitting to the Michaelis–Menten parameters. Data are mean ± s.e. (n = 3) as derived from nonlinear curve fitting to the Michaelis−Menten equation. Error in the relative *k*_cat_/*K*_M_ was obtained from propagation.was characterized using constructs.

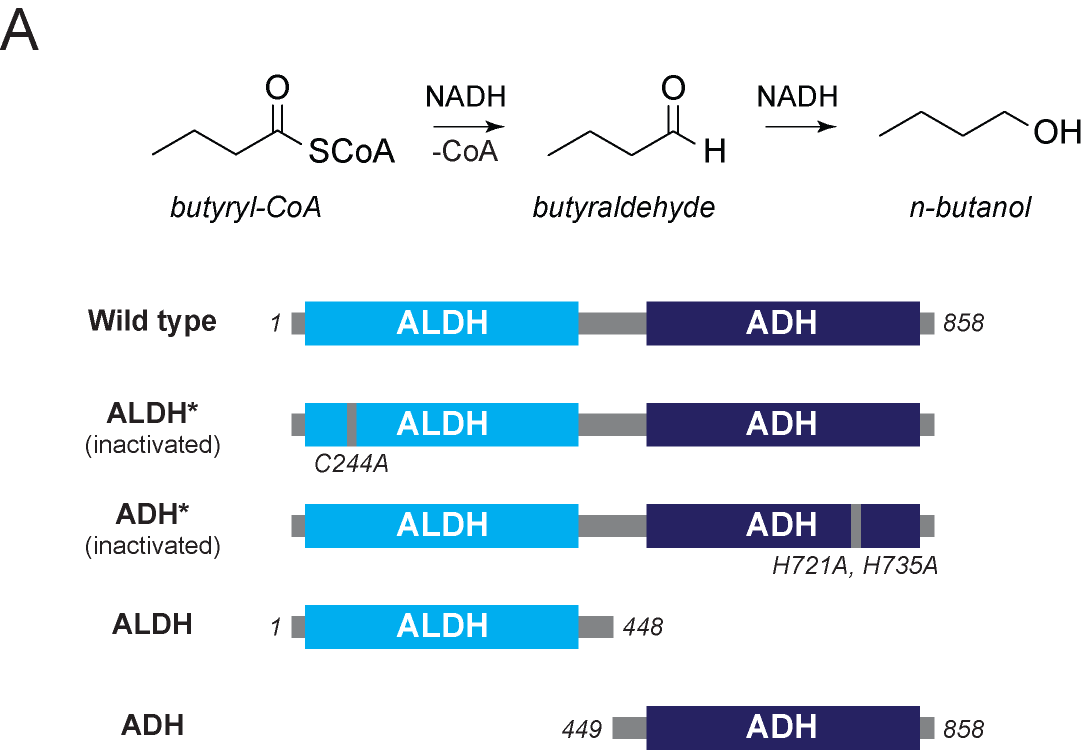

B

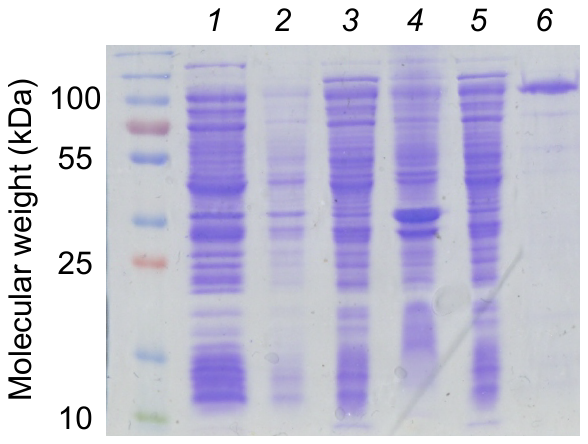

C

| **Enzyme** | **Substrate** | ***k*_cat_ (s^-1^)** | ***K*_M_ (µM)** | ***k*_cat_/*K*_M_ (M^-1^ s^-1^)** |
| --- | --- | --- | --- | --- |
| **WT AdhE2** | |  |  |  |
| NADH oxidation | butyryl-CoA | 2.2 ± 0.1 | 23 ± 2 | (9.6 ± 0.3) x 10^4^ |
| (both domains) | acetyl-CoA | 3.3 ± 0.1 | 250 ± 15 | (1.3 ± 0.1) x 10^4^ |
| CoA release | butyryl-CoA | 1.2 ± 0.1 | 10 ± 1 | (1.1 ± 0.1) x 10^5^ |
| (ALDH domain) | acetyl-CoA | 1.3 ± 0.1 | 100 ± 10 | (1.3 ± 0.2) x 10^4^ |
| Aldehyde substrate | butyraldehyde | 2.9 ± 0.1 | 4000 ± 400 | (7.0 ± 0.2) x 10^2^ |
| (ADH domain) | acetaldehyde | 5.6 ± 0.1 | 4500 ± 300 | (1.2 ± 0.1) x 10^3^ |
| **Domain mutants** | |  |  |  |
| ALDH inactive | butyraldehyde | 18.7 ± 1.3 | 2500 ± 500 | (7.5 ± 0.7) x 10^3^ |
|  | acetaldehyde | 19.8 ± 1.0 | 2800 ± 400 | (7.1 ± 0.3) x 10^3^ |
| ADH inactive | butyryl-CoA | 0.3 ± 0.1 | 4 ± 1 | (9.0 ± 1.0) x 10^4^ |
|  | acetyl-CoA | 1.3 ± 0.1 | 70 ± 10 | (1.9 ± 0.4) x 10^4^ |
| **Domain truncations** | |  |  |  |
| ALDH | butyryl-CoA | < 0.1 | n.d. | n.d. |
| ADH | butyraldehyde | 0.2 ± 0.1 | 300 ± 50 | (5.8 ± 0.4) x 10^2^ |
|  | acetaldehyde | 0.2 ± 0.1 | 2000 ± 700 | (1.1 ± 0.2) x 10^2^ |

**Figure S4. Phylogenetic analysis of ALDH domains.** (A) The biochemical literature was surveyed for ALDH domains characterized to have preference for acyl-CoA substrates larger than acetyl-CoA. Both monofunctional and bifunctional ALDH domains were considered, including the atypical ALDH DmpF that is a fusion with an aldolase domain. In the case of bifunctional ALDHs, only the specificity of the ALDH domain was considered. (B) A phylogenetic tree incorporating biochemically characterized ALDH domains was assembled to identify homologs that may have greater than C_4_ substrate specificity. Sequences in purple have higher *k*_cat_/*K*_M_ for acyl-CoAs larger than acetyl-CoA, and sequences in green have higher *k*_cat_/*K*_M_ for acetyl-CoA. (C) The branching pattern of the biochemically informed tree was applied to all sequences in the ALDH family. Green sequences denote ALDH domains of bifunctional enzymes and red sequences denote monofunctional ALDH domains.

A

| **Preference** | **Gene** | **Accession No.** | **Organism** | **Reference** |
| --- | --- | --- | --- | --- |
| **C_4_ substrates** | AdhE2 | NP_149199 | *Clostridium acetobutylicum* | This study |
|  | ALD | AAD31841 | *Clostridium beijerinckii* | J Toth *et al* [*16*] |
|  | ALDH | YP_001310903 | *Clostridium beijerinckii* | RT Yan *et al* [*17*] |
|  | PduP | BAG26139 | *Lactobacillus reuteri* | LH Luo *et al* [*18*] |
|  | ALDH | YP_003687877 | *Propionibacterium freudenreichii* | N Hosio *et al* [*19*] |
| **C_2_ substrates** | AdhE | NP_415757 | *Escherichia coli* | S Atsumi *et al* [*20*] |
|  | AdhE | AAC47539 | *Giardia intestinalis* | LB Sánchez *et al* [*21*] |
|  | AdhE | AAV66076 | *Leuconostoc mesenteroides* | OK Koo *et al* [*22*] |
|  | DmpF | CAA43226 | *Pseudomonas sp.* CF600 | J Powlowski *et al* [*23*] |

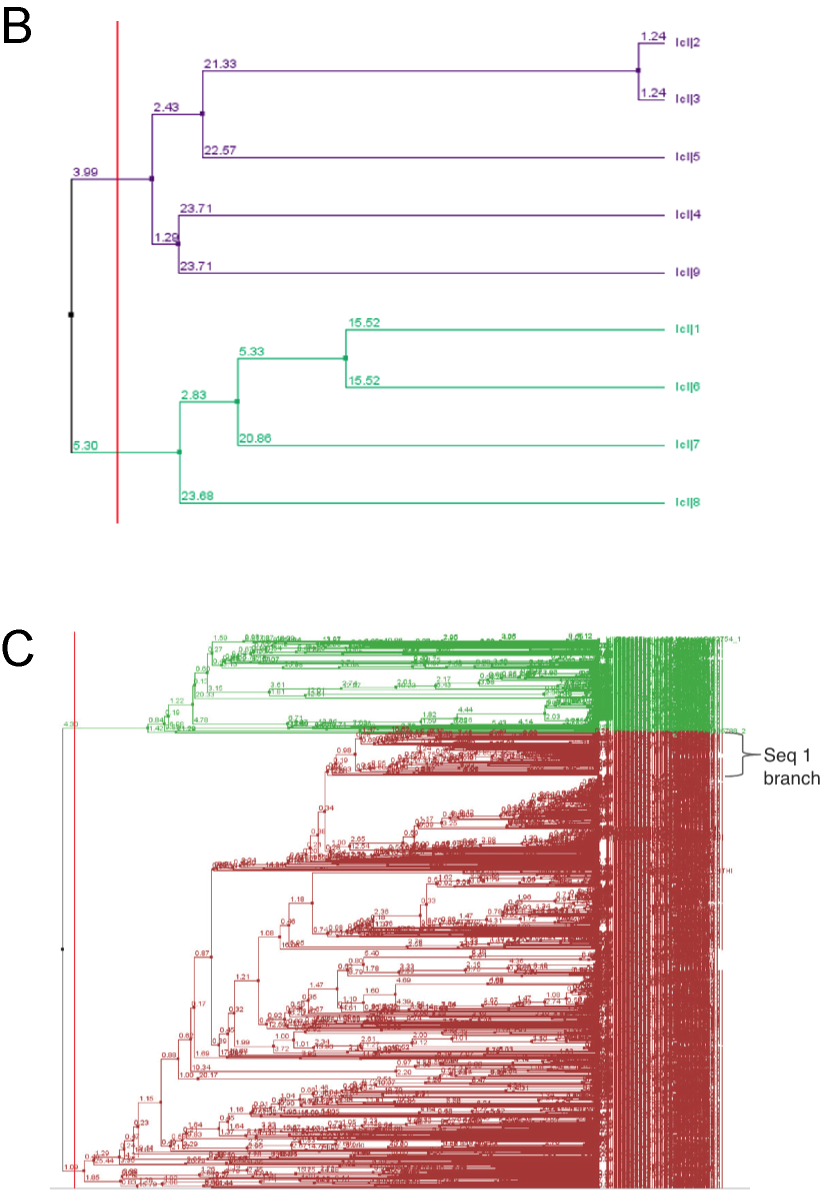

**Figure S5. Screening of AdhE2s.** AdhEs were screened *in vivo* by co-transforming DH1 pBT33-Bu2 [*6*] and the pCDF3-ter.AdhE2 variants 1–96 or homologs 29–43. All strains were grown anaerobically in TB with 2.5% (*w*/*v)* glucose media for 3 d post induction before production titers were analyzed by GC-FID. (A) An initial panel of 96 variants of AdhE2 (AdhE2.1-96) was designed by Calysta to incorporate approximately 40 mutations selected from the natural sequence diversity of AdhE2 homologs. Each variant contained 3–5 mutations and every mutation was present in multiple variants. Around half of the variants produced minimal *n*-butanol, indicating that particular combinations of mutations resulted in non-functional enzymes. The remaining variants produced a wide range of titers in comparison to the wild type sequence. Some variants produced higher *n*-butanol titers, but upon further inspection, exhibited only mild improvements in *n*-butanol:ethanol ratio, suggesting the primary effect was an overall activity increase without shifting substrate specificity. Data are mean ± s.d. of biological replicates (n = 3). (B) AdhE2 homologs (AdhE29-43) sampled from the putative C_4_ branch of the phylogenetic tree. Comparison of *n*-butanol and ethanol titers do not show any increased selectivity compared to WT (AdhE2 from *C. acetobutylicum*). Data are mean ± s.d. of biological replicates (n = 3).

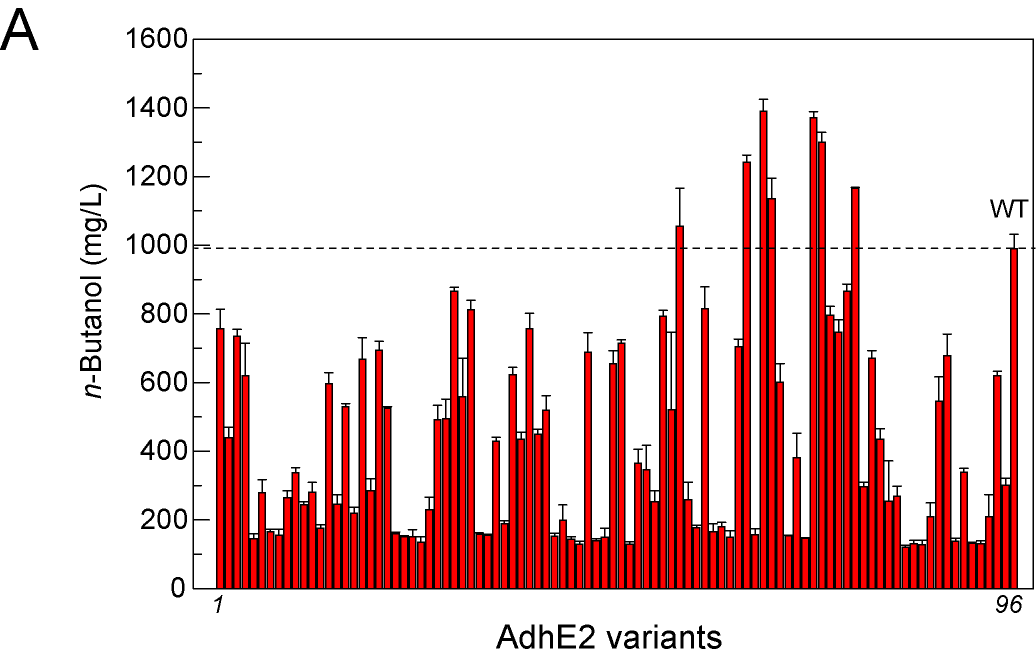

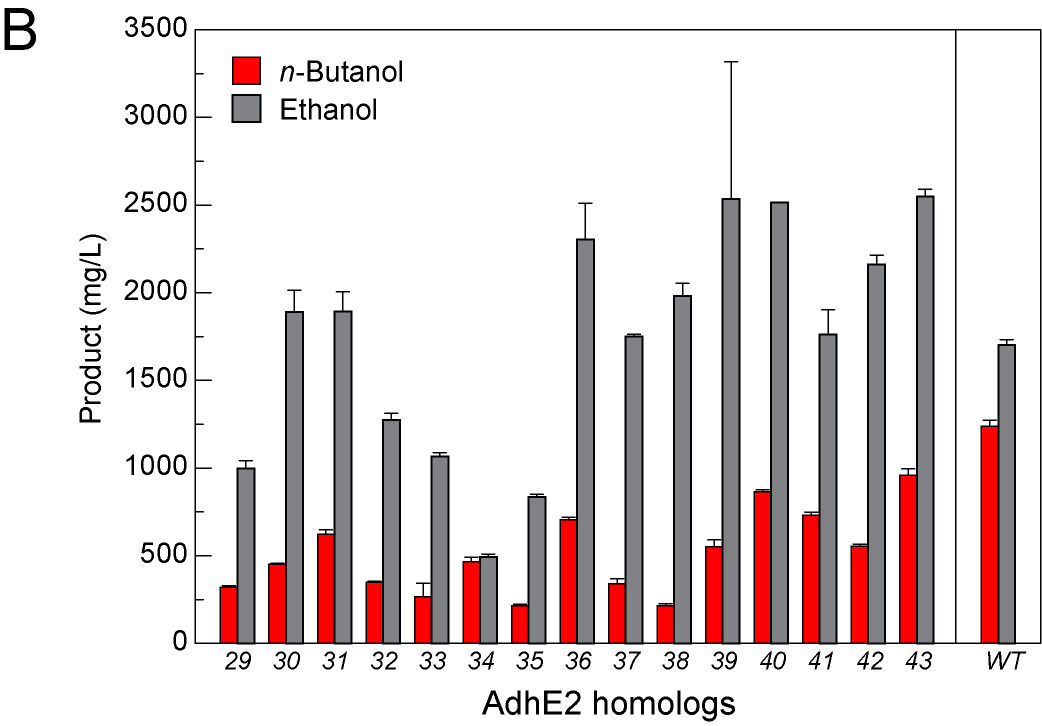

**Figure S6. Screening of monofunctional ALDHs for C_4_ selectivity.** Using the existing ALDH phylogenetic tree, the hypothetical C_4_-selective branch was widely sampled for monofunctional ALDHs to incorporate the full diversity of the branch in a small number of sequences. These sequences (ALDH1-15 and 46) were screened *in vivo* by co-transforming DH1Δ*5* pBT33-Bu2 and the pCDF3-ter.ALDH plasmid. All strains were grown anaerobically in TB with 2.5% (*w*/*v)* glucose media for 3 d post induction before analyzing production titers by GC-FID. 15 of 16 monofunctional ALDHs produced more *n*-butanol than ethanol. Data are mean ± s.d. of biological replicates (n = 3).

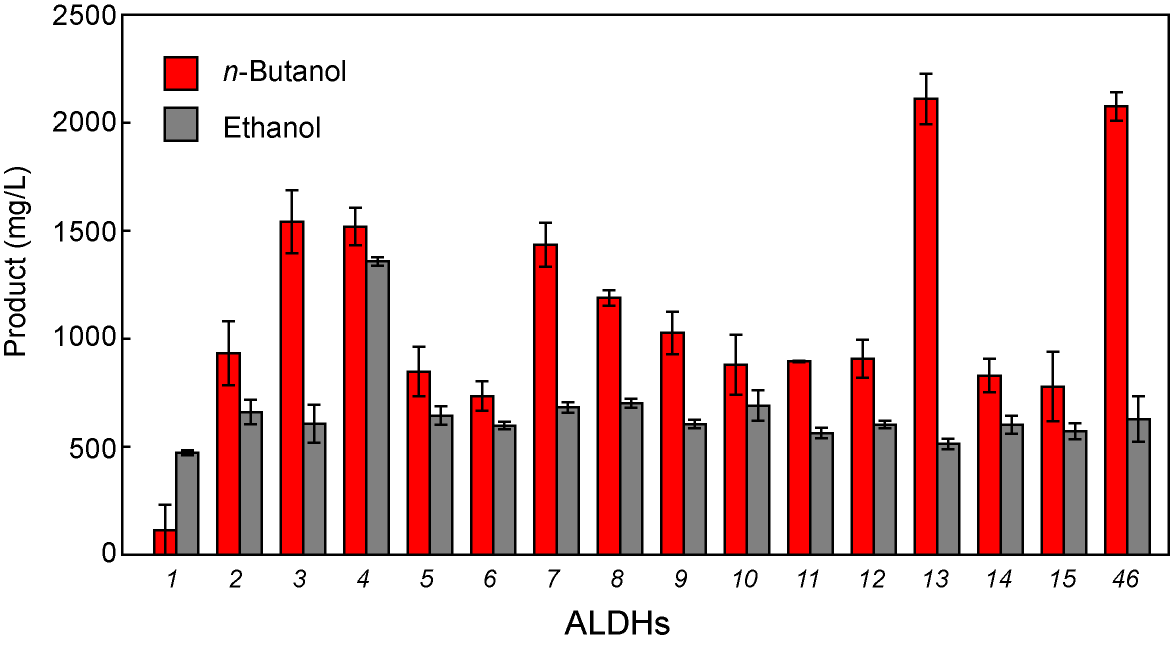

**Figure S7. Screening monofunctional ALDHs for production of 1,3-butanediol (BDO).** (A) The monofunctional ALDHs (ALDH1-16 and 46) were also screened *in vivo* for their ability to produce BDO. DH1Δ*5* was co-transformed with pBT33-Bu2 and the pCDF3-ter.ALDH plasmid for *n*-butanol production or pT5T33-phaA.hbd and pCDF3-ALDH for BDO production. (B) The stereochemical preference of the ALDHs were then tested by screening for the production of (3*R*)- compared to (3*S*)-BDO by using either pT5T33-phaA.phaB or pT5T33-phaA.hbd upstream of the ALDH enzyme on pCDF3-ALDH to produce either (*R*)-3-hydroxybutyryl-CoA or (*S*)-3-hydroxybutyryl-CoA, respectively. These data suggest that little stereochemical preference at the ALDH is observed. All strains were grown anaerobically in TB with 2.5% (*w*/*v*) glucose media for 3 d post induction before analyzing production titers. Screening was performed using GC-MS for product peak identification. The same method as described for *n*-butanol anlysis was used and the amount of product was quantified by comparison to a standard. Data are mean ± s.d. of biological replicates (n = 3).

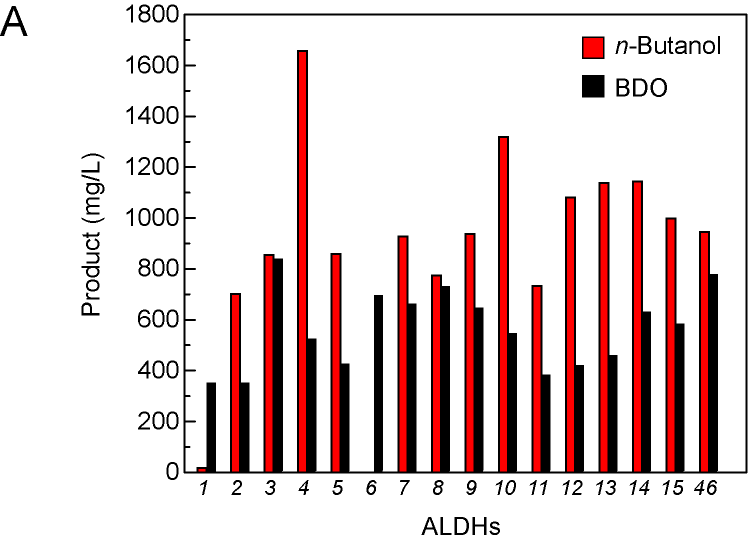

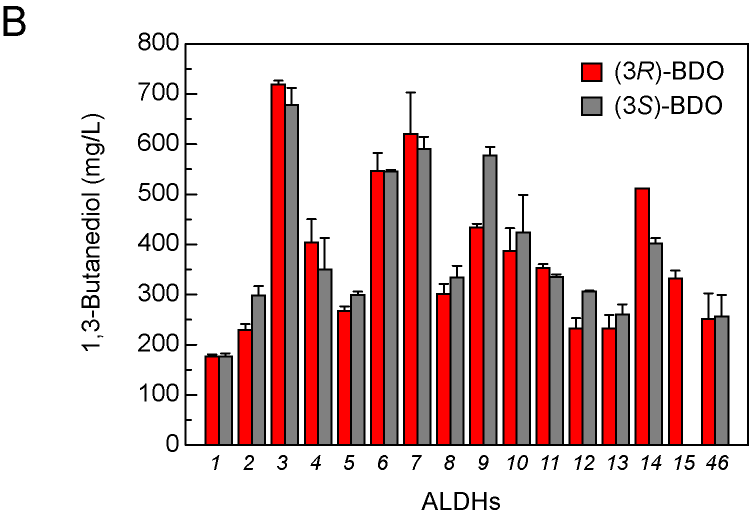

**Figure S8. Identification and screening of ADHs for BDO production.** (A) A sequence similarity network of the ADH family was generated to identify alcohol dehydrogenases that would efficiently reduce 3-hydroxybutyraldehyde to 1,3-butanediol. ADH sequences were blasted against each other and then clustered at increasingly stringent e-values, such that subfamilies become apparent. This network was then overlaid with ADHs of known substrate specificity as a frame of reference (BdhAB, butanol dehydrogenase; DhaT, 1,3-propanediol oxidoreductase; YqhD, <C_3_ aldehyde reductase) [*24–*26]. (B) Sequences from the network were broadly sampled to maximize diversity and increase the likelihood of identifying a highly active ADH. (C) ADHs were also screened *in vivo* under anaerobic conditions for their ability to produce BDO by co-transforming DH1Δ*5* pBBR1-aceEF.lpd pT533-phaA.phaB/hbd with a pCWori.trc-ter-aldh46.ADH plasmid (ADH1-22 and DhaT2-8). In this case, the ADHs appear to be selective for production of (3*R*)-BDO. All strains were grown anaerobically in TB with 2.5% (*w*/*v)* glucose media for 5 d post induction before analyzing production titers by GC-MS. Data are mean ± s.d. of biological replicates (n = 3).

A

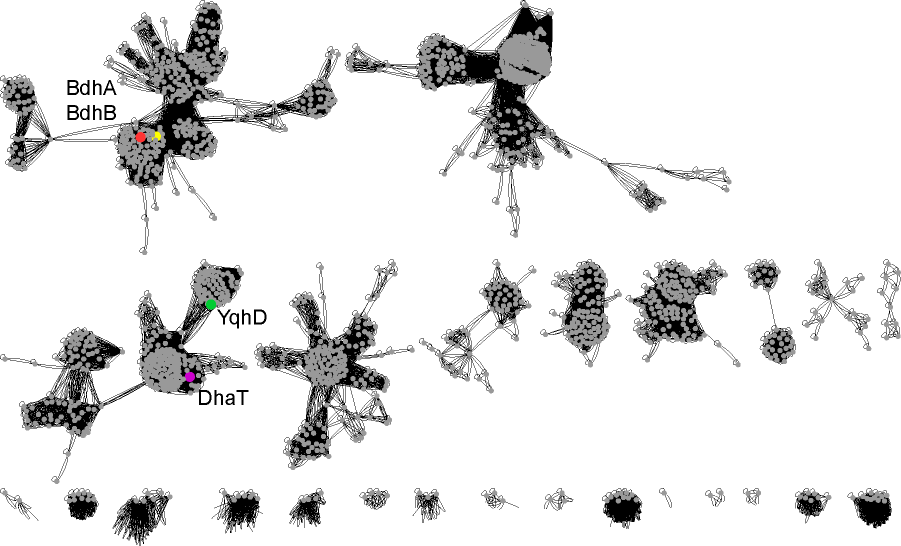

B

| **Gene** | **Accession No.** | **Organism** |
| --- | --- | --- |
| ADH1 | B6YQP9_AZOPC | *Azobacteroides pseudotrichonymphae genomovar* CFP2 |
| ADH2 | A0RQF7_CAMFF | *Campylobacter fetus* subsp. *fetus* 82-40 |
| ADH3 | G5F136_9ACTN | *Olsenella* sp. oral taxon 809 F0356 |
| ADH4 | B1C7G7_9FIRM | *Anaerofustis stercorihominis* DSM 17244 |
| ADH5 | YUGK_BACSU | *Bacillus subtilis* 168 |
| ADH6 | A8SGI9_9FIRM | *Faecalibacterium prausnitzii* M21/2 |
| ADH7 | E2SQ66_9FIRM | *Erysipelotrichaceae bacterium* 3_1_53 |
| ADH8 | E1QYZ8_OLSUV | *Olsenella uli* ATCC 49627 |
| ADH9 | F5X0G1_STRG1 | *Streptococcus gallolyticus* ATCC 43143 / F-1867 |
| ADH10 | E6W4G5_DESIS | *Desulfurispirillum indicum* ATCC BAA-1389 / S5 |
| ADH11 | E6K7W2_9BACT | *Prevotella buccae* ATCC 33574 |
| ADH12 | B1C4Z8_9FIRM | *Clostridium spiroforme* DSM 1552 |
| ADH13 | G4L3E3_TETHN | *Tetragenococcus halophilus* strain DSM 20338 |
| ADH14 | E8LLW8_9GAMM | *Succinatimonas hippei* YIT 12066 |
| DhaT2 | E4RKV2_HALSL | *Halanaerobium hydrogeniformans* |
| DhaT3 | Q15G22_CITFR | *Citrobacter freundii* |
| DhaT4 | A0PY50_CLONN | *Clostridium novyi* NT |
| DhaT5 | Q3A1K9_PELCD | *Pelobacter carbinolicus* DSM 2380 / Gra Bd 1 |
| DhaT6 | A5D4X5_PELTS | *Pelotomaculum thermopropionicum* DSM 13744 |
| DhaT7 | B1V2D9_CLOPF | *Clostridium perfringens* D JGS1721 |
| DhaT8 | E3H9G9_ILYPC | *Ilyobacter polytropus* DSM 2926 / CuHBu1 |
| ADH15 | Q1JYE4_DESAC | *Desulfuromonas acetoxidans* DSM 684 |
| ADH16 | B5YIE2_THEYD | *Thermodesulfovibrio yellowstonii* ATCC 51303 |
| ADH17 | D2BSS7_DICD5 | *Dickeya dadantii* Ech586 |
| ADH18 | F0ERB1_HAEPA | *Haemophilus parainfluenzae* ATCC 33392 |
| ADH19 | G5IQ05_9ENTE | *Enterococcus saccharolyticus* 30_1 |
| ADH20 | B2V5D0_CLOBA | *Clostridium botulinum* Alaska E43 / Type E3 |
| ADH21 | E2SME8_9FIRM | *Erysipelotrichaceae* sp. 3_1_53 |
| ADH22 | B0NYL0_9CLOT | *Clostridium* sp. SS2/1 |

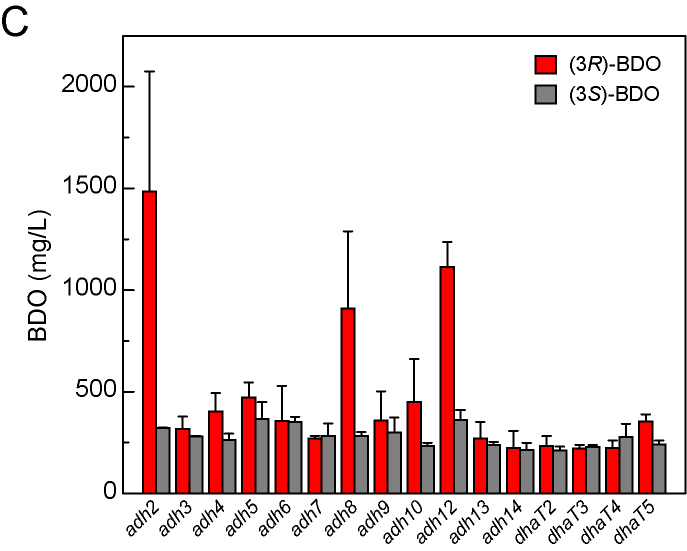

**Figure S9. Optimizing ALDH-ADH pairs for BDO vs. 4-hydroxybutanone (HB) production.** Having identified a number of ALDHs and ADHs competent for high titer butanediol production, we screened the combinatorial set of candidate enzymes to find optimal combinations. (A) A preliminary *in vivo* screen of DH1Δ*5* pT533-phaA.phaB pCWori.trc-ter-ALDH.ADH identified *aldh7.adh2* as the best overall performer for both BDO and HB production. Strains were grown anaerobically in TB with 2.5% (*w*/*v)* glucose media for 2 d post induction before analyzing production titers by LC-MS. Data are mean ± s.d. of biological replicates (n = 3). (B) To further optimize pathway selectivity, additional ADHs from the sequence similarity network were sampled to identify enzymes with greater specificity that would not enable hydroxybutanone production. The subfamilies containing ADH2, 8, and 12 were sampled at greater depth as these ADHs were shown to be most active in the initial screen. These screen identified a new optimal pair, *aldh3.adh22*, capable of capturing a large fraction of the C_4_ product pool as butanediol and producing 3 g/L of total products. Strains were grown anaerobically in TB with 2.5% (*w*/*v)* glucose media for 3 d post induction before production titers were analyzed by LC-MS. Data are mean ± s.d. of biological replicates (n = 3).

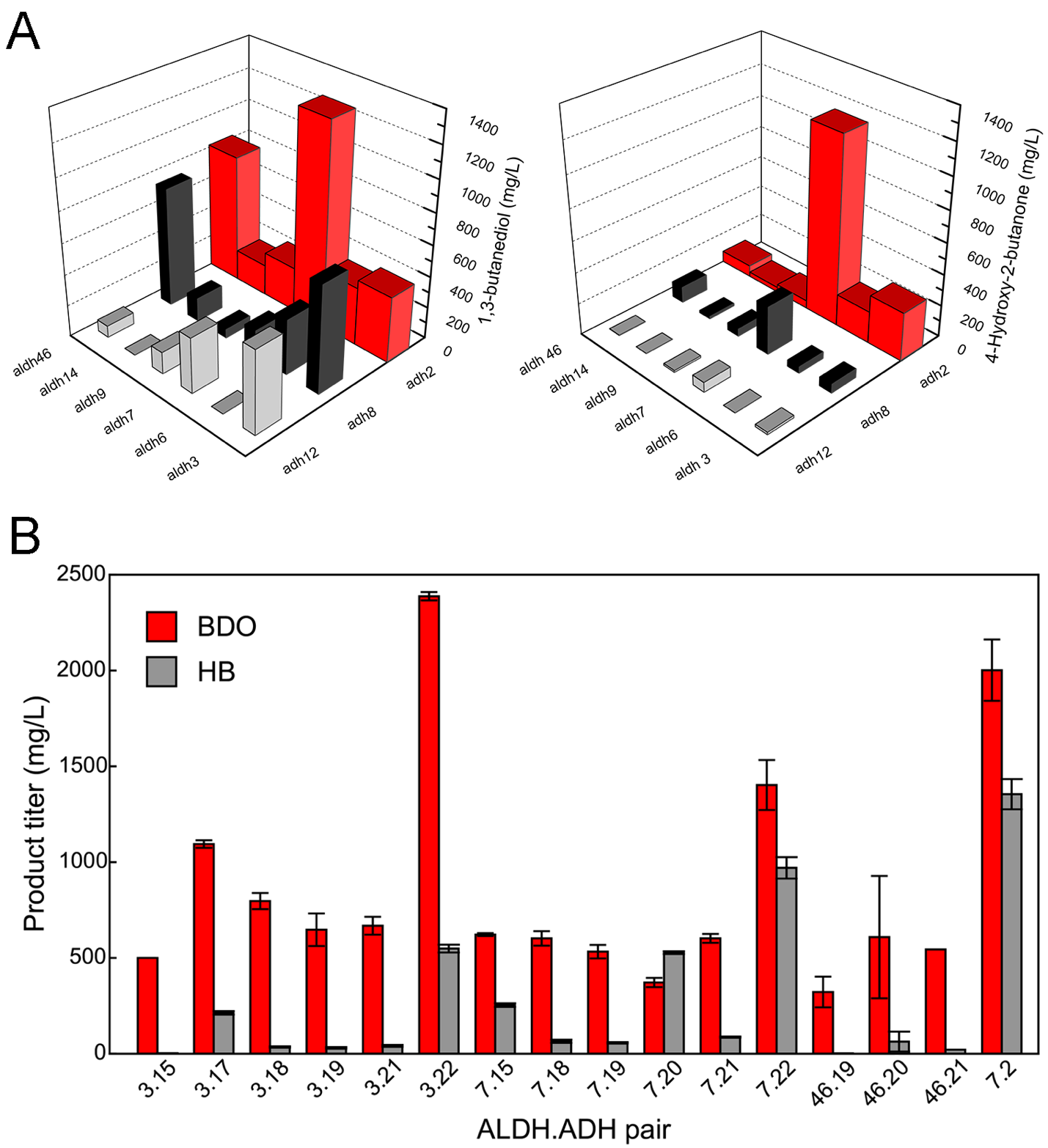

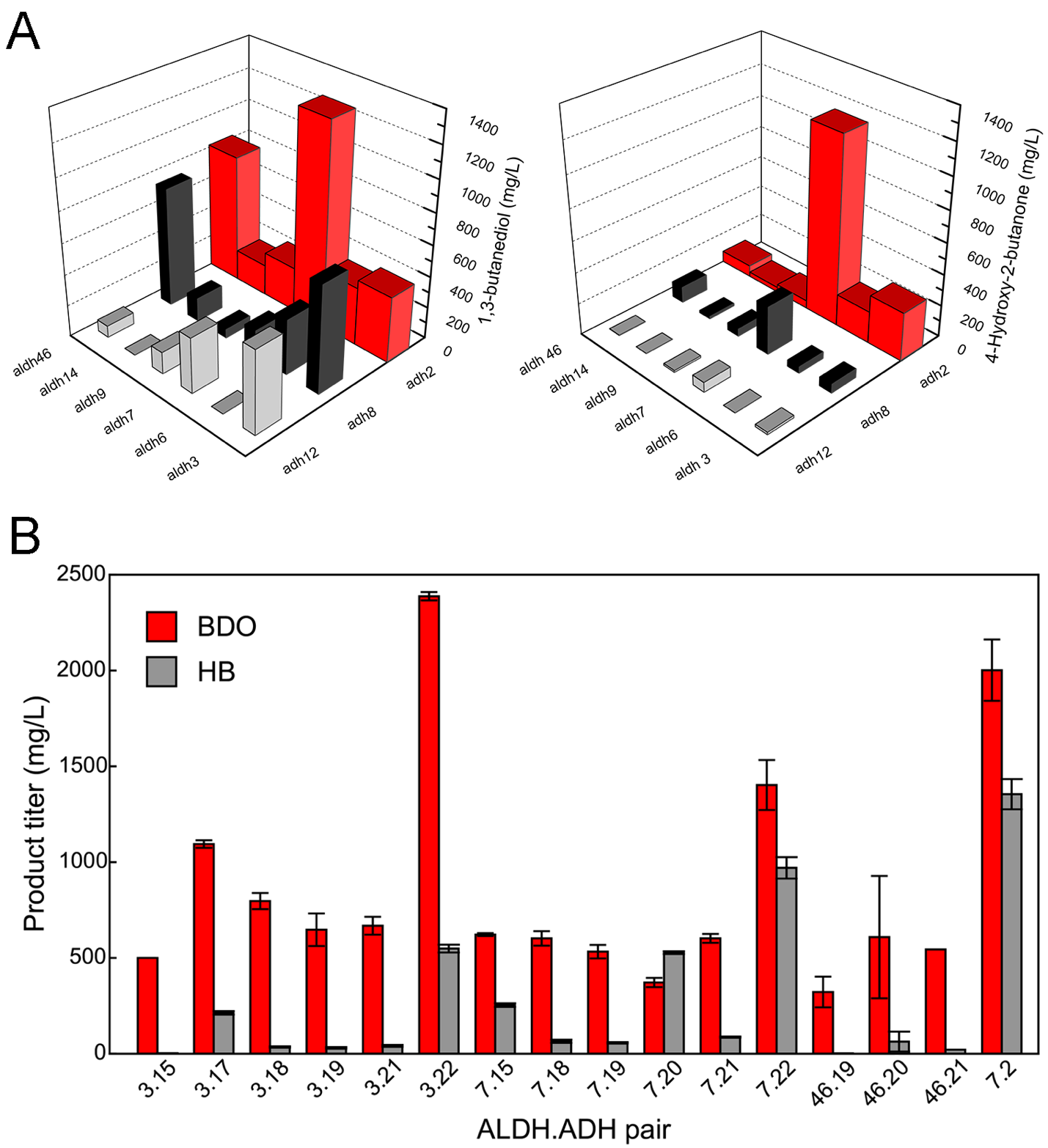

**Figure S10. Screening of secondary ADHs (sADHs) to increase BDO selectivity.** (A) A strategy for increasing the selectivity of BDO production is to use a secondary alcohol dehydrogenase (sADH) to reduce HB to BDO. (B) A list of sADHs that were either reported to reduce 4-hydroxy-2-butanone to 1,3-butanediol or have broad specificity for similar substrates was identified. A substantial number of these enzymes has been reported in bacteria, yeast, and parasitic protozoa. These enzymes are generally classified as zinc or iron-alcohol dehydrogenases and maximum percent identity within the sequences represented here range from 27–76%. (C) The identified sADHs were screened *in vivo* to characterize their BDO:HB selectivity by co-transforming DH1Δ*5* pBBR1-aceEF.lpd pT533-phaA.phaB with a pCWori.trc-sadhX-aldh7.adh2 plasmid (sADH1-15). The parent plasmid pCWori.trc-ter-aldh7.adh2 produced an even mixture of BDO and HB and served as the negative control. At least four sADHs enabled butanediol production of 2 g/L with hydroxybutanone production limited to <250 mg/L or less. This pathway design appears to maximize BDO production as the dead-end pathway from acetoacetyl-CoA reduction is eliminated. Strains were grown anaerobically in TB with 2.5% (*w*/*v)* glucose media for 4 d post induction before analyzing production titers by LC-MS. Data are mean ± s.d. of biological replicates (n = 3).

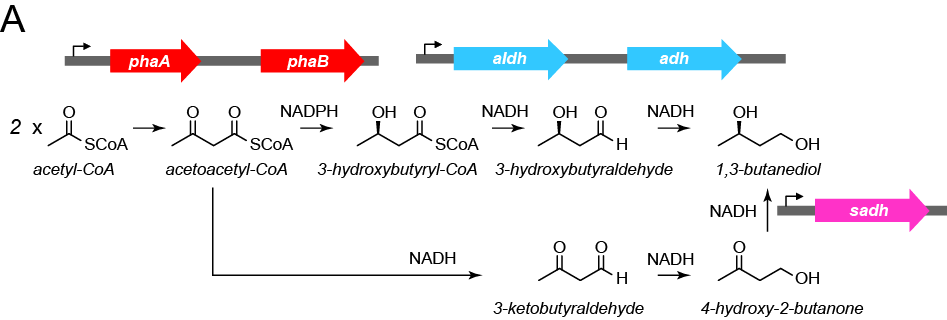

B

| **Gene** | **Accession No.** | **Organism** | **Reference** |
| --- | --- | --- | --- |
| sADH1 | KGK36767.1 | *Pichia kudriavzevii* | R. C. Zheng *et al* [*27*] |
| sADH2 | WP_011011186.1 | *Pyrococcus furiosus* DSM 3638 | R. Machielsen and J. Oost *et al* [*28, 29*] |
| sADH3 | WP_011614641.1 | *Cupriavidus necator* | A. Steinbüchel *et al* [30] |
| sADH4 | P14941.1 | *Thermoanaerobacter brockii* | R. J. Lamed *et al* [*31*] |
| sADH5 | AAA23199.2 | *Clostridium beijerinckii* | A. A. Ismaiel *et al* [*32*] |
| sADH6 | XP_455102.1 | *Kluyveromyces lactis* NRRL Y-1140 | T. Oda *et al* [*33*] |
| sADH7 | AAP39869.1 | *Phytomonas* sp. ADU-2003 | S. M. Molinas *et al* [*34*] |
| sADH8 | Q0KDL6.1 | *Ralstonia eutropha* H16 | A. Steinbüchel *et al* [*35*] |
| sADH9 | XP_001580601.1 | *Trichomonas vaginalis* G3 | R. Sutak and D. Leitsch *et al* [*36,37*] |
| sADH10 | AJP52792.1 | *Pseudomonas fluorescens* | C. T. Hou *et al* [*38*] |
| sADH11 | WP_011835462.1 | *Lactococcus lactis* | N. García-Quintáns *et al* [*39*] |
| sADH12 | AAC04974.1 | *Saccharomyces cerevisiae* | E. González *et al* [*40*] |
| sADH13 | WP_000374004.1 | *Escherichia coli* | H. Zhang *et al* [*41*] |
| sADH14 | BAD32689.1 | *Zygoascus ofunaensis* | K. Yamada Onodera *et al* [*42*] |
| sADH15 | BAA24528.1 | *Candida parapsilosis* | H. Yamamoto and H. Man *et al* [*43,44*] |
| sADH16 | BAN45671.1 | *Cyberlindnera jadinii* | T. Yang *et al* [*45*] |
| sADH17 | CAD36475.1 | *Rhodococcus ruber* | B. Kosjek *et al* [*46*] |

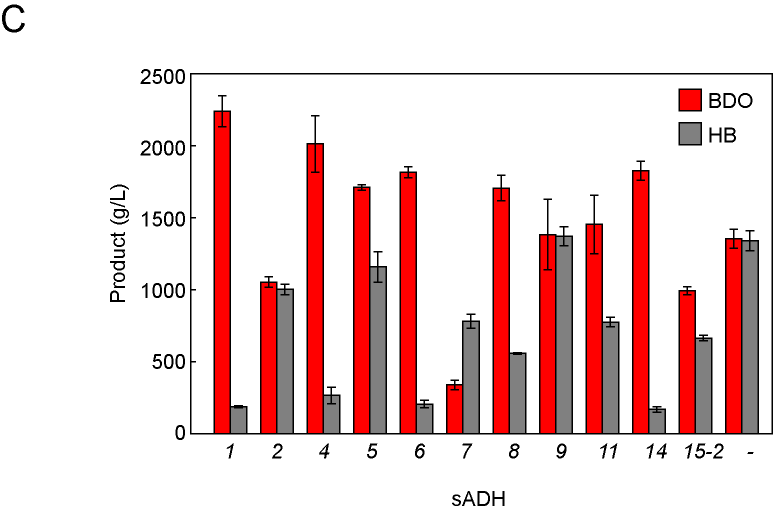

**Figure S11. Development of a genetic selection for *n*-butanol production.** Enrichment of higher productivity strains under anaerobic conditions. A small fraction of medium production strains (*E. coli* DH1Δ*5* pBT-0.3crt pCDF3-ter.aldh46) were mixed in a large excess of a low production strain (*E. coli* DH1Δ*5* pBT-0.03hbd pCDF3-ter.aldh46) to simulate a mutagenized library in which most mutations are neutral or deleterious. Over time the small fraction of medium production cells were found to proliferate and dominate the culture as monitored by qPCR (*Figure 3*). The length of the lag phase appears to correlate with the initial abundance of medium production cells. Strains were grown anaerobically in TB with 2.5% (*w*/*v)* glucose media over the course of 5 d, cultures containing the medium production strain were competent to attain high OD_600_ and butanol titer.

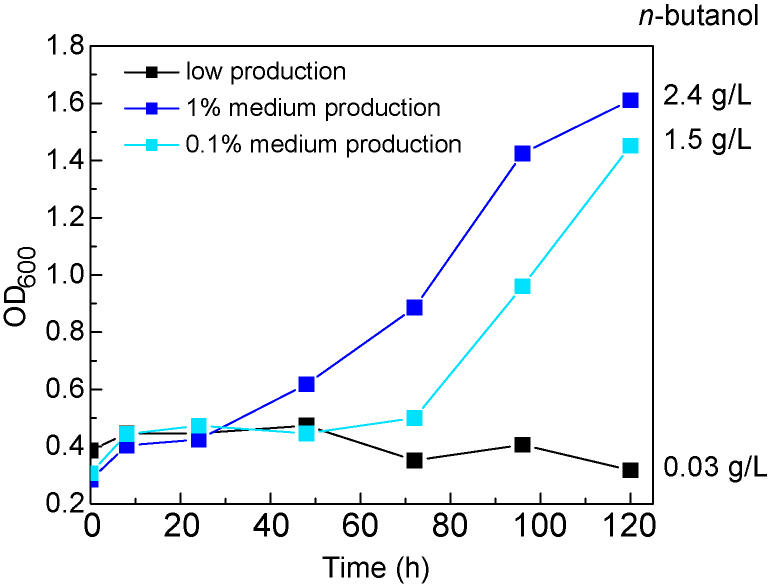

**Figure S12. Selection of *n*-butanol strains using EMS mutagenesis under anaerobic conditions.** *E. coli* DH1Δ*5* pBBR1-aceEF.lpd and pBT33-Bu2 pCWori.trc-ter-aldh46.adh was treated with EMS for 60 min and then subjected to enrichment in M9 media under anaerobic conditions. A representative evolutionary trajectory is found below. Although improvements were observed, the gain in product titer plateaued within two rounds of mutagenesis. Genome sequencing of genomic DNA isolated from the pool identified ~150 SNPs with 95% overlap, suggesting both that a large number of neutral mutations occurred and that cultures progressed rapidly to genetic homogeneity under the selection conditions. *n*-Butanol titers were measured using GC-MS. Data are mean ± s.d. of biological replicates (n = 3).

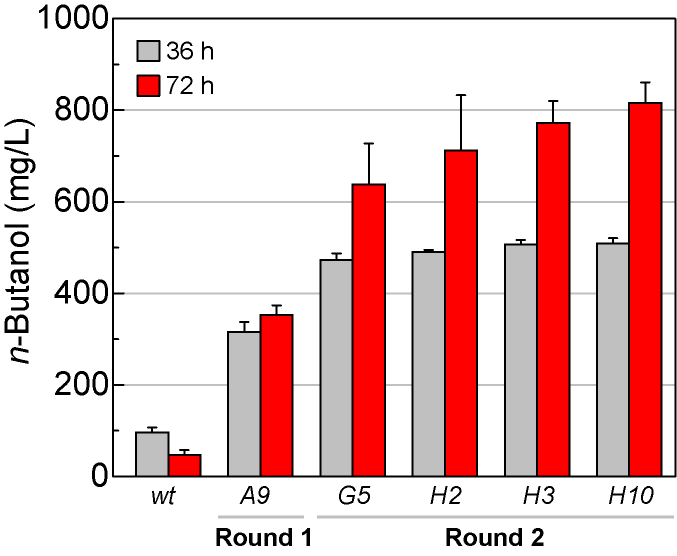

**Figure S13. Characterization of adaptive evolution of *n*-butanol strains under anaerobic conditions.** All selections were performed in triplicate with cultures supplemented with 2.5% (*w/v*) glucose. OD_600_ for each flask was measured before every dilution. Production titers were validated in the selection media and controls represent *E. coli* parent strains freshly transformed with the appropriate plasmids. Strain labels indicate plasmids/flask-dilution number-clone number. Numbers above bars correspond to a unique identifier number for the sequenced strain with a shape indicating specific genetic loci mutated (*Table S1.A2*). (A) Adaptive evolution with *E. coli* DH1Δ*5* as the host in LB media with three different *n*-butanol pathways. All strains contained the pBBR1-aceEF.lpd and pT5T33-Bu2 plasmids with different downstream plasmids (A, pCWori.trc-ter-aldh46.adh2; B, pCWori.trc-ter-aldh46.adh8; C, pCWori.trc-ter-aldh21.adh2). (B) Adaptive evolution with *E. coli* DH1Δ*5* as the host in M9 media supplemented with 10% LB (*v*/*v*). The parent strains for this evolution were derived from the selection in LB media: A35-D35-2 (2622), C1-D41-1 (2625), C3-D35-1 (2628), and C3-D41-1 (2629). (C) Adaptive evolution with *E. coli* BW21153Δ*5* as the host in M9 media supplemented with 10% LB (*v*/*v*). All strains contained the pBBR1-AceEF.Lpd and pT5T33-Bu2 plasmids with different downstream plasmids (A, pCWori.trc-ter-aldh46.adh2; B, pCWori.trc-ter-aldh46.adh8; C, pCWori.trc-ter-aldh21.adh2). (D) Adaptive evolution with various evolved strains in M9 media.

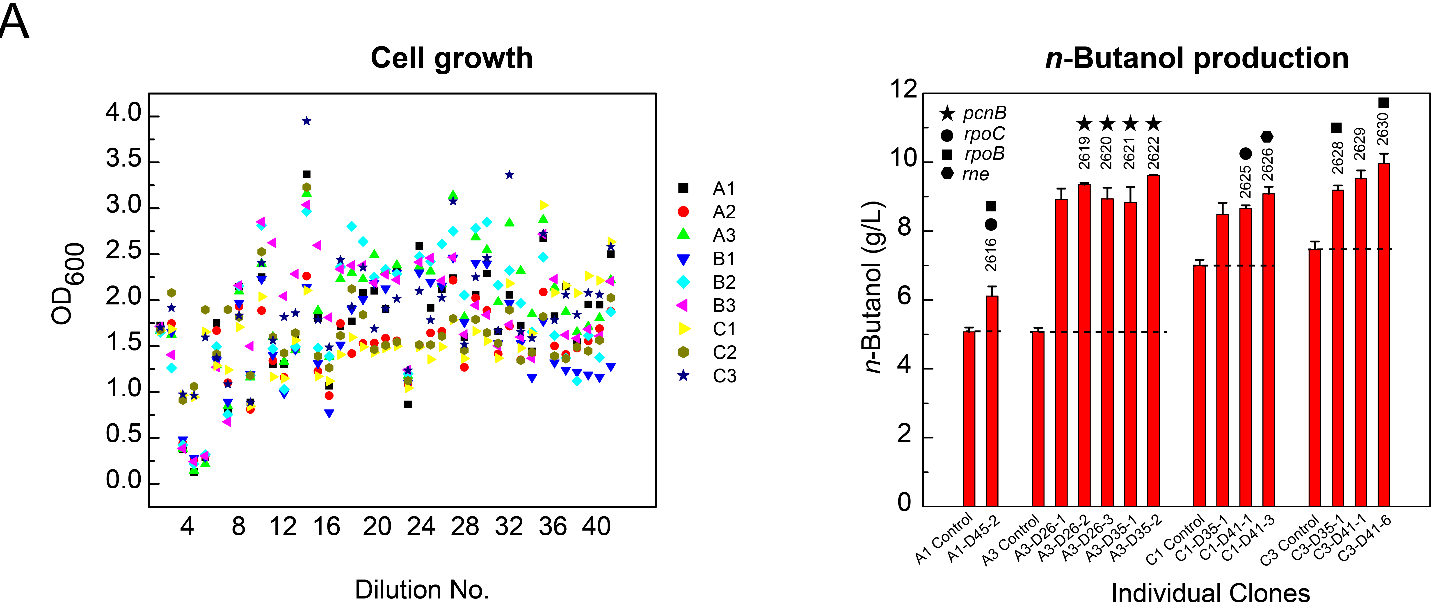

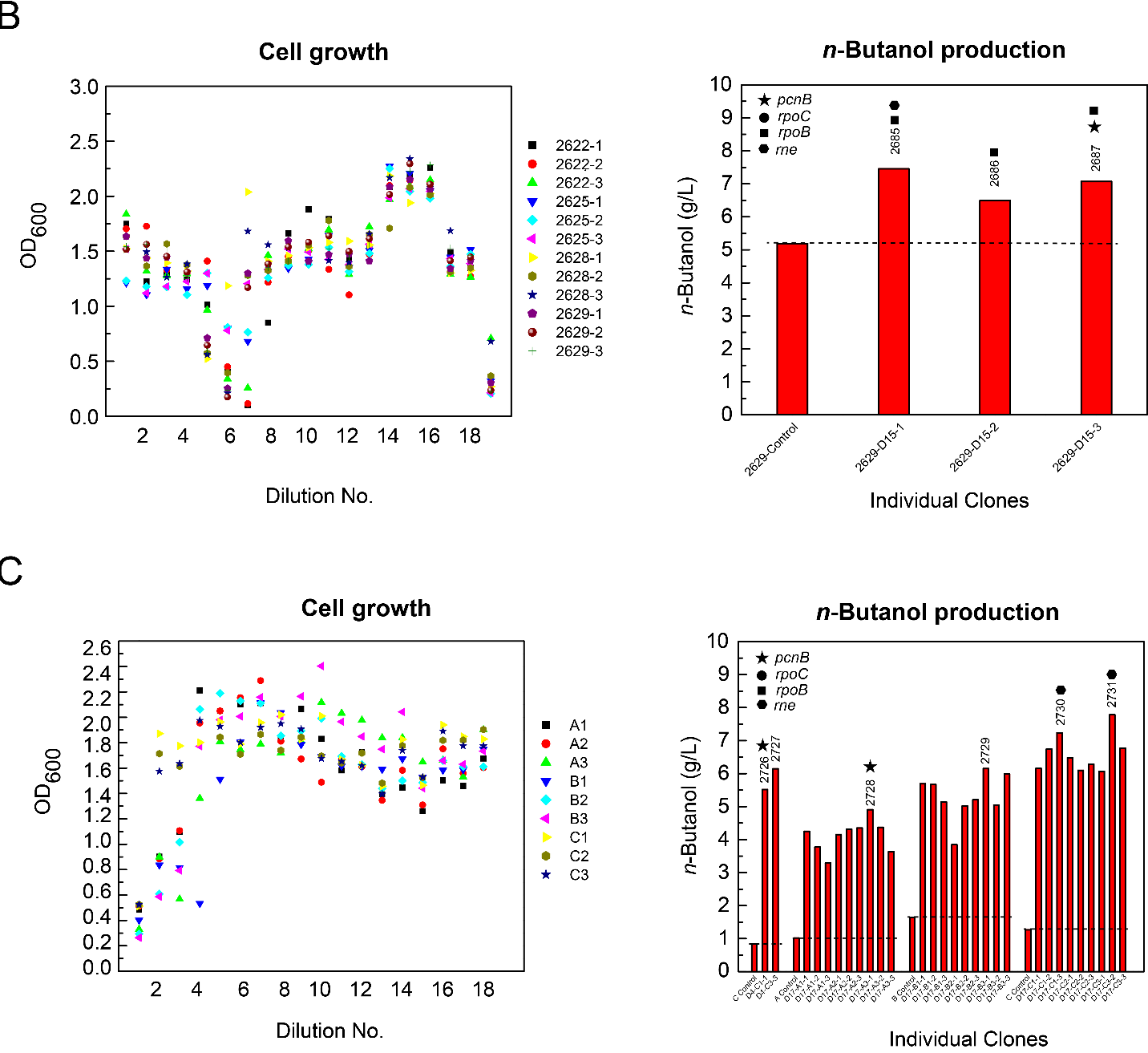

D

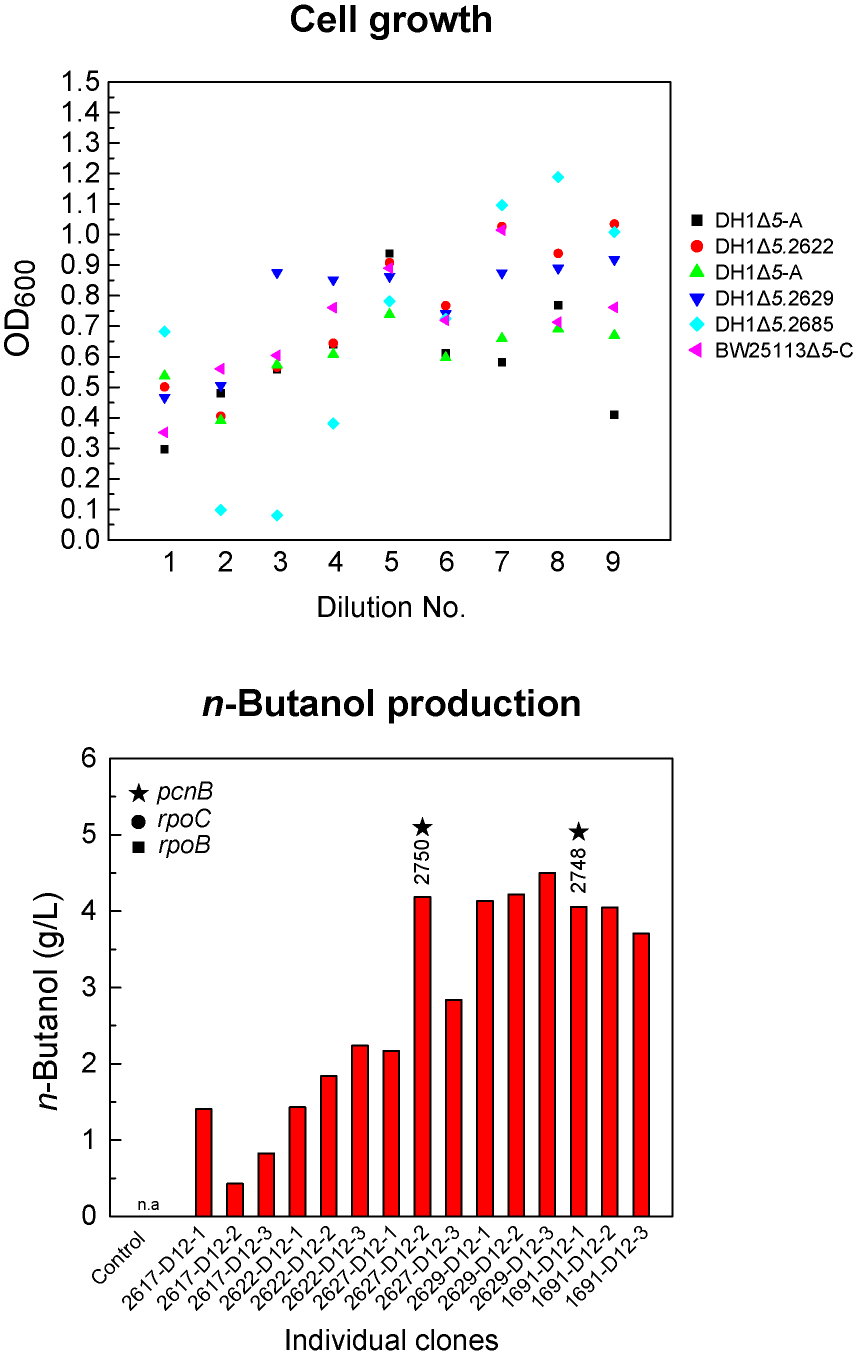

| **Parent strain** | **Description** | **Label** |
| --- | --- | --- |
| DH1Δ5 | Pathway A | 2617 |
| DH1Δ5.2622 | Isolated Pathway A clone from LB selection (*Figure S14A*: A3-D35-2) | 2622 |
| DH1Δ5 | Pathway C | 2627 |
| DH1Δ5.2629 | Isolated Pathway C clone from LB selection (*Figure S14A*: C3-D41-1) | 2629 |
| DH1Δ*5*.2685 | Isolated Pathway C clone from M9/LB selection (*Figure S14B*: 2629-D15-1) | 2685 |
| BW25113Δ*5* | Pathway C | 1691-C |

**Figure S14. Characterization of adaptive evolution of BDO and HB strains under anaerobic conditions.** All selections were performed in triplicate with TB cultures supplemented with 2.5% (*w/v*) glucose. OD_600_ for each flask was measured before every dilution. Production was validated in the selection media and controls represent *E. coli* parent strains freshly transformed with the appropriate plasmids. Strain labels indicate plasmids/flask-dilution number-clone number. Numbers above bars correspond to a unique identifier number for the sequenced strain with a shape indicating specific genetic loci mutated (*Table S1.A2*). (A) Growth curves of adaptive of HB and BDO evolution with (HB, strain A) DH1Δ*5* pBBR1-aceEF.lpd pT533-phaA pCWori.trc-ter-aldh7.adh2, (BDO, strain B) DH1Δ*5* pBBR1-aceEF.lpd pT533-phaA.phaB pCWori.trc-ter-aldh7.adh2, (BDO, strain C) DH1Δ*5* pBBR1-aceEF.lpd pT533-phaA.phaB pCWori.trc-sadh1-aldh7.adh2. Cultures were grown (B) Control BDO and HB production with plasmids extracted from evolved strains and transformed into a clean parental *E. coli* DH1Δ*5* host. (Strains A, B, and C, see Panel A description; Strain D, DH1Δ*5* pBBR1-aceEF.lpd pT533-phaA.phaB pCWori.trc-aldh3.adh22). The similar production compared to fresh plasmids indicates that mutations responsible for increasing product titer are likely found on the chromosome. Strain numbers for evolved strains are indicated above each bar in the figure. Data are mean ± s.d. of biological replicates (n = 3).

A

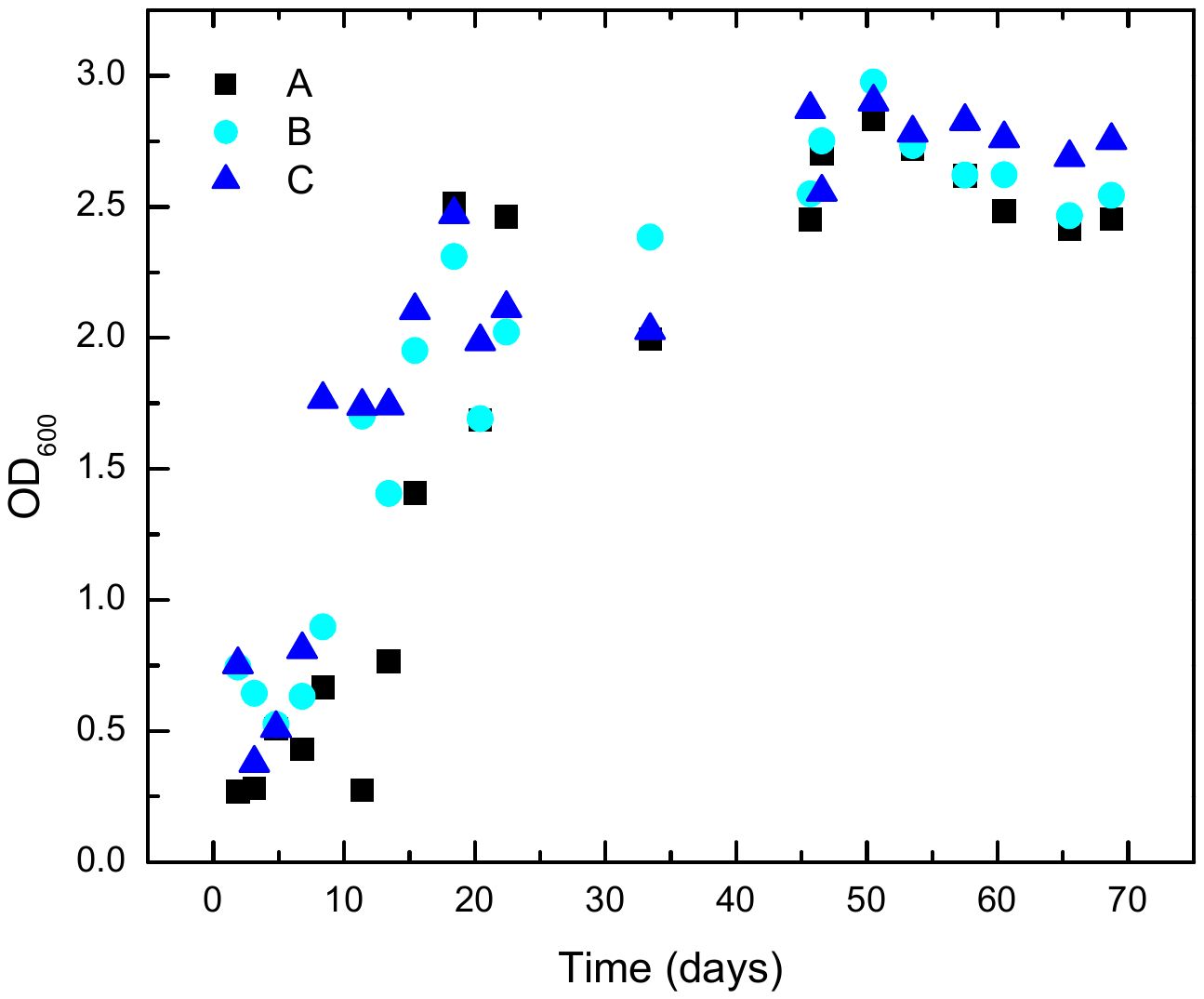

B

**Evolved parent strains**

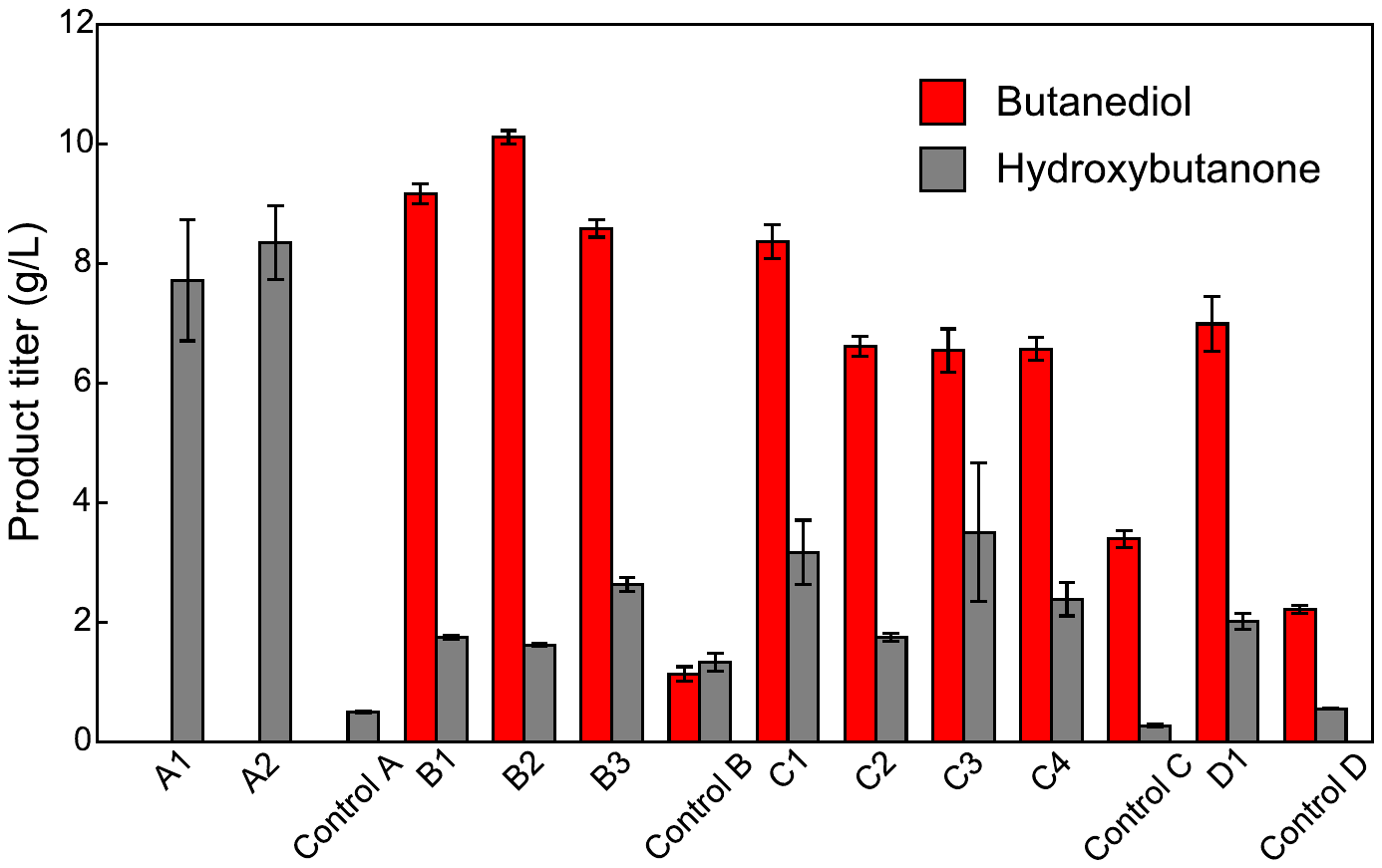

**Extracted plasmids transformed into clean host**

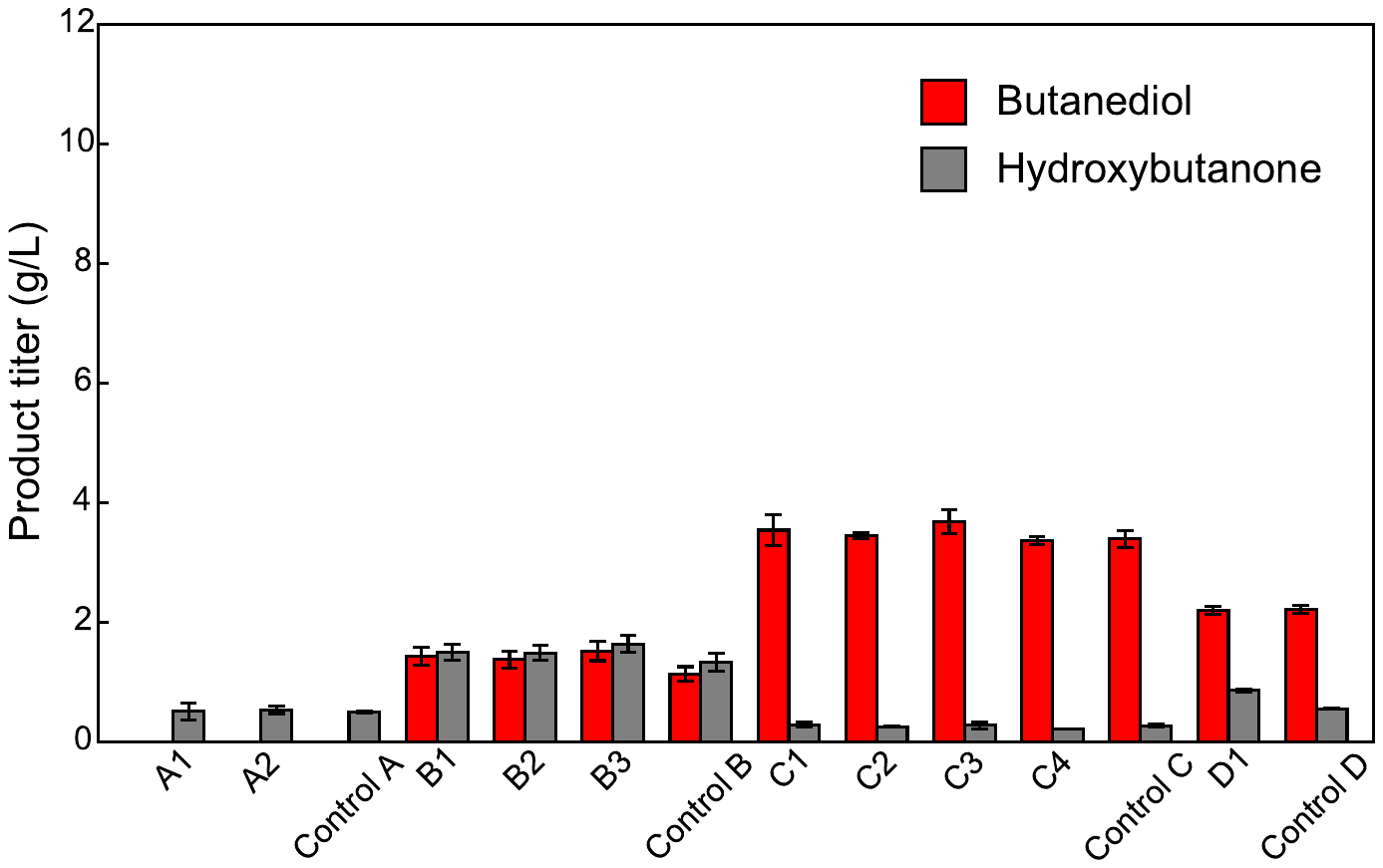

**Figure S15. Production titers of C4 monomers compared to parent strains.** Production titers in the parent DH1Δ*5* strain compared to the evolved DH1Δ*5*.2622 strains both bearing the production plasmids pBBR1-aceEF.lpd pT5T33-Bu2 pCWori.trc-ter-aldh46.adh2 plasmids. Cells were cultured in TB media supplemented with 8% (*w/v*) glucose with a 20% (*v/v*) oleyl alcohol overlay. Data are mean ± s.d. of biological replicates (n = 3). (A) *n*-Butanol production by DH1Δ*5*.2622 pBBR1-aceEF.lpd pT5T33-Bu2 pCWori.trc-ter-aldh46.adh2 compared to the parental strain. The theoretical yield for this experiment is 37.6 g L^-1^, indicating that the evolved strain can utilize carbon sources other than glucose to produce *n*-butanol if it is consumed. (B) HB production in DH1Δ*5*.2403 pBBR1-aceEF.lpd pT5T33-phaA pCWori.trc- aldh7.adh2 and BDO production in DH1Δ*5*.2406 pBBR1-aceEF.lpd pT5T33-phaAB pCWori.trc-sadh1-aldh7.adh2 compared to the parental strains.

A

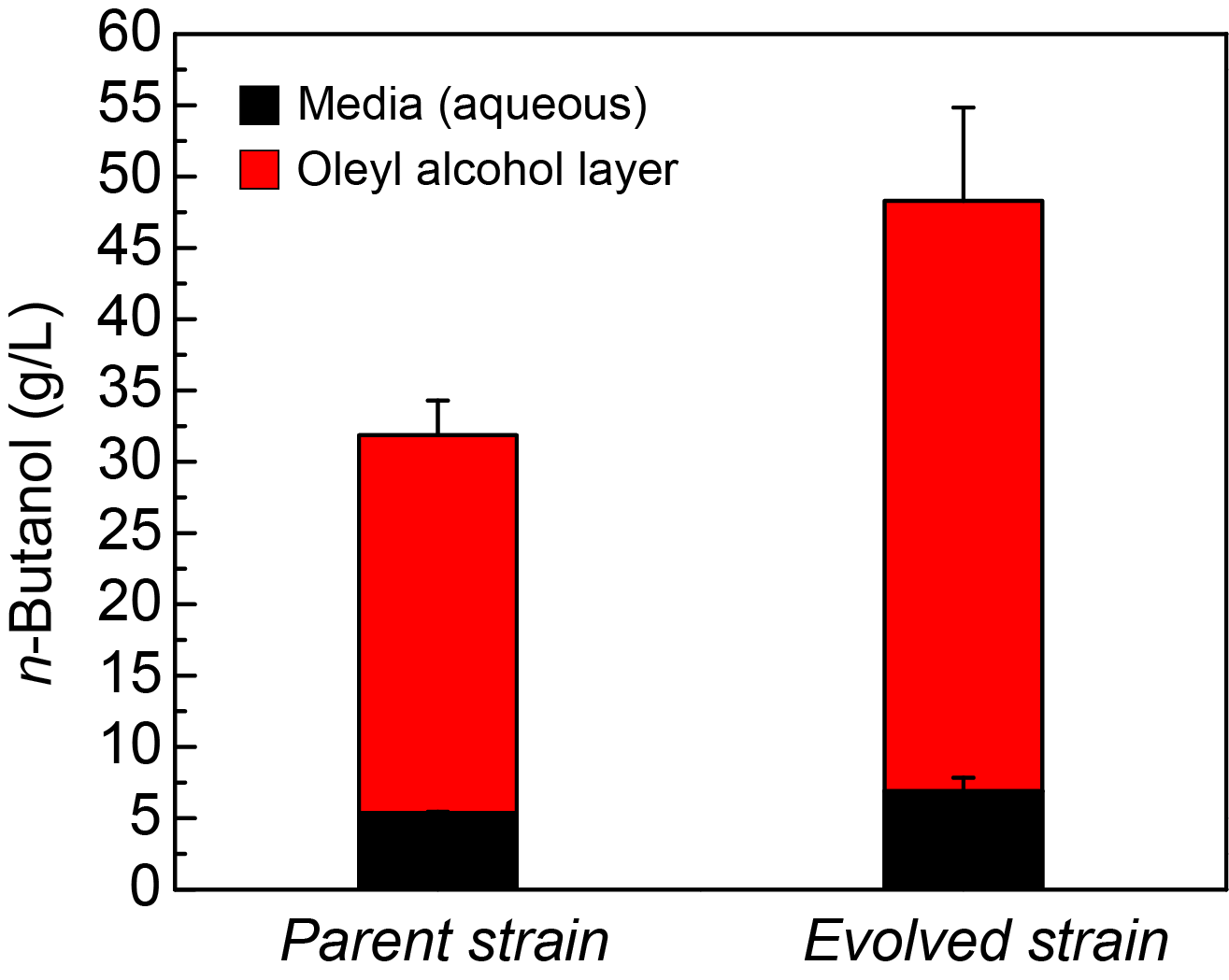

B

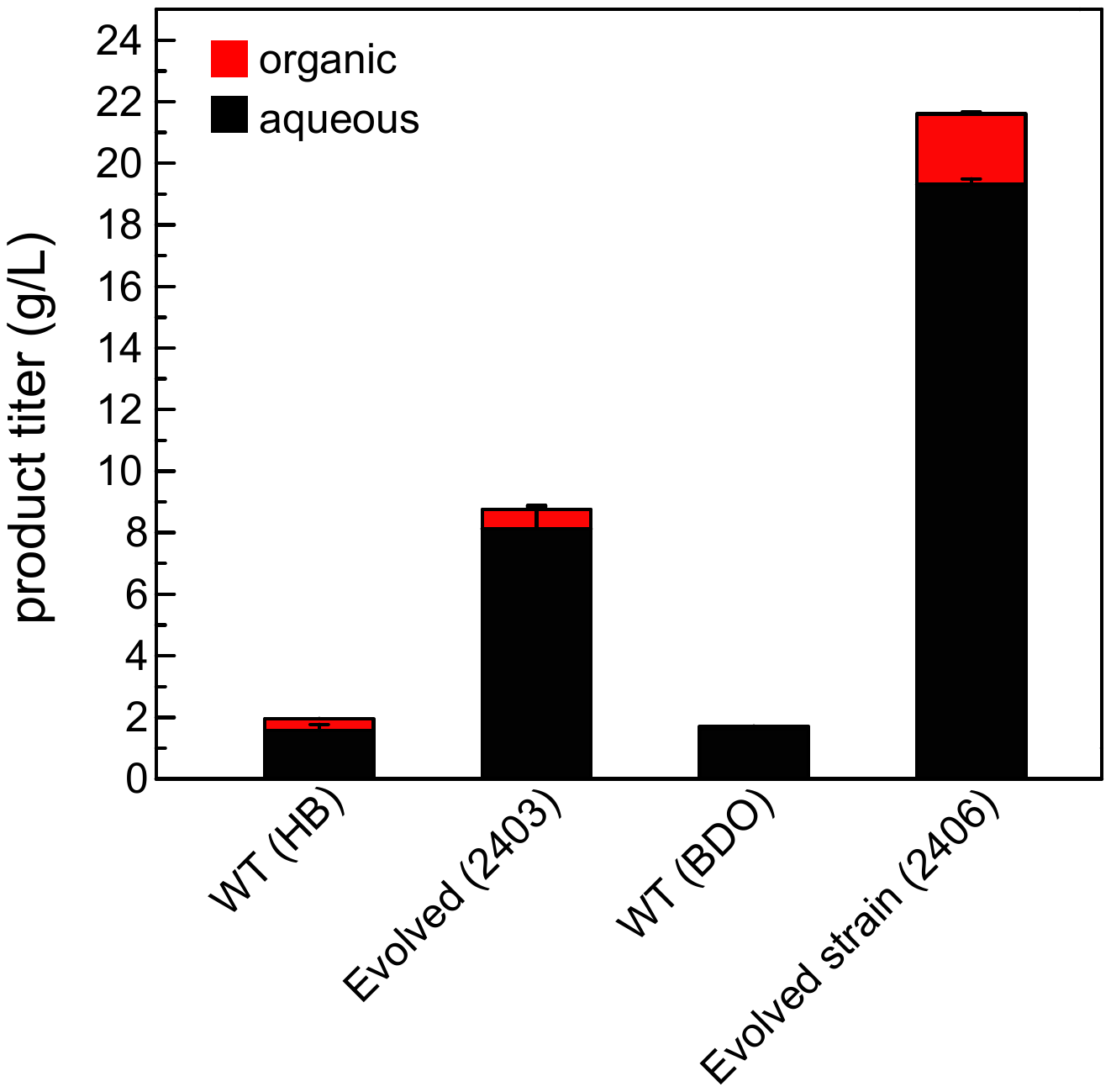

**Table S2. Genome sequencing of evolved strains.** 31 different evolved strains were sequenced along with the DH1Δ5 or BW25115Δ5 parent strain. Reads were mapped to the reference genome of DH1 or BW25113 and analyzed for changes including SNPs as well as rearrangements using Breseq. (A) Predicted point mutations in the coding region. (B) Predicted point mutations in intergenic regions. (C) Unassigned new junctions evidence. Each new junction consisits of two row, one describing one side of the junction in the reference sequence. (D) Missing coverage evidence. Full tables of differences between parent and evolved strains.

A

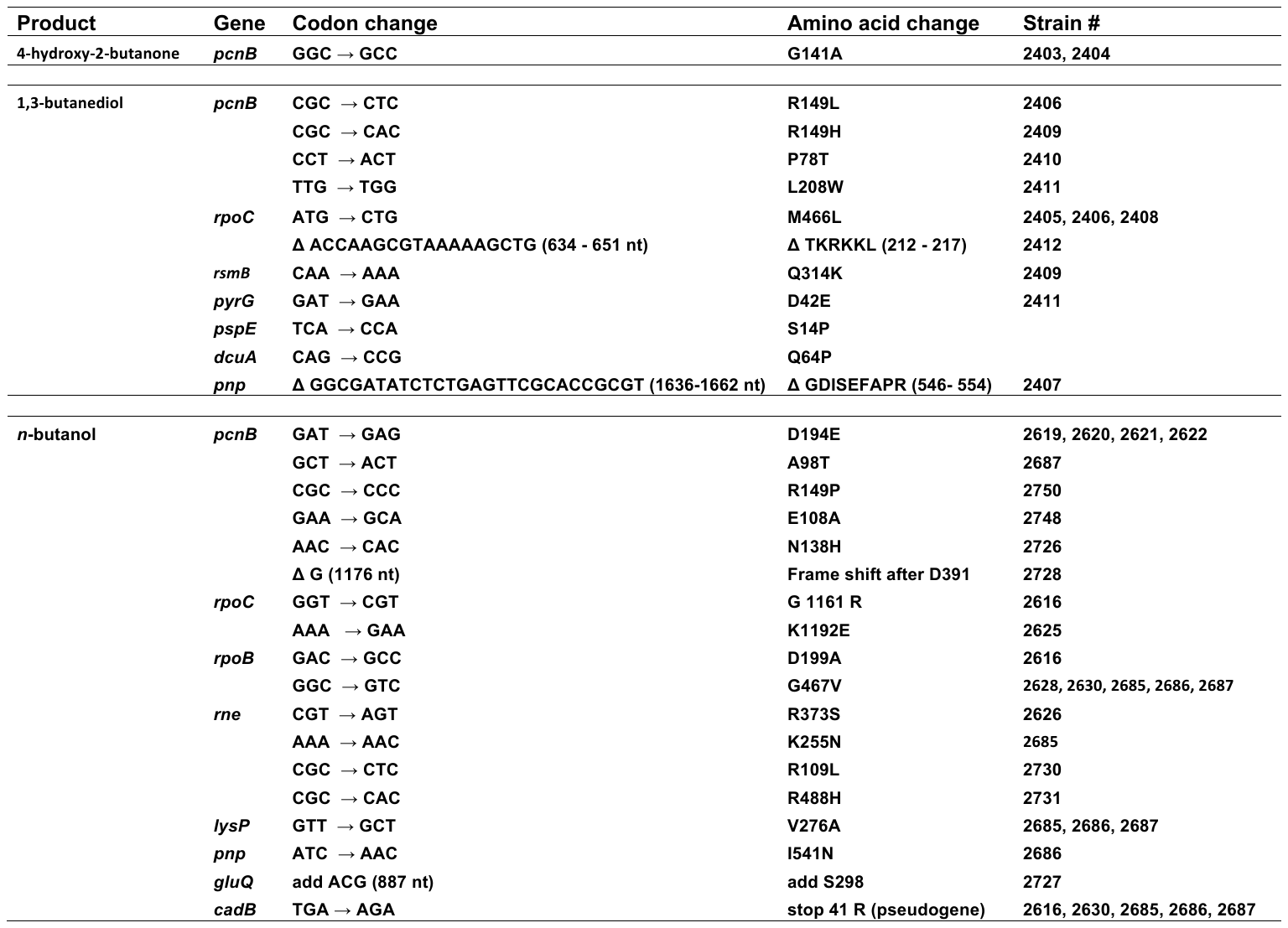

B

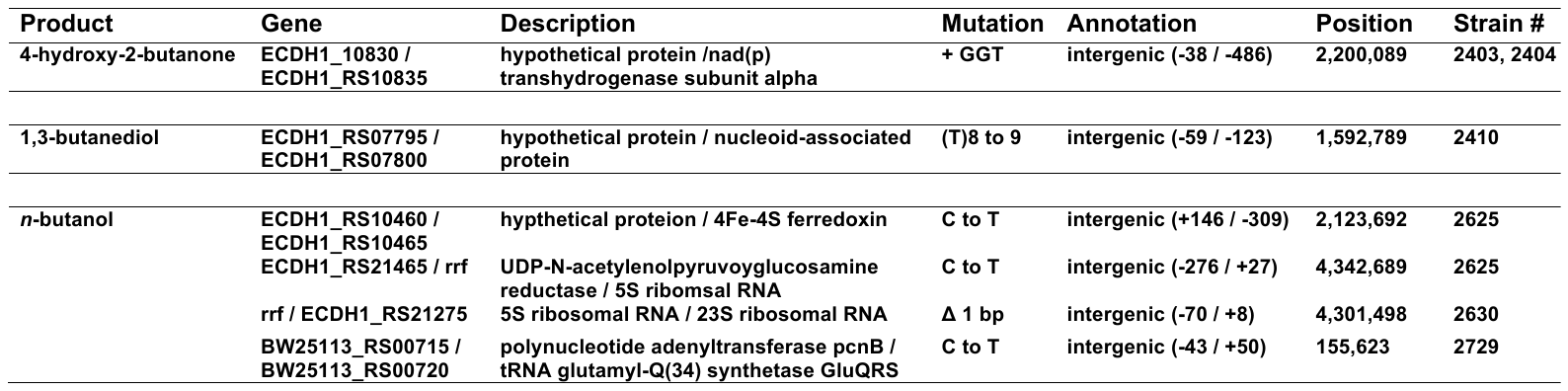

C

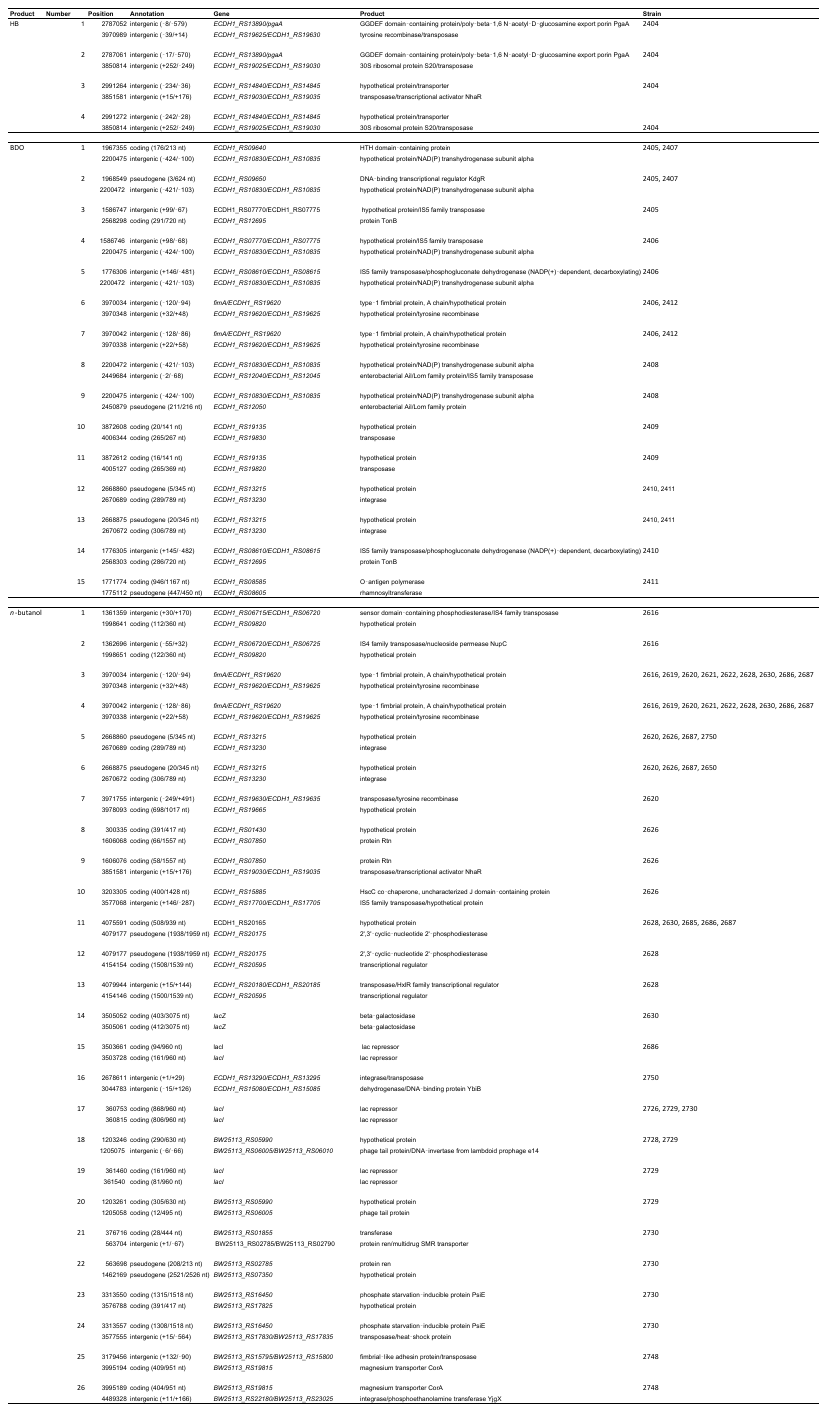

D

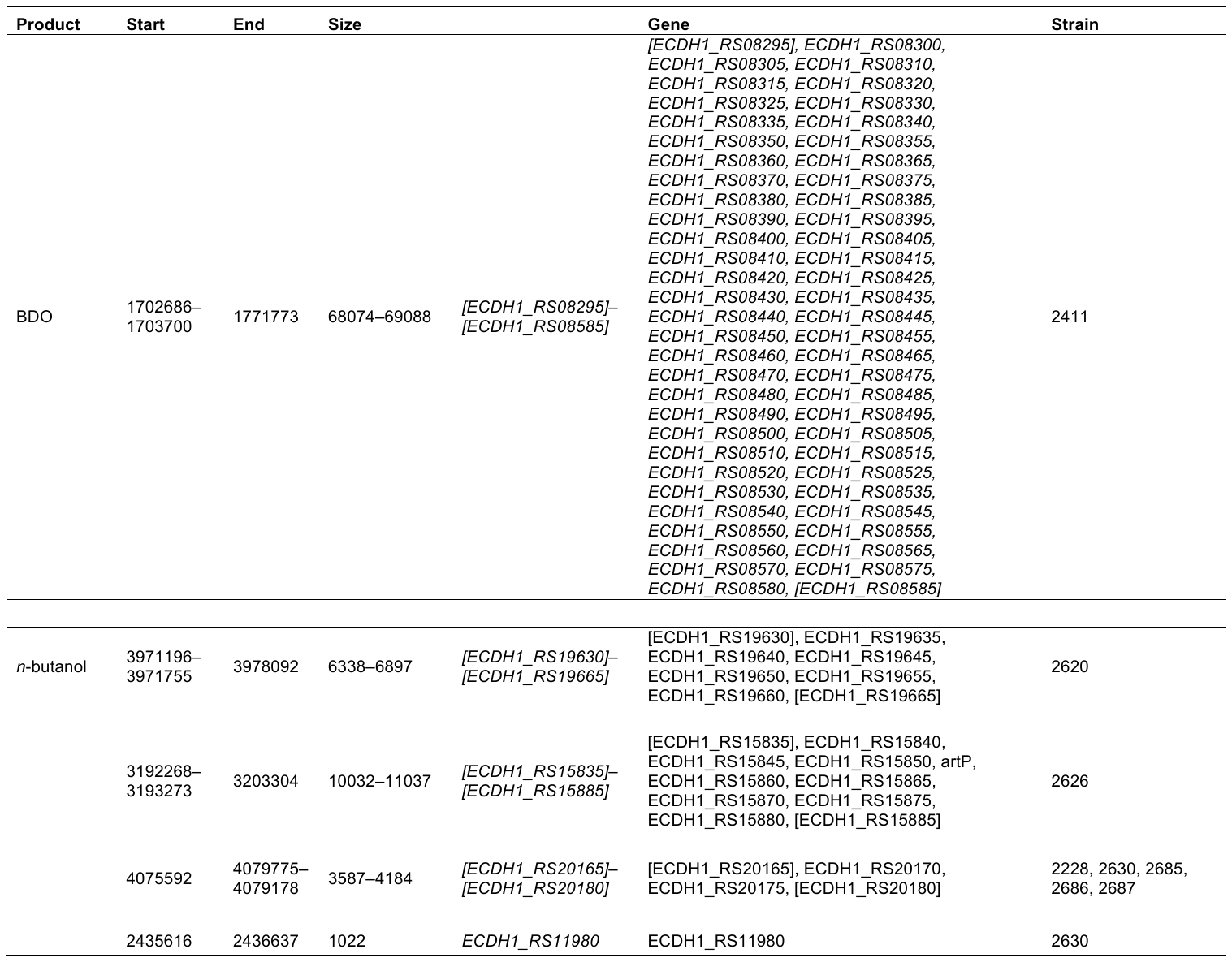

**Figure S16. Cell lysate enzyme activities of *n*-butanol pathway enzymes for parent and evolved strains.** Enzyme activities were measured in cell lysate of parent and evolved strains to examine whether increased heterologous expression of pathway enzymes could the source for increases in *n*-butanol titer. DH1Δ*5.*2616 was compared to its parent, DH1Δ*5* pBBR1-aceEF.lpd pT5T33-Bu2 pCWori.trc-ter-aldh46.adh2. BW25113Δ*5*.2730 was compared to its parent, BW25113Δ*5* pBBR1-aceEF.lpd pT5T33-Bu2 pCWori.trc-ter-aldh21.adh2. There is no significant activity differences for the four enzymes tested between the parent and evolved strains, leading us to conclude that differential pathway enzyme expression is not a major factor. Data are mean ± s.d. of biological replicates (n = 3).

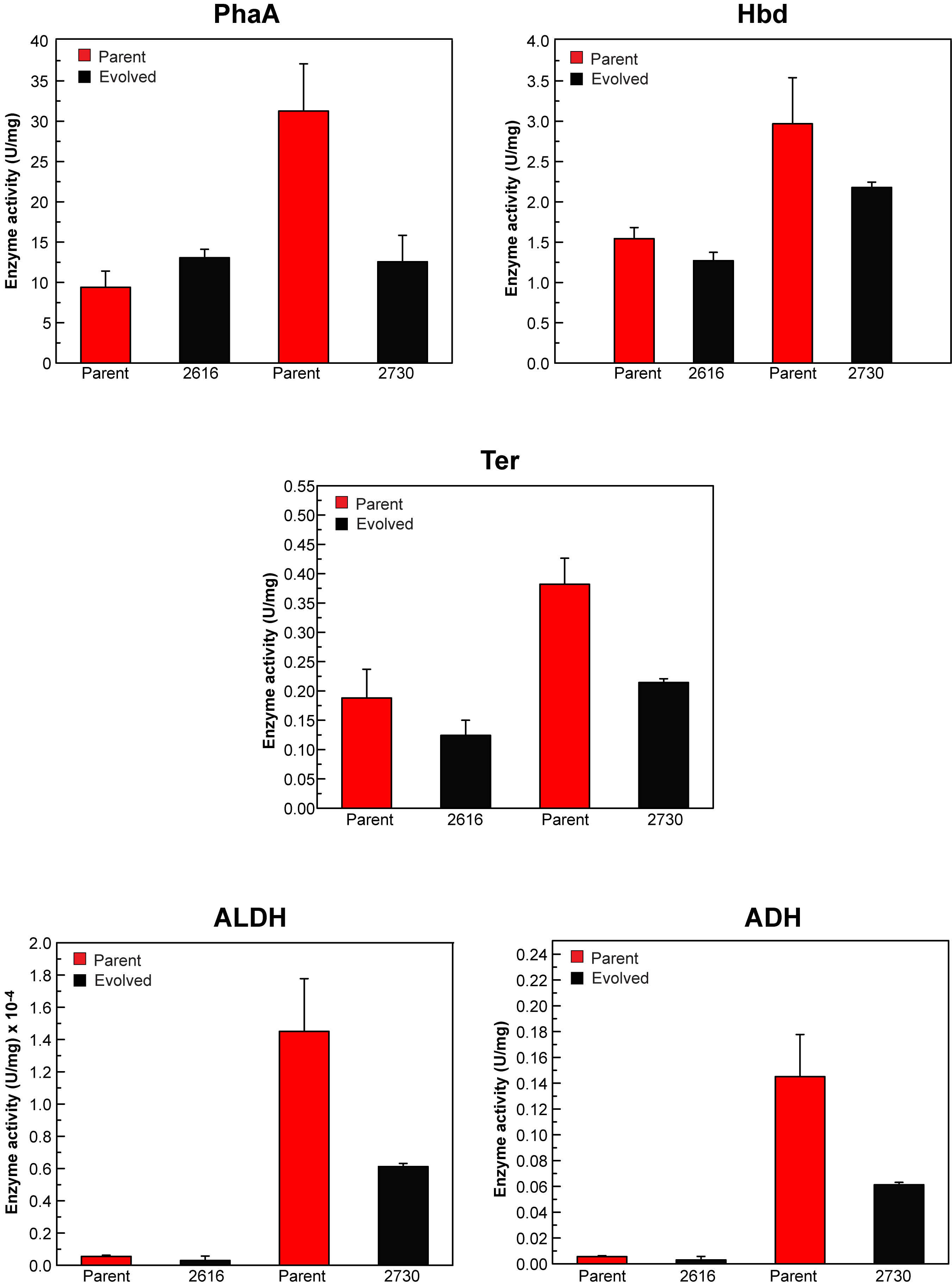

**Figure S17. RNA-Sequencing analysis of parent and evolved BDO strains.** Volcano plot of RNA sequencing data of the parent DH1Δ*5* pBBR1.aceEF.lpd pT533-phaA.phaB pCWori.trc-ter-aldh7.adh2 compared to the evolved strain (DH1Δ*5*.2406), generated by Sleuth.

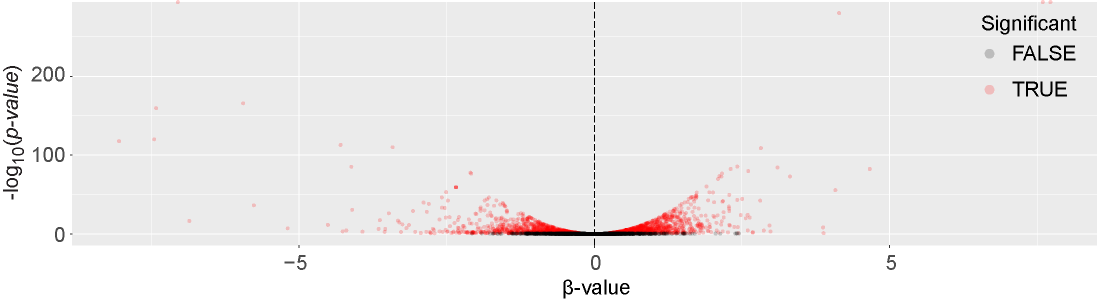

**Figure S18. Acetyl-CoA pool analysis between parent, mutant, and the evolved strain.** Biomass was harvested after 24 h growth under anaerobic conditions after IPTG induction. Metabolites were extracted using freeze and thaw cycles and supernatants were ran on LC-MS/MS for acetyl-CoA quantification. Data showed introducing the *pcnB* and *rpoC* mutations found in DH1Δ*5*.2406 in a clean genetic background (DH1Δ*5* parent) resulted in a ~2 fold increased in acetyl-CoA pool compared to the parent strain, which captures the same phenotype observed in the evolved strain. All three hosts carried pT533-phaA.phaB, pCWori.trc-aldh7.adh2, and pBBR1-aceE.F.lpd. Parent: DH1Δ*5;* PcnB_R149L RpoC_M466L: DH1Δ*5* _2406_*pcnB*(R149L)_*rpoC*(M466L); Evolved strain : DH1Δ*5*.2406. Data are mean ± s.d. of biological replicates (n = 5).

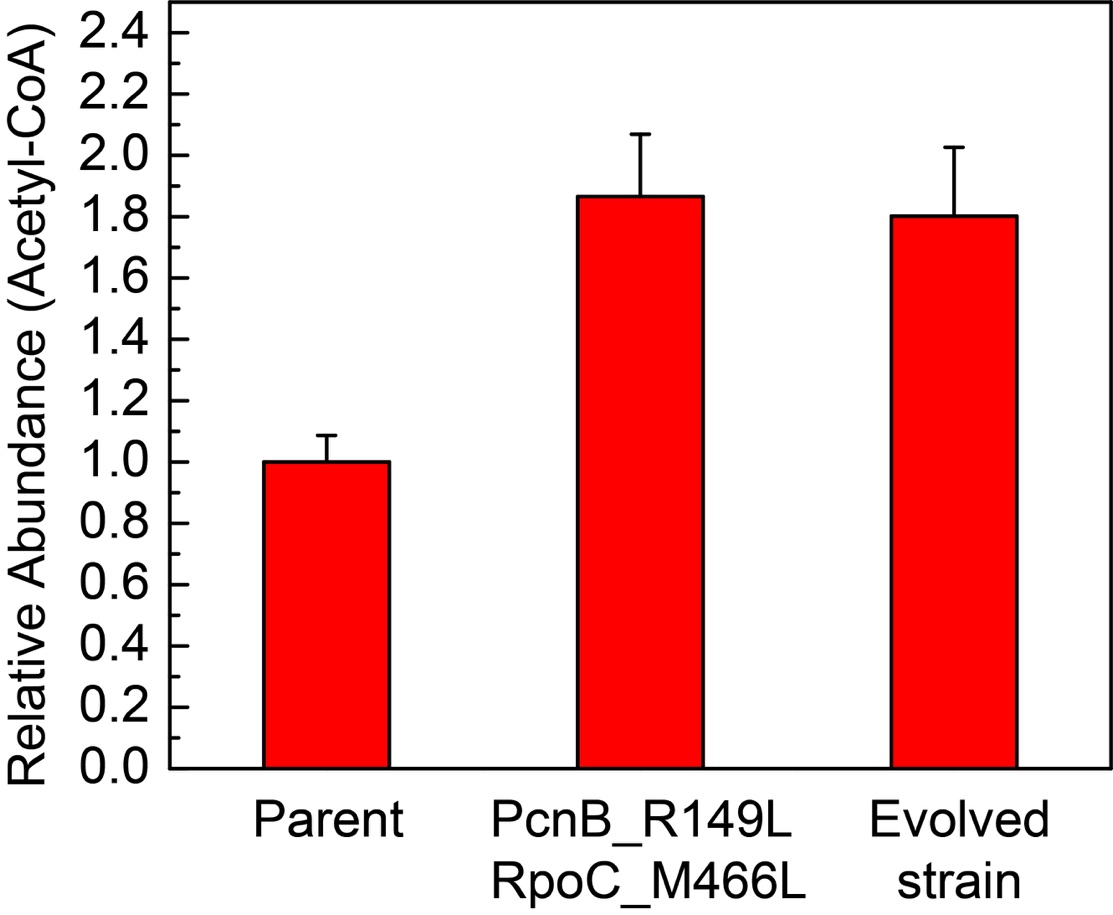
